# A global screen for assembly state changes of the mitotic proteome by SEC-SWATH-MS

**DOI:** 10.1101/633479

**Authors:** Moritz Heusel, Max Frank, Mario Köhler, Sabine Amon, Fabian Frommelt, George Rosenberger, Isabell Bludau, Simran Aulakh, Monika I. Linder, Yansheng Liu, Ben C. Collins, Matthias Gstaiger, Ulrike Kutay, Ruedi Aebersold

**Author notes:** Senior authors (Ulrike Kutay, Ruedi Aebersold). Corresponding author & Lead contact.

## Abstract

Living systems integrate biochemical reactions that determine the functional state of each cell. Reactions are primarily mediated by proteins that have in systematic studies been treated as independent entities, disregarding their higher level organization into complexes which affects their activity and/or function and is thus of great interest for biological research. Here, we describe the implementation of an integrated technique to quantify cell state-specific changes in the physical arrangement of protein complexes, concurrently for thousands of proteins and hundreds of complexes. Applying this technique for comparison of human cells in interphase and mitosis, we provide a systematic overview of mitotic proteome reorganization. The results recall key hallmarks of mitotic complex remodeling and discover new events, such as a new model of nuclear pore complex disassembly, validated by orthogonal methods. To support the interpretation of quantitative SEC-SWATH-MS datasets, we extend the software *CCprofiler* and provide an interactive exploration tool, *SECexplorer-cc*.

**Highlights:**

- Quantification of proteome assembly state changes using SEC-SWATH-MS
- Systems-wide analysis of assembly state changes in the mitotic proteome
- Discovery and validation of a novel mitotic disassembly intermediate of the nuclear pore complex
- Higher sensitivity and information content compared to thermostability-based approaches for global measurement of proteome states
- *SECexplorer*, an online platform to browse results and investigate proteins newly implicated in cell division

## Introduction

Living systems are characterized by a large number of biochemical functions that are tightly interconnected and coordinated (Hartwell et al., 1999; Ideker et al., 2001). Classical biochemical analyses have led to the association of many biochemical functions with proteins and protein complexes. The function of proteins and protein complexes depends on a defined 3D structure of individual proteins, as well as the composition and specific steric arrangement of proteins into protein-protein complexes. Detailed studies on tetrameric hemoglobin have shown that changes in the composition, arrangement or structure of the complex changes its activity (Pauling 1949), a seminal finding that has since become one of the hallmarks of molecular biology. Whereas much of our biochemical understanding is based on in-depth studies of specific complexes, these are time consuming and more importantly, disregard interactions and coordination between different complexes.

Driven primarily by genomics, life science research has been transformed by high throughput, data-driven approaches. Proteomics is the embodiment of this approach for proteins. To date, most proteomic studies have been carried out by *bottom up proteomics*, where proteins are denatured and cleaved into peptides which are then analyzed by mass spectrometry. Whereas this technology has reached a high level of maturity, information about the structure, the composition and steric arrangement of components in a complex are lost. Therefore, for the most part, proteomics has treated proteins as unstructured biopolymers, disregarding structure and organization into complexes as an important layer of function and control.

Recently, several techniques have been proposed that attempt to extend the large scale analysis of proteins towards the detection of conformational changes between different states ((Becher et al., 2018; Dai et al., 2018; Leuenberger et al., 2017; Liu and Fitzgerald, 2016; Schopper et al., 2017; Tan et al., 2018), towards the organization of proteins into complexes (Heusel et al., 2019; Kristensen et al., 2012; Scott et al., 2017; Wan et al., 2015) and towards assigning protein subcellular localization (Dunkley et al., 2004; Foster et al., 2006). Conformational changes of proteins have been detected by using changes in physico-chemical properties including thermostability ((Becher et al., 2018; Dai et al., 2018; Tan et al., 2018)), stability towards denaturing conditions (Xu et al., 2014) or altered protease susceptibility (Leuenberger et al., 2017; Liu and Fitzgerald, 2016; Schopper et al., 2017) as proxy. Inference of complex composition and subcellular localization has been based on chromatographic fractionation, typically by size exclusion chromatography (SEC), of native complexes (Kristensen et al., 2012; Larance et al., 2016; Liu et al., 2008) and subcellular fractions (Dunkley et al., 2004; Foster et al., 2006; Itzhak et al., 2016), followed by the mass spectrometric analysis of the resulting fractions. Pioneering, comparative analyses of co-fractionation patterns of native complexes have revealed extensive re-organization of the modular proteome across metazoans (Wan et al., 2015) and following induction of apoptosis (Scott et al., 2017). However, the co-fractionation approach has been beset by limited SEC resolution, and limitations inherent in data dependent analysis (DDA) mass spectrometry, the method almost universally used in co-fractionation studies. These include limited proteomic depth and accuracy of quantification and stochastic peptide sampling (Aebersold and Mann, 2016). Collectively, these limitations resulted in the need for multidimensional separation to assign proteins to specific complexes and, for the most part, unknown error levels of complex assignments (Kristensen et al., 2012; Scott et al., 2017; Stacey et al., 2017; Wan et al., 2015). Recently, we demonstrated increased selectivity and overall performance in co-fractionation-based profiling of cellular complexes using a workflow that is based on single dimension fractionation by high resolution size exclusion chromatography, quantitative measurement of polypeptide elution profiles by SWATH mass spectrometry, an instance of data independent acquisition (DIA)(Gillet et al., 2012, 2016) and the introduction of a complex-centric data analysis strategy (Heusel, Bludau et al., 2019). We also described a software tool *CCprofiler*, that implements a complex-centric strategy to infer protein complexes and uses a target-decoy model to assign a probability to each complex (Heusel, Bludau et al., 2019).

Here, we apply SEC-SWATH-MS to detect rearrangements in the modular proteome in HeLa CCL2 cells in two cell cycle states, the interphase and prometaphase. We developed a quantification module for the *CCprofiler* that supports the differential, quantitative analysis of thousands of proteins and their association with complexes. We benchmark the reproducibility of the integrated wet lab/computational method and compare its performance to state-of-the art thermostability-based methods (Becher et al., 2018; Dai et al., 2018). We validate the method by showing that it recapitulates known complex remodeling events between the different states tested. We discover and validate by orthogonal methods a new model of nuclear pore complex disassembly, thus demonstrating the potential of the method to discover new biology. To support additional exploration of the present dataset and future differential SEC-SWATH-MS datasets we provide an online tool, *SECexplorer-cc*.

We expect that the parallel quantification of abundance and compositional changes of hundreds of protein complexes will significantly advance our understanding of biochemical mechanisms and processes.

## Results

### Generation of a SEC-SWATH-MS dataset for the detection of changes in the organization of the mitotic proteome

As basis for our study into mitotic changes in the organization of the proteome, we applied the previously described SEC-SWATH-MS workflow, in conjunction with targeted analysis (Heusel, Bludau et al., 2019) in triplicate to cells synchronized in either cell cycle state as illustrated in Figure 1A. Mild cellular extracts containing native proteins and protein complexes were fractionated by high resolution SEC and the proteins in each fraction were digested and quantitatively profiled across the chromatographic fractions by SWATH mass spectrometry (Collins et al., 2017; Gillet et al., 2012; Röst et al., 2014). The samples tested were Hela CCL2 cells synchronized in interphase or mitosis. We inferred changes in complex composition and quantity from the resulting protein apparent size distribution patterns. The mitotic arrest of the respective synchronized cell populations was documented by microscopic assessment of cell shape and the detection of mitosis-specific electrophoretic mobility shifts of hyper-phosphorylated Nup53 and Histone H3 phosphorylation (**Supplemental Figure S1 A-C**). For each of the three replicates performed for either condition, 65 consecutive SEC fractions were collected and for each fraction the proteins were digested and analyzed, by SWATH-MS, generating a quantitative dataset consisting of a total of 390 SEC fractions. The resulting dataset was computationally analyzed using the OpenSWATH software suite and a project specific spectral library as prior information (see Figure 2A and experimental procedures for details). Overall, the analysis identified 70,445 peptides associated with 5514 proteins at a TRIC target FDR of 5%. Upon SEC-informed filtering as described previously (Heusel, Bludau et al., 2019), 60,891 peptides and 5,044 proteins were quantified with high confidence across the chromatographic fractions and with an overall decoy-estimated protein level false discovery rate of below 0.4% (see methods for details). Per mitotic condition, we quantified 52,718 and 56,553 peptides resulting in 4,438 and 4,798 protein profiles in interphase and mitosis, respectively (Figure 1B). The SEC conditions we used resolved proteins and protein complexes ranging from ca. 5 MDa to 10 kDa with a peak capacity to baseline-separate ca. 20 peaks and showed good reproducibility as apparent from UV/Vis spectrometric traces from the respective samples (Figure 1C). The large absorbance at 280 nm observed in the low molecular weight range (fractions 63-80) originated from detergents employed for mild lysis, as evidenced by the drop off of protein-level MS intensities beyond fraction F55, in line with the 30 kDa molecular weight cutoff employed for sample workup (Figure 1D). Even though a comparably high resolution SEC method was used, we detected in the range of ca. 1,200 to 2,000 proteins per fraction (Figure 1D**, lower panel**). Whereas most proteins were detected in both conditions, 4,192 (83%) proteins appeared more readily extractable from mitotic cells (Figure 1B). This is apparent from the lower cumulative ion intensities across most fractions (Figure 1D) and is potentially a consequence of mitotic reorganization including nuclear envelope breakdown. The thus generated set of protein abundance profiles was the basis for the further analyses.

**Figure 1:**
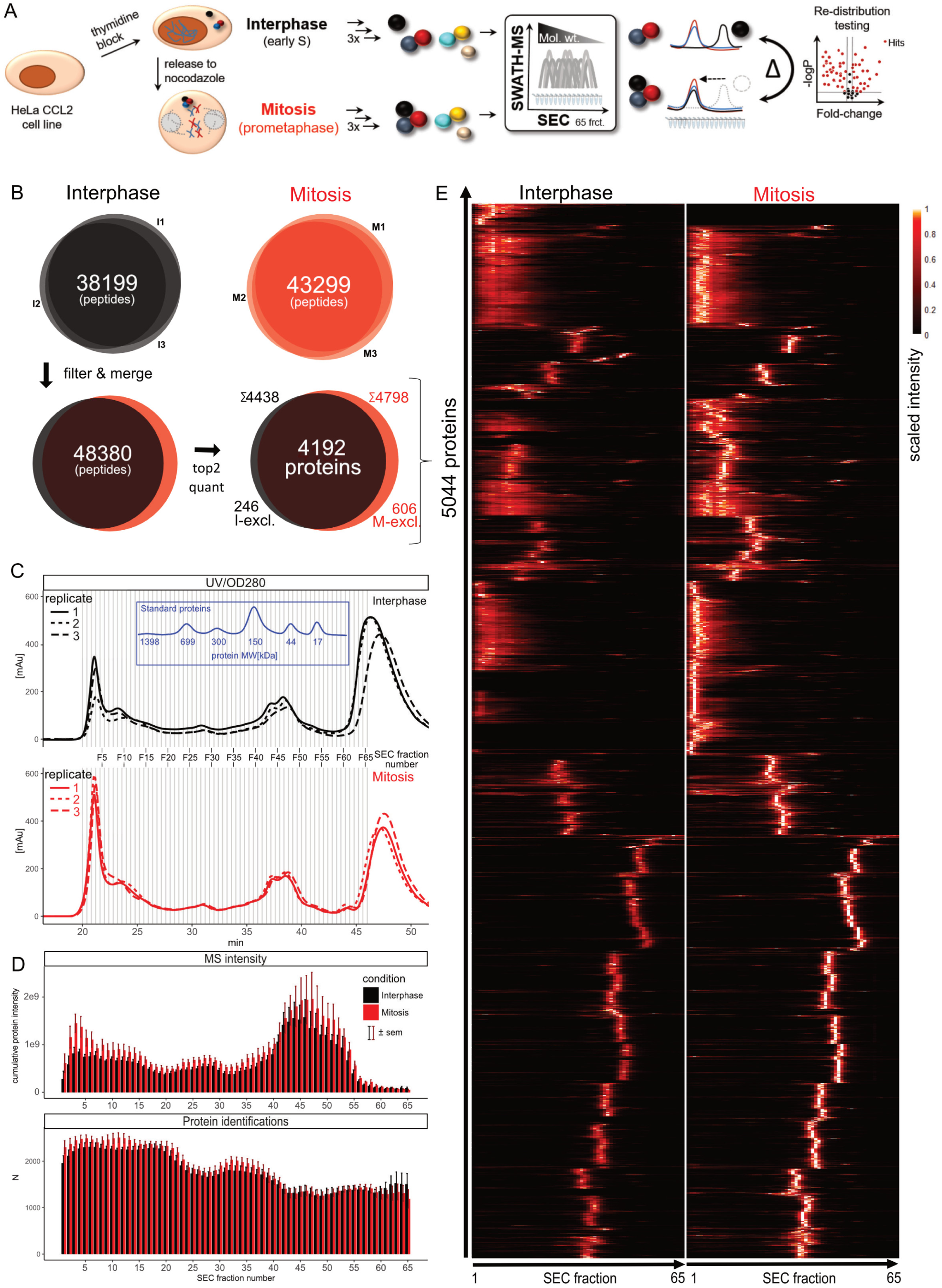
Proteome rearrangement screening by SEC-SWATH-MS: Workflow and dataset properties. **A** Scheme of the SEC-SWATH-MS workflow to screen for mitotic proteome rearrangement. HeLa CCL2 cells were synchronized in interphase or mitosis by chemical treatment, followed by triplicate extraction of complexes by mild lysis, fractionation by size exclusion chromatography and quantitative profiling of eluting proteins. Differential association scoring via *CCprofiler* reveals proteins with altered complex association states between conditions. **B** Peptide identifications across the three experimental repeats and summary per condition on peptide and protein level, giving rise to the dataset overview in panel E. Of the total 5044 observed proteins, most were detected independently in both conditions 4,192 (83%). **C** Semi-preparative scale SEC of interphasic and mitotic complex preparations and size reference protein mix monitored by UV/Vis spectroscopy. Elution of standard proteins calibrates the fraction number to apparent molecular weight mapping in the study. **D** Summary of MS-observed protein level intensity and number of confidently identified proteins along the 65 fractions across the repeats and after normalization (See data processing methods). The large absorbance at 280 nm observed in the low molecular weight range originates from detergent employed for mild lysis and does not reflect protein mass that was not sampled in the fractionation scheme, as can be extrapolated from protein-level MS intensities dropping beyond F55, in line with the 30 kDa molecular weight cutoff employed in sample workup. **E** Dataset overview heat map summarizing the data to 5,044 conditional protein elution patterns observed. Mean intensities of the top 2 cumulatively highest-intense peptides per protein were summarized from 3 replicate measurements and scaled from 0 to 1 per protein for visualization in heat map. Differential analysis is however performed at the level of individual, protein-specific peptides (See Figure 2).

**Figure 2:**
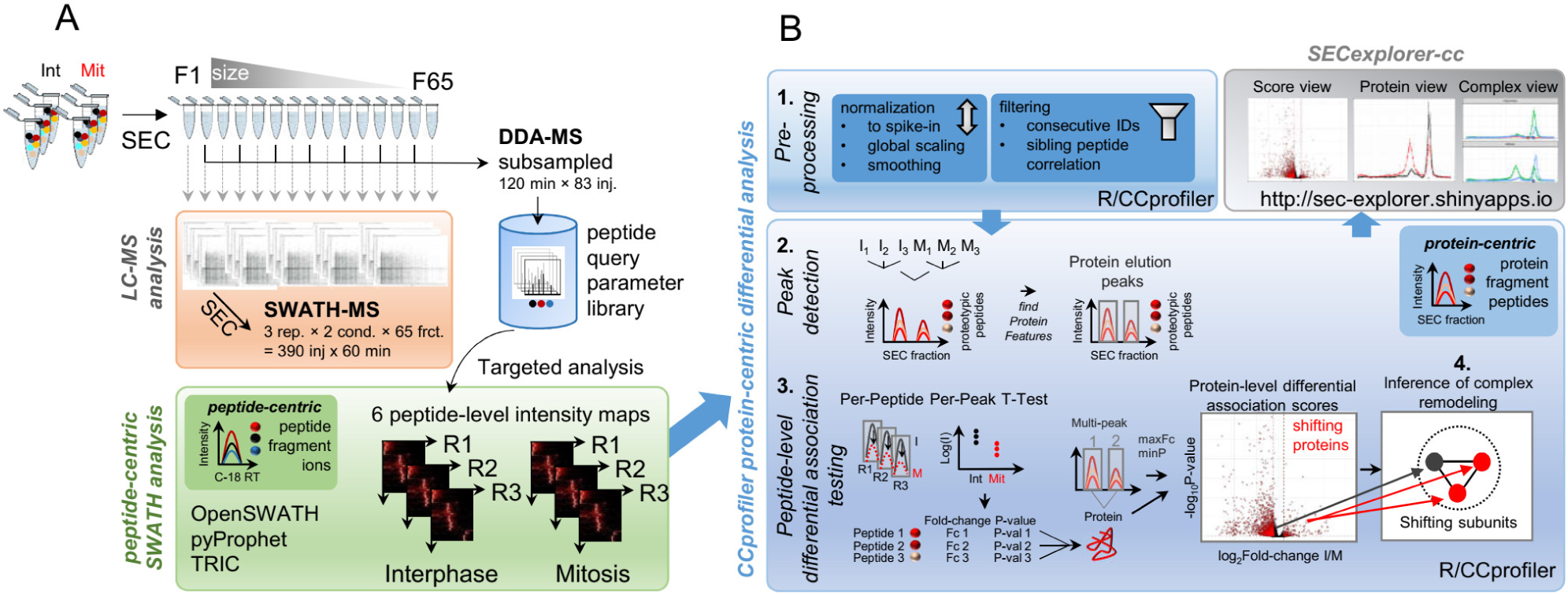
Proteome rearrangement screening by SEC-SWATH-MS: Data acquisition and processing. **A** LC-MS/MS analysis of tryptic peptides generated from 65 fractions per replicate and condition by data-independent acquisition SWATH mass spectrometry (SWATH-MS, n runs = 390). A subsampled set of fractions were subjected to longer 120 min gradient length analysis acquiring data in data-dependent mode to deepen coverage of the sample-specific peptide spectrum and query parameter libraries (DDA-MS, n = 83). Quantitative signals of the targeted peptides is then extracted from the SWATH-MS maps in peptide-centric analysis where peptide analytes are evidenced based on peptide fragment ion signal peak groups along C-18 chromatographic retention time. This produces six peptide-level quantitative matrices as basis for extracting information on protein complex association changes in subsequent differential analysis via *CCprofiler*. **B** Detailed scheme of protein-centric differential analysis of the quantitative peptide-level data via *CCprofiler* involving 1. preprocessing, 2. protein-centric elution peak detection where protein analytes are evidenced based on proteolytic peptide (‘protein fragment’) signal peak groups along SEC retention time/fraction number and 3. peak-resolved peptide-level statistical scoring to detect differentially associating proteins with shifting mass distribution across SEC fractions. In step 4, from the protein/subunit-level hits protein complex remodeling is inferred in a complex-centric fashion. The results are made available in easily browsable form via *SECexplorer-cc*.

### Quantification of protein association state changes from SEC-SWATH-MS data

The distribution of protein intensities per fraction for either condition provides a bird’s eye view of the acquired dataset (Figure 1E). To detect proteins that show significant changes with respect to their association with specific protein complexes, we applied a scoring system that quantifies protein mass re-distribution across distinct elution ranges based on two or more proteotypic peptides quantified by SWATH-MS in each peak. The association of proteins with protein complexes was carried out by the *CCprofiler* (Heusel, Bludau et al., 2019) a software tool that implements a complex-centric strategy using prior knowledge of protein complex composition. To detect quantitative changes of protein complex quantity and composition in either cellular state, we added to the *CCprofiler* tool a new module supporting differential quantification. It performs the sequential steps schematically shown in Figure 2B.

In the first step, signal intensities of SEC-fractions are normalized to a spike-in standard, missing values are imputed using the background signals from neighboring SEC fractions and SEC traces are aligned across experiments (Figure 2B, panel 1, for details and tools used see experimental procedures). The result of the first step is a calibrated and refined list of peptides and their respective intensities per fraction.

In the second step, the peptide level data are used to detect protein elution peaks along the SEC dimension. This is achieved by selecting a high quality set of peptide traces, by summing peptide intensities across replicates and conditions and by then employing the *CCprofiler* protein-centric analysis module to infer protein elution peaks. An elution peak is derived from the observation of co-eluting peaks of groups of sibling peptides derived from the same parent protein (See Figure 2B, step 2. and peak detection summary and example in **Supplemental Figure S4A**&**B**). This analysis resulted in a total of 6,040 elution peaks of 4,515 of the identified proteins. Accordingly, protein-centric analysis successfully detects peptide co-elution peak groups from 90% of the identified proteins and their peptide SEC profile sets, with no high quality elution signal detectable for the remaining proteins at the confidence threshold (q-value < 0.05). The distinctive elution peaks represent unique complex assembly states of the respective proteins. Consistent with previous protein-centric analyses of the proteome of cycling HEK293 cells (Heusel, Bludau et al., 2019), the majority of the observed elution peaks in the present dataset fell into a SEC separation range consistent with the association of the protein with a complex.

The distinctive protein elution peaks and their signal intensities computed in step two provide the basis for the third step we term ‘differential association testing’. Here, we calculated the log2-transformed abundance of each peptide per replicate for each observed protein elution peak in the six samples, resulting in six quantitative measurements per peptide per elution peak. For each peptide, differential abundance was then tested for each elution event using a t-statistic (Figure 2B, step 3. and exemplified for BAF53 in **Supplemental Figure S4B**). To obtain a significance measure of differential abundance of individual protein elution peaks, the median peptide level p-value of all peptides per protein per elution peak were integrated based on a scoring scheme assuming a beta distribution of the respective values, as described (Suomi and Elo, 2017). To generate the final differential protein association map, proteins are represented by the peak with highest fold-change and shifting proteins assigned based on cut-offs along the Benjamini-Hochberg-adjusted p-value (pBHadj score) and absolute SEC-localized fold-change (Figure 2B, step 3, right panel).

In the last step, differential association to chromatographic peaks detected per protein is then interpreted in the context of reference complexes to infer instances of protein complex remodeling using the complex-centric concept (Heusel, Bludau et al., 2019) (Figure 2B, step 4 and exemplified in **Supplemental Figure S4C**). The results are finally visualized and browsable via the web tool *SECexplorer-cc* as detailed further below. The extension of the *CCprofiler* toolset by the quantification module thereby supports the automated detection of altered protein association states and inferred protein complex remodeling from SEC-SWATH-MS data and enables the detection of altered protein complexes that are at the core of the present study.

### Benchmarking the differential SEC-SWATH-MS workflow and software tool using the mitotic dataset

The application of the method described above to the triplicate data obtained from two cell cycle states indicated substantial rearrangement of the proteome. Specifically, 2,189 SEC elution peaks of 1,793 proteins showed significant changes in abundance (pBHadj score ≤ 0.01, absolute SEC-localized fold-change ≥ 1.5, Figure 3A) and 1,626 shifts in the SEC elution range of assembled higher-order complexes. In the following, we further assessed the results at three levels: First, the technical reproducibility of data generation and analysis, second, recall of rearrangements of complexes known to be altered between cell cycle states and third, comparison of the results with those obtained from an orthogonal method from samples in comparable cell cycle states. In particular, we compared the differential SEC-SWATH results with results obtained from the parallel measurement of protein thermostability (Becher et al., 2018; Dai et al., 2018) from which changes in protein complexes were inferred.

**Figure 3:**
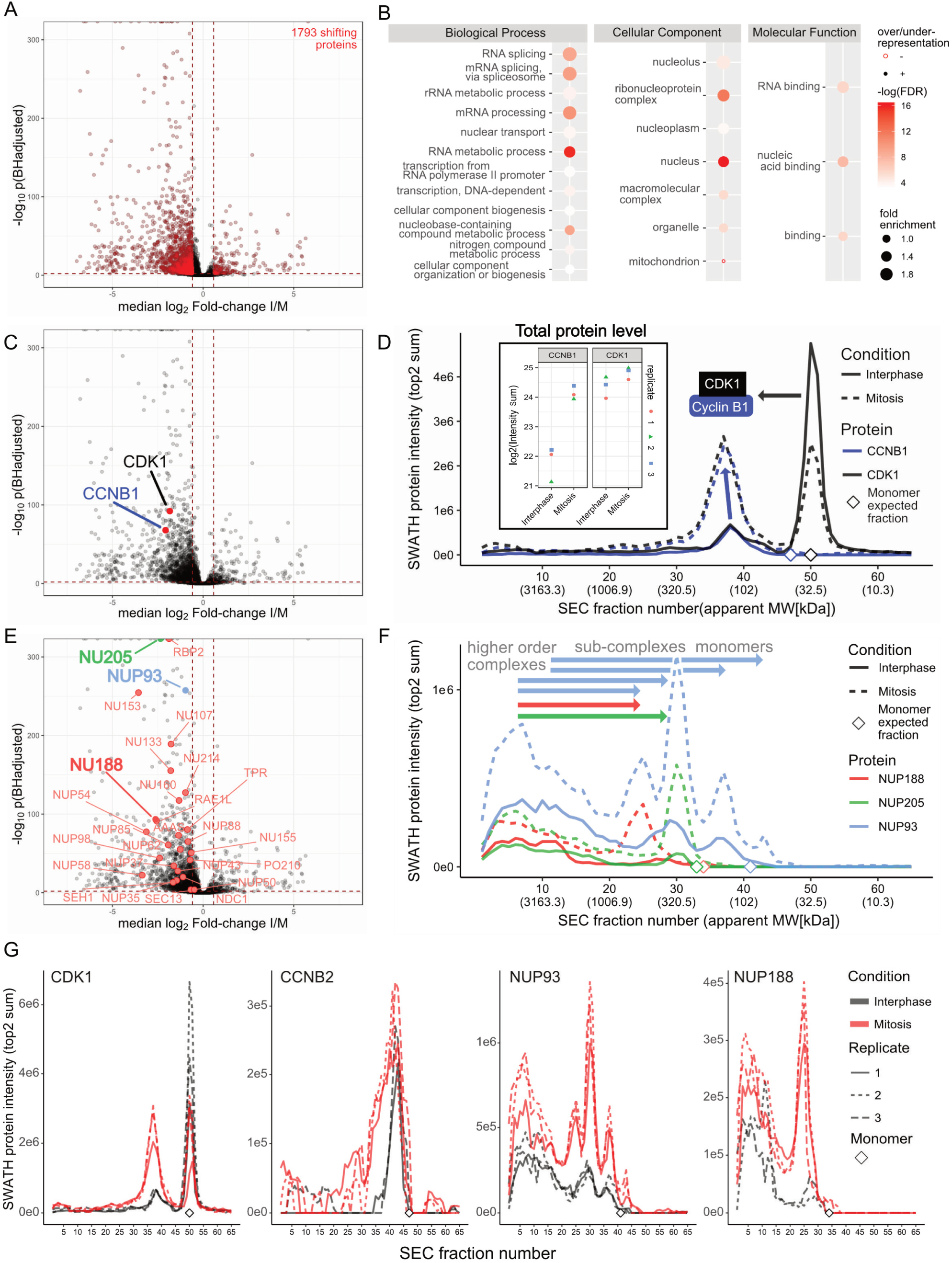
Benchmarking part 1: Recall of expected mitotic complex remodeling events. **A** Protein-level differential association score map highlighting 1,784 proteins shifting quantitative distribution across apparent sized assessed via SEC-SWATH. Benjamini-Hochberg adjusted P-value cutoff, ≤ 0.01; absolute SEC-localized fold-change ≥ 1.5 -fold. Reproducibility of SEC peptide and protein identification and quantification is assessed in Figure 1B-D, protein level error chromatograms (**Supplemental item 1**) and protein intensity correlation (**Supplemental Figure 2A**). **B** Gene ontology analysis of the total set of shifting proteins (PANTHERdb, against the background of all 5,044 proteins detected) suggests activity in processes related to cell cycle progression. **C** Protein-level differential association scores for a ‘true positive’ detection of an instance of mitotic complex remodeling among cell cycle regulators cyclin-dependent kinase 1 (CDK1) and cyclin B1 (CCNB1). **D** Conditional protein-level SEC chromatograms for CDK1 and CCNB1 capture and quantitatively characterize mitosis-specific recruitment of 69 % of the CDK1-derived MS-signal (right panel) to the CCNB1-assembled state. Inset: As expected, stable levels of CDK1 are observed while CCNB1 appears induced in mitosis (Total intensity observed across the 65 SEC fractions). Diamonds mark the SEC fraction expected for respective monomers based on naked sequence average molecular weight and external size calibration based on reference protein fractionation (compare Figure 1C). **E** Protein-level differential association scores for a second instance of ‘true positive’ detection of mitotic complex remodeling, nuclear pore complex disassembly. 26 of the 27 detectable subunits (out of 32 total, for assignment see experimental procedures) are detected to significantly shift their SEC elution patterns (as assigned in panel **A** and see main text and methods). **F** Conditional protein-level SEC chromatograms for an exemplary subset of protein components of the nuclear pore complex disassembling upon mitotic entry. Protein-centric differential association scores are highlighted in panel **E**. Chromatographic profiles of NUP188, NUP205 and NUP93 reveal protein mass re-distribution from high molecular weight NPC complexes (fraction 5, apparent MW ≥ 5 MDa) to lower MW signals representative of smaller NPC sub-complexes or likely a monomeric pool in the case of NUP93. Diamonds mark fractions where monomer elution would be anticipated.

#### Reproducibility

The availability of three replicate SEC-SWATH-MS measurements of either cell cycle state allowed us to assess the reproducibility of the method. Specifically, we evaluated technical cross-replicate variability at the level of i) size exclusion chromatography by the UV/Vis photospectrometric traces, ii) SWATH-MS by peptide and inferred protein identities and their relative abundance and iii) the overall SEC-SWATH-MS workflow by the reproducibility of protein-level SEC chromatograms. SEC fractionation was well-reproducible as is apparent from the UV absorbance profiles (λ = 280 nm) shown in Figure 1C. Further, SWATH-MS identified > 80% of detected peptides in all three replicates per cell cycle state and 48,380 peptides of 4,192 proteins in both cell cycle states. A total of 5,044 proteins were profiled across both cell cycle states (Figure 1B). SWATH-MS quantified proteins with good reproducibility as protein intensities were highly correlated across replicates and adjacent fractions with an average Pearson’s R > 0.98 between replicate fractions of the same biological condition (**Supplemental Figure S2B**). The reproducibility of protein abundance measurements was longitudinally affected by deteriorating mass spectrometer performance. However, these progressive effects were efficiently compensated by normalization based on reference spike-in peptides (**Supplemental Figure S2A** and see experimental procedures). The high degree of reproducibility achieved for the overall workflow was further apparent from protein-level SEC chromatograms reconstructed from the independent experimental repeats (See replicate SEC chromatograms of a select set of proteins given in Figure 3G and all protein-level chromatograms with error bars provided in **Supplemental Item 1** with source data in **Supplemental Table 1**). Overall, these metrics demonstrate the level of reproducibility of the SEC-SWATH-MS workflow towards the detection of differential protein associations between the two cell cycle states.

#### Recall of known biology of cell cycle states

Mitotic processes have been extensively studied. We therefore related the results of this study to known mechanisms of mitotic biology, first at the level of general patterns and second at the level of specific complexes. First, we calculated the over representation in gene ontology annotations of the 1793 proteins with shifts detected by SEC. In agreement with previous knowledge the results indicate cell cycle state-dependent changes in the functional groups “RNA splicing and mRNA binding events” (Dominguez et al., 2016), “cellular reorganization” (Gong et al., 2007) and “ribonucleoprotein and macromolecular complexes” (Linder et al., 2017) (Figure 3B). Second, we related the results obtained in this study to specific complexes that are known to be present at different assembly states in the cell cycle states tested (Vermeulen et al., 2003). The best confirmed and generally accepted events of this type include the mitotic activation of CDK1 by binding to its partner cyclin B1 (Gavet and Pines, 2010) and the mitotic disassembly of nuclear pore complexes (NPCs) (Linder et al., 2017). We found both events confirmed by the SEC-SWATH data. Figure 3C shows the rewiring of the CDK1 - cyclin B1 (CCNB1) module, supported by high differential association testing scores. Specifically, the size-calibrated subunit elution profiles closely reflected the formation of CDK1-CCNB1 complexes in the cell cycle state and indicated that CDK1 subunits of the CDK1-CCNB1 complex were recruited from the monomer pool, whereas the overall expression level of CDK1 across cell cycle states remained stable (Figure 3D). In contrast, CCNB1 subunits showed increased expression in mitosis (Figure 3D, insert), consistent with current models of CDK1 regulation by periodic expression of CCNB1 (Vermeulen et al., 2003). Notably, only part of mitotic CDK1 transitioned to the complex-assembled form, with 69% detected in the assembled and 31% of the total MS signal detected in the monomeric range. Further, the dataset confirmed the mitotic disassembly of NPCs (Figure 3E&F) with 26 of the 27 observed canonical subunits detected as SEC-shifting (out of 32 bona fide components as defined by (Hoelz et al., 2016)). Nucleoporin SEC profiles suggested protein mass re-distribution from a high molecular weight (MW) population of higher order nucleoporin complexes (fraction 5, void volume peak with apparent MW ≥ 5 MDa) to lower MW signals representative of NPC sub-complexes. This is exemplified by the SEC elution profiles of the inner ring complex members NUP188, NUP205 and NUP93 that are known to be part of mitotic sub-complexes of the NPC (Linder et al., 2017) (Figure 3E). Sub-complexes eluted in earlier fractions of elevated molecular weight compared to the respective monomers (Figure 3F, note monomer expected fraction markers). The observed profiles were highly reproducible across the experimental repeats (Figure 3G).

#### Validation via orthogonal method

A significant strength of the present method is its ability to quantify changes in protein complex abundance and chromatographic retention in a highly multiplexed manner. Recently, orthogonal methods have been described that assess thermal stability variation. In these methods changes in protein thermostability are used as a proxy for changes in protein interaction and activity. This notion is supported by the observation of strikingly similar thermostability profiles among subunits of the same complex (Tan et al., 2018). Two such studies explored altered thermostability across the cell cycle (Becher et al., 2018; Dai et al., 2018). They used chemical synchronization in early S and prometaphase, thus matching the biological conditions analyzed in the present study. We therefore compared the results obtained by the two orthogonal methods, SEC-SWATH-MS and thermal profiling, represented by two instances of the approach termed Cellular Thermal Shift Assay (CETSA) (Dai et al 2018) and Thermal Protein Profiling (TPP) (Becher et al, 2018).

First, we compared the proteome coverage achieved by the respective methods. The SEC-SWATH-MS dataset identified 5,044 proteins. Of these 4,515 showed detectable SEC elution peak(s) of which after statistical filtering 4,480 protein elution profiles were scored (for details on dropouts see experimental procedures). This number is comparable to that achieved by TPP (n = 4,780). CETSA achieved a markedly lower coverage at n = 2,773 proteins. More than 600 proteins were exclusively characterized by SEC-SWATH-MS (Figure 4A).

**Figure 4:**
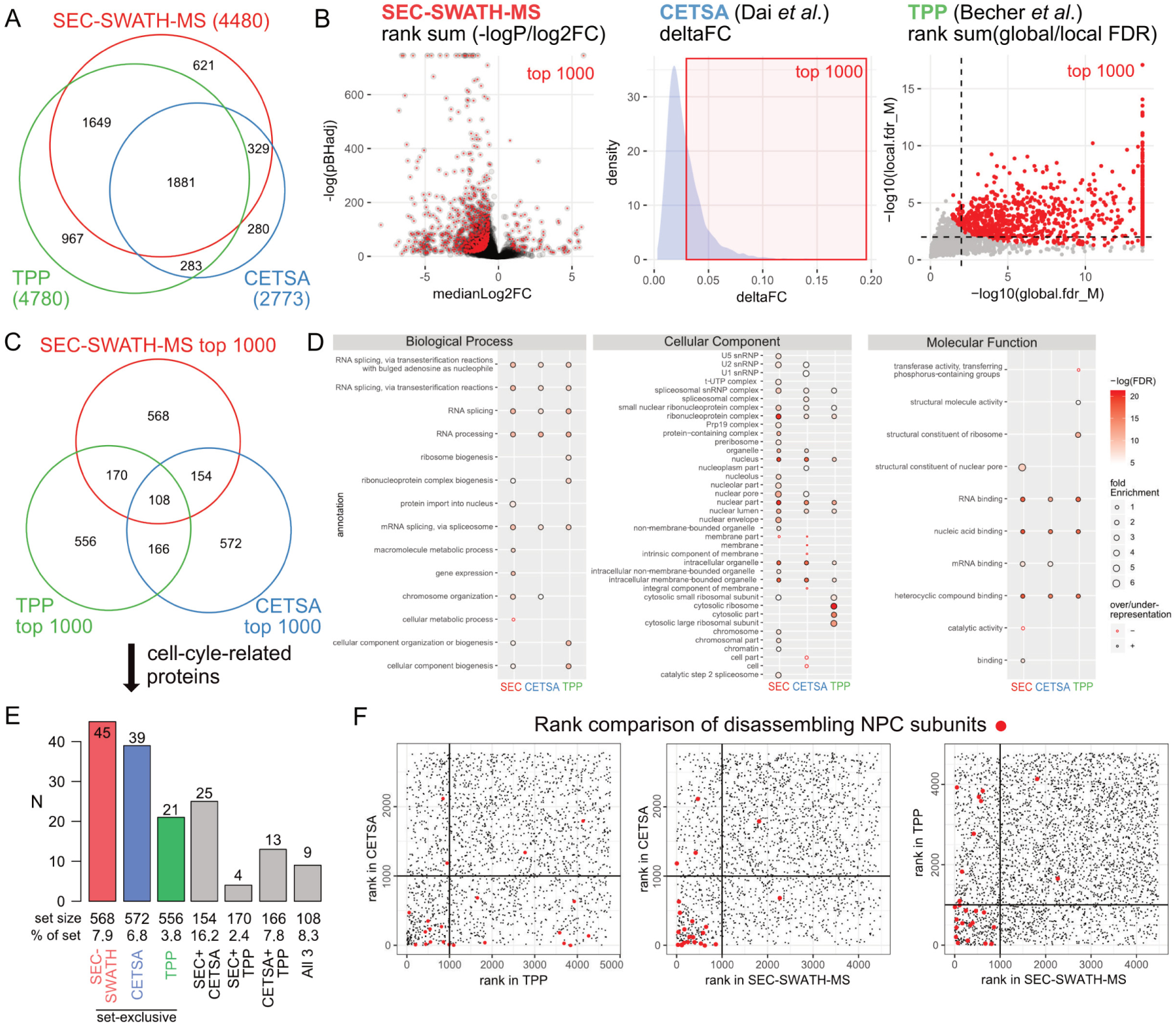
Benchmarking part 2: Performance of measuring proteome state dynamics by apparent size vs. thermostability. **A** Comparison of the total set of proteins characterized in SEC-SWATH (n = 4,480, this study), CETSA (Dai et al., 2018) (n = 2,773) and TPP (Becher et al., 2018) (n = 4,780). **B** Classification of top 1000 proteins to compare method performance. Proteins were ranked according to method-specific scores that intend to capture alterations in proteome association or thermostability state changes between cell populations chemically synchronized in distinct cell cycle stages (Comparison: interphase and prometaphase). All protocols employ thymidine block and nocodazole release experimental regimes. In SEC-SWATH-MS, top 1000 proteins are classified by linear combination of score ranks of equally weighted log_10_(pBHadj) and median log_2_ fold-change and selection along rank sum. Protein thermostability workflows employ two different scores (CETSA: delta fold-change; TPP: local and global FDR). The scores of the top 1000 proteins selected per method are indicated. To obtain protein ranks from the two scores in the TPP dataset, we combined the ranks along local and global FDR to select proteins along the rank sum. This procedure is equivalent to the protein ranking and selection procedure used for the SEC-SWATH-MS results. Dashed lines indicate the FDR cut-offs employed by the authors of the original study to select the 923 hits reported (Becher et al., 2018). **C** Comparison of top 1000 association- or stability-changing proteins detected in either approach shows degree of orthogonality and unexpectedly high dissimilarity of the protein sets reported to alter thermostability. **D** Gene ontology annotation overrepresentation testing of the top 1000 proteins per result set obtained from the three methods. Pathway enrichment is given in **Supplemental Figure S3**. **E** Comparison of method sensitivity based on method-exclusive recovery of ‘true positive’ proteins which function in relation to the cell cycle. Protein sets from Venn diagram in **C**. Cell-cycle-related proteins were defined by Uniprot annotation column ‘Function’ parsed on ‘cell cycle’. Of these 289 proteins, 156 were covered by one or multiple of the compared methods. Recovered numbers and rates with respect to set size suggest highest sensitivity of SEC-SWATH-MS and CETSA with broader proteome coverage of SEC-SWATH-MS (4,480 vs 2,773 proteins in SEC-SWATH-MS vs. CETSA). **F** Comparison of ranks of rearranging NPC component proteins in the three methods. Lowest rank means strongest signal in the respective method. SEC-SWATH-MS ranks truly re-arranging proteins highest.

Second, for each method we selected the 1000 proteins that showed highest scores indicating mitotic change and compared the observability of expected patterns of mitotic change within this set. For SEC-SWATH-MS the scoring was based on pBHadj/FC rank sum; for CETSA on deltaFC and for TPP on local FDR and global FDR rank sum (Figure 4B and methods section). For TPP, the selection of proteins along the combined rank sum included the majority of the proteins reported as hits in the original study (**Supplemental Figure S4C**). Unexpectedly, our analysis did not indicate higher similarity between the results of the two thermostability based studies than between either of these studies and the SEC-SWATH-MS derived data (274 proteins, 15.9%, shared between CETSA and TPP vs. 262 proteins, 15.1%, shared between CETSA and SEC-SWATH-MS and 278 proteins, 16.1 %, shared between TPP and SEC-SWATH-MS, Figure 4C). Whereas the three sets of top ranked proteins showed a small overlap of 108 of 2,294 proteins, the functional and pathway enrichment patterns of the three protein sets were in good agreement. All workflows uncovered strongest activity changes in RNA processing and splicing processes and corresponding ribonuclear complexes and the RNA binding machinery (Figure 4D).

The differential SEC-SWATH-MS analysis preferentially retrieves proteins in the GO categories “nuclear transport”, “proteins forming complexes” and “proteins of the nuclear envelope” including components of the nuclear pore, while TPP exclusively retrieved proteins associated with the ribosomal machinery. Both, SEC-SWATH-MS and CETSA detected proteins associated with chromosome segregation, whereas this activity was not detected by TPP. Membrane proteins appear underrepresented among the hits reported by SEC-SWATH-MS and CETSA, but not TPP. Similarly, metabolic and enzymatic functions appear slightly underrepresented in hits from SEC-SWATH-MS and TPP, but not CETSA (Figure 4D).

SEC-SWATH-MS exclusively showed a tendency to uncover proteins known to engage in binding activities (Figure 4D, Molecular Function). Pathway enrichment analysis showed a higher number of similar enrichments between SEC-SWATH-MS and CETSA and a more strongly diverging pattern for the TPP results (**Supplemental Figure S3A**). To control for biases introduced by our rank-based selection of the 1000 most regulated proteins from the TPP results, we also included the hit list reported in the original study (897 Uniprot proteins mapping unambiguously to the 923 reported hit gene names (Becher et al., 2018), Table S2, see **Supplemental Figure S3B** for a comparison of the selected protein sets). Specifically, SEC-SWATH-MS and CETSA both showed significant enrichment of pathway terms “Cell Cycle / Mitotic Cell Cycle” and only SEC-SWATH-MS captured altered functional states of the core cell-cycle regulator anaphase promoting complex APC/C and activation of the mitotic spindle assembly checkpoint SAC (**Supplemental Figure S3A**). To explore potential differences in the protein activities inferred from the protein properties measured by the different methods (stability or size), we analyzed the biological process annotation of protein sets reported exclusively by SEC-SWATH-MS (n = 568) and by both stability-based approaches (n = 166) (**Supplemental Figure S3C**), respectively. Interestingly, the different protein sets converge at the level of core regulated processes (RNA splicing & -processing) and highly related processes. For instance, both protein sets point towards the assembly of protein complexes, whereas changes in thermal stability were observed preferably in proteins of ribosome and ribonucleoprotein complex biogenesis and changes in assembly state and size were observed in proteins involved in protein-containing complex subunit assembly and - organization. The measurement of both, thermal stability and size indicated alterations in proteins from different metabolic processes, of which alterations in the category “organic substance metabolic processes” were detected by either method. This may suggest that alterations in metabolic processes can manifest in either protein stability or the assembly state of complexes or both properties.

Third, we evaluated the recovery of proteins with known function in the cell-cycle process in the respective datasets (UniProt functional annotation parsed on ‘cell cycle’, Figure 4E). The SEC-SWATH-MS data showed the highest sensitivity for the measurement of altered protein (association) states among the compared methods (Figure 4E, comparing sets from Venn diagram in Figure 4C).

Fourth, we compared the sensitivity of the methods to recapitulate known biochemical events of mitotic disassembly of NPCs. In this comparison, SEC-SWATH-MS showed the highest degree of sensitivity as nucleoporins were ranked highest in the priority lists compared to the priority lists generated by the other methods (Figure 4F). This comparison validates the SEC-SWATH-MS differential workflow to generate biological insights similar to those obtainable via CETSA and at extended proteome coverage comparable to that achieved in the TPP workflow.

Overall, these three levels of benchmarking showed high performance of the SEC-SWATH-MS differential workflow, including the extended *CCprofiler* tool, to reveal altered protein association states in biological samples with high sensitivity and broad proteomic coverage. In addition, the chromatographic profiles contain extended layers of information such as the specific composition and abundance of distinct complexes and also indicate quantitative changes in protein abundance.

### Inference of cell-cycle dependent complex remodeling

We used the quantitative, complex-centric SEC-SWATH-MS technique to detect changes in complex quantity and composition between cell cycle states. These analyses are based on the SEC profiles of 4,515 proteins forming 6,040 distinct peaks (*CCprofiler* protein-centric q-value = 5%, see methods for details) and constitute a global ‘master map’ set of observable protein features across replicates and conditions (See Figure 2B and **Supplemental Figure S4A-B**). The chromatographic elution profile of each protein in the master map was analyzed with respect to the following dimensions of information. First, the number of peaks in the chromatographic range covering complex associated proteins indicated a minimal number of distinct complexes a protein was associated with. Second, changes in chromatographic elution between conditions identified proteins with significant changes of complex association, and third, the correlation of peaks in the elution patterns of different proteins confirmed the presence of specific complexes by complex-centric analysis. The data indicated that proteins observed in two or more distinct complex-assembled states were enriched in signaling factors, proteins with known binding functions and proteins involved in modulating post-translational modification such as acetylation and phosphorylation (**Supplemental Figure S4D**). The data further indicated that in most cases in which a protein was associated with different complexes the SWATH-MS signals for independent peptides strongly correlated between peaks, indicating overall very robust signal quality. Outlier peptides, i.e. peptides for which the between-peak correlation deviated from the correlation of other peptides, likely indicate cases of post-translational modifications resulting in the differential association of proteoforms to different complexes.

The ensemble of protein SEC elution profiles further provided a base to estimate the fraction of the proteome detected in monomeric or assembled form in either state. We designated proteins as detected in an assembled state if their apparent MW based on the SEC elution was minimally twofold larger than the predicted MW of the protein in monomeric form. The protein SEC elution profiles were interrogated from two perspectives. First, we performed a naïve assignment of protein intensity to either complex-assembled or monomeric state. To make these assignments, we used the MS signals of the two most-abundant proteotypic peptides per protein. In line with previous observations of the HEK293 proteome assembly state via SEC-SWATH-MS (Heusel, Bludau et al., 2019), the major fraction of the interphasic and mitotic HeLa CCL2 proteome mass was observed in complex-assembled state (57 ± 6% and 58 ± 4%, respectively). In terms of protein numbers 70 ± 2% and 72 ± 2% of the proteins were detected at least in part in complex assembled state in interphase and mitosis, respectively.

To next explore differences in protein profiles observed between conditions, we applied protein-centric peak detection per each cell cycle state. To increase signal-to-noise we merged the three replicates and detected 5,291 and 5,637 distinct elution peaks for 4,083 and 4,264 proteins in interphase and mitosis, respectively. Based on the apex SEC fraction and associated apparent molecular weight, the 1-5 elution peaks observed per protein were assigned to likely assembled or monomeric pools of the total protein population (Figure 5A). According to these assignments, 29 and 25% of proteins were observed eluting exclusively in monomeric form, while 71 and 75% of proteins were observed in at least one complex-assembled form in interphase and mitosis, respectively. A significant fraction of proteins was further observed eluting in both, monomeric and complex assembled form(s) (12% in both interphase and mitosis, respectively). These results indicate that in either cell cycle state a significant fraction of the proteome was associated with complexes that are accessible to differential quantification of protein association state changes via SEC-SWATH-MS and *CCprofiler*.

**Figure 5:**
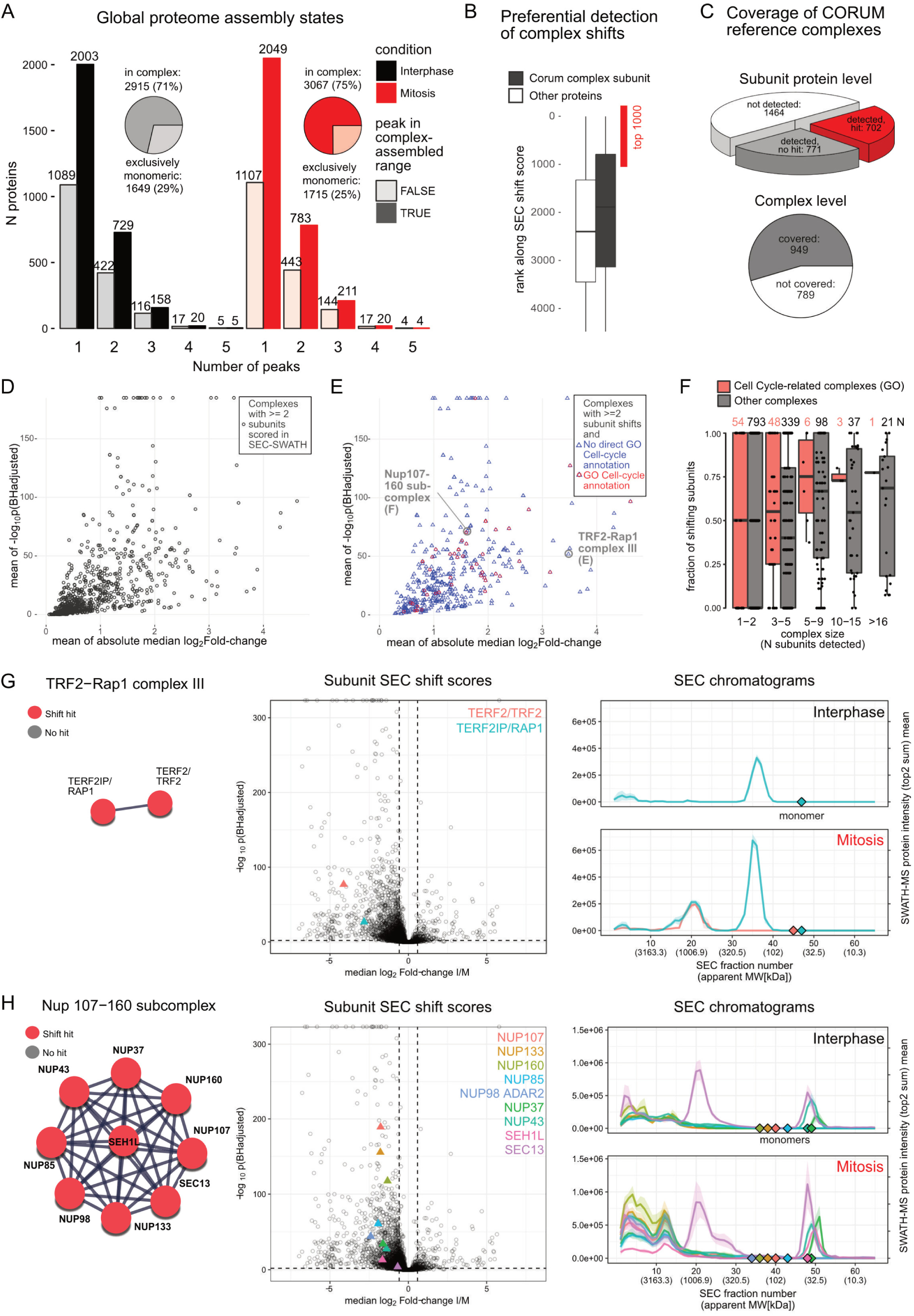
Inference of cell-cycle dependent complex remodeling. **A** Global proteome assembly states observed in interphase and mitosis. Bar plots show numbers of proteins eluting in one to five distinct peaks. Pie charts show that the majority of proteins peak at least once in the likely complex-assembled range (apparent molecular weight twice or larger than the annotated monomer molecular weight). In either cell cycle state, 12% of the proteins are observed in both monomeric and complex-assembled state and ca. 30 % of the proteins elute in two or more distinct peaks, in line with previous observations on the HEK293 proteome modularity profiled by SEC-SWATH-MS (Heusel, Bludau et al., 2019). Peak detection was strictly error-controlled (q-value FDR estimate of 5 %) against randomized peptide-to-protein associations. For details, see experimental procedures. **B** Preferential detection of shifts in proteins that are subunits of known reference complexes of the CORUM database displayed based on the proteins SEC shift score ranks. Proteins that under certain conditions integrate into complexes display lower ranks, with rank 1 representing the highest SEC shift score observed. **C** Coverage of CORUM reference complexes in SEC-SWATH-MS. Upper chart: Coverage on the level of complex component subunits. Lower chart: Coverage on the level of complexes (covered if two or more of the annotated subunits were among the SEC-SWATH-MS results). **D** The 949 complexes covered in the dataset with two or more subunits. For visualization purposes, the complexes are represented by the complex-level means of subunit-level differential SEC shift scores. **E** The 432 complexes that were detected to undergo remodeling in mitosis vs. interphase, based on significant SEC shifts of minimally two of their component subunit proteins. For visualization purposes, the complexes are represented by the complex-level means of subunit-level differential SEC shift scores. The coloring indicated whether the complex is annotated with “cell cycle” in the database-contained gene ontology terms (GO). **F** For the changing complexes, the fraction of shifting subunits (of those detected by SEC-SWATH-MS) was plotted as function of complex size (n subunits, detected by SEC-SWATH-MS) and whether or not the complex bears the GO annotation ‘cell cycle’. Shift completeness is higher among cell cycle-related assemblies. **G** Example of a complex that is remodeling along the cell cycle states but not annotated with ‘cell cycle’, TRF2-Rap1 complex III (CORUM ID 1205), showing a representative spokes model, subunit protein level SEC-shift scores in the context of all observed shift scores and their quantitative elution along SEC as profiled by SWATH-MS. Both subunits display significant SEC shifts. The analysis detects a co-elution signal indicating the presence of a complex of ca. 950 kDa (Apex fraction 21) in mitosis but not in interphase, where only RAP1 is detected. **H** Equivalent to panel G. Second example of a complex that is remodeling along the cell cycle states but not annotated with ‘cell cycle’, Nup107-160 subcomplex (CORUM ID 87), showing a representative spokes model, subunit protein level SEC-shift scores in the context of all observed shift scores and their quantitative elution along SEC as profiled by SWATH-MS. All 9 subunits display significant SEC shifts. The Nup107-160 sub-complex appears specifically in mitosis and with an apparent molecular weight of 2.8 MDa (Apex fraction 13, right panel). Subunit SEC13 shows an additional peak at ca. 10 MDa (Apex fraction 20-21) where it appears bound to its partner subunit in its alternative context in the COPII complex, SEC31 (**Supplemental Figure S6**).

We further used the ensemble of protein patterns to determine which proteins and associated functions displayed a change in protein complex association between the two cell cycle states tested. The data indicated substantial rearrangement of the proteome. Specifically, 2,189 SEC elution peaks of 1,793 proteins showed significant abundance shifts. Of these, 1,626 shifts were in the SEC elution range of complex-assembled proteins suggesting significant rearrangements in the underlying complex(es) (Figure 3A, pBHadj score ≤ 0.01, absolute SEC-localized fold-change ≥ 1.5). Proteins with altered complex association states were predominantly associated with functions in “transcriptional and splice regulatory machinery” and “cellular component organization” (Figure 3B). The 1000 top-ranking proteins selected for comparative benchmarking analyses further revealed reorganization in the MAPK cascade (Figure 4D, Biological Process) and rearrangements involving central cell-cycle-associated modules such as the APC/C, NPC and mitotic spindle checkpoint and mitotic anaphase pathways, among others (**Figure S3A**).

Whereas these enrichment analyses already pointed at specific complexes undergoing mitotic change, we next evaluated our results using the CORUM reference set of complexes as prior information (Ruepp et al., 2010). Initially, we evaluated the rearrangements detected by SEC-SWATH-MS that occur preferentially among protein complexes with known involvement in mitotic processes. Indeed, we detected a higher frequency of changed patterns of proteins that are known to associate into complexes, compared to a control group not known to be affiliated with a complex (Figure 5B). On the level of individual proteins, the SEC-SWATH-MS dataset covered over half of the subunits of the reference CORUM complexes (1473 of 2937). Of these, 702 showed significant changes in protein complex association between states (Figure 5C, upper panel). More than half of the reference complexes were detected based on two or more subunit proteins in our analysis (n = 949, Figure 5C, lower panel and Figure 5D). Overall, 432 complexes showed evidence of remodeling based on significant SEC shifts observed for minimally two of their subunits (Figure 5E). Among these changing complexes, we observed preferential and more complete detection of changes in complexes with known involvement in the cell cycle (Figure 5F). While many of the complexes with evidence of remodeling are associated with known functions in relation to the cell cycle, a significant fraction is not and thus present opportunities for further exploration. The remodeled complexes and their functional annotation are summarized in **Supplemental Table 2**.

The results in Figure 5G&H illustrate two complexes without GO annotation for “cell cycle” but with strong evidence for mitotic changes. For both complexes all CORUM subunits were detected. The first is the cell-cycle-dependent assembly of the TRF2-RAP1 complex in mitosis observed by strong SEC shift scores of both subunits and co-elution of RAP1 and TRF2 with apex in SEC fraction 21 (ca. 950 kDa) in mitotic cells but not interphasic cells. TRF2 was detected only in mitotic cells. In interphasic cells, RAP1 and showed no peak in the SEC fraction range of the TRF2-RAP1 complex (Figure 5G). The results are in line with the previously reported recruitment of RAP1 to telomeres via TRF2 and consistent with the co-elution of the two proteins in the void volume (fractions 1-5, analytes >10 MDa) (Figure 5G, right panels). However, RAP1 was detected in the void volume of interphasic cells but without the concurrent detection of TRF2. Notably, we did not observe the TRF2 and RAP1-containing shelterin complex known to occupy telomeres in interphase (Liu et al., 2004), likely due to low abundance and/or low recovery in mild lysis of interphase cells. In either cell cycle state, RAP1 was observed with an apparent molecular weight of ∼180 kDa, in line with preassembly into the tetrameric form in which it participates to form the octameric TRF2-RAP1 complex composed of four copies of each protein (Arat and Griffith, 2012). In mitotic cells, the strongest peak group containing RAP1 and TRF2 centered around 950 kDa, ca. 2.5x the weight expected for TRF2-RAP1 hetero-octamers, suggesting the association of the octamer with as yet unknown proteins. Notably, RAP1 has been known to associate with the I-kappa-B-kinase (IKK) complex to enhance NF-kappa-B target gene expression (Teo et al., 2010). To test whether the observed 950 kDa signal reflected the RAP1 pool likely engaged in an interaction with the IKK complex, we considered in addition the elution profiles of IKK subunits CHUK and IKKB (**Figure S5A**). The CHUK and IKKB profiles suggest the presence of two distinctly sized and only partly SEC-resolved variants of the IKK complex, one of ca. 2.5 MDa (apex fraction 12) and one of ca. 1.7 MDa (apex fraction 16) (**Figure S5A**). The majority of the RAP1 signal at ca. 950 kDa appeared independent from the two distinct populations of IKK complex variants observed. However, a peak shoulder in the RAP1 signal at elevated molecular weight (estimated apex fraction 16) conformed with a small fraction of RAP1 bound to the 1.7 MDa but not the 2.5 MDa variant of the IKK complex (**Figure S5A,** lower panel, fractions 14-18).

As a second example, we observed striking changes in the elution profile of the Nup107-160 sub-complex of the NPC (Corum ID 87, Figure 5H). All 9 subunits were detected with significant SEC shifts (Figure 5H, middle panel). In mitotic cells, the complex was observed based on a co-elution peak group formed by all subunits at a molecular weight of 2.8 MDa (Apex fraction 13, Figure 5H, right panel). In interphase cells, no defined co-elution peak was detected in this size range, suggesting the presence of the Nup107-160 sub-complex exclusively in mitotic extracts. Interestingly, the subunit SEC13 was observed in a second peak at ∼ 1 MDa in both cell cycle states, suggesting its presence in an additional complex resolved by SEC. We surmised that this peak may represent SEC13 in the context of its alternative functional role in COPII vesicle-mediated transport (Tang et al., 1997). To test this hypothesis, we overlayed the elution profiles of the SEC13 partners in the coatomer complex and, indeed, observe co-elution with its partner SEC31 but not with the adaptor proteins SEC23A/B and SEC24/B (See **Supplemental Figure S6**). These observations demonstrate the capacity of our method to capture mitotic liberation of Nup107-160 sub-complexes from NPCs in and to resolve protein engagement across different functional contexts. Mitotic disassembly of NPCs is a hallmark of mitotic progression (Linder et al., 2017) but this event has not been annotated in the respective GO terms. This insight into complex dynamics at sub-complex resolution led us to explore whether SEC-SWATH-MS chromatographic profiling could reveal further and potentially novel aspects of mitotic NPC disassembly (see below).

In summary, these results show that a major portion of the proteome changes at the level of complex association between mitotic states and that hundreds of specific complex remodeling events were apparent from the data. Insights at sub-complex resolution warrant in-depth analysis of chromatographic profiles not only for newly implicated proteins but may also reveal novel or additional roles of proteins with known functions in cell cycle progression.

### Discovery and independent validation of NPC disassembly intermediates

The discovery of compositional rearrangements of protein complexes between cell cycle states allowed us to propose testable changes of biochemical processes. Among these is the mitotic disassembly of the NPC (Hoelz et al., 2016). The ensemble of SEC protein profiles analyzed in this study contained 27 of the 32 bona fide NPC components (Hoelz et al., 2016), shown in Figure 6A in either state. The patterns show a general, distinctive change towards complexes of lower MW in the mitotic sample, consistent with NPC disassembly into sub-complexes. All but one of the detected subunits (NUP50) showed shifts in protein-centric differential scoring. For eight of the subunits, elution peaks in the monomeric range were detected, equally across both cell cycle states. The observation of monomeric pools in the SEC experiment may indicate the presence of a subset of NPC components that seem to be present in the cell as assembly-competent monomeric forms, to potentially bind other partners to fulfill other functions or liberated from partner Nups during preparation of the cell extracts.

**Figure 6:**
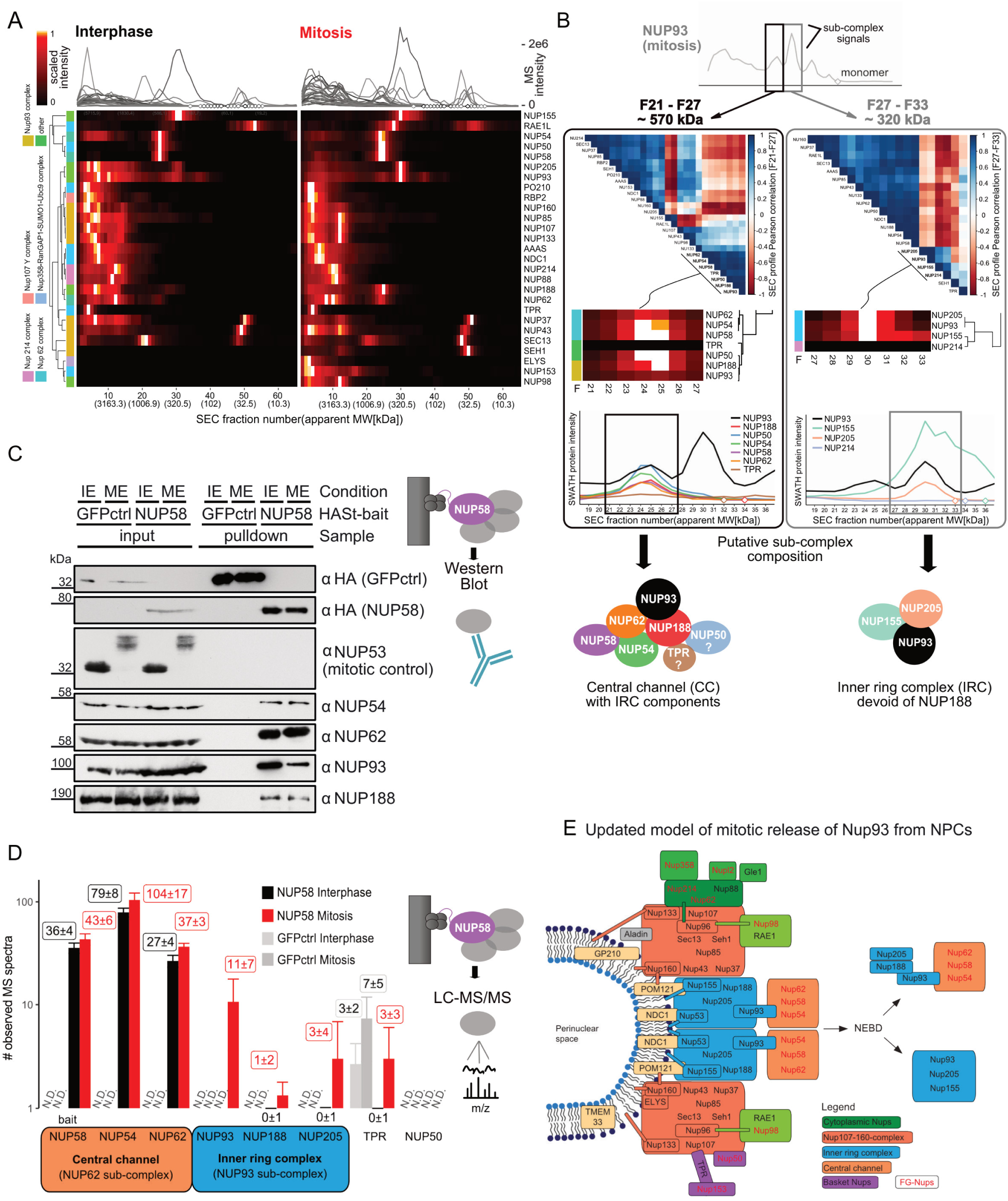
Discovery and independent validation of NPC disassembly intermediate. **A** Quantitative elution patterns of NPC subunits in interphase and mitosis display protein mass re-distribution from large nucleoporin complexes in interphase with void volume elution (> 5MDa) towards lower MW ranges larger than individual components, in line with NPC disassembly into defined sub-complexes. Also compare Figure 3F. **B** Targeted search for co-eluting proteins as candidate co-complex members, using as basis two of the mitotically induced sub-complex signals arising from the inner ring complex component NUP93. Left panel: MS signal correlation analysis in the first peak (apex fraction, 25; apparent MW, ca. 570 kDa; elution range, F21-F27) nominates candidate members of the sub-complex eluting in signal 1 based on co-elution. Right panel, co-elution-based nomination of additional sub-complex membership candidates for signal 2 (apex fraction, 30; apparent MW, 320 kDa; elution range F27-F33). Bottom, putative sub-complex composition subjected to validation experiments (see panels C&D). Note that targeted search for co-eluting proteins (on global dataset scale) is a core function of our online data interrogation tool SECexplorer-cc. **C** Testing mitotic sub-complex composition based on co-purification with central channel complex captured via NUP58 by immunoblotting. As mitotic control NUP53 was included and complete size shifts indicate high homogeneity of cell cycle state of the analyzed cell populations in interphase (IE) and mitosis (ME). As negative control to control for non-specifically bound background protein, green fluorescent protein was included. Both NUP93 and NUP188 co-purify with the central channel in interphase and continue to do so in mitosis, confirming the results from SEC-SWATH-MS. **D** Testing mitotic sub-complex composition based on co-purification of Nups with the central channel complex captured via NUP58 by mass spectrometry (AP-MS). The number of identified mass spectra serves as semi-quantitative measure to estimate protein retrieval from cells in either cell cycle state and respective controls. The results confirmed the co-purification of NUP93 and NUP188 and also showed the presence of the inner ring complex component NUP205. which is detected in SEC but does not form a defined elution peak in the signal 1 range. Neither TPR nor NUP50 are recovered in amounts above background binding level and are thus likely not part of the mitotic NPC sub-complex. **E** Model of mitotic NPC disassembly and storage of NUP93 in distinct mitotic sub-complex reservoirs before re-formation of daughter cell nuclear envelopes. A substantial fraction of NUP93 is stored in the newly identified mitotic sub-complex composed of central channel components NUP54, NUP58 and NUP62 as well as inner ring complex components NUP93, NUP188 and NUP205.

Due to the distinctive pattern changes between the two states tested, we focused on the key inner ring complex component NUP93, which functions as an adaptor between the NPC scaffold and the central channel FG Nup62-Nup58-Nup54 sub-complex. In interphase extracts, NUP93 was principally detected in a wide high MW peak (fractions 1-15, > 10 MDa to ca. 1.8 MDa) apart from minor amounts detected in two lower MW populations (apex fraction 29, ∼ 360 kDa and apex fraction 37, ∼145 kDa). In the mitotic state, NUP93 was detected in three distinct lower MW SEC peaks, with a new signal at an apparent MW of ca. 570 kDa (elution range F21-F27) and two signals of increased intensity when compared to the interphase pattern at apparent MW of ca. 320 kDa (elution range F27-F33) and 145 kDa (elution range F35-F39) (Figure 3F, **Supplemental Figure S5B** and schematically illustrated in Figure 6B). The observed peak pattern indicates that in mitotic cells NUP93 associates with complexes of different size that elute markedly earlier than monomeric NUP93 (93.5 kDa, expected elution fraction 41), thus suggesting the formation of distinctive NUP disassembly intermediates containing NUP93. To infer the composition of these complexes, we locally correlated the elution pattern of NUP93 with the elution patterns of other Nups (Figure 6B). For the peak at ca. 320 kDa, this analysis suggested NUP205, NUP155 and NUP214 as NUP93 interaction partners (F27-33, Figure 6B, right panels). Similarly, for the peak at ca. 570 kDa, (Figure 6B left panel), six proteins, namely the central channel FG Nups NUP62, NUP54 and NUP58 as well as NUP188, NUP50 and TPR eluted in the same peak as NUP93.

The relative mass spectrometric signal intensities of peptides can be used to estimate the abundance of a protein in a SEC peak (Ludwig and Aebersold; Rosenberger et al., 2014). For the peak at ca. 320 kDa quantification of the respective protein signal intensities identified the inner ring proteins NUP155 and NUP205 as the predominant binding partners of NUP93. In contrast, NUP214, whose quantitative pattern correlates well in the SEC dimension, is present at much lower signal intensity which is not consistent with stoichiometric participation in the complex. The results thus suggest a complex dissociating from the NPC holo-complex in mitosis that is composed of NUP93, NUP155 and NUP205. Notably, NUP188 displays only low local correlation with NUP93 in the queried target range, suggesting that NUP188 does not participate in the complex of ∼320 kDa detected between F27 and F33 (Figure 6B, right panel). The observed MW of 320 kDa is smaller than the cumulative weight of a stoichiometric hetero-trimer (476 kDa). This discrepancy could be due to compact shape and/or interactions with the stationary phase. Similarly, we used the protein intensities to also estimate composition and the abundances of the proteins in the peak at ca. 570 kDa (Figure 6B left panel). The data suggest that in this complex the inner ring component NUP93 was associated with the central channel sub-complex NUP62, NUP54 and NUP58, as well its inner ring complex partner subunit NUP188. Based on the absolute signal intensity, stoichiometric participation of TPR in the ∼570 kDa assembly appears unlikely. These findings are supported by current models of NPC structure (Beck and Hurt, 2017; Hoelz et al., 2016; Lin et al., 2016) in which central channel Nups are coordinated by adaptor Nups of the inner ring complex. Based on the holo-complex model, the recovery of both TPR and NUP50 as part of NUP93-containing complexes appears unlikely, whereas interaction between NUP93 and central channel appears probable because NUP93 serves as anchor point for the central channel within the NPC holo-complex (Chug et al., 2015; Hoelz et al., 2016; Lin et al., 2016).

We validated the existence of the previously unknown mitotic sub-complex consisting of NUP93, NUP188 and the trimeric central channel sub-complex NUP62-NUP58-NUP54 by co-precipitation coupled to immunoblotting or mass spectrometry as orthogonal methods. We inducibly expressed HA-St-tagged NUP58 in HeLa cells synchronized in either interphase or mitosis. As a control for the completeness of mitotic arrest, we performed immunoblots of NUP53 in the input samples, demonstrating its efficient mitotic hyper-phosphorylation, as previously reported (Linder et al., 2017). We isolated the native complex associated with the tagged Nup58 under mild conditions and tested the isolate for the presence of the suggested complex components by immunoblotting and LC-MS/MS in either cell cycle state (Figure 6C&D, **Supplemental Figure S1D & E**). Indeed, immunoblotting confirmed co-purification of both the immediate partner Nups (NUP62, NUP54) and the inner ring complex components NUP93 and NUP188 with NUP58 from interphase cells. Importantly, in mitosis, this connectivity was maintained, in agreement with our SEC-SWATH-MS results (Figure 3C). These data confirm the presence of a mitotic sub-complex involving both central channel and inner ring complex components NUP93 and NUP188. The observed relative signal intensities in immunoblotting show a reduced recovery of NUP93 with the central channel Nups in mitosis compared to interphase. This is consistent with mitotic partitioning of the NUP93 protein pool across multiple macromolecular entities, as indicated by SEC-SWATH-MS (Figure 3C, rightmost two lanes and compare Figure 6B, right panel).

Next, we analyzed proteins co-isolated with Nup58 by mass spectrometry and used the number of identified mass spectra as semi-quantitative measure to estimate protein retrieval from cells in either cell cycle state and respective controls. The results further confirmed the co-purification of NUP93 and NUP188 and also showed the presence of the inner ring complex component NUP205 (Figure 6D). Notably, NUP205 was consistently detected in the SEC elution range under investigation (F21-F27) but did not show a distinctive co-elution peak. Neither NUP50 nor TPR were detected as significant components of the isolates and are unlikely to represent *bona fide* components of the new sub-complex. (Figure 6D). Thus, Nup50 is likely part of a different, independent protein complex eluting at a similar position in SEC, which can motivate future research on the mitotic fate of this Nup.

Together, these results indicate that, in contrast to preceding models of NPC disassembly, a fraction of the inner ring complex components NUP93 and NUP188 and likely NUP205 remain attached to the central channel sub-complex composed of NUP62, NUP58 and NUP54 after mitotic entry. This configuration can explain how central channel Nups are efficiently reincorporated into reforming NPCs along with their partner scaffold Nups during mitotic exit. Further, a second population of NUP93 is stored in a mitotic sub-complex with the canonical IRC components NUP155 and NUP205 in absence of NUP188 (See model in Figure 6E).

### Browsing dynamic complex association maps in SECexplorer-cc

SEC-SWATH-MS portrays the process of mitosis from the angle of protein mass re-distribution across differently sized stable complexes resolved by SEC. There remains a lot to be learned from the rich dataset generated, with the prospect of discovering new proteins involved in cell cycle regulation and of identifying new components of both static as well as dynamic assemblies. To support community-based mining and interpretation of our dataset, we provide a web tool, *SECexplorer-cc* (https://sec-explorer.shinyapps.io/hela_cellcycle/, Figure 7) which offers four functionalities: (i) Interactive viewing of protein SEC fractionation profiles in interphase and mitosis (compare Figure 3D and F). (ii) Search for locally co-eluting proteins to identify putative new binding partners showing strong co-elution within a certain range of target protein elution (compare Figure 6B). (iii) Interactive display and protein selection from the differential association score map (compare Figure 3A). (iv) Display of one or multiple protein’s fractionation profiles in reference to the profiles of immediate interaction and/or functional partners dynamically retrieved from StringDB (Szklarczyk et al., 2017). We expect that *SECexplorer-cc* will support a community effort to fully leverage the rich information encoded by the mitotic proteome rearrangement SEC-SWATH-MS data, which, ideally, will support better understanding of modular proteome function along cell division.

**Figure 7:**
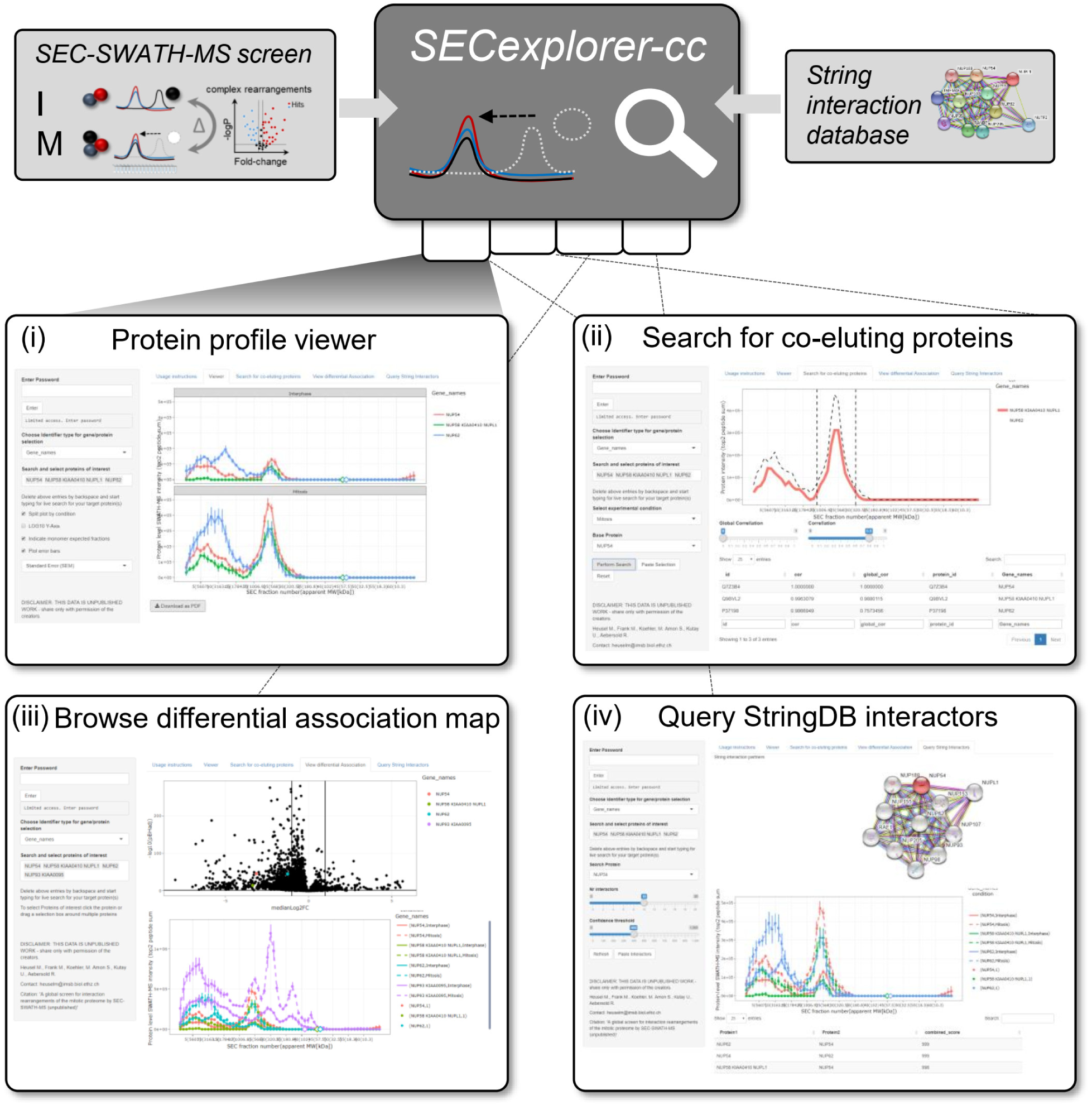
Browsing dynamic complex association maps in SECexplorer-cc. Overview of *SECexplorer-cc*, a web tool that allows to browse and rapidly interrogate differential SEC-SWATH-MS datasets to support researches in customized hypothesis testing. SECexplorer-cc currently combines the dynamic cell-cycle proteome association map reported here and integrates it with information from the String database (Szklarczyk et al., 2017). *SECexplorer-cc* features four core functionalities. **(i)** Interactive display and hit protein selection from the statistical SEC shift score map. **(ii)** Semi-targeted search for locally co-eluting proteins to identify putative new binding partners showing strong co-elution within a certain range of target protein elution. **(iii)** Display of custom protein sets conditional fractionation profiles. **(iv)** Display of one or multiple hit protein’s fractionation profiles in reference to its immediate functional and/or physical interaction partners obtained StringDB to discover potential physical association among functionally related proteins and to add cellular context information.

## Discussion

The biochemical state of a cell is the result of the ensemble of biochemical functions which in turn are largely carried out by macromolecular complexes. Changes in the quantity, composition, subunit topology or structure of such complexes are thus assumed to reflect altered functional states of the cell and the systematic detection of such alterations is therefore of great interest for basic and translational research.

In this study we introduce a new software module to the *CCprofiler* suite (Heusel, Bludau et al, 2019) that supports the differential, quantitative analysis of thousands of proteins and their association with complexes from SEC-SWATH-MS datasets, based on a complex-centric analysis strategy. We benchmarked the tool and applied it to detect rearrangements protein complexes in a differential analysis of human cells in two chemically induced cellular states, interphase and mitosis/early prometaphase. The described technique is based on the co-fractionation and protein correlation profiling rationale (Dong et al., 2008; Foster et al., 2006; Liu et al., 2008; Wessels et al., 2009) and includes the improvements with regard to chromatographic resolution, accurate data-independent mass spectrometry (Gillet et al., 2012; Röst et al., 2014) and complex-centric data analysis strategy implemented in SEC-SWATH-MS (Heusel and Bludau et al., 2019). Through the addition of the quantification module, the workflow now supports the systems-wide differential analysis of > 5,000 of proteins and their association with complexes. The data show that the system shows a high degree of reproducibility and performs favorably when benchmarked against protein thermostability measurements, an orthogonal state-of-the-art method to infer changes in protein complexes (Becher et al., 2018; Dai et al., 2018). Application of the method to HeLa CCL2 cells recapitulated known cell cycle-dependent complex remodeling events between the cell cycle states tested and suggested a new model of nuclear pore complex disassembly which was subsequently validated by orthogonal methods, thus demonstrating the potential of the method and dataset to reveal new biology. To support additional exploration of the present data set and future differential SEC-SWATH datasets, we provide an online tool, SECexplorer-cc.

To retrieve information on changes in protein complex quantity and composition, we devised a protein-centric differential analysis strategy that detects altered SEC distribution patterns of ca. 5,000 proteins. Changes in these elution patterns between conditions indicated changes in the association of the respective protein with different complexes and such events could be quantitatively compared between states using the new quantification module implementing a strategy we term differential association testing. This computational pipeline was devised to achieve (i) optimal sensitivity to recall changes based on all available peptide-level information and (ii) maximal coverage by not restricting the differential analysis on previously known protein complexes. With more than 90% of the identified proteins being represented as quantitative SEC elution peaks, the method achieved broad coverage of biological functions. The protein-centric differential analysis pipeline is implemented as additional module of our R/CCprofiler toolset, available at https://github.com/CCprofiler/CCprofiler/tree/helaCC).

To benchmark the method we compared the differential SEC-SWATH-MS data obtained in this study with results of two studies that investigated cell-cycle-associated changes in protein complexes through inference from protein thermostability changes measured in similarly perturbed cell systems (TPP and CETSA, (Becher et al., 2018; Dai et al., 2018)). The TPP and SEC studies used HeLa cells (TPP; HeLa Kyoto; SEC, HeLa CCL2) and the CETSA study used K562 cells. Under the assumption of strong conservation of the mitotic regulatory circuitry between the cells tested, our comparison suggests favorable sensitivity to recall known cell cycle dependent complex rearrangement events of the SEC-SWATH-MS method. Despite the relatively small overlaps between the protein sets showing strongest mitotic changes in either method (15 - 16 %, see Figure 4C), the protein sets largely converged at the level of biological processes and pathways that were associated with the selected protein sets. Our results give first insights into the biological information gleaned from the two approaches, thermal stability and SEC-SWATH-MS, respectively, to infer mitotically altered protein states. However, generic conclusions on the relative merits of detecting alterations in apparent size vs. thermostability of protein complexes are not possible with certainty as in no case the two thermostability-based methods agree on an enriched term that was not also retrieved by SEC-SWATH-MS. More detailed analysis of the approach-specific hits suggests that certain instances of changes in enzymatic and metabolic activities are better-observable based on protein stability rather than size, which could arise from allosteric regulation by small molecule binding which is expected to be better detectable by altered thermostability than by altered complex size. Furthermore, the measurement of thermostability via TPP showed smaller bias against capturing of changes in membrane- and DNA-associated proteins than the results from the SEC-SWATH-MS and CETSA workflows (Figure 4D). Overall our comparison indicates higher sensitivity of the SEC-SWATH-MS method to uncover changes in core cell cycle machinery (Figure 4E), such as e.g. increased formation of translation initiation complexes or the inactivation of APC/C complex (**Supplemental Figure S3A**), when compared to the stability-based approaches. Overall the data suggest that SEC, as implemented in the SEC-SWATH-MS workflow, currently offers a preferable combination of analytical breadth and sensitivity.

Our study generated new and confirmed known biological insights. For example, only a subset of the CDK1 pool is recruited to upregulated CCNB1 in mitosis (31% appear to remain monomeric, compare Figure 3D). This effect cannot be attributed to incomplete synchronization of the analyzed cell populations. We surmise that the degree of assembly acts as an additional layer of CDK1 activity control by defining the cellular concentration of assembled CCNB1-CDK1 complexes that can be transferred to an active state by protein phosphorylation (Nigg, 1993). This notion is relevant for ongoing drug development programs that target the respective interaction interfaces as alternative anticancer strategy (Peyressatre et al., 2015).

Differential SEC-SWATH-MS data also illustrated the process of mitotic disassembly of NPCs into distinctive sub-complexes. This allowed us to identify a novel mitotic sub-complex and to extend the model for mitotic NPC disassembly (Laurell et al., 2011; Linder et al., 2017). These observations from the SEC-SWATH dataset were further confirmed with orthogonal methods. Even though several publications reported mechanistic insights into the phosphorylation-driven process of mitotic NPC disassembly (Laurell et al., 2011; Linder et al., 2017), the overall process remains incompletely understood. NPC disassembly must liberate Nups in a state in which they are readily available for rapid reassembly during mitotic exit. Our discovery of a mitotic protein complex between central channel FG-Nups and their anchoring scaffold Nups indicates that the efficient incorporation of the FG Nups may occur together with their anchoring partners. Such mechanism would be ideally suited to explain correct and rapid reassembly during mitotic exit. In this scenario, reintegration will rely on a larger set of protein interactions directed by the scaffold subunits. Detailed knowledge about the mechanistic rules of mitotic NPC disassembly and interphasic re-assembly may well prove helpful in the design of future therapies aiming to modulate the process of cell division.

The specific complexes discussed in this paper, including the novel NPC disassembly intermediates only represent a small fraction of the information contained in the dataset of 5,044 proteins of which 1,793 showed significantly different association with complexes across the two cell cycle states analyzed. To support further in depth interrogation of the dataset we disseminate it in an easily browsable form via *SECexplorer-cc*. To infer strong hypotheses on novel players that justify at times costly and lengthy follow-up experiments, it is important to bear in mind inherent limitations of and potential confounding effects in the experimental system. These include i) differential extractability of proteins due to cellular re-organization, exemplified by breakdown of the nuclear envelope upon mitotic entry, ii) indirect effects, i.e. rearrangement as a mere consequence rather than cause of the altered state of the biological system, iii) confounding effects of the experimental procedure, such as e.g. stress responses triggered by chemical treatment rather than cell cycle stage, as has been suggested in comparisons of quantitative proteome profiles employing different protocols to achieve cell cycle synchronization (Ly et al., 2015). Further, co-elution may be observed among proteins that participate in physically independent complexes that co-elute in chromatography, as exemplified by NUP50 co-elution with but not participation in the newly identified mitotic NPC sub-complex reported here. Further, experimentally induced disassembly of labile complexes is expectable under SEC-induced dilution and will likely result in an overestimation of the proteins present in monomeric form.

Multiple extensions or optimizations of the SEC-SWATH-MS methodology can be envisioned. For example, given the diverse properties of cellular assemblies methodological adaptations could benefit more focused studies of a given complex class of interest, such as e.g. ribosomal complexes (Yoshikawa et al., 2018). Further, limited sample throughput of the method currently precludes time course or cohort analyses (Compare 473 LC-MS measurements and 556 hours of gradient time spent here), a bottleneck which may be bypassed by integrating ultra-fast liquid chromatography setups (Bache et al., 2018) in the SWATH/DIA-MS workflow. Such adaptations will likely render the analysis of proteomes including their organizational state a routine procedure in the near future.

Another thrust for further development of the method is bioinformatic information retrieval. We here deliberately chose an approach that makes use of all peptide level information to detect rewiring proteins with maximal sensitivity and at optimal breadth. However, alternative computational strategies can be envisaged that will support the retrieval of additional information from SEC-SWATH-MS data. These include (i) increased sensitivity to detect proteins that exchange interactors without effect on the SEC-SWATH-MS signal (net-0 interactome changes) by interaction-network-centered approaches, (ii) improved method throughput by reducing the need for replication and rate of SEC sampling, (iii) improved information retrieval on the level of protein complexes and their precise composition across functional states and (iv) retrieval of information on post-translationally modified and alternatively spliced gene products and the respective impact on complex assembly and dynamics thereof. With developments along those lines ongoing, it appears particularly promising to re-interrogate the SEC-SWATH-MS data presented here from these additional perspectives in the future.

In contrast to classical proteomics studies that reduced proteomes to a quantitative list of component parts, SEC-SWATH-MS facilitates studies of the proteome including its biophysical arrangement into complexes as a core functional layer. This enables deep, systems-wide surveys of changes in protein association- and inferred activity states from virtually any experimental system of interest as the method is independent from genetic engineering. The datasets encode protein assembly and interaction states, the composition of complexes and alternate roles of distinct proteoforms. Systematic measurements of protein attributes closely correlated to protein function bear profound potential for the discovery of yet unknown players and mechanisms at the core of (disease) phenotypes generated by biomolecular networks and systems.

## Supporting information

Supplemental Item SI1 Protein chromatogram plots

## Acknowledgments

We would like to thank the Scientific IT Support (ID SIS) of ETH Zurich for support and maintenance of the Aebersold lab-internal computing infrastructure and the ScopeM facility for continuous support of microscopy at ETH Zurich. The project was supported by the SystemsX.ch projects PhosphoNetX PPM and project TbX to R.A., the Swiss National Science Foundation (grant no. 3100A0-688 107679 to R.A. and 310030_184801 to U.K.), and the European Research Council (ERC-20140AdG 670821 to R.A. and ERC AdG grant NucEnv_322582 to U.K). M.G. acknowledges the support by the Innovative Medicines Initiative project ULTRA-DD (FP07/2007-2013, grant no. 115766). MH was supported by a grant from Institut Mérieux. B.C.C. was supported by a Swiss National Science Foundation Ambizione grant (PZ00P3_161435).

## Author Contributions

Conceived and supervised project, UK, BCC, MG and RA; Performed experiments, MK, SAm, FF and MH, Generated and characterized the HeLa cell line expressing tetracycline-inducible HA-Strep Nup58, MIL, Performed data analysis, MF, MH, SAu; Developed and implemented data analysis tools MF, MH, IB, GR; Writing – original draft, MH; Writing – review & editing, MF, MK, SAm, GR, FF, IB, SAu, BCC, MG, UK, RA.

## Tables

There are no main text tables. All tables related to our work are included in the form of supplemental tables (See Supplemental Information).

## Experimental Procedures (initial submission)

### Cell culture and mitotic arrest

HeLa CCL2 cells were obtained from the ATCC collection and cultured in modified complete Dulbecco Modified Eagle Medium (DMEM) at 37°C and 5% CO2. Cells in interphase (early S-phase) were obtained by double-thymidine block. Cells in mitosis (early prometaphase) were obtained by single thymidine block followed by nocodazole treatment and mitotic

shake-off, as follows. Cells were grown to a confluency of approximately 60% and arrested in S phase by addition of 3 mM thymidine (Sigma) for 20 h. After overnight incubation the cells were thoroughly washed twice with warm PBS and left for recovery in complete DMEM medium for 2 h prior to addition of 3 mM thymidine (Sigma) and incubation at 37°C for 13 h (interphase-arrest by double thymidine-block), or, for synchronization in mitosis, cells were treated with 100 ng/ml nocodazole at 37°C for 13 h before harvest via mitotic shake-off to ensure recovery of only mitotic cells. Cells arrested in interphase were harvested using EDTA, before cells were pelleted and snap-frozen in liquid nitrogen. Synchronization in interphase and mitosis was confirmed by monitoring phosphorylation of NUP53 by Western blotting and the observed weight gain of mitotic, poly-phosphorylated NUP53 (See Figure 3C and **Figure S1A**).

### Tandem affinity purification of central channel complexes

HeLa cell lines carrying tetracycline-inducible HA-Strep-tagged NUP58 and HA-Strep-tagged GFP were generated using a HeLa Flp-In T-REx cell line (Häfner et al., 2014). Expression of HASt-NUP58 and HASt-GFP were induced by addition of 0.1 μg/mL tetracycline (Sigma). Correct expression and localization of affinity-tagged Nup58 was validated by immunofluorescence and Western blotting (see below). For affinity purifications, bait expression was induced for 48 h before synchronizing the cells in interphase or prometaphase as described above. After washing and pelleting, cells were snap-frozen in liquid nitrogen. Thawed cells were lysed by resuspension in lysis buffer containing 25 mM Tris-HCl pH 7.6, 125 mM NaCl, 2 mM MgCl2, 1 mM DTT, 0.5% NP-40, protease and phosphatase inhibitors and sonicated. After lysate clearance by centrifugation (15’000 rpm, 30 min, 4°C), HA-Strep-Nup58 and HA-Strep-GFP were purified by affinity chromatography with StrepTactin sepharose (IBA) for 45 min at 4°C. Beads were washed three times with lysis buffer and one time with lysis buffer without NP40 and protease inhibitors. Bound protein was eluted with elution buffer containing 25 mM Tris-HCl pH 7.6, 125 mM NaCl, 2 mM MgCl2, 1 mM DTT and 2.5 mM D-biotin (Sigma). Elutions and input samples were further analysed using SDS-PAGE followed by Western blotting.

### Western Blotting

Cell pellets from synchronized HeLa CCL2 cells and samples from pulldown experiments were resuspended in SDS-sample buffer and briefly denatured at 95°C. Protein was resolved by SDS-PAGE and transferred to nitrocellulose blotting membranes (GE Healthcare). Membranes were blocked over night with 5% skim milk powder in PBS-T (PBS containing 0.1% Tween 20). Subsequently, membranes were incubated at RT for 1 h with indicated antibodies diluted in 5% milk-PBS-T. Primary rabbit polyclonal antibodies directed against NUP188, NUP93, NUP53 and NUP54 have been described (Linder et al., 2017). Antibodies directed against actin (Sigma, cat no. A1978), HA (Roche), pH3 (Cell signaling, cat no. 9701S) and NUP62 (Abcam, cat no. ab188413) are commercially available. After three washing steps with TBS-T secondary antibody solutions were applied in 5% milk-PBS-T and membranes kept shaking for 1 h at RT. Subsequent washing was followed by detection. HRP-conjugated secondary antibodies used to detect primary antibodies included goat anti–rabbit IgG and goat anti-mouse IgG (Sigma-Aldrich). Chemiluminescence was initiated using ECL detection reagent (GE Healthcare) and the signal was detected using Fuji RX film (Fujifilm)

### Validation of cell cycle arrest and HASt-NUP58 HeLa cell line by immunofluorescence

For immunofluorescence, cells were fixed with 4% paraformaldehyde in PBS, washed with PBS and permeabilized for 5 min in 0.1% Triton X-100. Immunostaining was performed as described previously(Zemp et al., 2009). Briefly, cells were blocked with blocking solution (2% BSA in PBS) at RT for 45 min. Anti-HA antibody (Enzo) was 1:500 diluted and anti-p-Histone H3S10 (Cell Signalling Technology) 1:400 in blocking solution and fixed cells were exposed at RT for 1 hr. Subsequently, cells were washed with blocking solution and stained with fluorescently labelled secondary antibody at a dilution of 1:300. After three washing steps with blocking solution, DNA was stained using Hoechst 33342 (ThermoFisher) 1:5000 diluted in blocking solution. Eventually, coverslips were mounted in Vectashield mounting medium (Vector laboratories Inc.) on a glass slide. Localization and expression levels were analysed using a Zeiss LSM 880 upright microscope with a 63x 1.4NA, oil, DIC Plan-Apochromat objective, and Western blotting respectively.

### Preparation of native proteomes and fractionation for MS analysis

Ca. 5e7 HeLa CCL2 cells were mildly lysed by freeze-thawing into 0.5% NP-40 detergent- and protease and phosphatase inhibitor containing buffer, essentially as described(Collins et al., 2013), albeit without the addition of Avidin. Lysates were cleared by 15 minutes of ultracentrifugation (100,000×g, 4 °C) and buffer was exchanged to SEC buffer (50 mM HEPES pH 7.5, 150 mM NaCl) over 30 kDa molecular weight cut-off membrane at a ratio of 1:25 and concentrated to 25-35 mg/ml (estimated by OD280). After 5 min of centrifugation at 16,900 ×g at 4 °C, the supernatant was directly subjected to fractionation on a SRT-C-SEC 500 column (dimensions 300×21.2 mm, pore size 500 Å, particle size 5 µm, Sepax-Tech, DE, USA). Per SEC run, 7.25 mg of native proteome extract (estimated by OD280) was injected and fractionated at 2 ml/min flow rate on ice (0-4 °C), collecting 90 fractions at 0.4 min per fraction from 20 to 56 min post-injection, fractions 1-65 (20 – 46 min elution time, 40 – 92 ml elution volume) of which were considered relevant proteome elution range and considered for quantitative analysis. For library generation purposes, a few fractions from an extended elution range were analyzed (up to fraction 89). In order to minimize time-sensitive artifacts of complex disassembly under dilution, samples were processed independently to achieve a fixed processing time-to-column of 1.5-2 h. Apparent molecular weight per fraction was log-linearly calibrated based on the apex elution fractions of a 5-protein standard sample with known protein mass analyzed with each experimental replicate (Column Performance Check Standard, Aqueous SEC 1, AL0-3042, Phenomenex, CA, USA). An aliquot of the unfractionated mild proteome extract was included in peptide sample preparation and LC-MS analysis.

### MS sample preparation

Size exclusion chromatographic fractions and an aliquot of the unfractionated mild proteome extract were processed as follows. Proteins were denatured by the addition of 1% sodium deoxycholate (SDC, Sigma-Aldrich) and heating to 95 °C for 5 min. Disulfide bonds were reduced by adding tris-(2-carboxyethyl)-phosphine to 5 mM and incubating at 37 °C for 30 min. Subsequent alkylation of free thiol groups was carried out by adding iodoacetamide to 10 mM and incubating in the dark for 30 min. Proteins were digested by 1.4 µg sequencing grade trypsin (Promega) per fraction at 37 °C overnight. 1% TFA and 1% ACN were added for stopping the digestion and precipitating SDC. Additionally, 0.4 pmol of E. coli β-galactosidase digest were added as internal standard, followed by centrifugation at 4,500 ×g for 15 min. Subsequently, samples were desalted by means of C18 reversed phase well plates (96-Well MACROSpin Plate, The Nest Group, MA, USA) according to the manufacturer’s recommendations and vacuum dried. For MS analysis, de-salted peptide samples were re-suspended in 20 µl LC solvent A (2% ACN, 0.1% FA) supplemented with internal retention time calibration peptides (iRT kit, Ki-3002-1, Biognosys AG, CH, used at 1:20 instead of 1:10 ratio as indicated in the manufacturer’s protocol).

Tandem affinity purified samples were processed as follows. To remove biotin, proteins were precipitated from the AP eluate by 25% TCA at -20°C overnight and protein pellets were washed three times with ice-cold acetone and dried. Proteins were denatured by dissolution in 30 µl of 8M urea in 100 mM NH_4_HCO_3_ at RT, followed by dilution to 0.8 M urea, reduction by 5 mM TCEP (37°C, 30 min) and alkylation (10 mM iodoacetamide in 100 mM NH_4_HCO_3_, 37°C, 30 min in the dark) before overnight digestion by sequencing grade trypsin (1 µg, Promega, 37°C). After adjustment of sample pH to ∼2 (using 5% TFA), peptides were de-salted on C-18 spin columns (3-30 µg capacity, The Nest group) using 0.1% TFA as acidifier in 2%/50% ACN for washing/elution. For MS measurements, peptides were dissolved in 20 µl 2% ACN. 0.1% FA.

### LC-MS measurements

Mass spectrometric analysis of the peptide samples generated from the chromatographic fractions was carried out on an Eksigent nanoLC Ultra 1D Plus and expert 400 autosampler system (Eksigent, Dublin, CA) coupled to a TripleTOF 6600 (Sciex, Ontario, Canada) equipped with a NanoSpray III ion source. The acquisition software was Analyst TF 1.7.1. A 75 um inner diameter PicoFrit emitter (New Objective, Woburn, MA) was packed in-house with Magic C18 AQ 3 um, 200 Å particles (Bruker, Billerica, MA) to a length of 40 cm. It was operated at a flow of 300 nl/min and at room temperature. The LC solvent A was composed of 98% ultra-high quality water, 2% acetonitrile and 0.1% formic acid, LC solvent B was 98% acetonitrile, 2% ultra-high quality water and 0.1% formic acid. 3 µl of sample were loaded onto the column and separated by a linear gradient from 5 to 35% B in 60 min (or 120 min for DDA and library generation purposes, see below).

For the quantitative analysis of the 390 SEC fractions constituting the core sample set of the experiment, the TripleTOF was operated in SWATH mode. One MS1 survey scan was followed by 64 SWATH scans in a looped fashion. The SWATH windows are listed in Supplementary Table 1, covering precursors in the range of 400-1200 m/z with their widths chosen to obtain similar precursor intensity densities within all SWATH windows, i.e. resulting in narrower windows for m/z regions with a high density of precursors. Adjacent SWATH windows overlapped by 1 m/z to accommodate for the Q1 isolation profile. The monitored m/z range was 360-1460 in the MS1 scan and 300-2000 in the SWATH scans, the accumulation time was 200 ms and 50 ms, respectively, which resulted in a cycle time of around 3.4 s. For fragmentation, a rolling collisional energy (calculated for a theoretical 2+ ion centered in the corresponding SWATH window) with a collisional energy spread of 15 eV was applied. The ion source was operated with the following settings: spray voltage, 2600 V; ion source gas flow, 16; curtain gas flow, 35; interface heater temperature, 75°C and declustering potential, 100.

For the generation of a sample-specific library of peptide query parameters, 83 of the 390 SEC samples, representing all replicates, elution ranges and conditions and fractions covering an extended elution range up to F89 in experimental replicate 1, were selected for analysis by data-dependent acquisition mode over prolonged LCMS gradients of 2 h per sample in order to allow for deeper coverage of the peptide query parameter library. For DDA runs, samples were separated by a 120 min linear gradient from 5 to 35% LC solvent B. One MS1 scan was followed by 20 MS2 scans with an accumulation time of 250 ms for MS1 and 100 ms for MS2. The monitored m/z range was 360-1460 for MS1 and 50-2000 for MS2. The dynamic exclusion time was set to 20 s.

AP-MS samples were analyzed by LC-MS/MS on an LTQ-Orbitrap XL system (Thermo Fisher Scientific) equipped with an EASY nLC II system (Proxeon) operated in DDA mode, acquiring over an LC gradient of 5-35% of LC solvent B (98% ACN, 0.1% FA) in LC solvent A (2% ACN, 0.1% FA) in 90 min up to 10 MS/MS spectra of the up to top10 most-abundant precursors selected from intermittent MS1 survey scans (m/z range, 350-1600 m/z). Precursors analyzed were dynamically excluded from re-selection for 30 s and with an exclusion list size of maximally 300 entries. The automatic gain control (AGC) target value was set to 2e5 for full scans (MS) and 3e4 (MS/MS) scans.

### Data processing

#### DDA-MS data processing

For generation of the peptide query parameter library DDA-MS data acquired in 2 h gradient time from 83 SEC fractions representative of all replicates, condition and SEC elution ranges was processed by spectrum-centric analysis and then processed into a spectrum- and reduced peptide query parameter-library essentially as described(Schubert et al., 2015), except for the following adaptations: MS spectra were searched for matches to the human UniProt/SwissProt reference database (reviewed, canonical entries, build: 2017-06-19, supplemented with typical contaminant (cRAP, https://www.thegpm.org/crap/) and E. coli β-galactosidase sequences) using trypsin cleavage, 50 ppm precursor and 0.1 Da fragment ion mass tolerance, carbamidomethyl (C) as static and oxidation (M) as variable modification and allowing up to two missed cleavages. The results from four independent searches using different search engines (X!Tandem 2013.06.15.1, Ommsa 2.1.9, MyriMatch 2.1.138 and Comet 2015.02 rev. 3) were integrated using iProphet of the Trans-Proteomic Pipeline ((TPP v4.7 POLAR VORTEX rev 0, Build 201403121010), filtering the results at 1% peptide FDR (0.978883 iprob) as determined using the tool Mayu(Reiter et al., 2009). We intentionally chose a less strict peptide-level FDR cut-off (compared to requiring 1% FDR on protein level) in order to increase sensitivity for the recovery of true positive peptide signals which would be lost as false negatives under strict protein level FDR control but that can, in an SEC experiment, be validated by quantitative agreement with high confidence sibling peptides along SEC fractionation as assessed in downstream data filtering. The spectra were then processed into a spectral library using the tool Spectrast(Lam et al., 2010) with iRT calibration followed by the generation of a peptide query parameter library, essentially as described(Schubert et al., 2015) and within the iPortal compute infrastructure(Kunszt et al., 2015). Query parameters were composed selecting the 6 most abundant fragment ion transitions per precursor from the b or y ion series within m/z range 350-2000 and allowing fragment charge states 1-2 and no mass gains or losses. The final library contains query parameters for 111,267 precursors of 90,932 peptides mapping to 9603 protein groups that were subsequently targeted for quantification in the 390 60 min gradient SWATH-MS runs of the 390 SEC fractions. Given the strict rules employed for downstream quantification (quantifying only single, unique proteins with at least 2 unique, proteotypic peptides) the number of maximally detectable analytes as constrained by the query parameter library drops to 102,629 precursors of 83,863 peptides mapping to 5,916 unique proteins. 60 min DDA-MS data were processed equivalently by spectrum-centric analysis to obtain spectral counts across chromatographic fractions as quantitative measure for technical comparisons (DDA search results and the spectral and peptide query parameter libraries are available via ProteomeXchange, see section data availability below).

#### SWATH-MS data processing

The SWATH -MS data were analyzed via targeted, peptide-centric analysis, querying 111,267 precursors from the sample-specific peptide query parameter library (see above) in the SWATH fragment ion chromatograms, using a modified OpenSWATH(Röst et al., 2014), PyProphet (Reiter et al., 2011; Teleman et al., 2015) and TRIC(Röst et al., 2016) workflow. First, one global classifier was trained on a subsampled set of SEC fractions across the experiment using pyProphet-cli (Rosenberger et al., 2017) (Specifically, fractions 3, 43 and 44 of each replicate and condition were analyzed jointly in order to generate a stable scoring function from the most analyte-rich measurements (F43 and F44) while including different analytes detected exclusively in the high MW range (F3)). Peptides from all fractions were then quantified and scored using the pre-trained scoring function using OpenSWATH, pyProphet and TRIC in the iPortal framework(Kunszt et al., 2015). TRIC was set to recover precursors at an experiment-wide assay/peptide query-level (TRIC target) FDR of 5%. The full result table (E1709051521_feature_alignment.tsv.gz) has been deposited, together with the MS raw data, to ProteomeXchange, see section data availability below.

#### Downstream SEC-SWATH-MS data processing and FDR estimation

Precursor-level results from E1709051521_feature_alignment.tsv were imported into *CCprofiler* extended by the protein-centric differential analysis module. Upon import, precursor intensity signals (summed intensity of the 6 most-abundant fragment ion XIC peak area) were summed per peptide (function: *importFromOpenSWATH*). Then, missing values flanked by at least two consecutive identifications were imputed by a spline fit (function: *findMissingValues*).Traces were scaled to the internal standard spike-in peptides of E. coli β-galactosidase (P00722, function: *normalizeToStandard*) followed by smoothing of total intensities according to a spline fit over total MS intensity sums (function: *smootheTraces*). The data were further filtered on chromatography-informed scores following final FDR estimation based on the simple target-decoy method. Specific filtering rules were, first, eliminating all values part of consecutive identification stretches below length three and, second, exclusion of peptides based on their quantitative fractionation pattern’s average dissimilarity to those of sibling peptides (originating from the same parent protein). Guided by the fraction of decoys remaining (target: 3% on unique protein identifier level), peptides with average sibling peptide correlation coefficient (spc) below [0.28 - 0.34] were removed, with cut-offs selected to achieve a decoy rate below 3% among the remaining protein entries. Subsequently, proteins were quantified by summing the top2 most-intense peptide signals.

Protein intensity across the replicates was averaged and standard errors of the mean calculated. Cumulatively, 5044 proteins were characterized. The FDR on protein level was estimated based on the simple target-decoy method, with correction for the fraction of false protein targets (equivalent to the fraction of absent proteins, π_A_(The et al., 2016)). We estimate π_A_ as the fraction of library-contained target proteins that were not detected with high confidence in the targeted analysis, i.e. π_A_ ≈ (5,916 – 5,044) / 5,916 = 0.147. This estimation is conservative because not all true positive proteins are recovered at high confidence, leading to overestimation of π_A_ and thus more conservative corrected FDR estimates. Accordingly, 135 decoy entries passing the filters together with the 5044 target proteins (decoy rate, 0.026) point to an estimated global protein level FDR of (135 * 0.147) / 5,044 = 0.0039, ≤ 0.4%.

#### Protein-centric differential SEC distribution testing

To detect proteins that significantly alter their quantitative SEC elution pattern we devised a protein-centric differential analysis pipeline that leverages all information on peptide level to identify proteins that shift elution behavior across distinct SEC ranges of elution (elution features). First, the global set of protein elution features apparent from the dataset were detected from an integrated set of peptide chromatograms (traces) using the SECprofiler framework, followed by statistical testing for differences based on the intensities observed in the different experimental replicates. Specifically, peptide traces were averaged within the conditions and then artificially combined into one summed set of 60891 traces (function *integrateTraceIntensities*). Then, high confidence protein elution features were detected as signals of co-peaking peptide traces, grouping the traces by parent protein identifier and including decoys with randomized peptide-to-protein mapping (functions: *findProteinFeatures*, *calculateCoelutionScore* and *calculateQvalue*). At a q-value cut-off of 5%, 6044 protein elution features were detected to define the ranges for differential abundance tests.

To detect differential SEC elution behavior, peptide abundances in the elution ranges were then tested for significant differences in abundance, using the functions *extractFeatureVals* and *testDifferentialExpression* and employing the paired t-test as statistical metric. All tests were based on the raw variability of the data, i.e. minimally processed data points (essentially, only scaled within conditions), to avoid biases introduced by data processing. Missing values were replaced by uniformly sampled intensities in the 5th percentile of quantified values. The test results were collected on protein level by deriving a fold-change adjusted median p-value from all peptide tests mapping to the respective parent proteins (Suomi and Elo, 2017). Using the function *aggregatePeptideTests* the protein level significance was calculated using a cumulative beta distribution parametrized on the number observed of peptides (Suomi and Elo, 2017). This resulted in protein p-values for 1-5 individual features per protein with separate scoring. To simplify visualization and reduce the results to one set of scores per protein, for each protein the lowest pBHadj and largest fold-change of which was selected for visualization in the screen hit volcano plot (See Figure 1 **D**). The R package *CCprofiler*, extended by the protein-centric differential analysis module is available via GitHub (https://github.com/CCprofiler/CCprofiler/tree/helaCC).

#### AP-MS data analysis

AP-MS DDA-MS data were analyzed by spectrum-centric analysis matching spectra against the human reviewed SwissProt reference sequence database (build 2018-08-01) using three search engines (X!Tandem, Myrimatch and Comet) and result integration via the Trans-proteomic pipeline (TPP), applying FDR cut-offs of 1% at the peptide level (iprophet-pepFDR, ≥ 0.6581 iprobability) and protein level (ProteinProphet), executed within the iportal framework (Kunszt et al., 2015). Searches allowed mass tolerances of 15 ppm and 0.4 Da on precursor and fragment ion level. Enzyme specificity was set to Trypsin, allowing up to two missed cleavages. Carbamidomethyl was set as static modification, no variable modifications were considered.

#### Assignment of Nuclear pore complex components

Nuclear pore complex members were assigned based on two consecutively applied criteria, i) Uniprot query (nuclear pore complex AND reviewed:yes AND organism:“Homo sapiens (Human) [9606]”) and ii) structural assignment in Table 1 by Hoelz et al. (Hoelz et al., 2016), resulting in 32 canonical subunits employed for benchmarking and exploration of NPC disassembly sub-complexes.

#### Statistical protein set enrichment and testing of over represented protein sets

Statistical over representation of protein functional groups among the protein sets of interest was performed in the PANTHER classification system (http://pantherdb.org/, Release 14.1) using the Statistical over representation test (Fisher’s exact test), testing against the background of all detected proteins (n = 6,010, Figure 3, Figure 4 and **Supplemental Figure S3A**) or the full human genome background (**Supplemental Figure S3C**).

#### Data availability

The mass spectrometry proteomics raw data and processing results have been deposited to the ProteomeXchange Consortium (http://proteomecentral.proteomexchange.org) via the PRIDE partner repository(Vizcaíno et al., 2012) with the dataset identifier PXD010288. The data are available in easily browsable and searchable form via https://sec-explorer.shinyapps.io/hela_cellcycle/ where the full dataset can be accessed using the password “ethsecexplorer”.

## Supplemental Information

The supplemental information entails six Supplemental Figures (S1-S6), one Supplemental Item (SI1) and three Supplemental Tables (ST1-3) with self-contained legends where applicable and the following titles.

Supplemental Figure S1: Validation of homogenous cell cycle arrest and of inducible Nup58-expressing HeLa cell lines

Supplemental Figure S2: Benchmarking the SEC-SWATH-MS workflow - correction of longitudinal effects and replicate intensity correlation

Supplemental Figure S3: Pathway enrichment among top 1000 proteins recovered by size or thermostability profiling along the cell cycle

Supplemental Figure S4: Protein elution peak detection and properties of multi-complex signal proteins

Supplemental Figure S5: Investigation of RAP1-TRF2-IKK cross-complex interaction and zoom-in to NUP93 subpopulations resolved across mitotic states

Supplemental Figure S6: Investigation of SEC13 moonlighting across Nup107-160 subcomplex and COPII vesicle transport related complexes.

Supplemental Item SI1: SEC-SWATH-MS protein chromatogram plots Supplemental Table ST1: Global protein quantification summary table

Supplemental Table ST2: Global protein elution peak detection and statistical scoring summary table

Supplemental Table ST3: Global complex remodeling summary table

## References

Aebersold, R., and Mann, M. (2016). Mass-spectrometric exploration of proteome structure and function. Nature 537, 347–355.

Arat, N.Ö., and Griffith, J.D. (2012). Human Rap1 Interacts Directly with Telomeric DNA and Regulates TRF2 Localization at the Telomere. J. Biol. Chem. 287, 41583–41594.

Bache, N., Geyer, P.E., Bekker-Jensen, D.B., Hoerning, O., Falkenby, L., Treit, P. V, Doll, S., Paron, I., Müller, J.B., Meier, F., et al. (2018). A Novel LC System Embeds Analytes in Pre-formed Gradients for Rapid, Ultra-robust Proteomics. Mol. Cell. Proteomics 17, 2284–2296.

Becher, I., Andrés-Pons, A., Romanov, N., Stein, F., Schramm, M., Baudin, F., Helm, D., Kurzawa, N., Mateus, A., Mackmull, M.T., et al. (2018). Pervasive Protein Thermal Stability Variation during the Cell Cycle. Cell 173, 1495–1507.e18.

Beck, M., and Hurt, E. (2017). The nuclear pore complex: understanding its function through structural insight. Nat. Rev. Mol. Cell Biol. 18, 73–89.

Chug, H., Trakhanov, S., Hülsmann, B.B., Pleiner, T., and Görlich, D. (2015). Crystal structure of the metazoan Nup62•Nup58•Nup54 nucleoporin complex. Science (80-.). 350, 106–110.

Collins, B.C., Gillet, L.C., Rosenberger, G., Röst, H.L., Vichalkovski, A., Gstaiger, M., and Aebersold, R. (2013). Quantifying protein interaction dynamics by SWATH mass spectrometry: application to the 14-3-3 system. Nat. Methods 10, 1246–1253.

Collins, B.C., Hunter, C.L., Liu, Y., Schilling, B., Rosenberger, G., Bader, S.L., Chan, D.W., Gibson, B.W., Gingras, A.-C., Held, J.M., et al. (2017). Multi-laboratory assessment of reproducibility, qualitative and quantitative performance of SWATH-mass spectrometry. Nat. Commun. 8, 291.

Dai, L., Zhao, T., Bisteau, X., Sun, W., Prabhu, N., Lim, Y.T., Sobota, R.M., Kaldis, P., and Nordlund, P. (2018). Modulation of Protein-Interaction States through the Cell Cycle. Cell 173, 1481–1494.e13.

Dominguez, D., Tsai, Y.-H., Weatheritt, R., Wang, Y., Blencowe, B.J., Wang, Z., Matese, J., Perou, C., Hurt, M., Brown, P., et al. (2016). An extensive program of periodic alternative splicing linked to cell cycle progression. Elife 5, 191–207.

Dong, M., Yang, L.L., Williams, K., Fisher, S.J., Hall, S.C., Biggin, M.D., Jin, J., and Witkowska, H.E. (2008). A “tagless” strategy for identification of stable protein complexes genome-wide by multidimensional orthogonal chromatographic separation and iTRAQ reagent tracking. J. Proteome Res. 7, 1836–1849.

Dunkley, T.P.J., Watson, R., Griffin, J.L., Dupree, P., and Lilley, K.S. (2004). Localization of organelle proteins by isotope tagging (LOPIT). Mol. Cell. Proteomics 3, 1128–1134.

Foster, L.J., de Hoog, C.L., Zhang, Y., Zhang, Y., Xie, X., Mootha, V.K., and Mann, M. (2006). A mammalian organelle map by protein correlation profiling. Cell 125, 187–199.

Gavet, O., and Pines, J. (2010). Activation of cyclin B1–Cdk1 synchronizes events in the nucleus and the cytoplasm at mitosis. J. Cell Biol. 189, 247–259.

Gillet, L.C., Navarro, P., Tate, S., Röst, H., Selevsek, N., Reiter, L., Bonner, R., and Aebersold, R. (2012). Targeted data extraction of the MS/MS spectra generated by data-independent acquisition: a new concept for consistent and accurate proteome analysis. Mol. Cell. Proteomics 11, O111.016717.

Gillet, L.C., Leitner, A., and Aebersold, R. (2016). Mass Spectrometry Applied to Bottom-Up Proteomics: Entering the High-Throughput Era for Hypothesis Testing. Annu. Rev. Anal. Chem. 9, 449–472.

Gong, D., Pomerening, J.R., Myers, J.W., Gustavsson, C., Jones, J.T., Hahn, A.T., Meyer, T., and Ferrell Jr., J.E. (2007). Cyclin A2 Regulates Nuclear-Envelope Breakdown and the Nuclear Accumulation of Cyclin B1. Curr. Biol. 17, 85–91.

Häfner, J., Mayr, M.I., Möckel, M.M., and Mayer, T.U. (2014). Pre-anaphase chromosome oscillations are regulated by the antagonistic activities of Cdk1 and PP1 on Kif18A. Nat. Commun. 5, 4397.

Hartwell, L.H., Hopfield, J.J., Leibler, S., and Murray, A.W. (1999). From molecular to modular cell biology. Nature 402, C47–52.

Heusel, M., Bludau, I., Rosenberger, G., Hafen, R., Frank, M., Banaei-Esfahani, A., Drogen, A., A. van, Collins, B.C., Gstaiger, M., and Aebersold, R. (2019). Complex-centric proteome profiling by SEC-SWATH-MS. Mol. Syst. Biol. 15, e8438.

Hoelz, A., Glavy, J.S., and Beck, M. (2016). Toward the atomic structure of the nuclear pore complex: when top down meets bottom up. Nat. Struct. Mol. Biol. 23, 624–630.

Ideker, T., Galitski, T., and Hood, L. (2001). A new approach to decoding life: systems biology. Annu. Rev. Genomics Hum. Genet. 2, 343–372.

Itzhak, D.N., Tyanova, S., Cox, J., and Borner, G.H. (2016). Global, quantitative and dynamic mapping of protein subcellular localization. Elife 5.

Kristensen, A.R., Gsponer, J., and Foster, L.J. (2012). A high-throughput approach for measuring temporal changes in the interactome. Nat. Methods 9, 907–909.

Kunszt, P., Blum, L., Hullár, B., Schmid, E., Srebniak, A., Wolski, W., Rinn, B., Elmer, F.-J., Ramakrishnan, C., Quandt, A., et al. (2015). iPortal: the swiss grid proteomics portal: Requirements and new features based on experience and usability considerations. Concurr. Comput. Pract. Exp. 27, 433–445.

Lam, H., Deutsch, E.W., and Aebersold, R. (2010). Artificial decoy spectral libraries for false discovery rate estimation in spectral library searching in proteomics. J. Proteome Res. 9, 605– 610.

Larance, M., Kirkwood, K.J., Tinti, M., Brenes Murillo, A., Ferguson, M.A.J., and Lamond, (2016). Global Membrane Protein Interactome Analysis using *In vivo* Crosslinking and Mass Spectrometry-based Protein Correlation Profiling. Mol. Cell. Proteomics 15, 2476– 2490.

Laurell, E., Beck, K., Krupina, K., Theerthagiri, G., Bodenmiller, B., Horvath, P., Aebersold, R., Antonin, W., and Kutay, U. (2011). Phosphorylation of Nup98 by Multiple Kinases Is Crucial for NPC Disassembly during Mitotic Entry. Cell 144, 539–550.

Leuenberger, P., Ganscha, S., Kahraman, A., Cappelletti, V., Boersema, P.J., von Mering, C., Claassen, M., and Picotti, P. (2017). Cell-wide analysis of protein thermal unfolding reveals determinants of thermostability. Science (80-.). 355, eaai7825.

Lin, D.H., Stuwe, T., Schilbach, S., Rundlet, E.J., Perriches, T., Mobbs, G., Fan, Y., Thierbach, K., Huber, F.M., Collins, L.N., et al. (2016). Architecture of the symmetric core of the nuclear pore. Science (80-.). 352, aaf1015–aaf1015.

Linder, M.I., Köhler, M., Boersema, P., Weberruss, M., Wandke, C., Marino, J., Ashiono, C., Picotti, P., Antonin, W., and Kutay, U. (2017). Mitotic Disassembly of Nuclear Pore Complexes Involves CDK1- and PLK1-Mediated Phosphorylation of Key Interconnecting Nucleoporins. Dev. Cell 43, 141–156.e7.

Liu, F., and Fitzgerald, M.C. (2016). Large-Scale Analysis of Breast Cancer-Related Conformational Changes in Proteins Using Limited Proteolysis. J. Proteome Res. 15, 4666– 4674.

Liu, D., Safari, A., O’Connor, M.S., Chan, D.W., Laegeler, A., Qin, J., and Songyang, Z. (2004). PTOP interacts with POT1 and regulates its localization to telomeres. Nat. Cell Biol. 6, 673–680.

Liu, X., Yang, W., Gao, Q., and Regnier, F. (2008). Toward chromatographic analysis of interacting protein networks. J. Chromatogr. A 1178, 24–32.

Ludwig, C., and Aebersold, R. Getting Absolute : Determining Absolute Protein Quantities via Selected Reaction Monitoring Mass Spectrometry. 80–109.

Ly, T., Endo, A., and Lamond, A.I. (2015). Proteomic analysis of the response to cell cycle arrests in human myeloid leukemia cells. Elife 4.

Nigg, E.A. (1993). Cellular substrates of p34(cdc2) and its companion cyclin-dependent kinases. Trends Cell Biol. 3, 296–301.

Peyressatre, M., Prével, C., Pellerano, M., and Morris, M.C. (2015). Targeting cyclin-dependent kinases in human cancers: from small molecules to Peptide inhibitors. Cancers (Basel). 7, 179–237.

Reiter, L., Claassen, M., Schrimpf, S.P., Jovanovic, M., Schmidt, A., Buhmann, J.M., Hengartner, M.O., and Aebersold, R. (2009). Protein identification false discovery rates for very large proteomics data sets generated by tandem mass spectrometry. Mol. Cell. Proteomics 8, 2405–2417.

Reiter, L., Rinner, O., Picotti, P., Hüttenhain, R., Beck, M., Brusniak, M.-Y., Hengartner, M.O., and Aebersold, R. (2011). mProphet: automated data processing and statistical validation for large-scale SRM experiments. Nat. Methods 8, 430–435.

Rosenberger, G., Ludwig, C., Röst, H.L., Aebersold, R., and Malmström, L. (2014). aLFQ: an R-package for estimating absolute protein quantities from label-free LC-MS/MS proteomics data. Bioinformatics 30, 2511–2513.

Rosenberger, G., Bludau, I., Schmitt, U., Heusel, M., Hunter, C.L., Liu, Y., MacCoss, M.J., MacLean, B.X., Nesvizhskii, A.I., Pedrioli, P.G.A., et al. (2017). Statistical control of peptide and protein error rates in large-scale targeted data-independent acquisition analyses. Nat. Methods 14, 921–927.

Röst, H.L., Rosenberger, G., Navarro, P., Gillet, L., Miladinović, S.M., Schubert, O.T., Wolski, W., Collins, B.C., Malmström, J., Malmström, L., et al. (2014). OpenSWATH enables automated, targeted analysis of data-independent acquisition MS data. Nat. Biotechnol. 32, 219–223.

Röst, H.L., Liu, Y., D’Agostino, G., Zanella, M., Navarro, P., Rosenberger, G., Collins, B.C., Gillet, L., Testa, G., Malmström, L., et al. (2016). TRIC: an automated alignment strategy for reproducible protein quantification in targeted proteomics. Nat. Methods 13, 777–783.

Ruepp, A., Waegele, B., Lechner, M., Brauner, B., Dunger-Kaltenbach, I., Fobo, G., Frishman, G., Montrone, C., and Mewes, H.-W. (2010). CORUM: the comprehensive resource of mammalian protein complexes--2009. Nucleic Acids Res. 38, D497–501.

Schopper, S., Kahraman, A., Leuenberger, P., Feng, Y., Piazza, I., Müller, O., Boersema, P.J., and Picotti, P. (2017). Measuring protein structural changes on a proteome-wide scale using limited proteolysis-coupled mass spectrometry. Nat. Protoc. 12, 2391–2410.

Schubert, O.T., Gillet, L.C., Collins, B.C., Navarro, P., Rosenberger, G., Wolski, W.E., Lam, H., Amodei, D., Mallick, P., MacLean, B., et al. (2015). Building high-quality assay libraries for targeted analysis of SWATH MS data. Nat. Protoc. 10, 426–441.

Scott, N.E., Rogers, L.D., Prudova, A., Brown, N.F., Fortelny, N., Overall, C.M., and Foster, (2017). Interactome disassembly during apoptosis occurs independent of caspase cleavage. Mol. Syst. Biol. 13, 906.

Stacey, R.G., Skinnider, M.A., Scott, N.E., and Foster, L.J. (2017). A rapid and accurate approach for prediction of interactomes from co-elution data (PrInCE). BioRxiv 152355.

Suomi, T., and Elo, L.L. (2017). Enhanced differential expression statistics for data-independent acquisition proteomics. Sci. Rep. 7, 5869.

Szklarczyk, D., Morris, J.H., Cook, H., Kuhn, M., Wyder, S., Simonovic, M., Santos, A., Doncheva, N.T., Roth, A., Bork, P., et al. (2017). The STRING database in 2017: quality-controlled protein–protein association networks, made broadly accessible. Nucleic Acids Res. 45, D362–D368.

Tan, C.S.H., Go, K.D., Bisteau, X., Dai, L., Yong, C.H., Prabhu, N., Ozturk, M.B., Lim, Y.T., Sreekumar, L., Lengqvist, J., et al. (2018). Thermal proximity coaggregation for system-wide profiling of protein complex dynamics in cells. Science 359, 1170–1177.

Tang, B.L., Peter, F., Krijnse-Locker, J., Low, S.H., Griffiths, G., and Hong, W. (1997). The mammalian homolog of yeast Sec13p is enriched in the intermediate compartment and is essential for protein transport from the endoplasmic reticulum to the Golgi apparatus. Mol. Cell. Biol. 17, 256–266.

Teleman, J., Röst, H.L., Rosenberger, G., Schmitt, U., Malmström, L., Malmström, J., and Levander, F. (2015). DIANA--algorithmic improvements for analysis of data-independent acquisition MS data. Bioinformatics 31, 555–562.

Teo, H., Ghosh, S., Luesch, H., Ghosh, A., Wong, E.T., Malik, N., Orth, A., de Jesus, P., Perry, A.S., Oliver, J.D., et al. (2010). Telomere-independent Rap1 is an IKK adaptor and regulates NF-κB-dependent gene expression. Nat. Cell Biol. 12, 758–767.

The, M., Tasnim, A., and Käll, L. (2016). How to talk about protein-level false discovery rates in shotgun proteomics. Proteomics 16, 2461–2469.

Vermeulen, K., Van Bockstaele, D.R., and Berneman, Z.N. (2003). The cell cycle: a review of regulation, deregulation and therapeutic targets in cancer. Cell Prolif. 36, 131–149.

Vizcaíno, J.A., Côté, R.G., Csordas, A., Dianes, J.A., Fabregat, A., Foster, J.M., Griss, J., Alpi, E., Birim, M., Contell, J., et al. (2012). The Proteomics Identifications (PRIDE) database and associated tools: status in 2013. Nucleic Acids Res. 41, D1063–D1069.

Wan, C., Borgeson, B., Phanse, S., Tu, F., Drew, K., Clark, G., Xiong, X., Kagan, O., Kwan, J., Bezginov, A., et al. (2015). Panorama of ancient metazoan macromolecular complexes. Nature.

Wessels, H.J.C.T., Vogel, R.O., van den Heuvel, L., Smeitink, J.A., Rodenburg, R.J., Nijtmans, L.G., and Farhoud, M.H. (2009). LC-MS/MS as an alternative for SDS-PAGE in blue native analysis of protein complexes. Proteomics 9, 4221–4228.

Xu, Y., Strickland, E.C., and Fitzgerald, M.C. (2014). Thermodynamic Analysis of Protein Folding and Stability Using a Tryptophan Modification Protocol. Anal. Chem. 86, 7041–7048.

Yoshikawa, H., Larance, M., Harney, D.J., Sundaramoorthy, R., Ly, T., Owen-Hughes, T., and Lamond, A.I. (2018). Efficient analysis of mammalian polysomes in cells and tissues using Ribo Mega-SEC. Elife 7.

Zemp, I., Wild, T., O’Donohue, M.-F., Wandrey, F., Widmann, B., Gleizes, P.-E., and Kutay, U. (2009). Distinct cytoplasmic maturation steps of 40S ribosomal subunit precursors require hRio2. J. Cell Biol. 185, 1167–1180.

