## Supplementary figures and images for "A global screen for assembly state changes of the mitotic proteome by SEC-SWATH-MS"

### SECchrom_O00764_PDXK_HUMAN_PDXK_C21orf124_C21orf97_PKH_PNK_.pdf

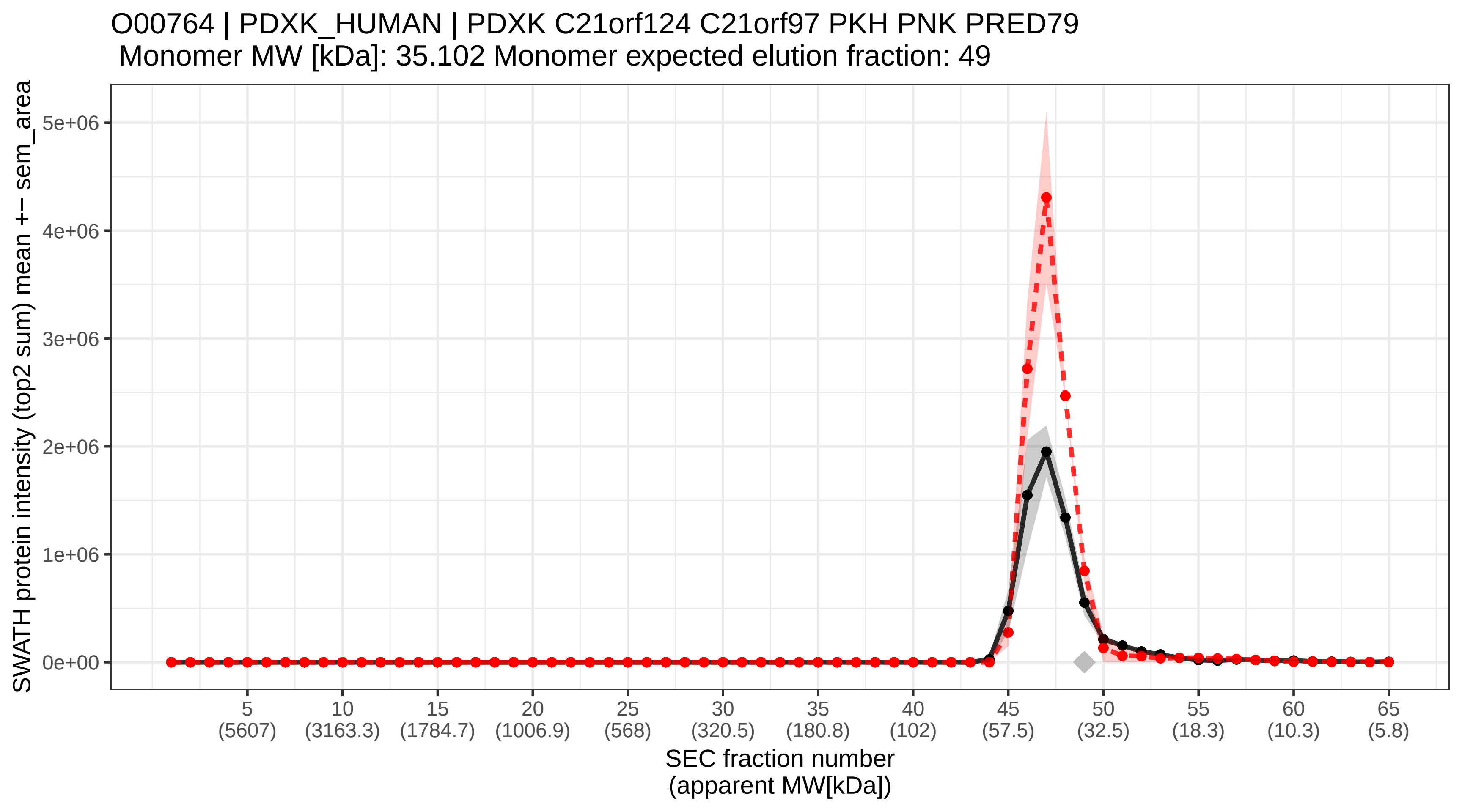

### SECchrom_O00767_ACOD_HUMAN_SCD.pdf

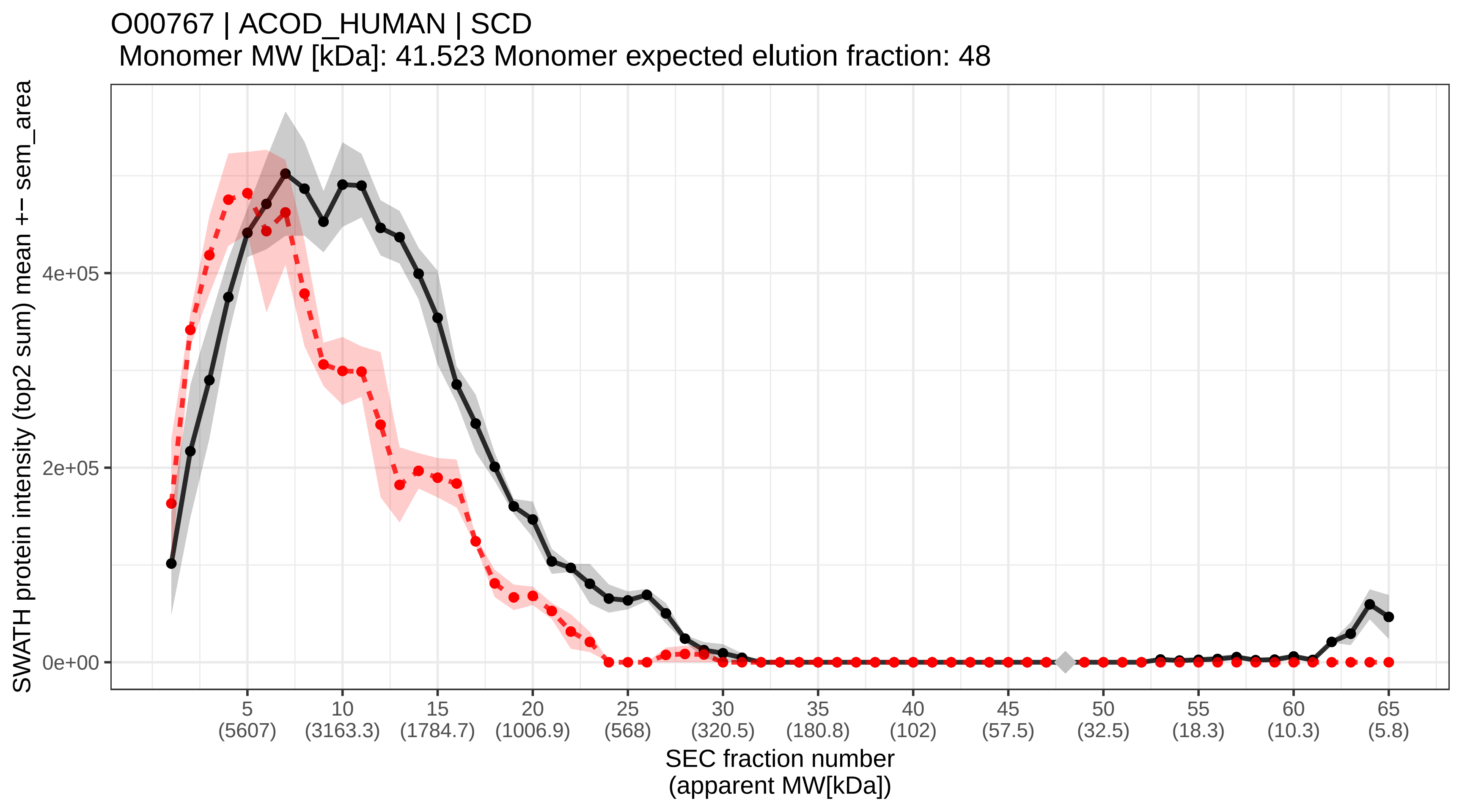

### SECchrom_O14497_ARI1A_HUMAN_ARID1A_BAF250_BAF250A_C1orf4_OS.pdf

O14497 | ARI1A_HUMAN | ARID1A BAF250 BAF250A C1orf4 OSA1 SMARCF1
Monomer MW [kDa]: 242.045 Monomer expected elution fraction: 32

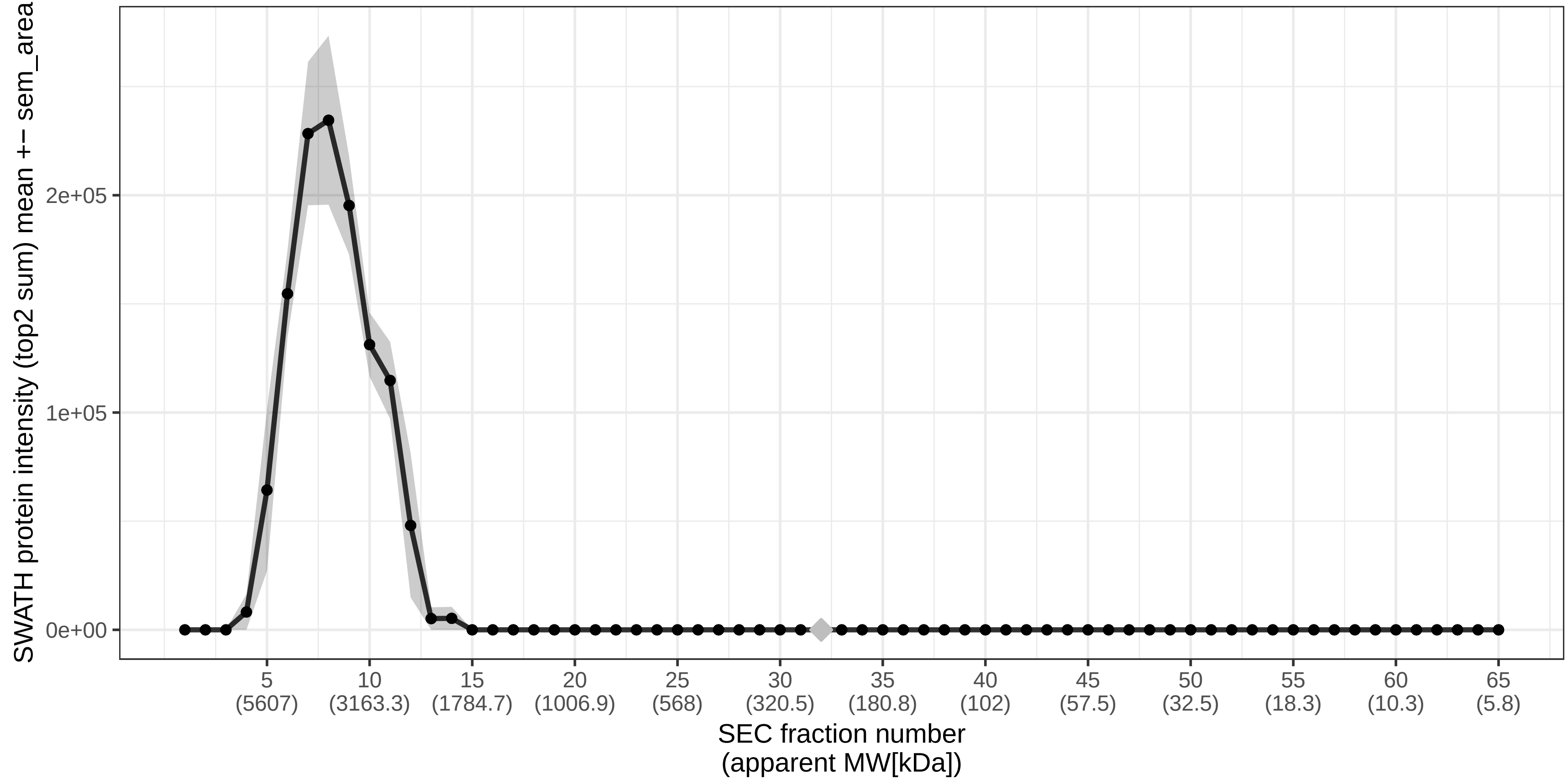

### SECchrom_O14524_NEMP1_HUMAN_NEMP1_KIAA0286_TMEM194_TMEM194A.pdf

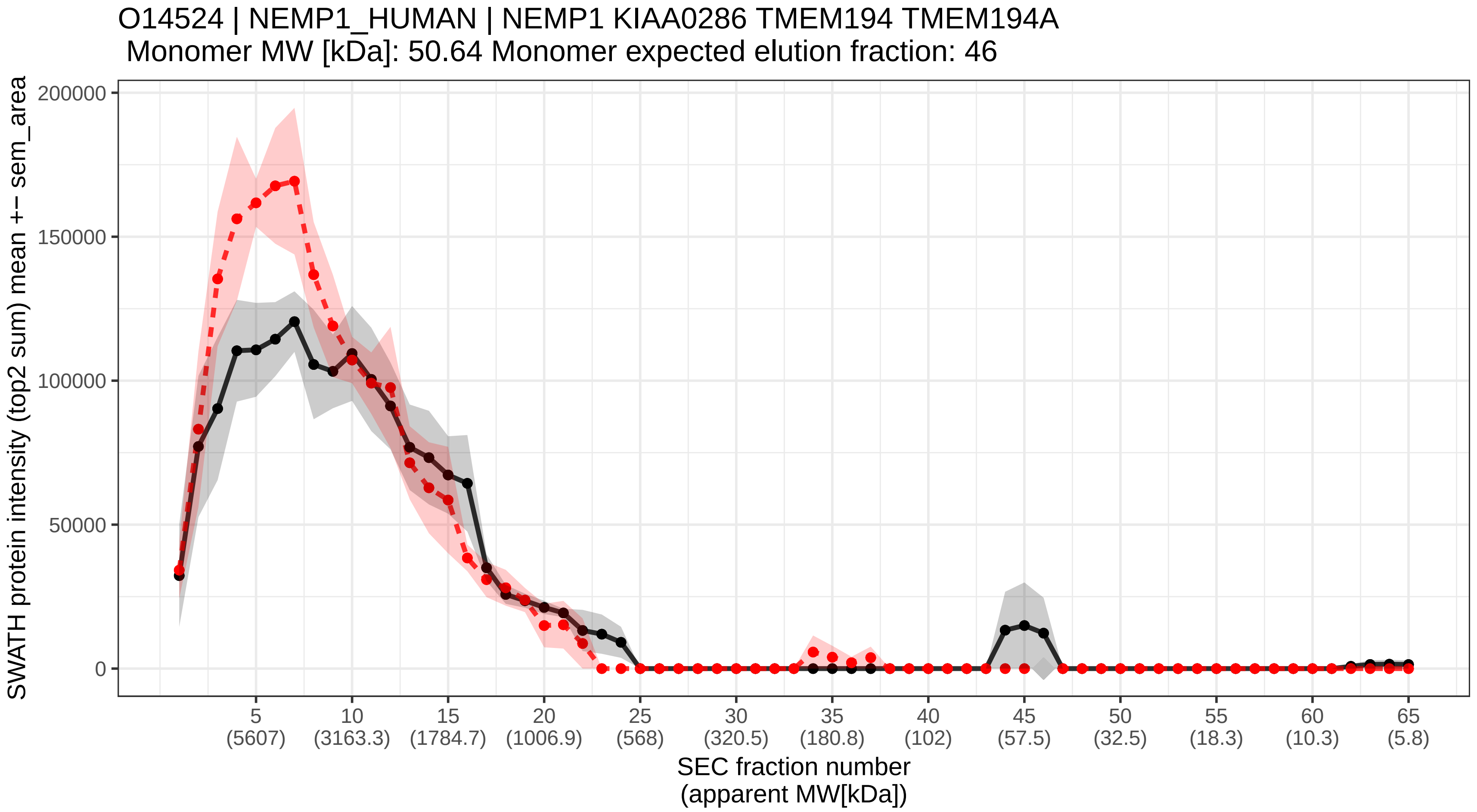

### SECchrom_O14530_TXND9_HUMAN_TXNDC9_APACD.pdf

O14530 | TXND9_HUMAN | TXNDC9 APACD
Monomer MW [kDa]: 26.534 Monomer expected elution fraction: 52

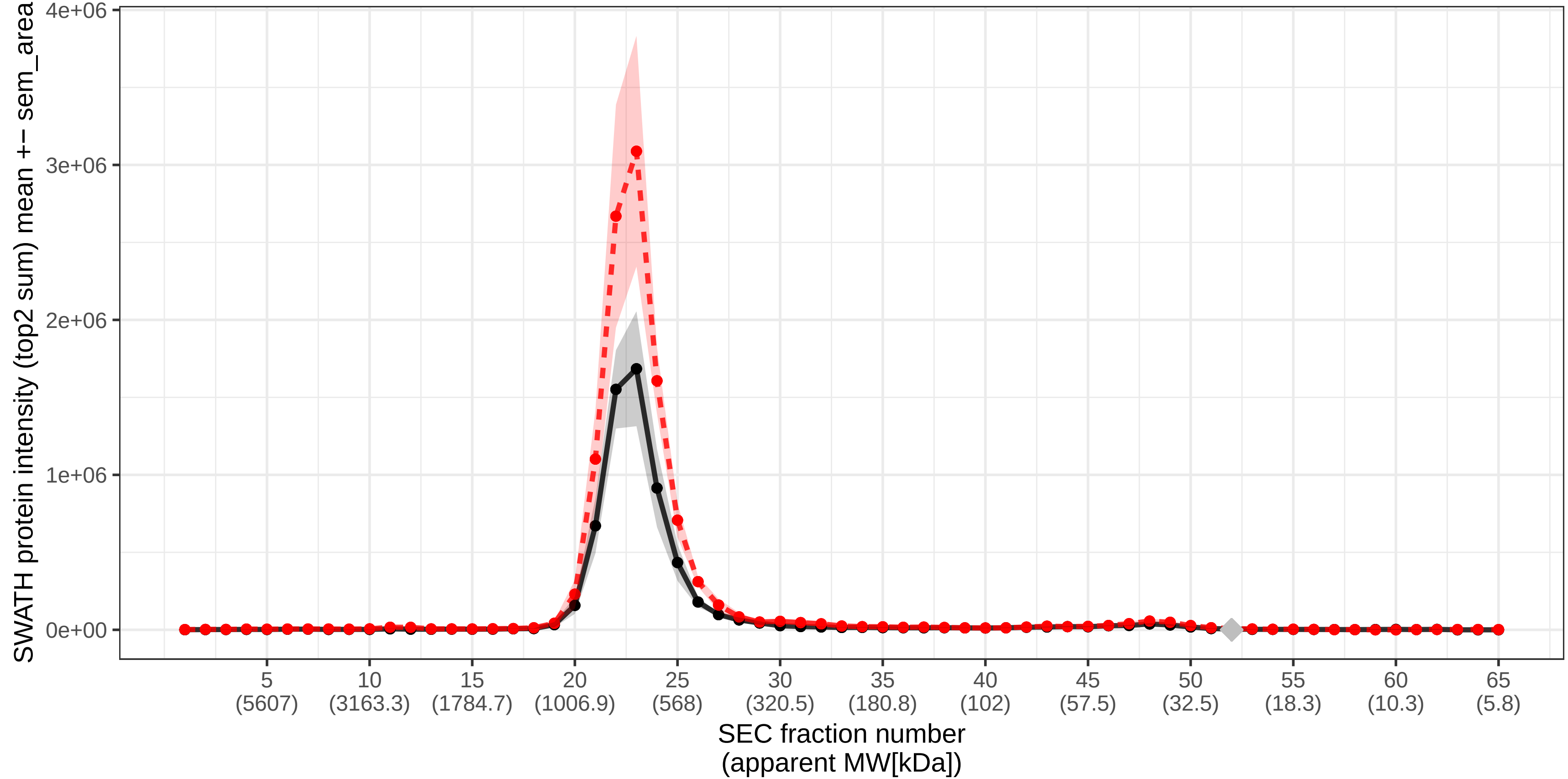

### SECchrom_O14545_TRAD1_HUMAN_TRAFD1_FLN29.pdf

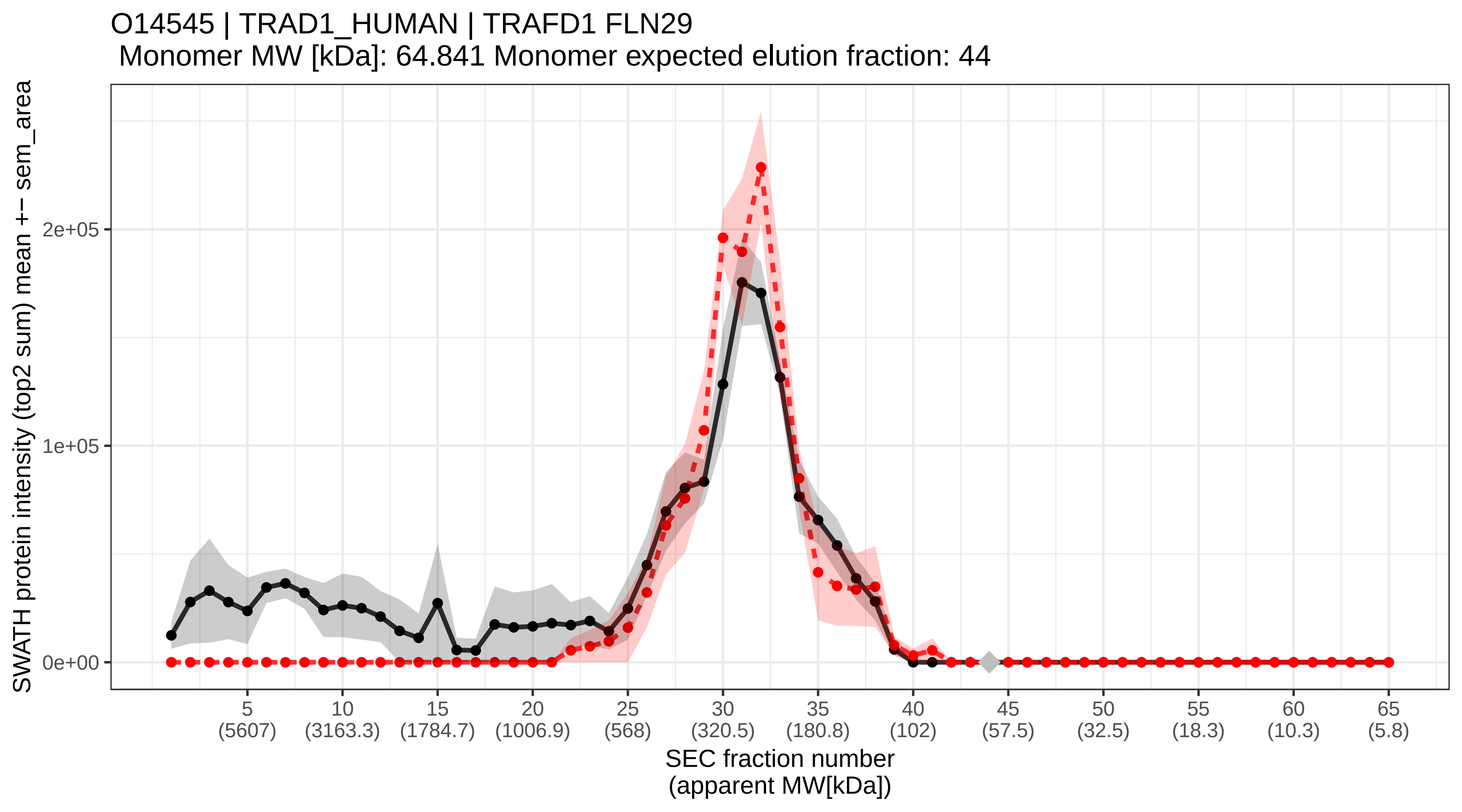

### SECchrom_O14548_COX7R_HUMAN_COX7A2L_COX7AR_COX7RP.pdf

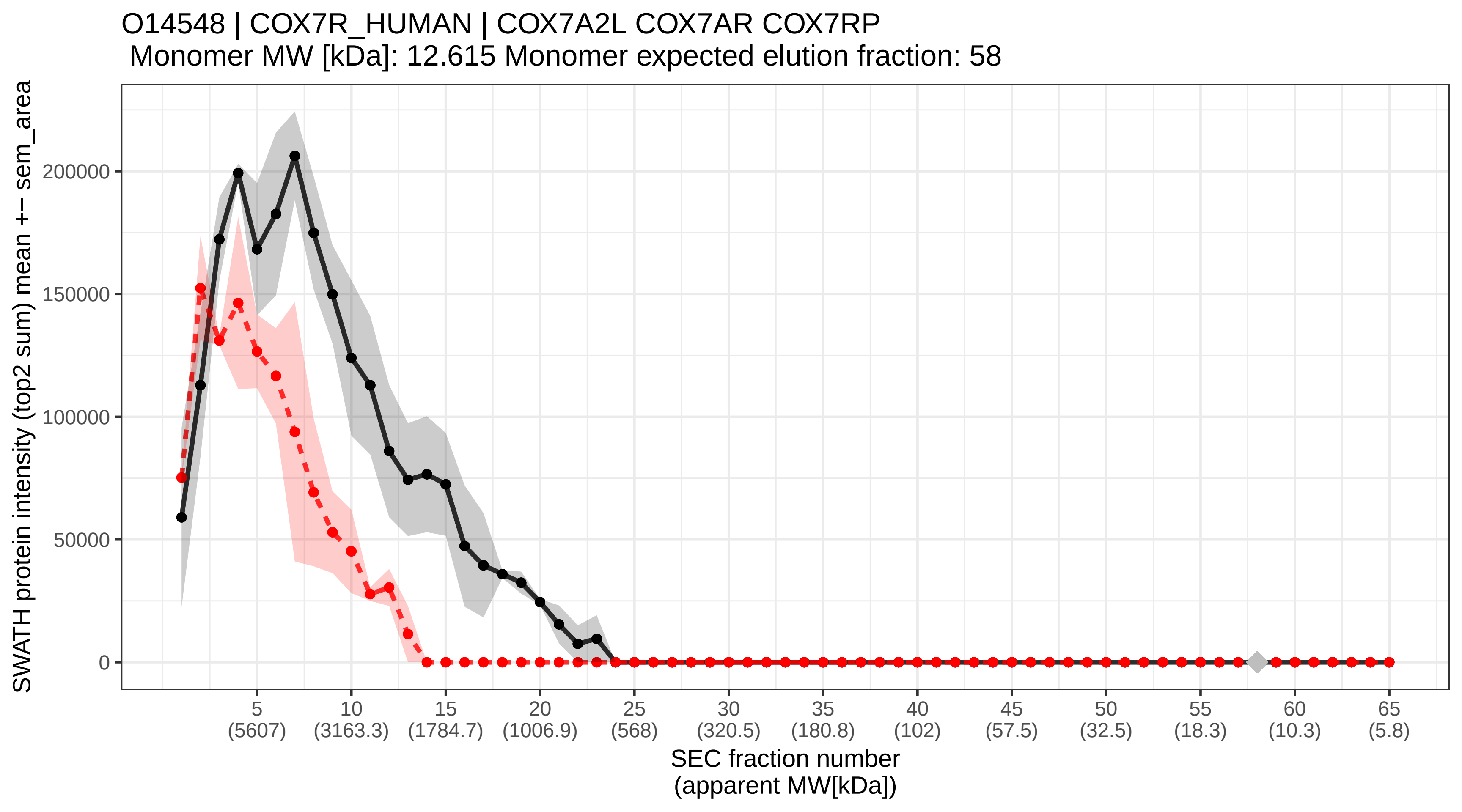

### SECchrom_O14561_ACPM_HUMAN_NDUFAB1.pdf

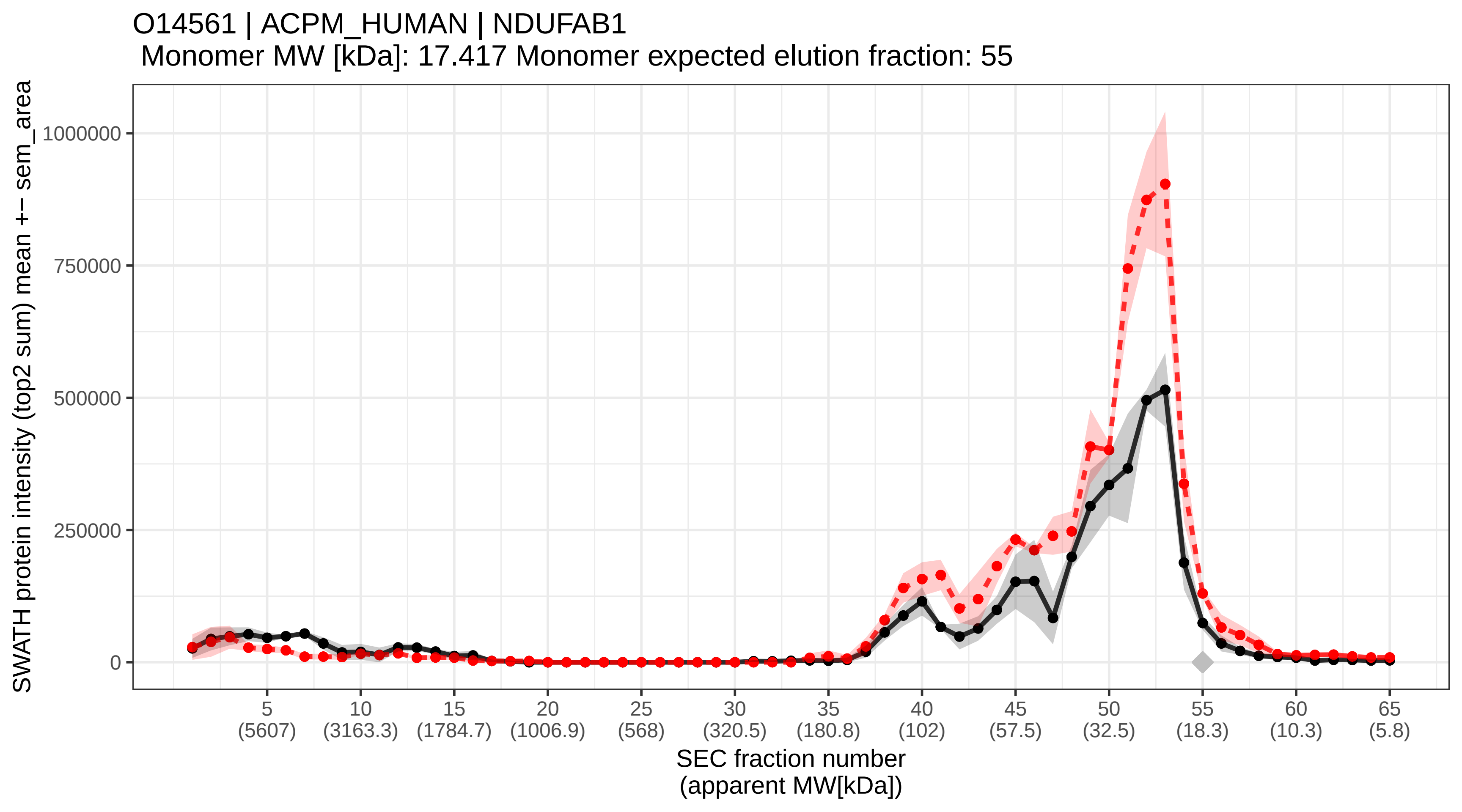

### SECchrom_O14562_UBFD1_HUMAN_UBFD1_UBPH.pdf

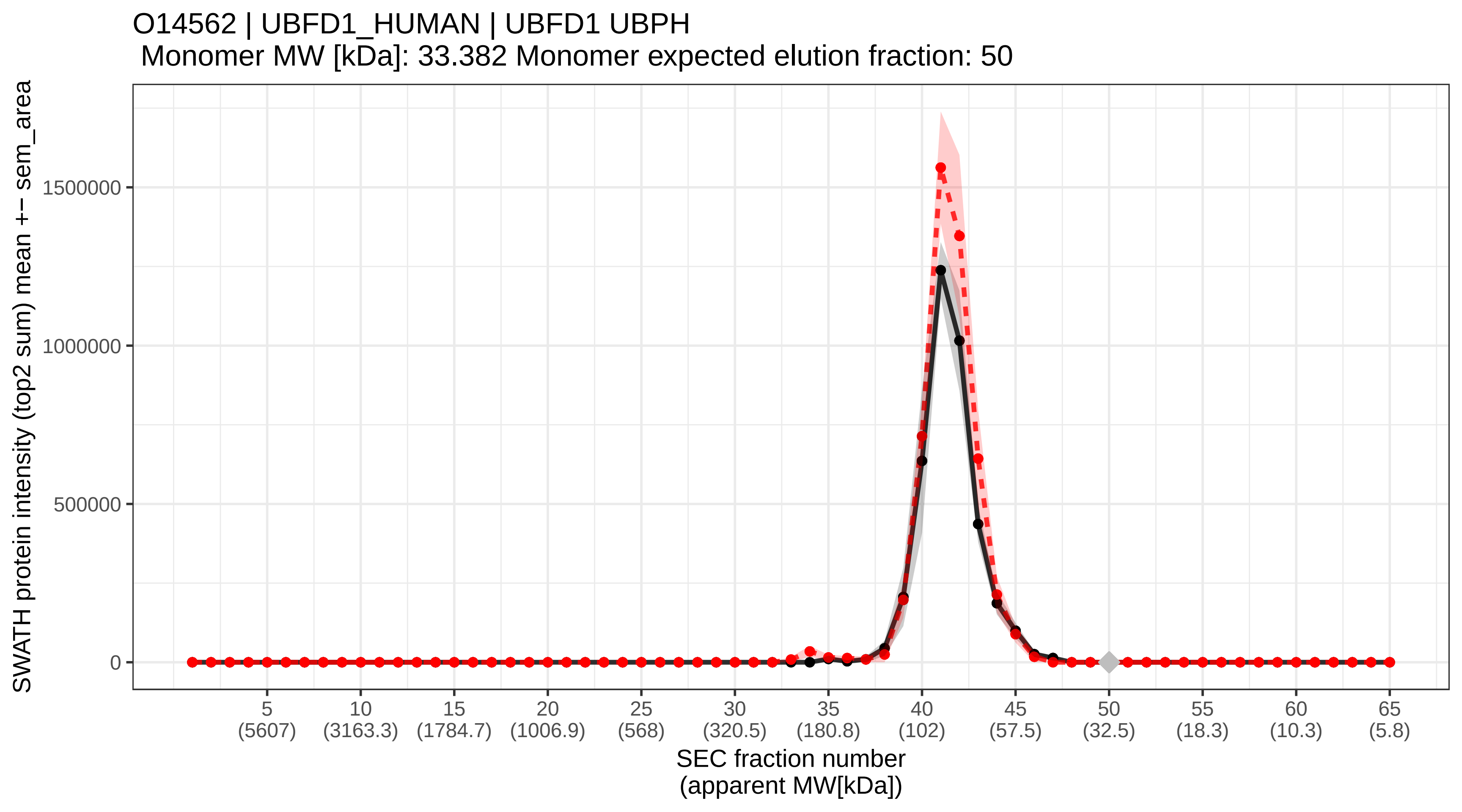

### SECchrom_O14578_CTRO_HUMAN_CIT_CRIK_KIAA0949_STK21.pdf

O14578 | CTRO_HUMAN | CIT CRIK KIAA0949 STK21
Monomer MW [kDa]: 231.431 Monomer expected elution fraction: 33

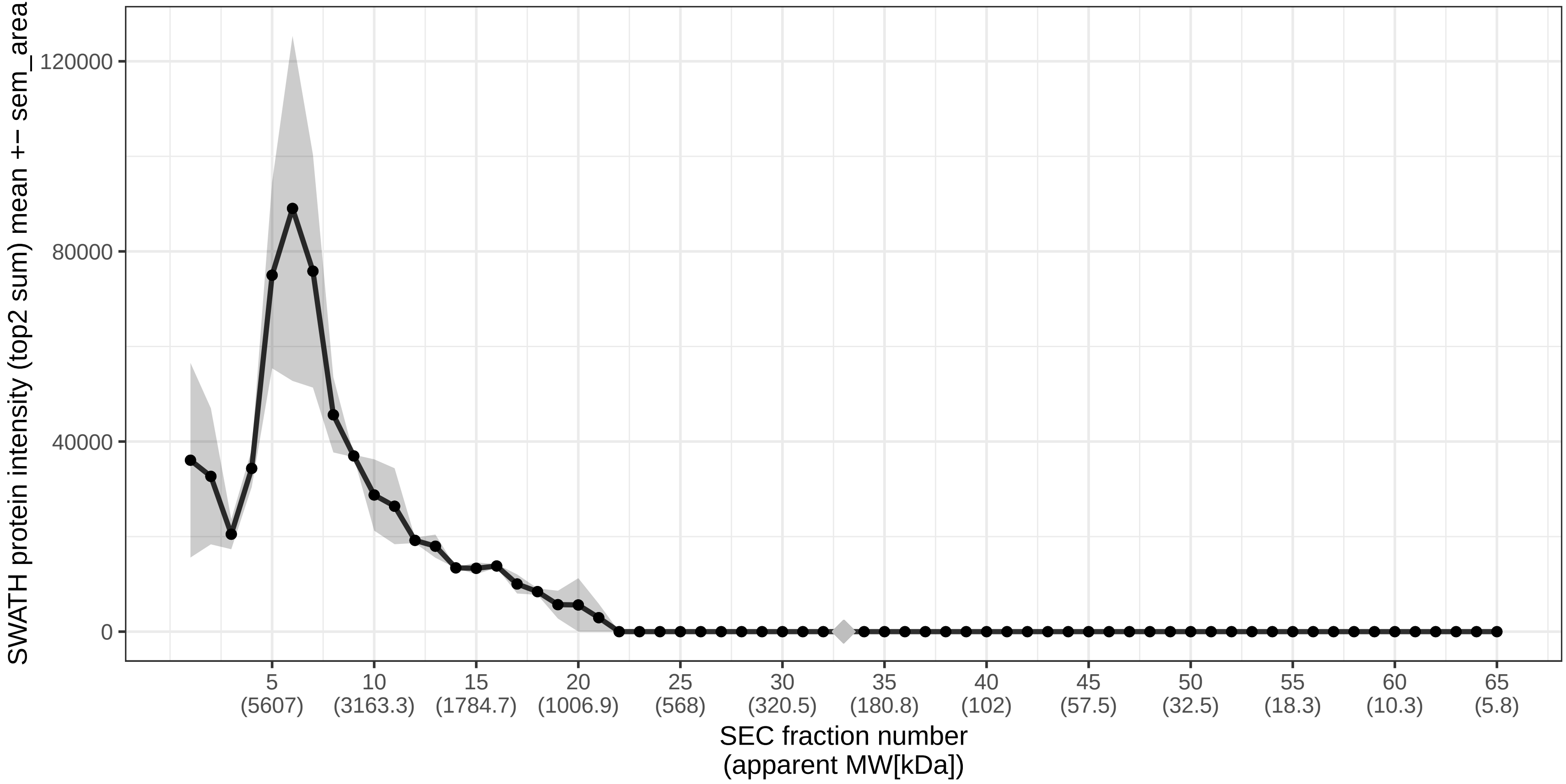

### SECchrom_O14579_COPE_HUMAN_COPE.pdf

O14579 | COPE_HUMAN | COPE
Monomer MW [kDa]: 34.482 Monomer expected elution fraction: 49

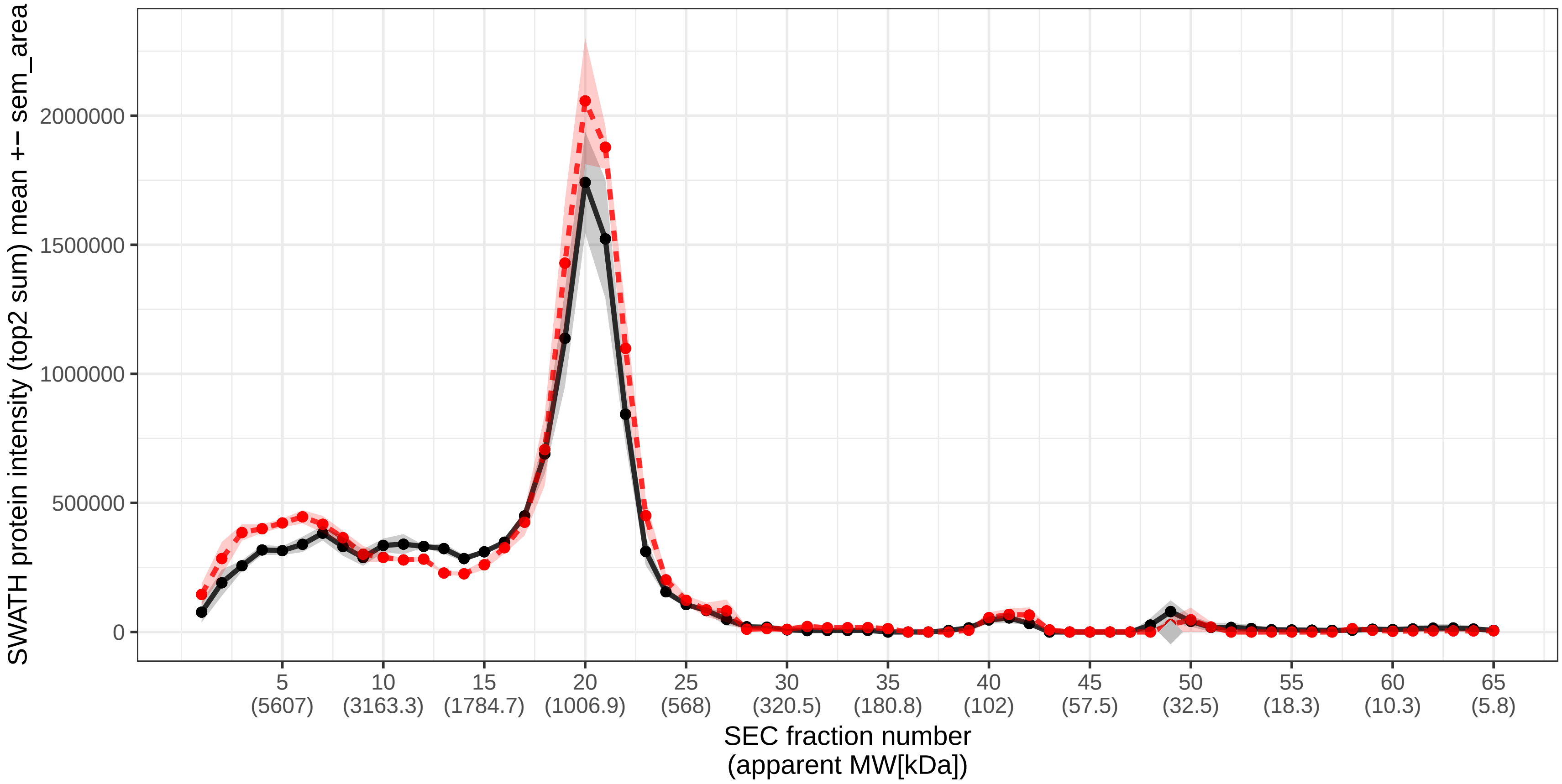

### SECchrom_O14617_AP3D1_HUMAN_AP3D1_PRO0039.pdf

O14617 | AP3D1_HUMAN | AP3D1 PRO0039
Monomer MW [kDa]: 130.158 Monomer expected elution fraction: 38

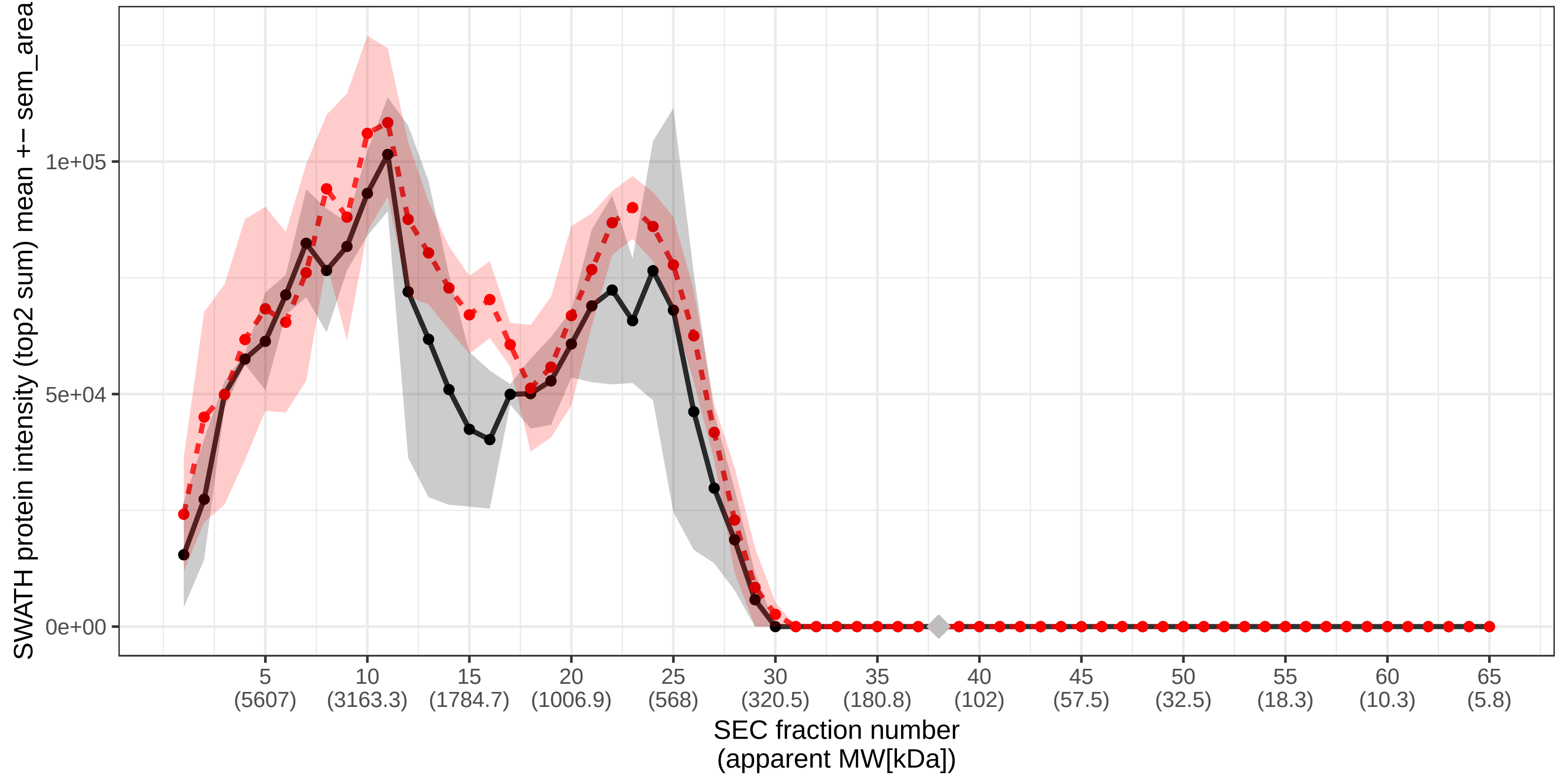

### SECchrom_O14618_CCS_HUMAN_CCS.pdf

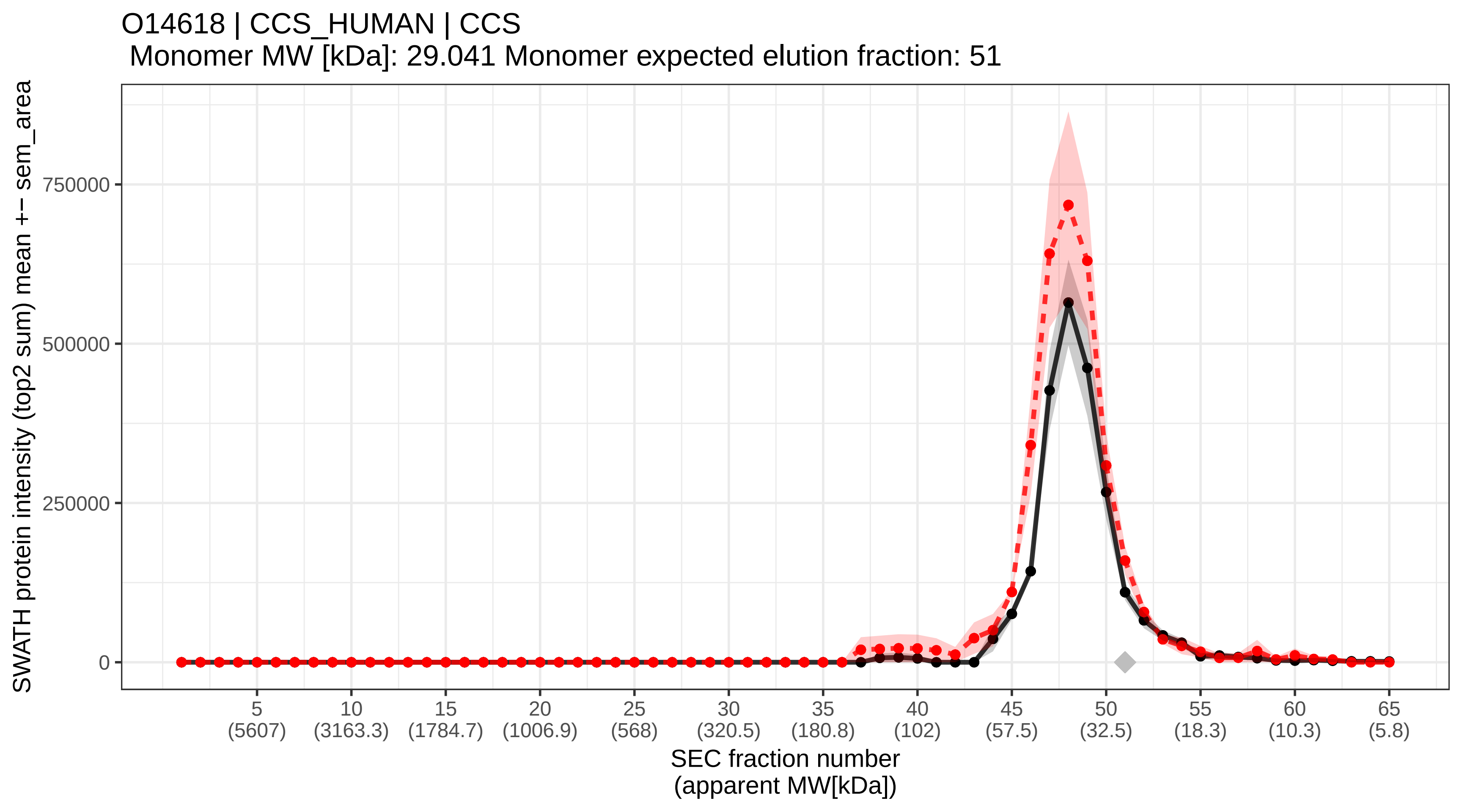

### SECchrom_O14639_ABLM1_HUMAN_ABLIM1_ABLIM_KIAA0059_LIMAB1.pdf

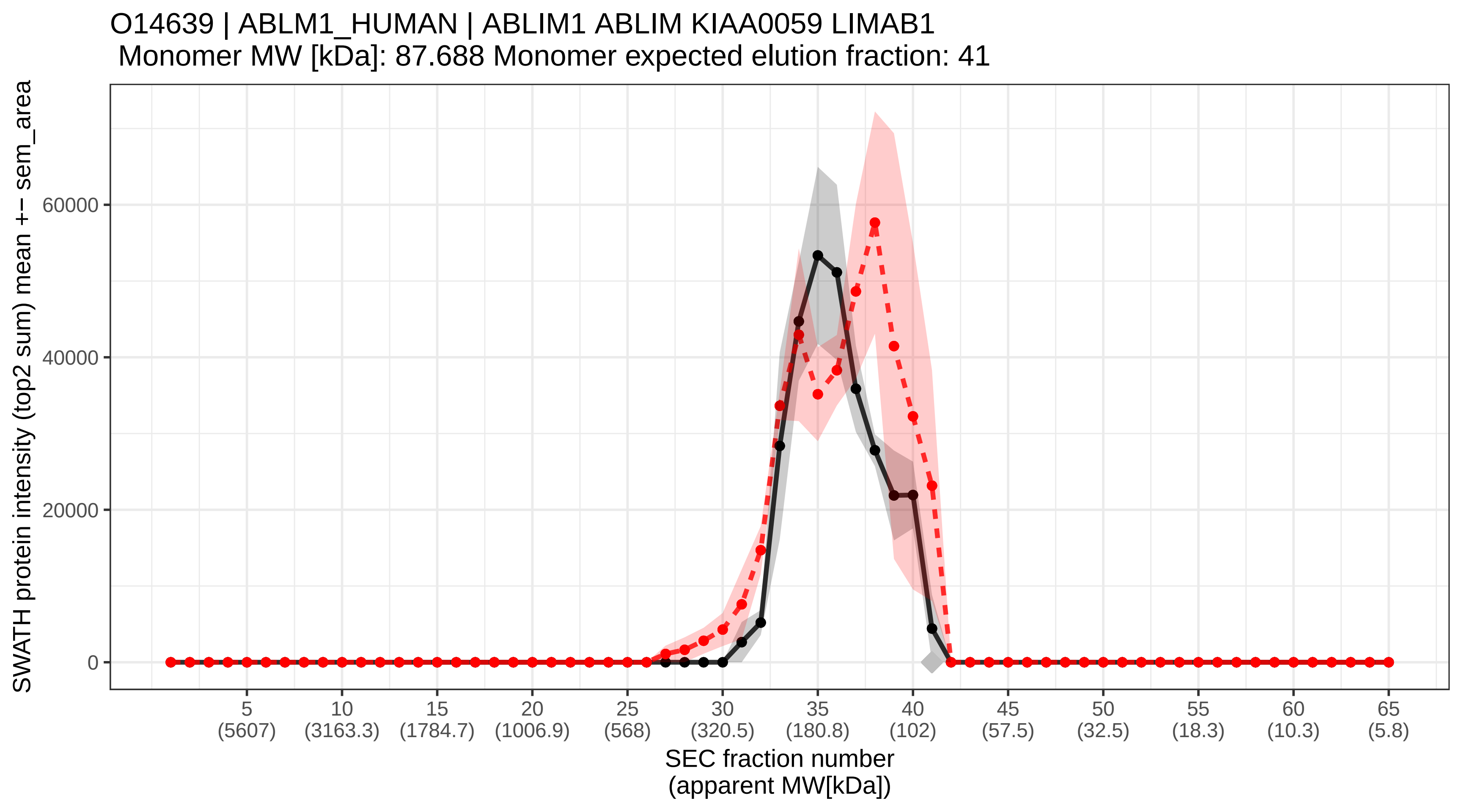

### SECchrom_O14641_DVL2_HUMAN_DVL2.pdf

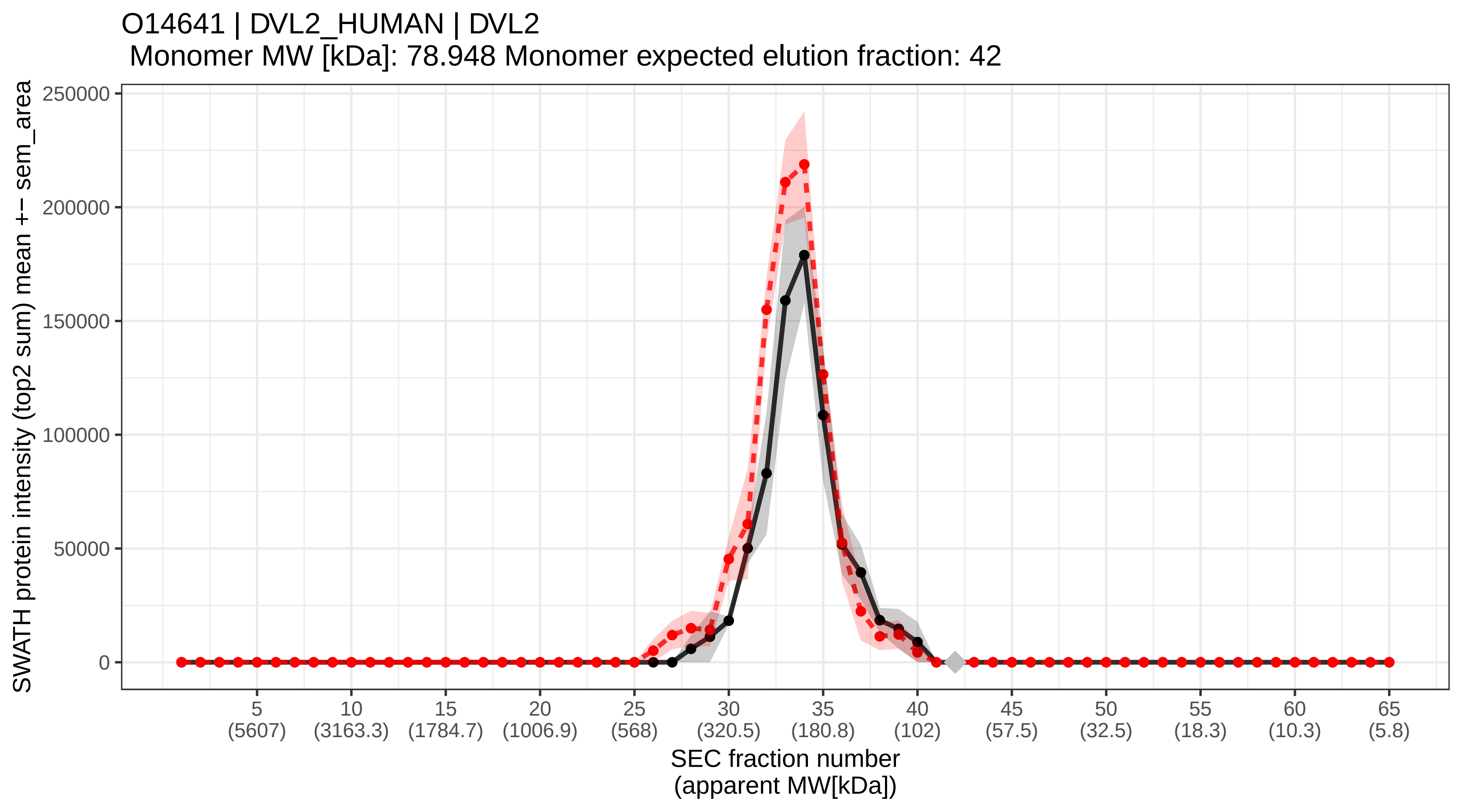

### SECchrom_O14656_TOR1A_HUMAN_TOR1A_DQ2_DYT1_TA_TORA.pdf

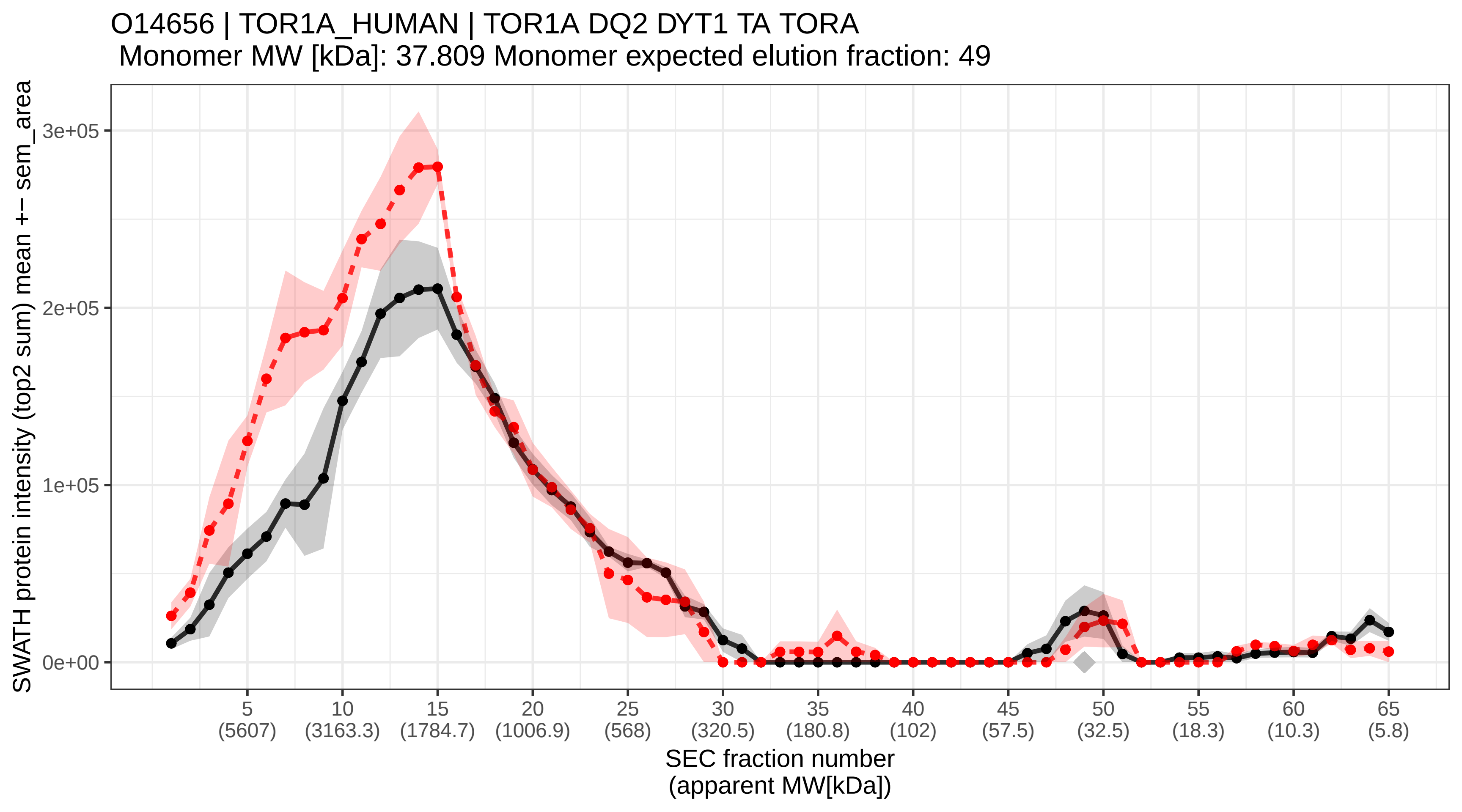

### SECchrom_O14657_TOR1B_HUMAN_TOR1B_DQ1_FKSG18.pdf

#### O14657 | TOR1B_HUMAN | TOR1B DQ1 FKSG18
Monomer MW [kDa]: 37.979 Monomer expected elution fraction: 49

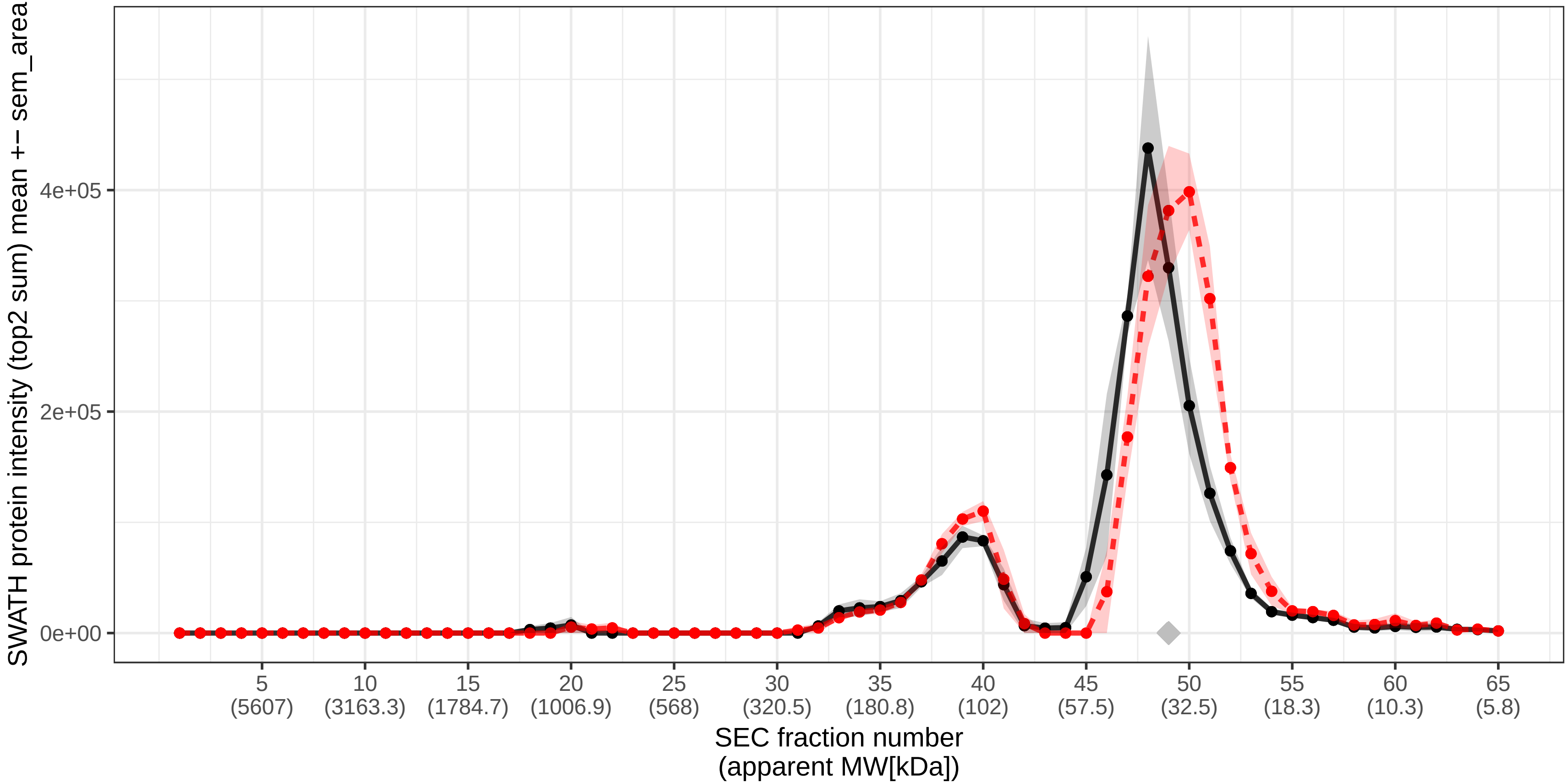

### SECchrom_O14662_STX16_HUMAN_STX16.pdf

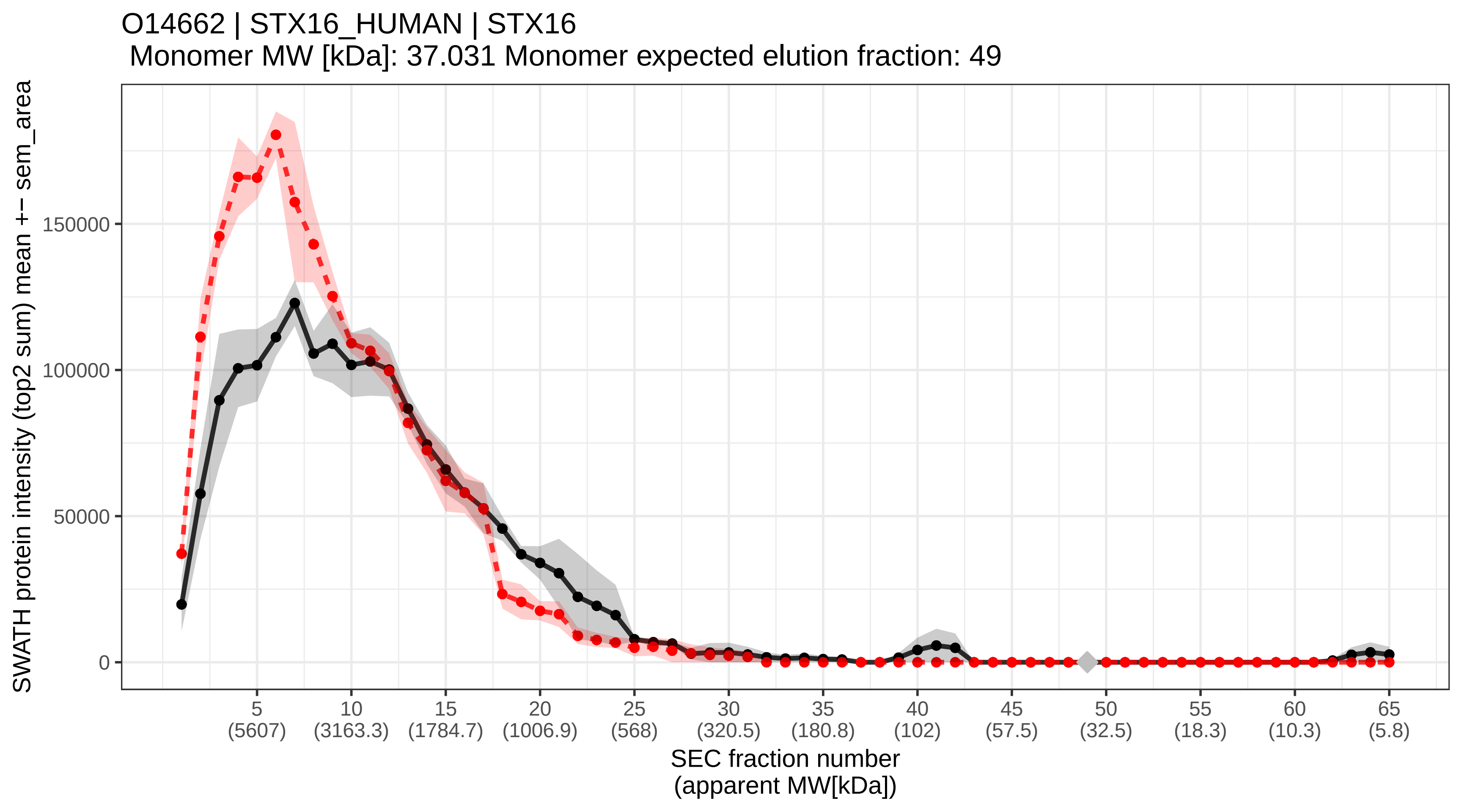

### SECchrom_O14672_ADA10_HUMAN_ADAM10_KUZ_MADM.pdf

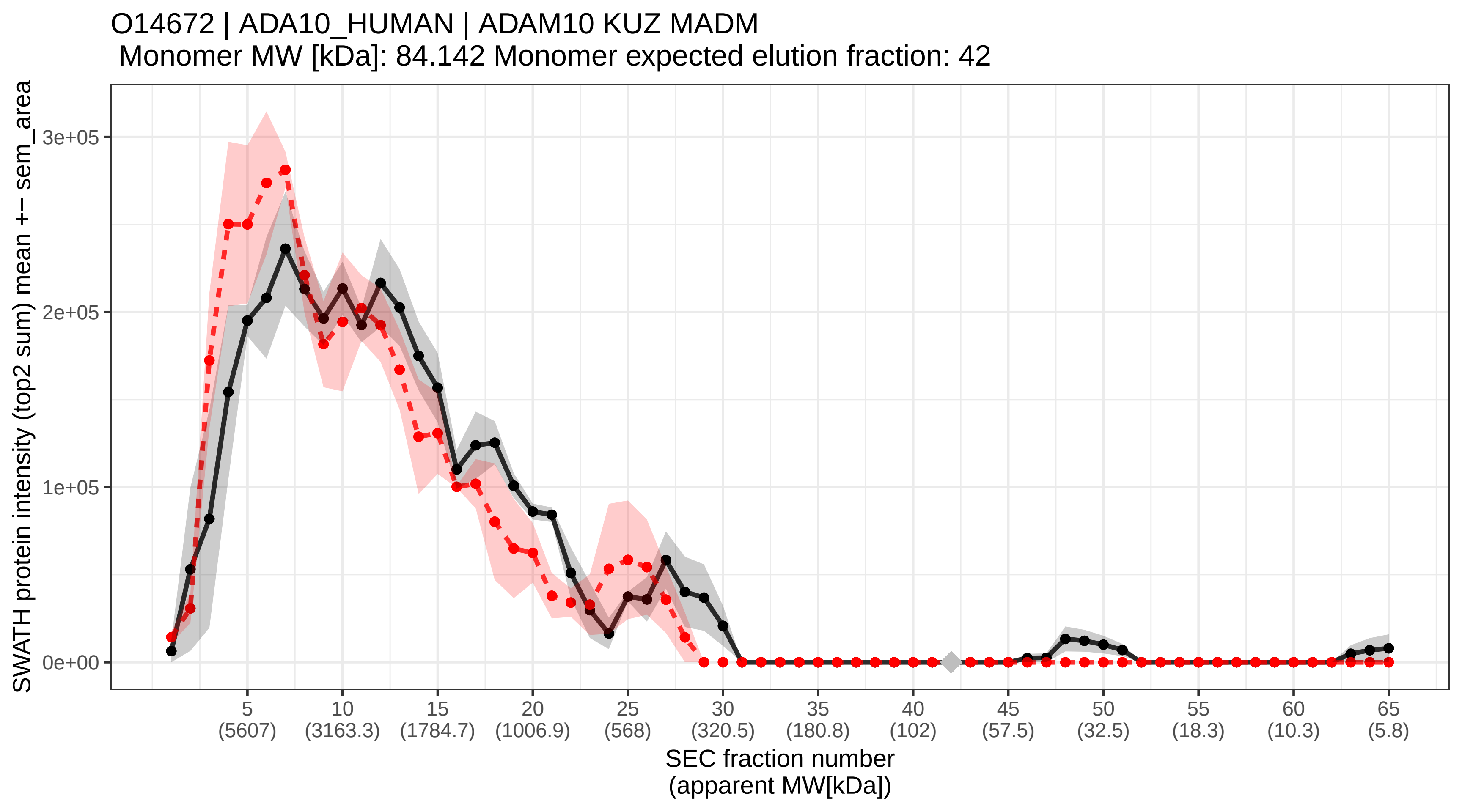

### SECchrom_O14684_PTGES_HUMAN_PTGES_MGST1L1_MPGES1_PGES_PIG12.pdf

O14684 | PTGES_HUMAN | PTGES MGST1L1 MPGES1 PGES PIG12
Monomer MW [kDa]: 17.102 Monomer expected elution fraction: 56

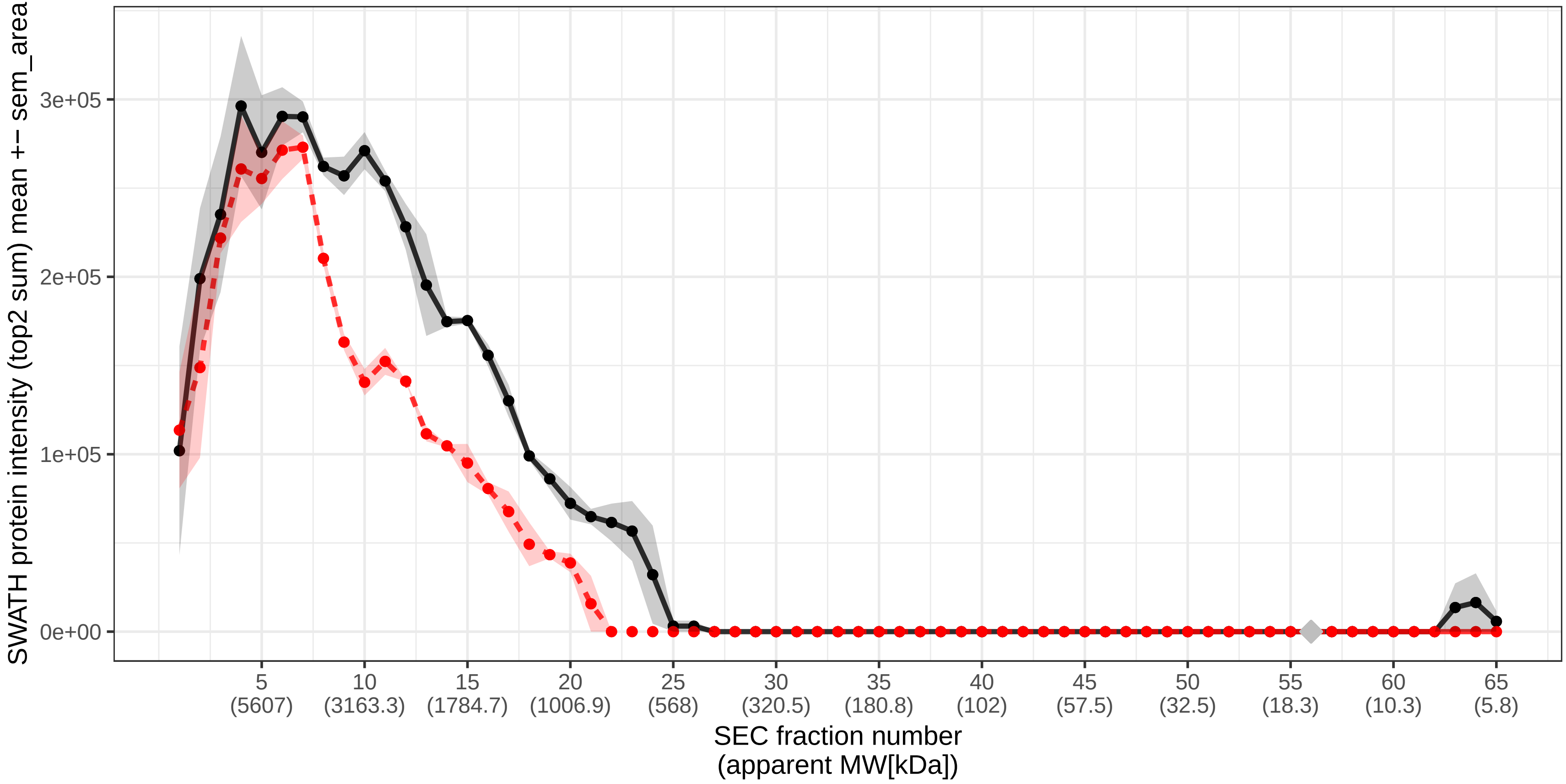

### SECchrom_O14727_APAF_HUMAN_APAF1_KIAA0413.pdf

O14727 | APAF_HUMAN | APAF1 KIAA0413
Monomer MW [kDa]: 141.84 Monomer expected elution fraction: 37

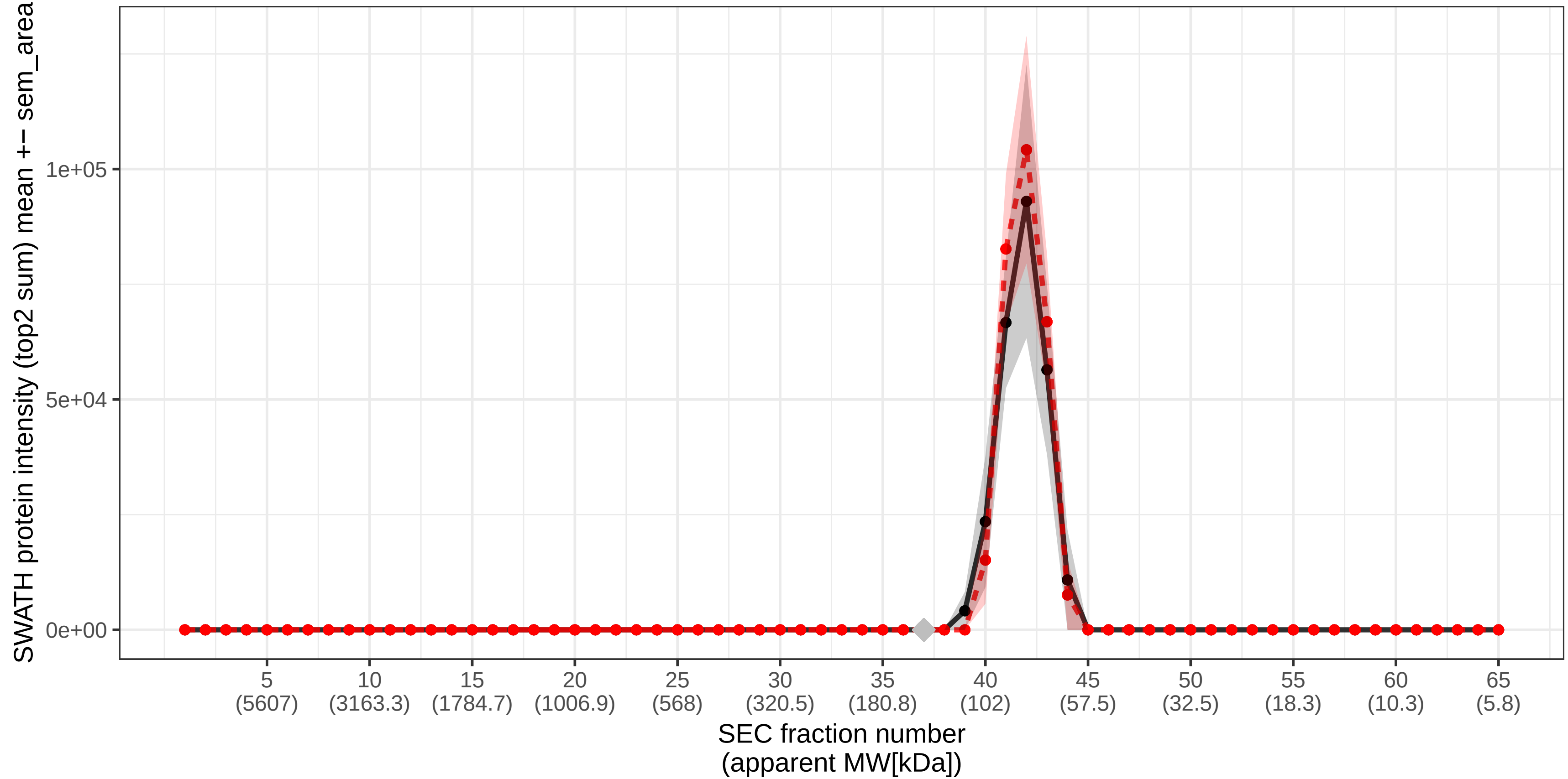

### SECchrom_O14732_IMPA2_HUMAN_IMPA2_IMP.18P.pdf

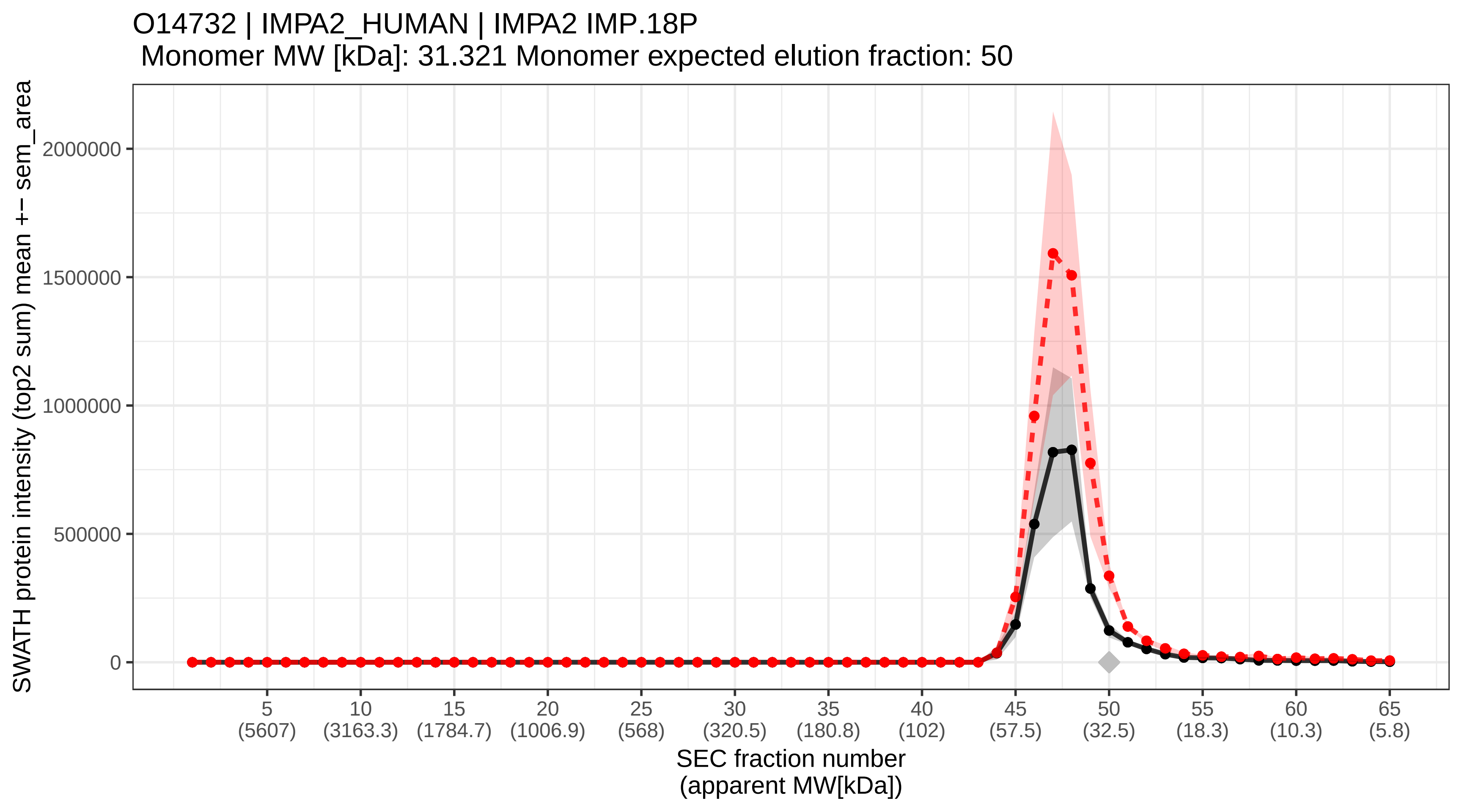

### SECchrom_O14733_MP2K7_HUMAN_MAP2K7_JNKK2_MEK7_MKK7_PRKMK7_S.pdf

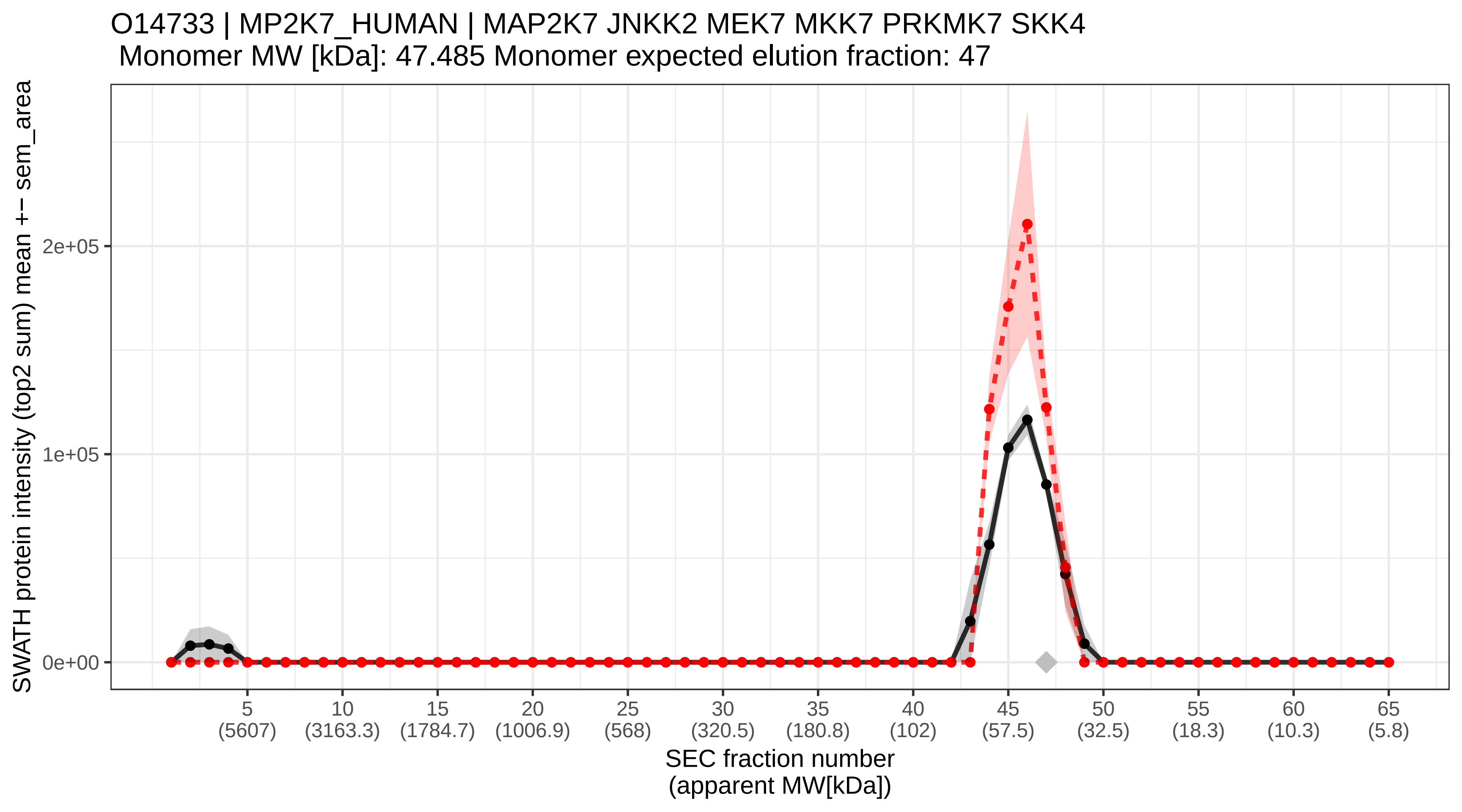

### SECchrom_O14734_ACOT8_HUMAN_ACOT8_ACTEIII_PTE1_PTE2.pdf

O14734 | ACOT8_HUMAN | ACOT8 ACTEIII PTE1 PTE2
Monomer MW [kDa]: 35.914 Monomer expected elution fraction: 49

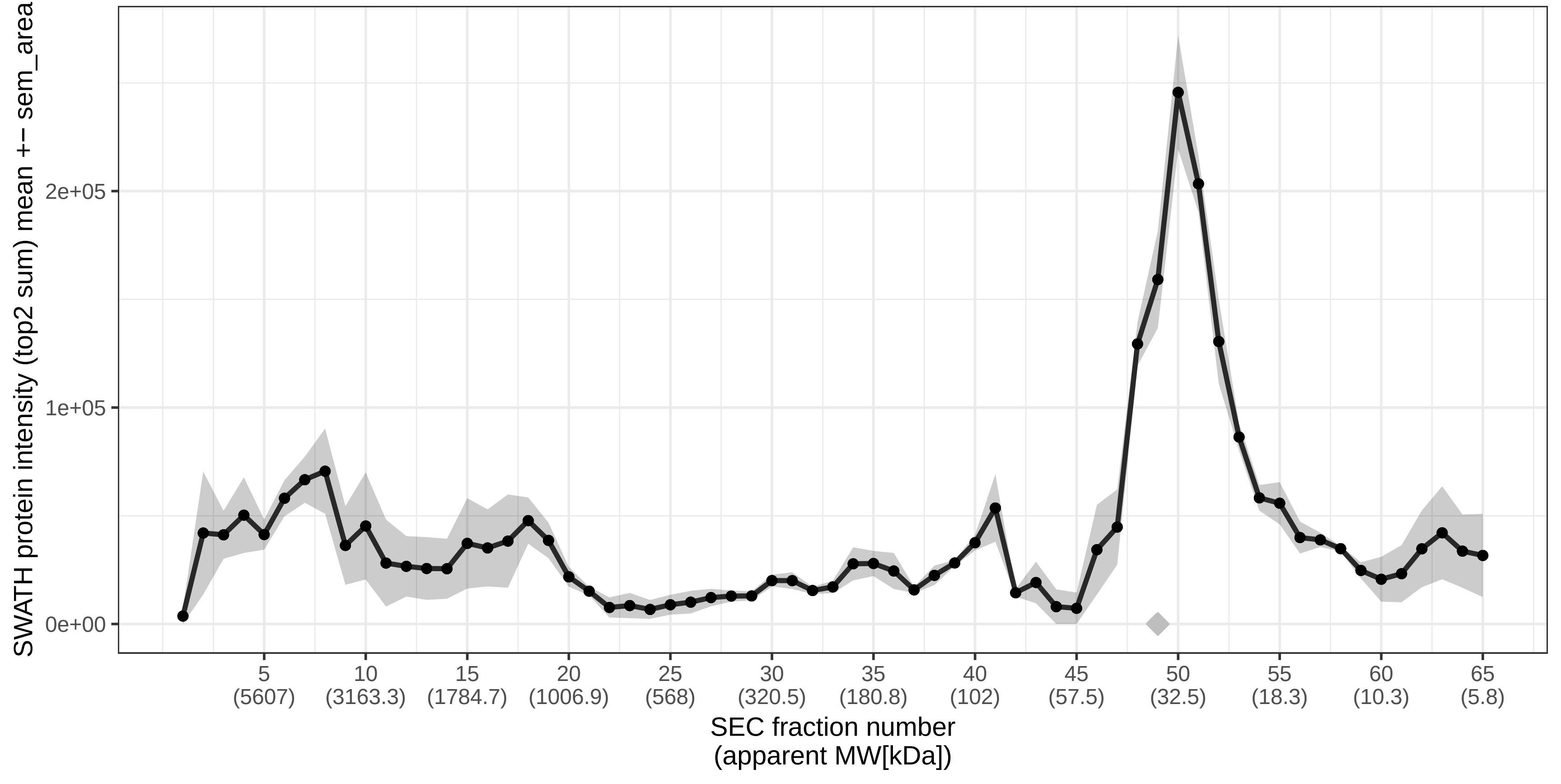

### SECchrom_O14735_CDIPT_HUMAN_CDIPT_PIS_PIS1.pdf

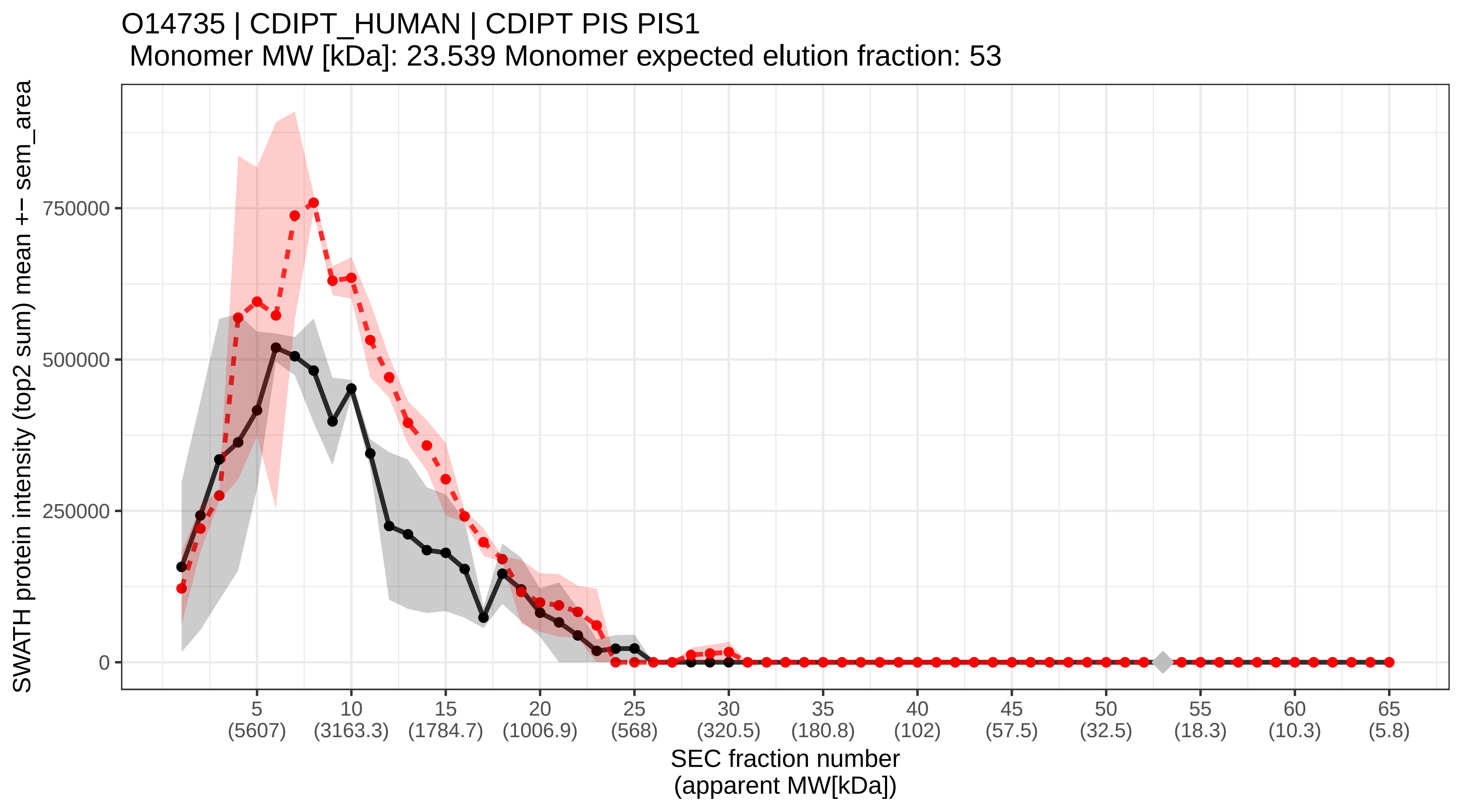

### SECchrom_O14737_PDCD5_HUMAN_PDCD5_TFAR19.pdf

O14737 | PDCD5_HUMAN | PDCD5 TFAR19
Monomer MW [kDa]: 14.285 Monomer expected elution fraction: 57

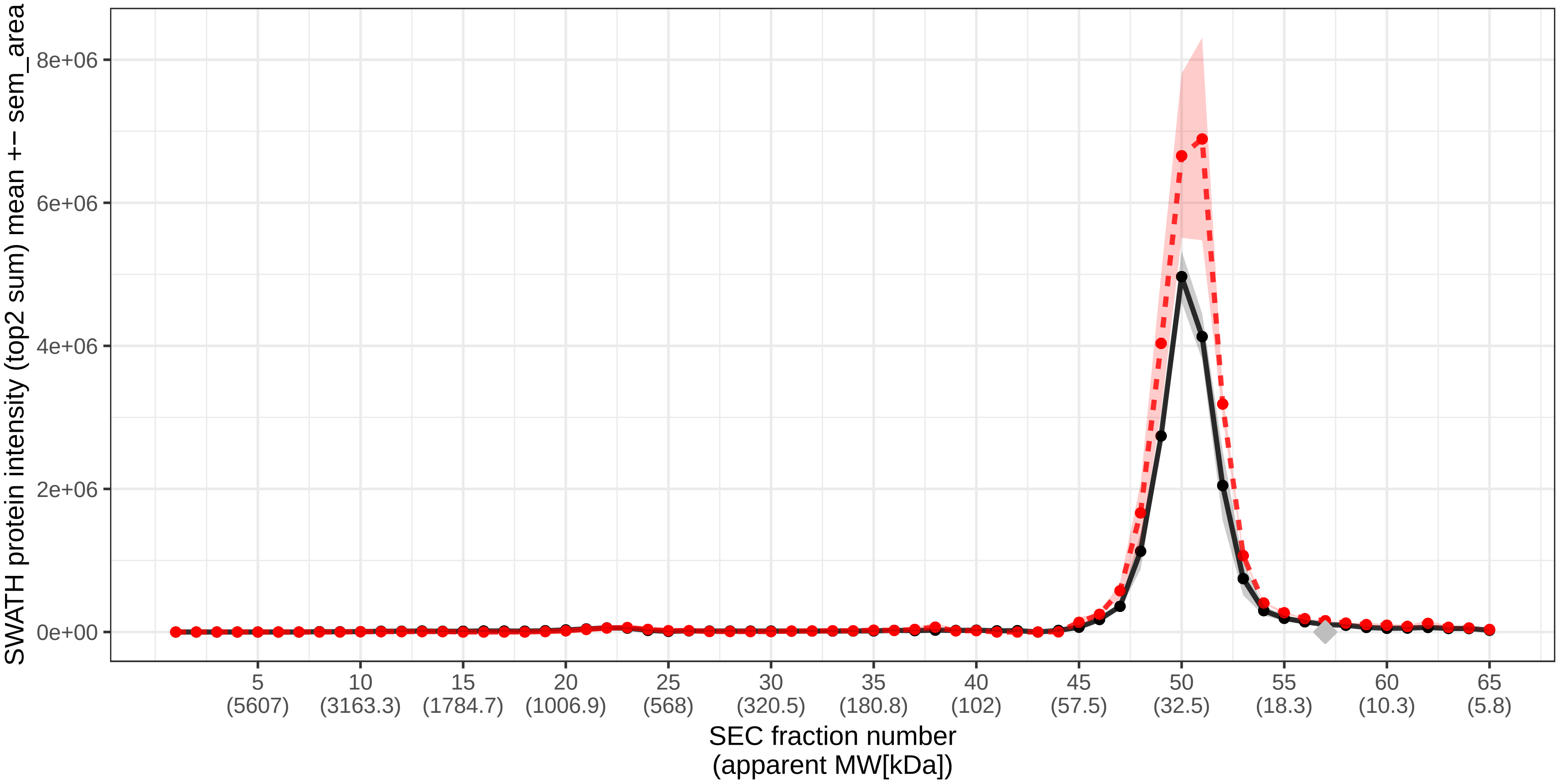

### SECchrom_O14744_ANM5_HUMAN_PRMT5_HRMT1L5_IBP72_JBP1_SKB1.pdf

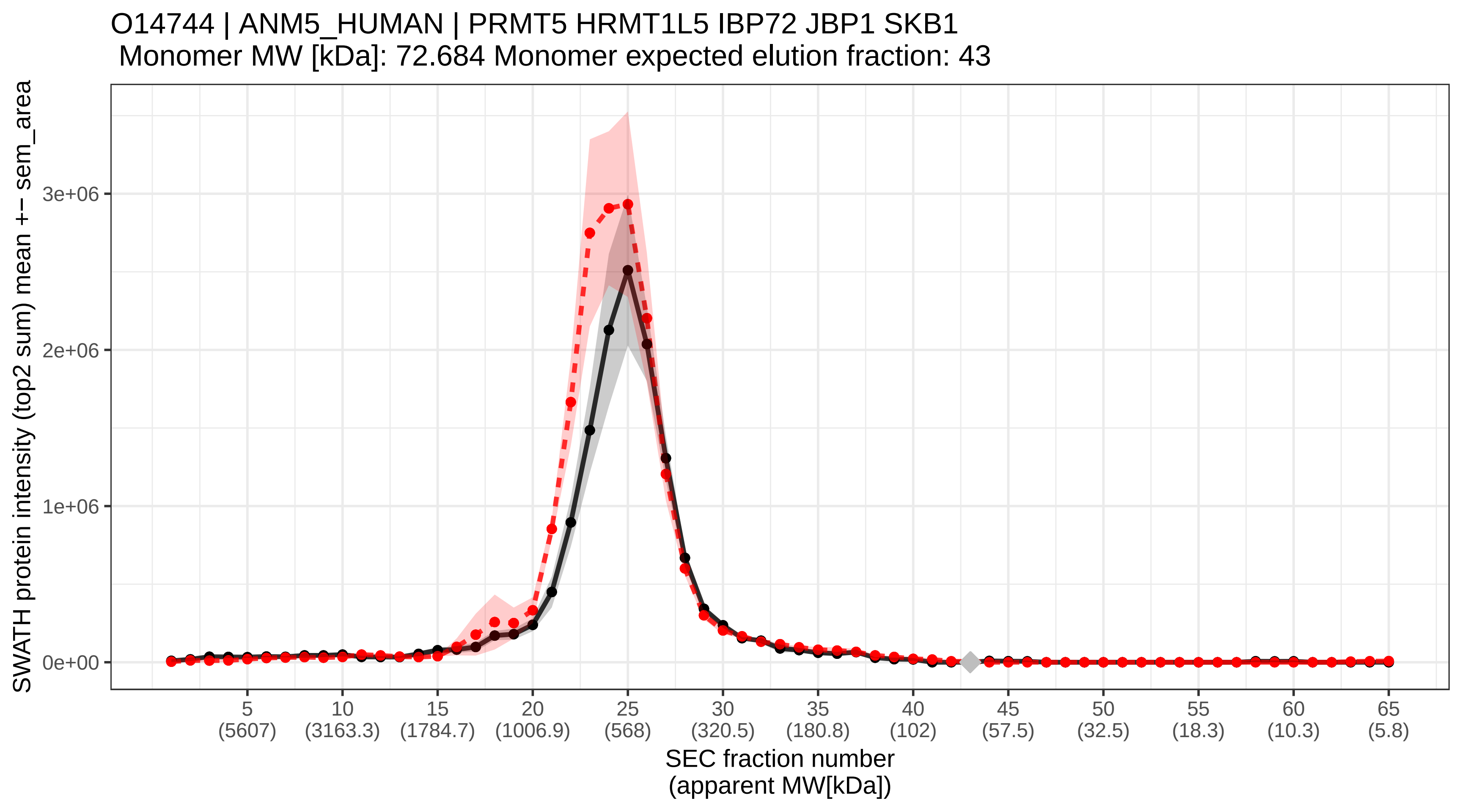

### SECchrom_O14745_NHRF1_HUMAN_SLC9A3R1_NHERF_NHERF1.pdf

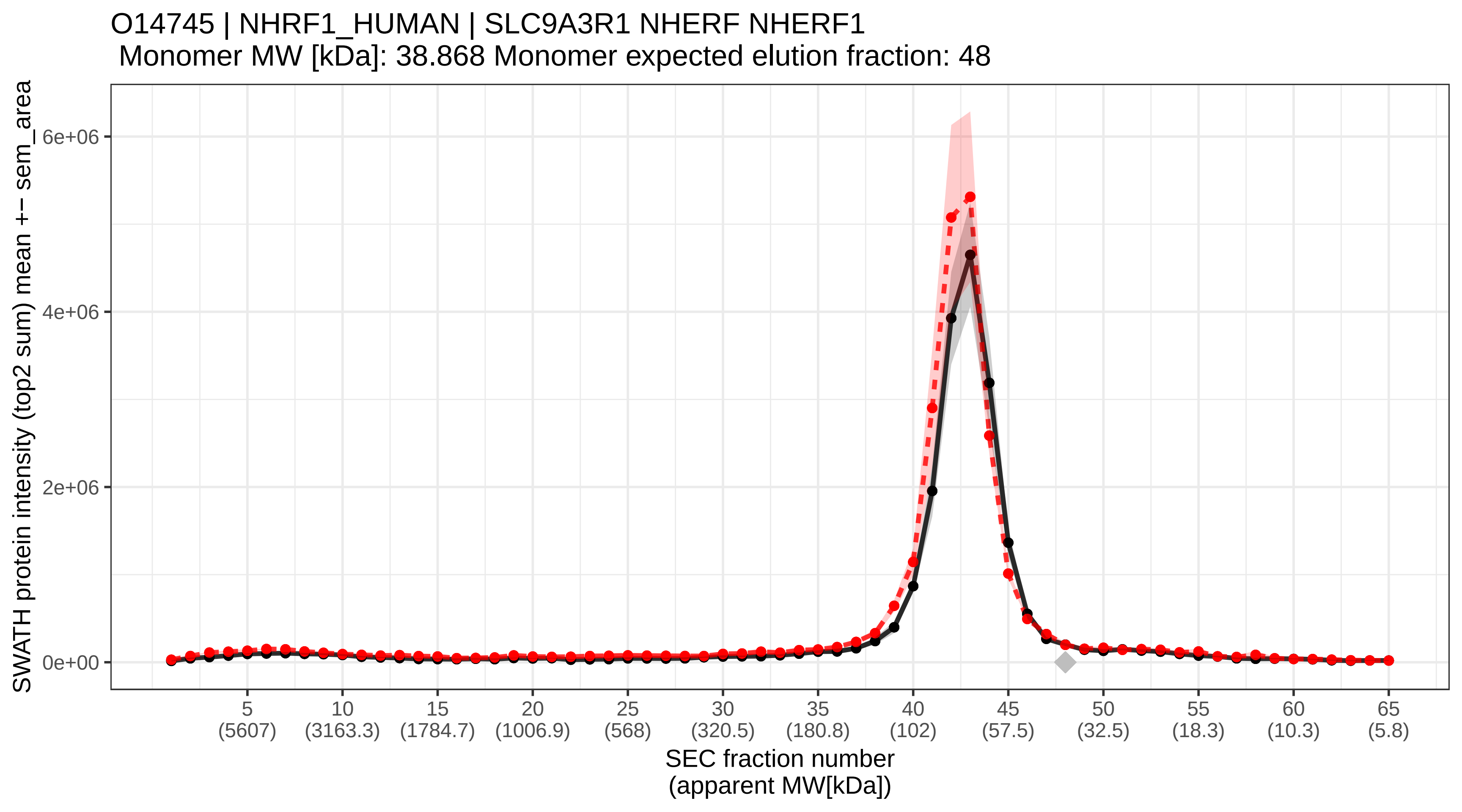

### SECchrom_O14757_CHK1_HUMAN_CHEK1_CHK1.pdf

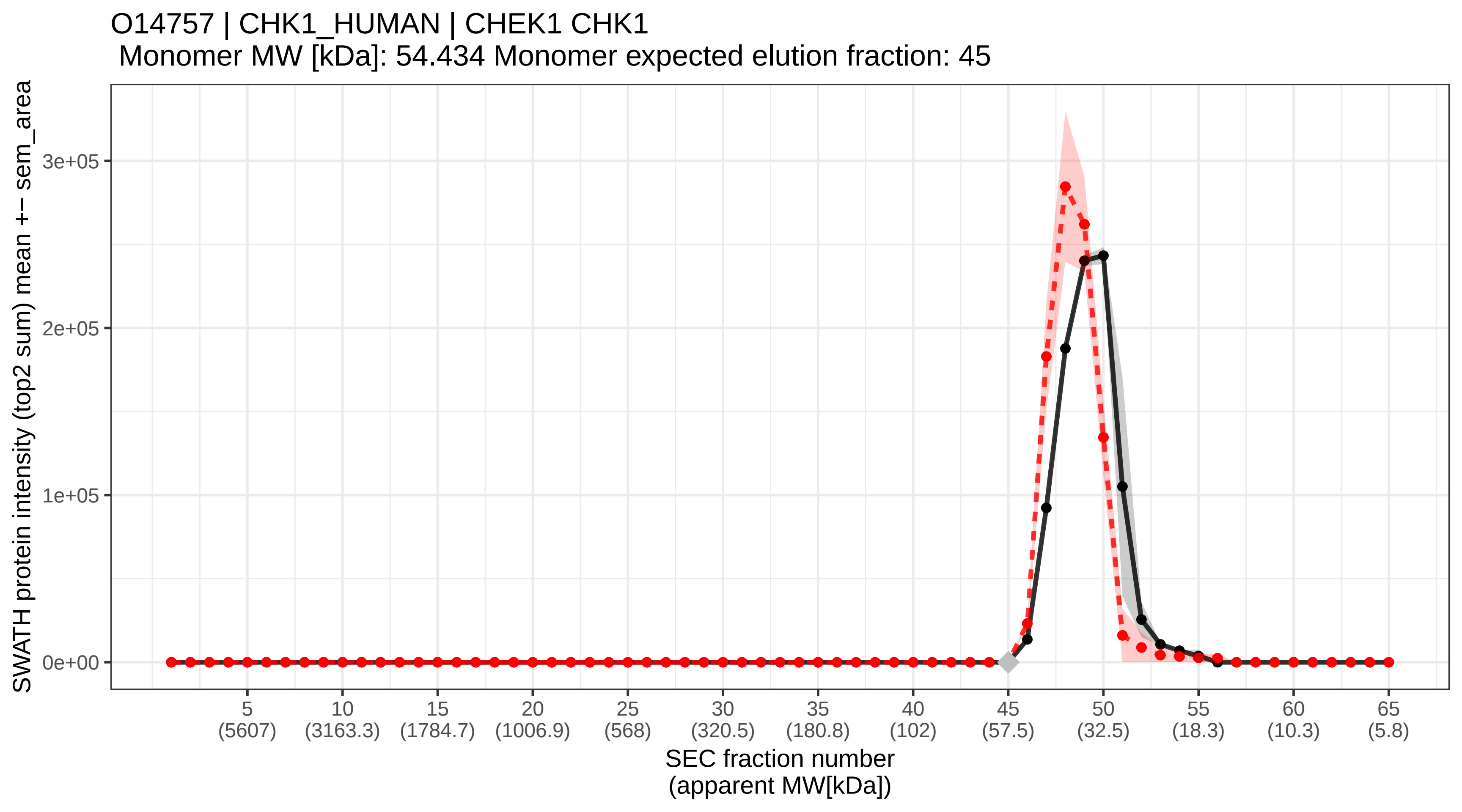

### SECchrom_O14776_TCRG1_HUMAN_TCERG1_CA150_TAF2S.pdf

O14776 | TCRG1_HUMAN | TCERG1 CA150 TAF2S
Monomer MW [kDa]: 123.901 Monomer expected elution fraction: 38

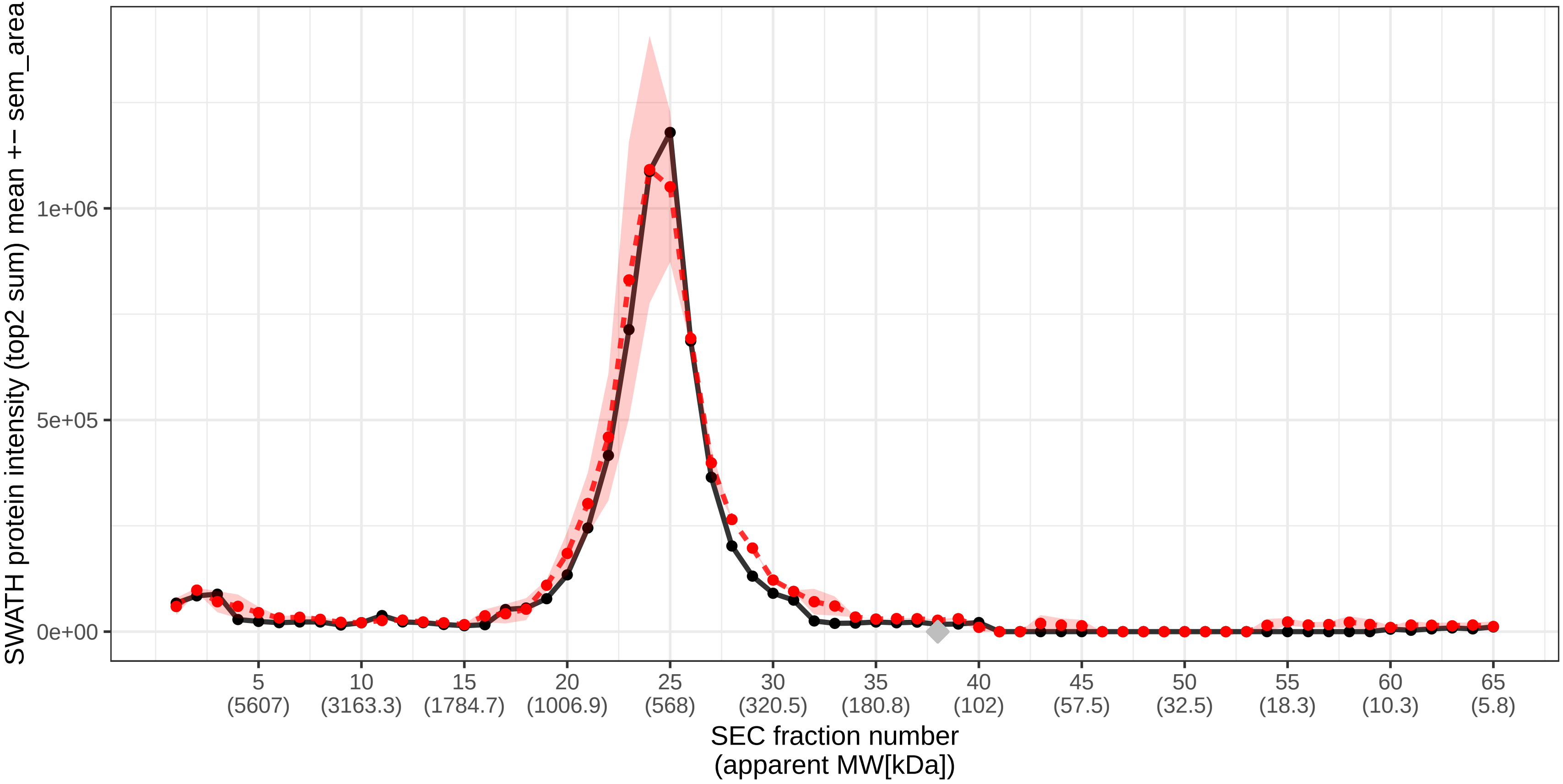

### SECchrom_O14777_NDC80_HUMAN_NDC80_HEC_HEC1_KNTC2.pdf

O14777 | NDC80_HUMAN | NDC80 HEC HEC1 KNTC2
Monomer MW [kDa]: 73.913 Monomer expected elution fraction: 43

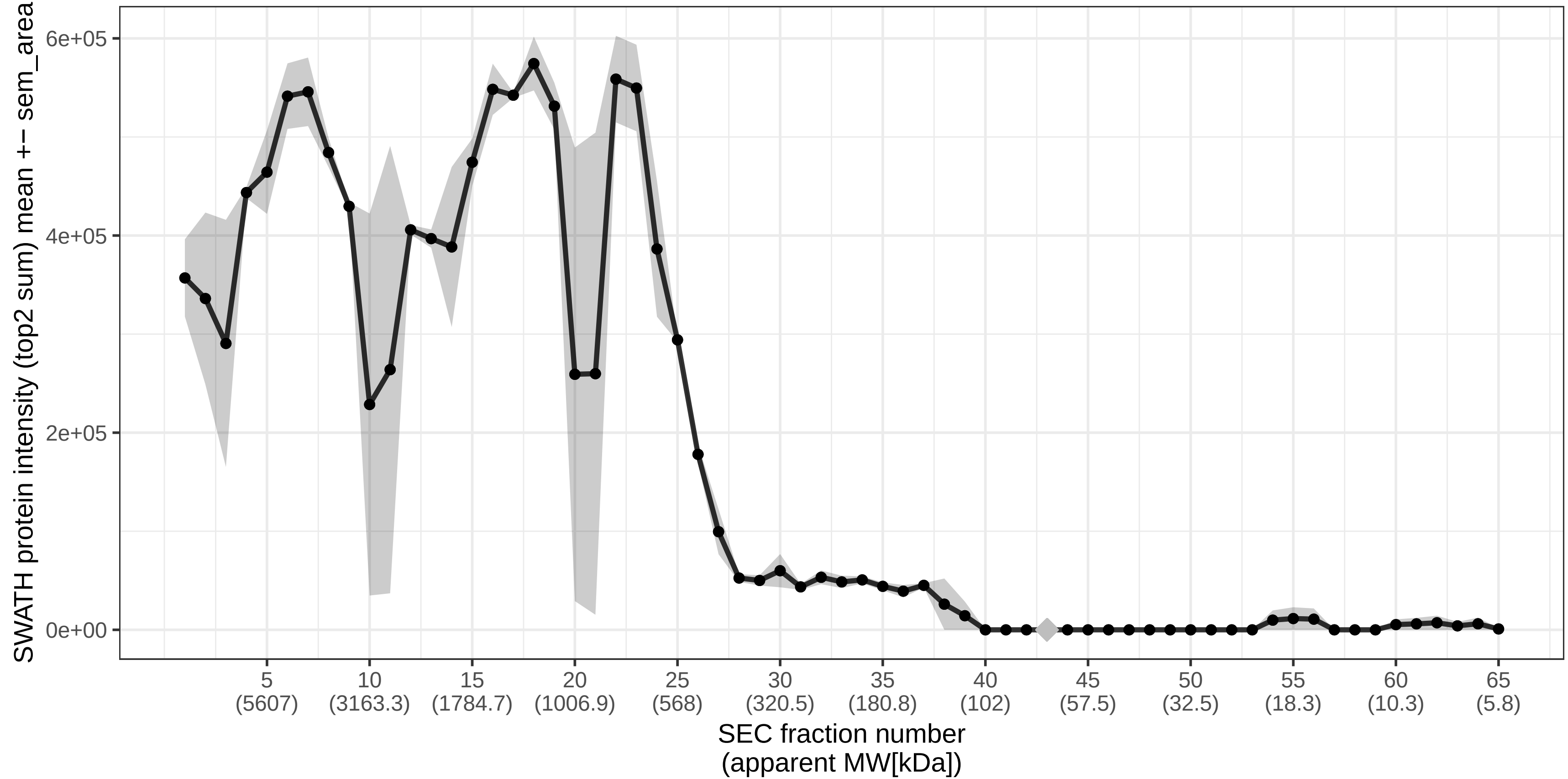

### SECchrom_O14787_TNPO2_HUMAN_TNPO2.pdf

O14787 | TNPO2_HUMAN | TNPO2
Monomer MW [kDa]: 101.388 Monomer expected elution fraction: 40

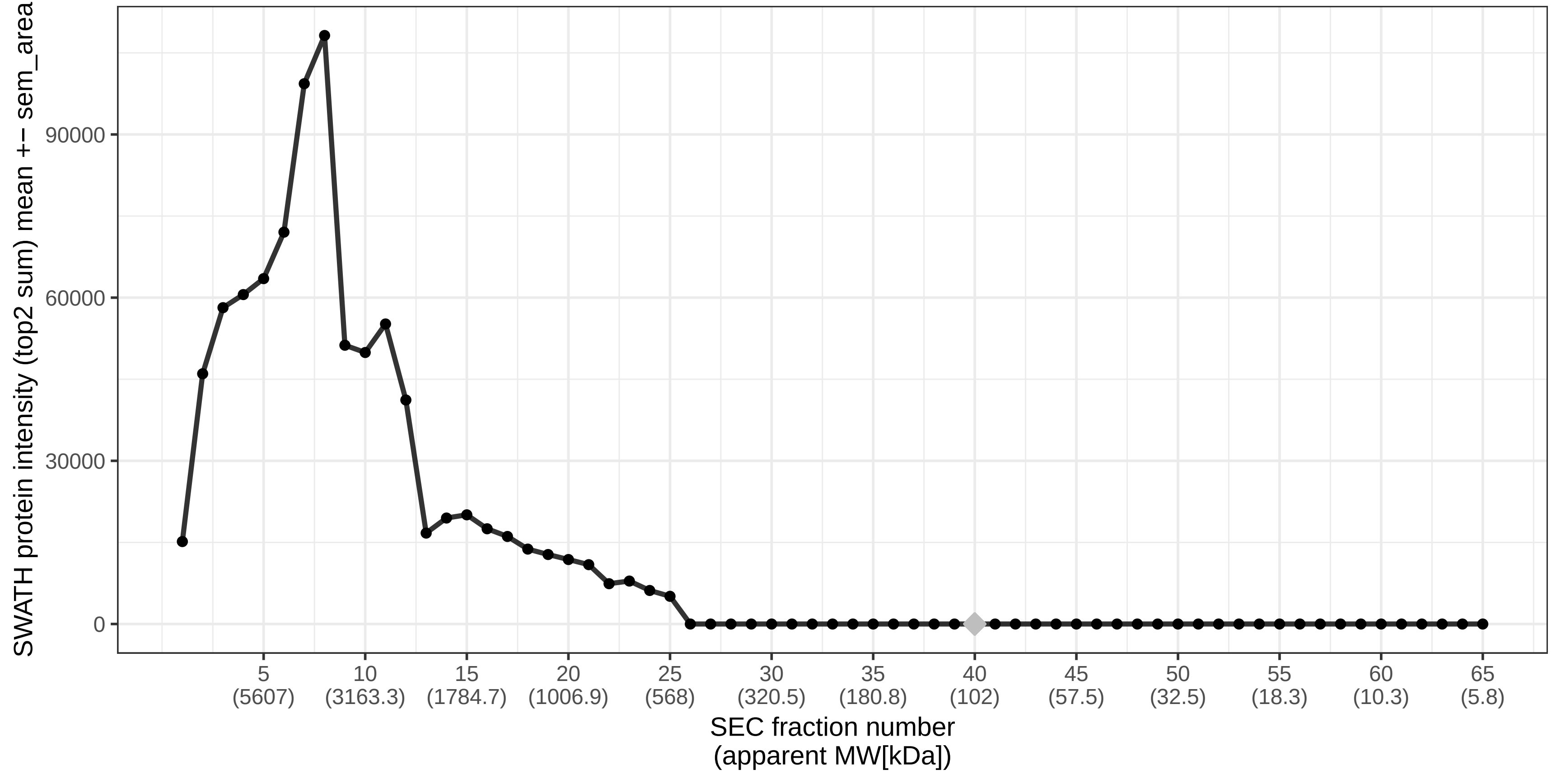

### SECchrom_O14795_UN13B_HUMAN_UNC13B_UNC13.pdf

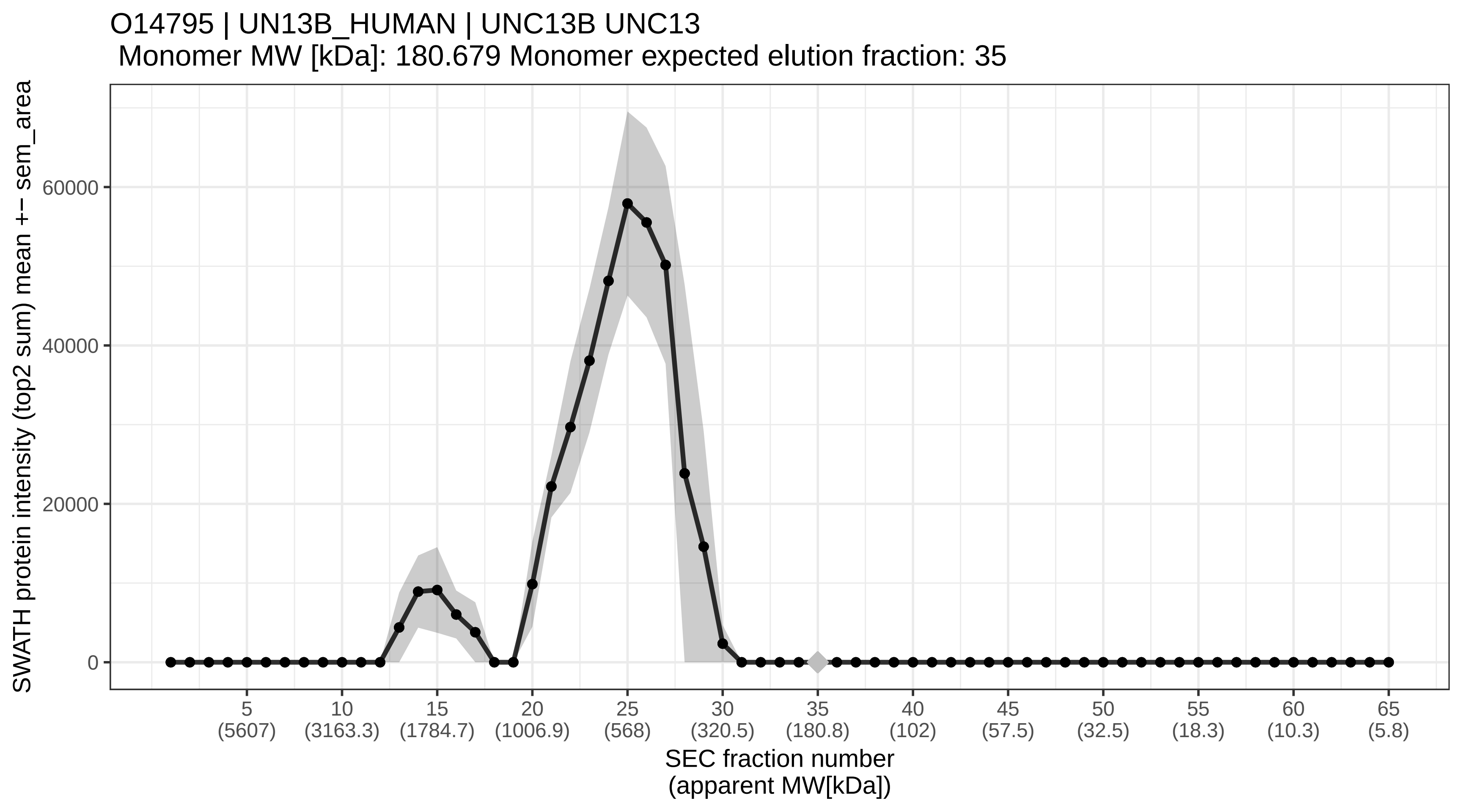

### SECchrom_O14802_RPC1_HUMAN_POLR3A.pdf

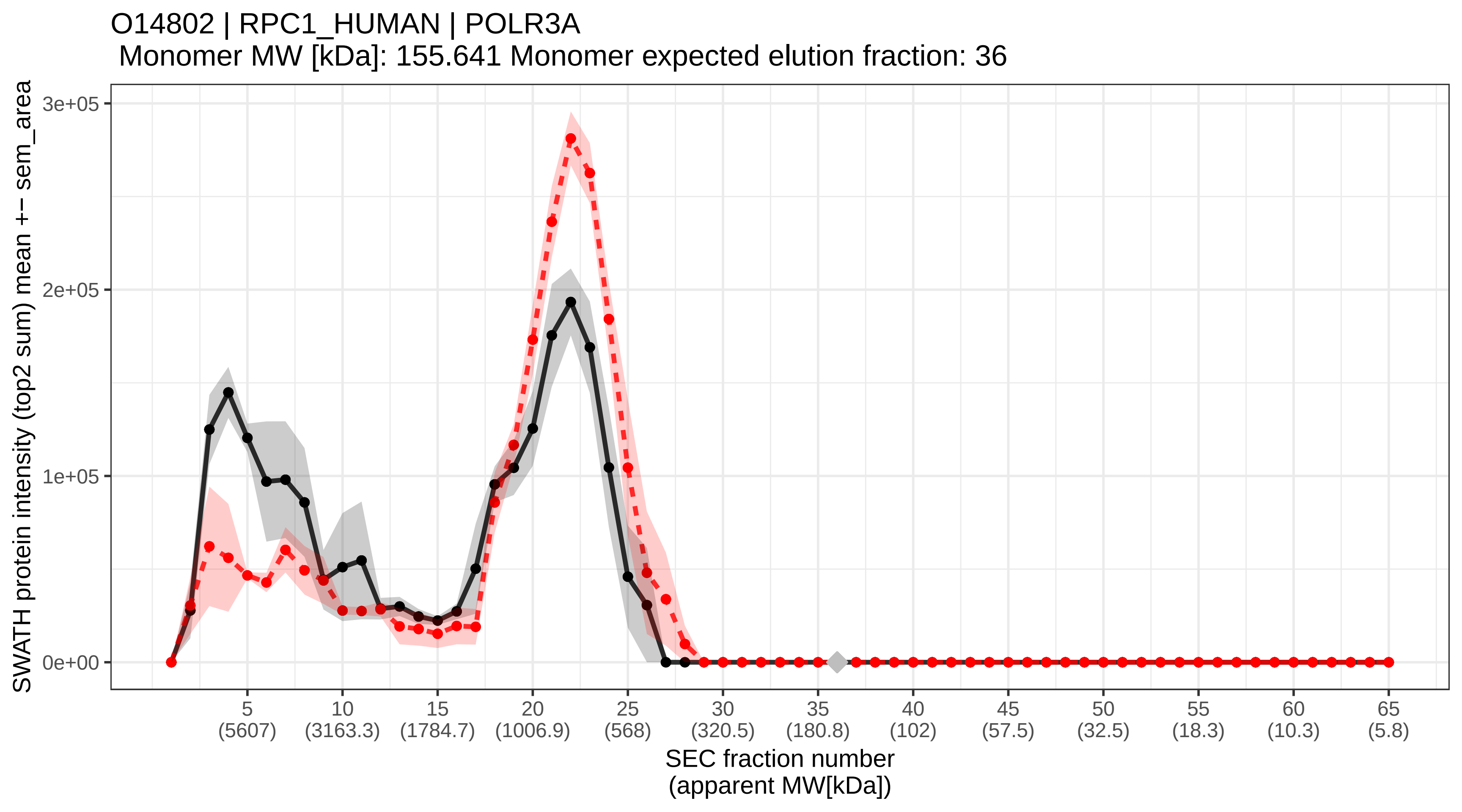

### SECchrom_O14818_PSA7_HUMAN_PSMA7_HSPC.pdf

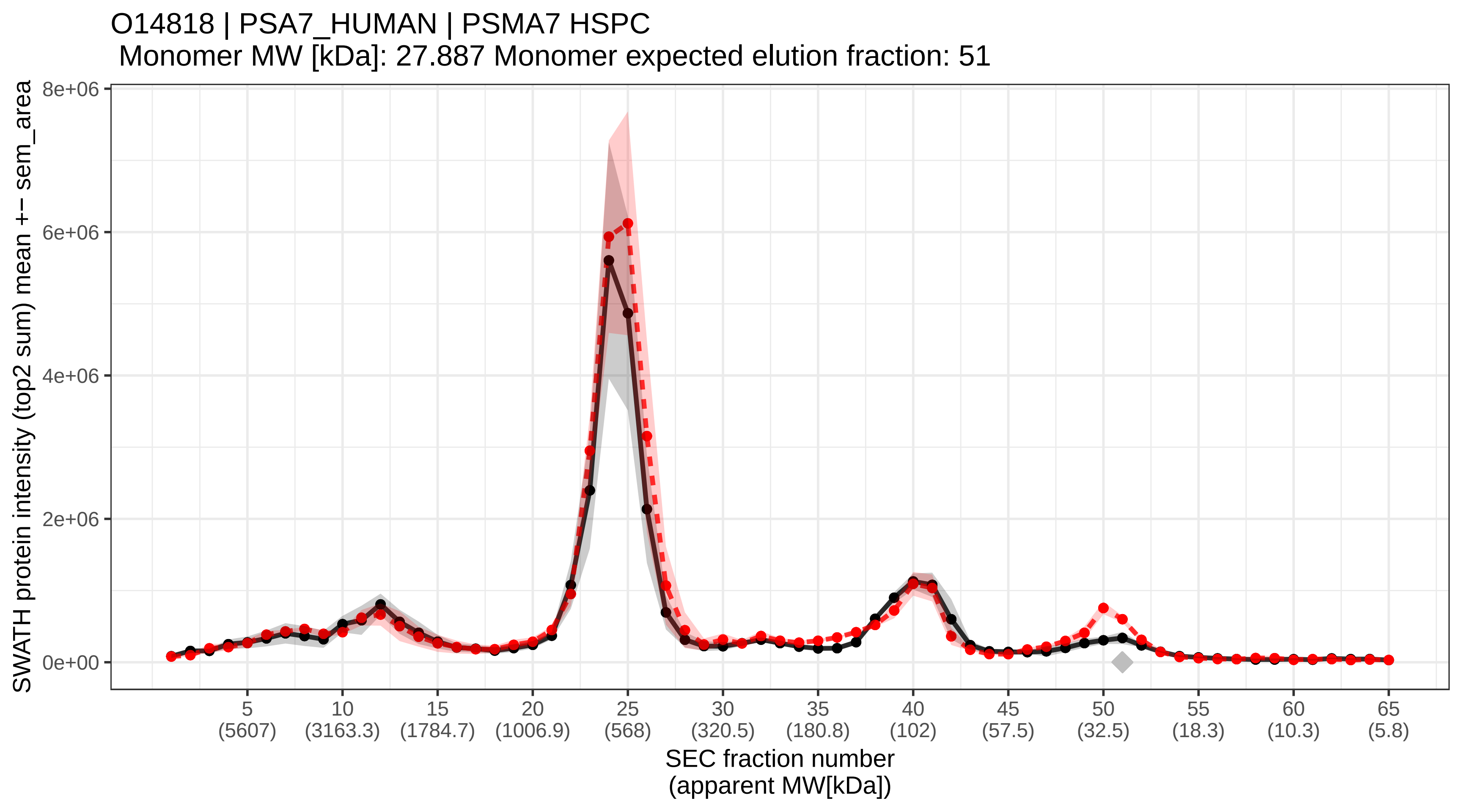

### SECchrom_O14828_SCAM3_HUMAN_SCAMP3_C1orf3_PROPIN1.pdf

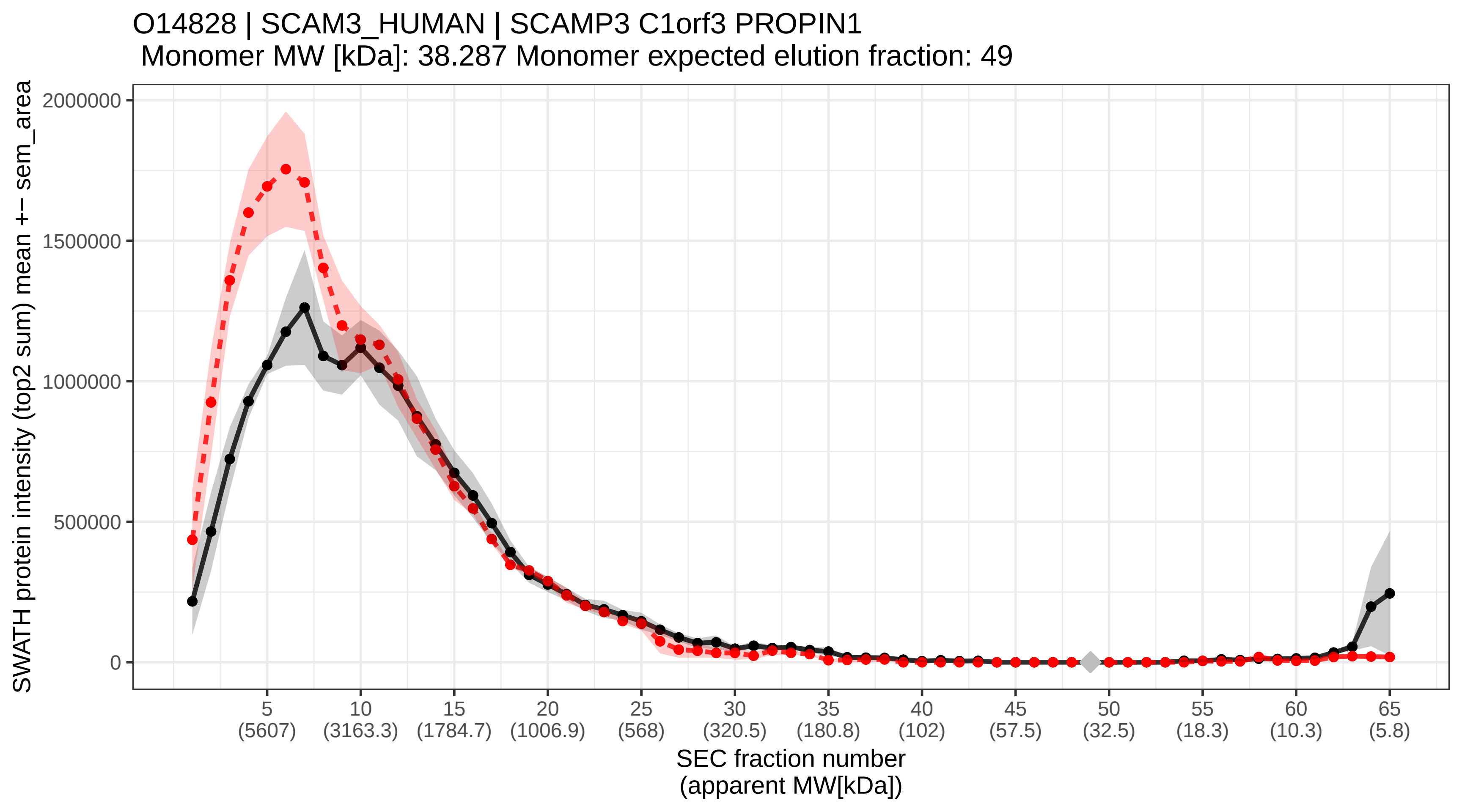

### SECchrom_O14832_PAHX_HUMAN_PHYH_PAHX.pdf

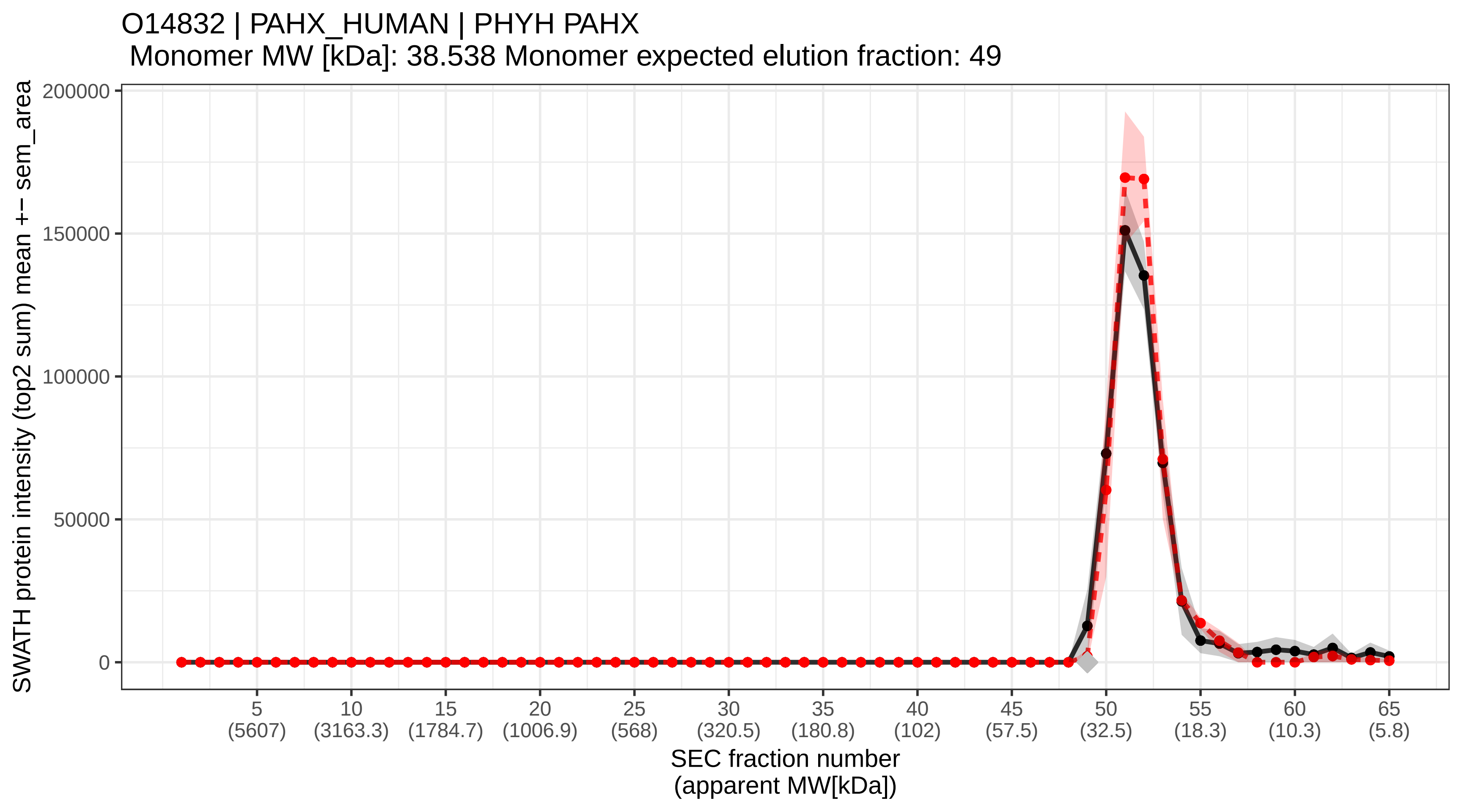

### SECchrom_O14841_OPLA_HUMAN_OPLAH.pdf

O14841 | OPLA_HUMAN | OPLAH
Monomer MW [kDa]: 137.457 Monomer expected elution fraction: 37

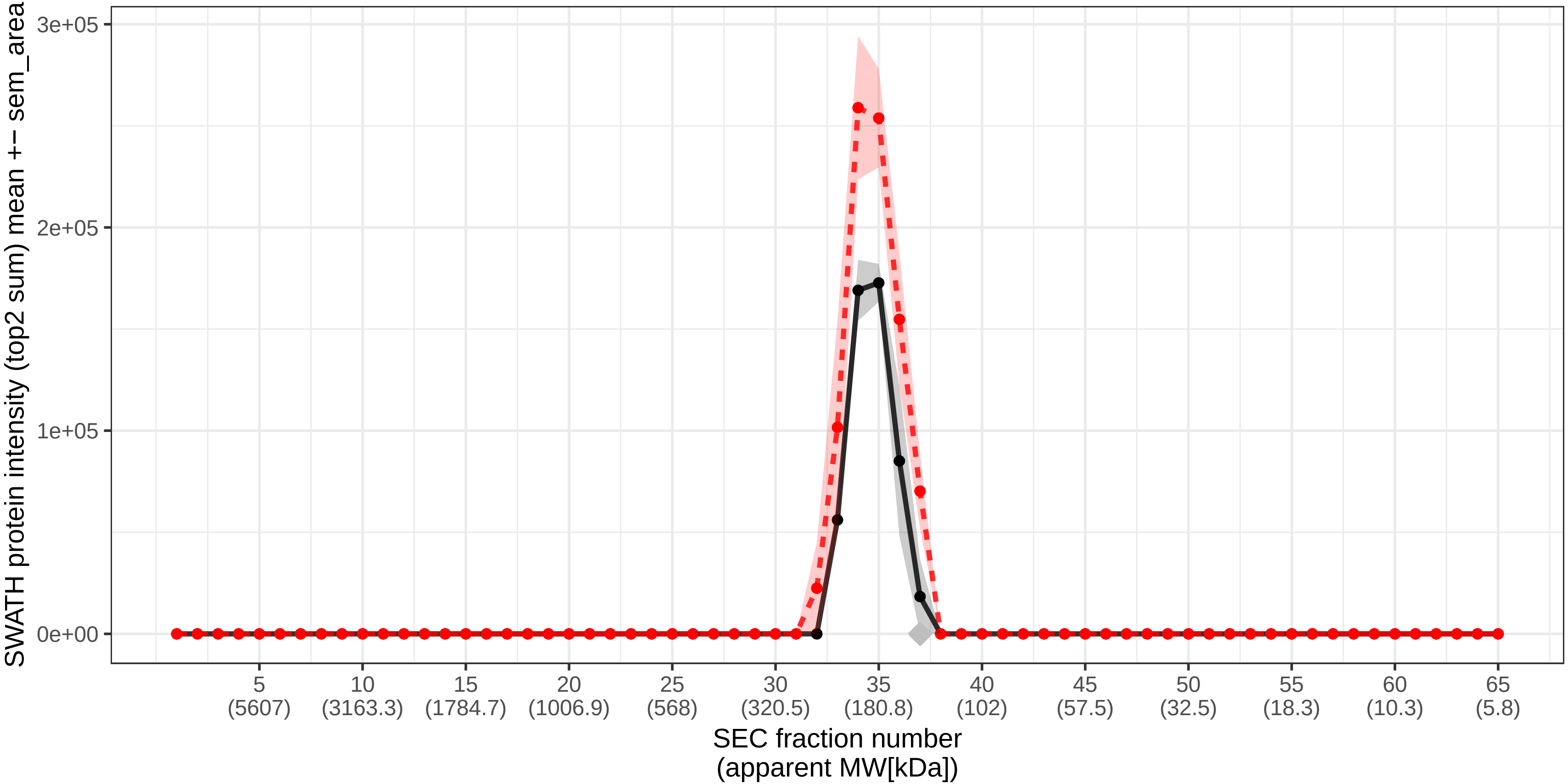

### SECchrom_O14879_IFIT3_HUMAN_IFIT3_CIG-49_IFI60_IFIT4_ISG60.pdf

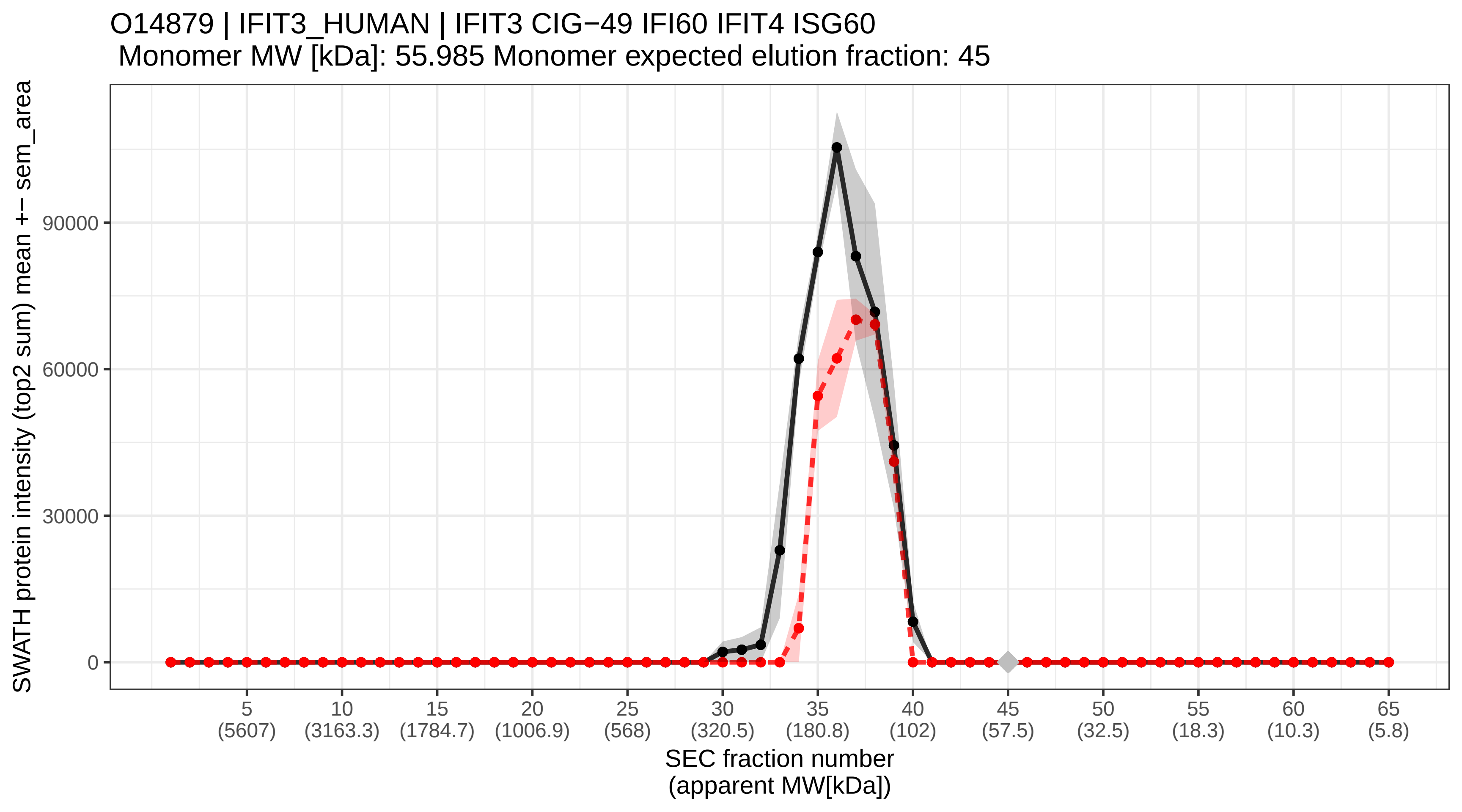

### SECchrom_O14880_MGST3_HUMAN_MGST3.pdf

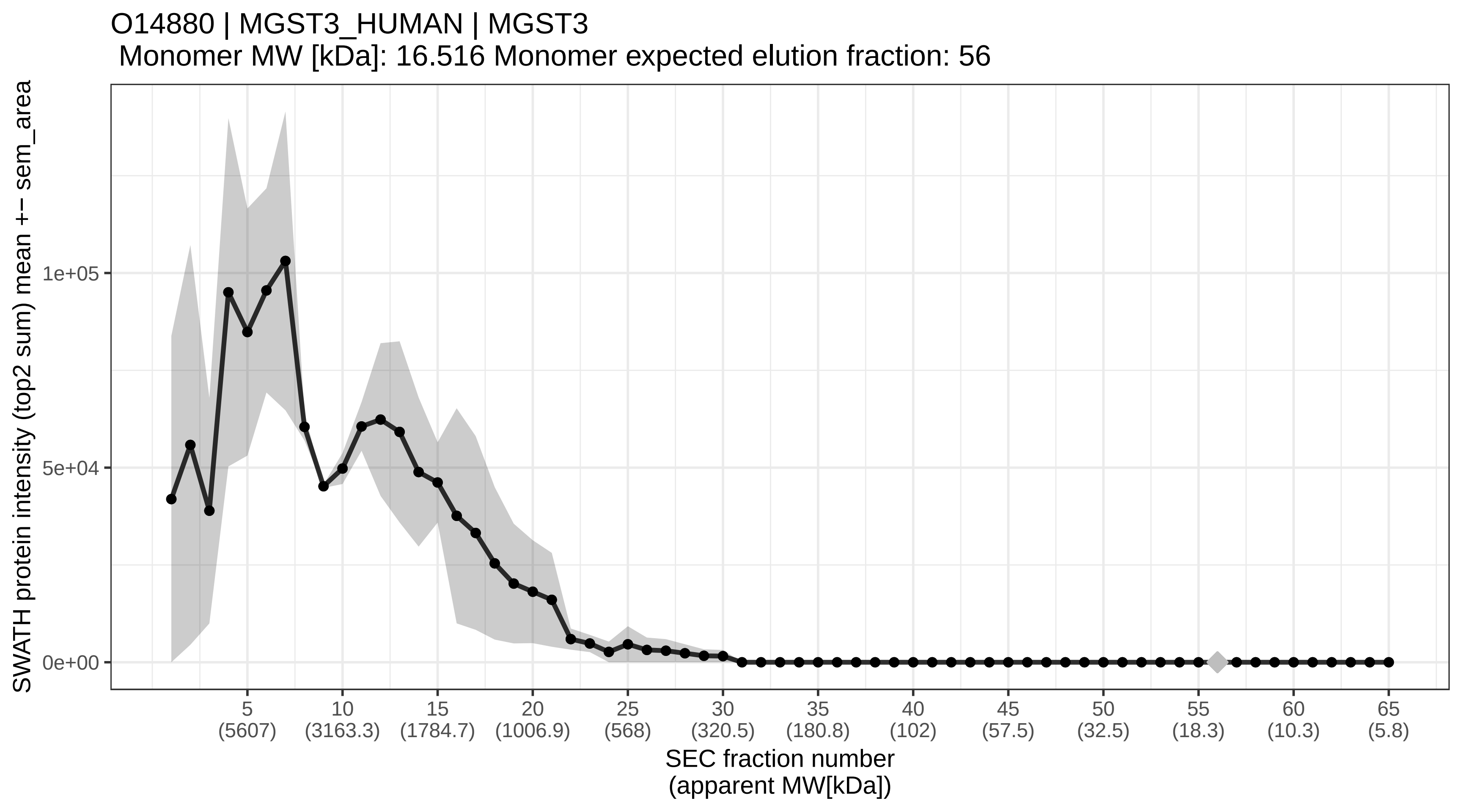

### SECchrom_O14893_GEMI2_HUMAN_GEMIN2_SIP1.pdf

O14893 | GEMI2_HUMAN | GEMIN2 SIP1
Monomer MW [kDa]: 31.585 Monomer expected elution fraction: 50

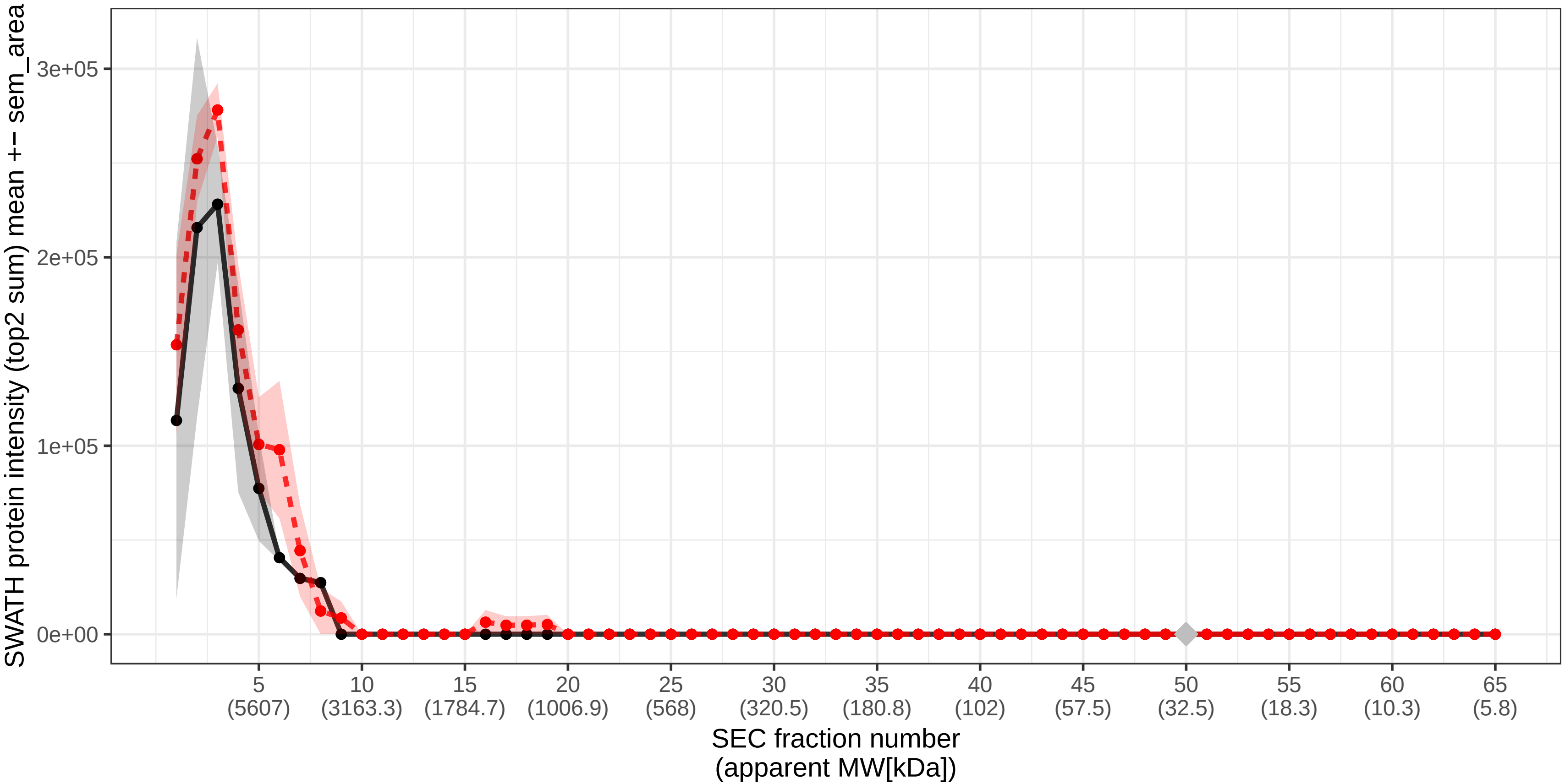

### SECchrom_O14907_TX1B3_HUMAN_TAX1BP3_TIP1.pdf

O14907 | TX1B3_HUMAN | TAX1BP3 TIP1
Monomer MW [kDa]: 13.735 Monomer expected elution fraction: 58

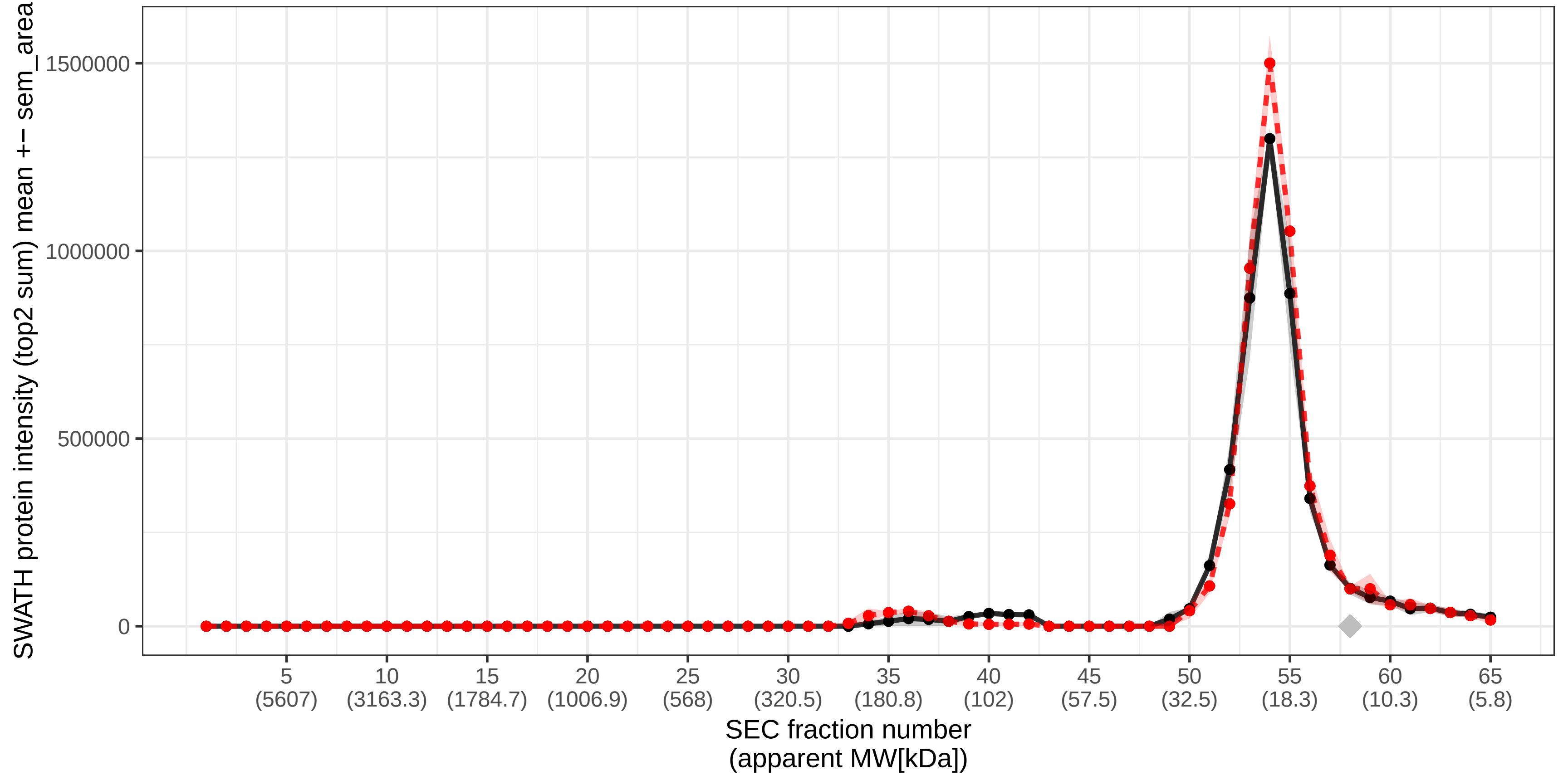

### SECchrom_O14908_GIPC1_HUMAN_GIPC1_C19orf3_GIPC_RGS19IP1.pdf

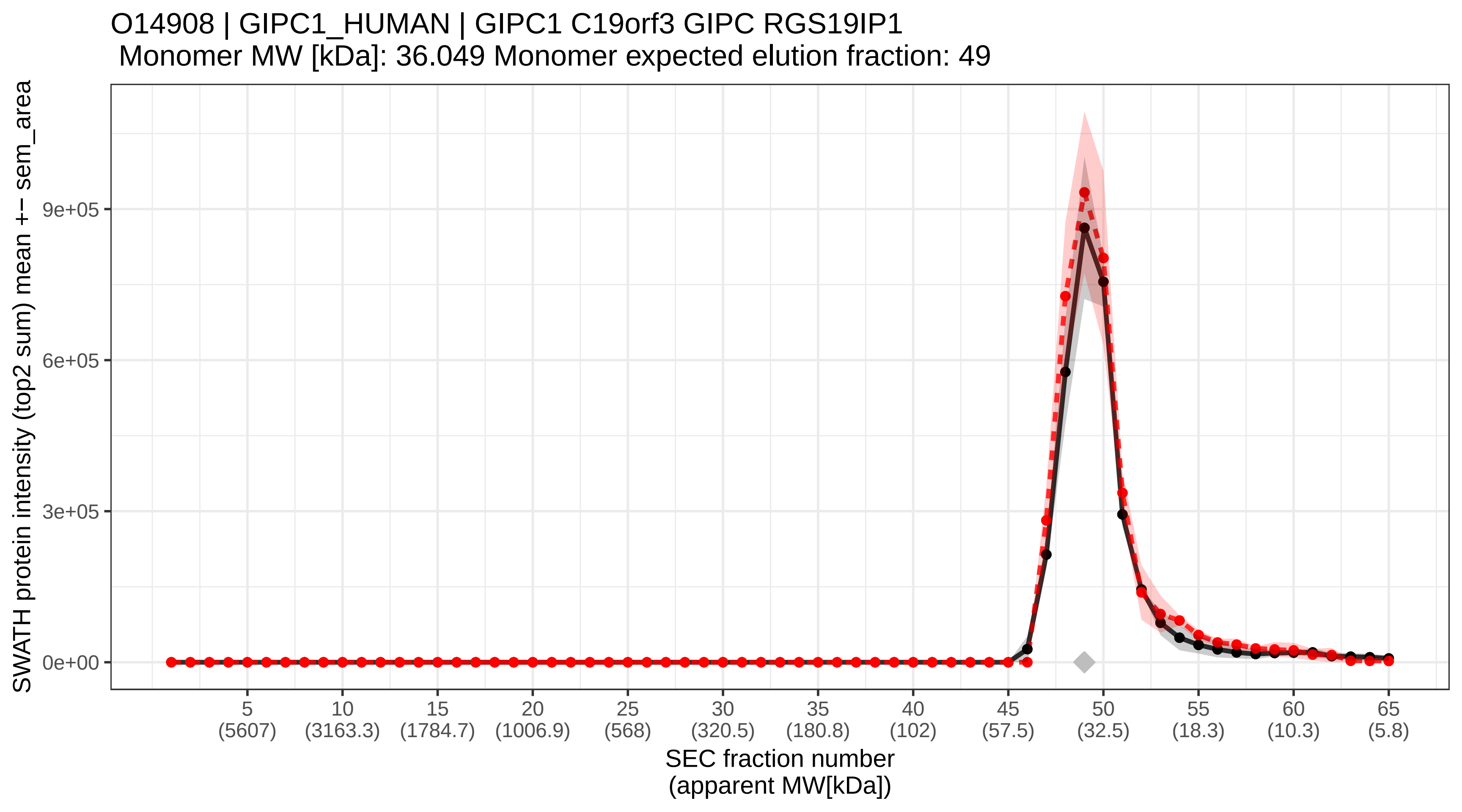

### SECchrom_O14920_IKKB_HUMAN_IKBKB_IKKB.pdf

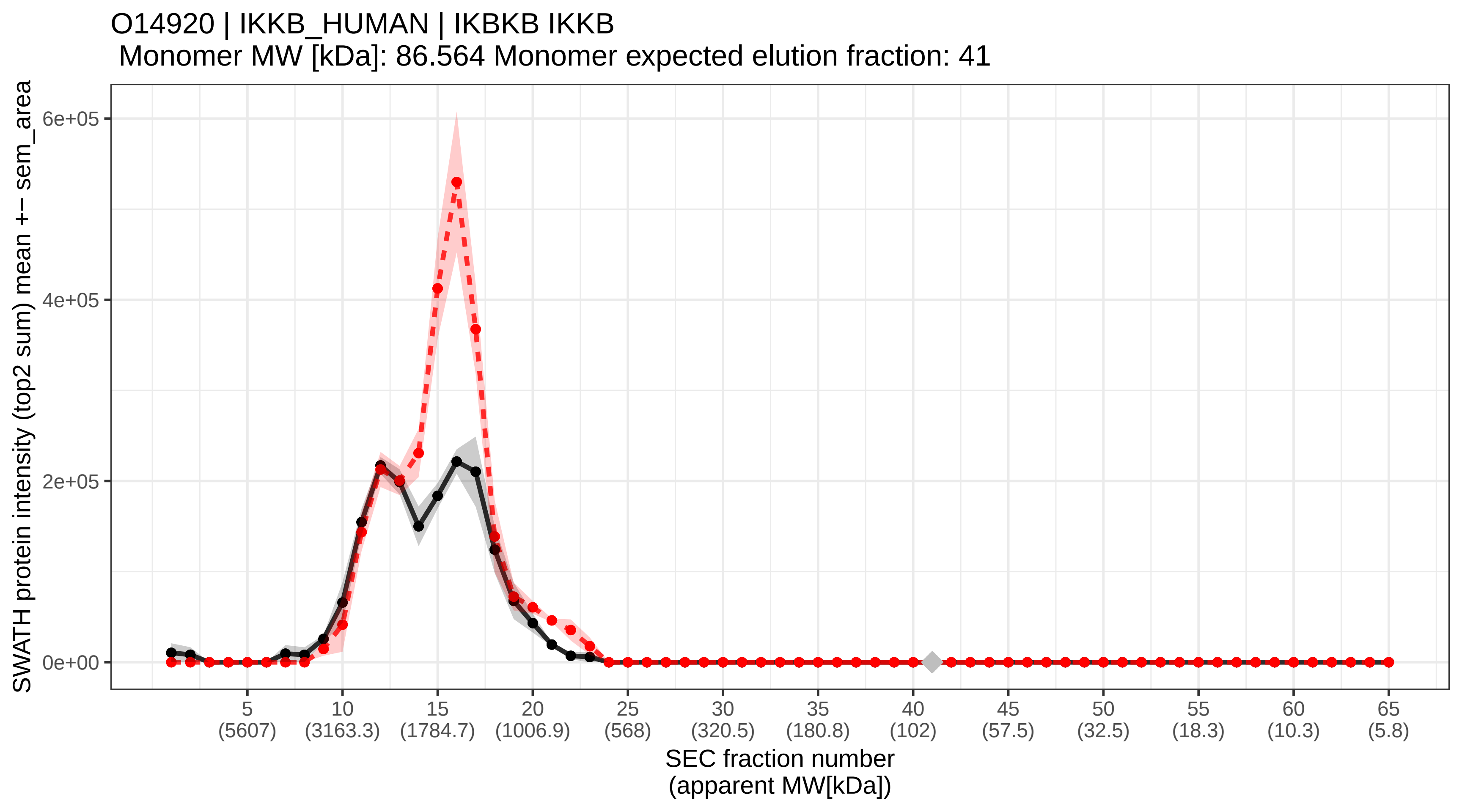

### SECchrom_O14929_HAT1_HUMAN_HAT1_KAT1.pdf

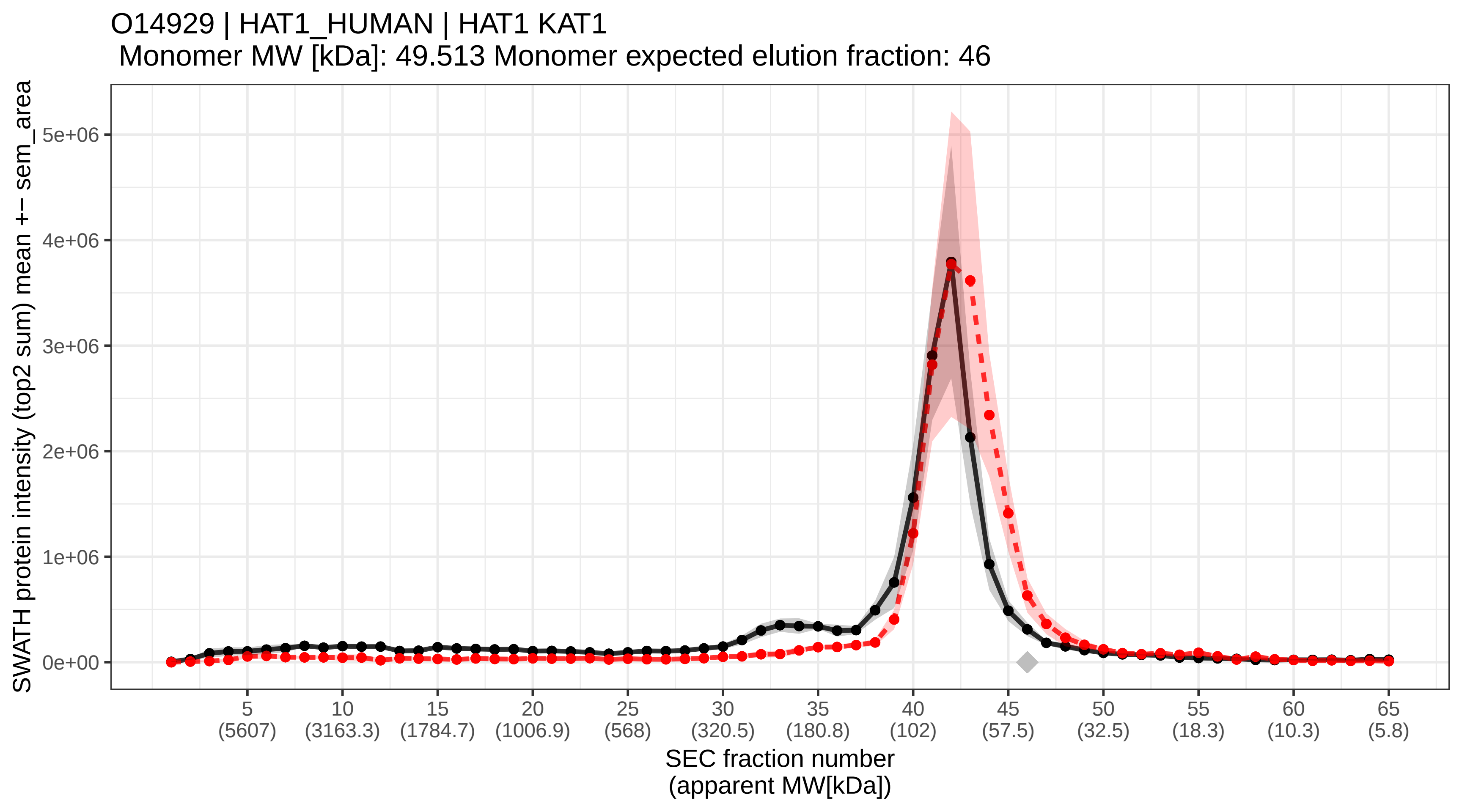

### SECchrom_O14936_CSKP_HUMAN_CASK_LIN2.pdf

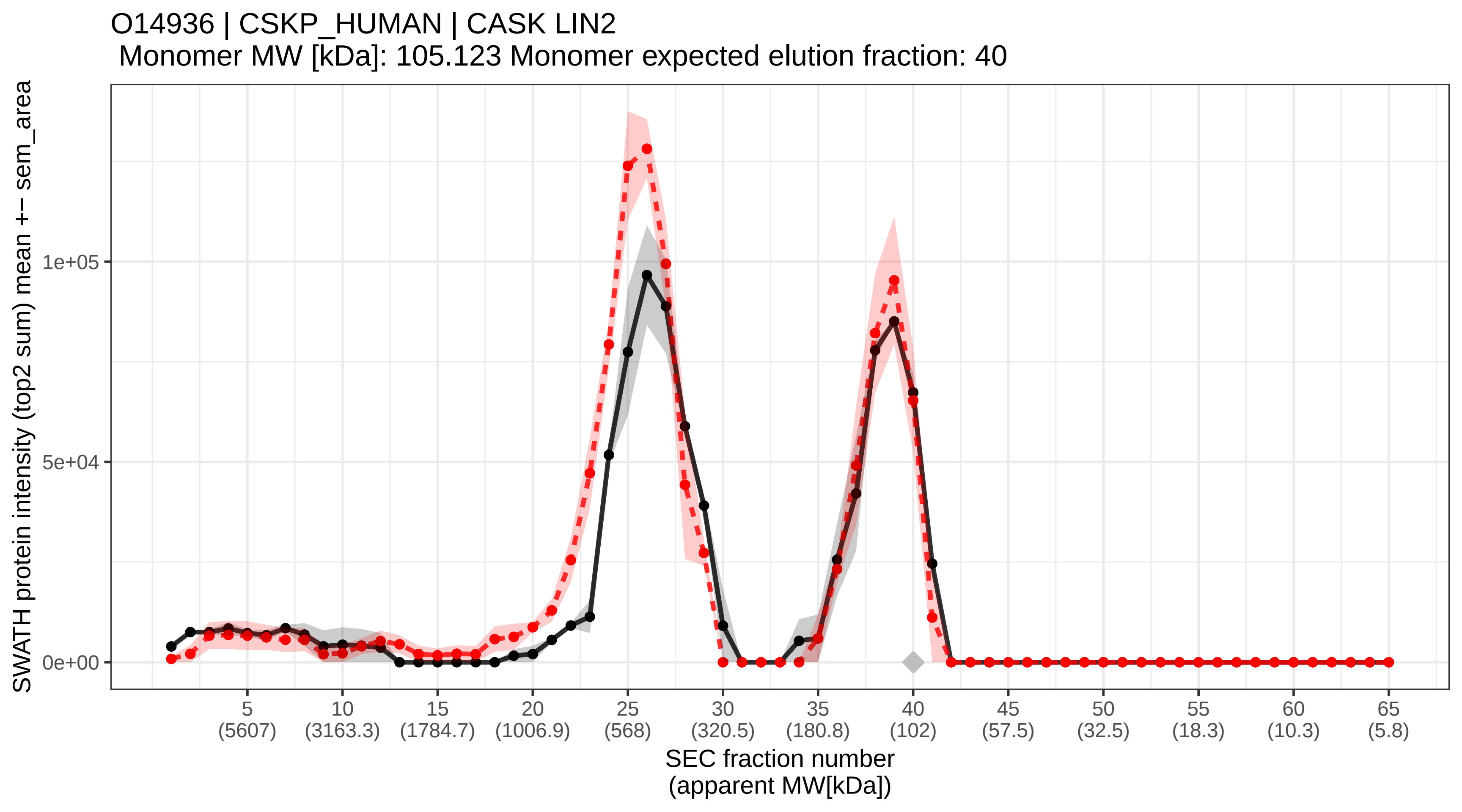

### SECchrom_O14949_QCR8_HUMAN_UQCRQ.pdf

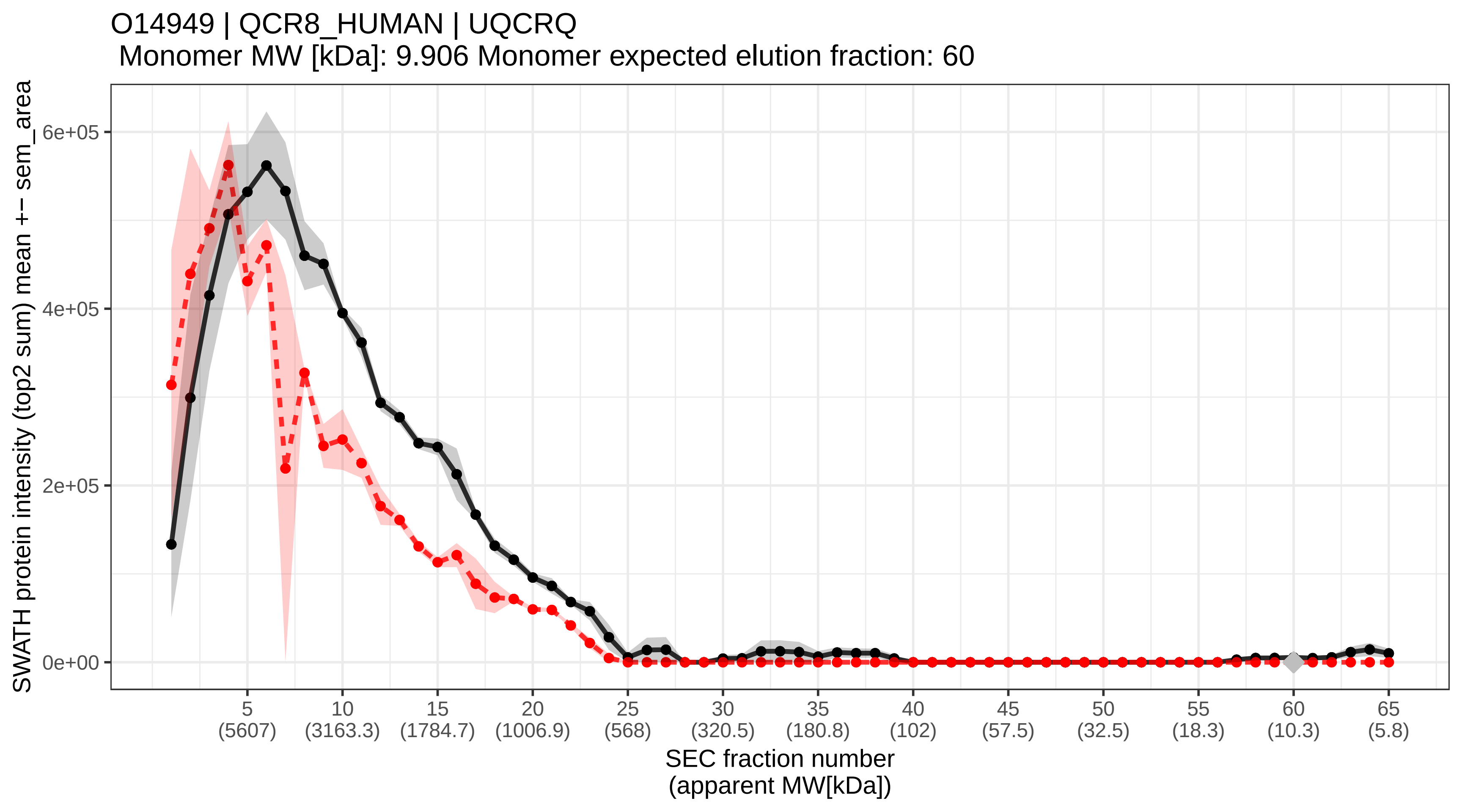

### SECchrom_O14964_HGS_HUMAN_HGS_HRS.pdf

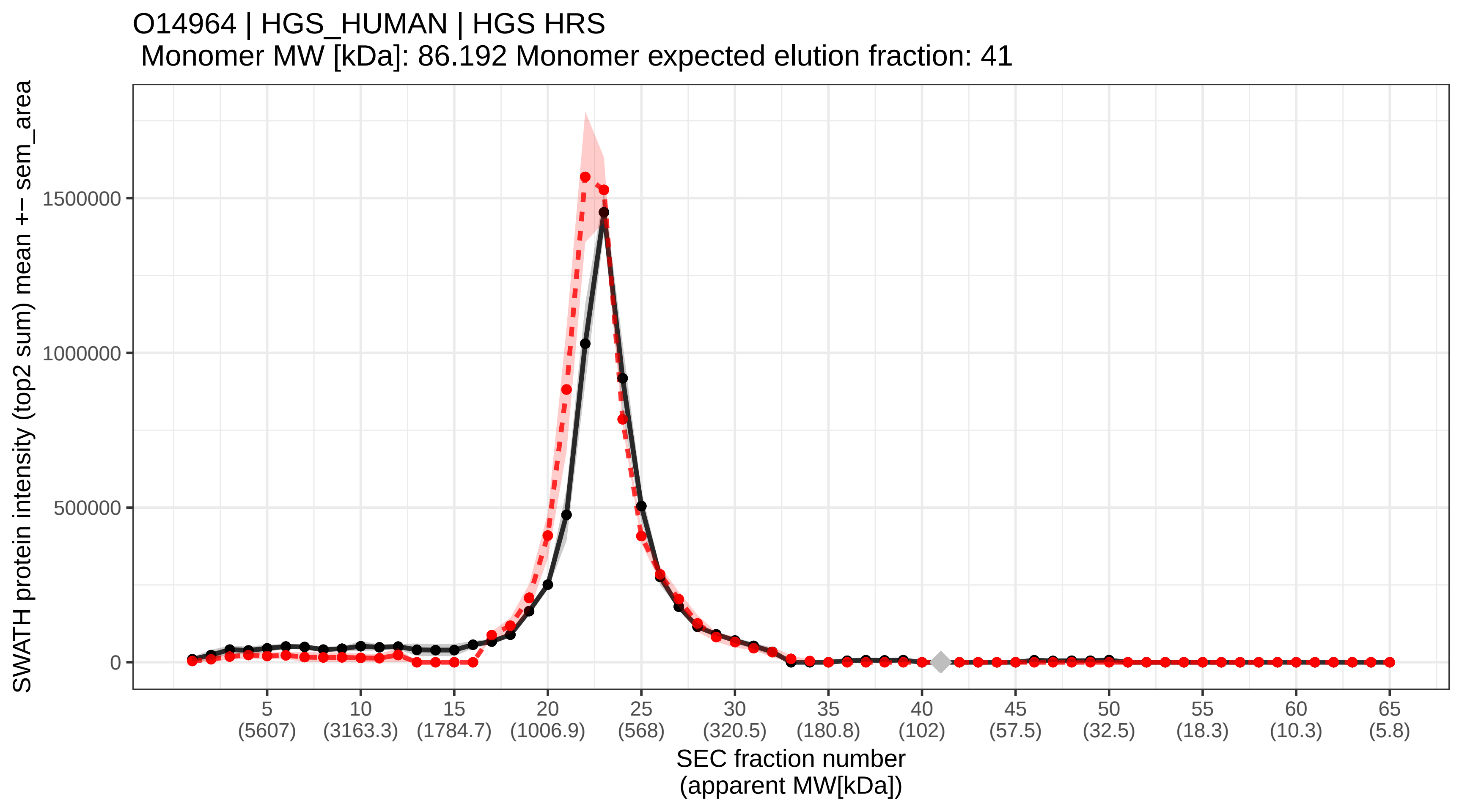

### SECchrom_O14967_CLGN_HUMAN_CLGN.pdf

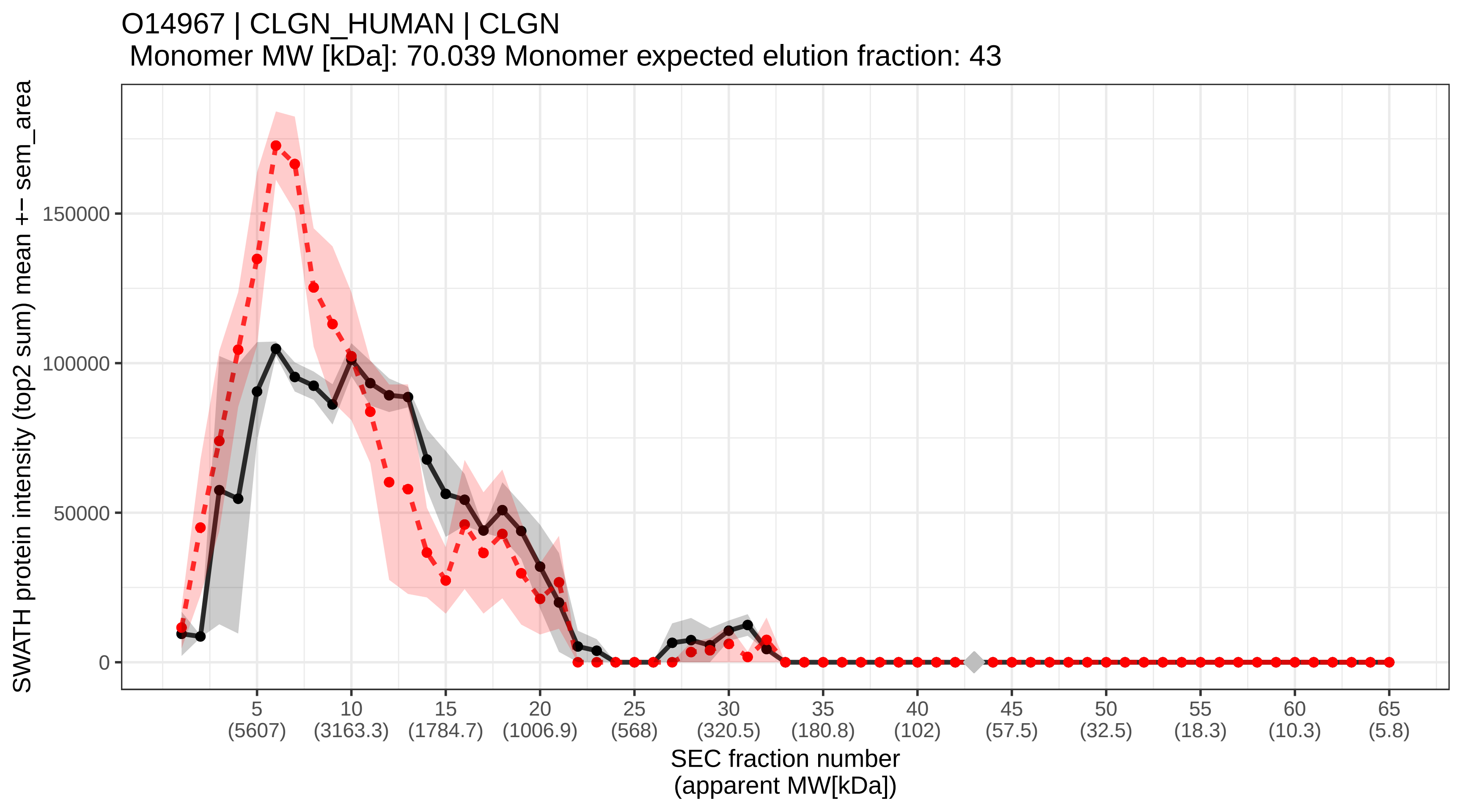

### SECchrom_O14972_DSCR3_HUMAN_DSCR3_DCRA_DSCRA.pdf

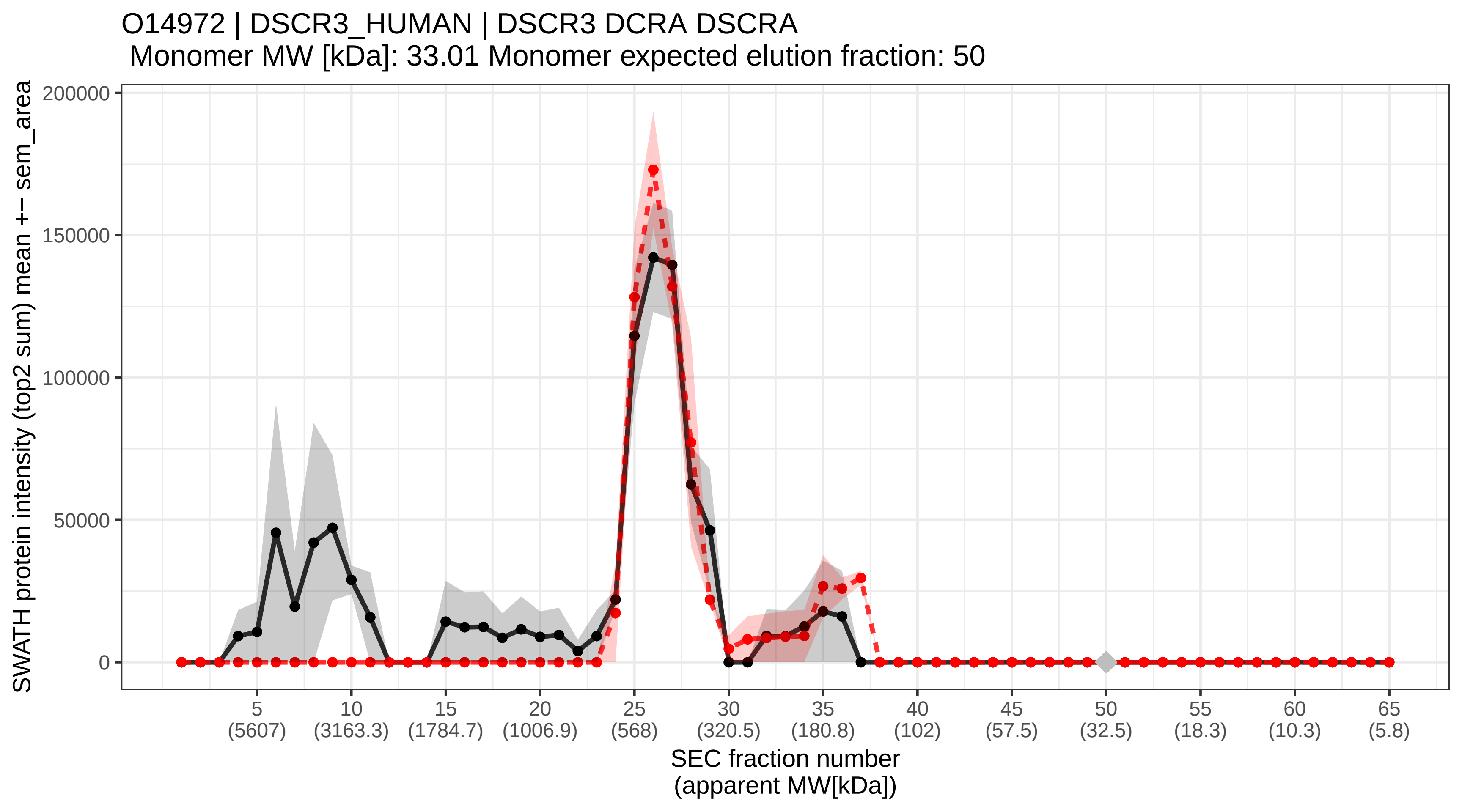

### SECchrom_O14974_MYPT1_HUMAN_PPP1R12A_MBS_MYPT1.pdf

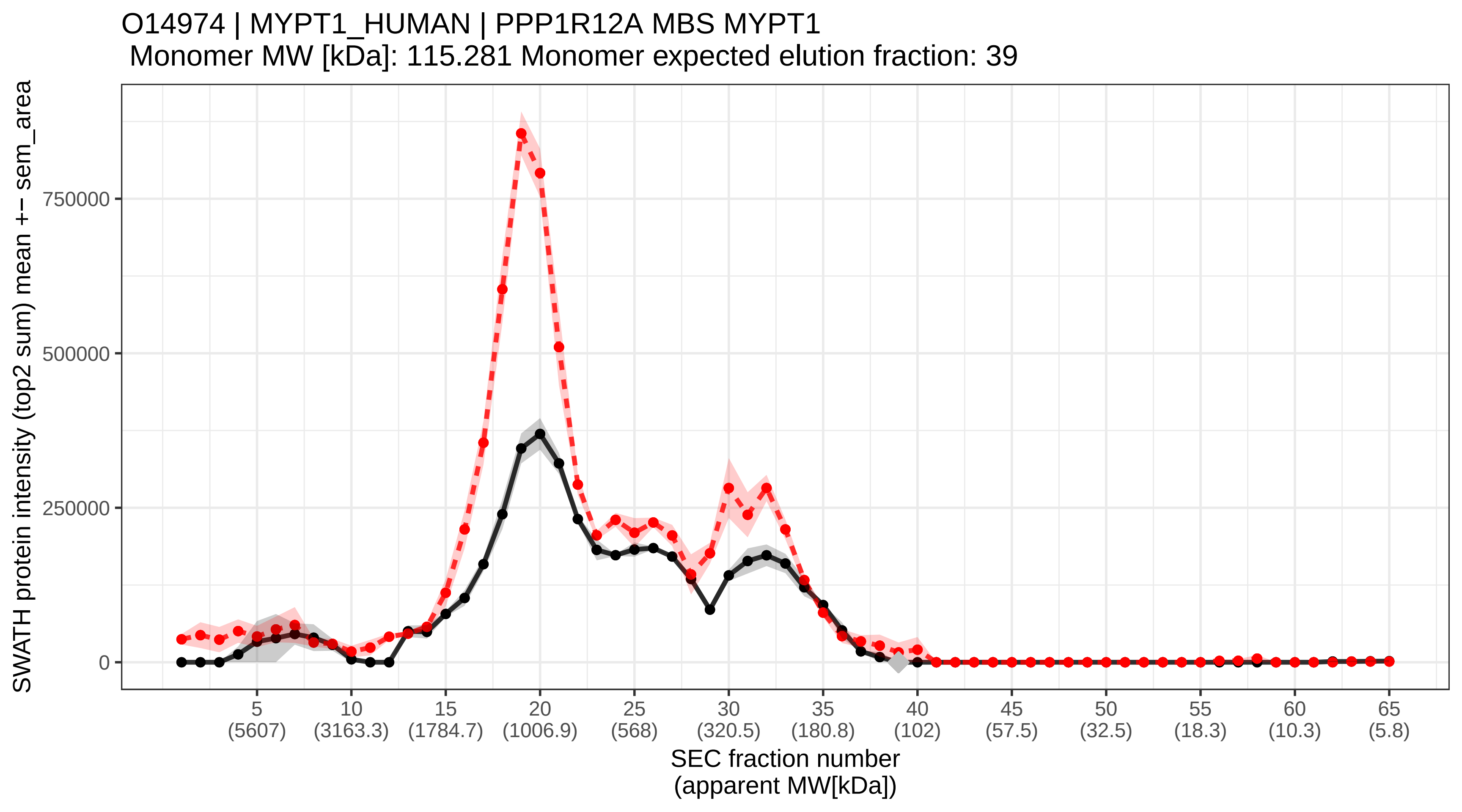

### SECchrom_O14975_S27A2_HUMAN_SLC27A2_ACSVL1_FACVL1_FATP2_VLA.pdf

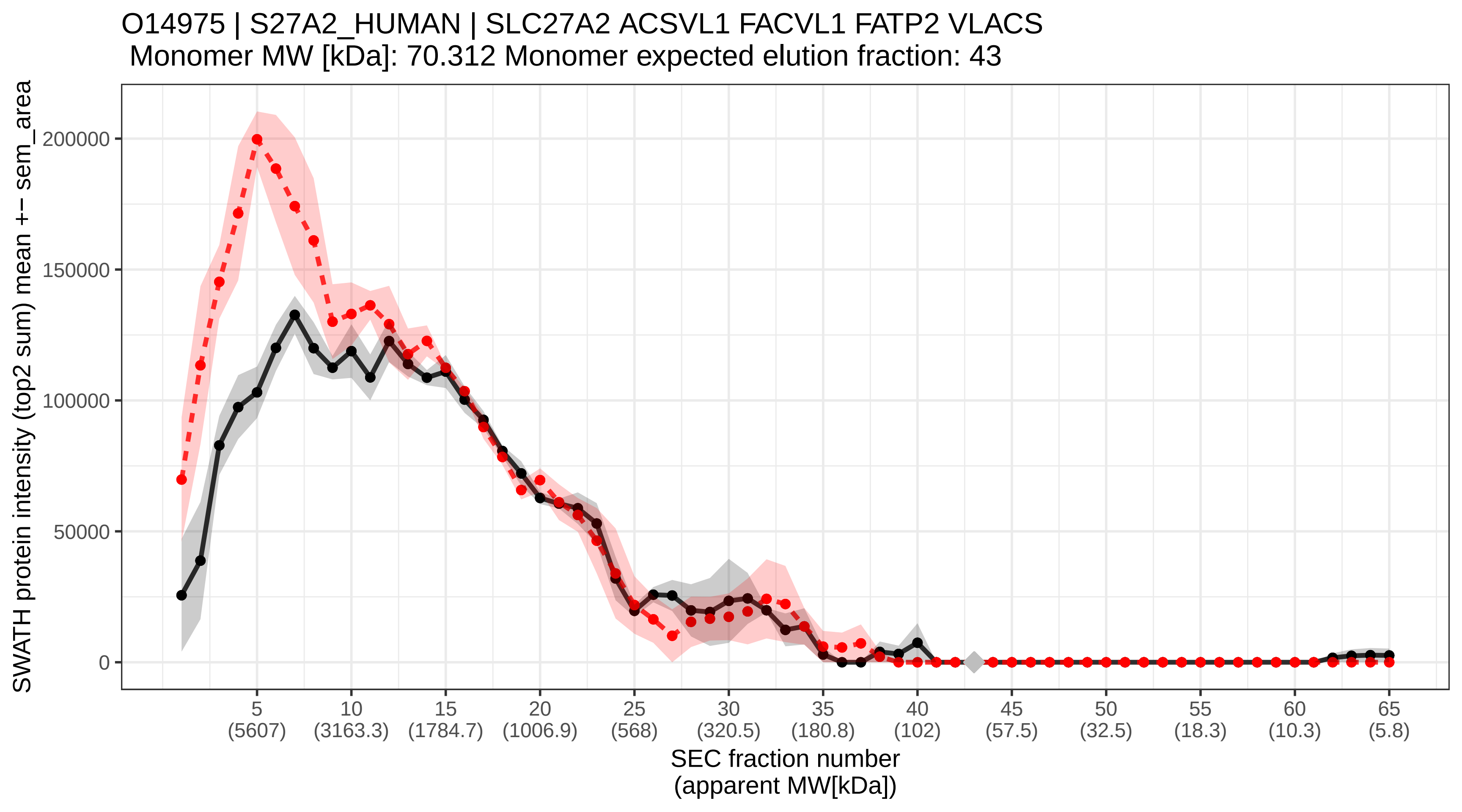

### SECchrom_O14976_GAK_HUMAN_GAK.pdf

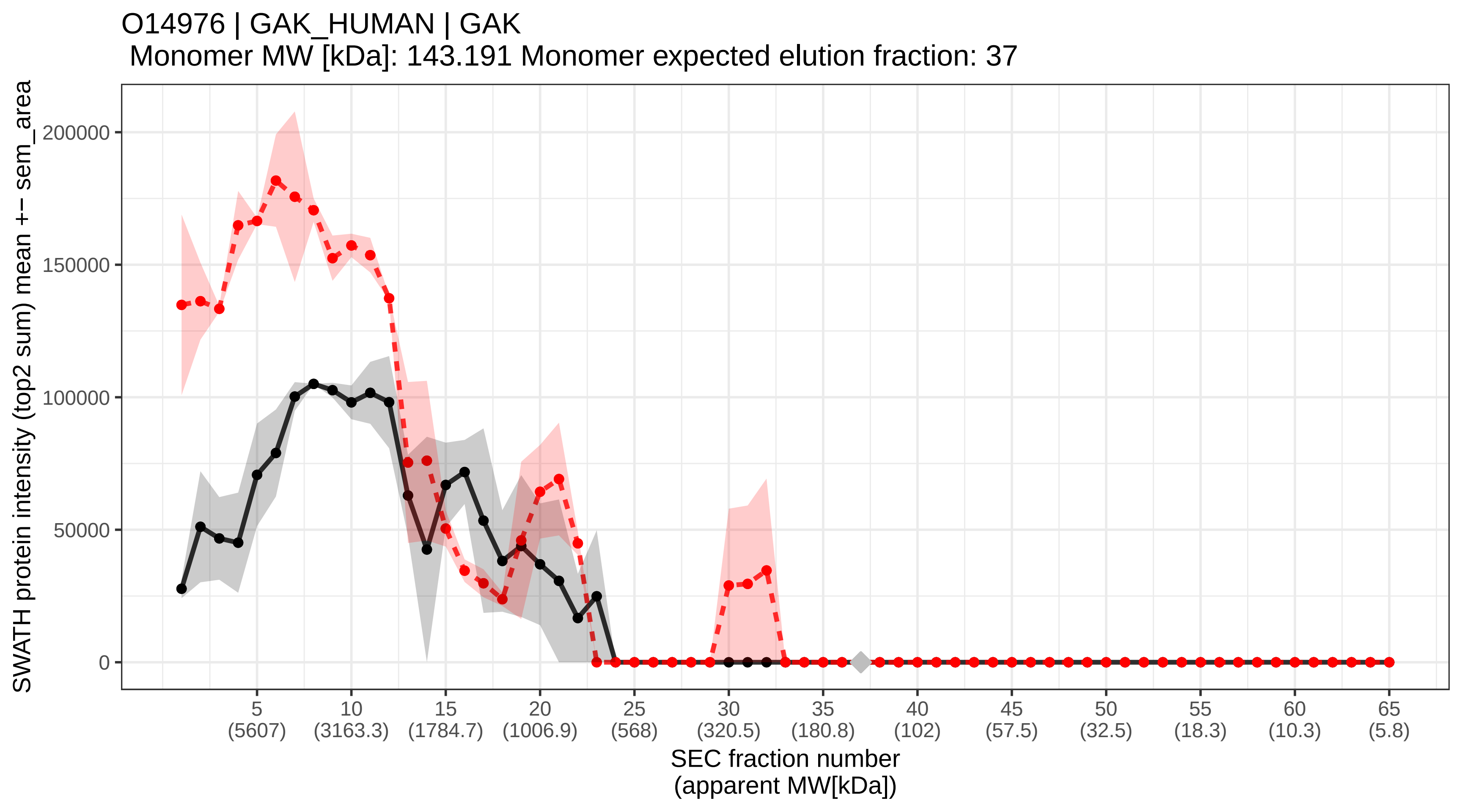

### SECchrom_O14979_HNRDL_HUMAN_HNRNPDL_HNRPDL_JKTBP.pdf

O14979 | HNRDL_HUMAN | HNRNPDL HNRPDL JKTBP
Monomer MW [kDa]: 46.438 Monomer expected elution fraction: 47

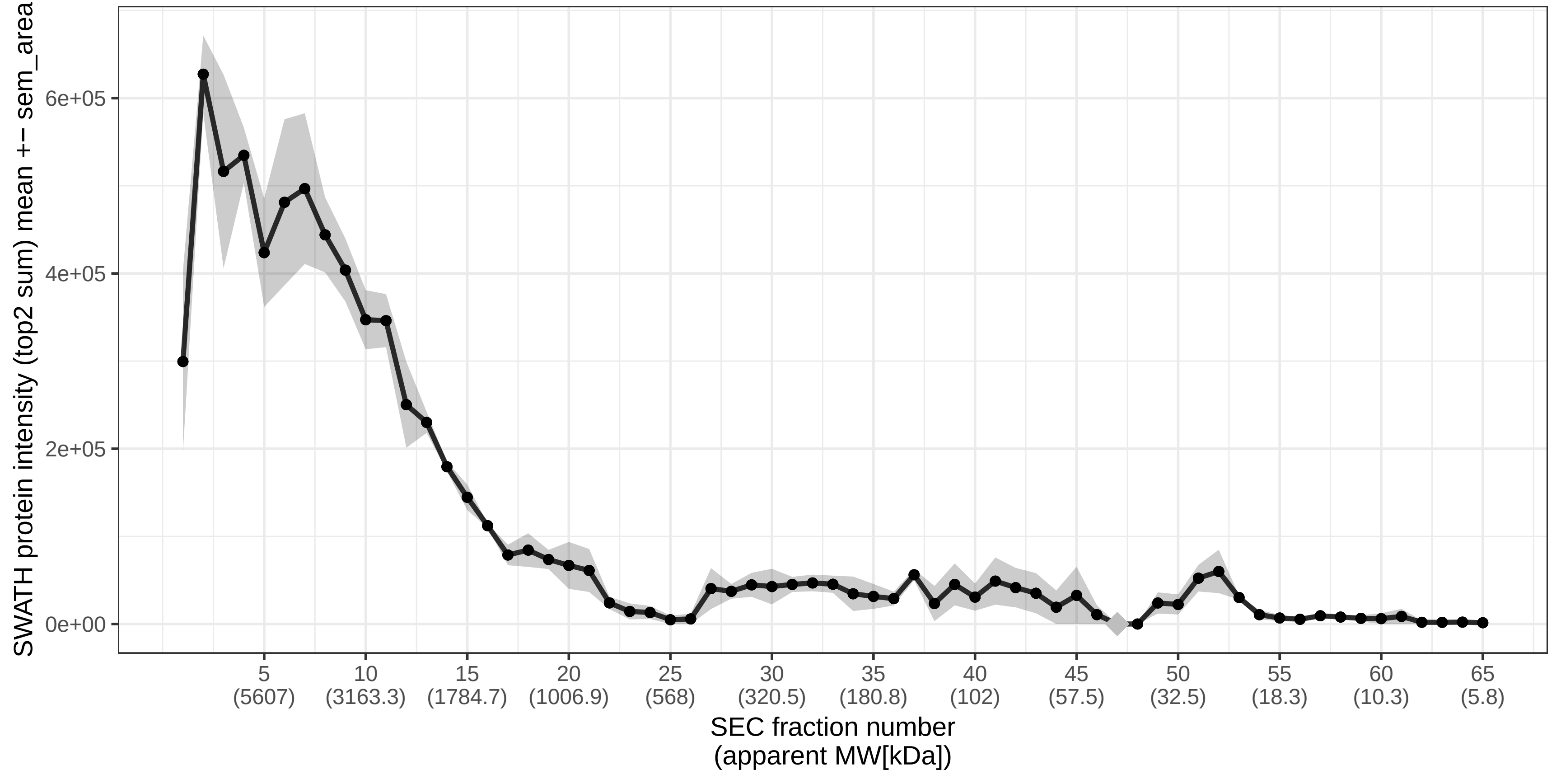

### SECchrom_O14980_XPO1_HUMAN_XPO1_CRM1.pdf

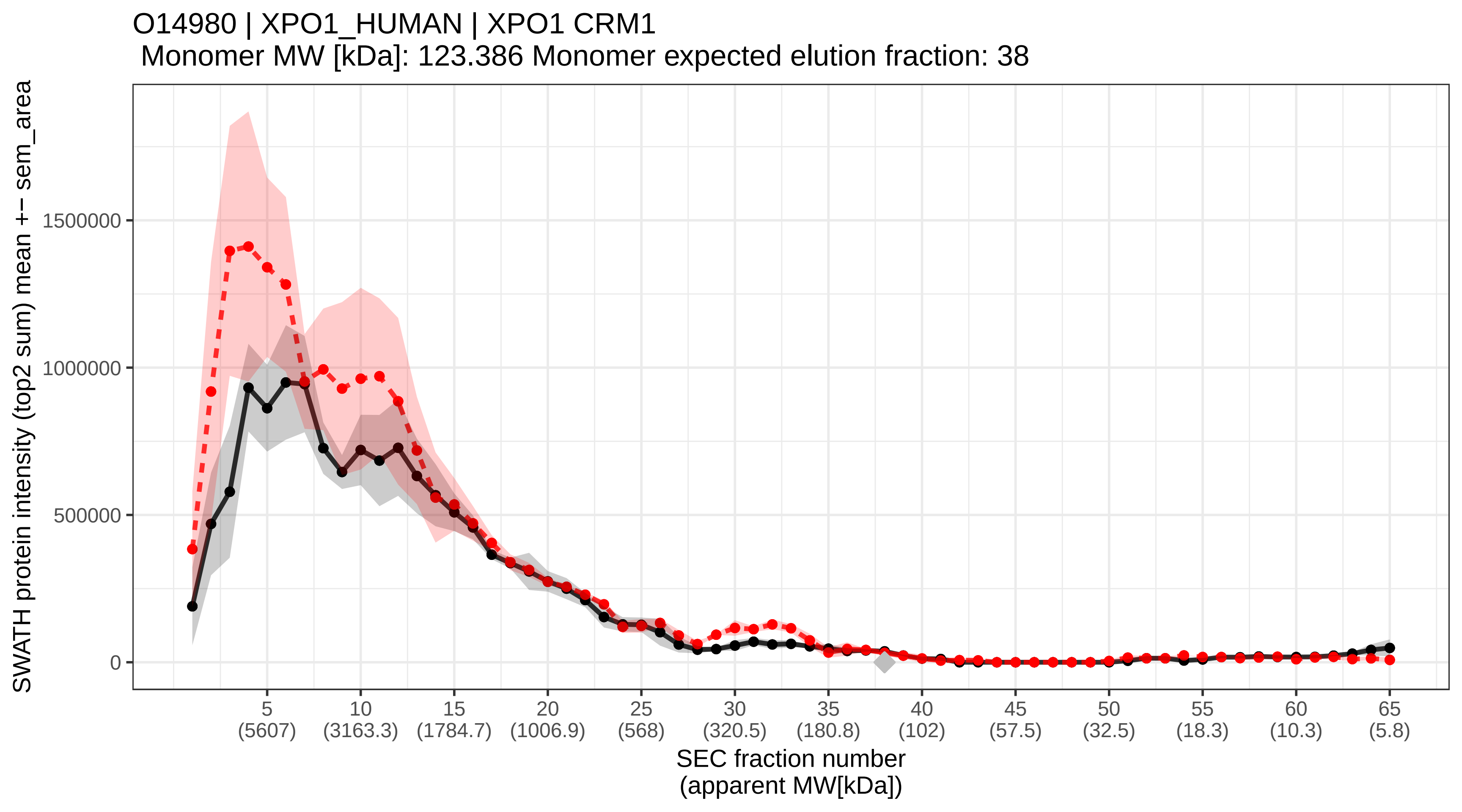

### SECchrom_O14981_BTAF1_HUMAN_BTAF1_TAF172.pdf

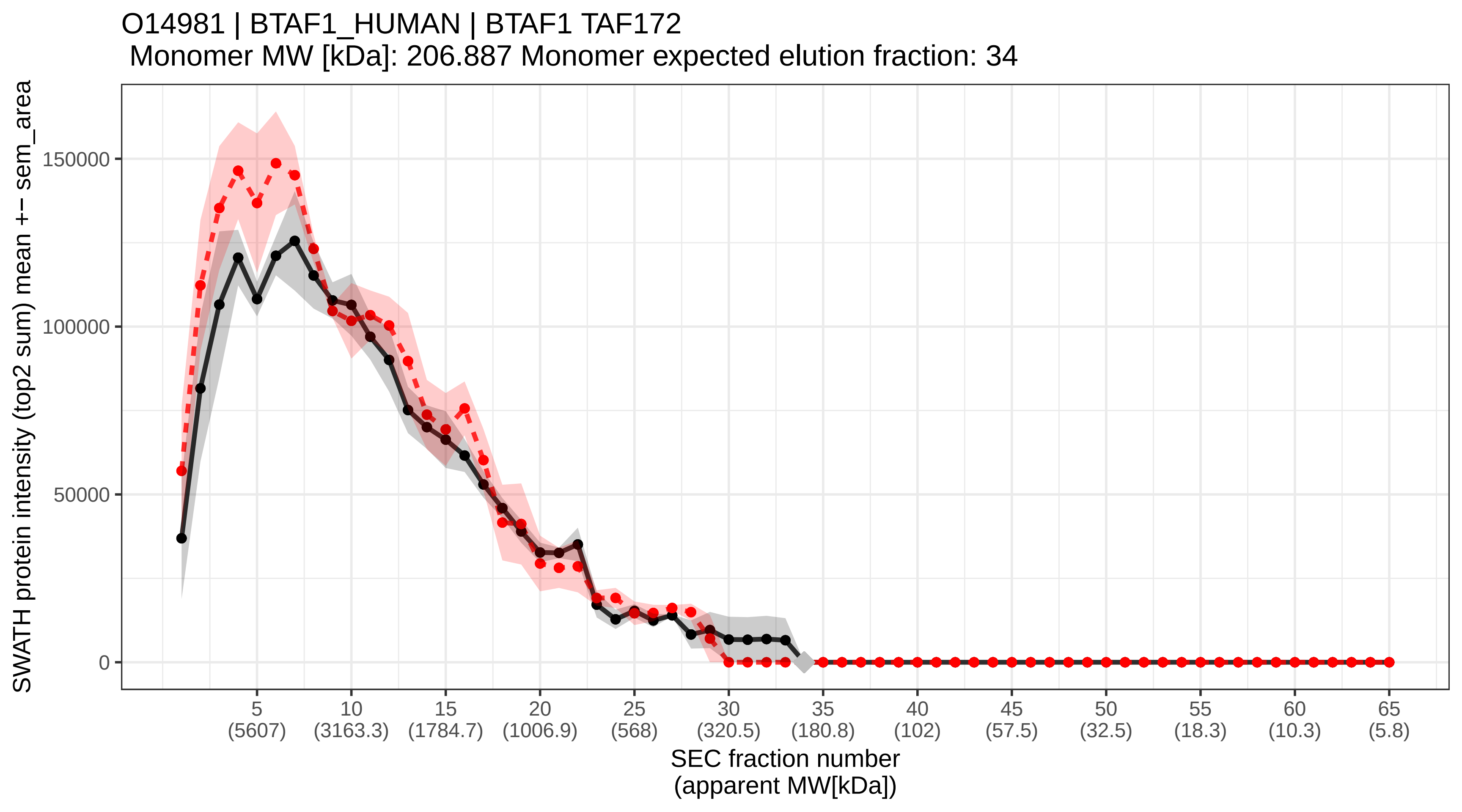

### SECchrom_O15013_ARHGA_HUMAN_ARHGEF10_KIAA0294.pdf

O15013 | ARHGA_HUMAN | ARHGEF10 KIAA0294
Monomer MW [kDa]: 151.612 Monomer expected elution fraction: 37

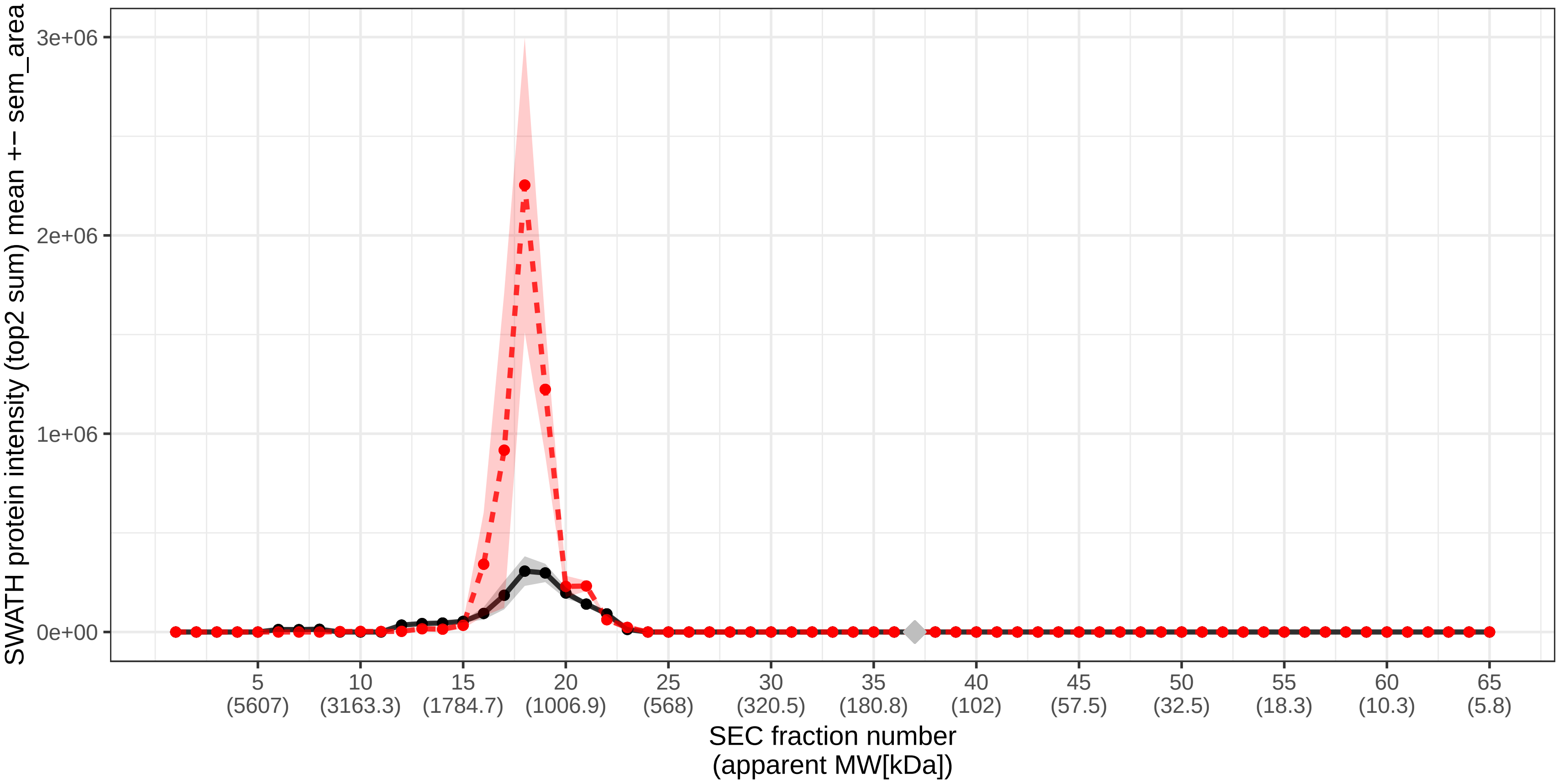

### SECchrom_O15020_SPTN2_HUMAN_SPTBN2_KIAA0302_SCA5.pdf

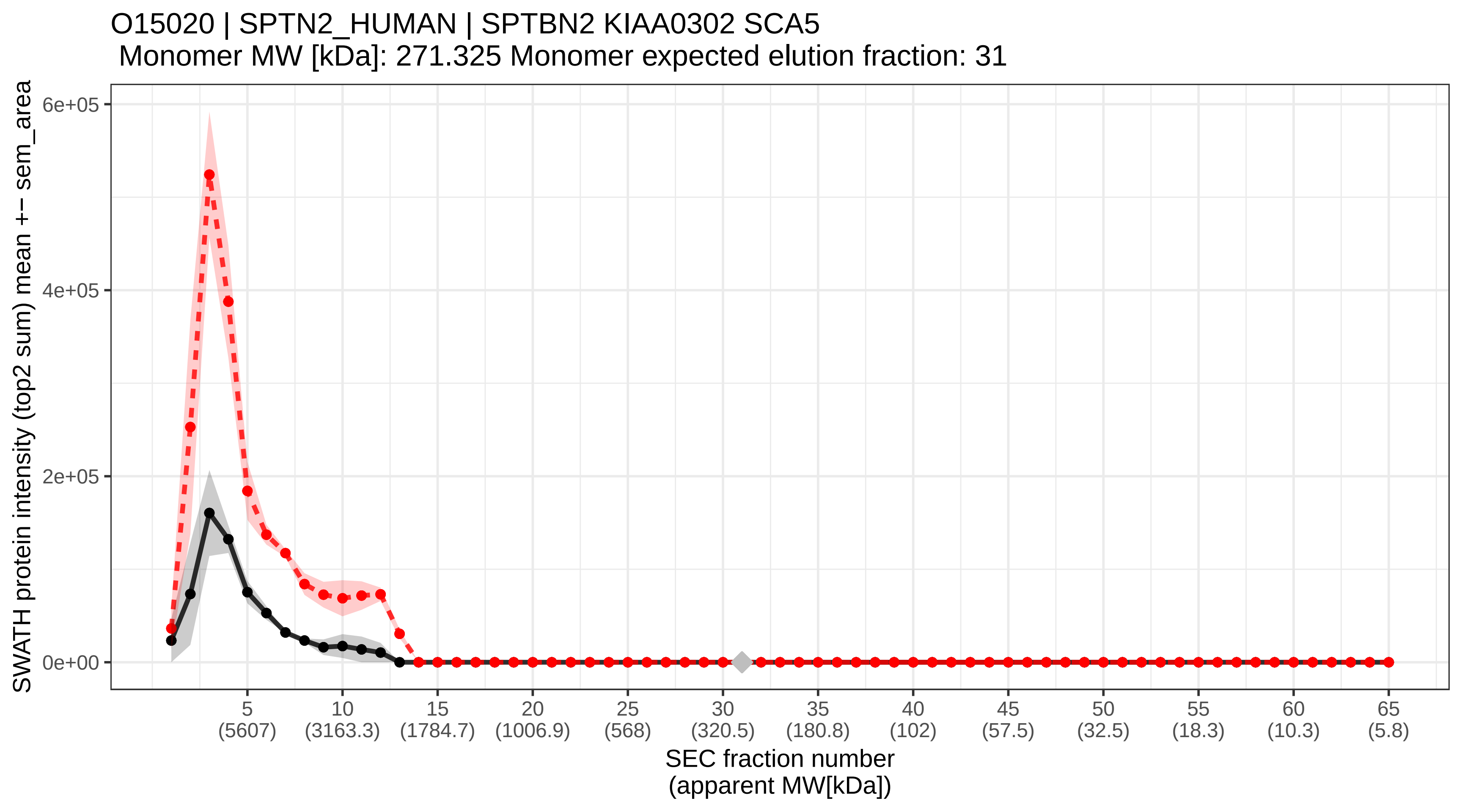

### SECchrom_O15027_SC16A_HUMAN_SEC16A_KIAA0310_SEC16_SEC16L.pdf

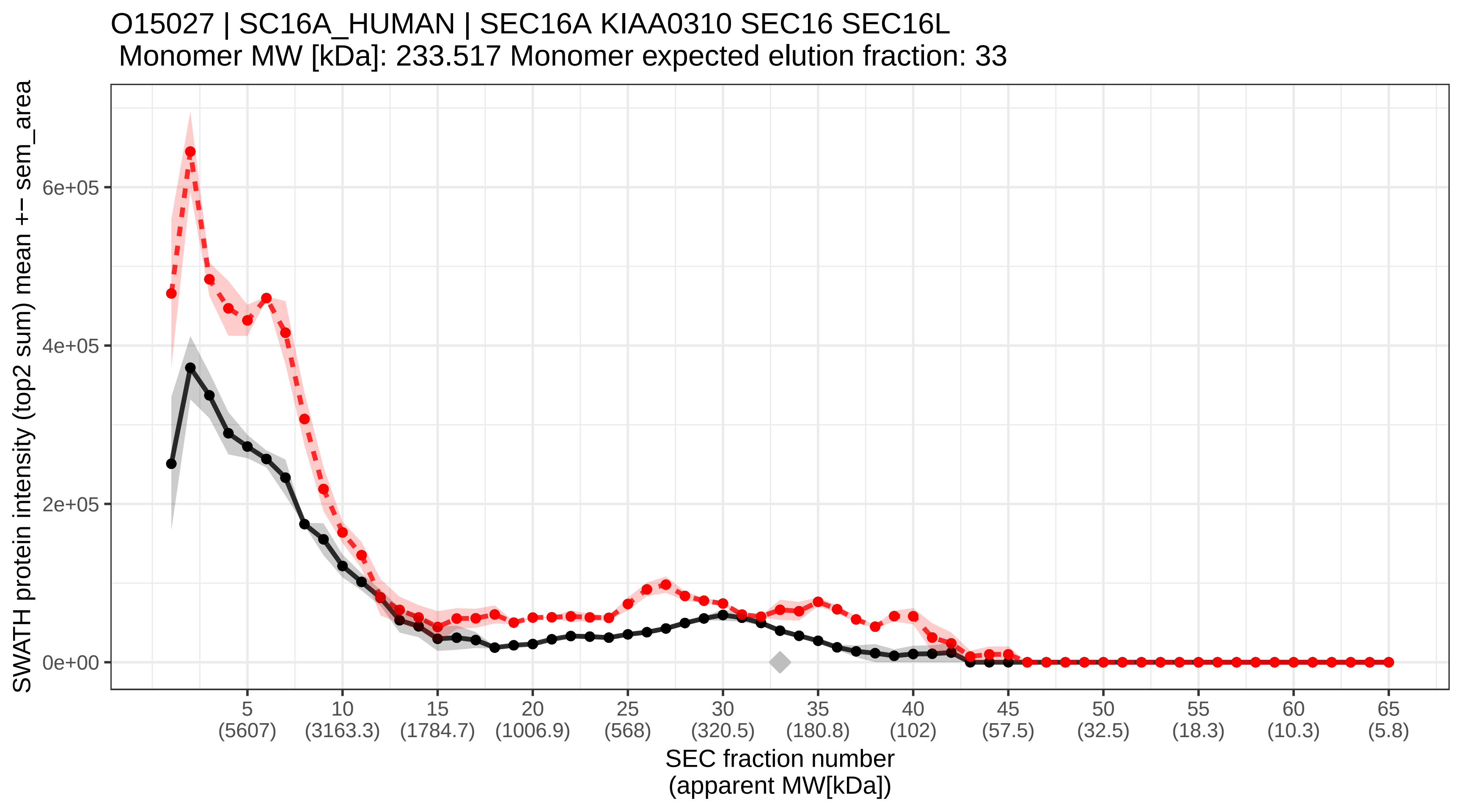

### SECchrom_O15031_PLXB2_HUMAN_PLXNB2_KIAA0315.pdf

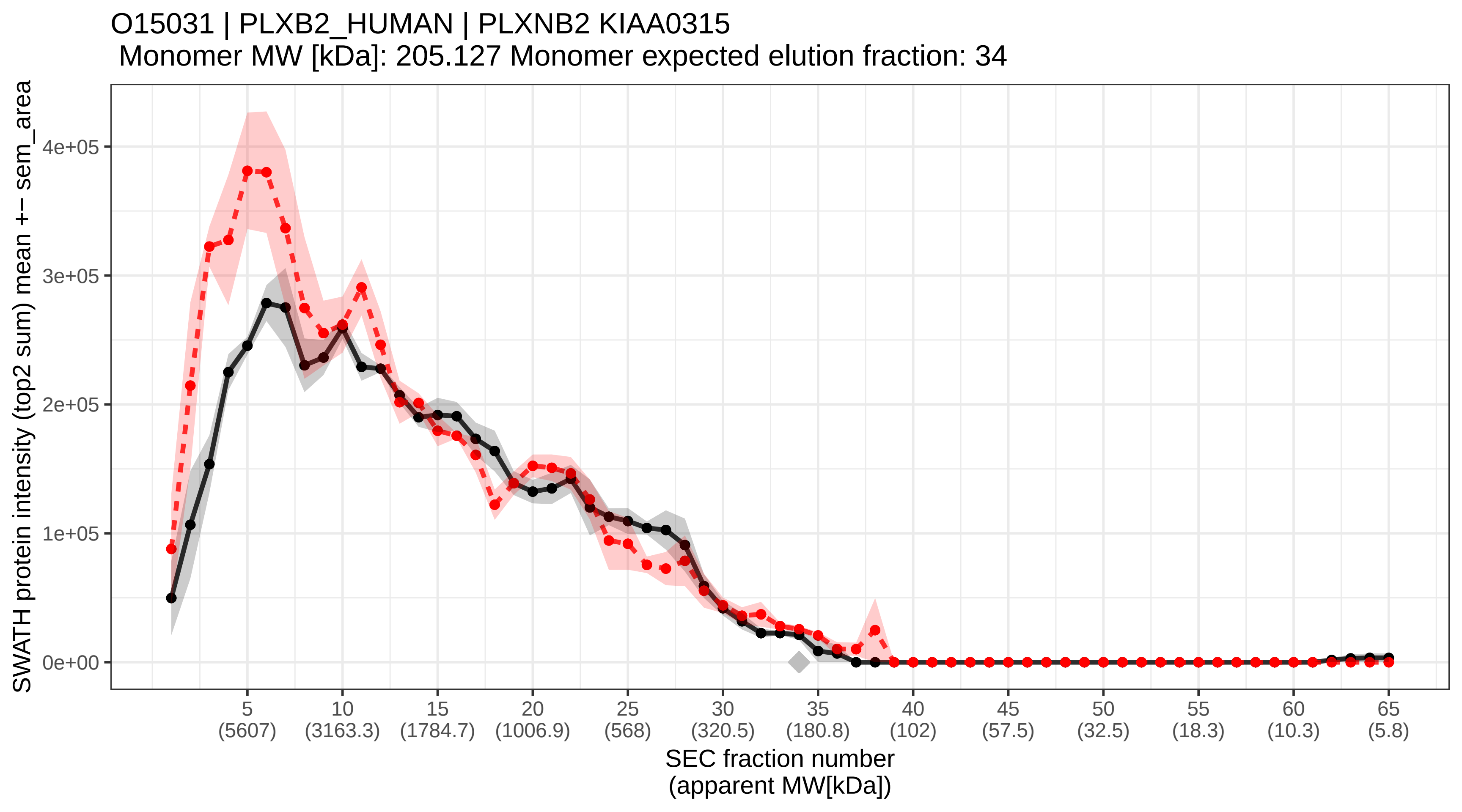

### SECchrom_O15037_KHNYN_HUMAN_KHNYN_KIAA0323.pdf

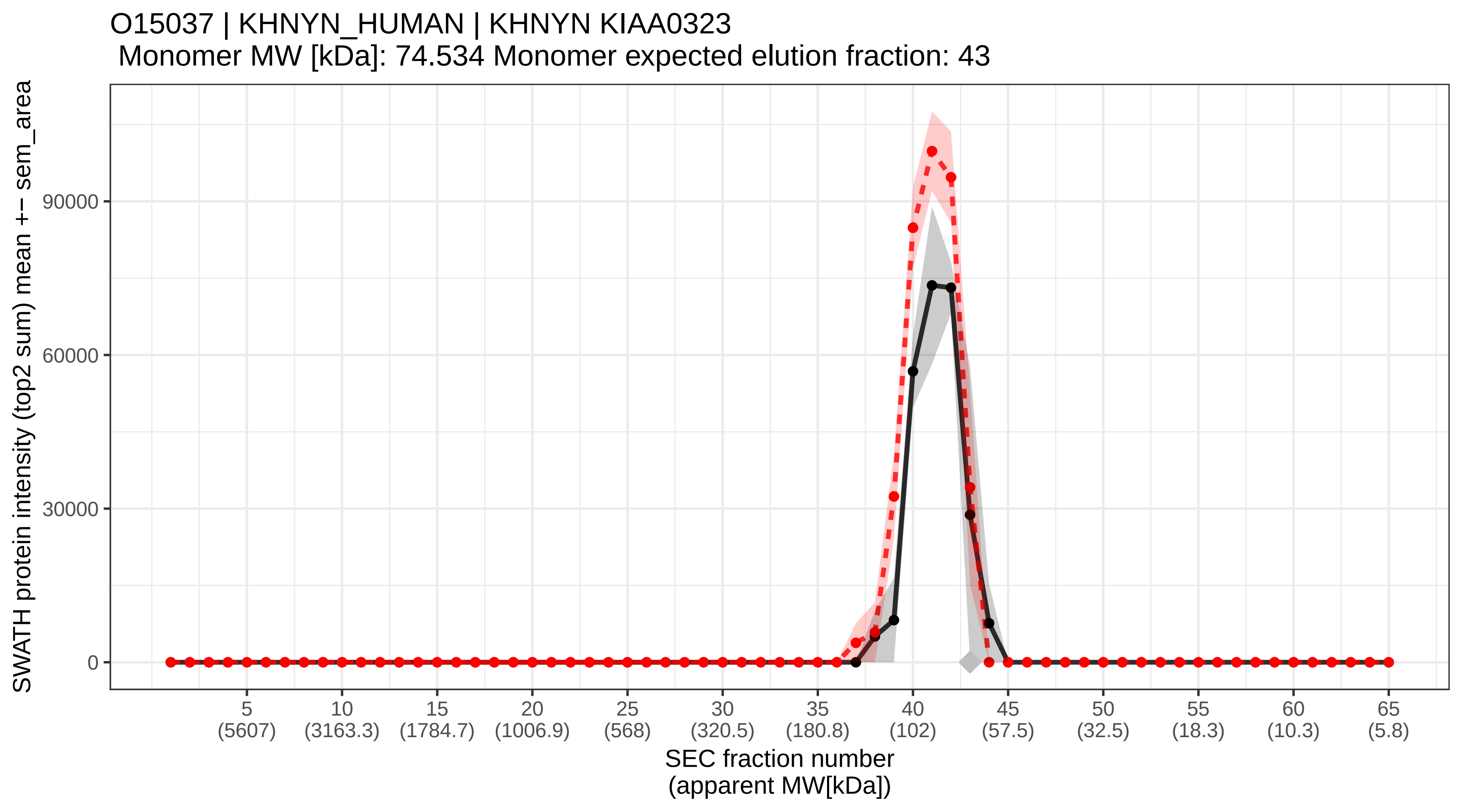

### SECchrom_O15042_SR140_HUMAN_U2SURP_KIAA0332_SR140.pdf

O15042 | SR140_HUMAN | U2SURP KIAA0332 SR140
Monomer MW [kDa]: 118.292 Monomer expected elution fraction: 39

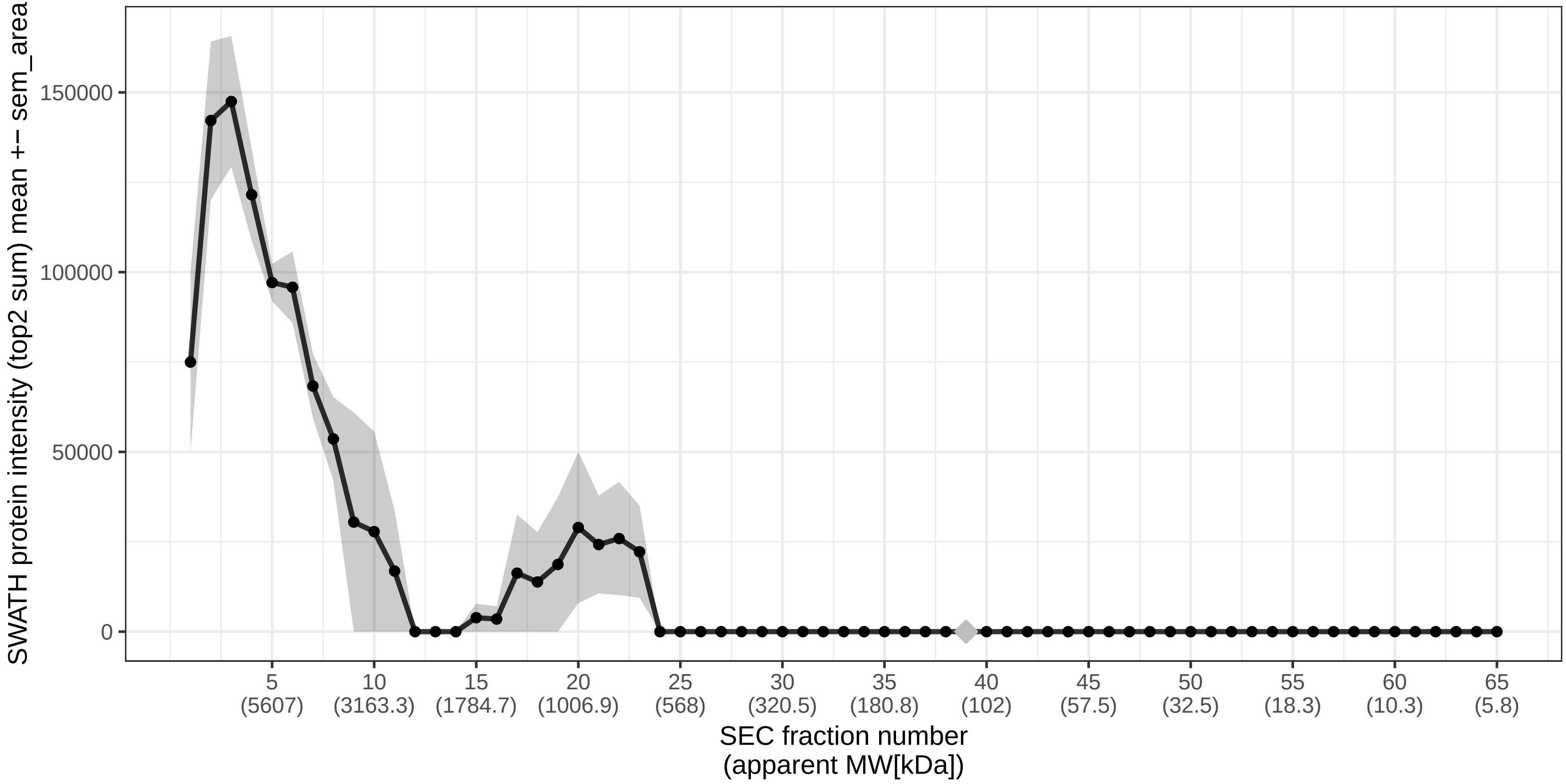

### SECchrom_O15047_SET1A_HUMAN_SETD1A_KIAA0339_KMT2F_SET1_SET1.pdf

O15047 | SET1A_HUMAN | SETD1A KIAA0339 KMT2F SET1 SET1A
Monomer MW [kDa]: 186.034 Monomer expected elution fraction: 35

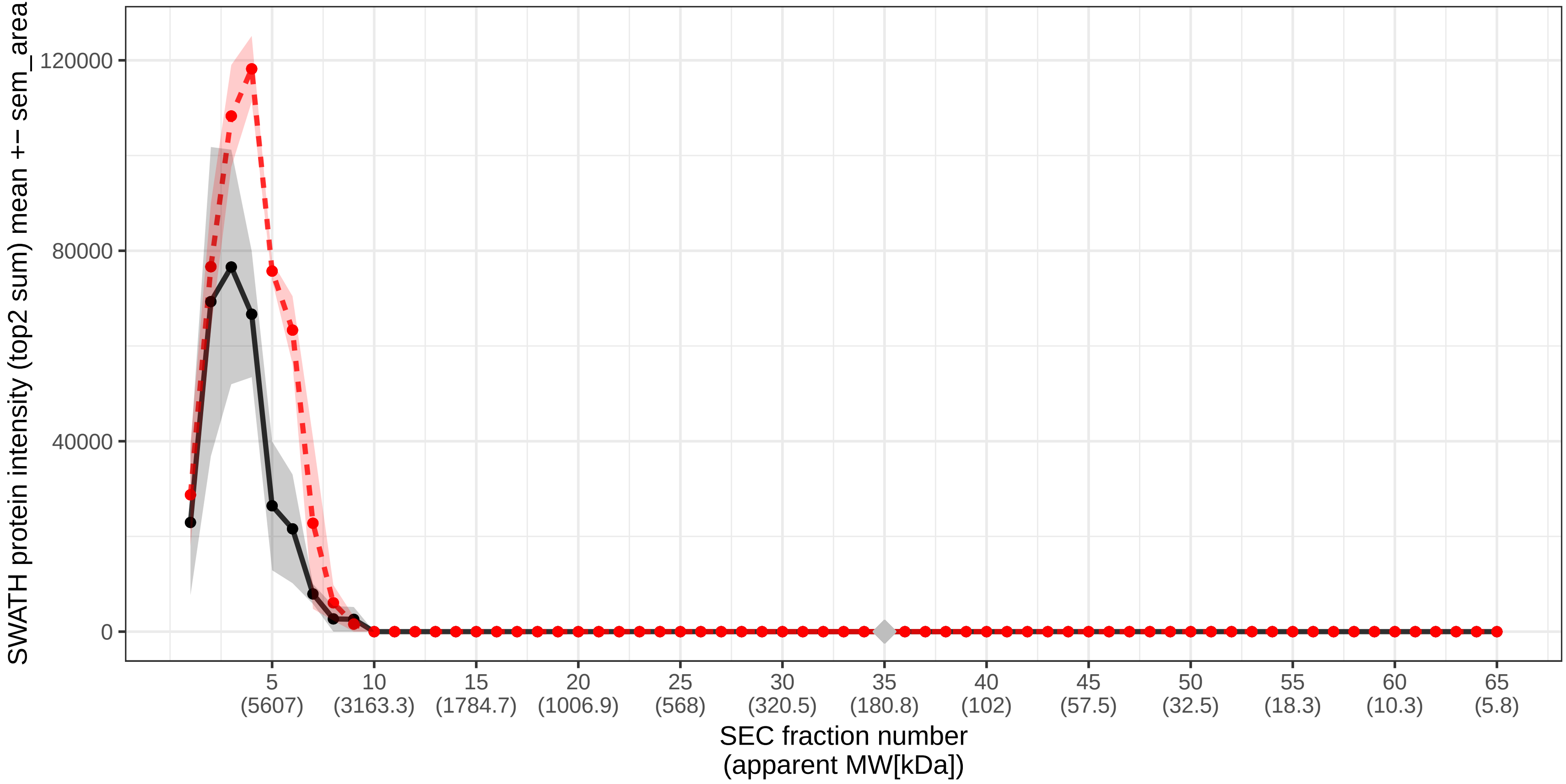

### SECchrom_O15056_SYNJ2_HUMAN_SYNJ2_KIAA0348.pdf

O15056 | SYNJ2_HUMAN | SYNJ2 KIAA0348
Monomer MW [kDa]: 165.538 Monomer expected elution fraction: 36

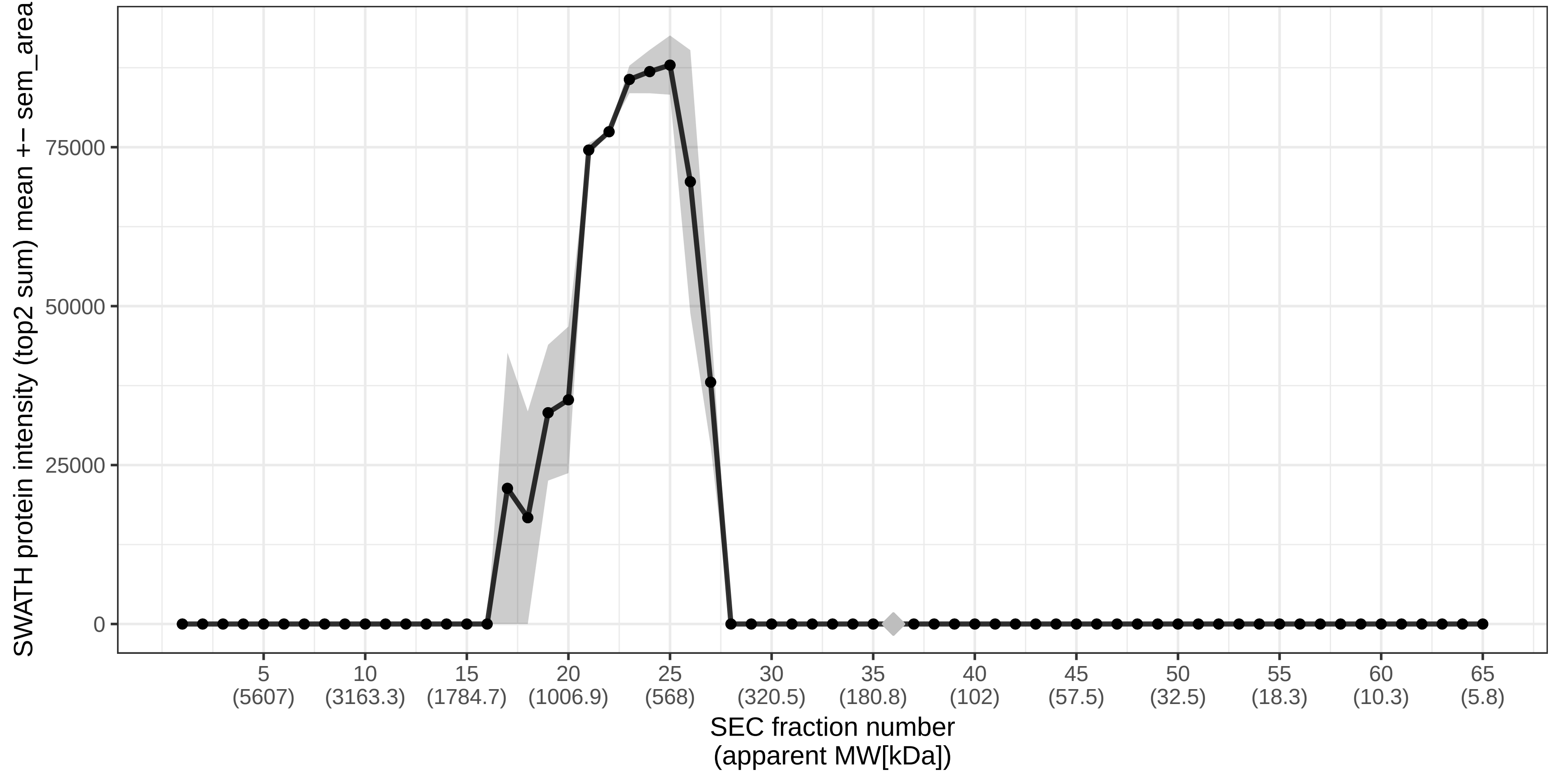

### SECchrom_O15061_SYNEM_HUMAN_SYNM_DMN_KIAA0353_SYN.pdf

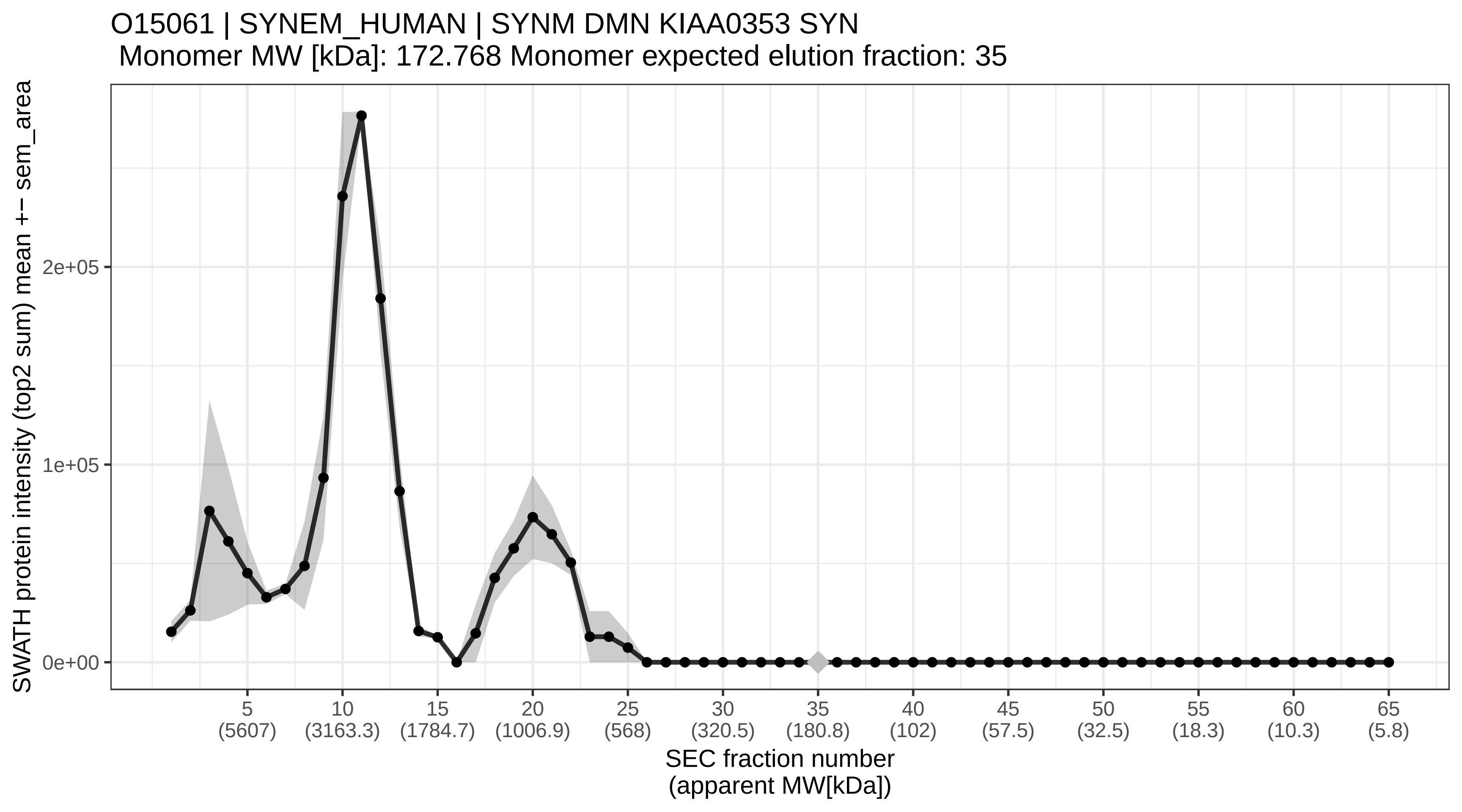

### SECchrom_O15066_KIF3B_HUMAN_KIF3B_KIAA0359.pdf

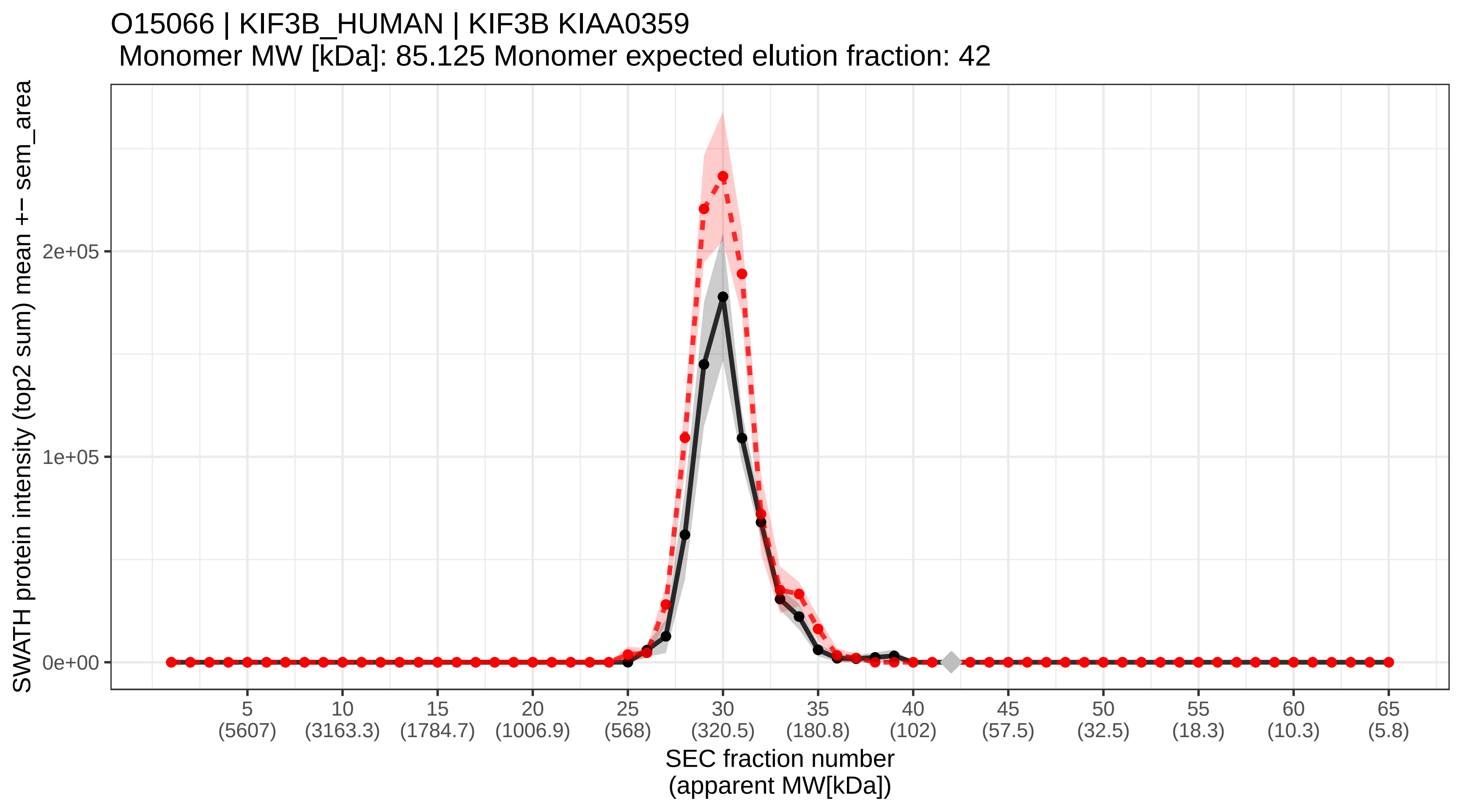

### SECchrom_O15067_PUR4_HUMAN_PFAS_KIAA0361.pdf

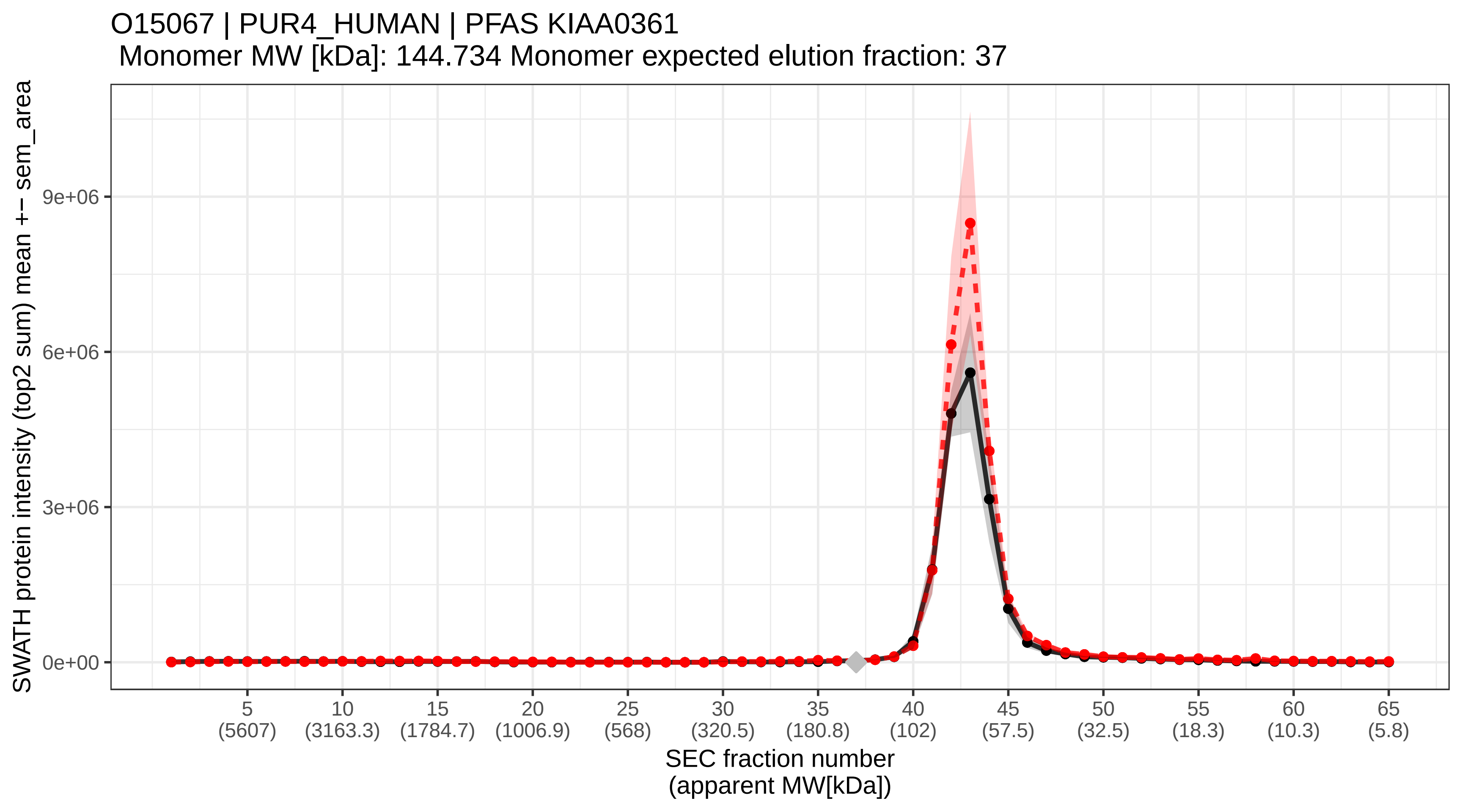

### SECchrom_O15084_ANR28_HUMAN_ANKRD28_KIAA0379.pdf

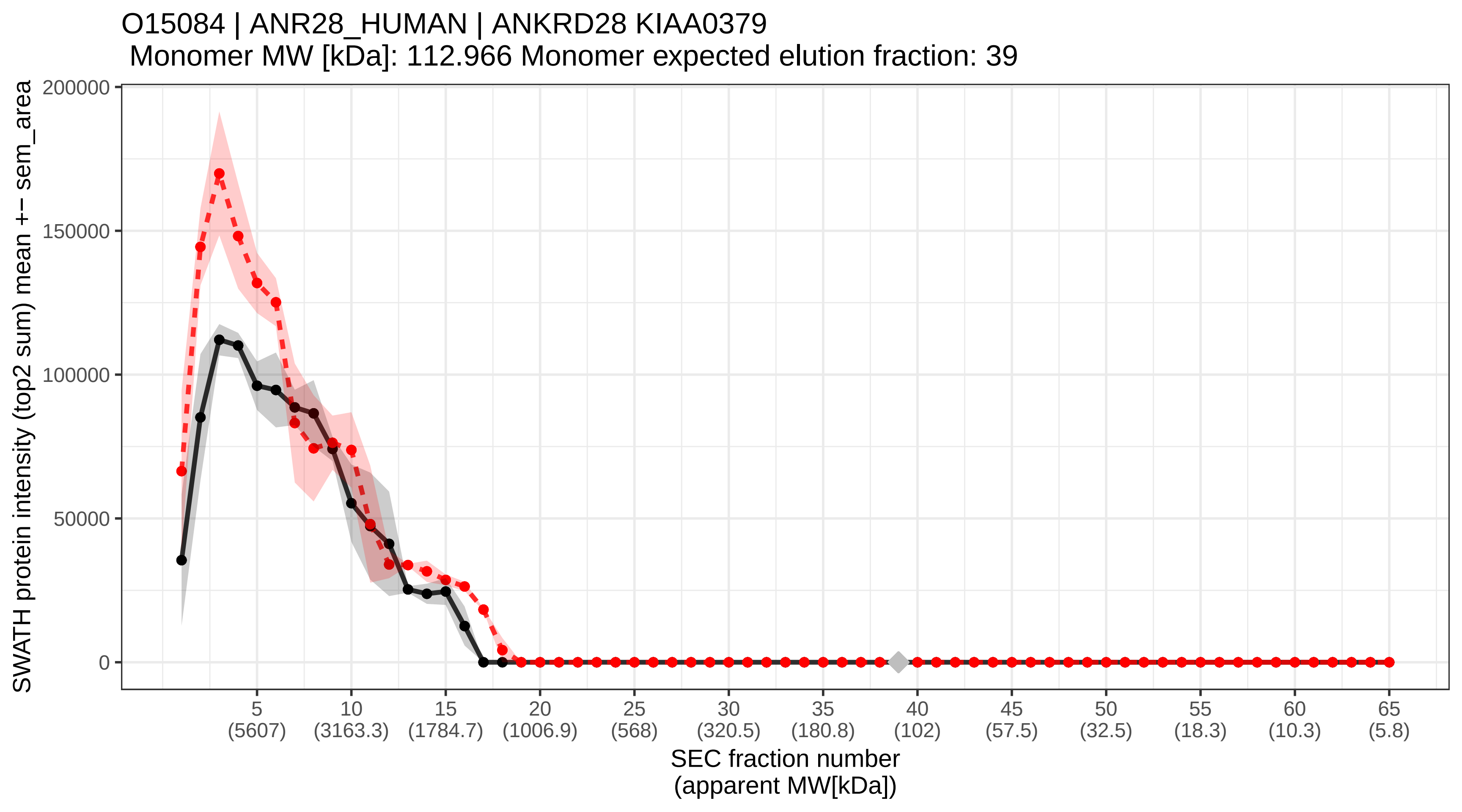

### SECchrom_O15111_IKKA_HUMAN_CHUK_IKKA_TCF16.pdf

O15111 | IKKA_HUMAN | CHUK IKKA TCF16
Monomer MW [kDa]: 84.64 Monomer expected elution fraction: 42

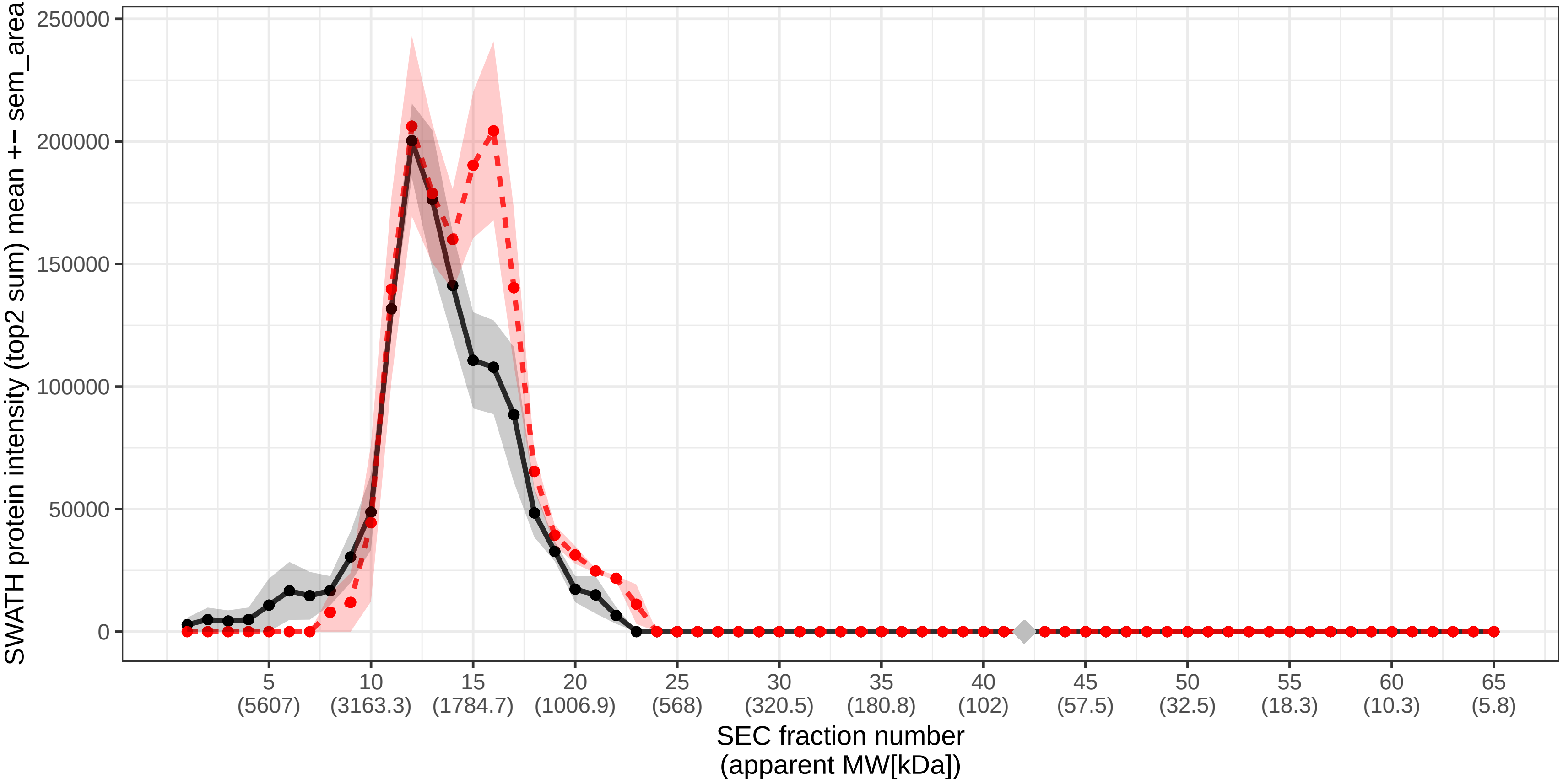

### SECchrom_O15116_LSM1_HUMAN_LSM1_CASM.pdf

O15116 | LSM1_HUMAN | LSM1 CASM
Monomer MW [kDa]: 15.179 Monomer expected elution fraction: 57

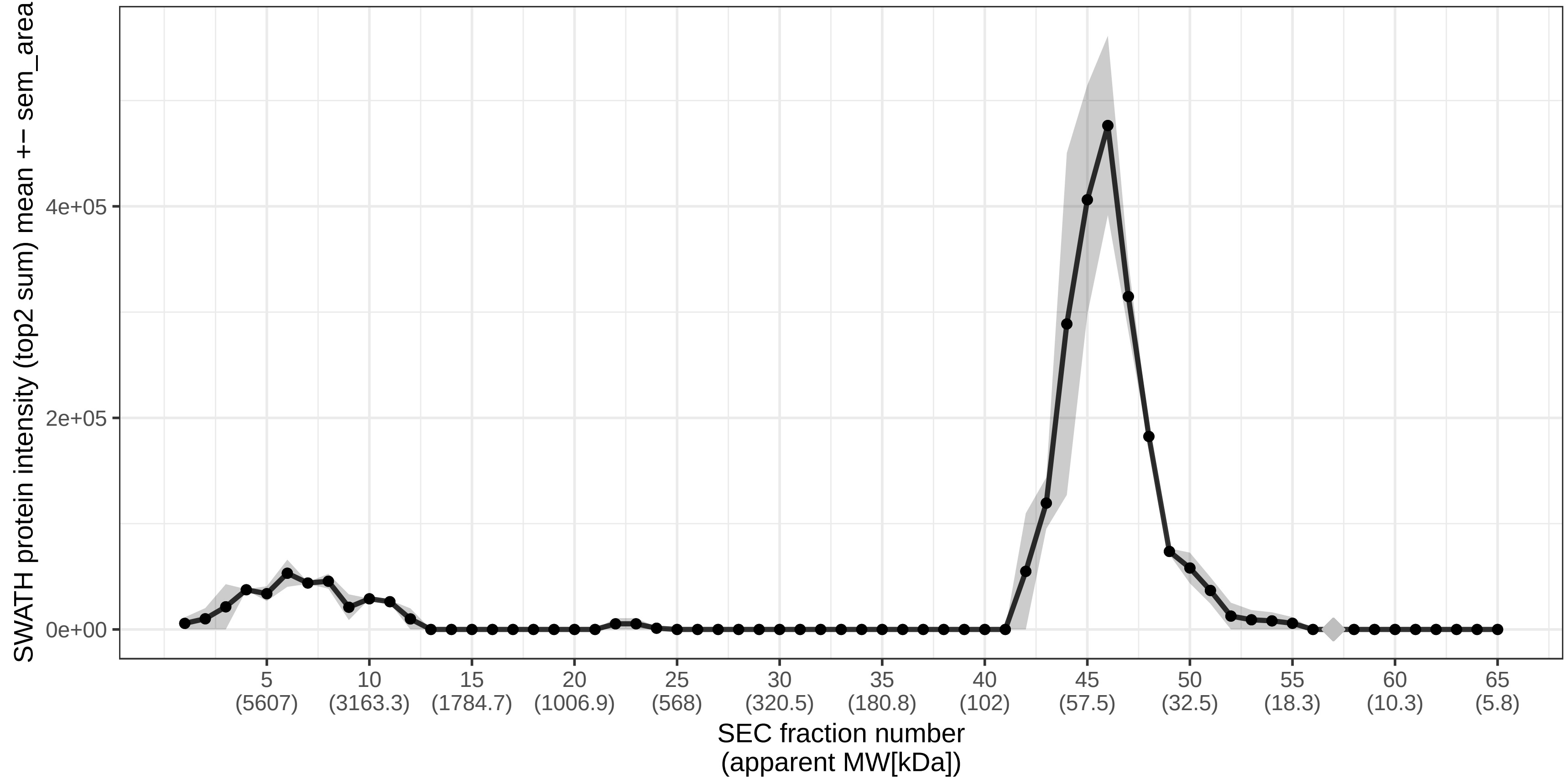

### SECchrom_O15118_NPC1_HUMAN_NPC1.pdf

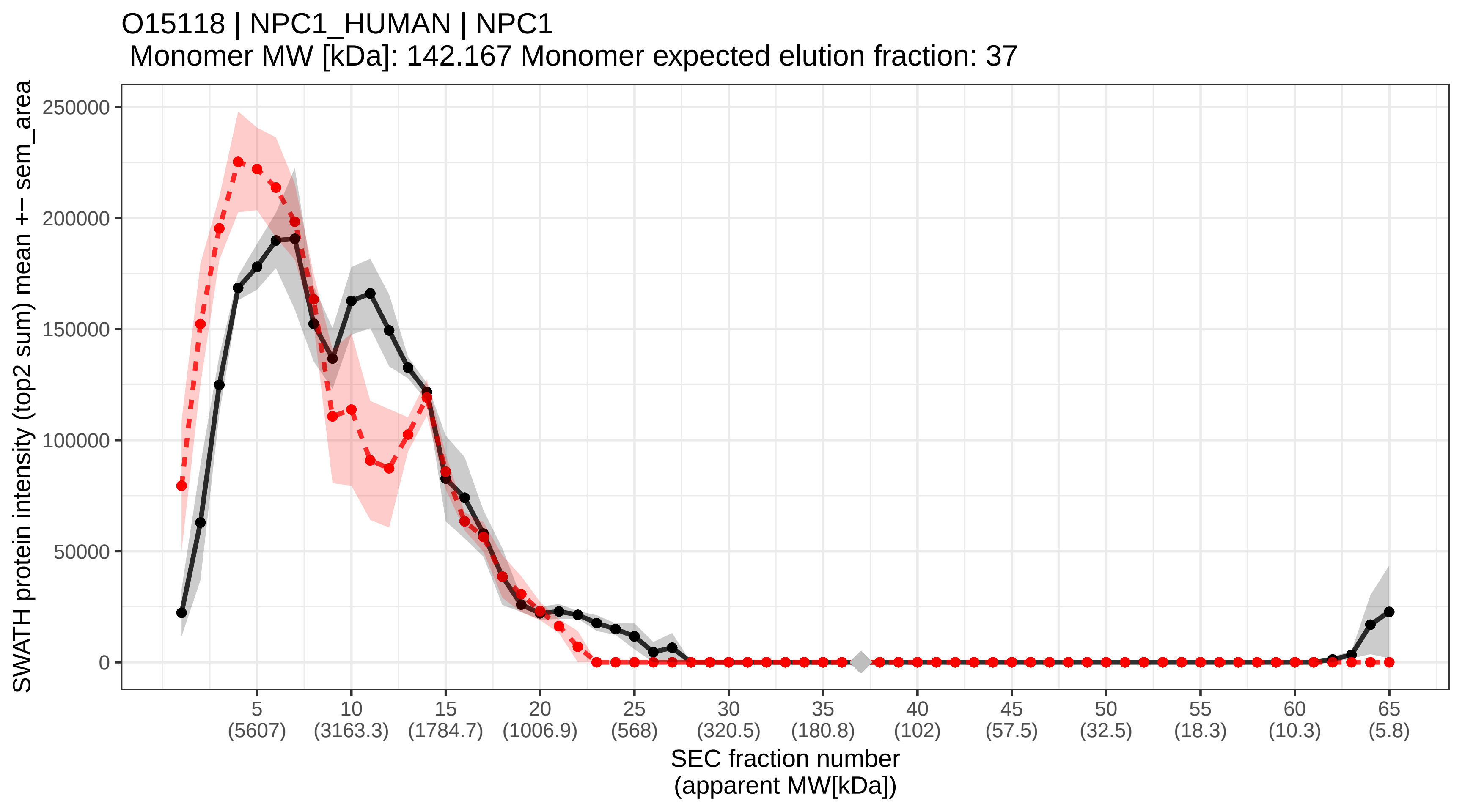

### SECchrom_O15121_DEGS1_HUMAN_DEGS1_DES1_MLD_MIG15.pdf

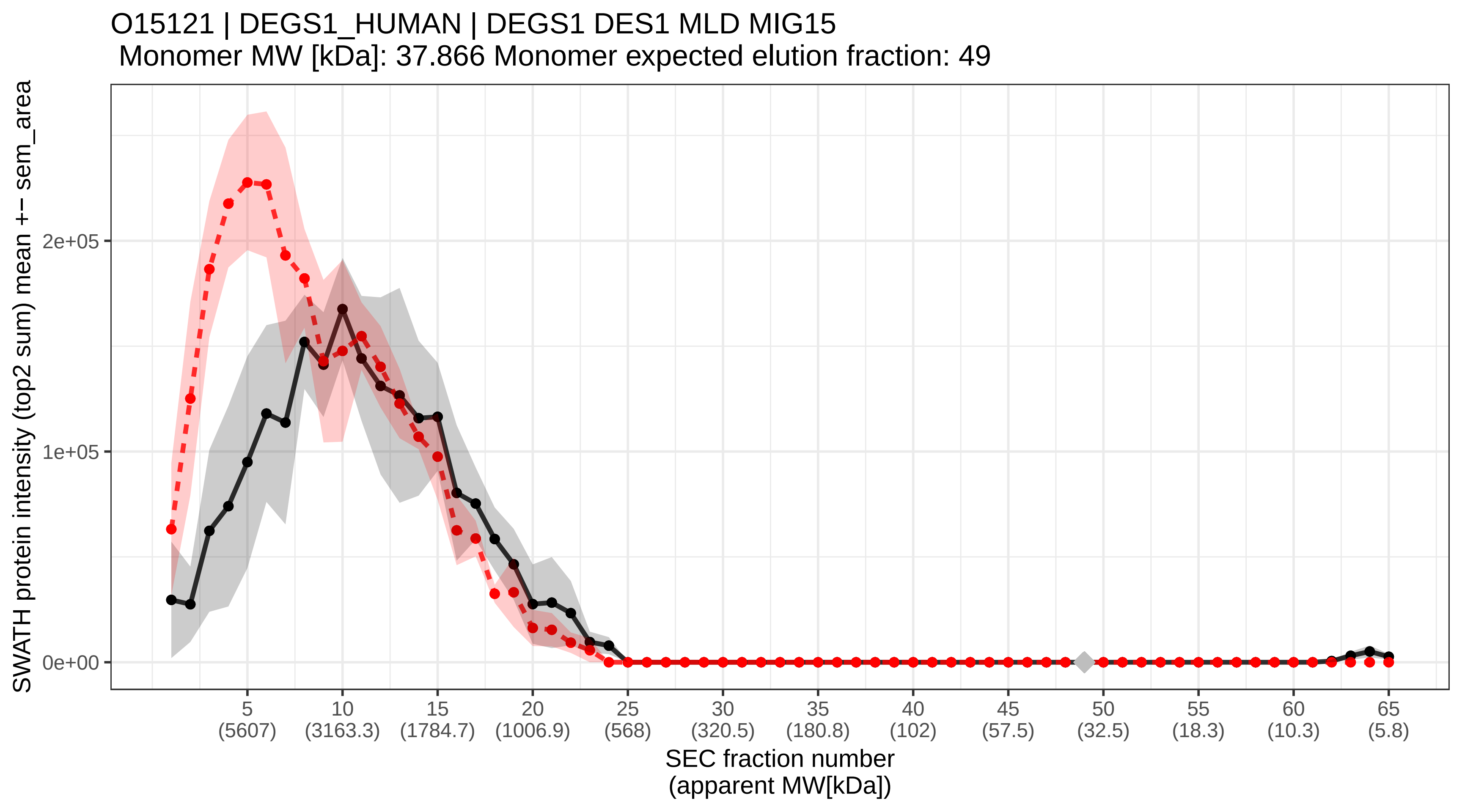

### SECchrom_O15126_SCAM1_HUMAN_SCAMP1_SCAMP.pdf

O15126 | SCAM1_HUMAN | SCAMP1 SCAMP
Monomer MW [kDa]: 37.92 Monomer expected elution fraction: 49

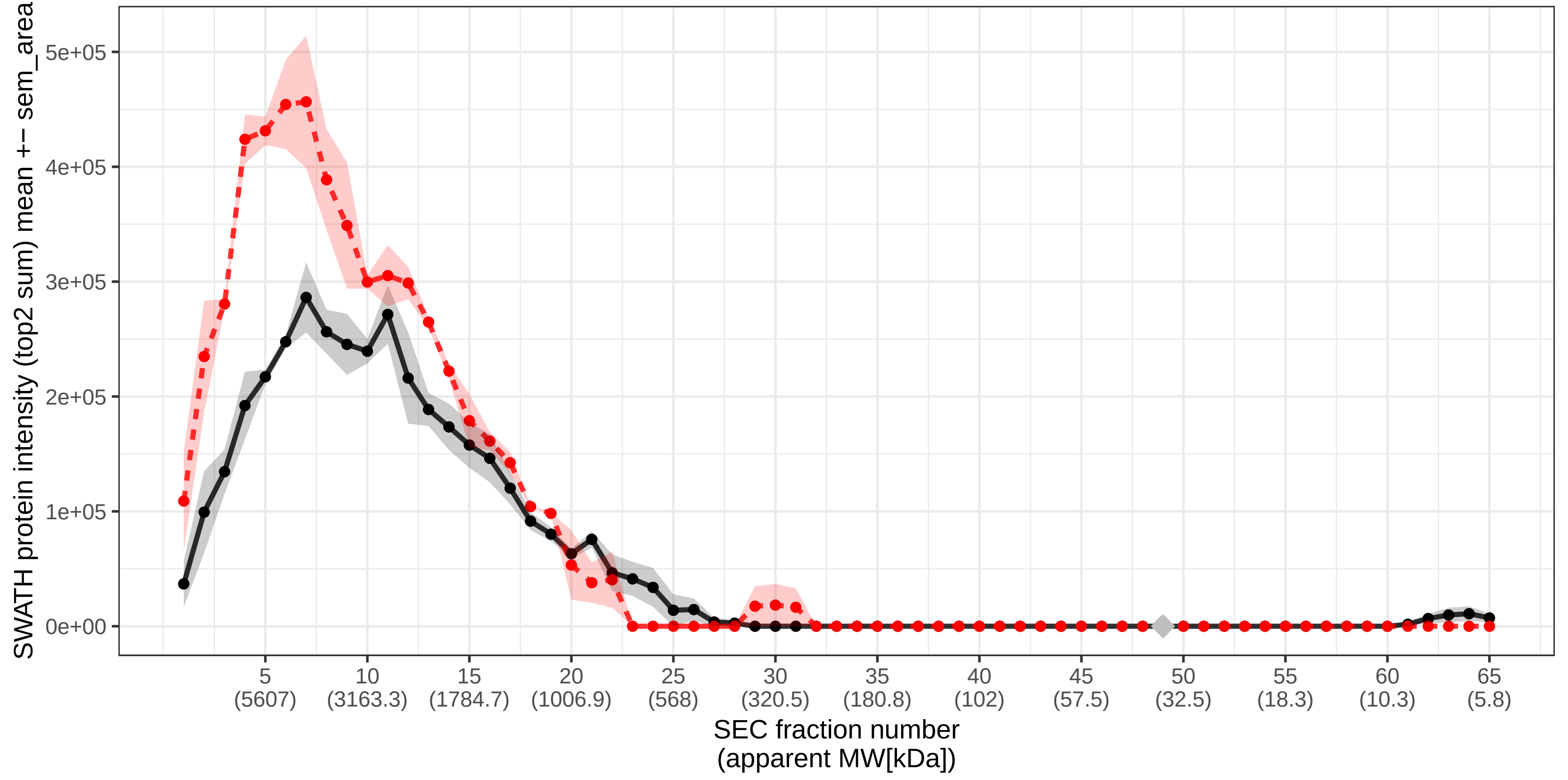

### SECchrom_O15127_SCAM2_HUMAN_SCAMP2.pdf

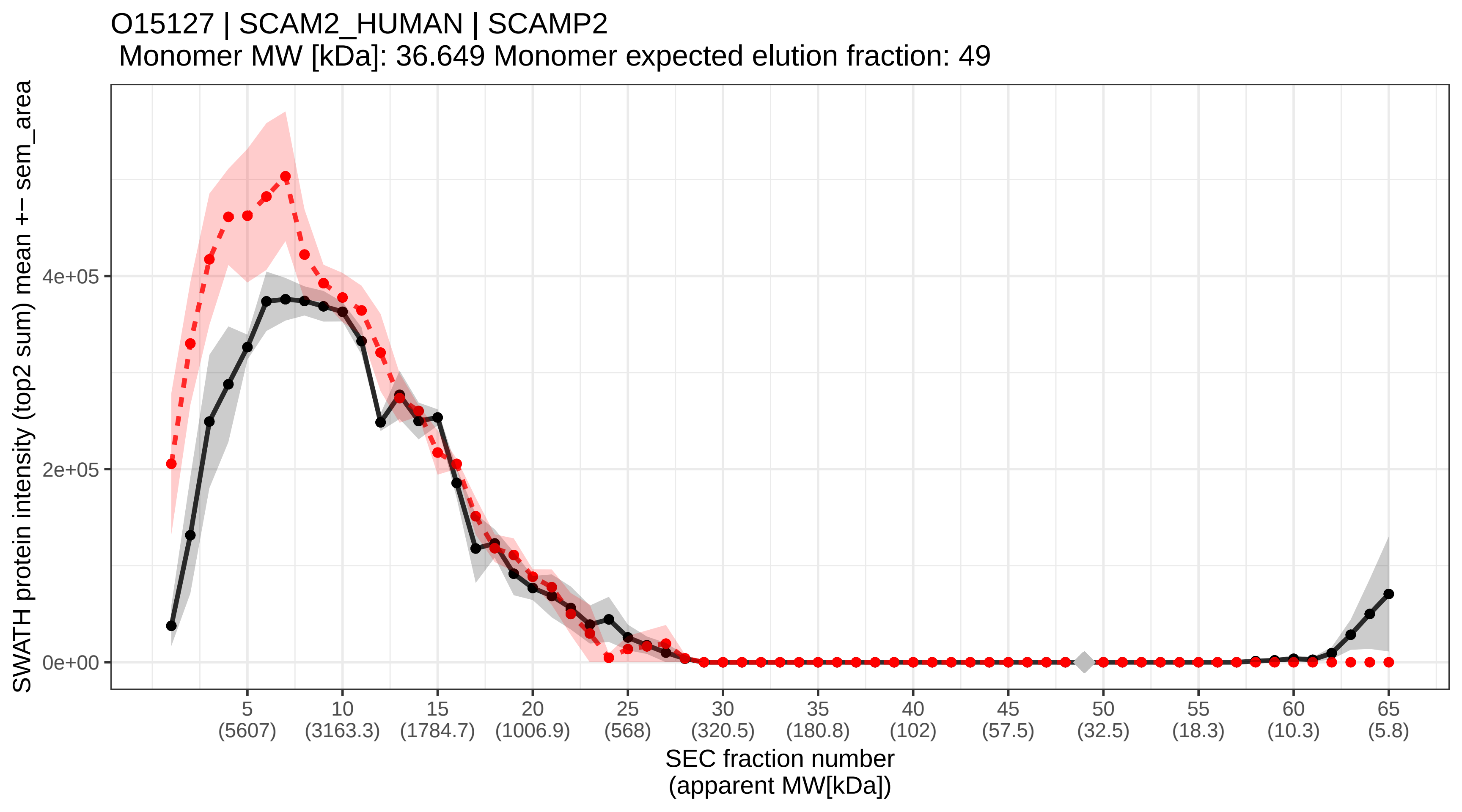

### SECchrom_O15143_ARC1B_HUMAN_ARPC1B_ARC41.pdf

O15143 | ARC1B_HUMAN | ARPC1B ARC41
Monomer MW [kDa]: 40.95 Monomer expected elution fraction: 48

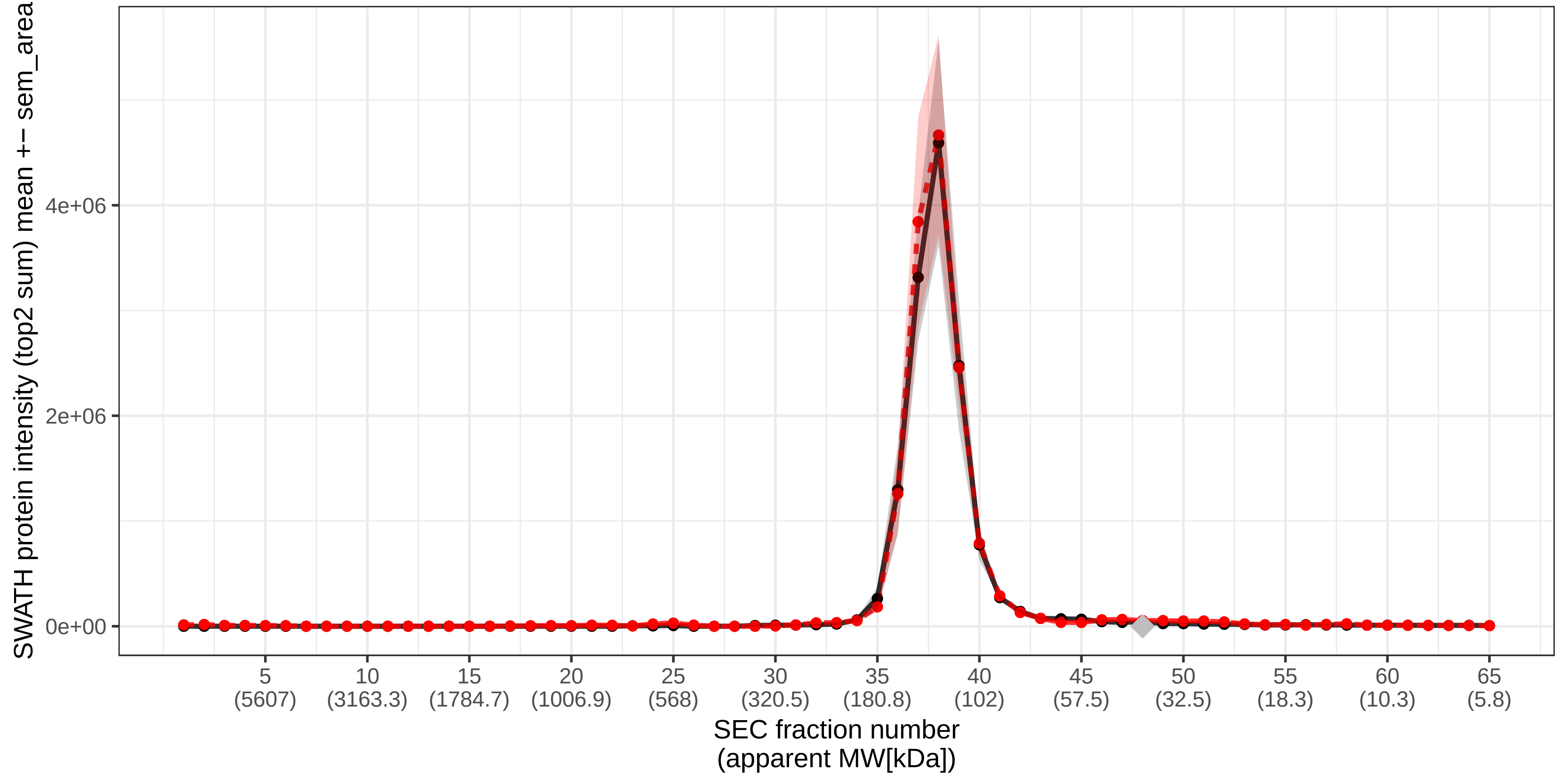

### SECchrom_O15144_ARPC2_HUMAN_ARPC2_ARC34_PRO2446.pdf

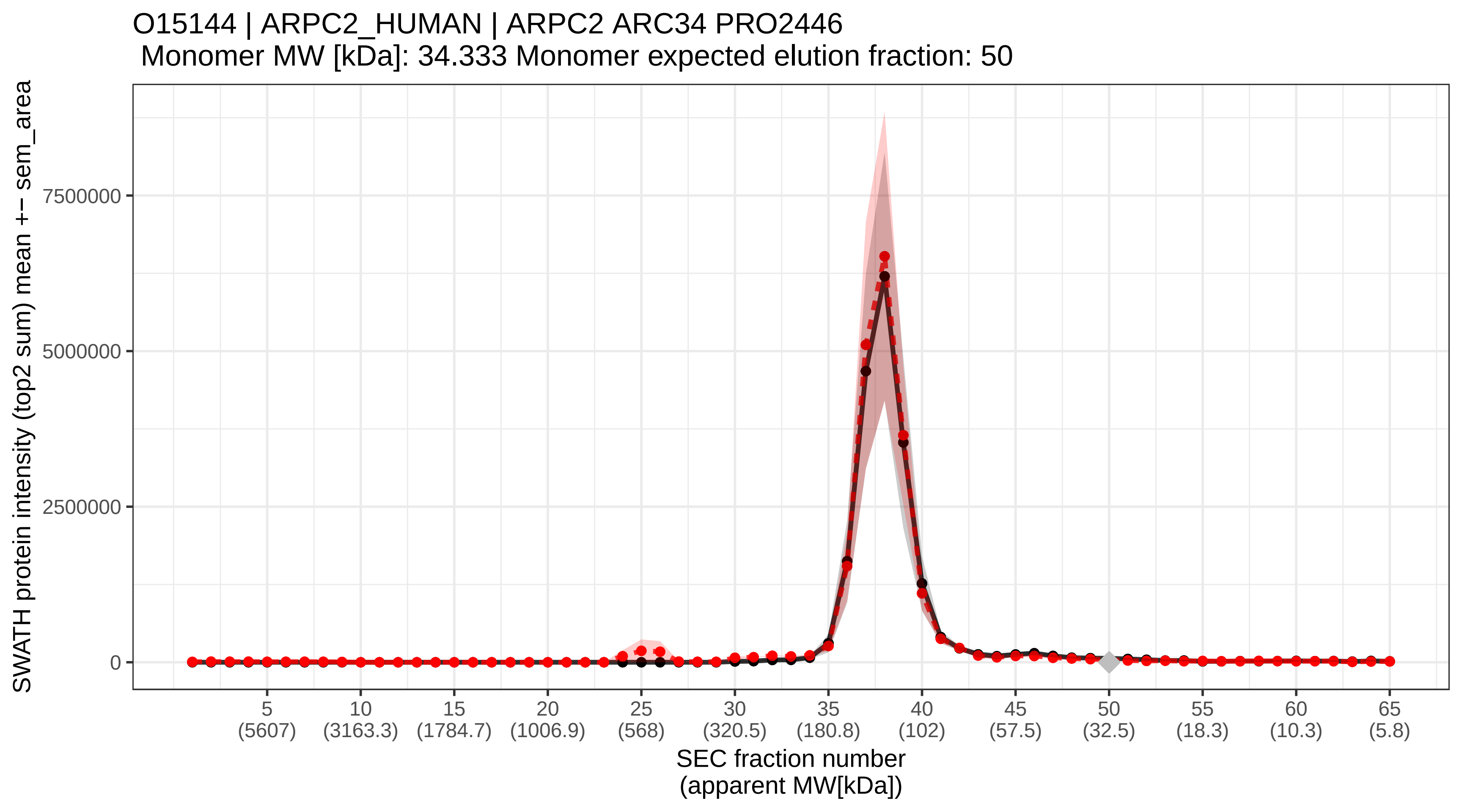

### SECchrom_O15145_ARPC3_HUMAN_ARPC3_ARC21.pdf

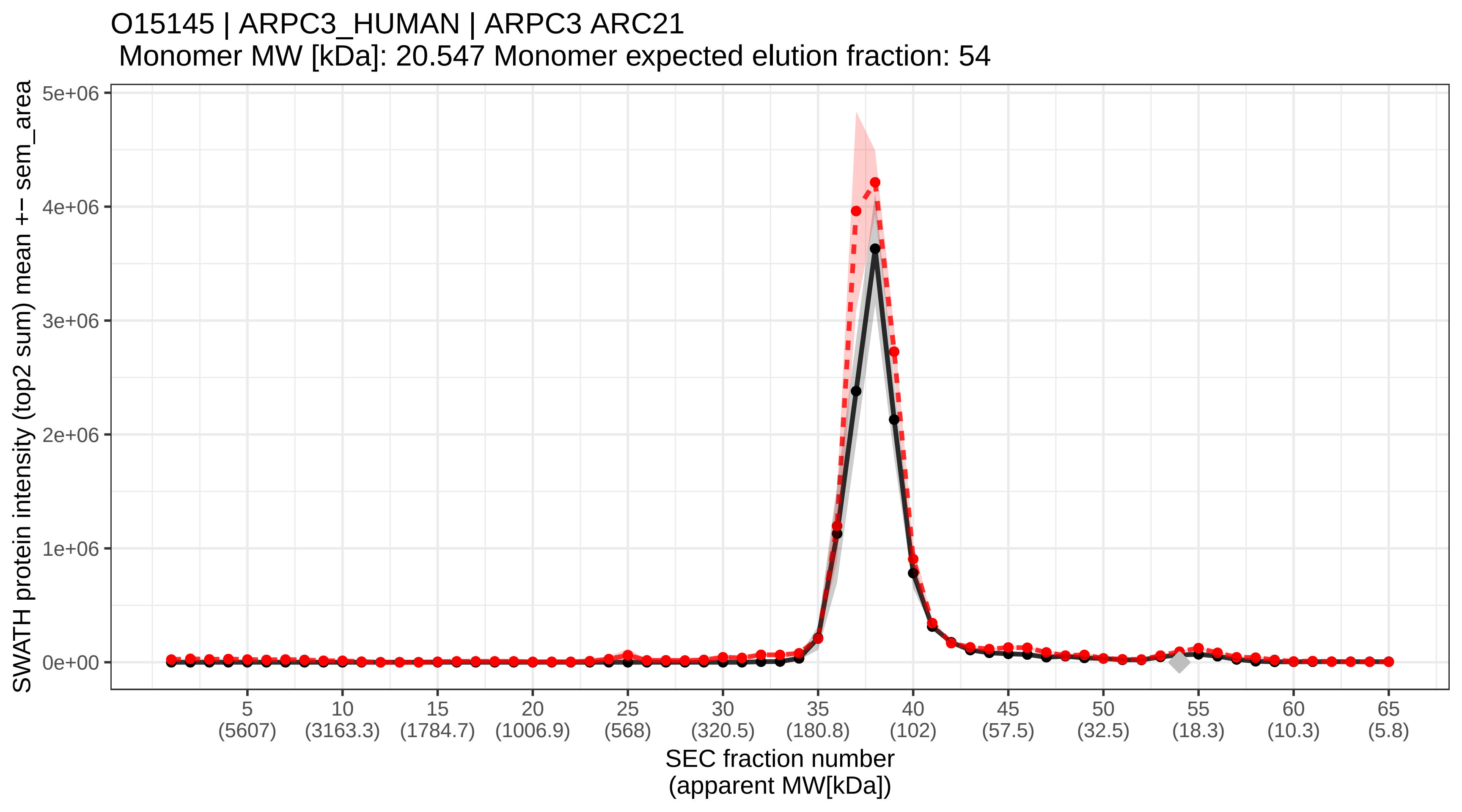

### SECchrom_O15155_BET1_HUMAN_BET1.pdf

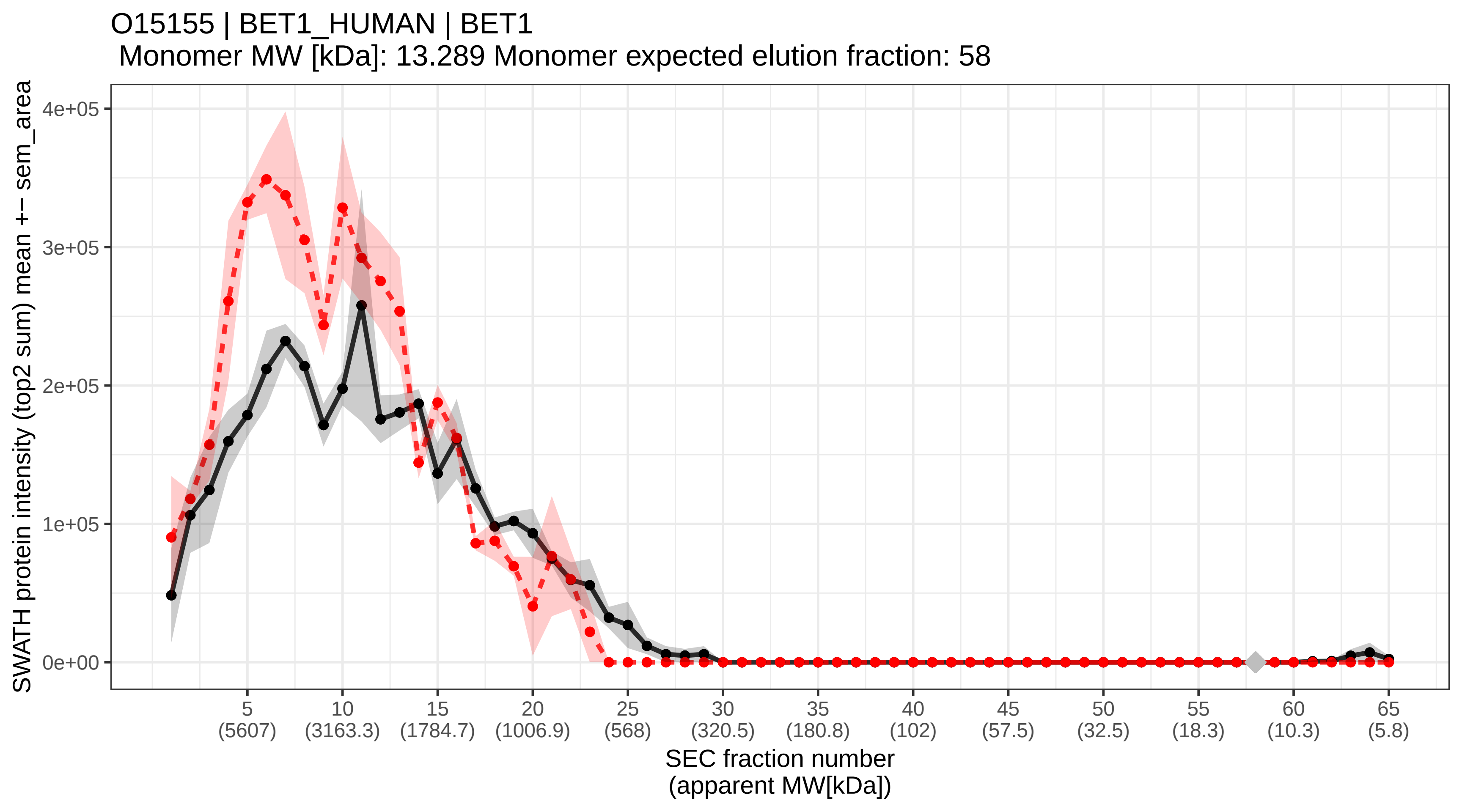

### SECchrom_O15160_RPAC1_HUMAN_POLR1C_POLR1E.pdf

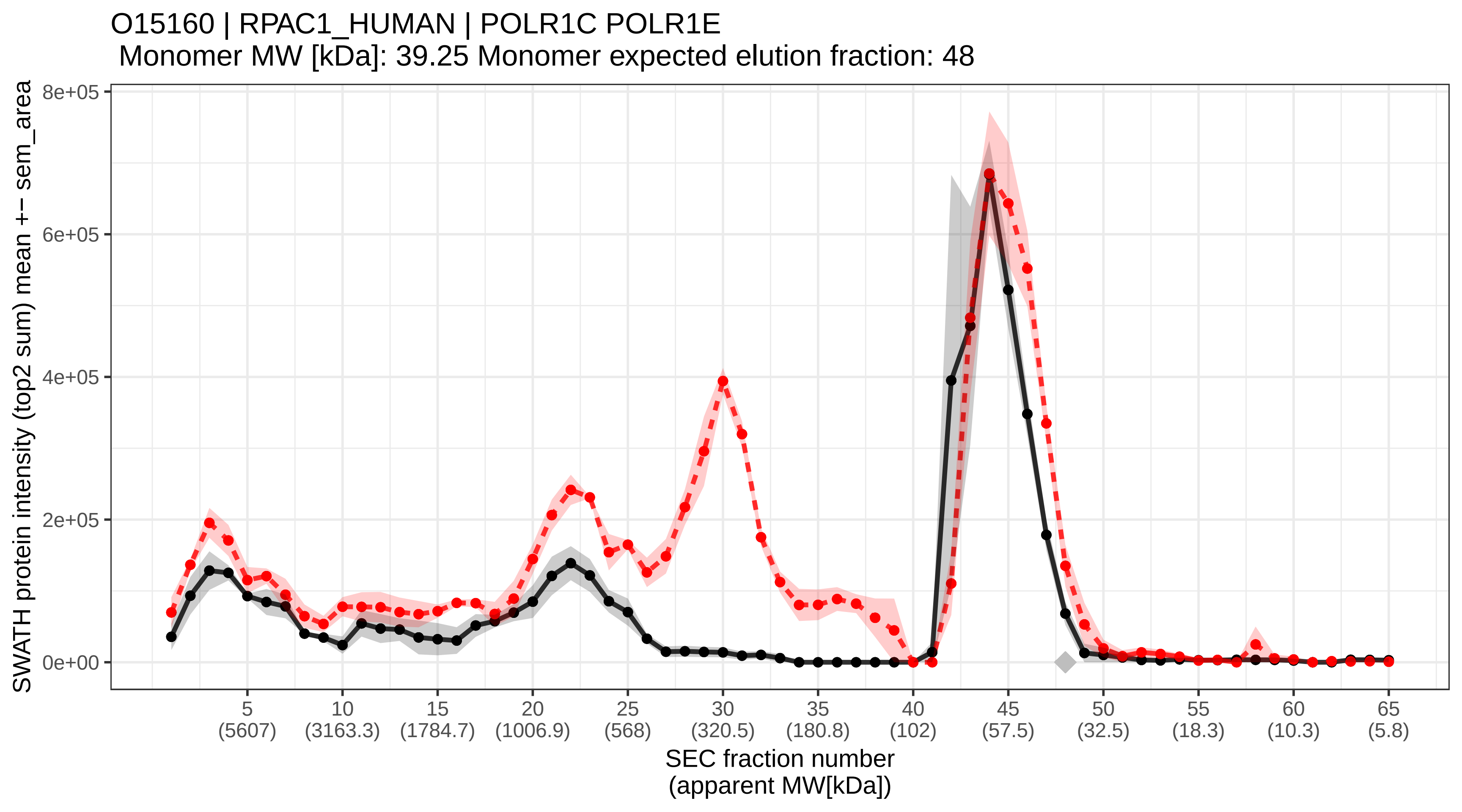

### SECchrom_O15164_TIF1A_HUMAN_TRIM24_RNF82_TIF1_TIF1A.pdf

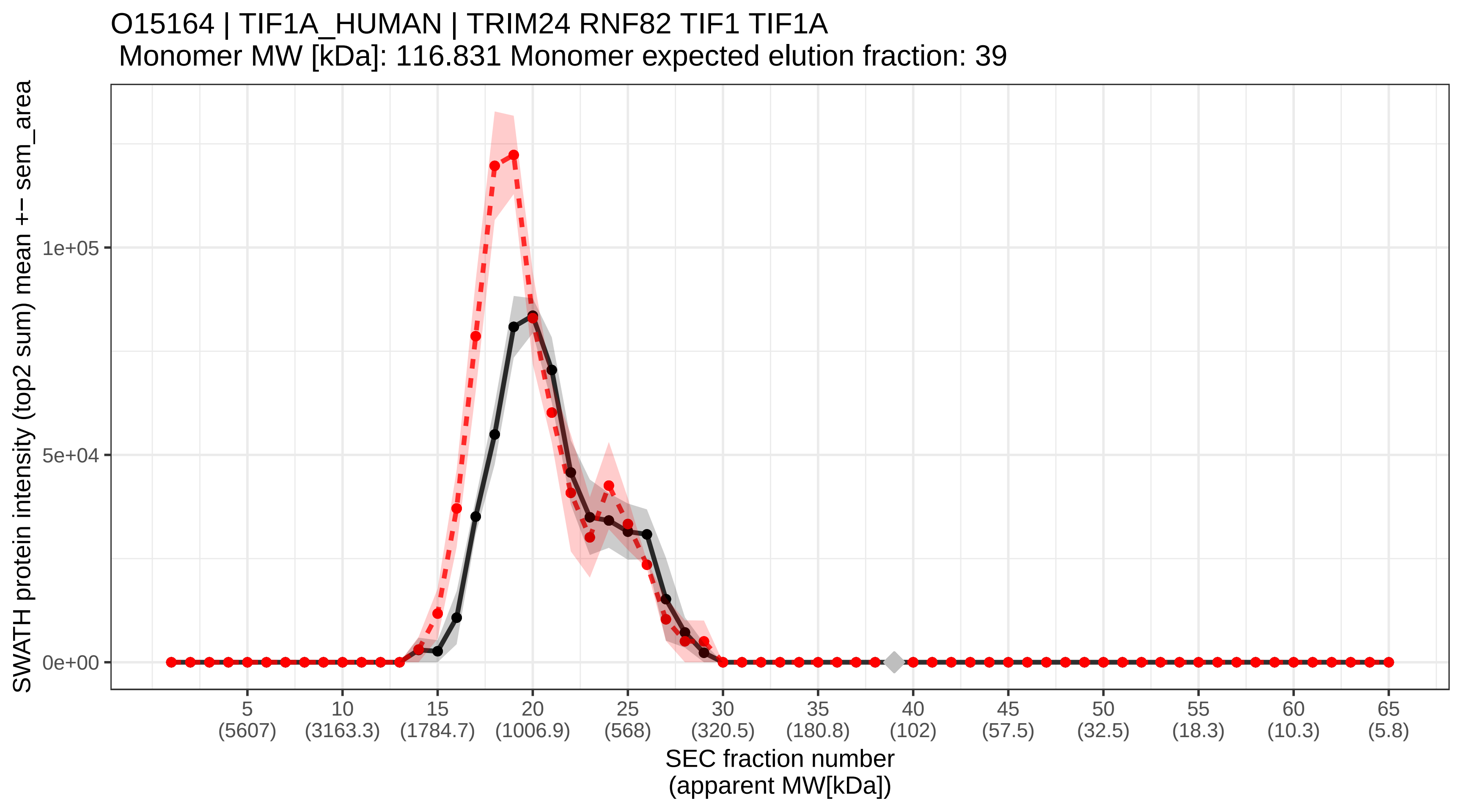

### SECchrom_O15173_PGRC2_HUMAN_PGRMC2_DG6_PMBP.pdf

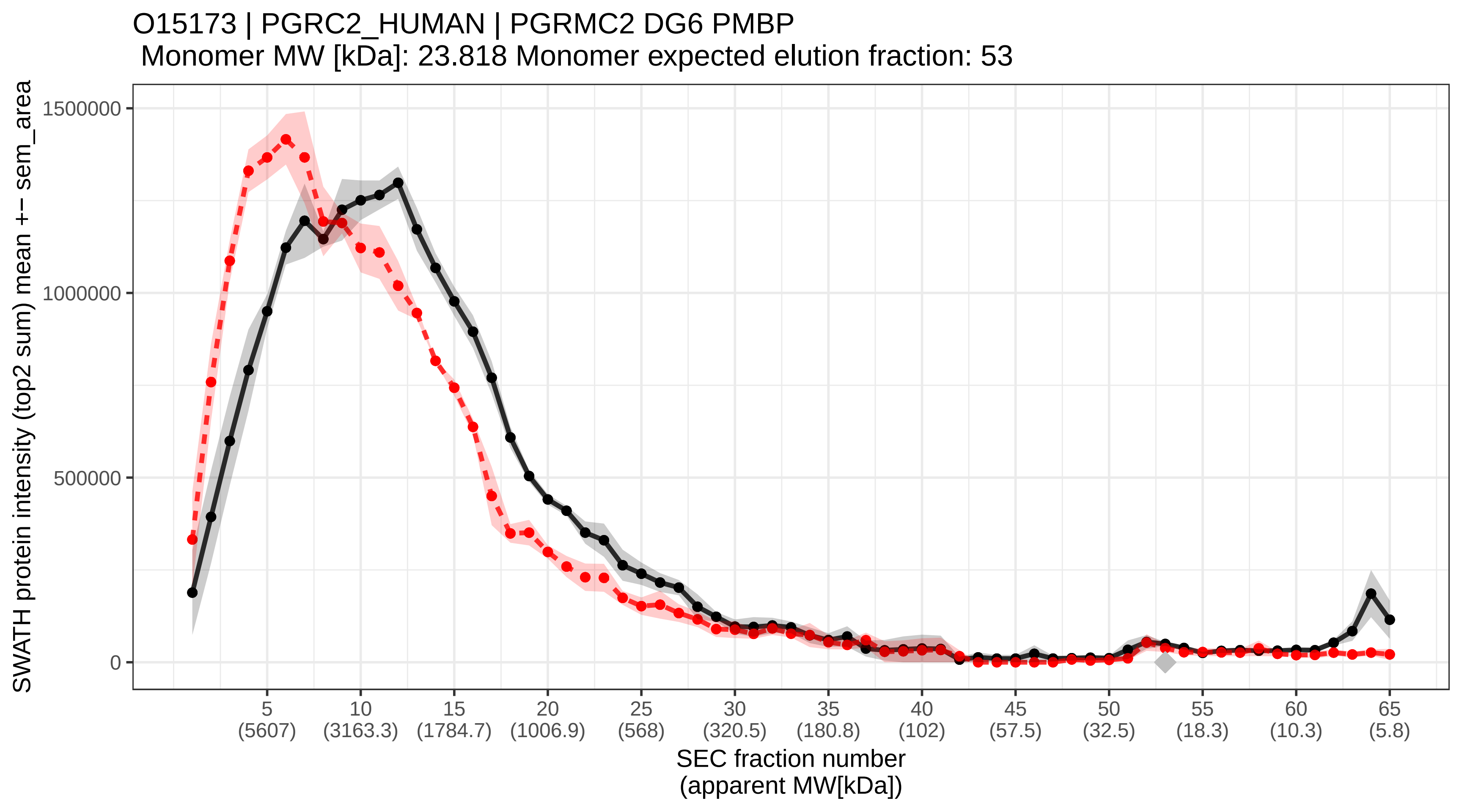

### SECchrom_O15212_PFD6_HUMAN_PFDN6_HKE2_PFD6.pdf

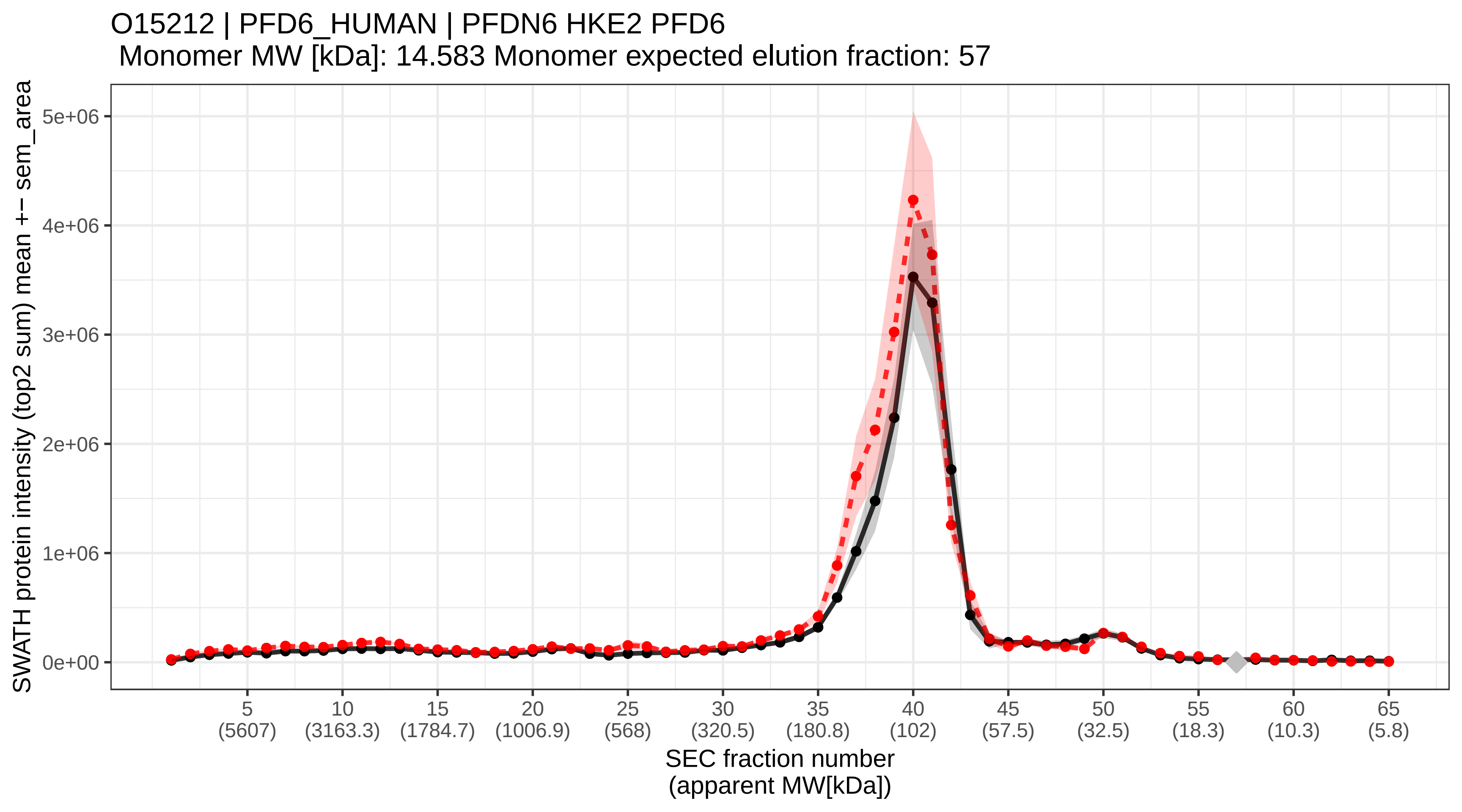

### SECchrom_O15226_NKRF_HUMAN_NKRF_ITBA4_NRF.pdf

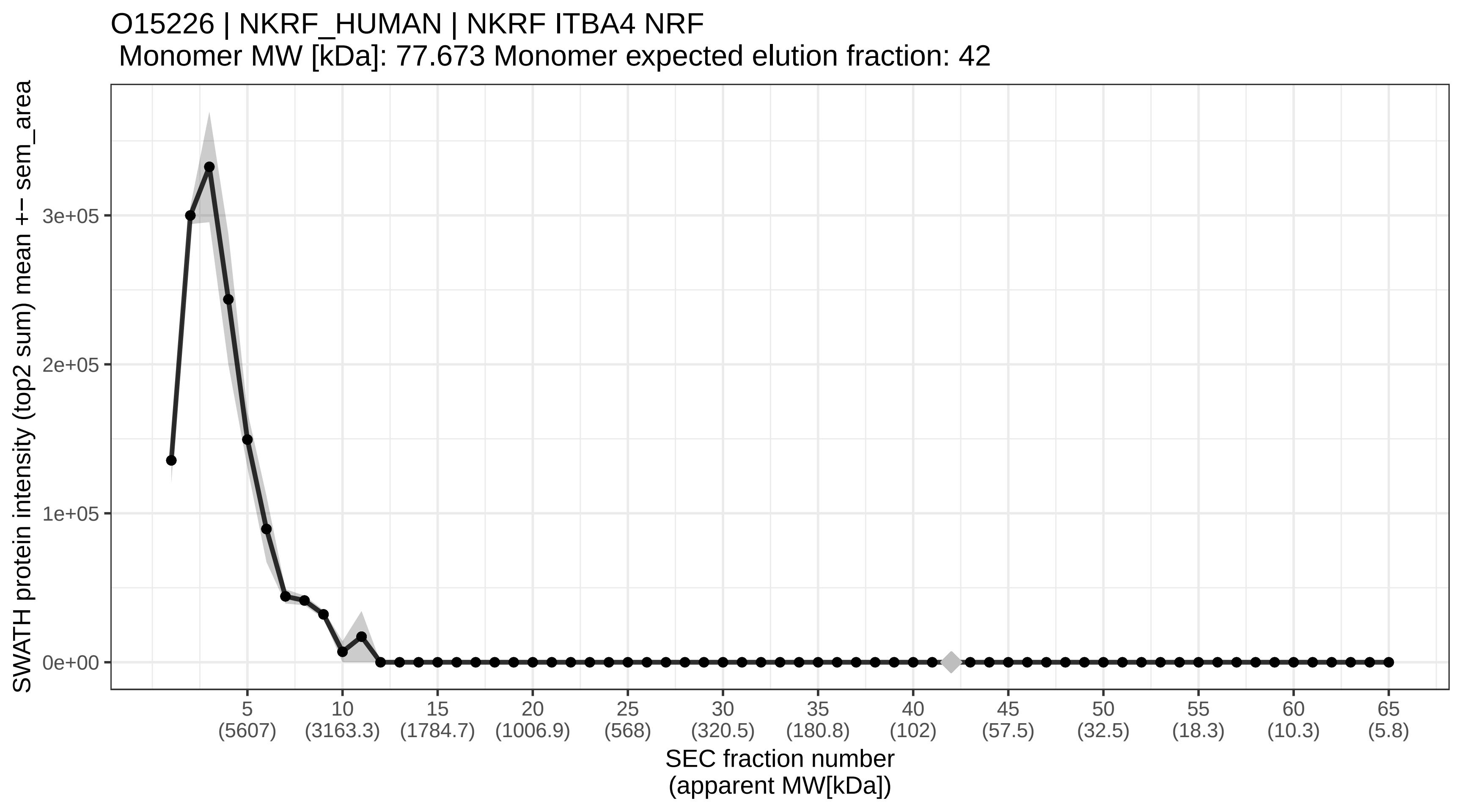

### SECchrom_O15228_GNPAT_HUMAN_GNPAT_DAPAT_DHAPAT.pdf

O15228 | GNPAT_HUMAN | GNPAT DAPAT DHAPAT
Monomer MW [kDa]: 77.188 Monomer expected elution fraction: 42

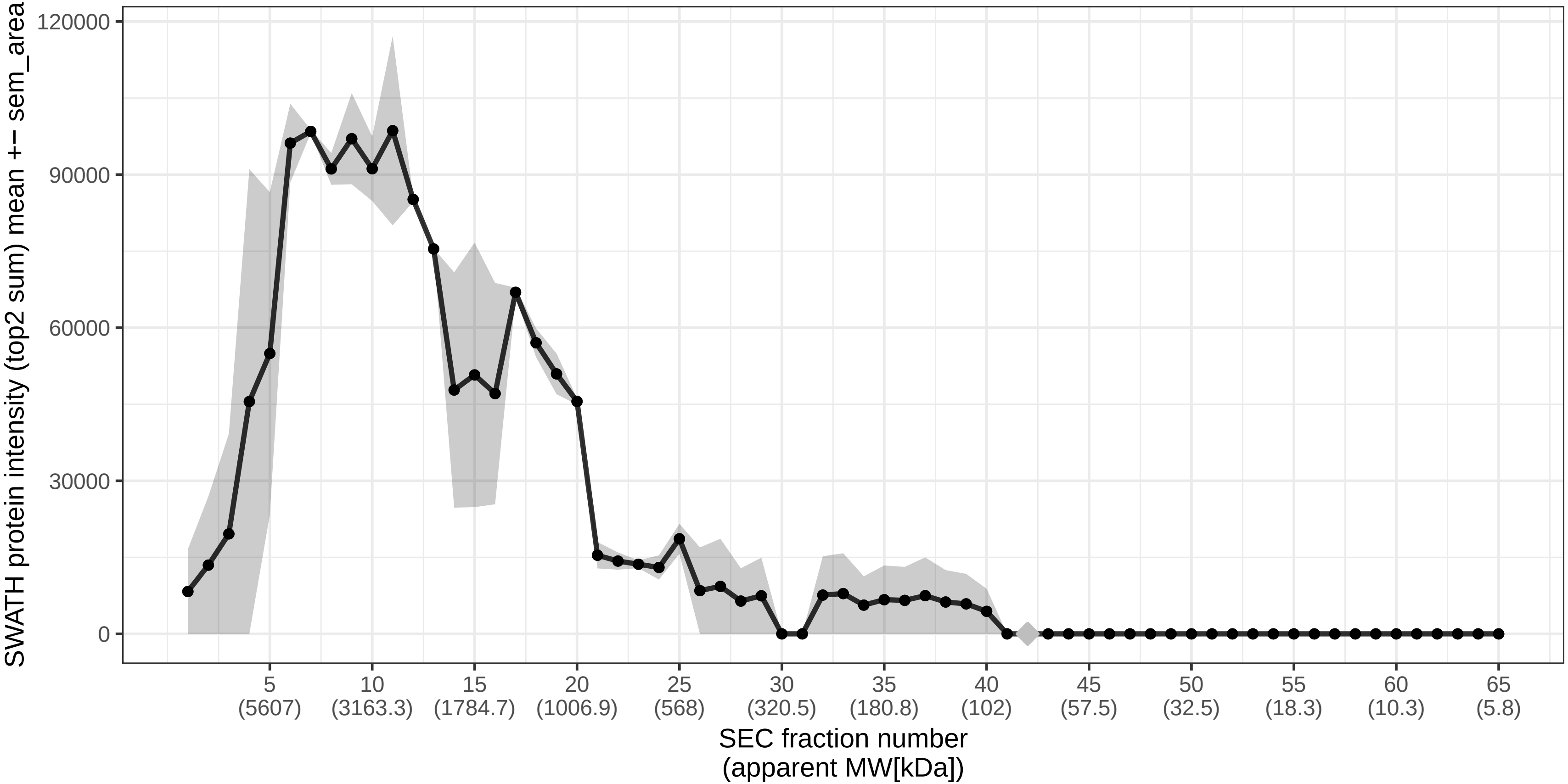

### SECchrom_O15230_LAMA5_HUMAN_LAMA5_KIAA0533_KIAA1907.pdf

O15230 | LAMA5_HUMAN | LAMA5 KIAA0533 KIAA1907
Monomer MW [kDa]: 399.737 Monomer expected elution fraction: 28

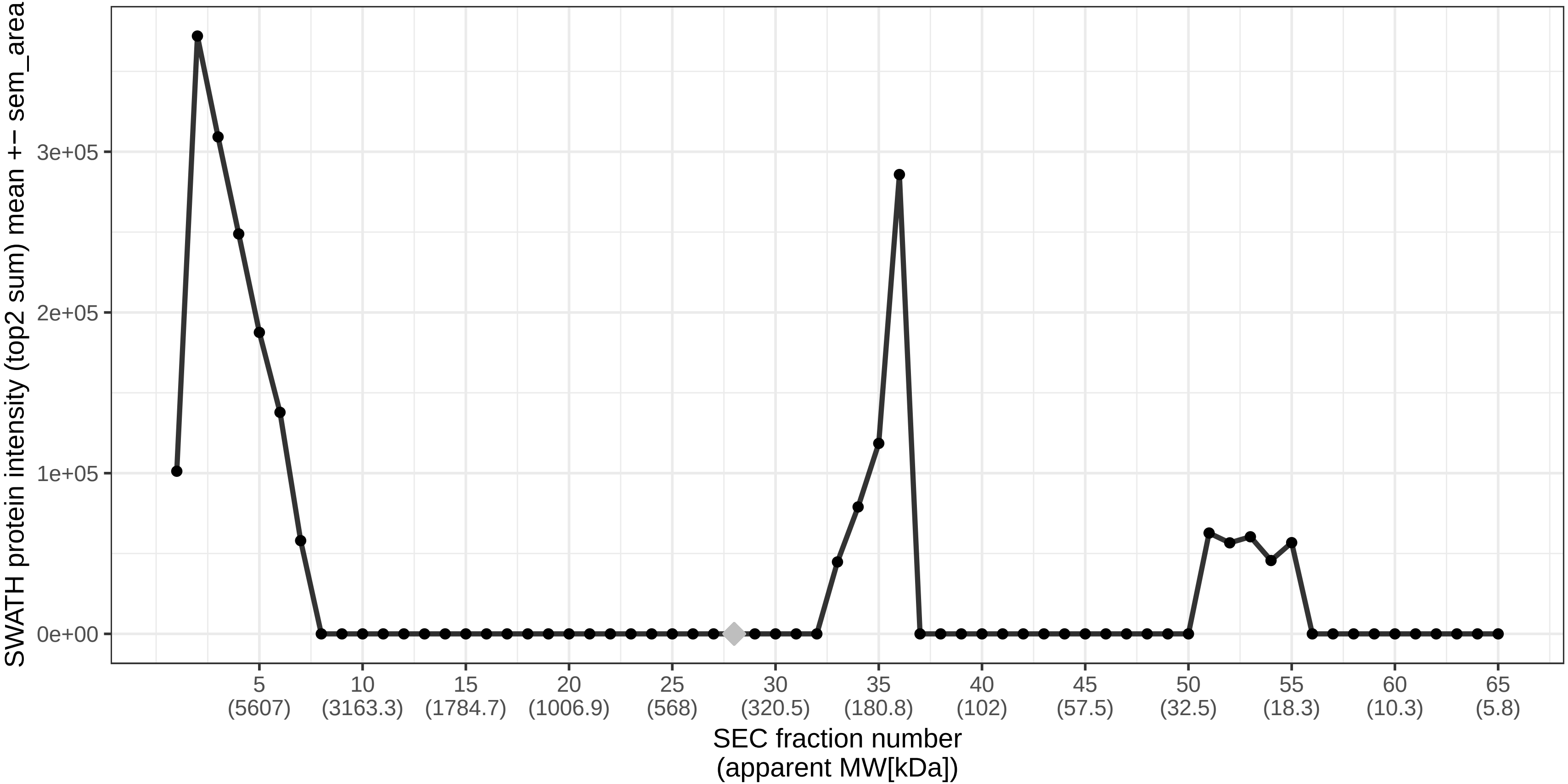

### SECchrom_O15234_CASC3_HUMAN_CASC3_MLN51.pdf

O15234 | CASC3_HUMAN | CASC3 MLN51
Monomer MW [kDa]: 76.278 Monomer expected elution fraction: 43

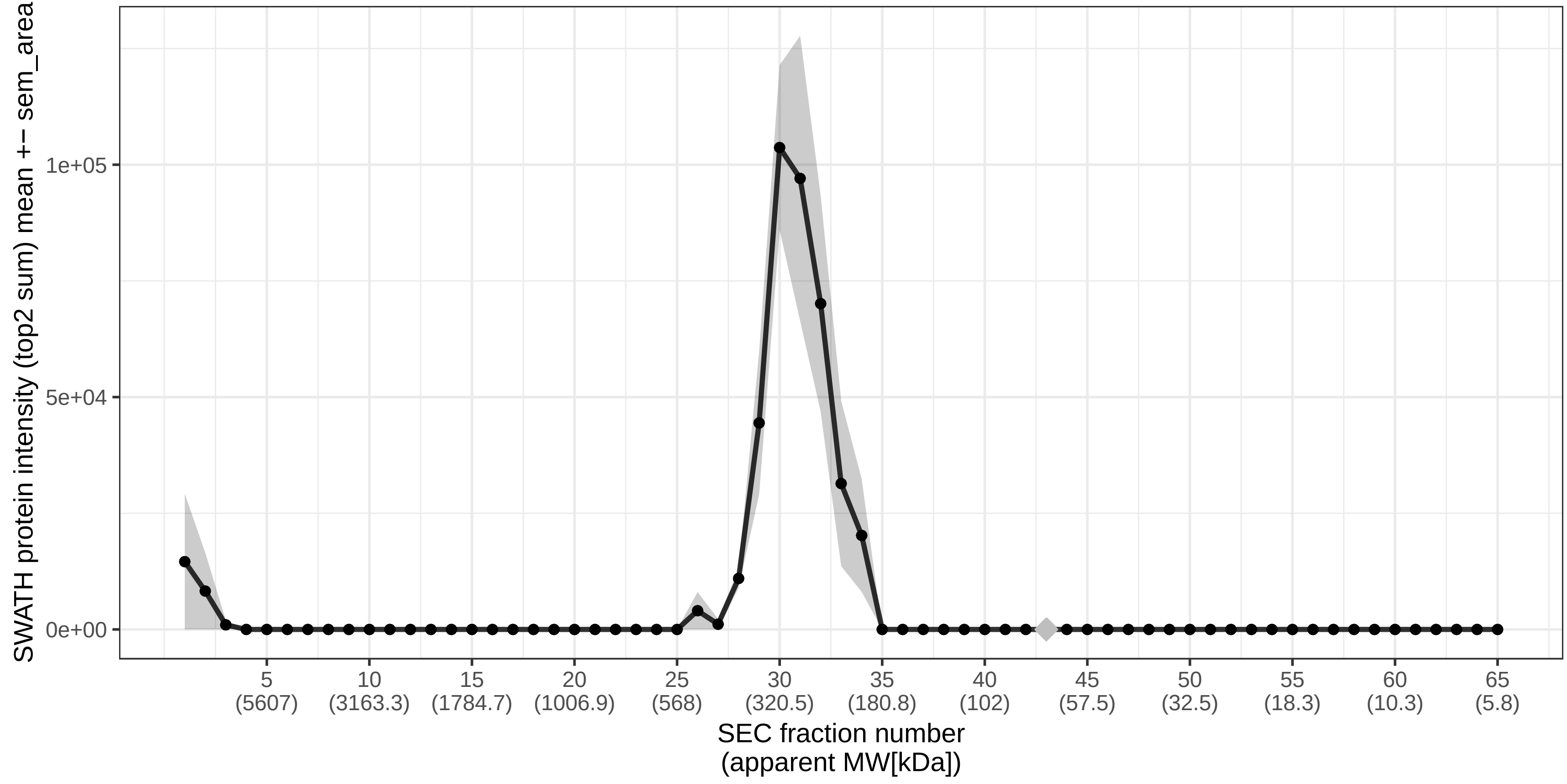

### SECchrom_O15254_ACOX3_HUMAN_ACOX3_BRCOX_PRCOX.pdf

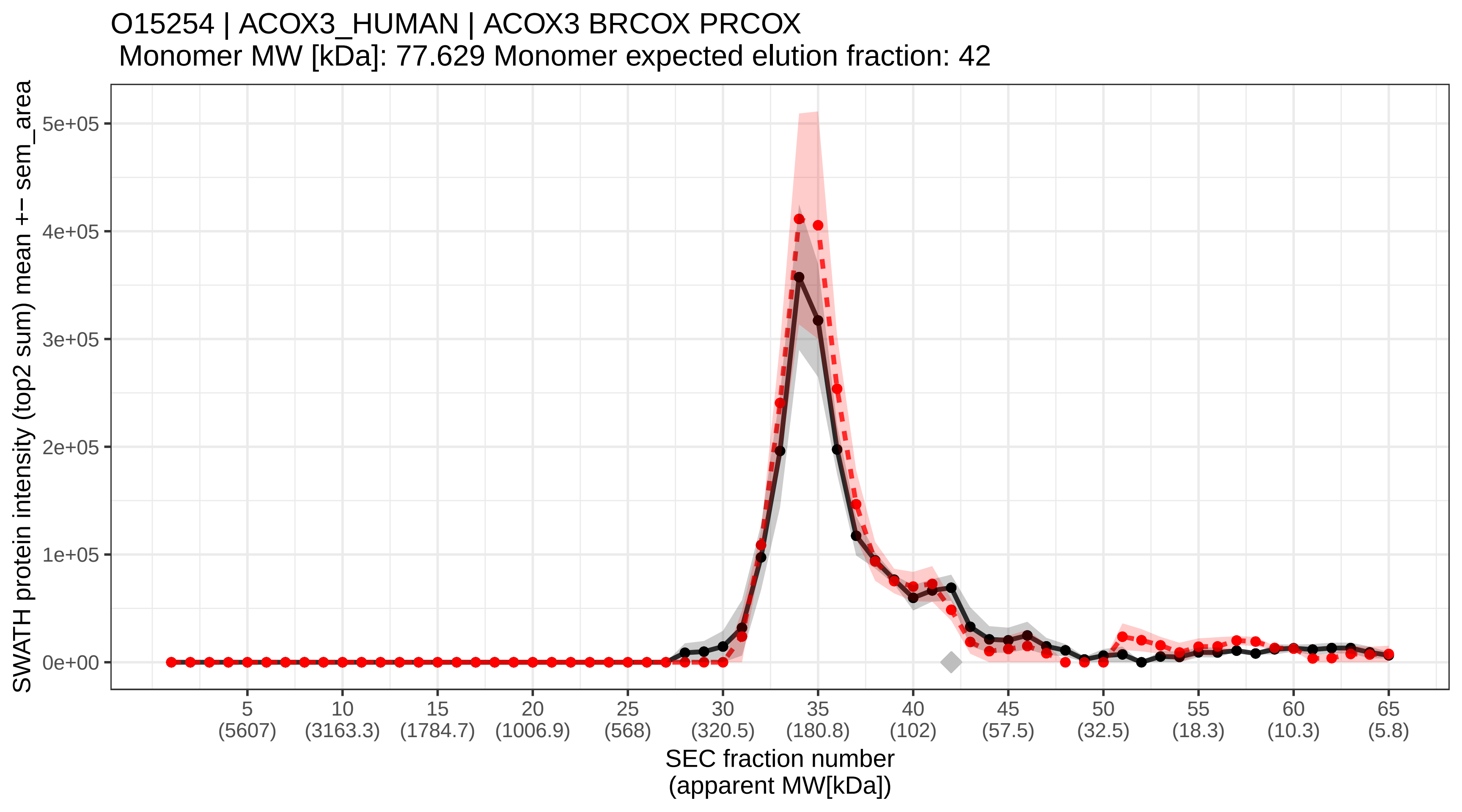

### SECchrom_O15258_RER1_HUMAN_RER1.pdf

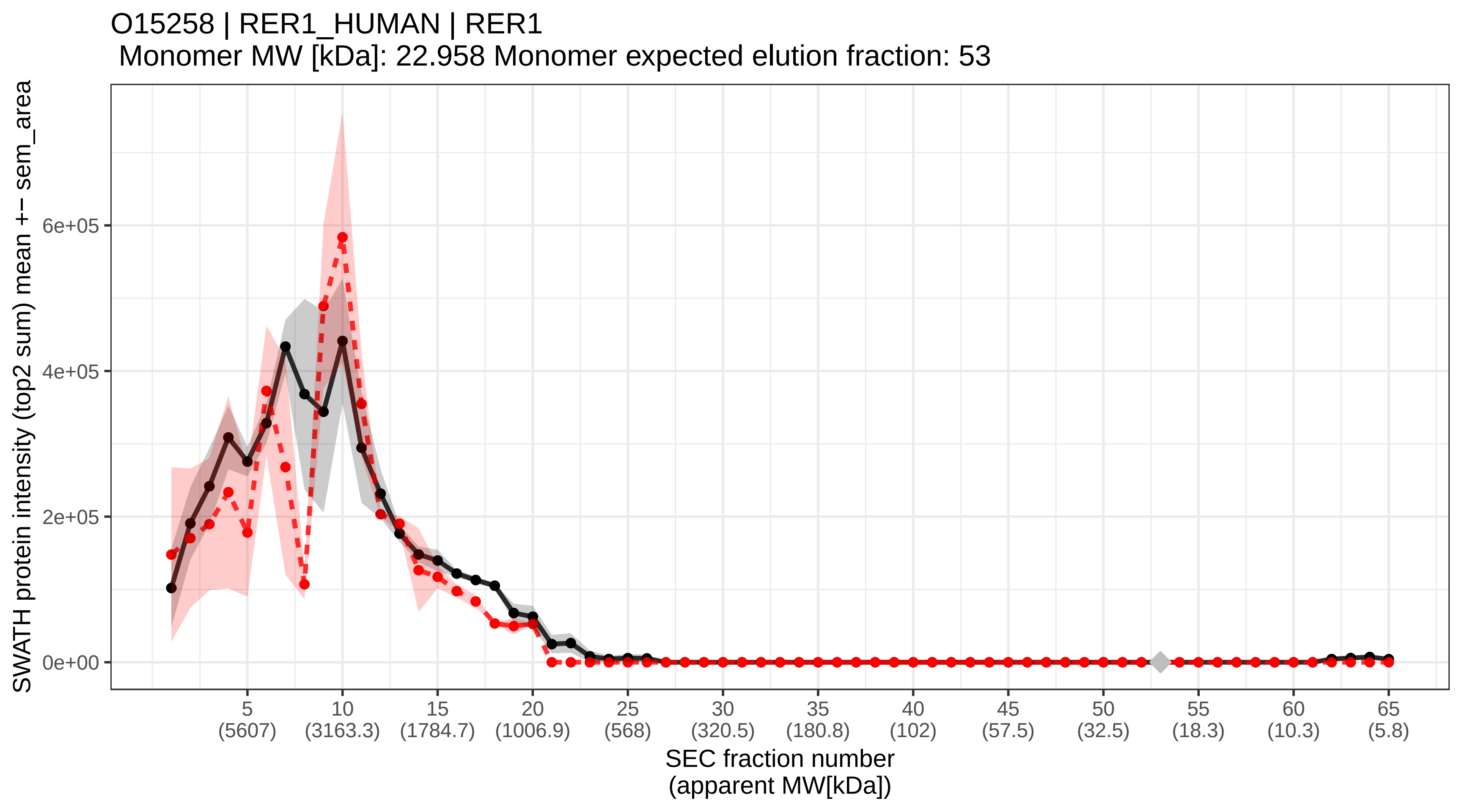

### SECchrom_O15260_SURF4_HUMAN_SURF4_SURF-4.pdf

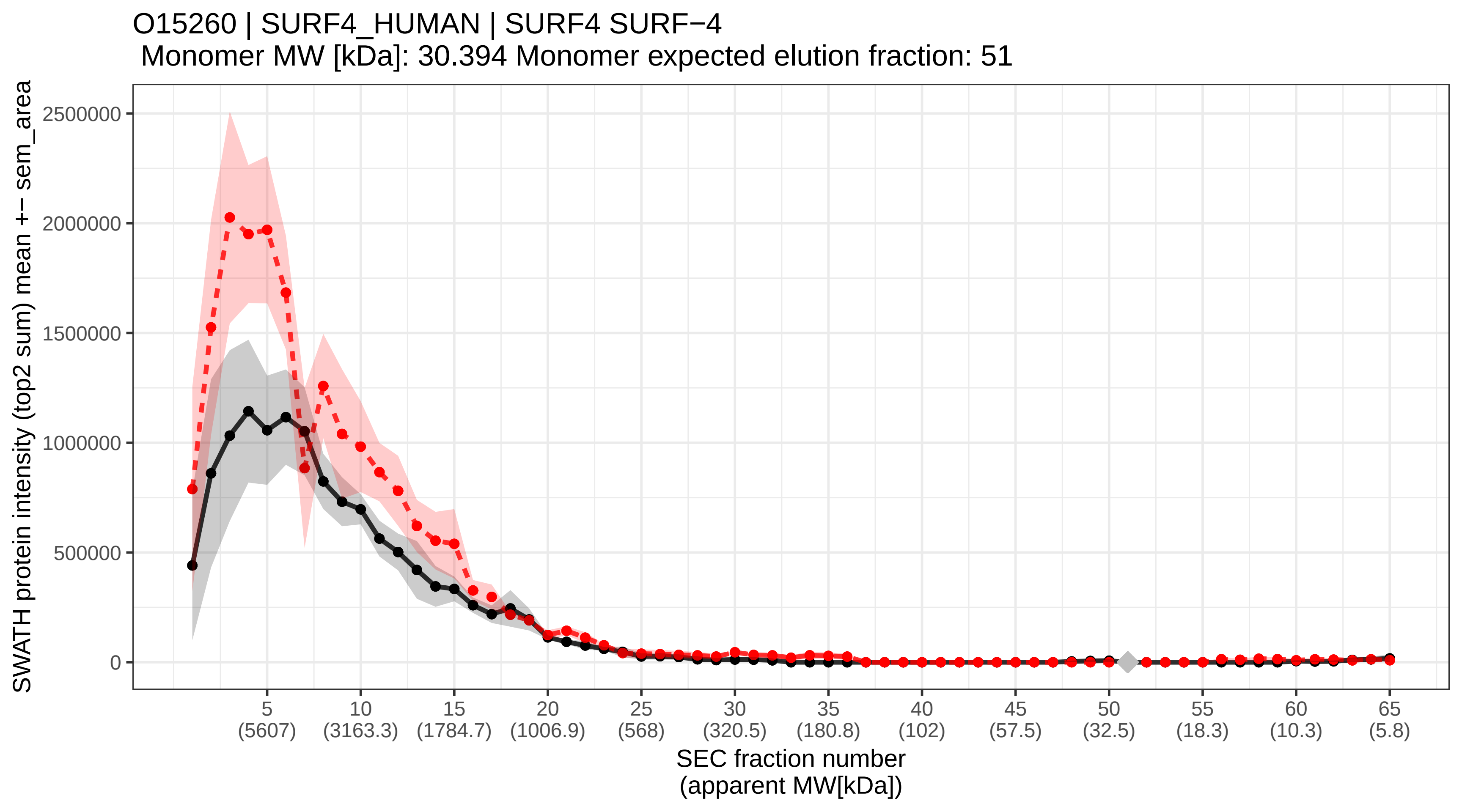

### SECchrom_O15269_SPTC1_HUMAN_SPTLC1_LCB1.pdf

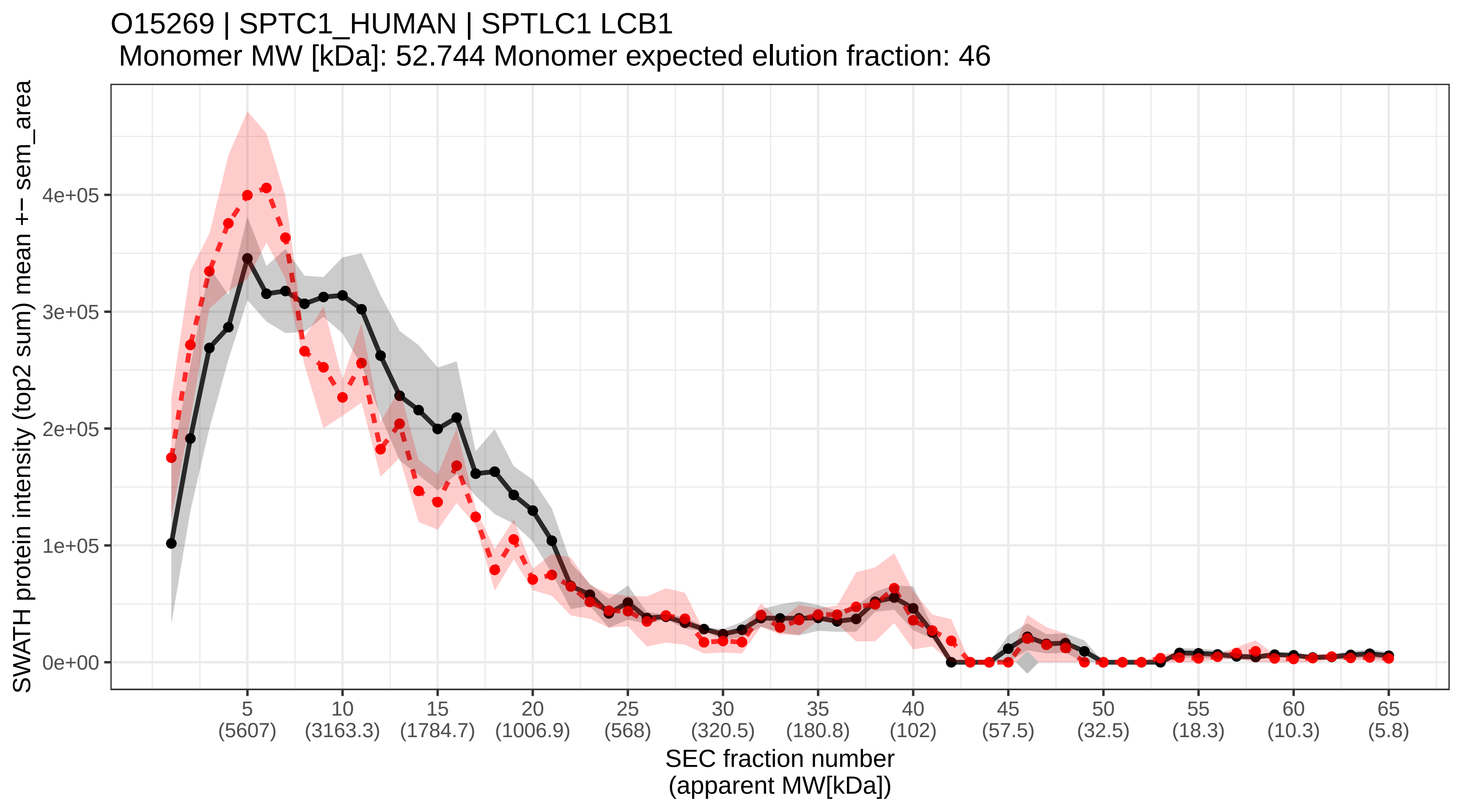

### SECchrom_O15270_SPTC2_HUMAN_SPTLC2_KIAA0526_LCB2.pdf

O15270 | SPTC2_HUMAN | SPTLC2 KIAA0526 LCB2
Monomer MW [kDa]: 62.924 Monomer expected elution fraction: 44

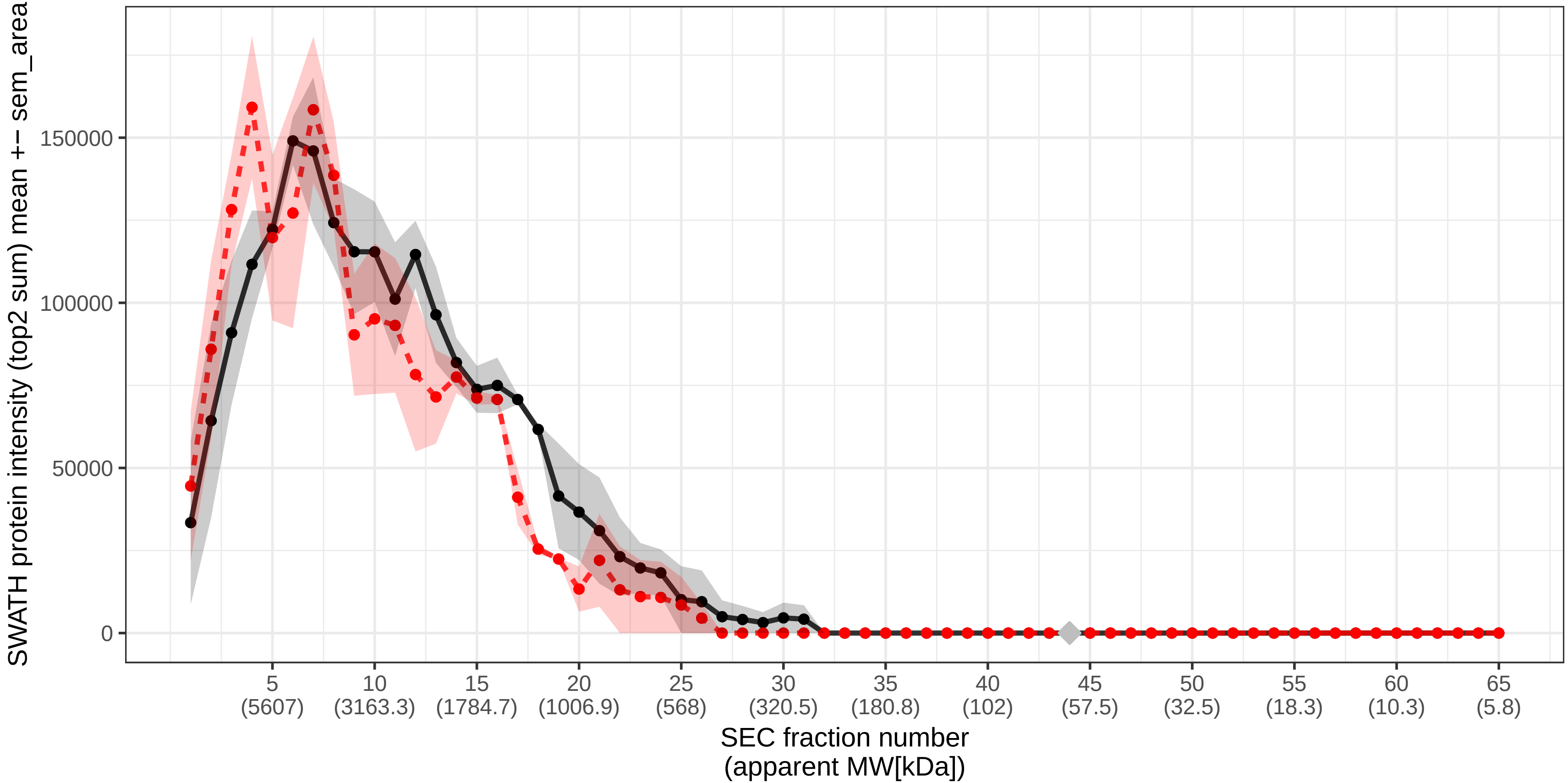

### SECchrom_O15294_OGT1_HUMAN_OGT.pdf

O15294 | OGT1_HUMAN | OGT
Monomer MW [kDa]: 116.925 Monomer expected elution fraction: 39

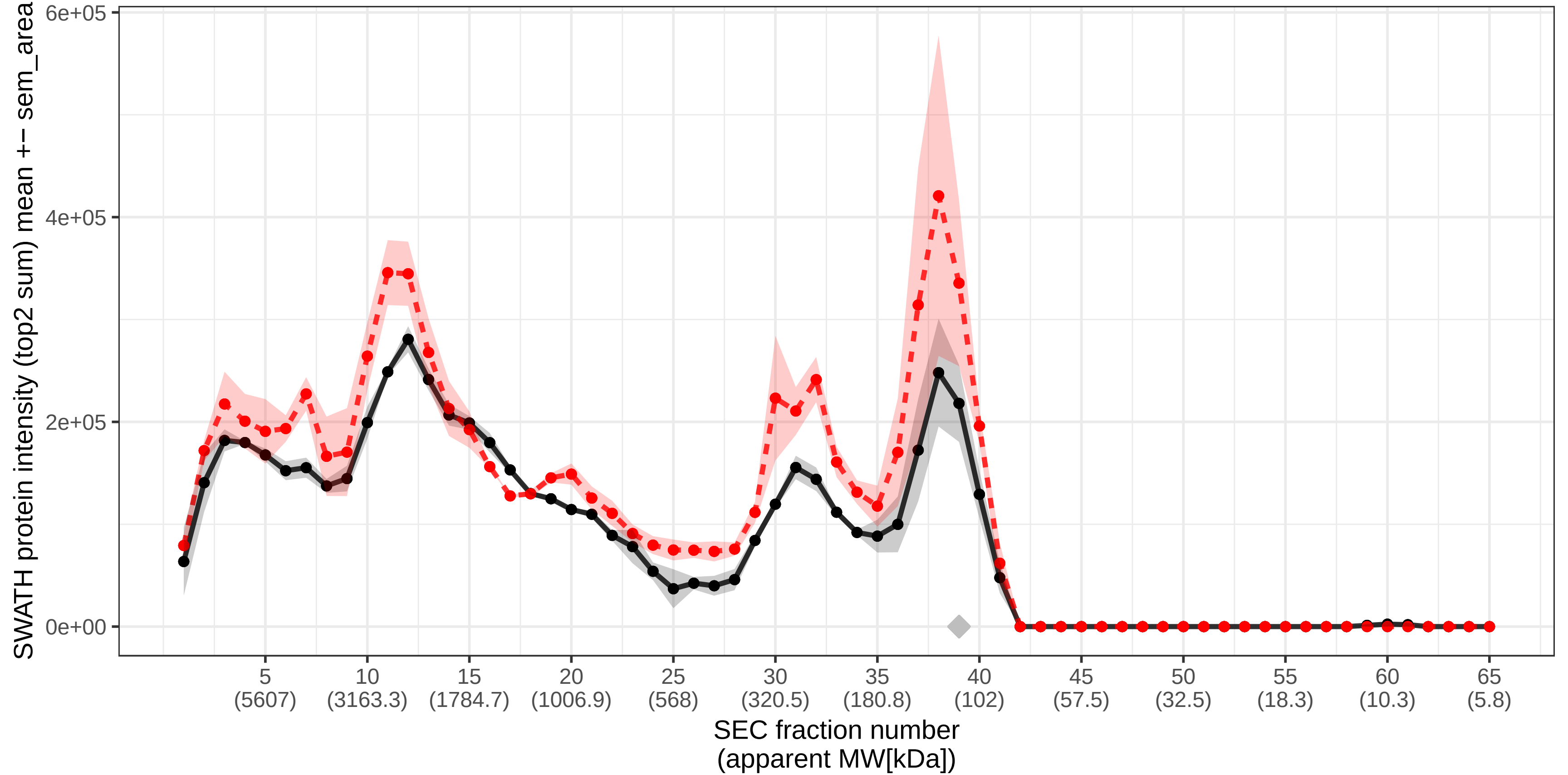

### SECchrom_O15305_PMM2_HUMAN_PMM2.pdf

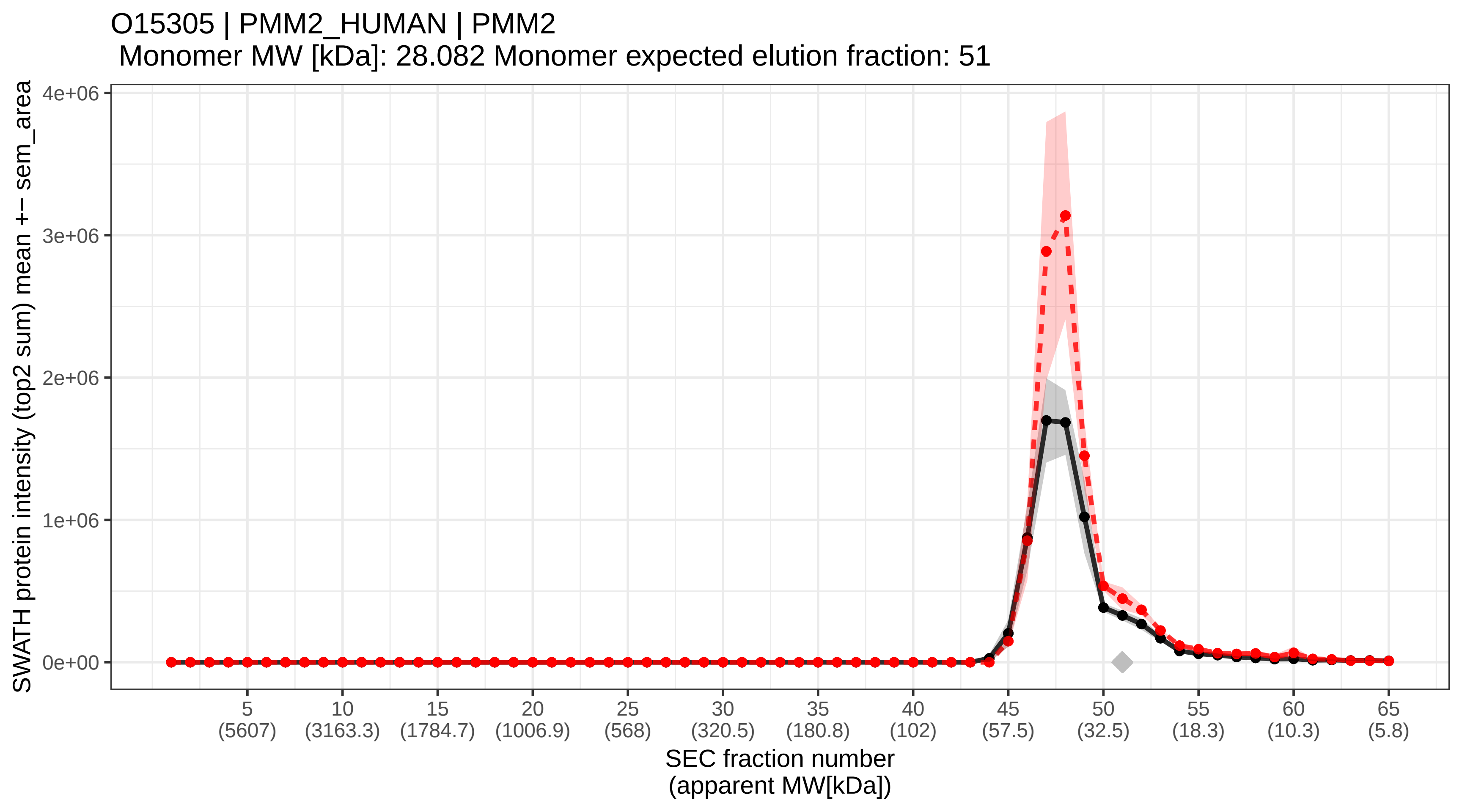

### SECchrom_O15318_RPC7_HUMAN_POLR3G.pdf

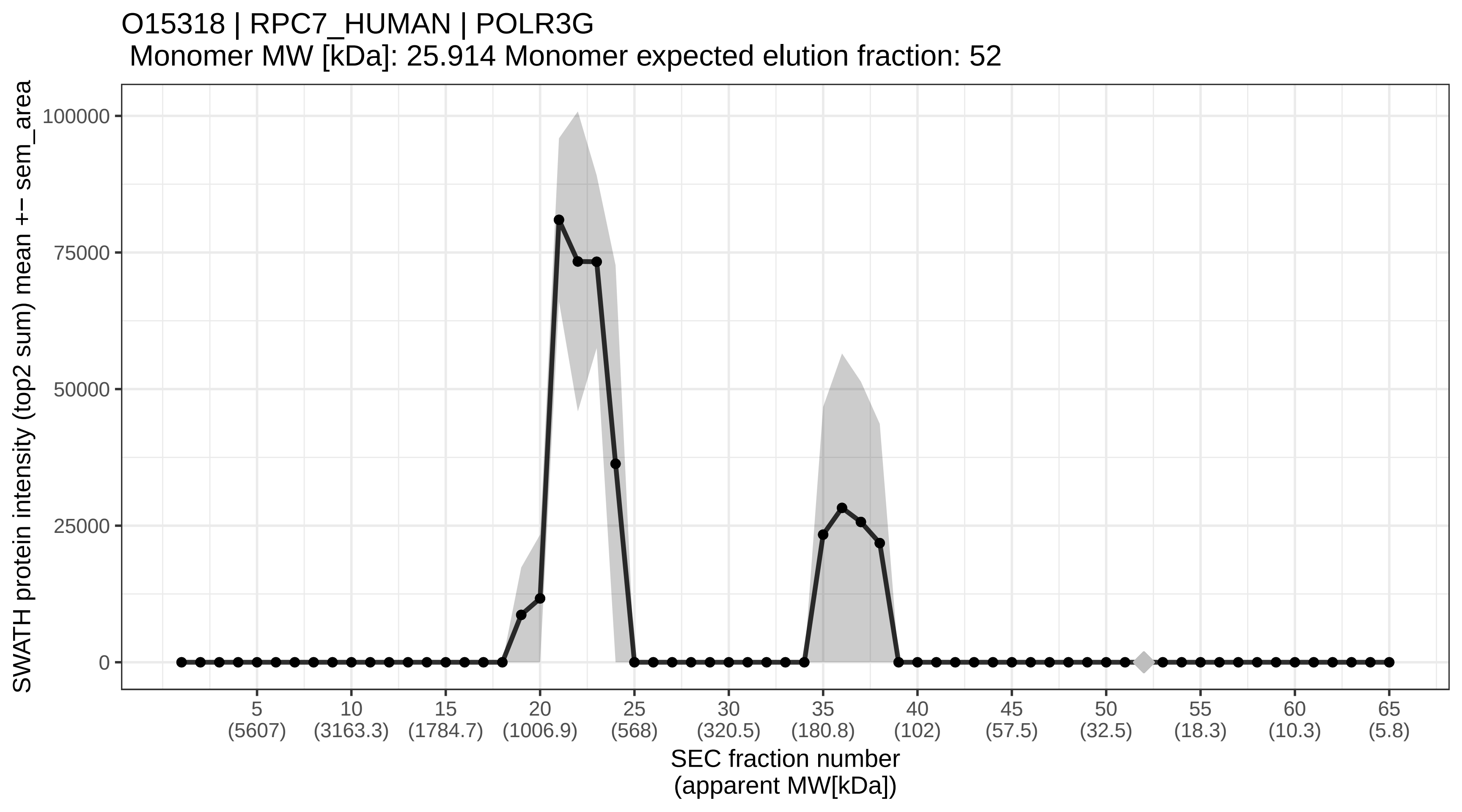

### SECchrom_O15321_TM9S1_HUMAN_TM9SF1.pdf

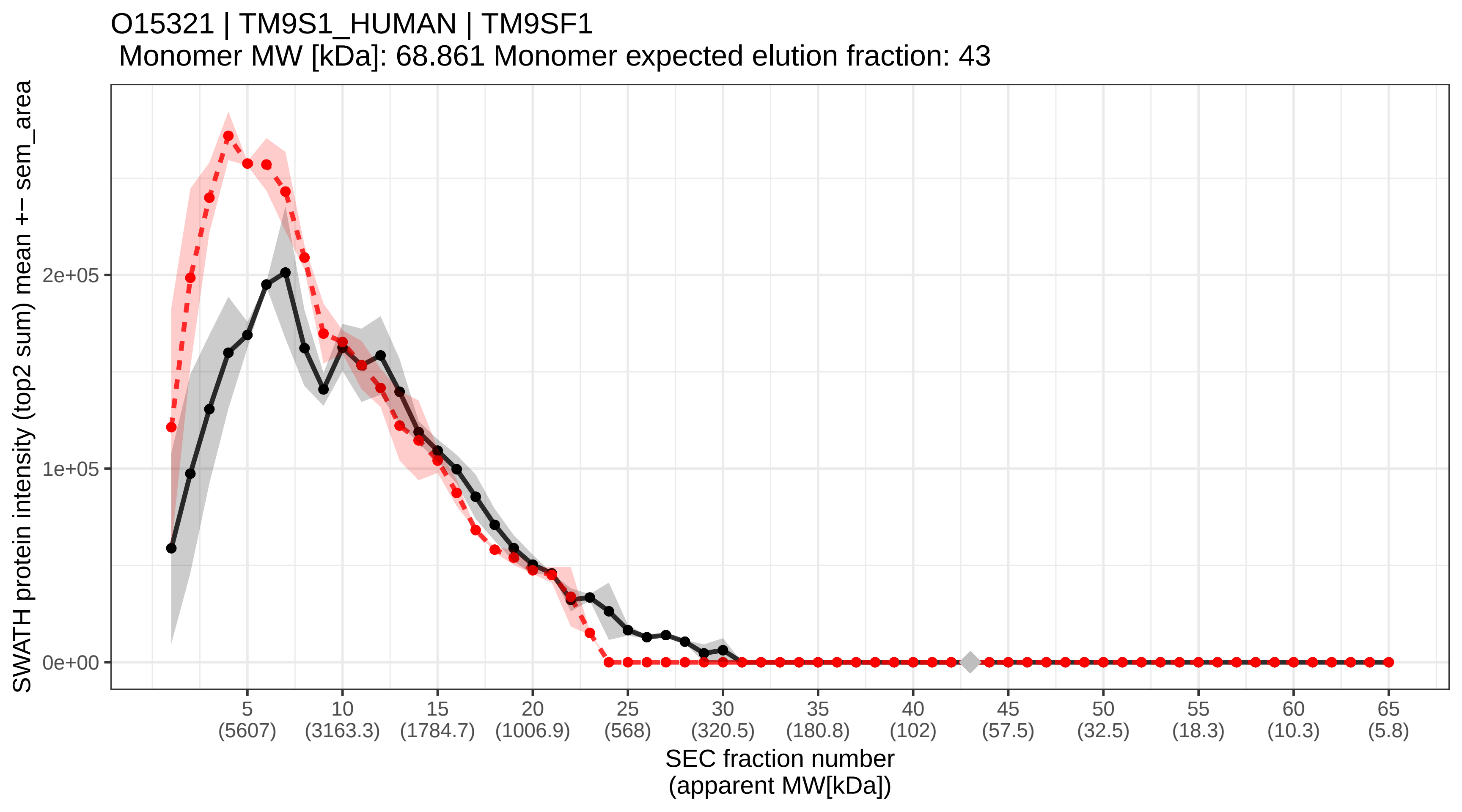

### SECchrom_O15327_INP4B_HUMAN_INPP4B.pdf

O15327 | INP4B_HUMAN | INPP4B
Monomer MW [kDa]: 104.738 Monomer expected elution fraction: 40

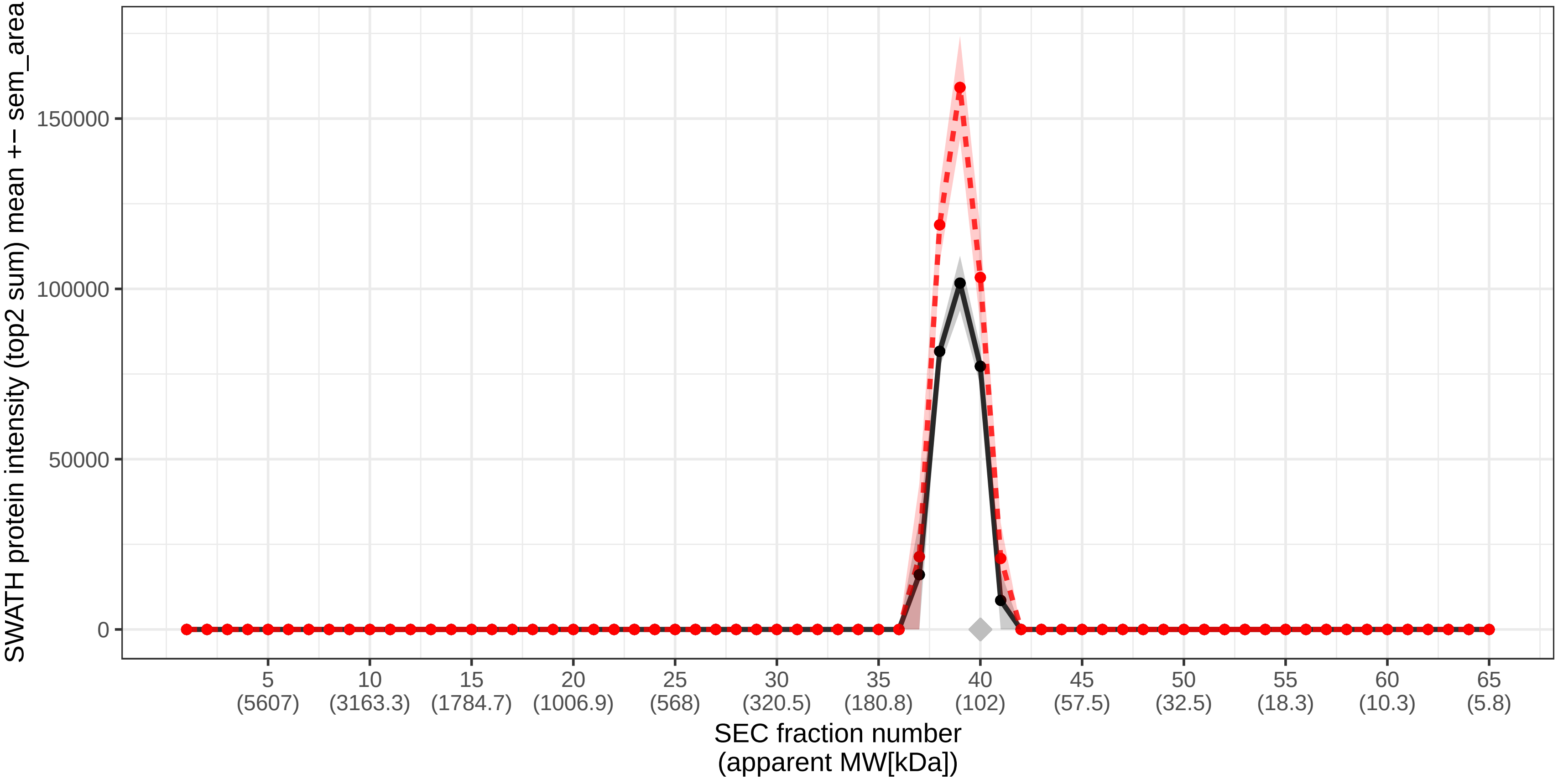

### SECchrom_O15344_TRI18_HUMAN_MID1_FXY_RNF59_TRIM18_XPRF.pdf

O15344 | TRI18_HUMAN | MID1 FXY RNF59 TRIM18 XPRF
Monomer MW [kDa]: 75.251 Monomer expected elution fraction: 43

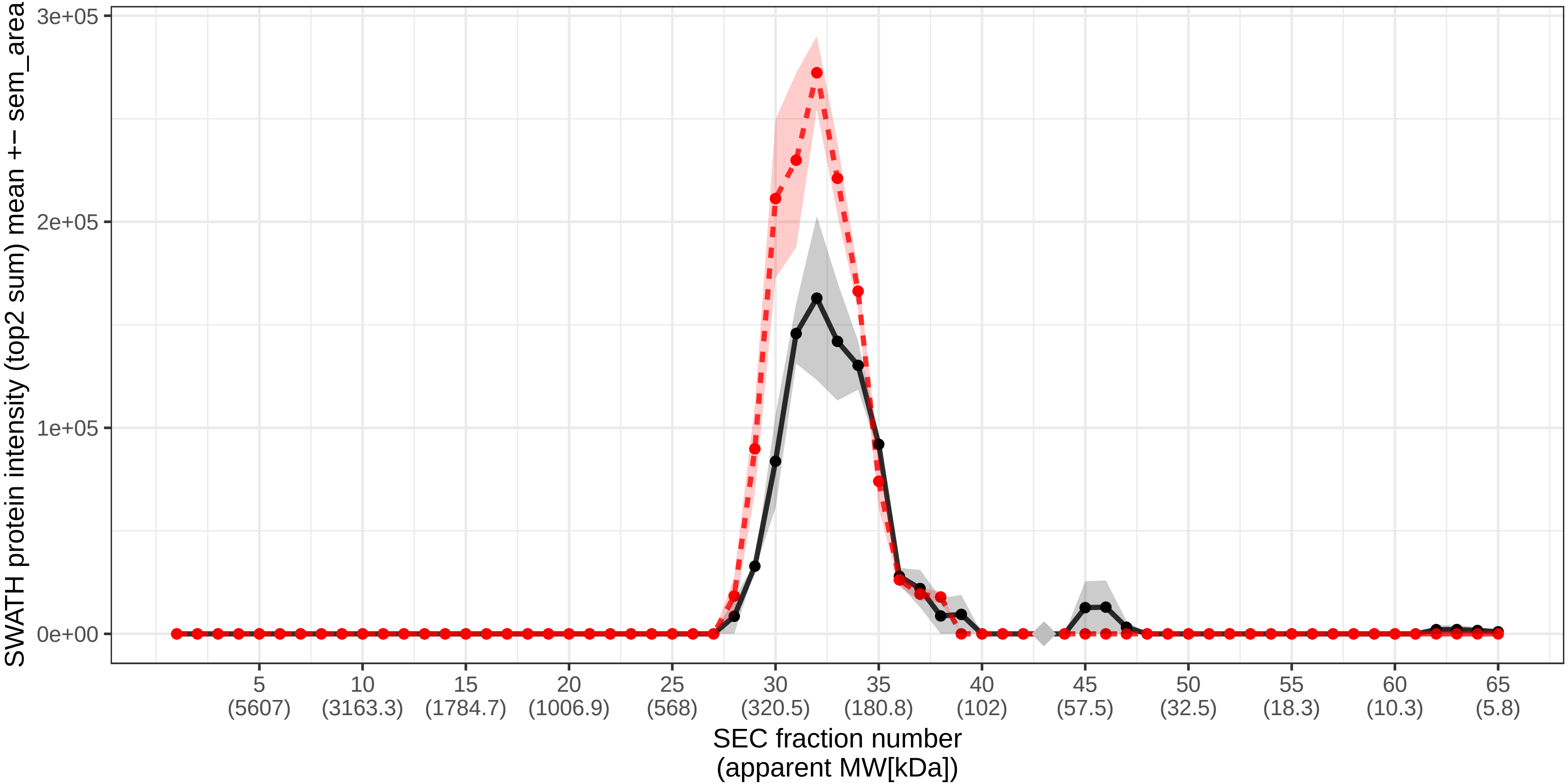

### SECchrom_O15347_HMGB3_HUMAN_HMGB3_HMG2A_HMG4.pdf

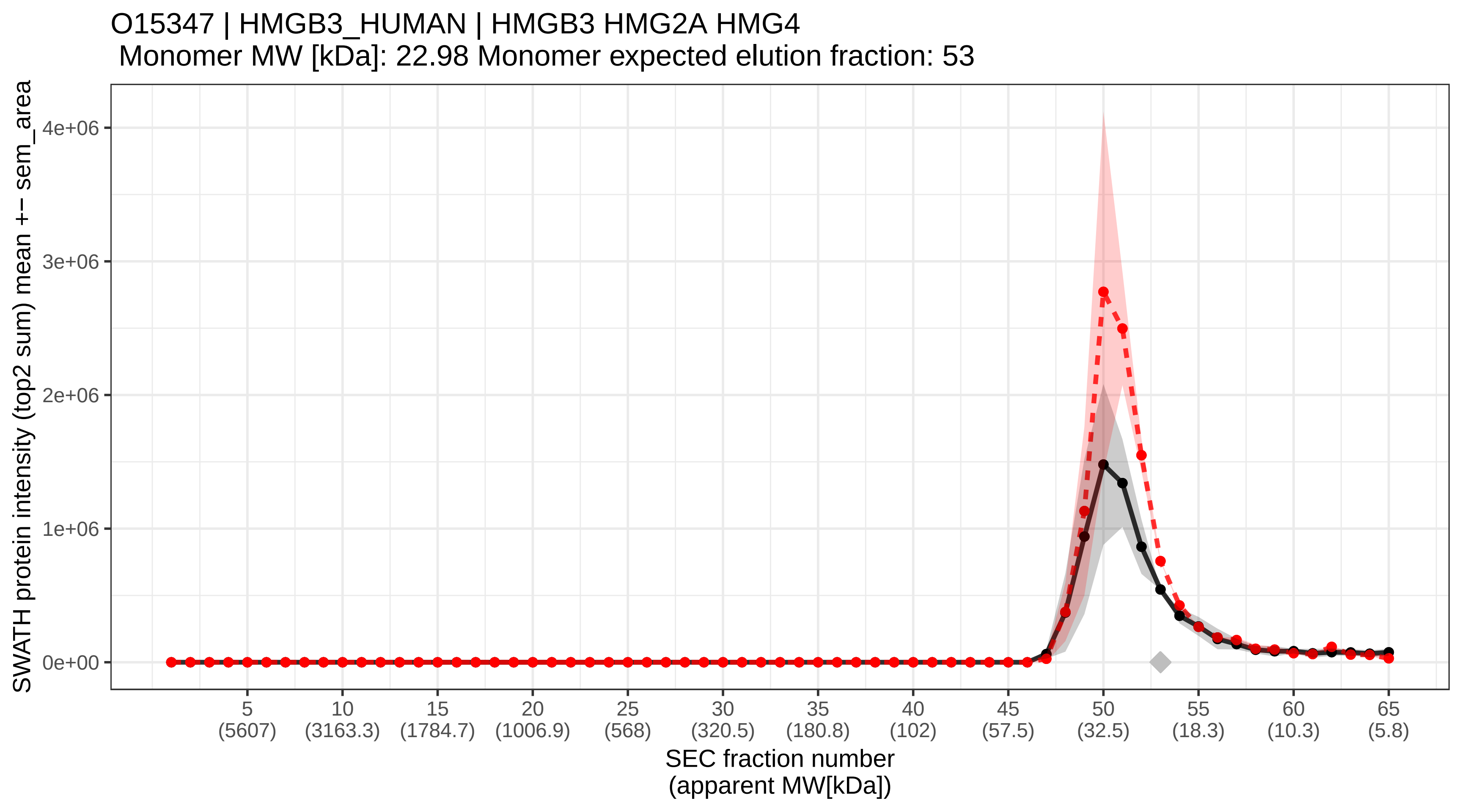

### SECchrom_O15355_PPM1G_HUMAN_PPM1G_PPM1C.pdf

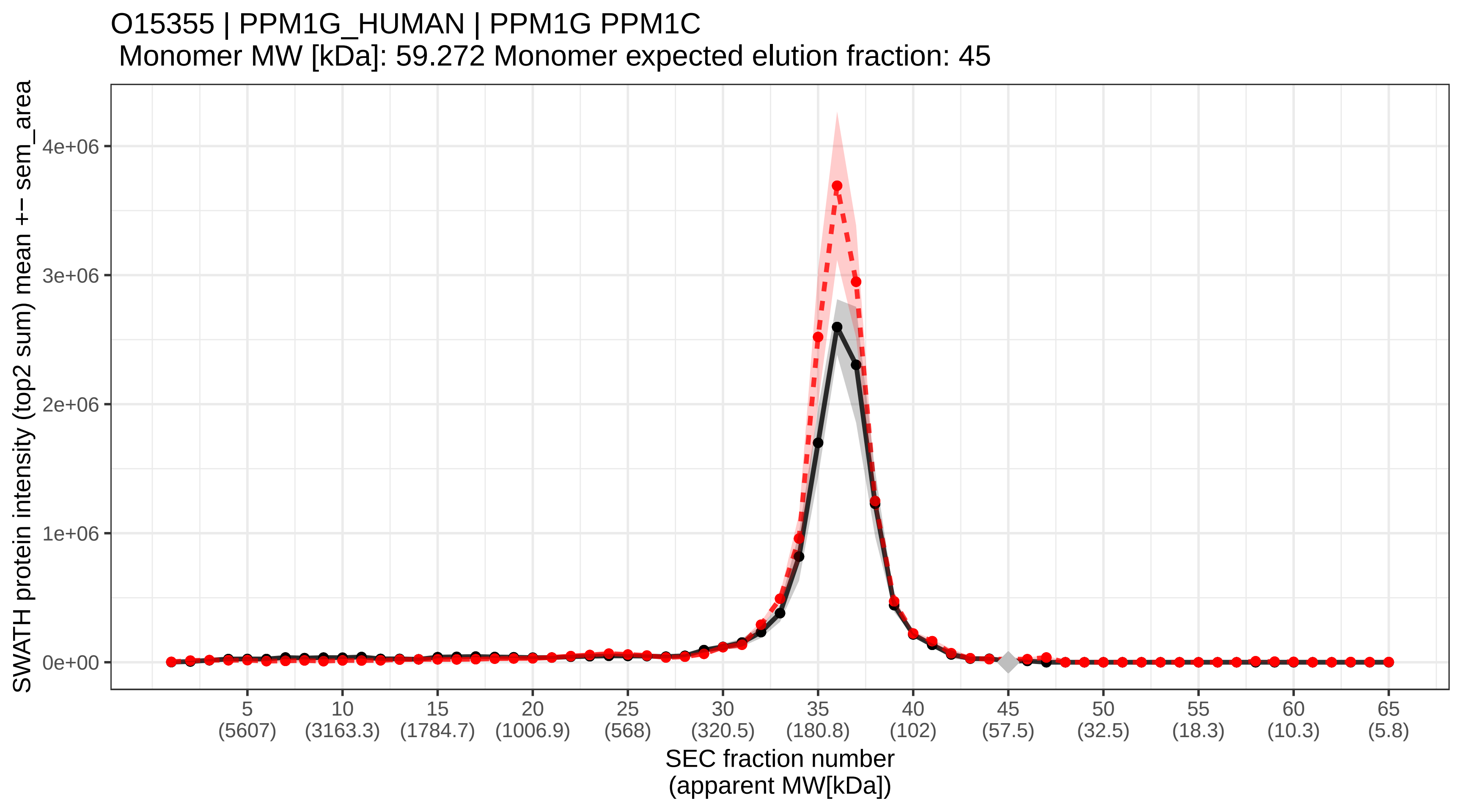

### SECchrom_O15357_SHIP2_HUMAN_INPPL1_SHIP2.pdf

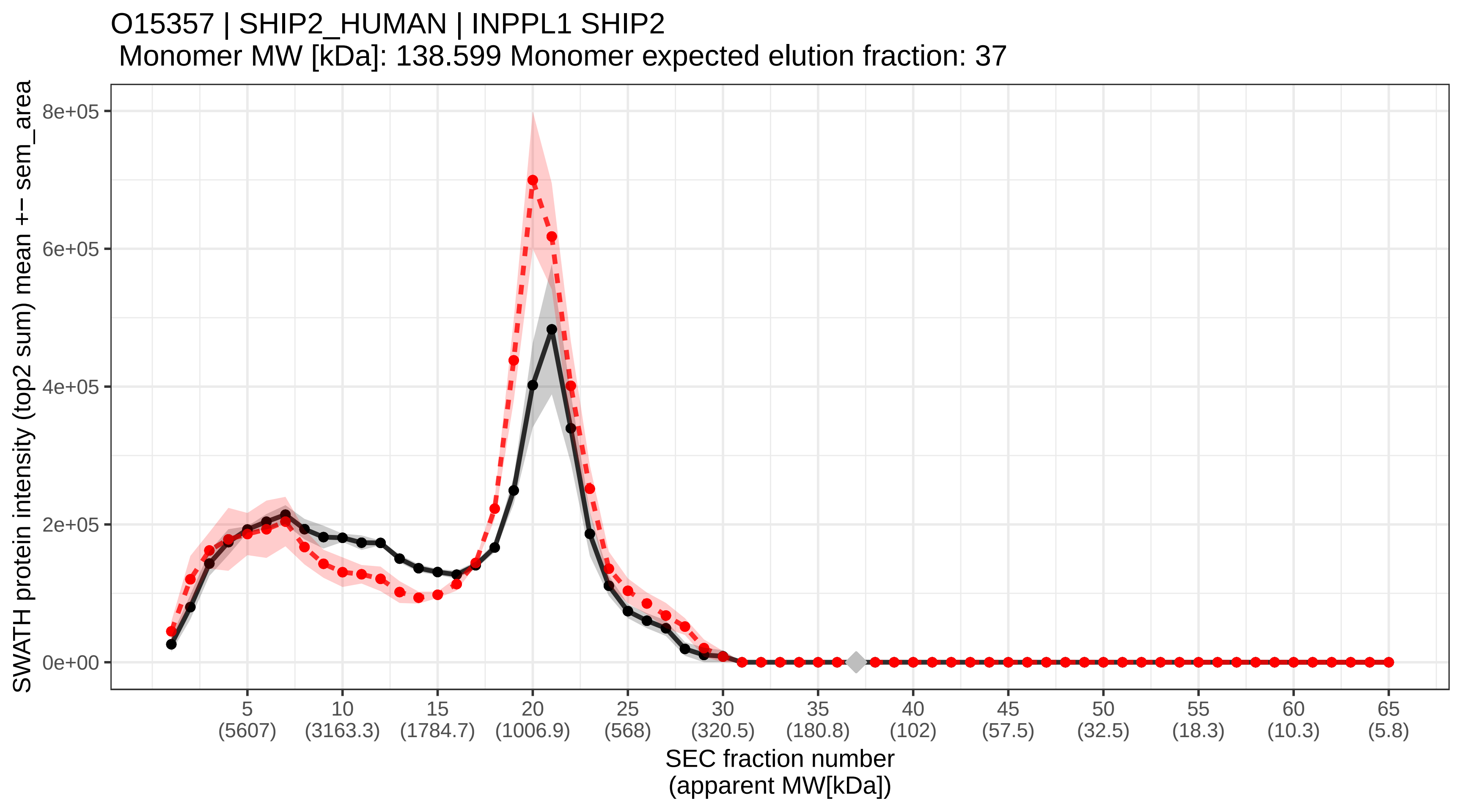
