## Supplementary figures and images for "A global screen for assembly state changes of the mitotic proteome by SEC-SWATH-MS"

### SECchrom_A0AVT1_UBA6_HUMAN_UBA6_MOP4_UBE1L2.pdf

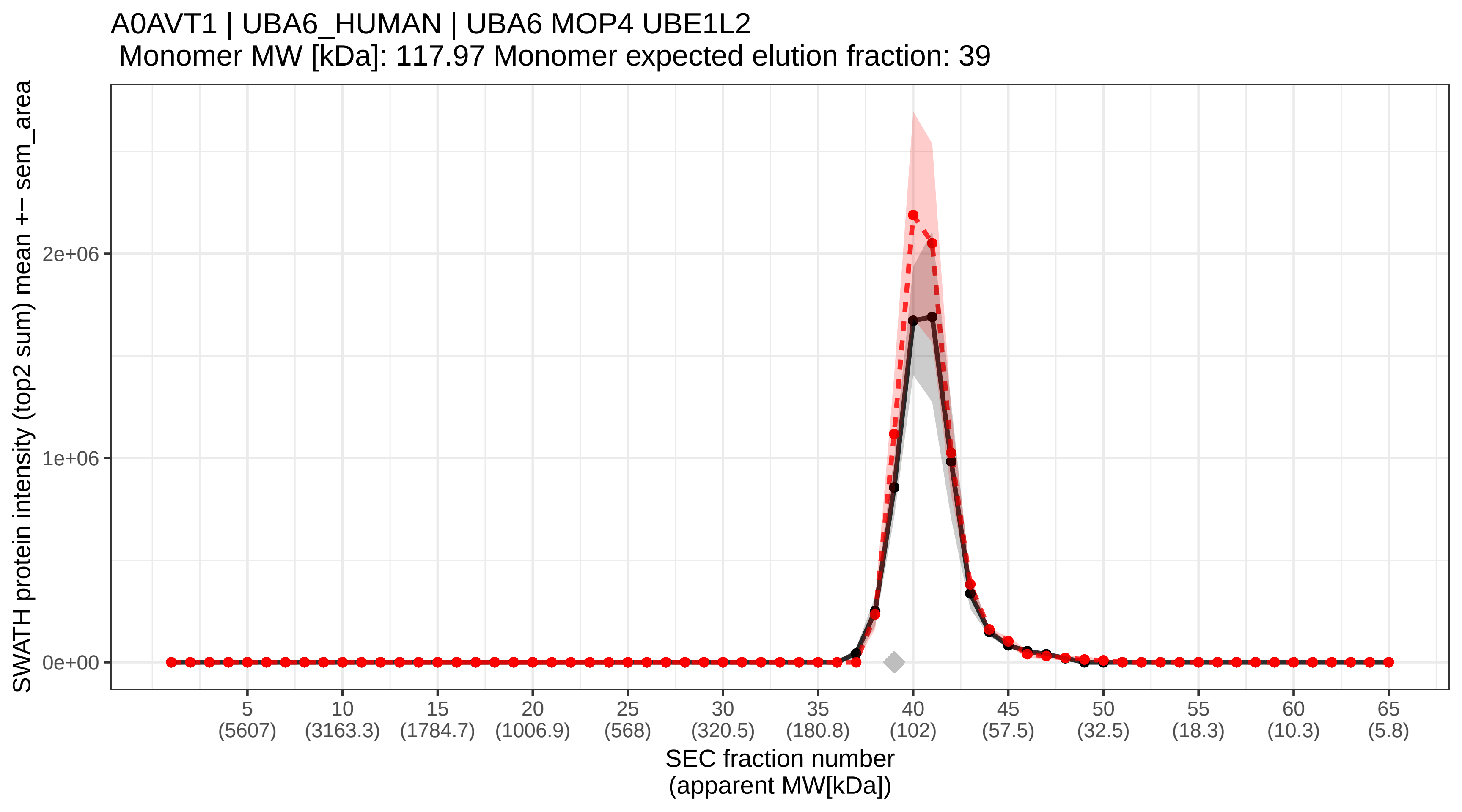

### SECchrom_A0FGR8_ESYT2_HUMAN_ESYT2_FAM62B_KIAA1228.pdf

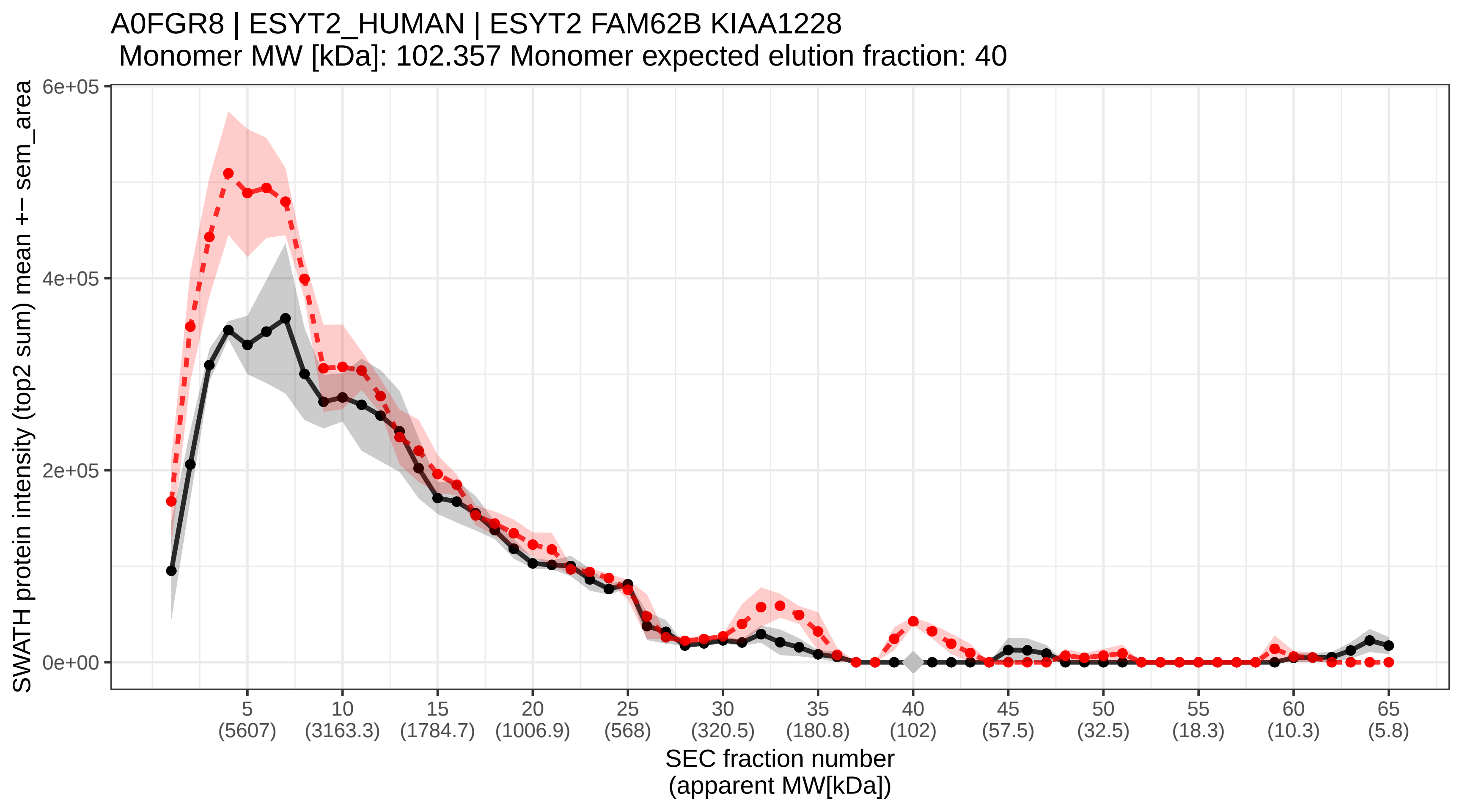

### SECchrom_A0MZ66_SHOT1_HUMAN_SHTN1_KIAA1598.pdf

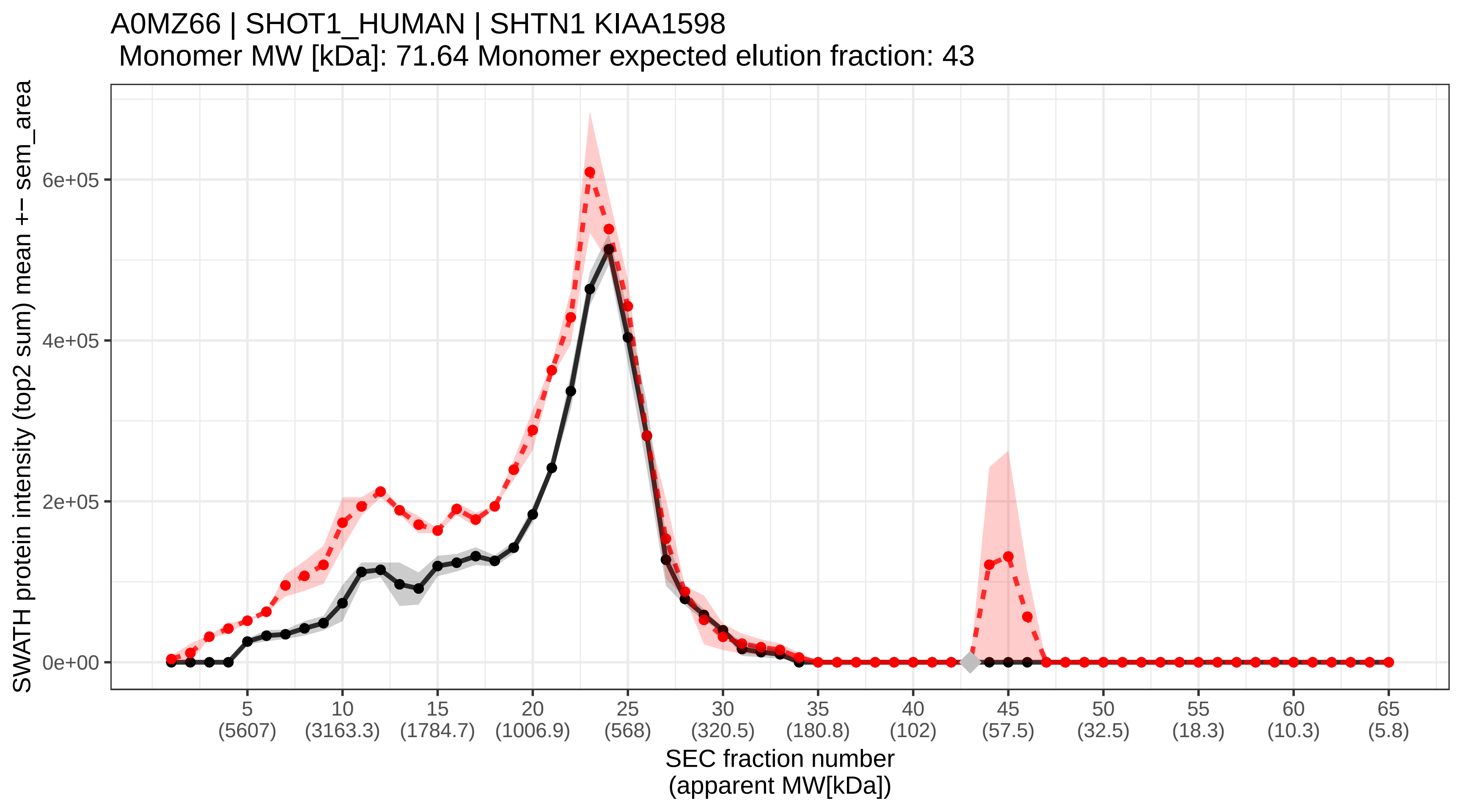

### SECchrom_A0PJW6_TM223_HUMAN_TMEM223.pdf

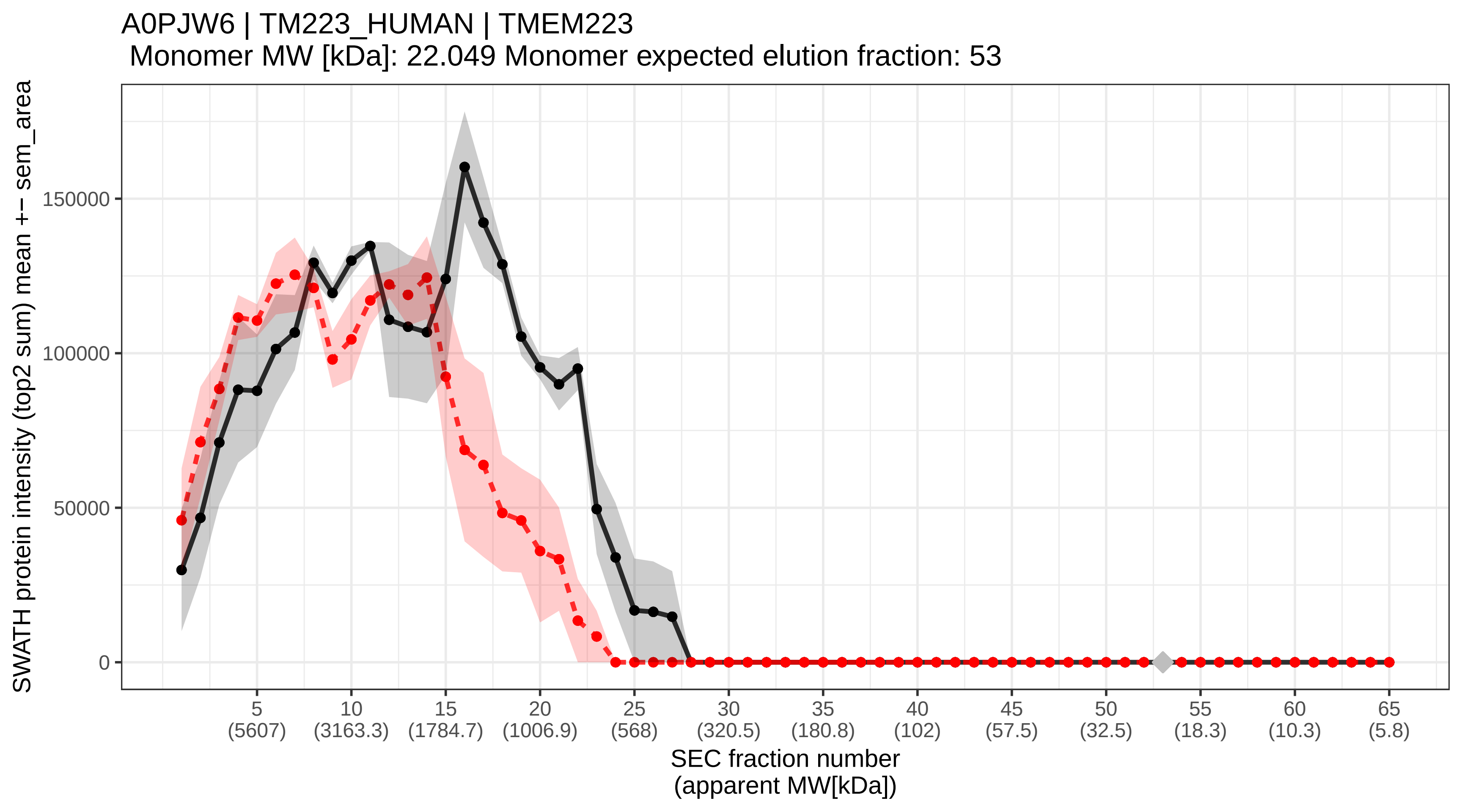

### SECchrom_A1L0T0_ILVBL_HUMAN_ILVBL_AHAS.pdf

A1L0T0 | ILVBL_HUMAN | ILVBL AHAS
Monomer MW [kDa]: 67.868 Monomer expected elution fraction: 44

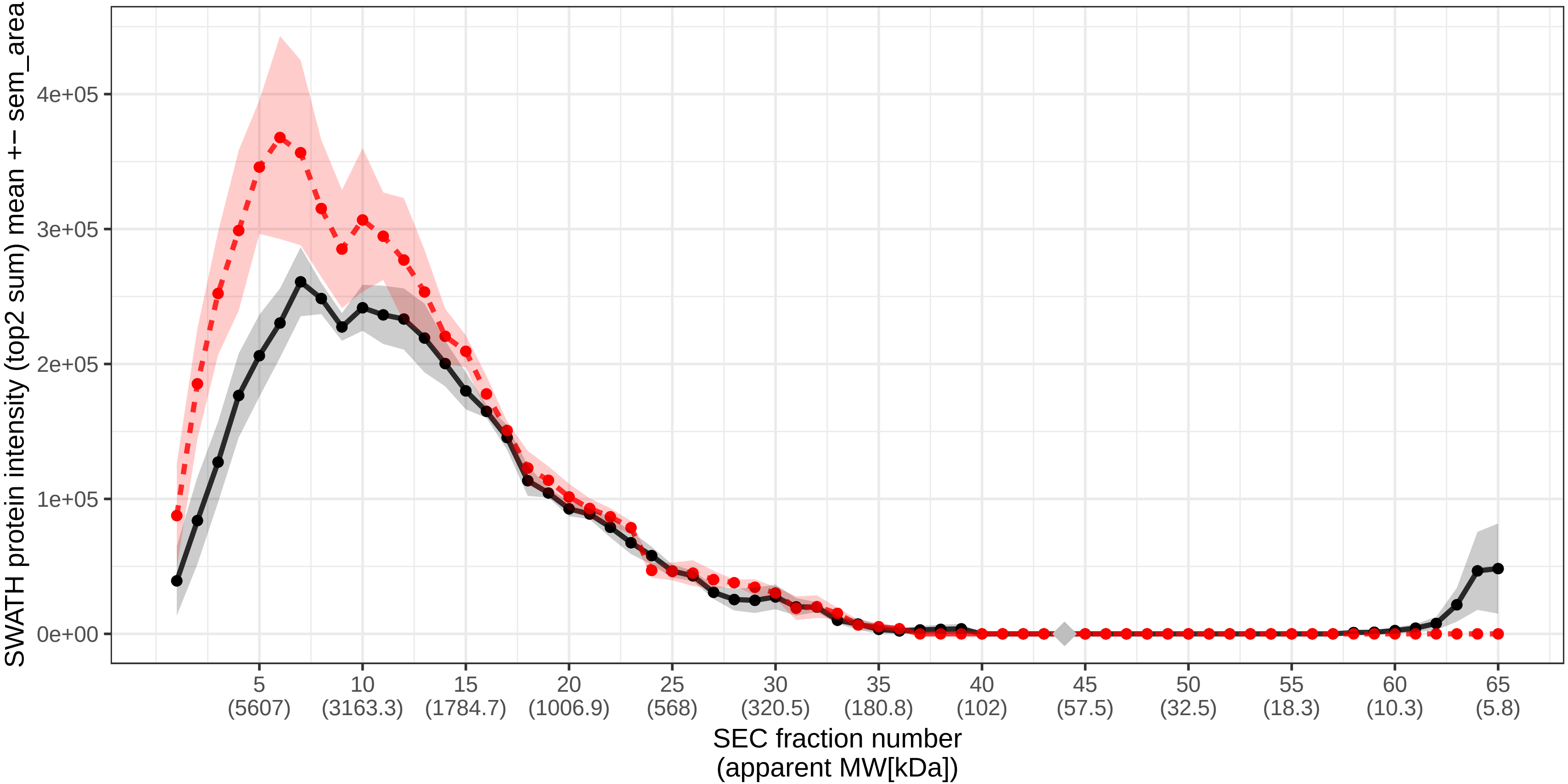

### SECchrom_A1X283_SPD2B_HUMAN_SH3PXD2B_FAD49_KIAA1295_TKS4.pdf

A1X283 | SPD2B_HUMAN | SH3PXD2B FAD49 KIAA1295 TKS4
Monomer MW [kDa]: 101.579 Monomer expected elution fraction: 40

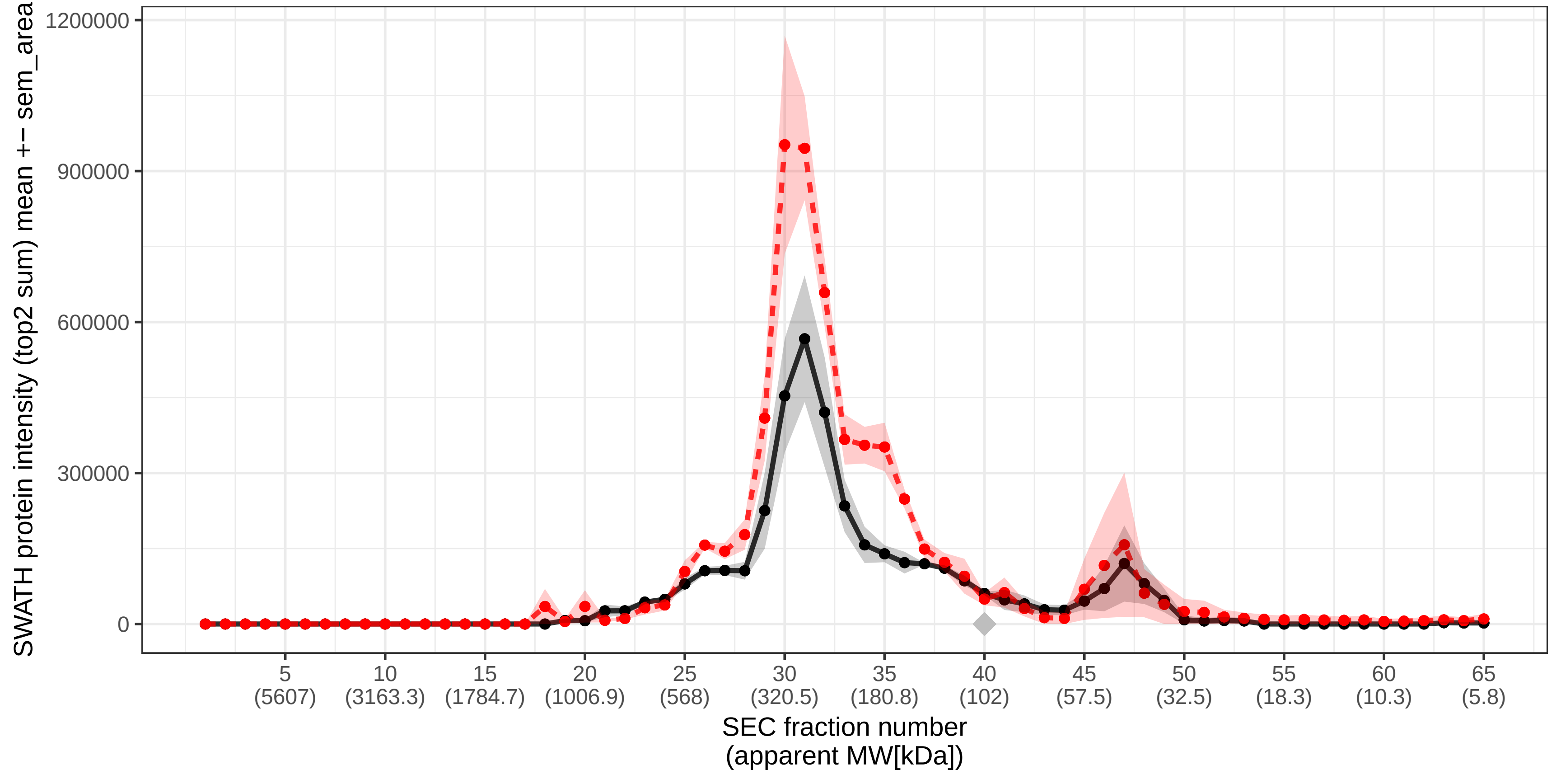

### SECchrom_A2RRP1_NBAS_HUMAN_NBAS_NAG.pdf

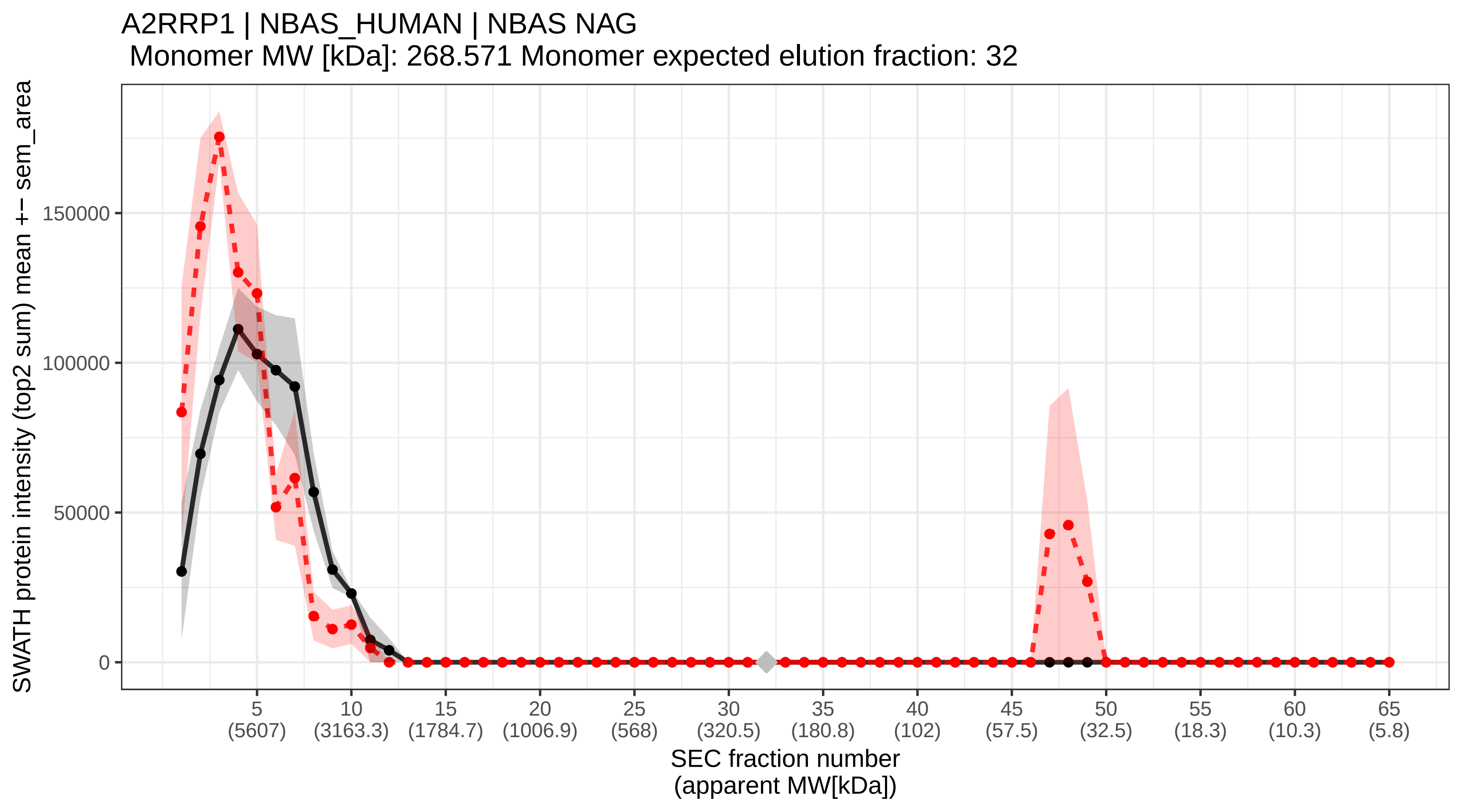

### SECchrom_A2RU67_F234B_HUMAN_FAM234B_KIAA1467.pdf

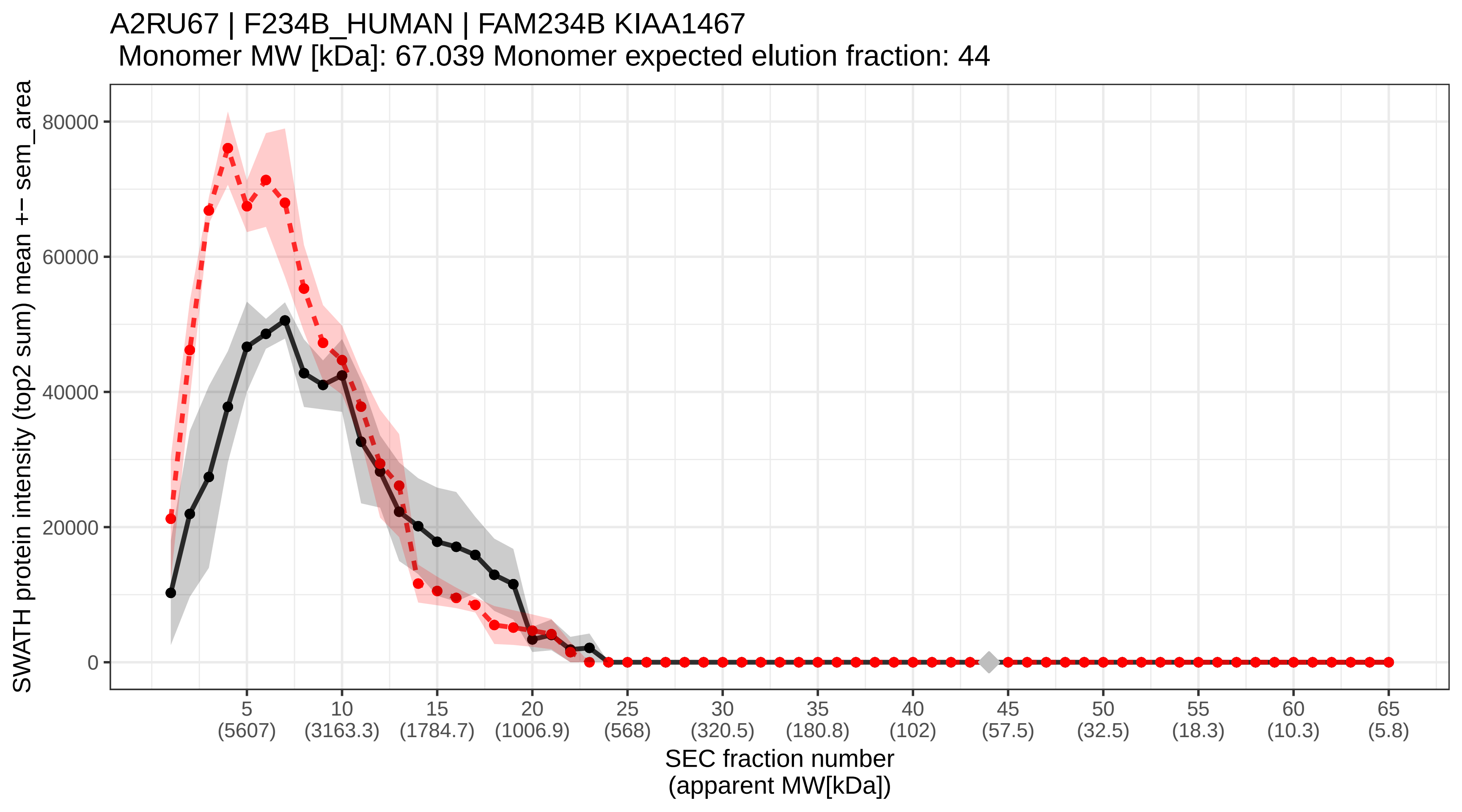

### SECchrom_A2RUC4_TYW5_HUMAN_TYW5_C2orf60.pdf

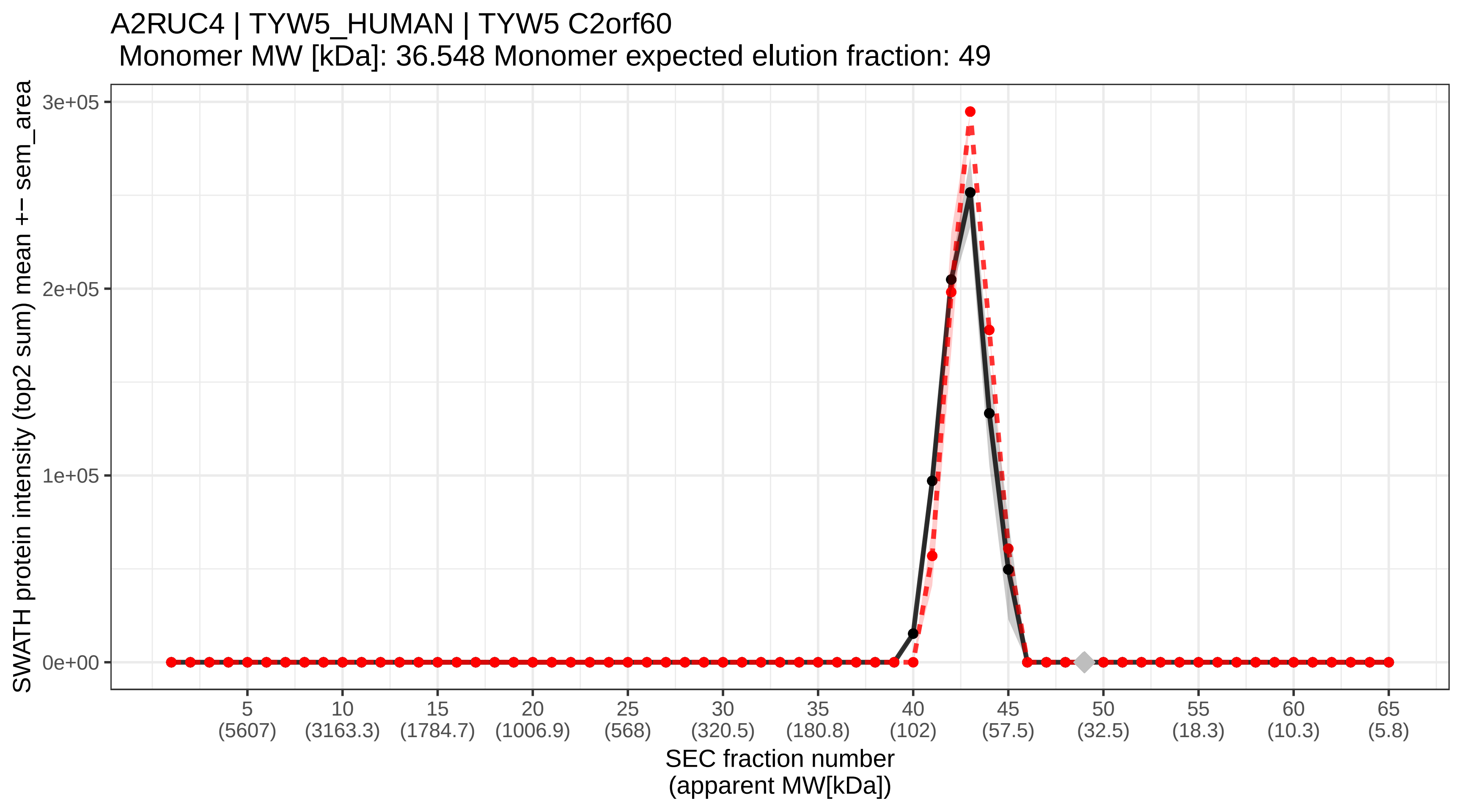

### SECchrom_A2RUS2_DEND3_HUMAN_DENND3_KIAA0870.pdf

A2RUS2 | DEND3_HUMAN | DENND3 KIAA0870
Monomer MW [kDa]: 135.89 Monomer expected elution fraction: 37

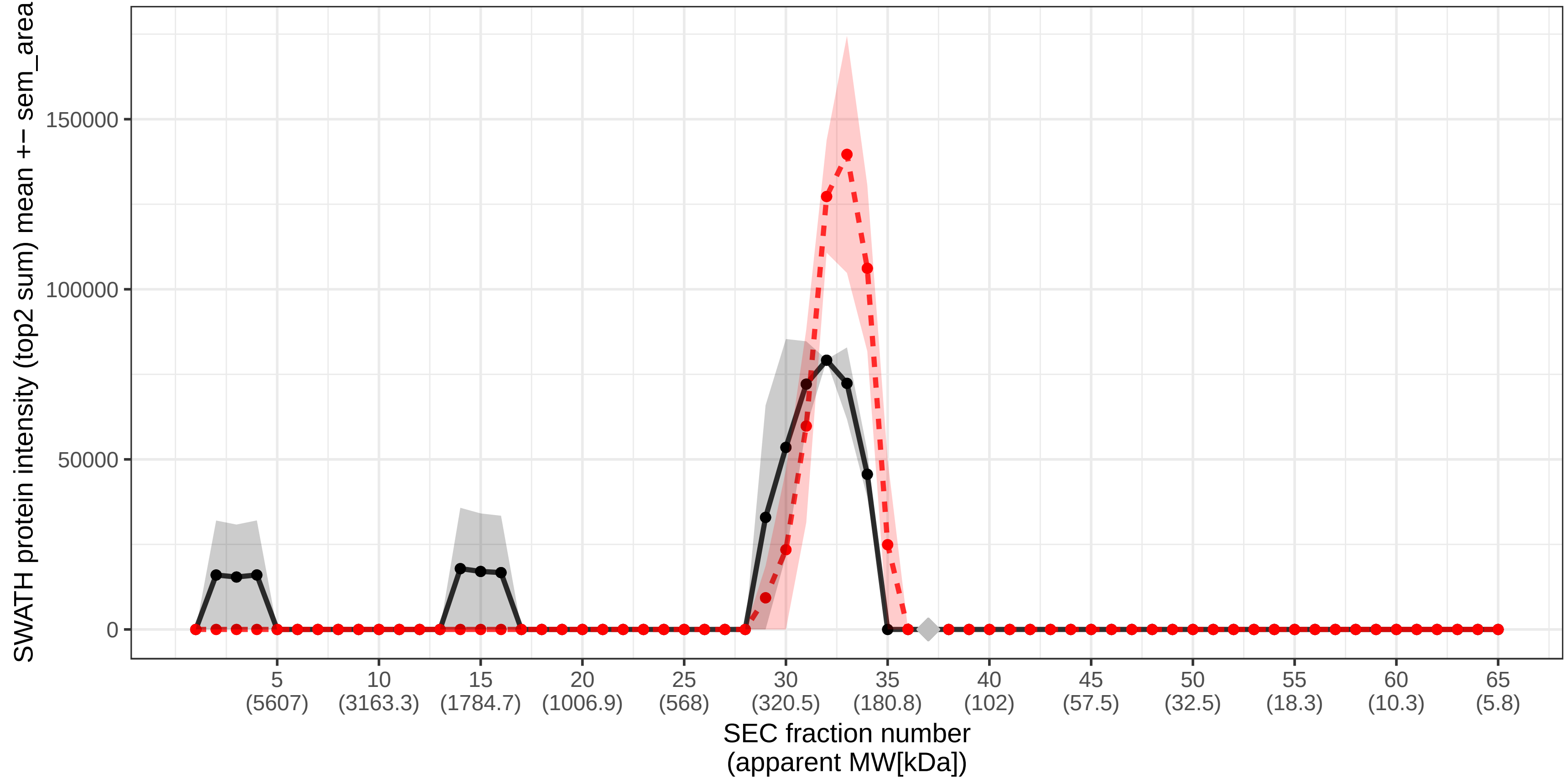

### SECchrom_A3KMH1_VWA8_HUMAN_VWA8_KIAA0564.pdf

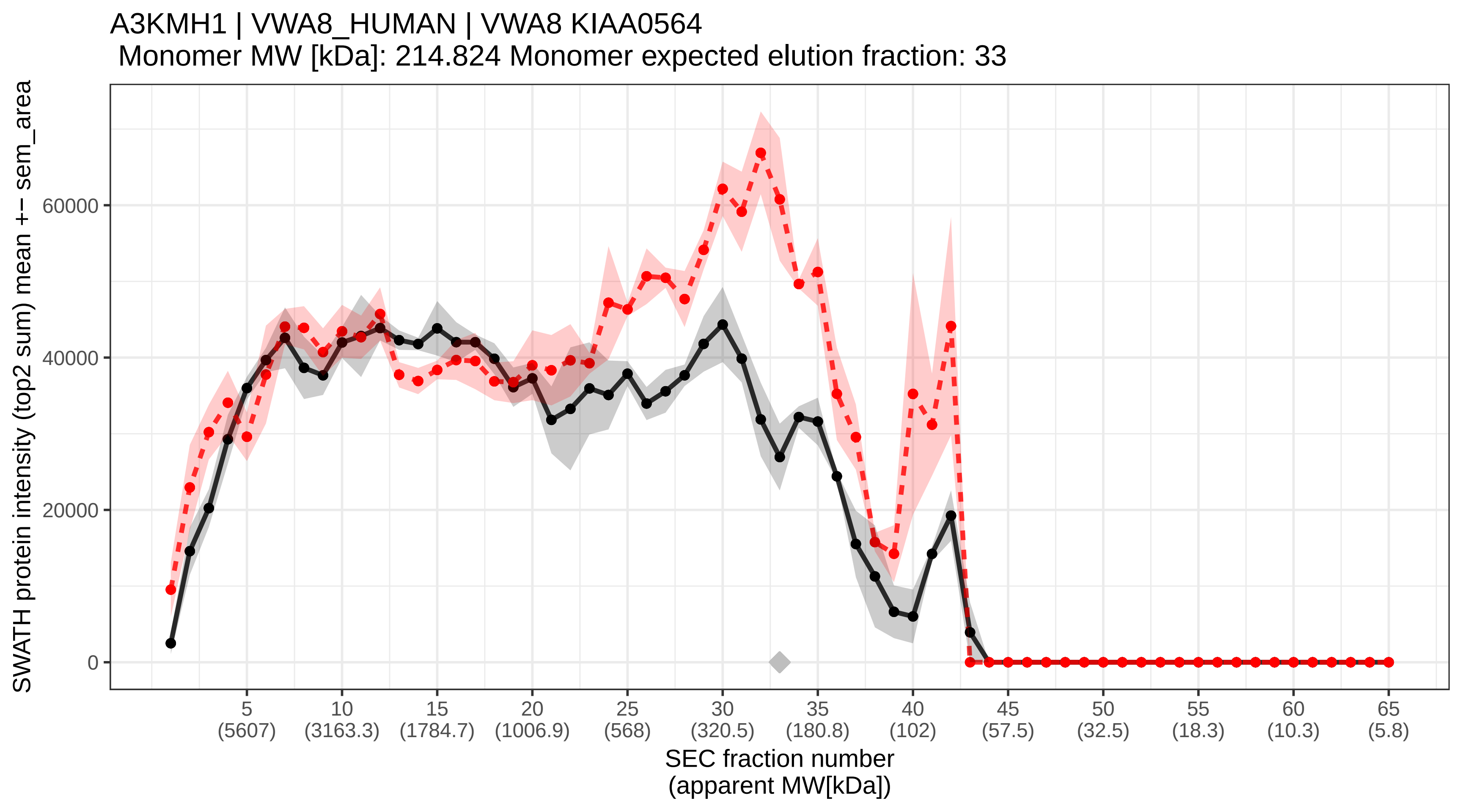

### SECchrom_A3KN83_SBNO1_HUMAN_SBNO1_MOP3.pdf

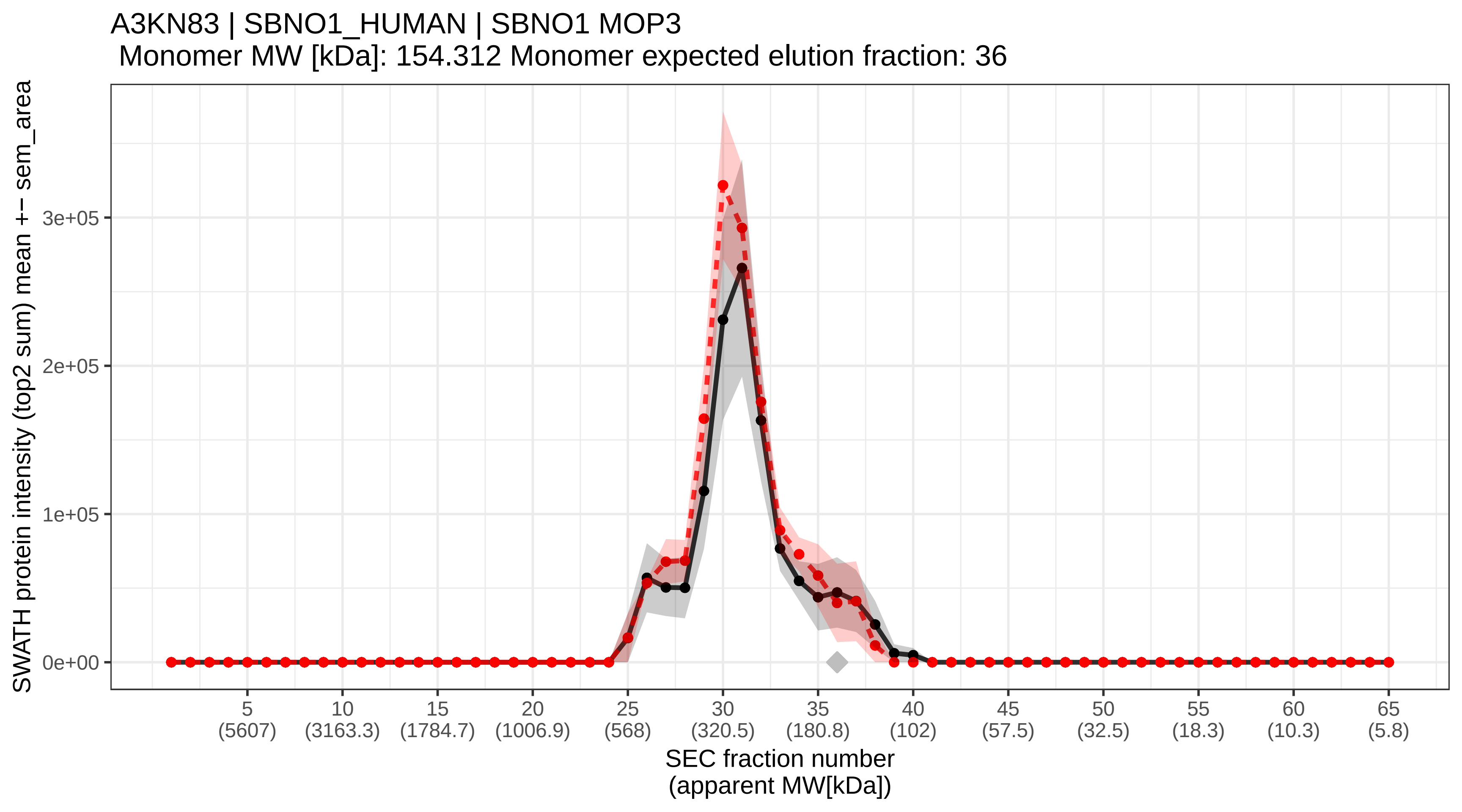

### SECchrom_A4D1P6_WDR91_HUMAN_WDR91_HSPC049.pdf

A4D1P6 | WDR91_HUMAN | WDR91 HSPC049
Monomer MW [kDa]: 83.344 Monomer expected elution fraction: 42

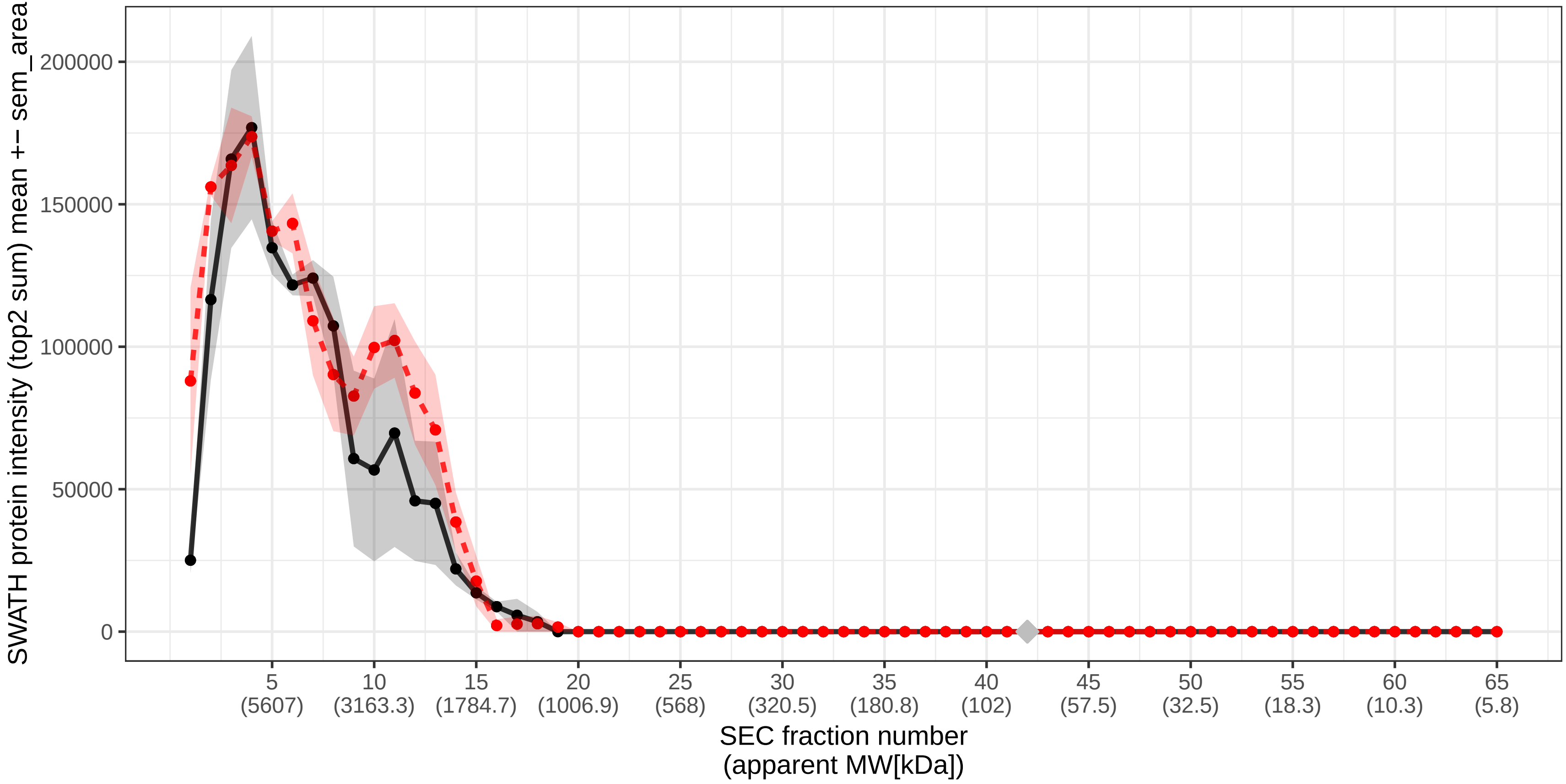

### SECchrom_A5A3E0_POTEF_HUMAN_POTEF_A26C1B.pdf

A5A3E0 | POTEF_HUMAN | POTEF A26C1B
Monomer MW [kDa]: 121.445 Monomer expected elution fraction: 38

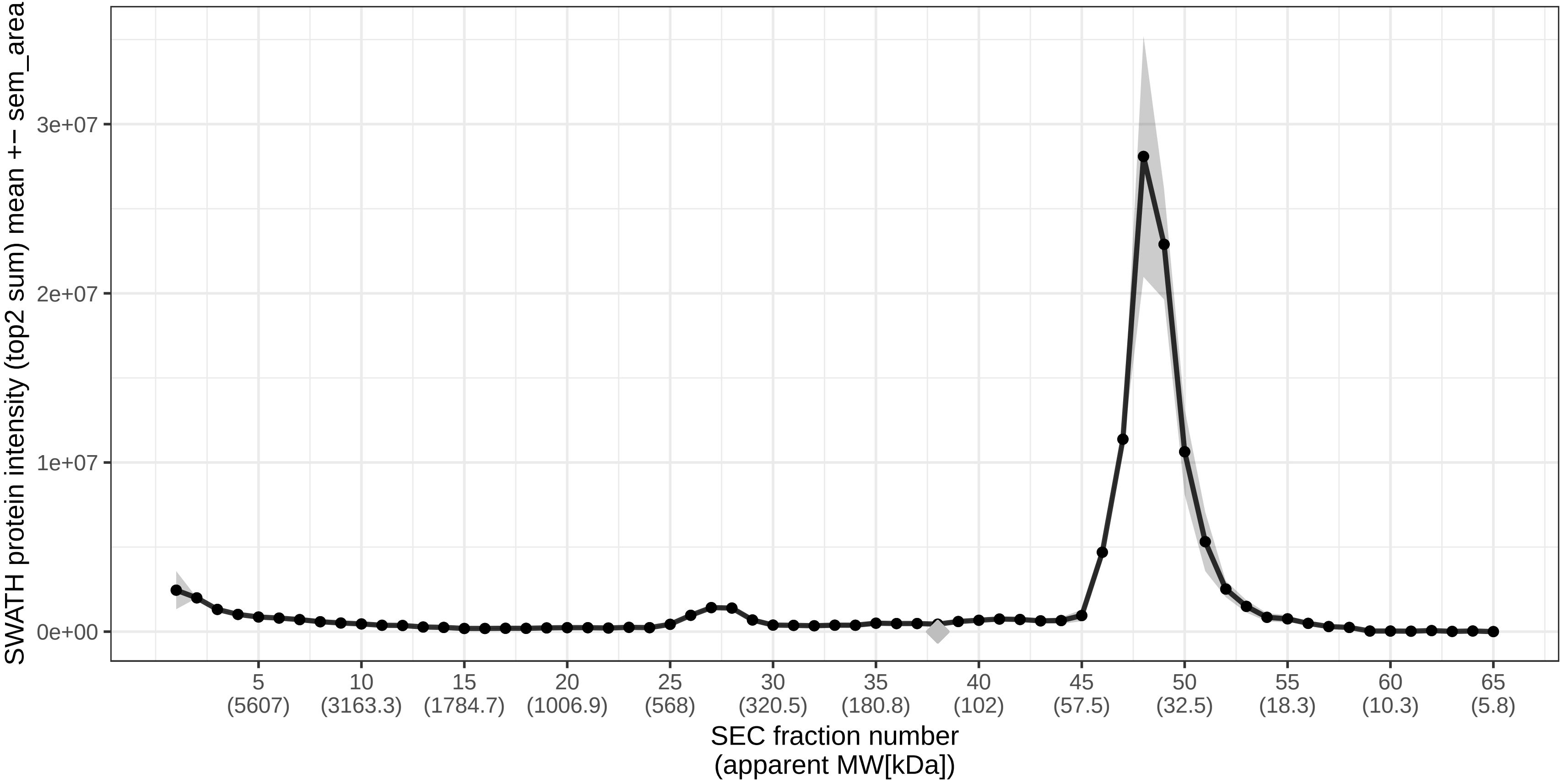

### SECchrom_A5D8V6_VP37C_HUMAN_VPS37C_PML39.pdf

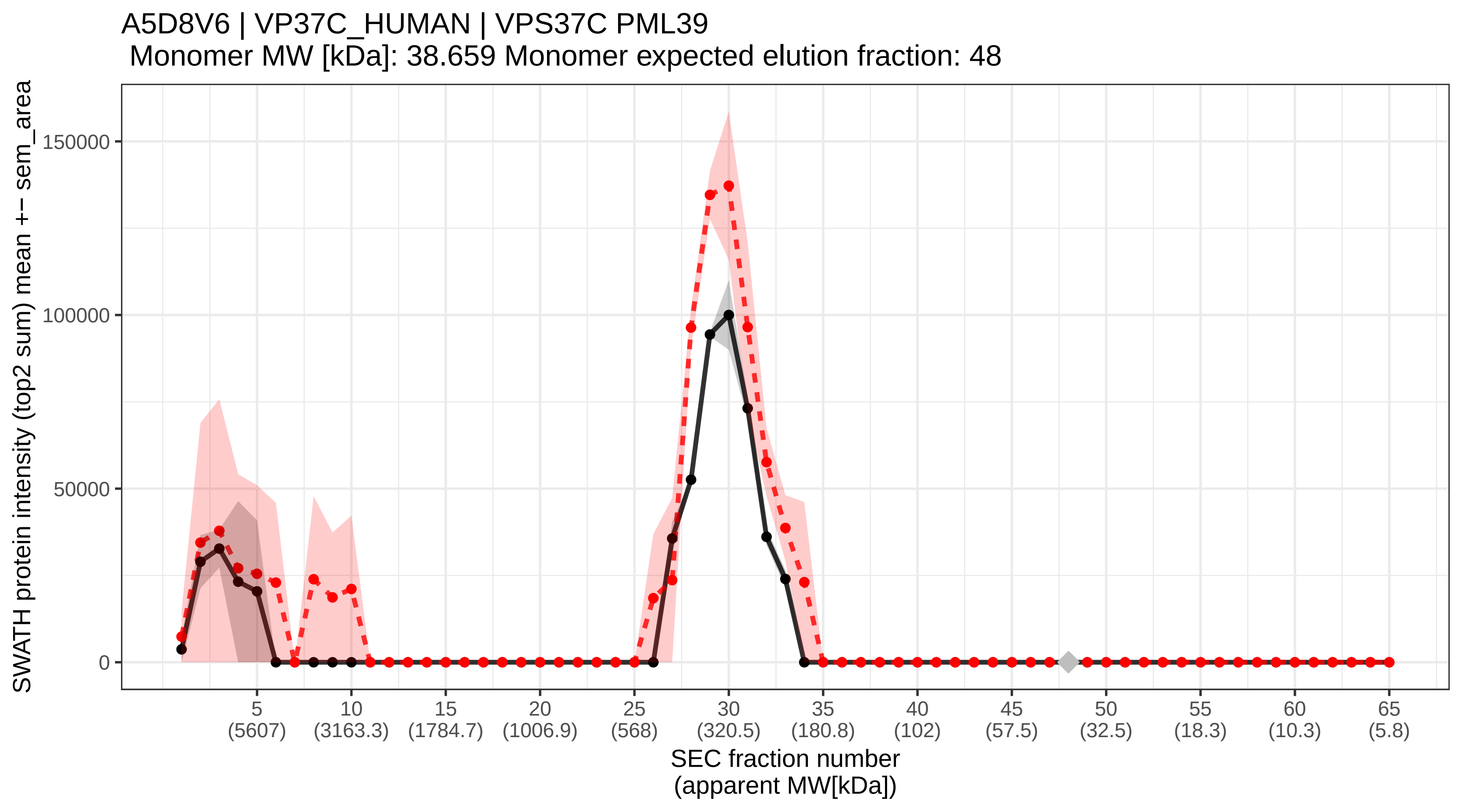

### SECchrom_A5PLL7_TM189_HUMAN_TMEM189_KUA.pdf

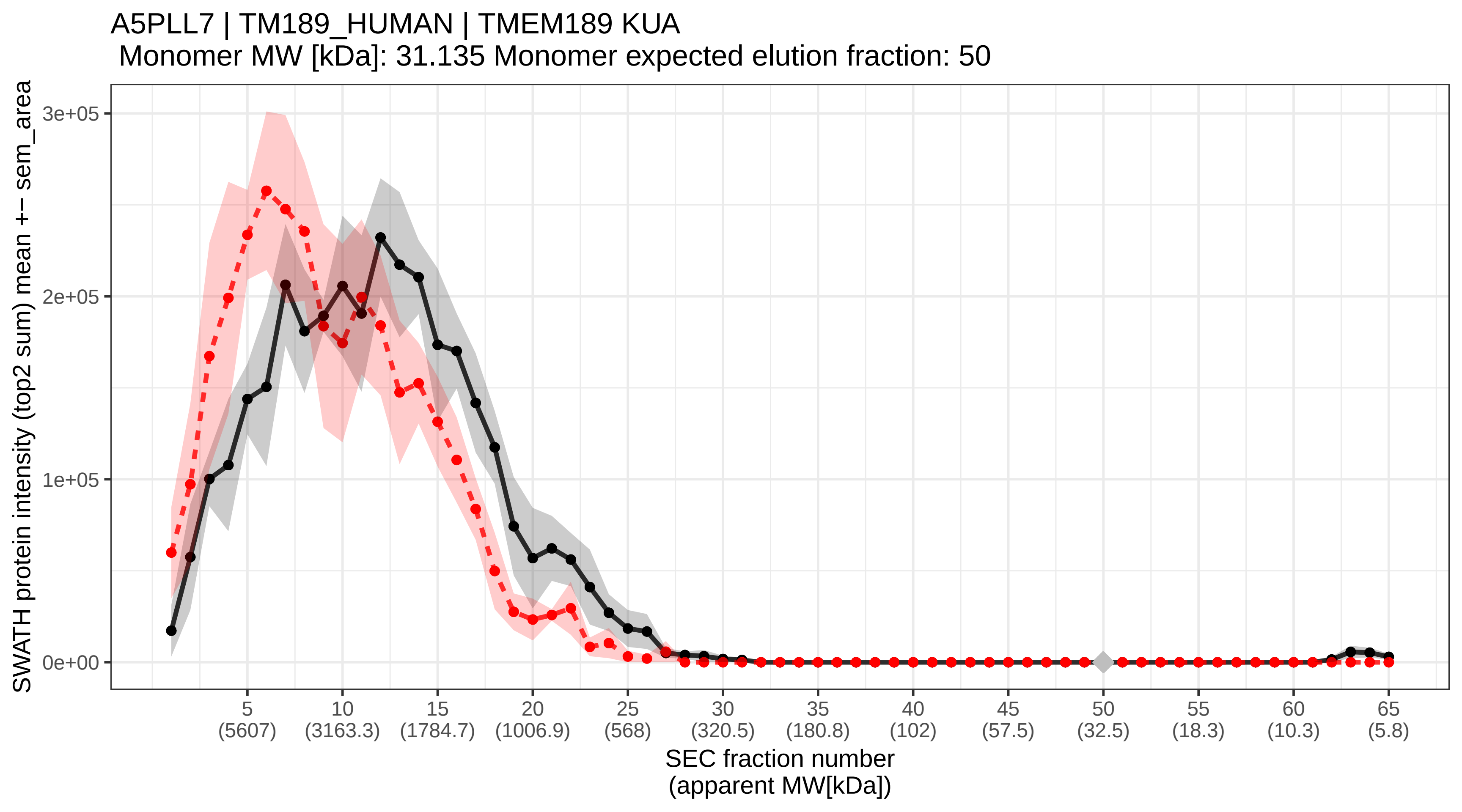

### SECchrom_A5PLN9_TPC13_HUMAN_TRAPPC13_C5orf44.pdf

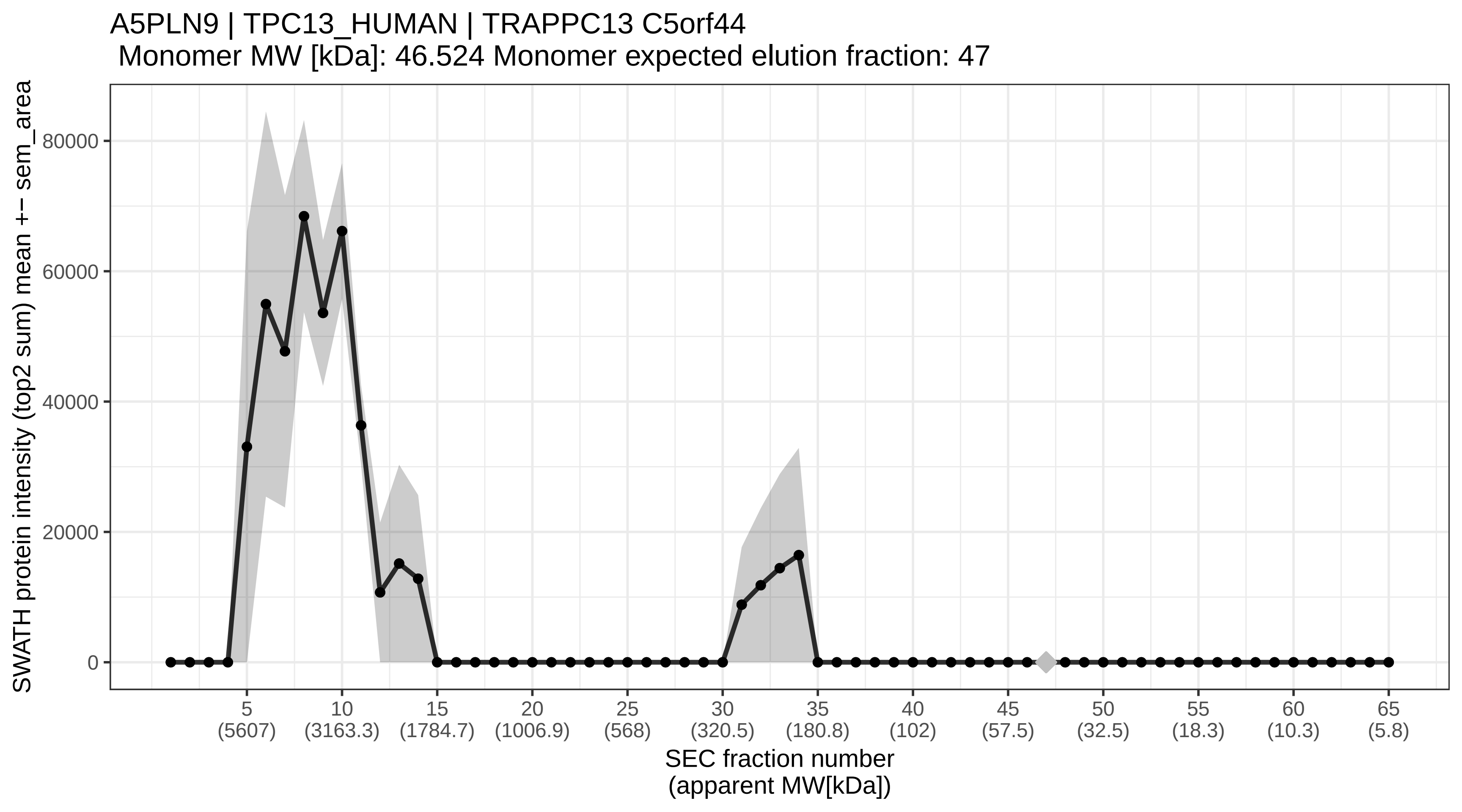

### SECchrom_A5YKK6_CNOT1_HUMAN_CNOT1_CDC39_KIAA1007_NOT1_AD-00.pdf

A5YKK6 | CNOT1_HUMAN | CNOT1 CDC39 KIAA1007 NOT1 AD−005
Monomer MW [kDa]: 266.939 Monomer expected elution fraction: 32

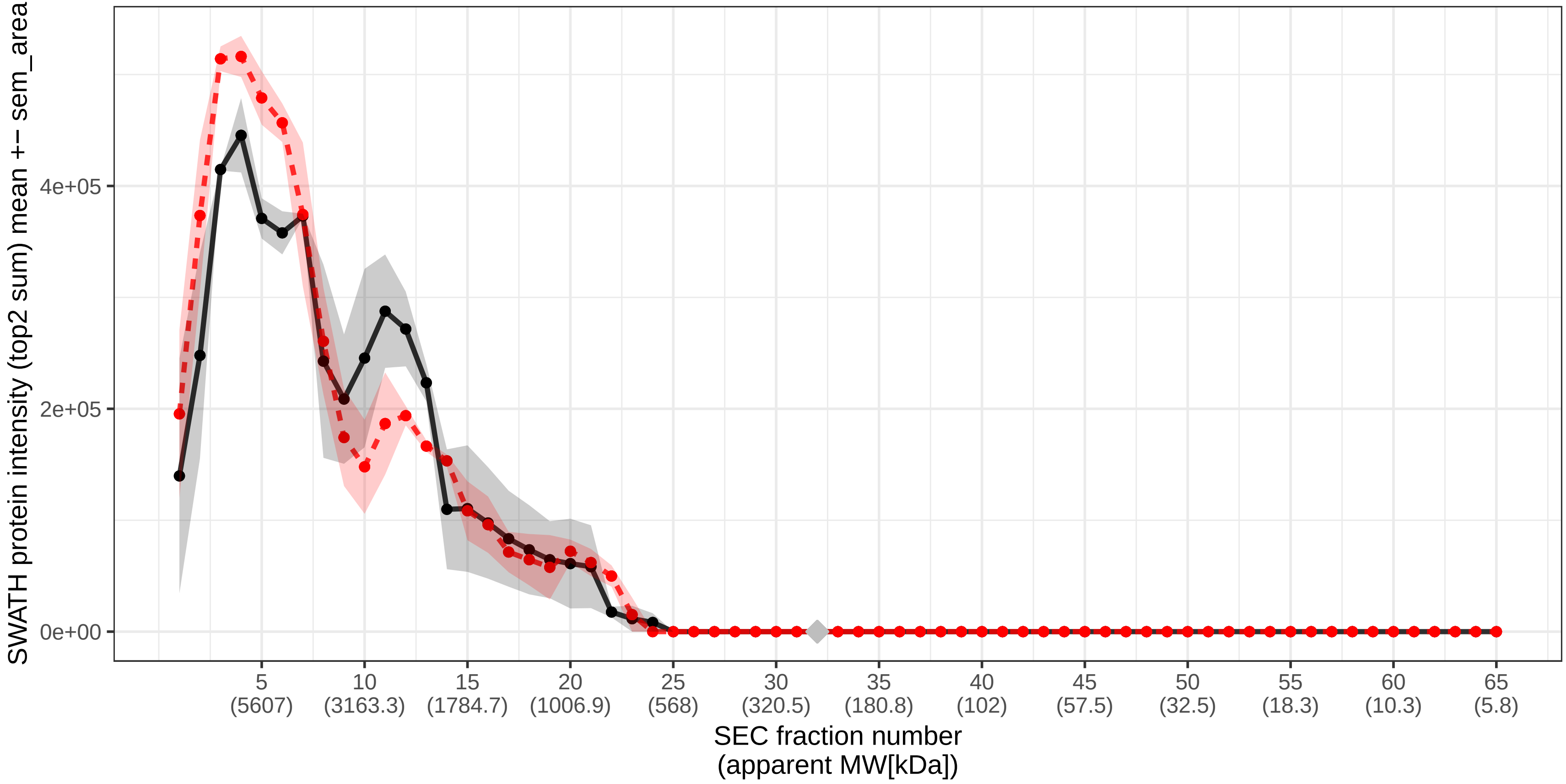

### SECchrom_A6NDG6_PGP_HUMAN_PGP.pdf

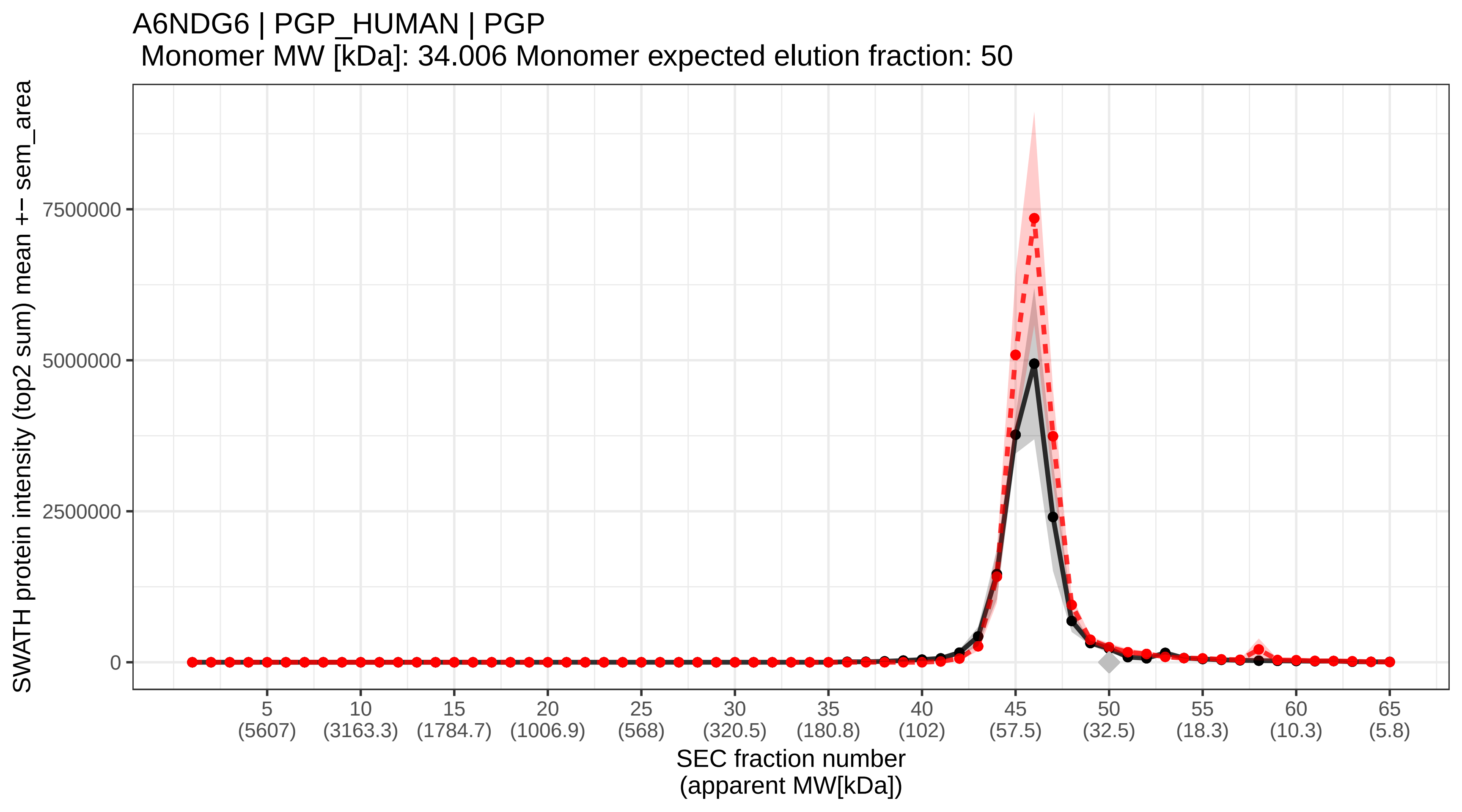

### SECchrom_A6NDU8_CE051_HUMAN_C5orf51.pdf

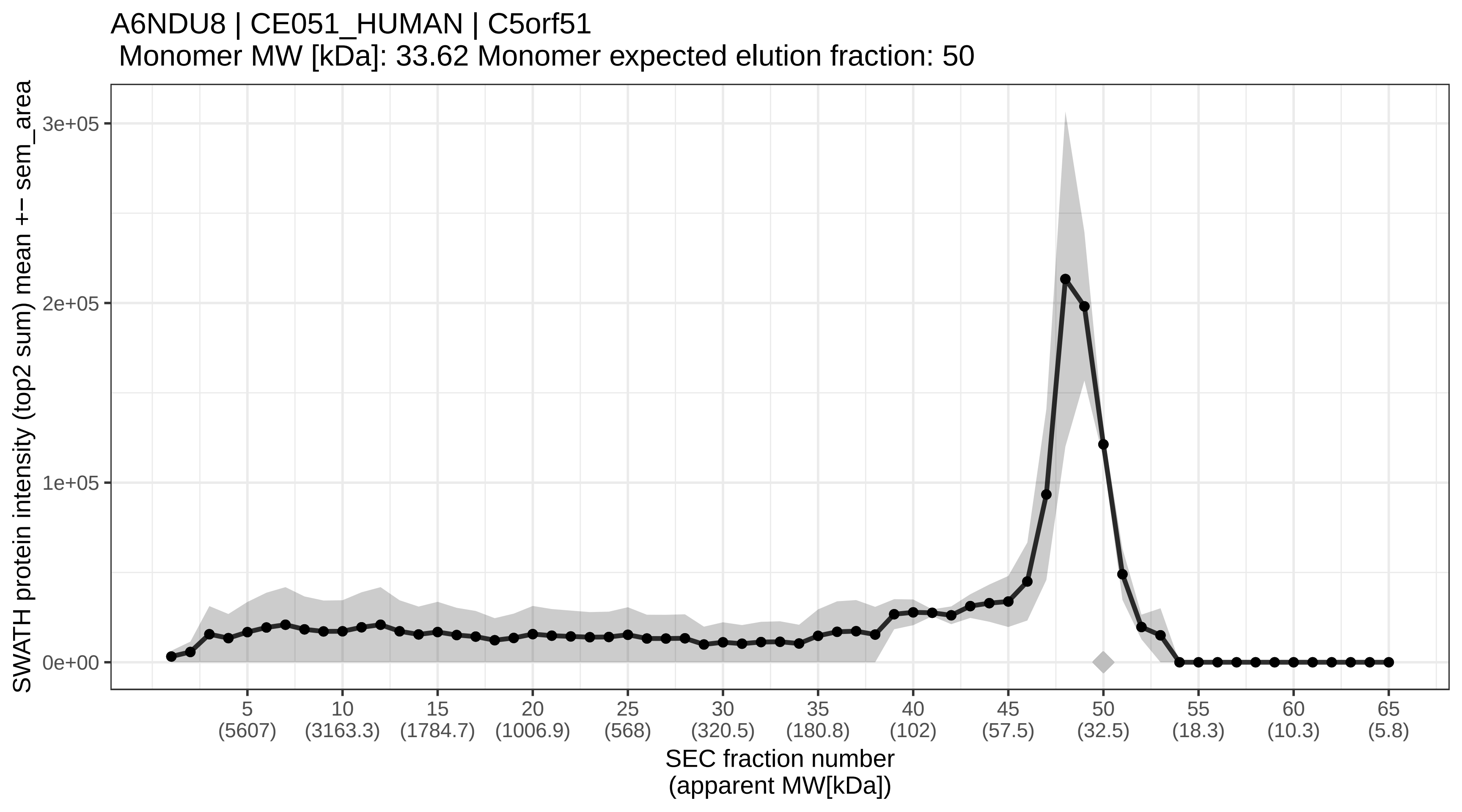

### SECchrom_A6NED2_RCCD1_HUMAN_RCCD1.pdf

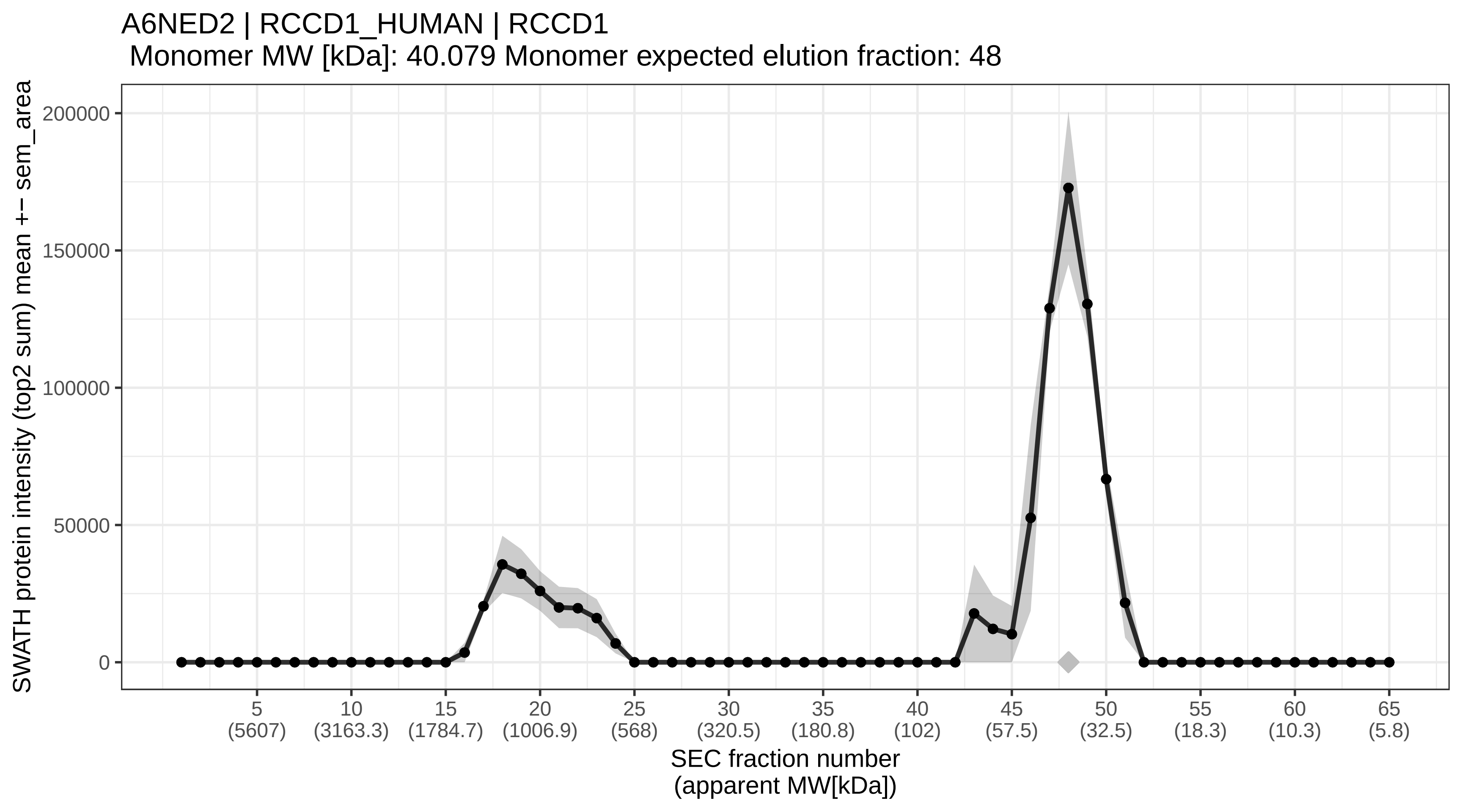

### SECchrom_A6NHR9_SMHD1_HUMAN_SMCHD1_KIAA0650.pdf

A6NHR9 | SMHD1_HUMAN | SMCHD1 KIAA0650
Monomer MW [kDa]: 226.374 Monomer expected elution fraction: 33

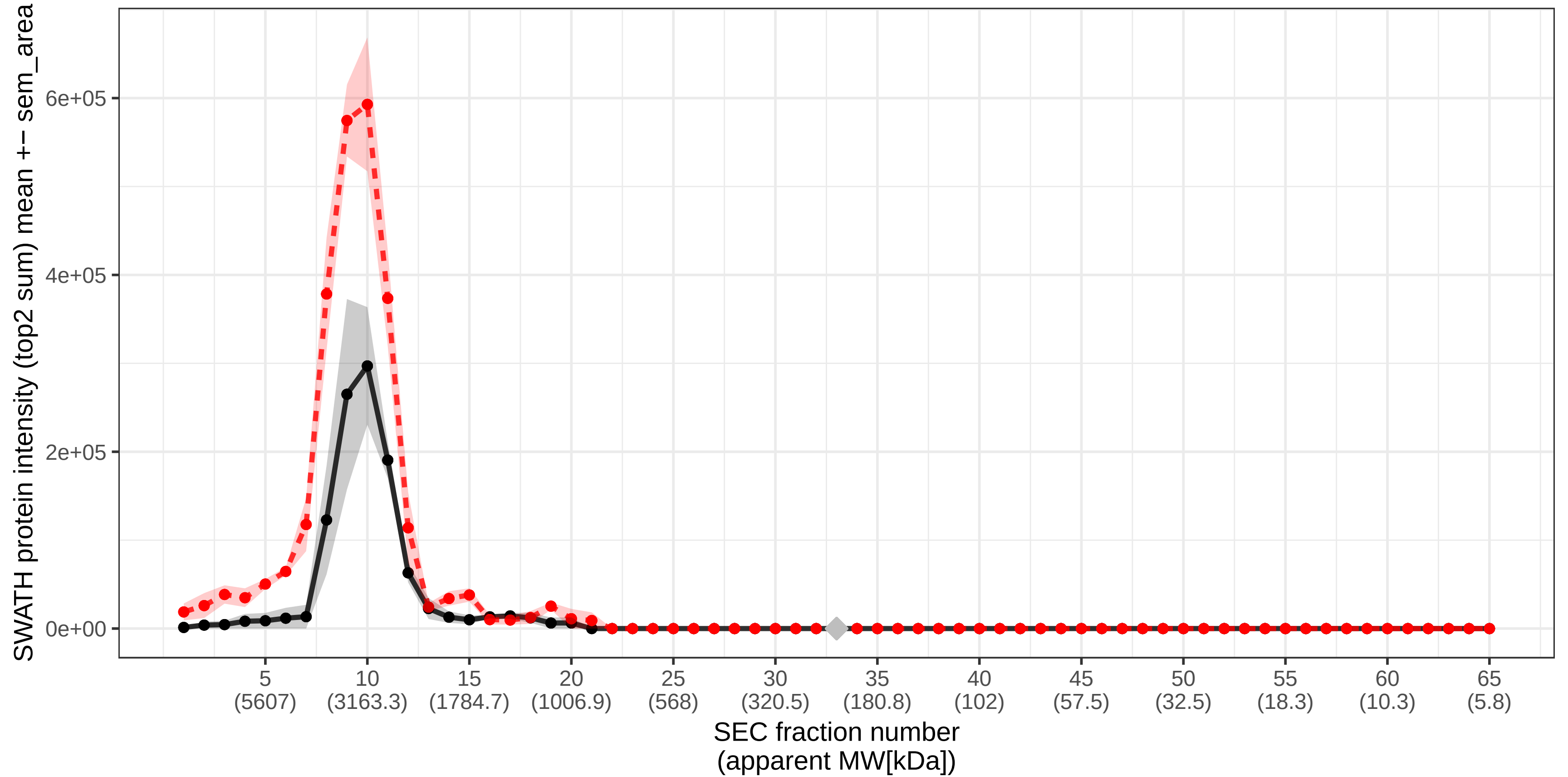

### SECchrom_A6NIH7_U119B_HUMAN_UNC119B.pdf

#### A6NIH7 | U119B_HUMAN | UNC119B
Monomer MW [kDa]: 28.137 Monomer expected elution fraction: 51

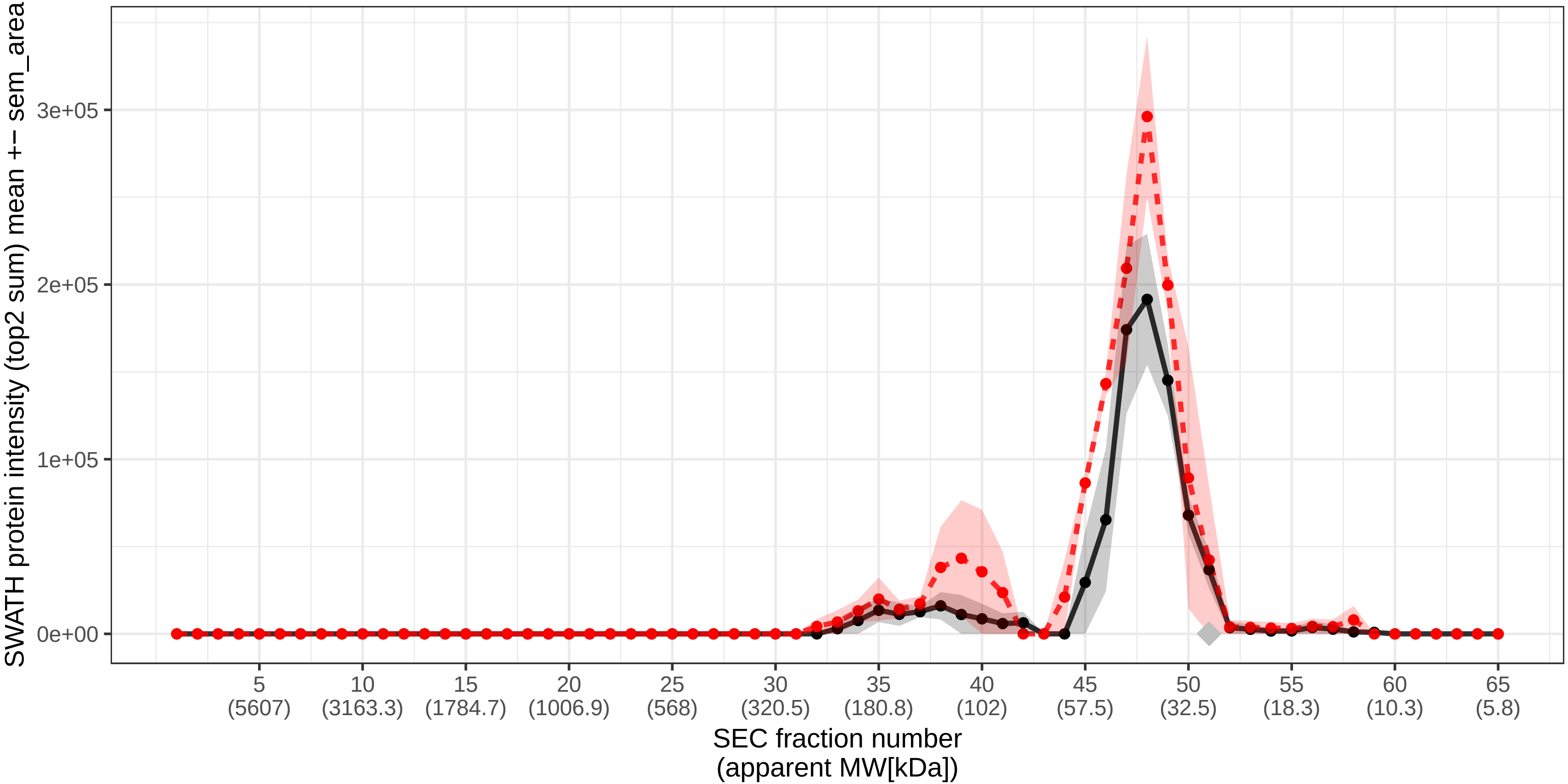

### SECchrom_A6NKD9_CC85C_HUMAN_CCDC85C.pdf

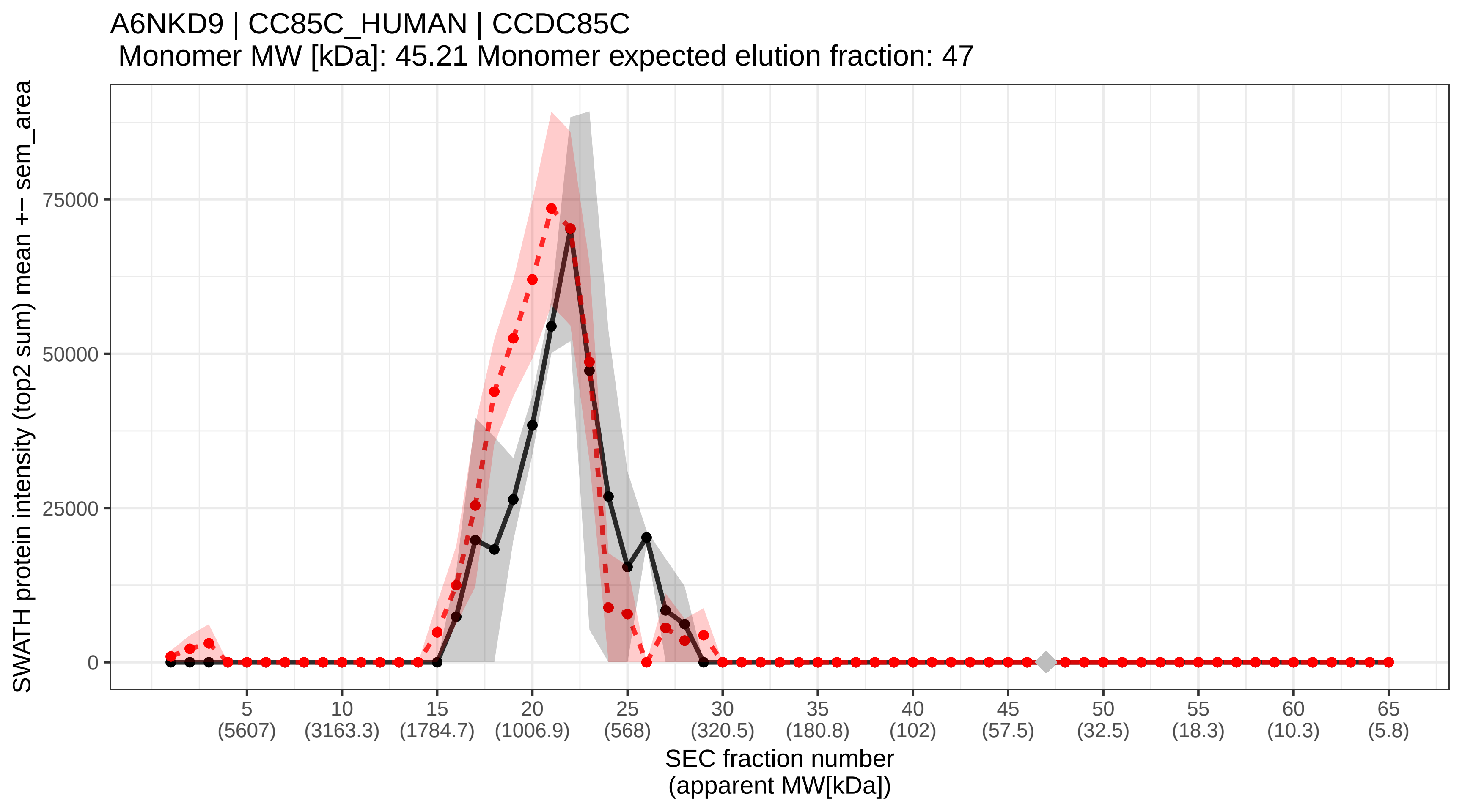

### SECchrom_A6NKF1_SAC31_HUMAN_SAC3D1_SHD1.pdf

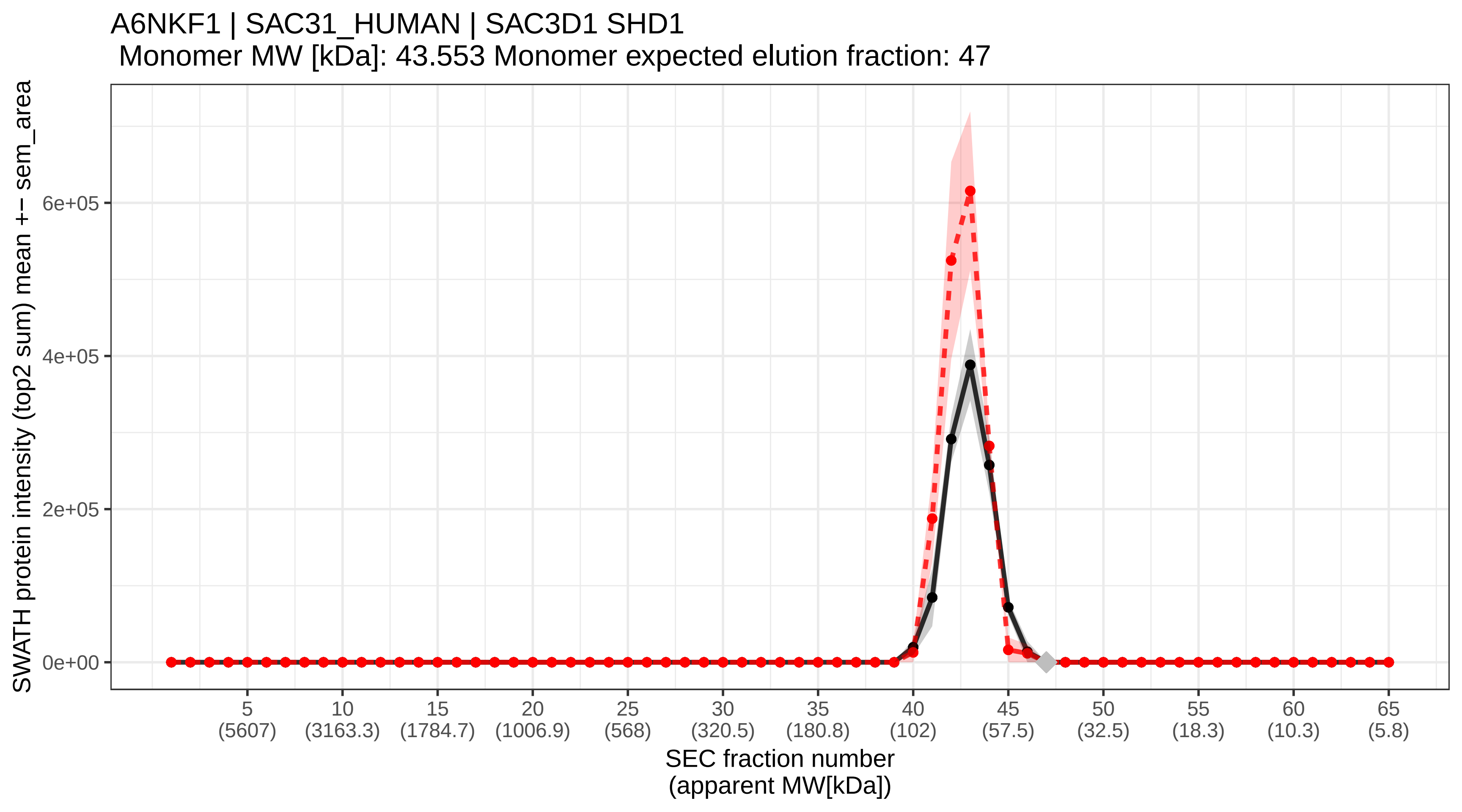

### SECchrom_A7E2V4_ZSWM8_HUMAN_ZSWIM8_KIAA0913.pdf

A7E2V4 | ZSWM8_HUMAN | ZSWIM8 KIAA0913
Monomer MW [kDa]: 197.297 Monomer expected elution fraction: 34

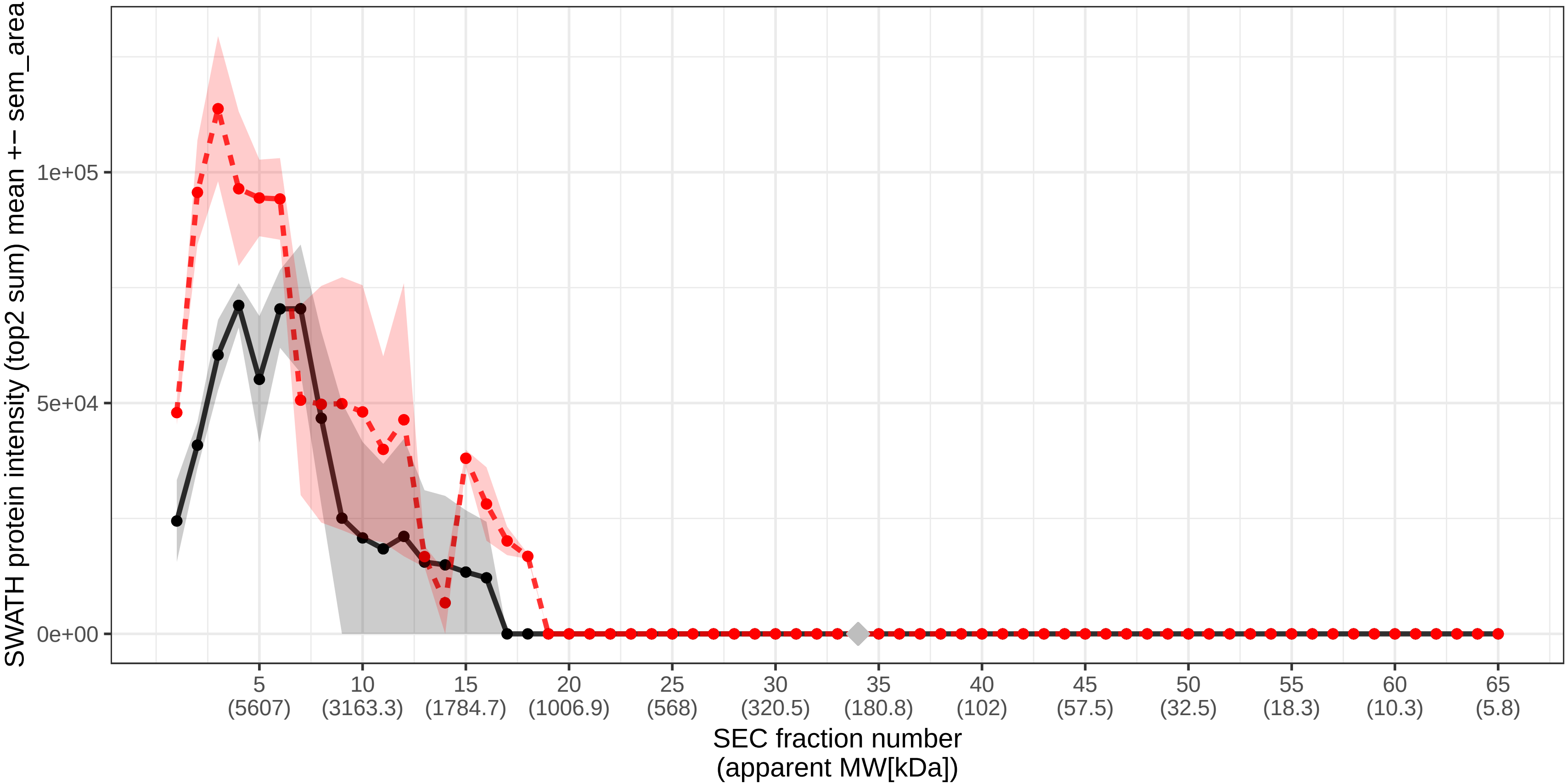

### SECchrom_A8MXV4_NUD19_HUMAN_NUDT19.pdf

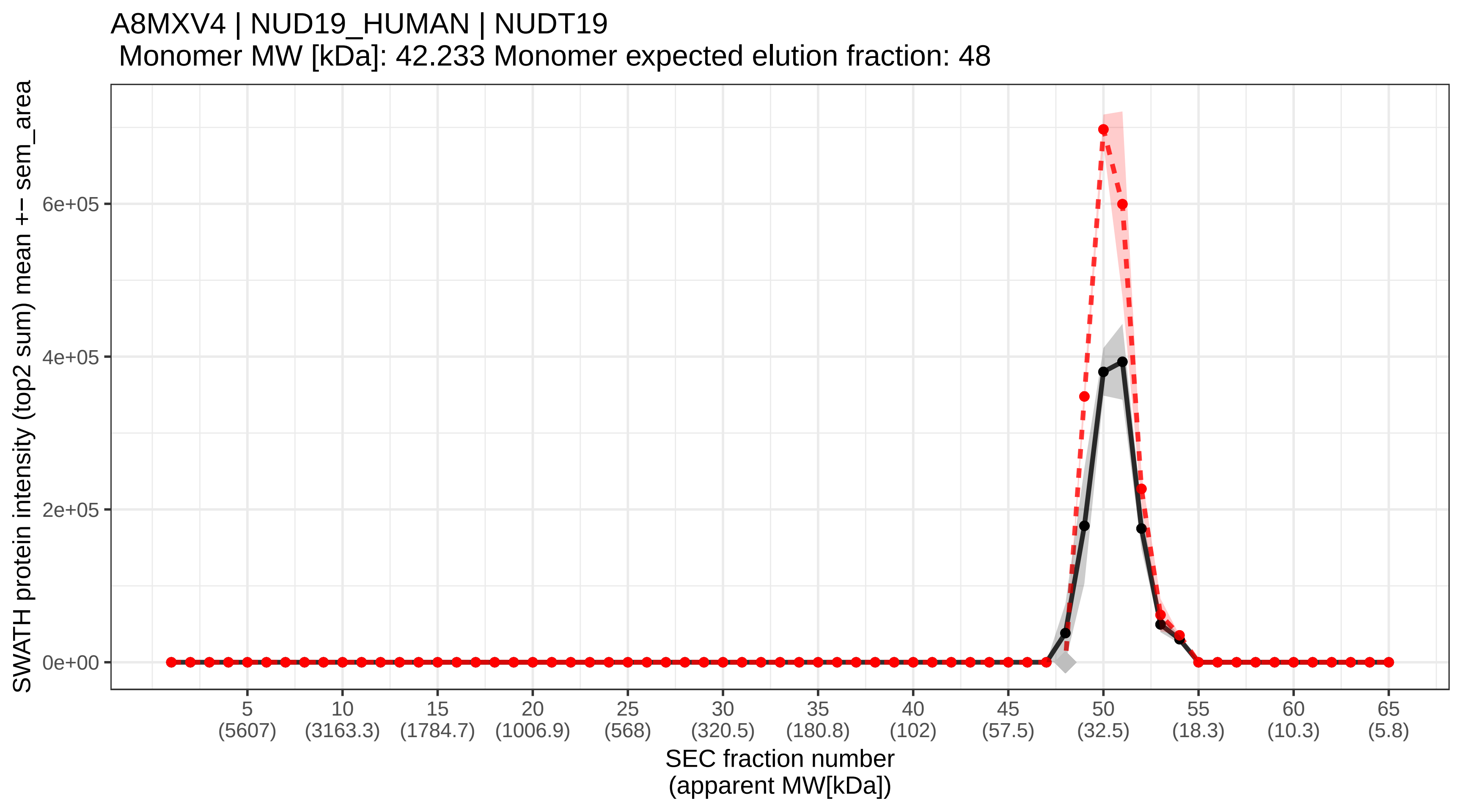

### SECchrom_A9UHW6_MI4GD_HUMAN_MIF4GD_SLIP1.pdf

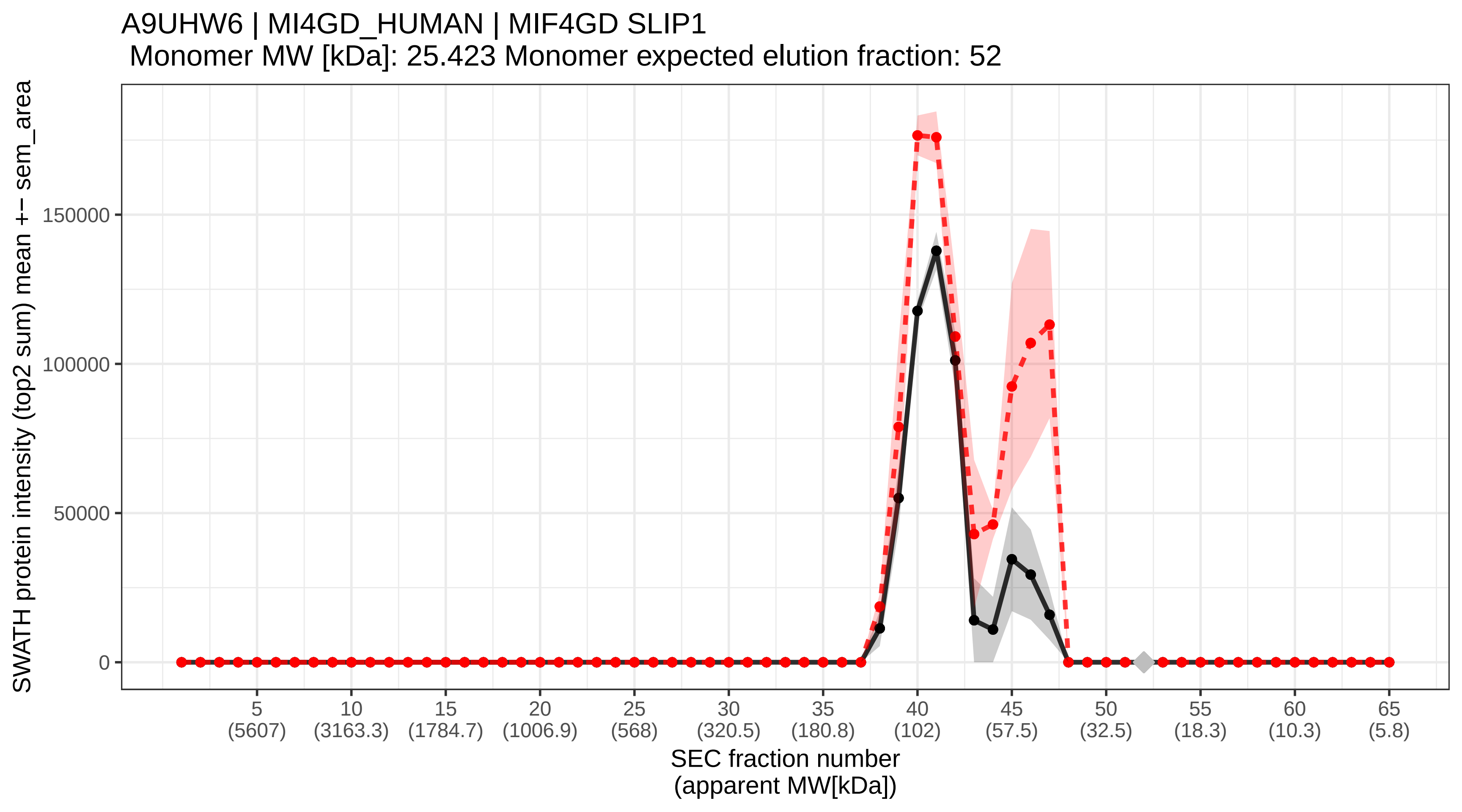

### SECchrom_C9JLW8_MCRI1_HUMAN_MCRIP1_FAM195B_GRAN2.pdf

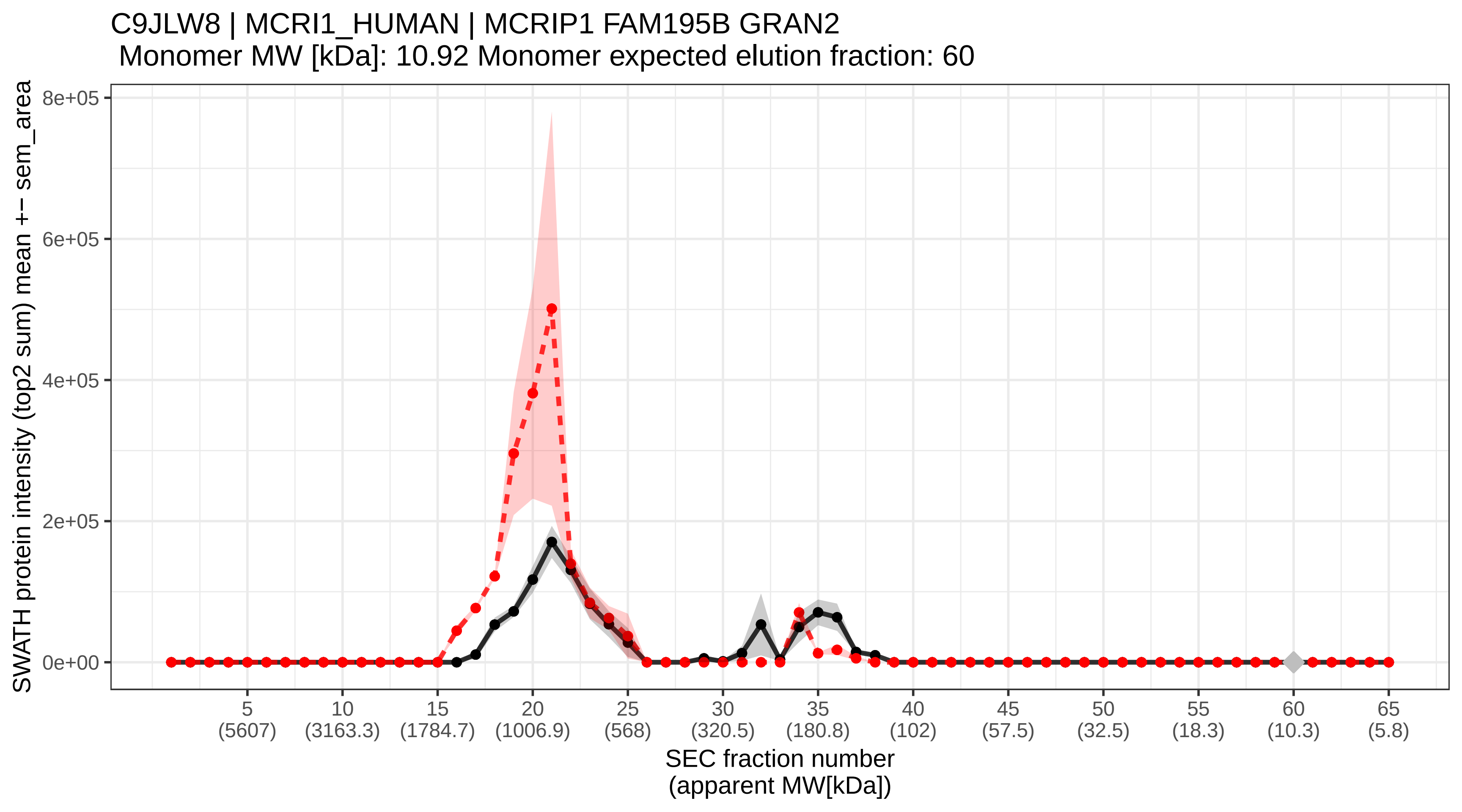

### SECchrom_cRAP_ALBU_BOVIN_NA_NA.pdf

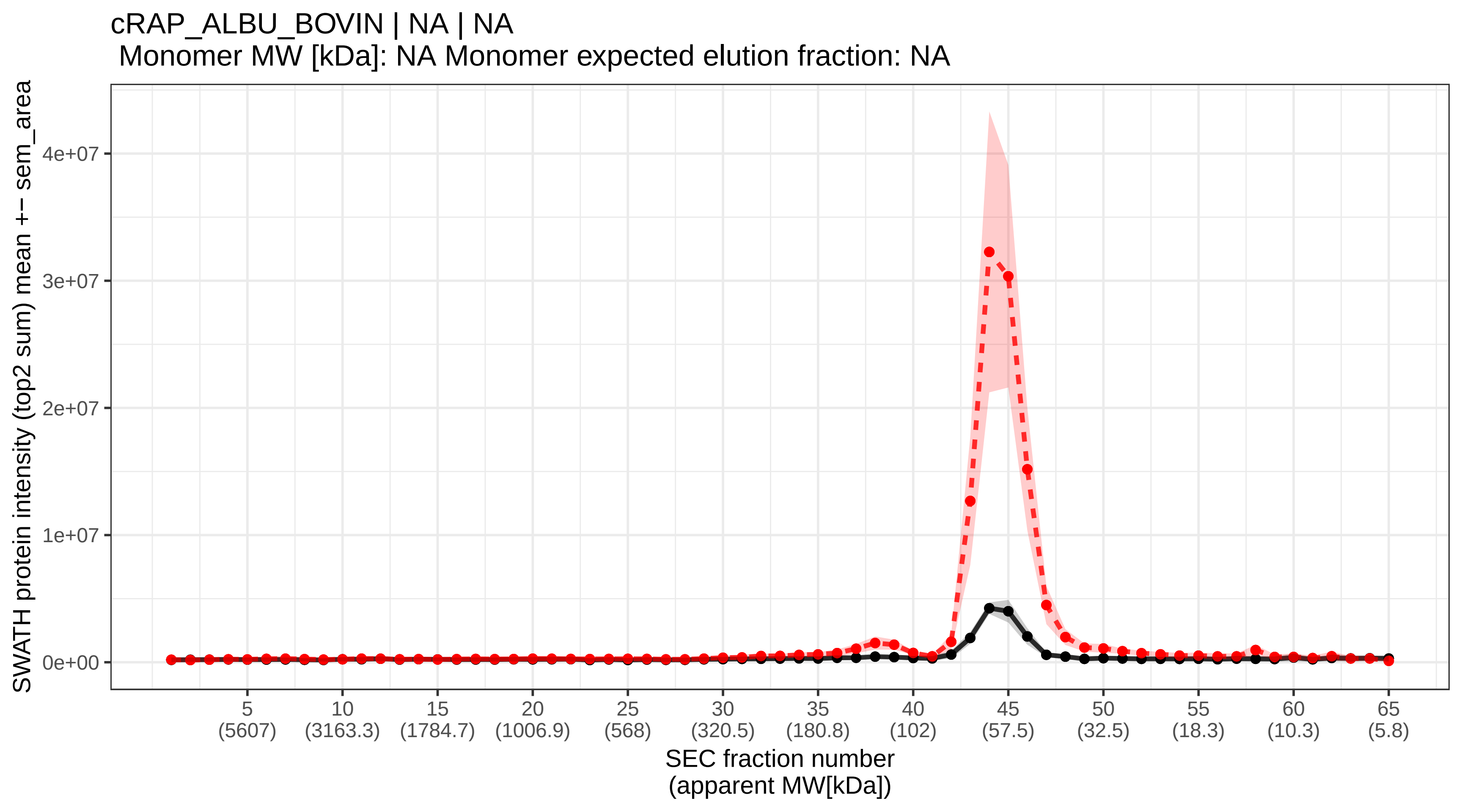

### SECchrom_cRAP_MYG_HORSE_NA_NA.pdf

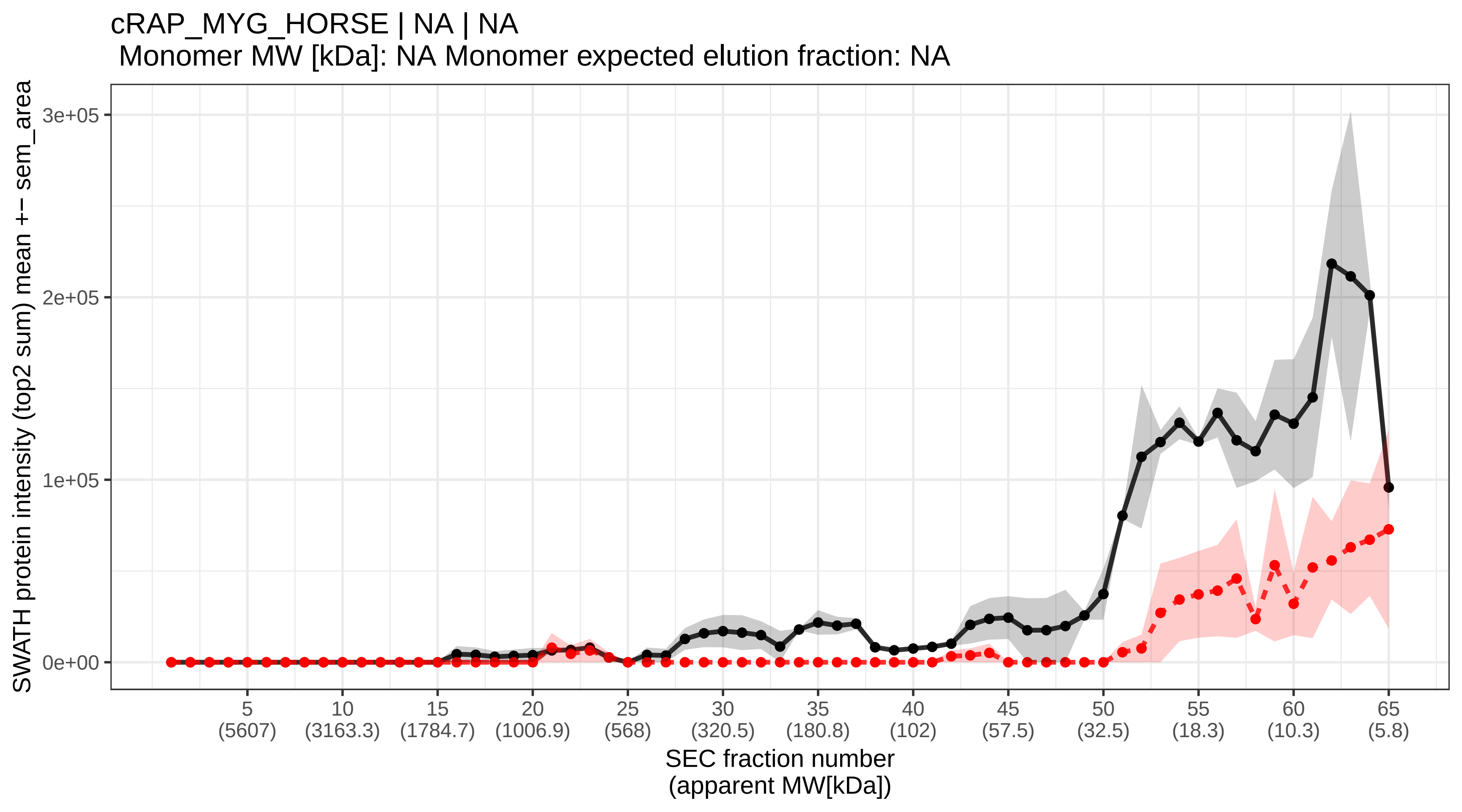

### SECchrom_cRAP_OVAL_CHICK_NA_NA.pdf

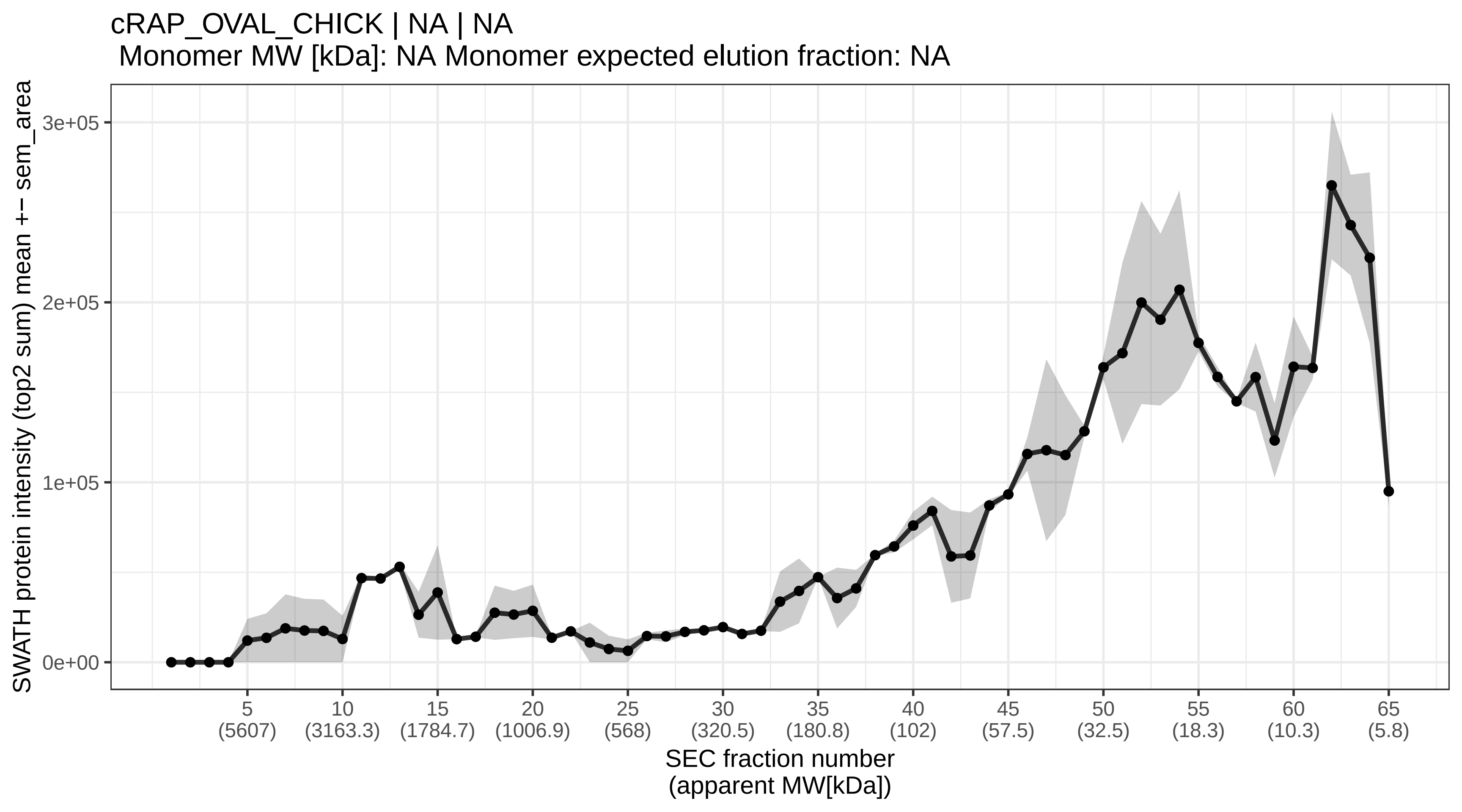

### SECchrom_iRT_protein_NA_NA.pdf

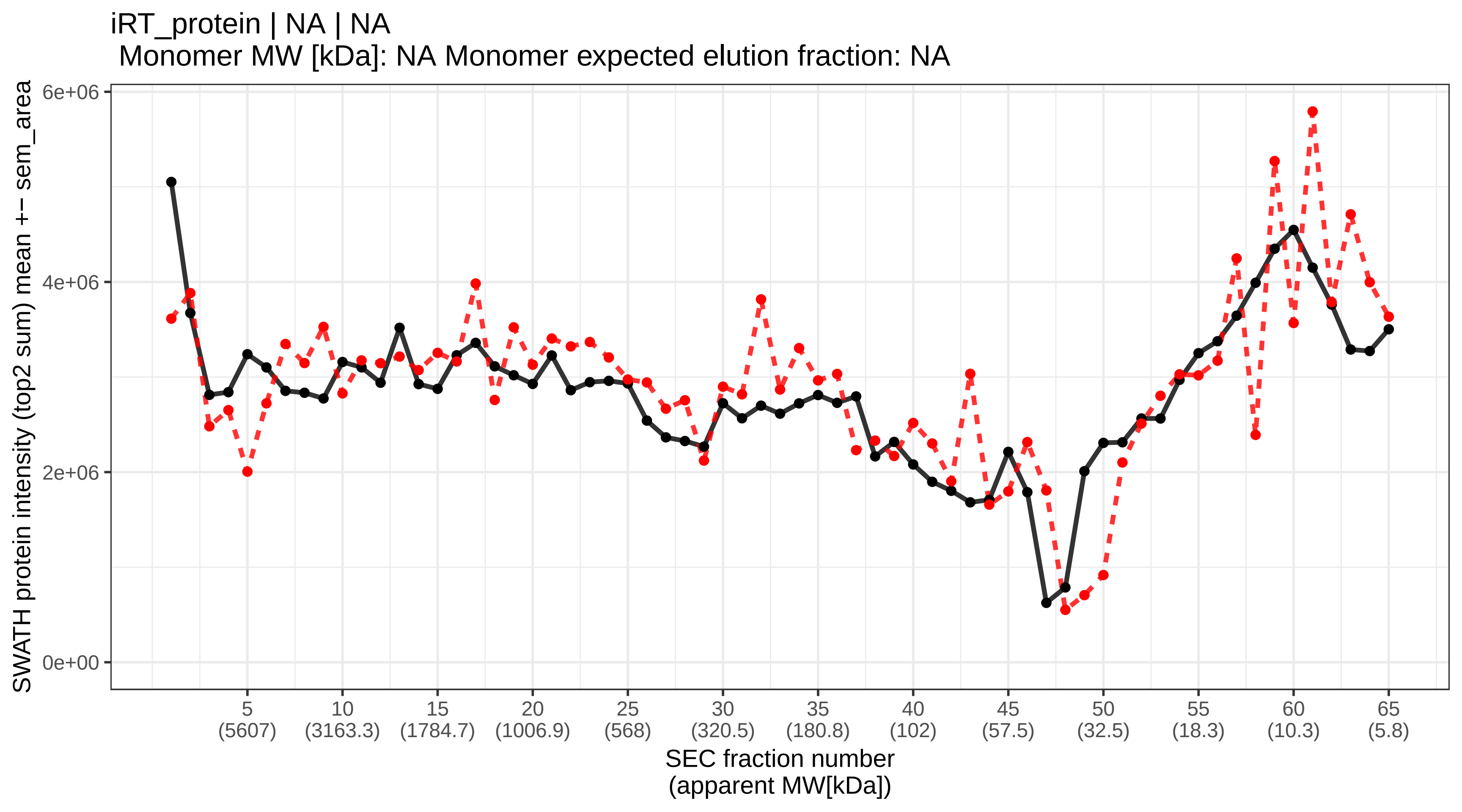

### SECchrom_L0R6Q1_NA_NA.pdf

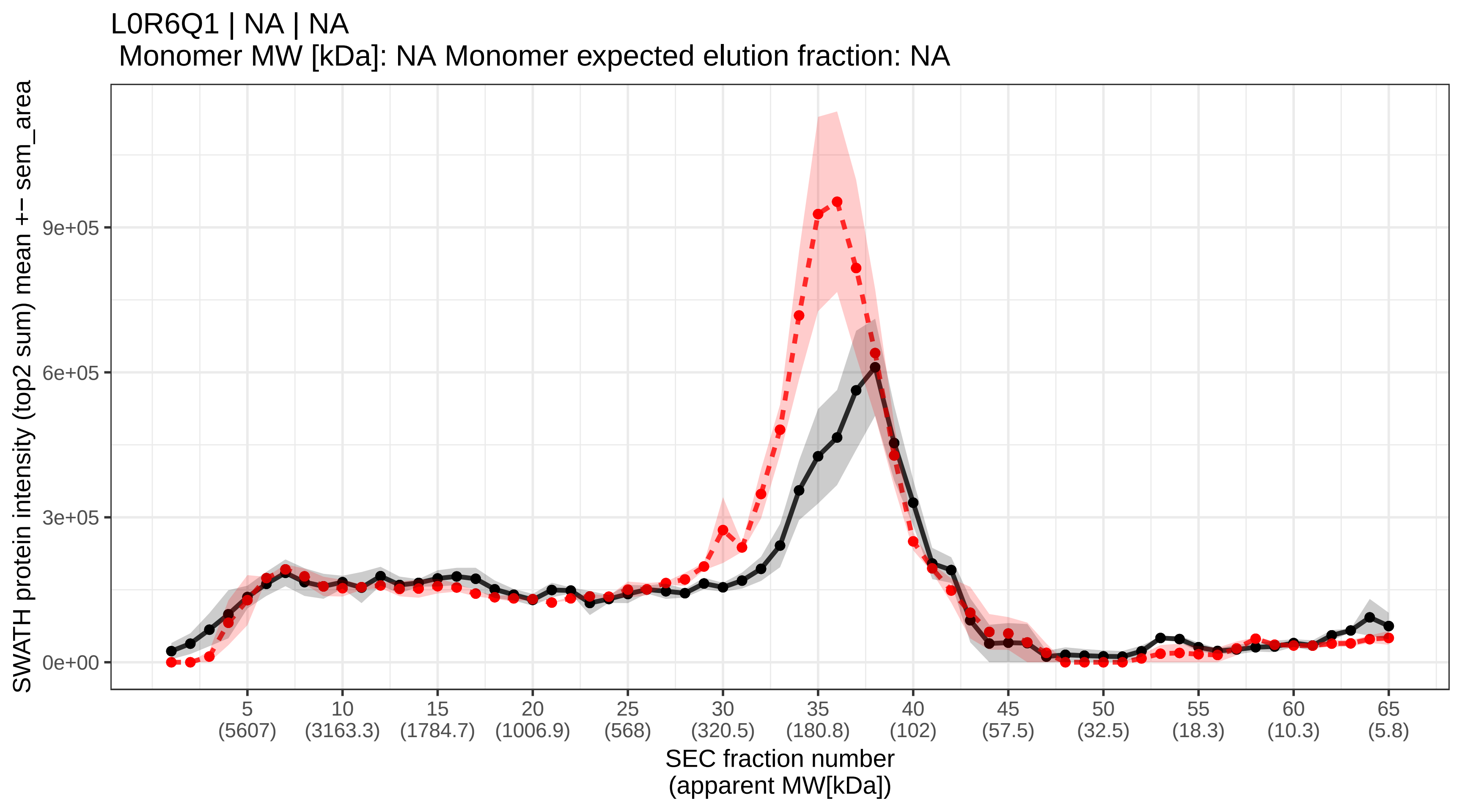

### SECchrom_O00115_DNS2A_HUMAN_DNASE2_DNASE2A_DNL2.pdf

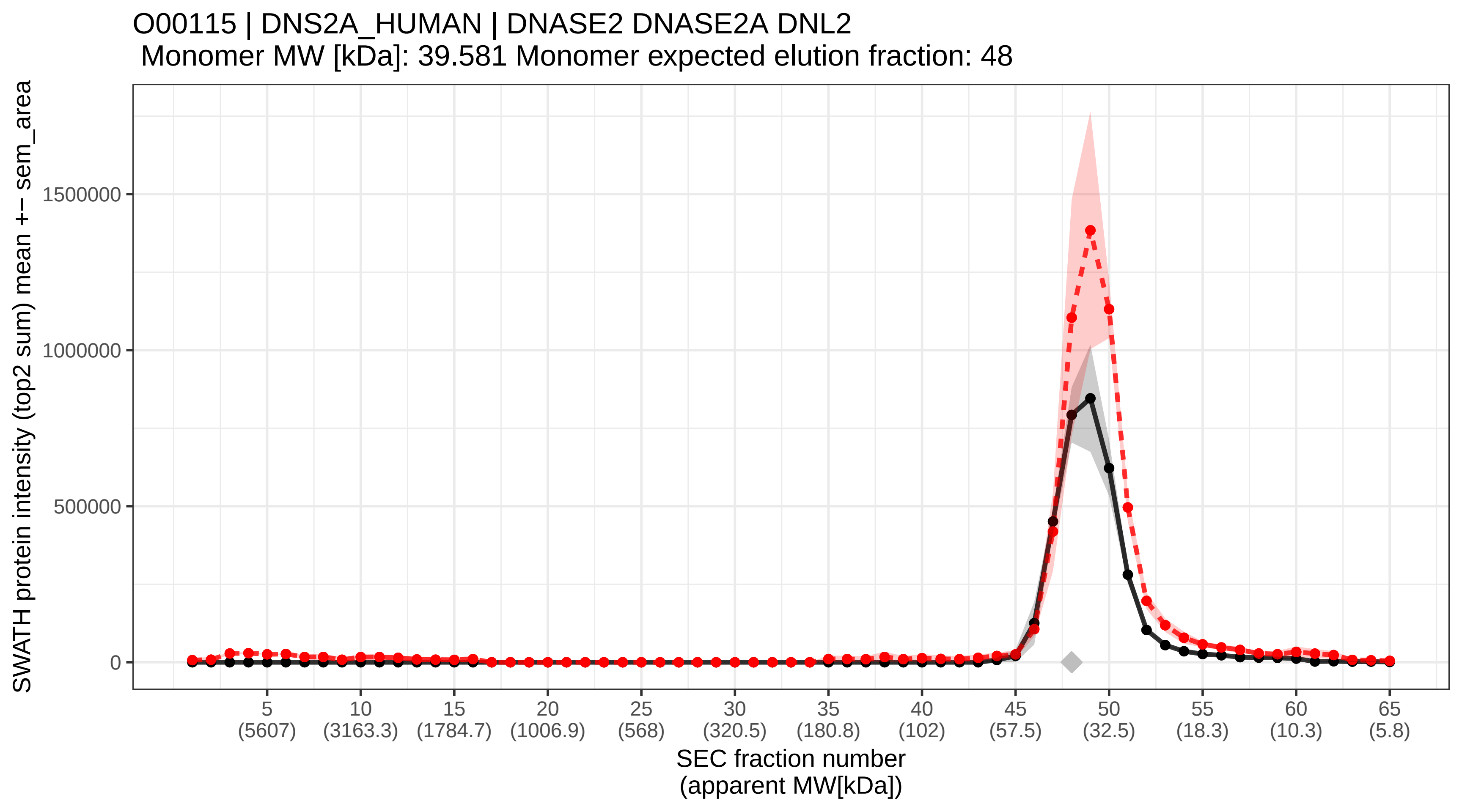

### SECchrom_O00116_ADAS_HUMAN_AGPS_AAG5.pdf

O00116 | ADAS_HUMAN | AGPS AAG5
Monomer MW [kDa]: 72.912 Monomer expected elution fraction: 43

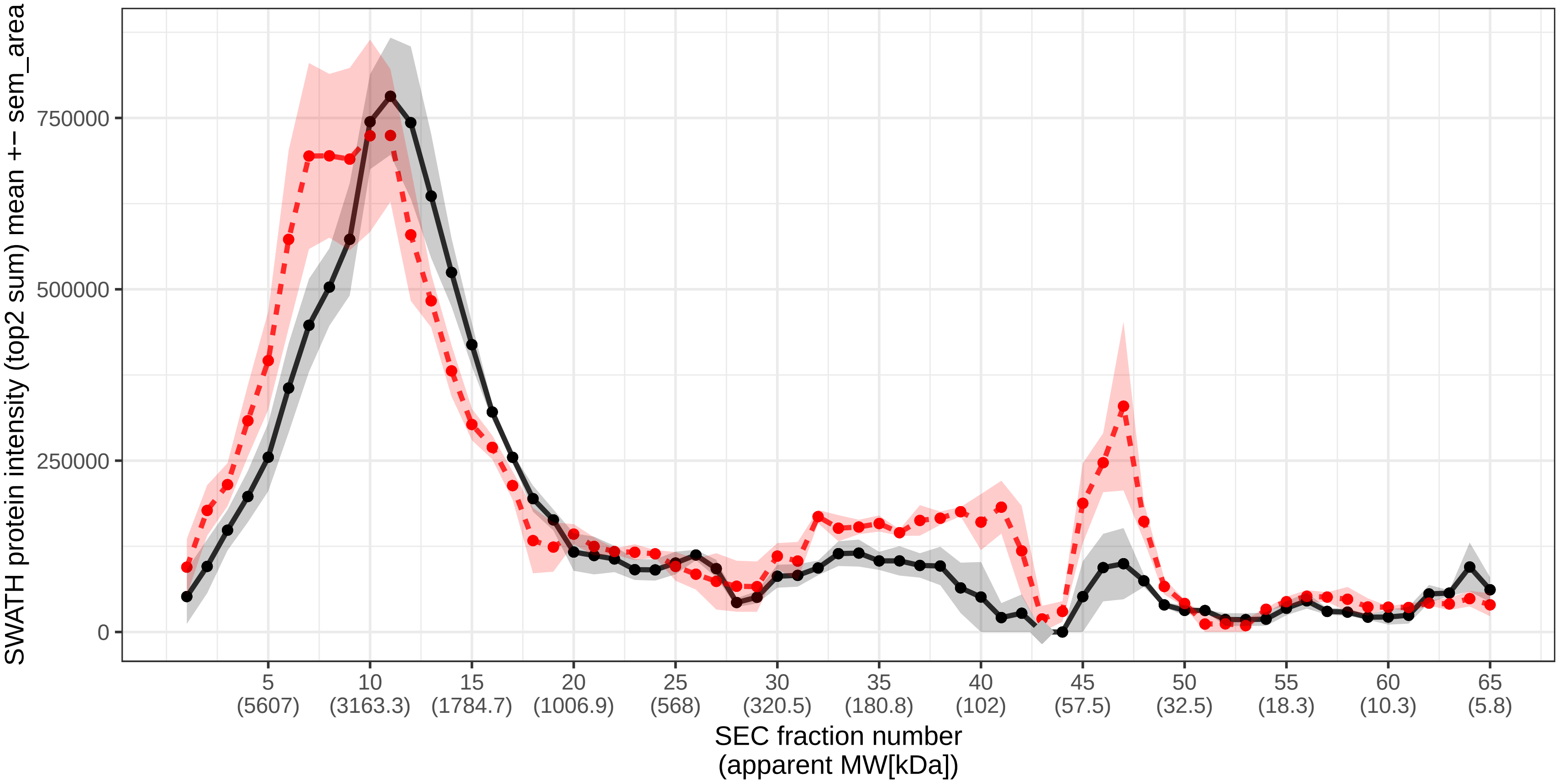

### SECchrom_O00139_KIF2A_HUMAN_KIF2A_KIF2_KNS2.pdf

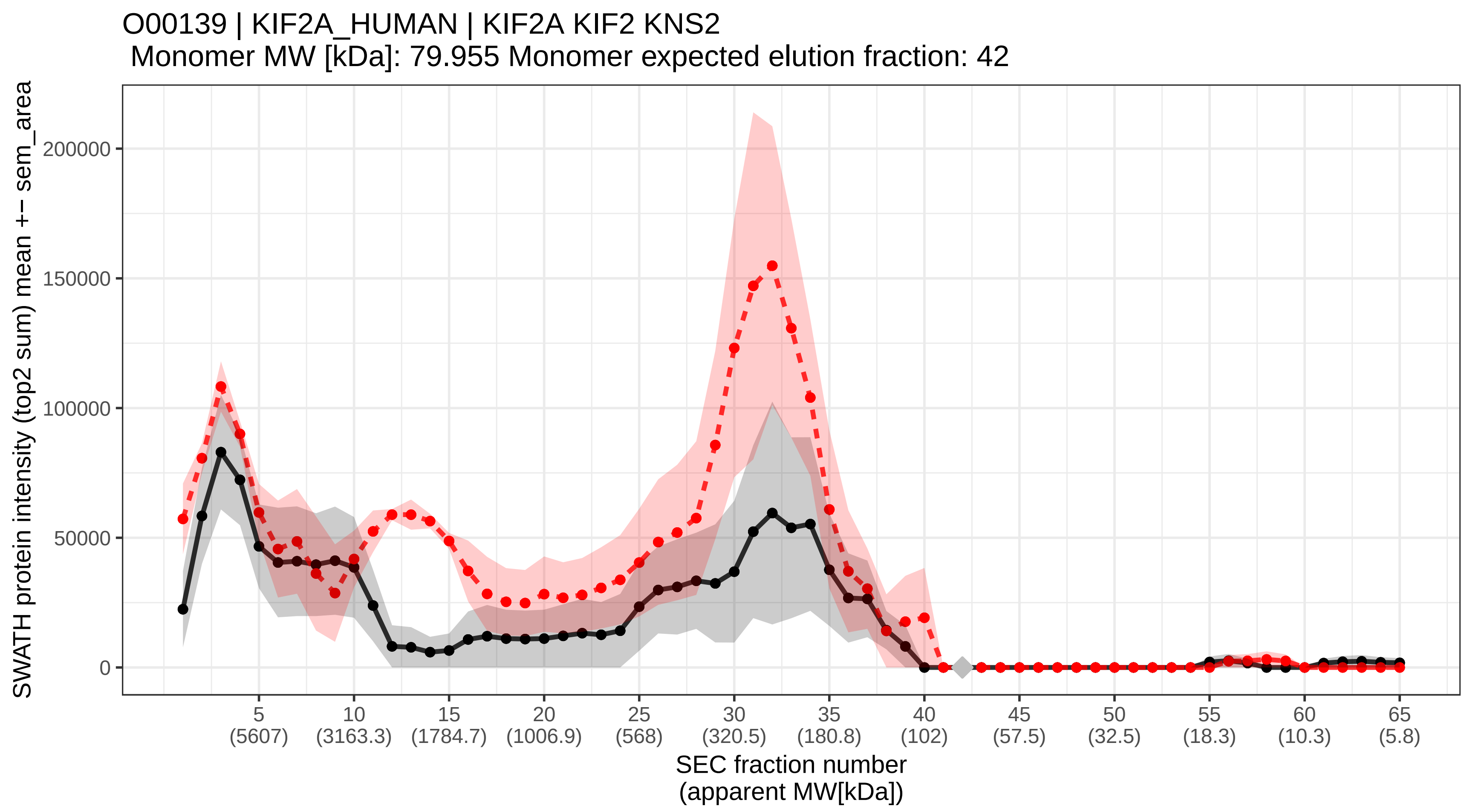

### SECchrom_O00148_DX39A_HUMAN_DDX39A_DDX39.pdf

#### O00148 | DX39A_HUMAN | DDX39A DDX39
Monomer MW [kDa]: 49.13 Monomer expected elution fraction: 46

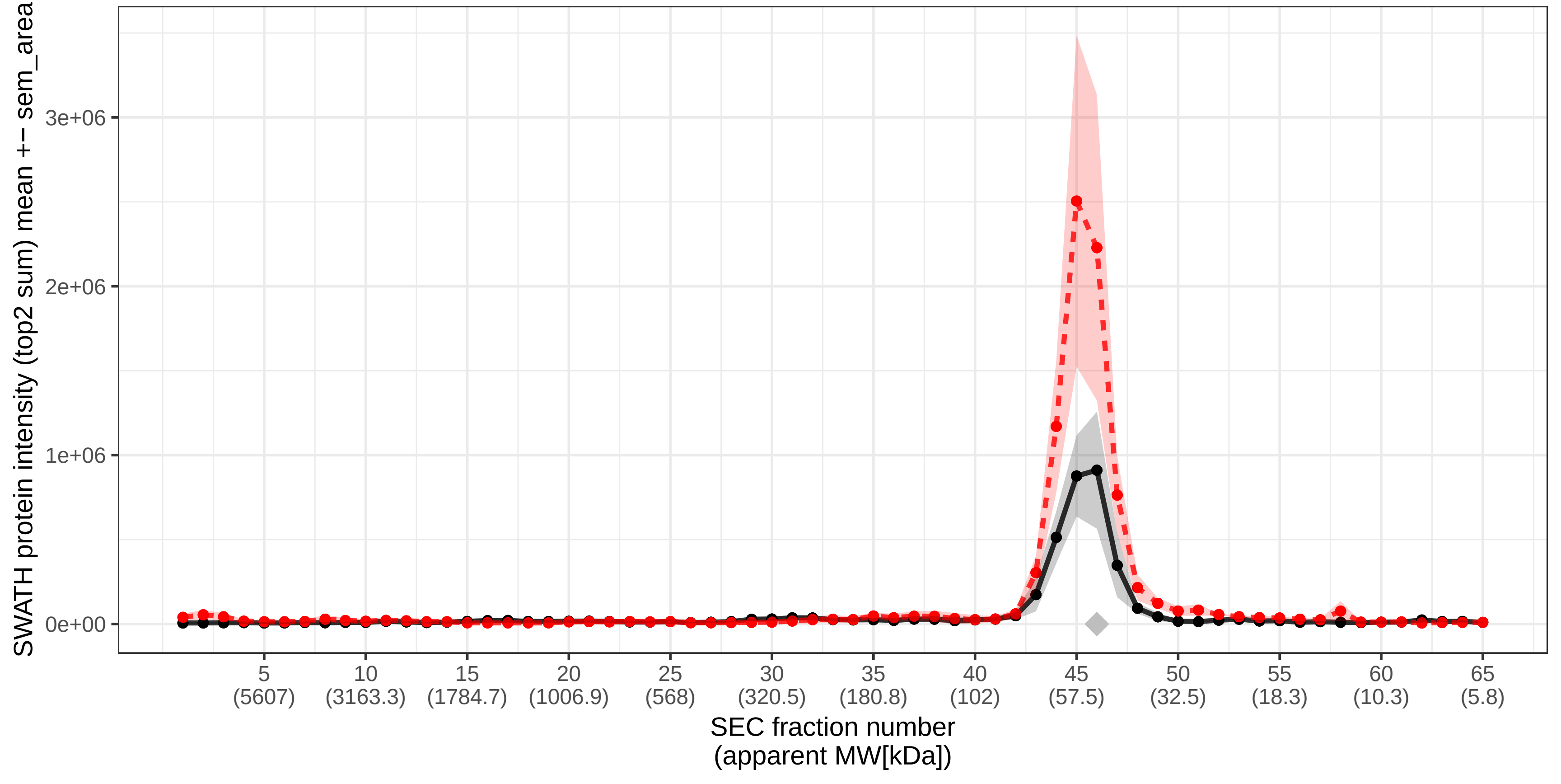

### SECchrom_O00151_PDLI1_HUMAN_PDLIM1_CLIM1_CLP36.pdf

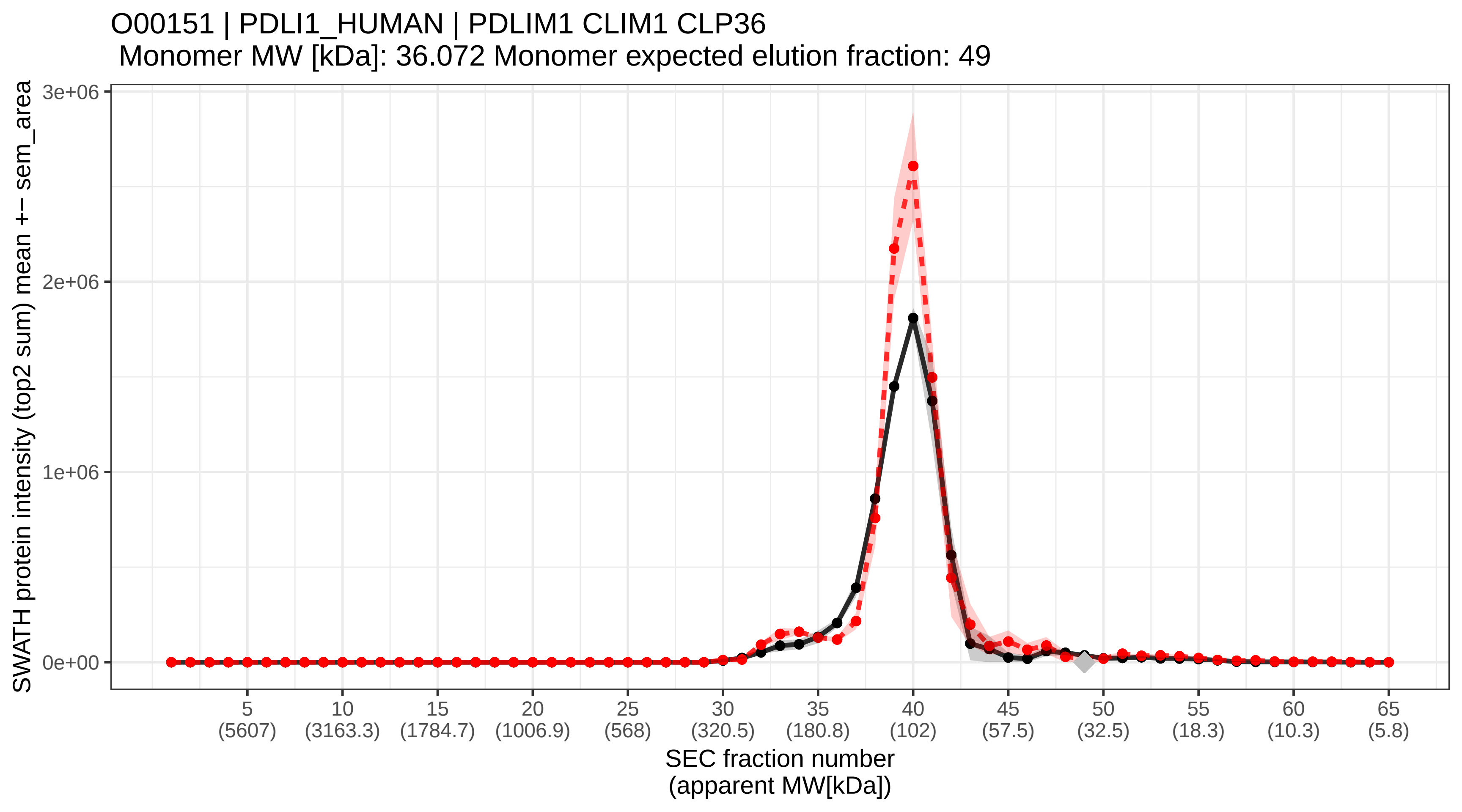

### SECchrom_O00154_BACH_HUMAN_ACOT7_BACH.pdf

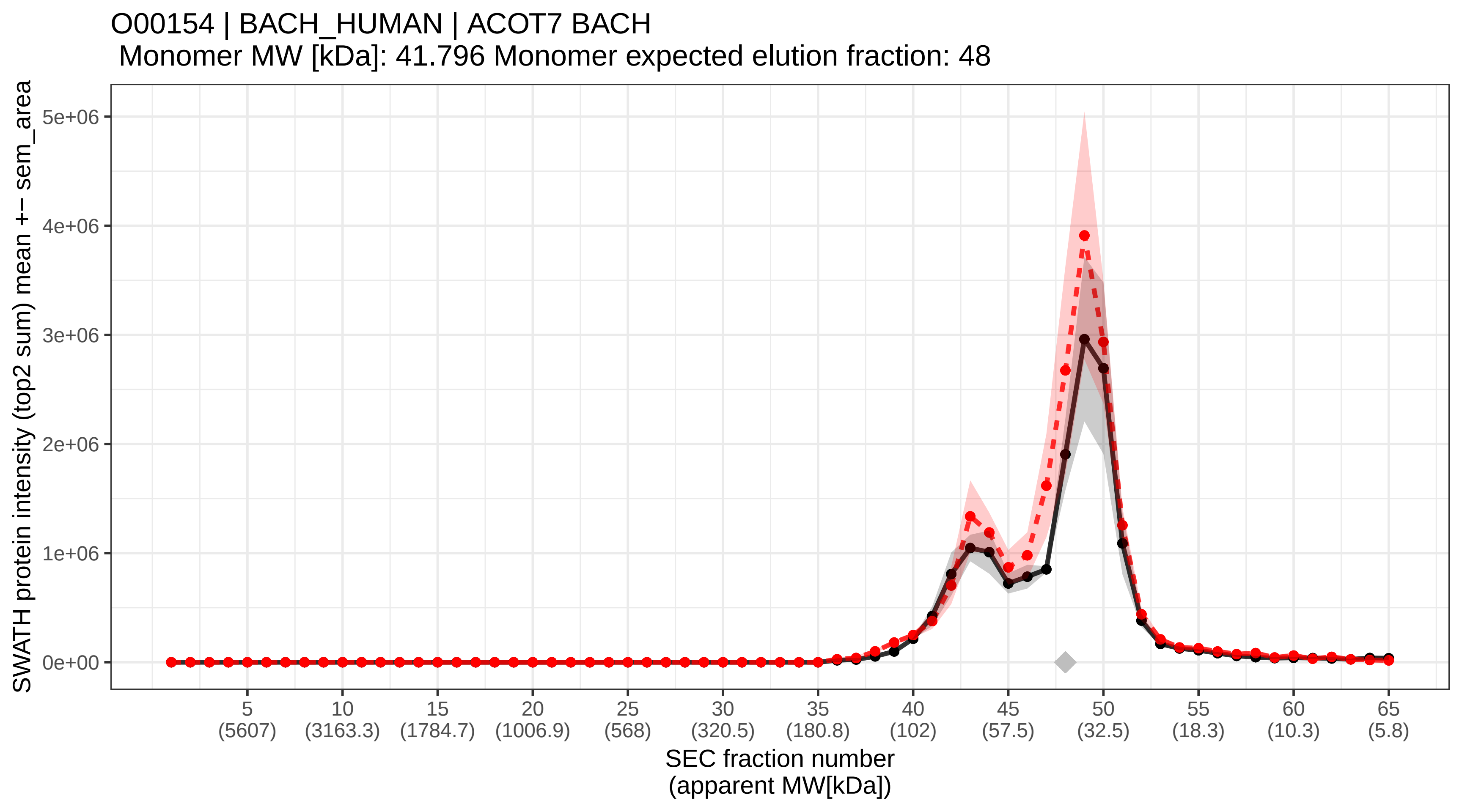

### SECchrom_O00159_MYO1C_HUMAN_MYO1C.pdf

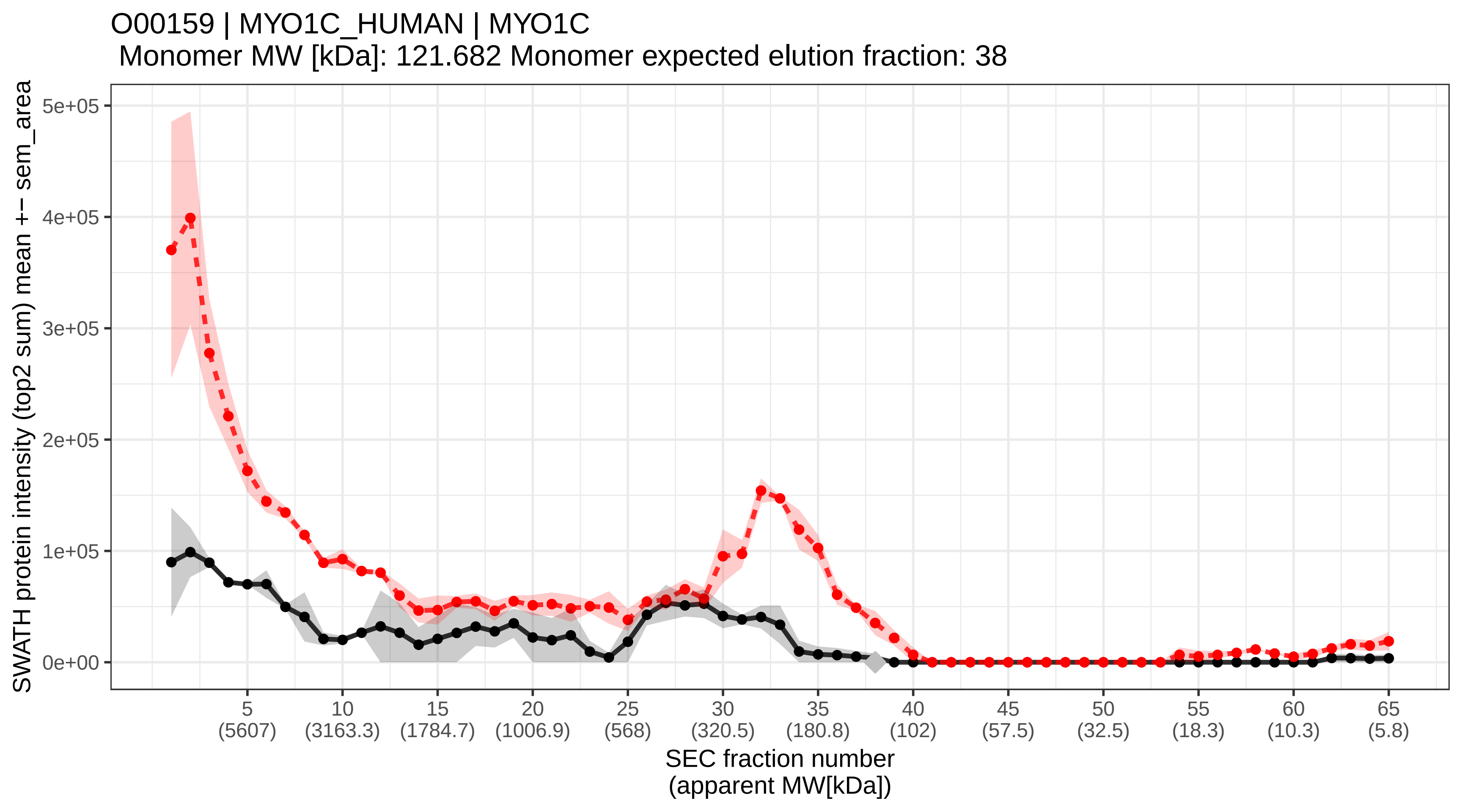

### SECchrom_O00161_SNP23_HUMAN_SNAP23.pdf

O00161 | SNP23_HUMAN | SNAP23
Monomer MW [kDa]: 23.354 Monomer expected elution fraction: 53

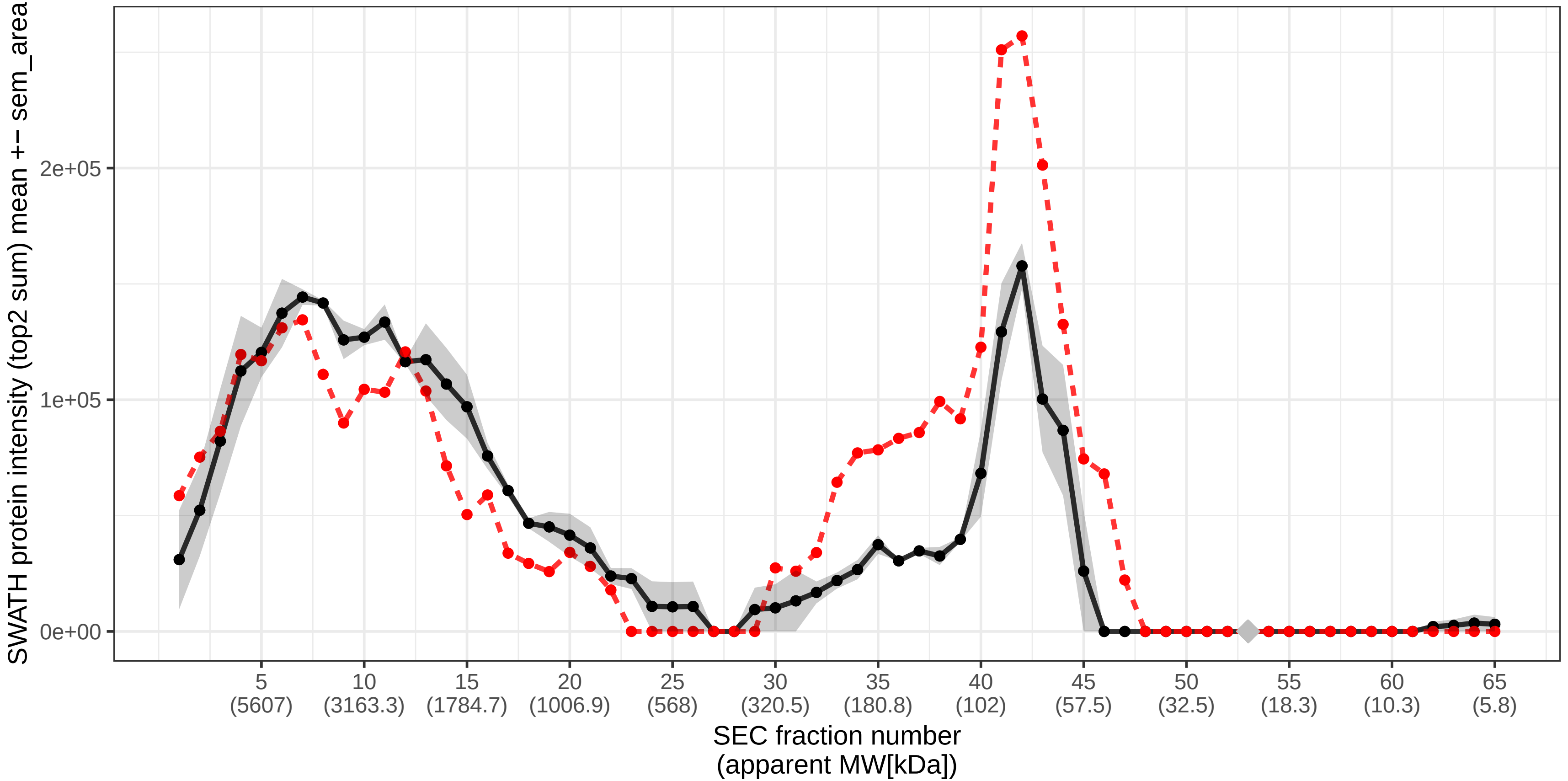

### SECchrom_O00165_HAX1_HUMAN_HAX1_HS1BP1.pdf

O00165 | HAX1_HUMAN | HAX1 HS1BP1
Monomer MW [kDa]: 31.621 Monomer expected elution fraction: 50

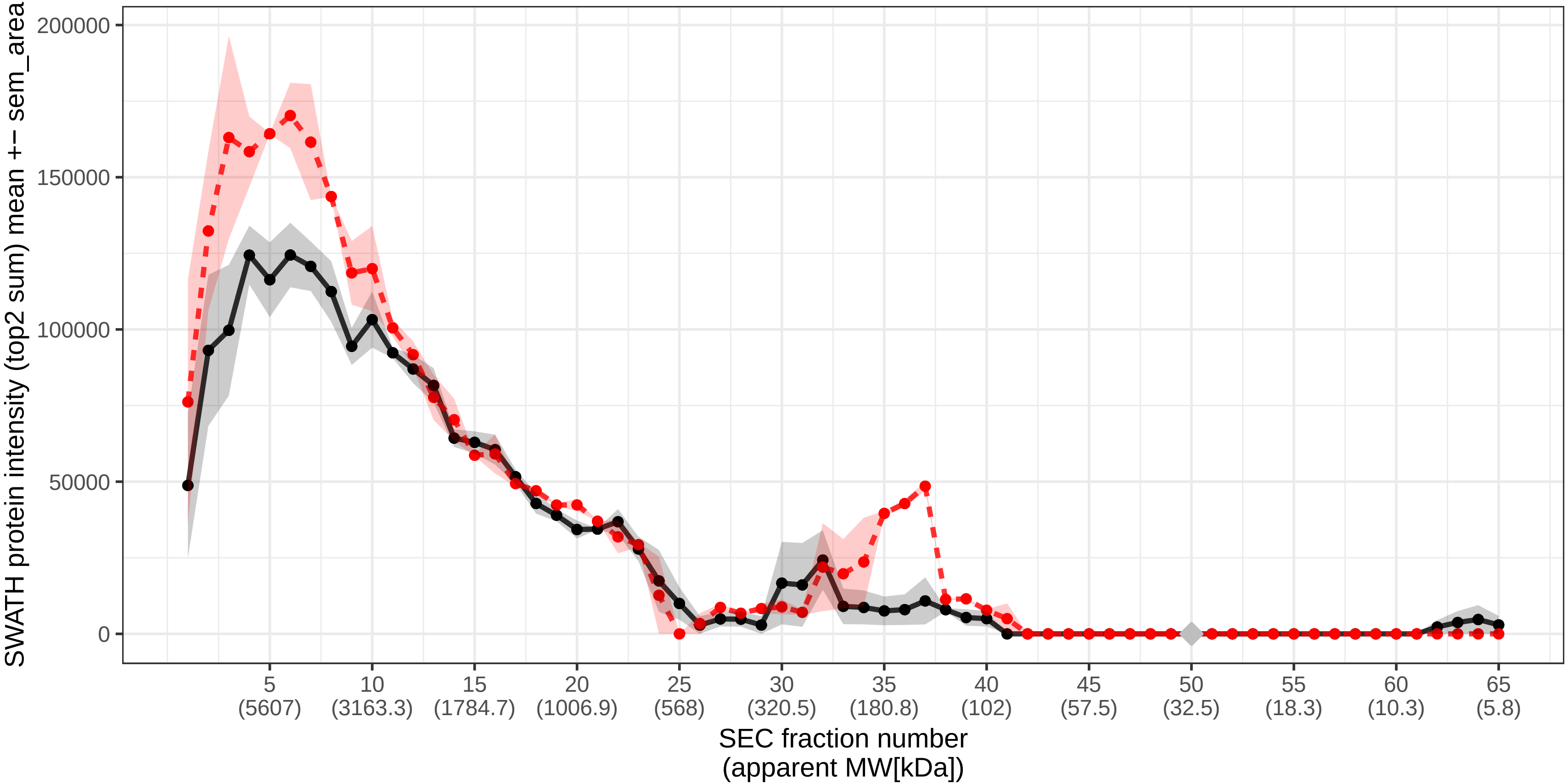

### SECchrom_O00170_AIP_HUMAN_AIP_XAP2.pdf

O00170 | AIP_HUMAN | AIP XAP2
Monomer MW [kDa]: 37.636 Monomer expected elution fraction: 49

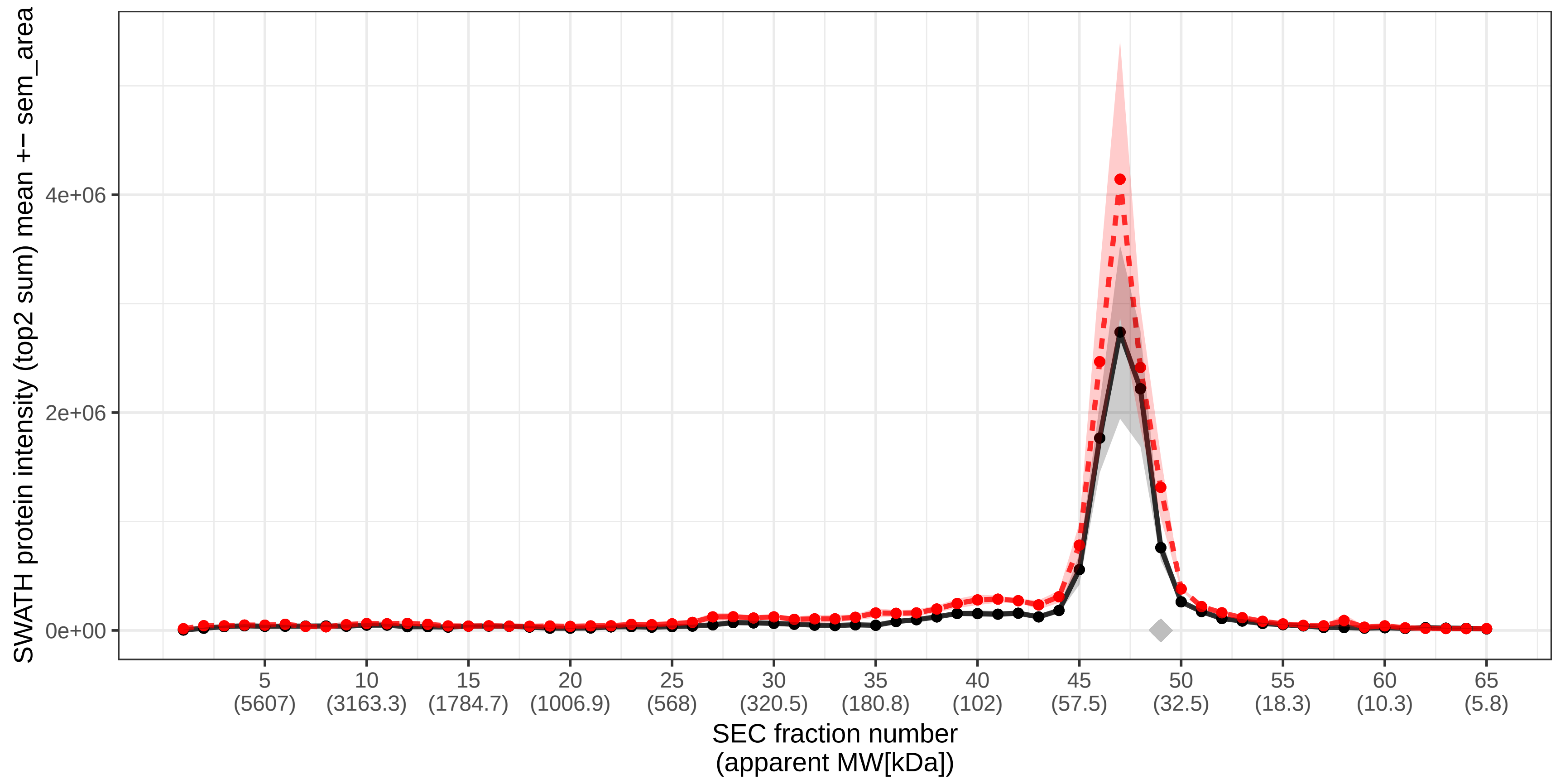

### SECchrom_O00178_GTPB1_HUMAN_GTPBP1.pdf

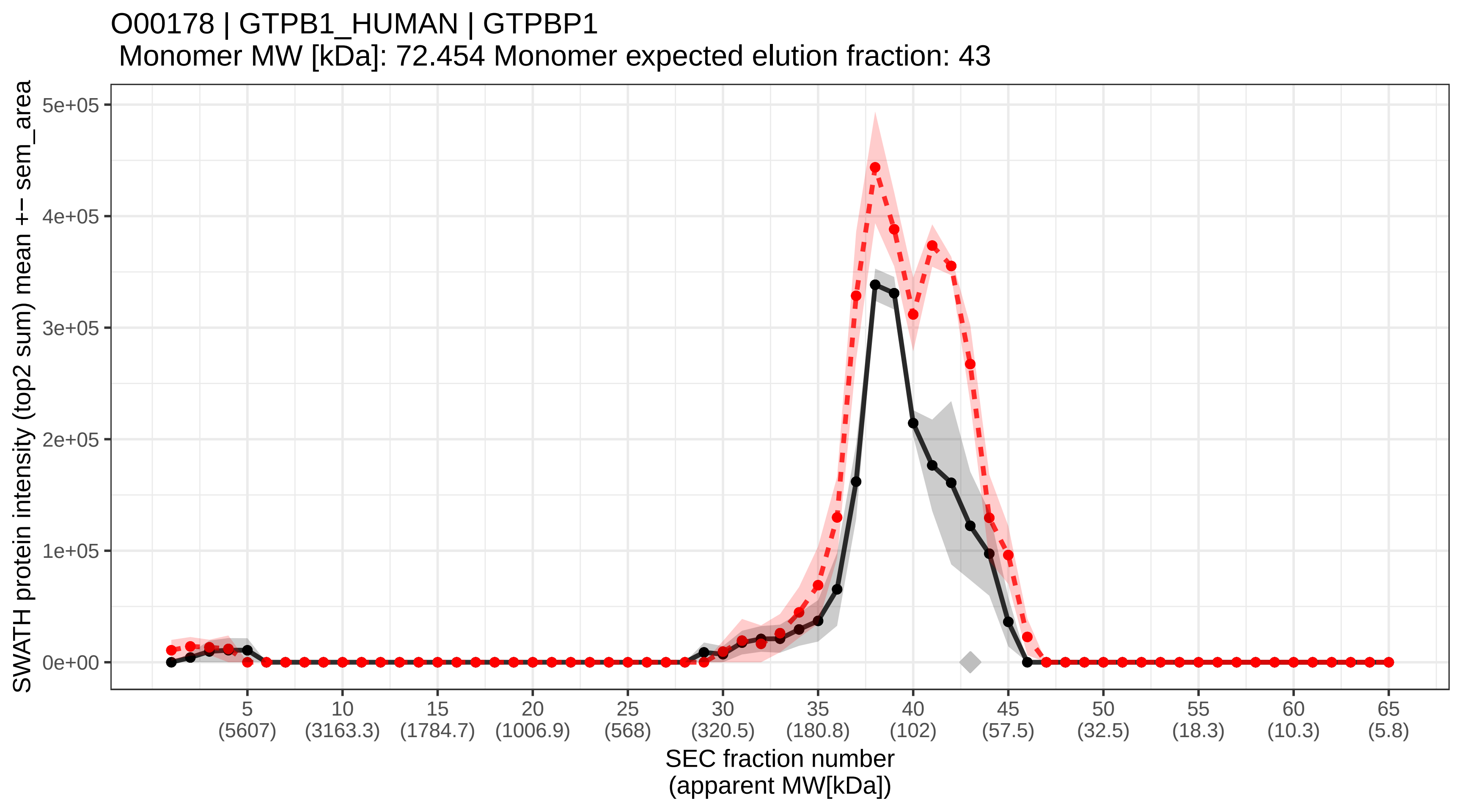

### SECchrom_O00186_STXB3_HUMAN_STXBP3.pdf

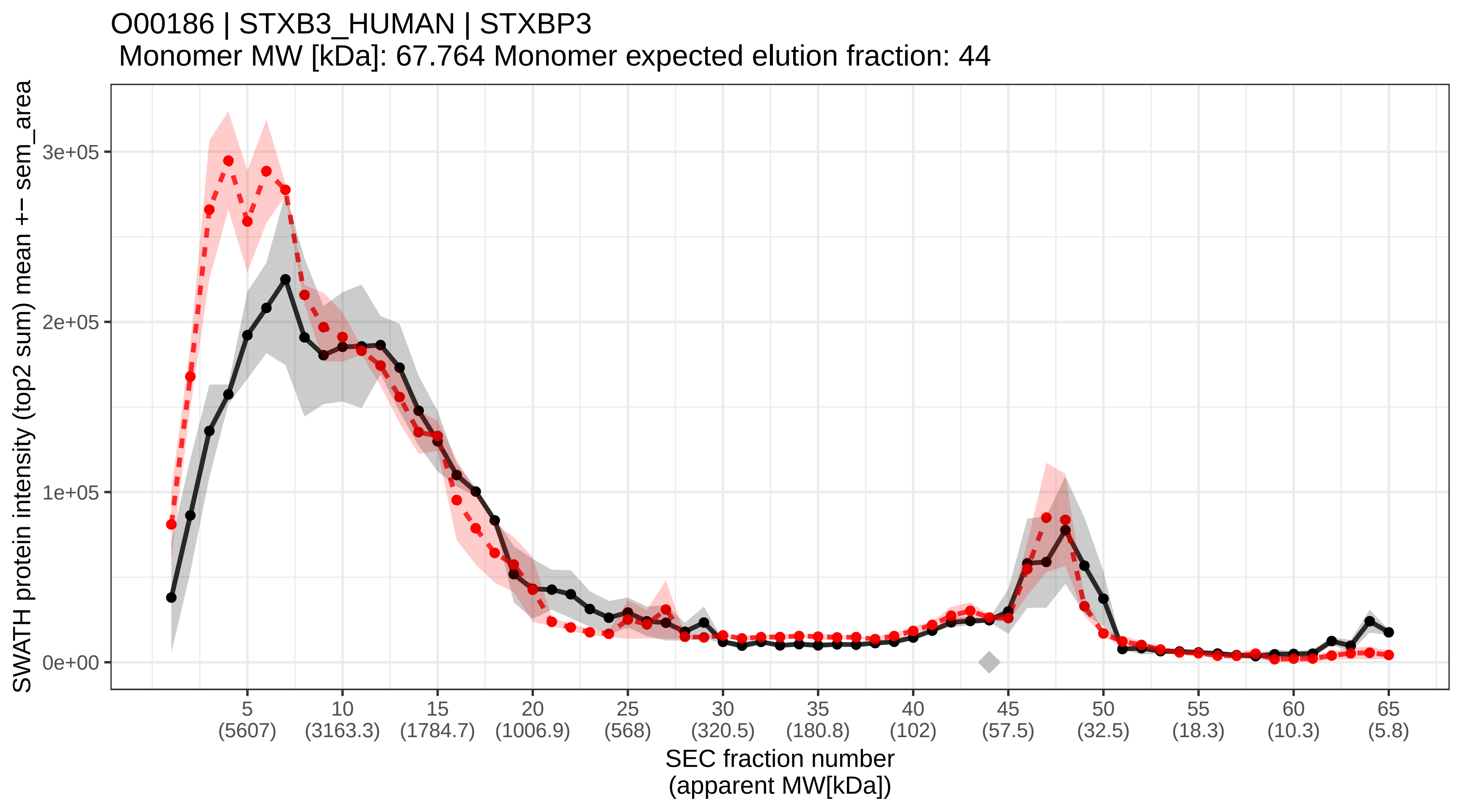

### SECchrom_O00189_AP4M1_HUMAN_AP4M1_MUARP2.pdf

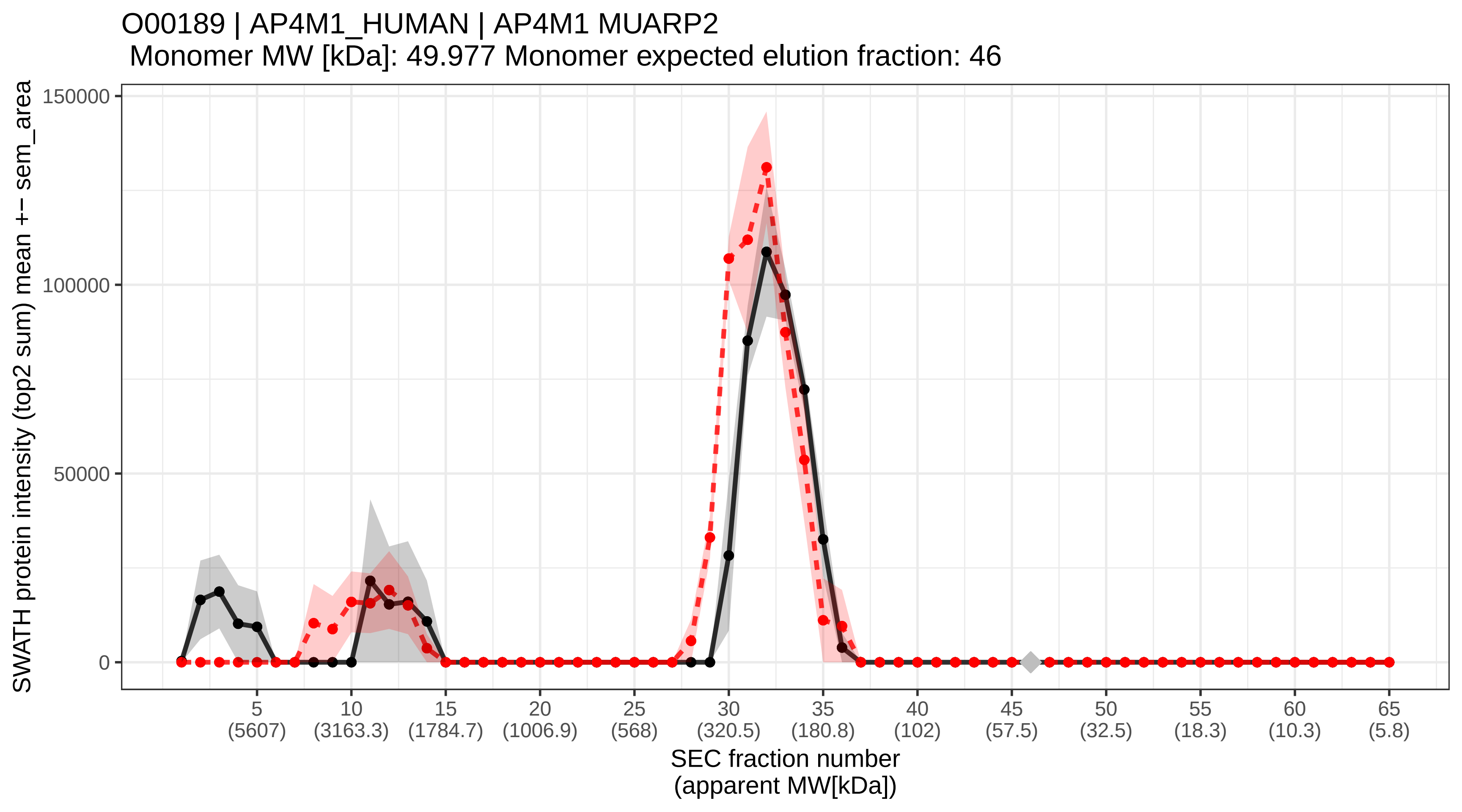

### SECchrom_O00193_SMAP_HUMAN_SMAP_C11orf58.pdf

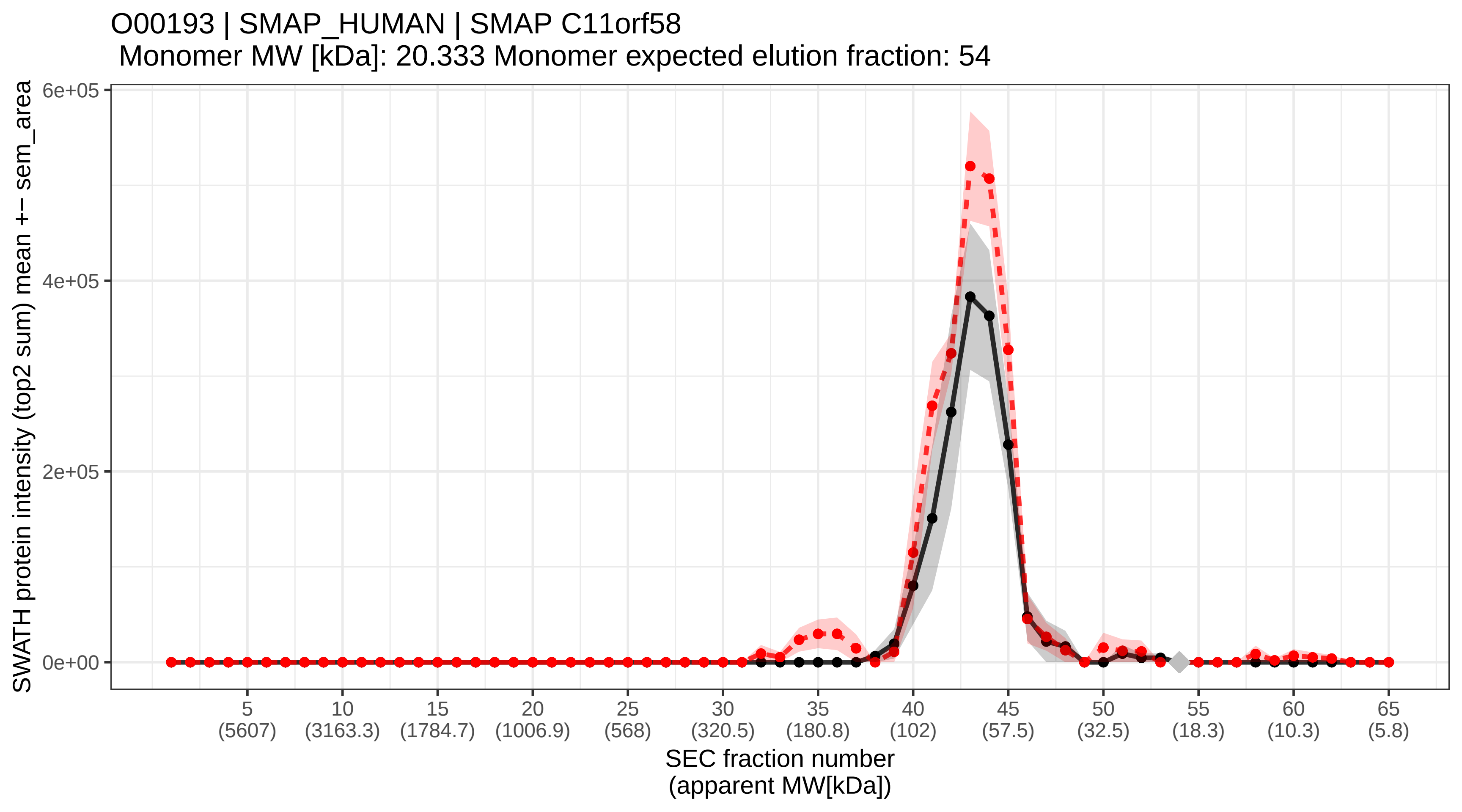

### SECchrom_O00203_AP3B1_HUMAN_AP3B1_ADTB3A.pdf

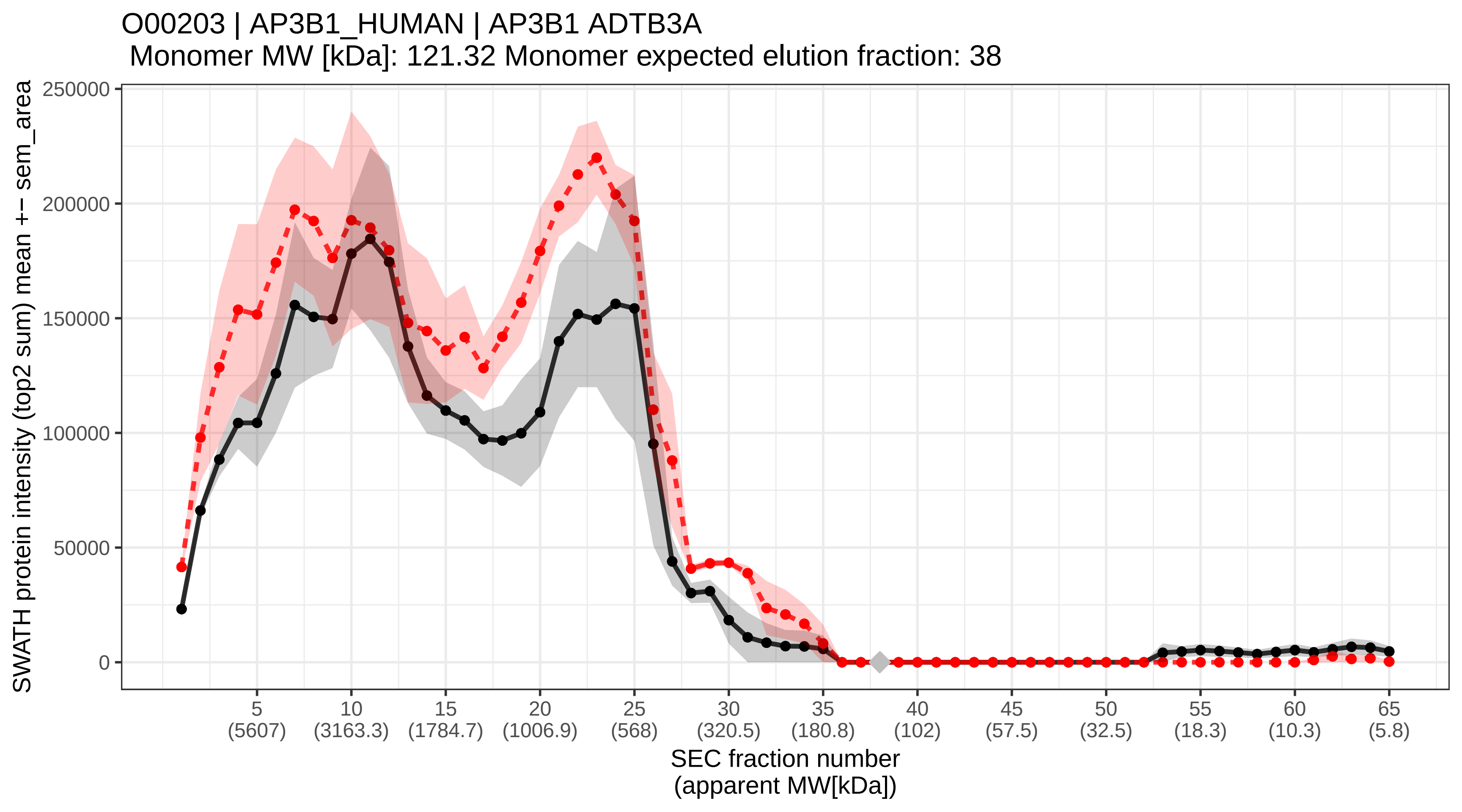

### SECchrom_O00213_APBB1_HUMAN_APBB1_FE65_RIR.pdf

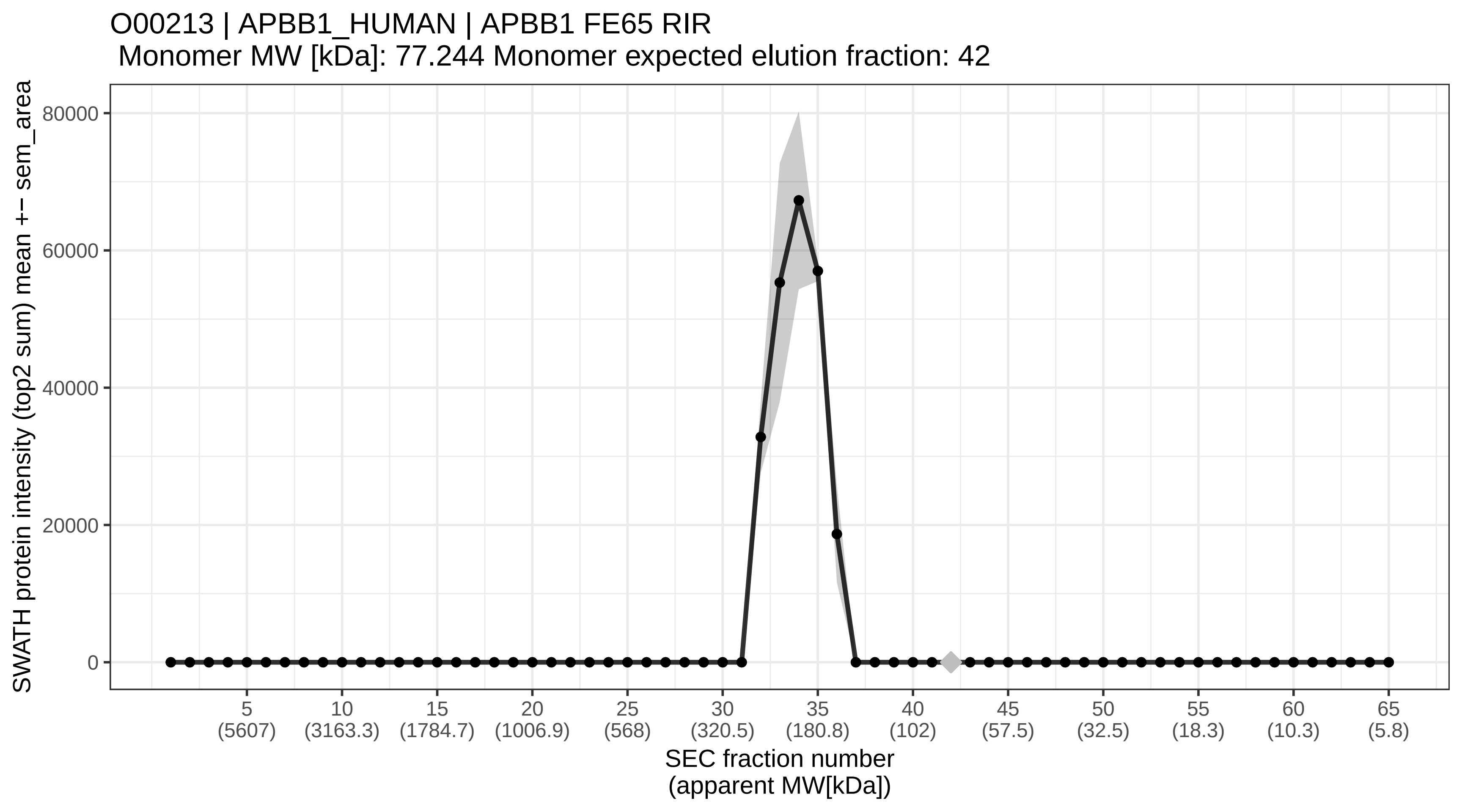

### SECchrom_O00217_NDUS8_HUMAN_NDUFS8.pdf

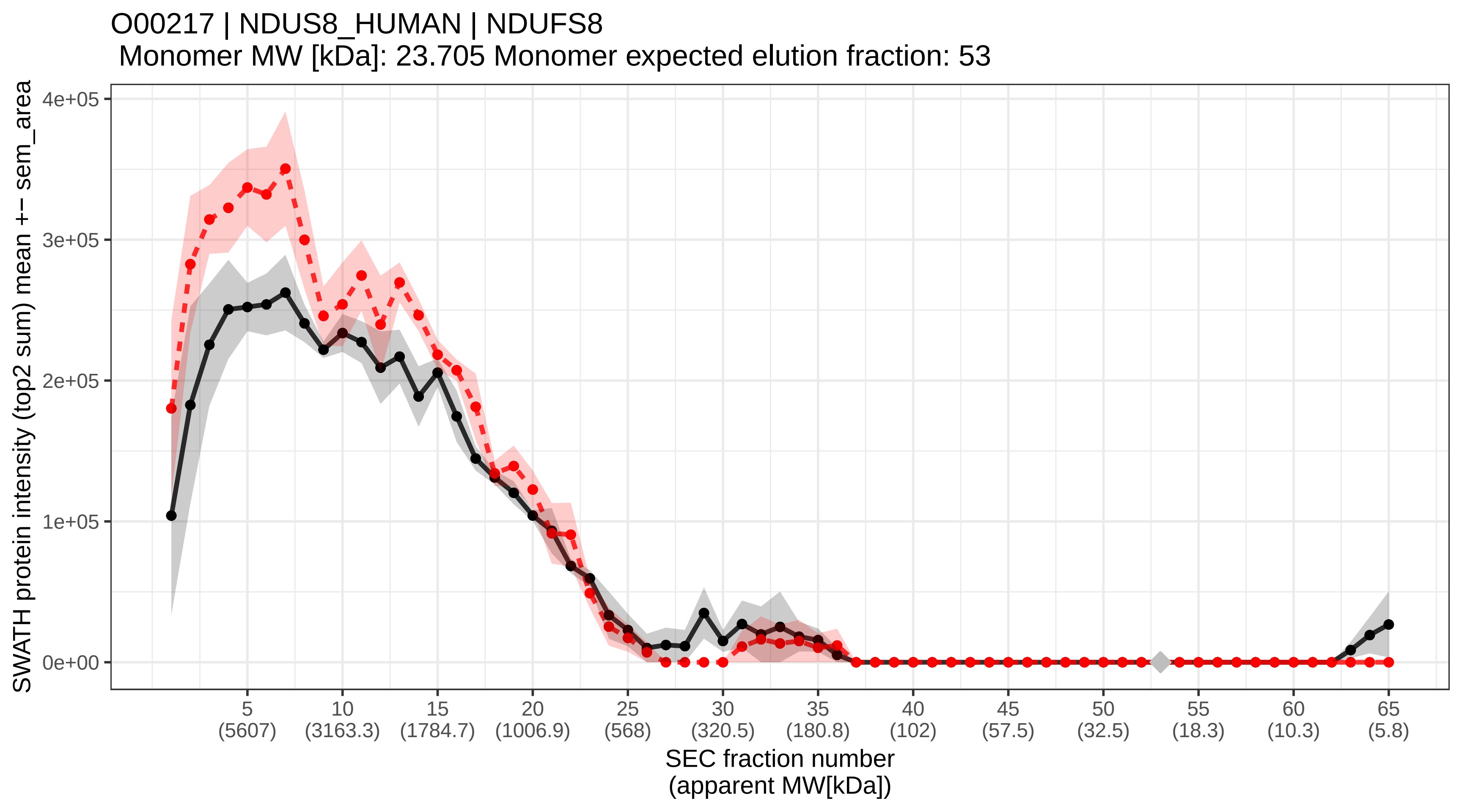

### SECchrom_O00220_TR10A_HUMAN_TNFRSF10A_APO2_DR4_TRAILR1.pdf

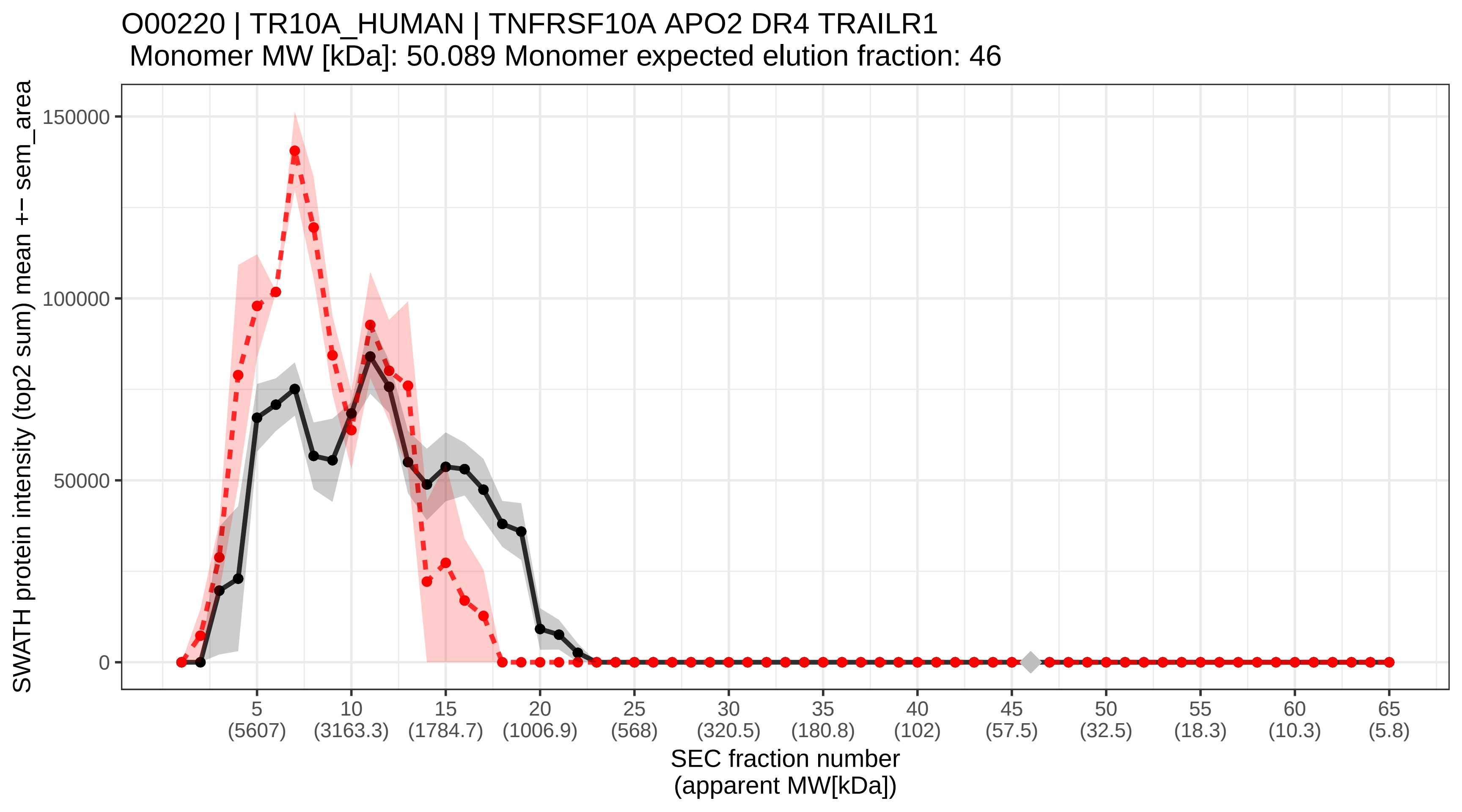

### SECchrom_O00221_IKBE_HUMAN_NFKBIE_IKBE.pdf

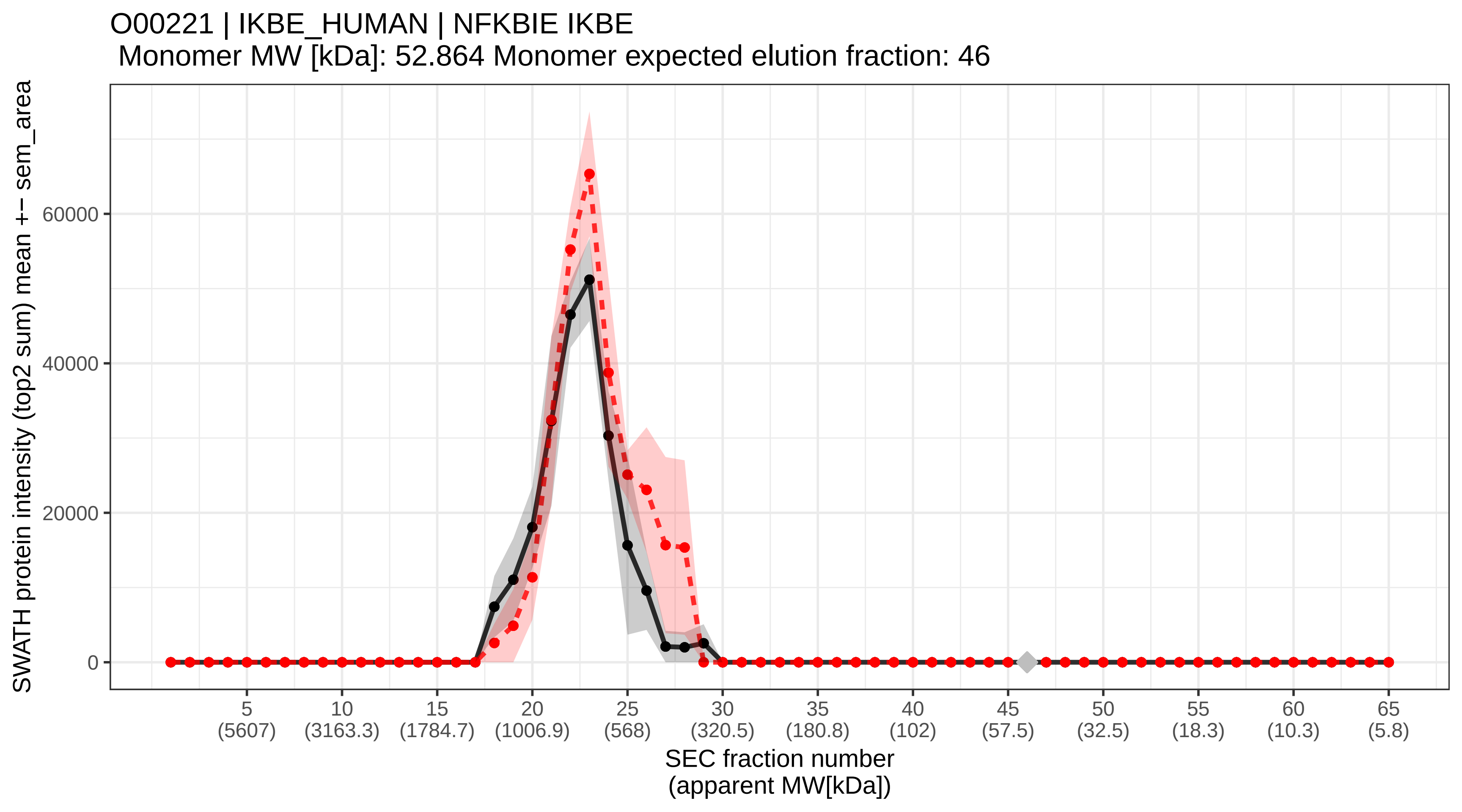

### SECchrom_O00231_PSD11_HUMAN_PSMD11.pdf

O00231 | PSD11_HUMAN | PSMD11
Monomer MW [kDa]: 47.464 Monomer expected elution fraction: 47

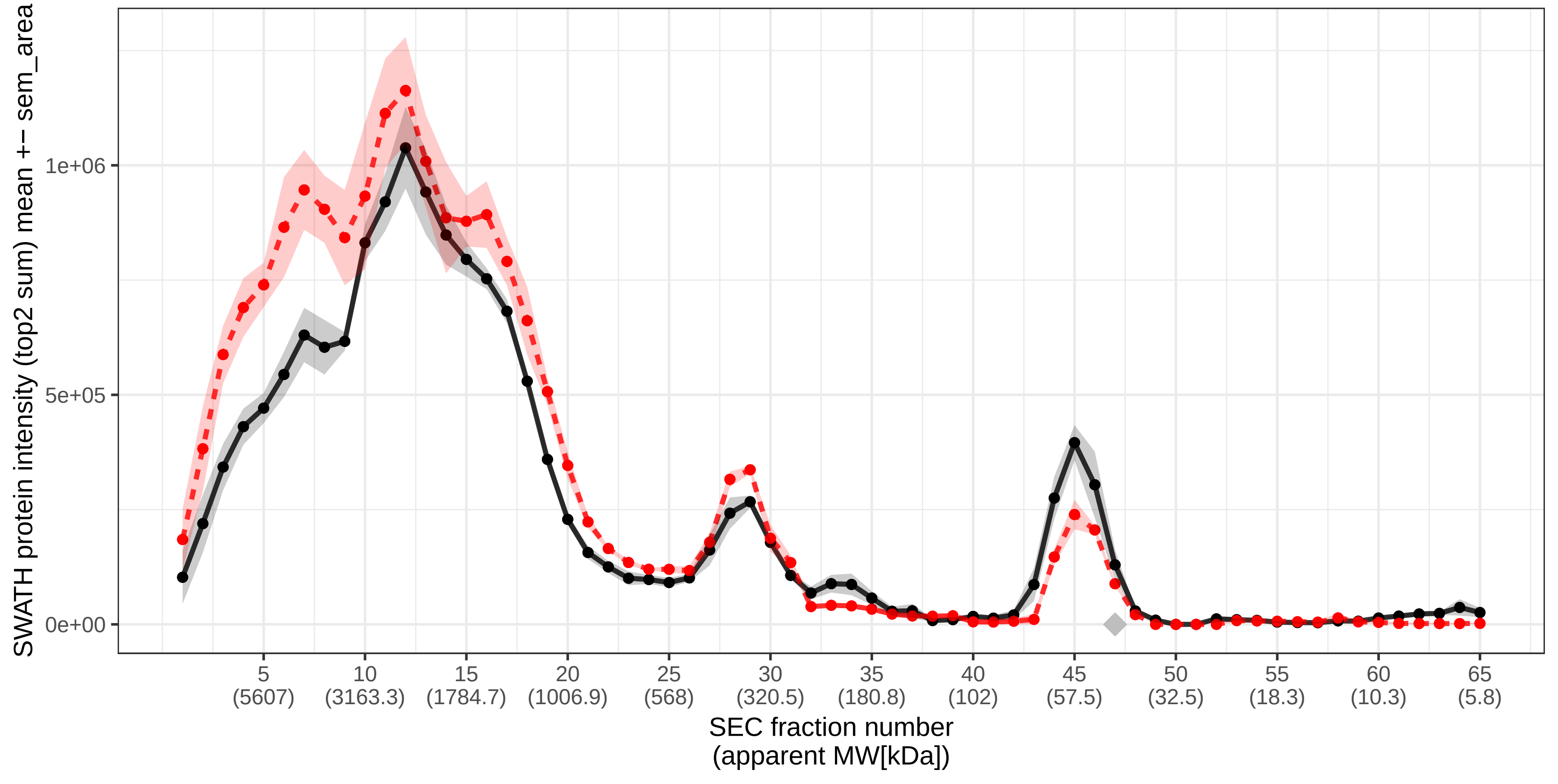

### SECchrom_O00232_PSD12_HUMAN_PSMD12.pdf

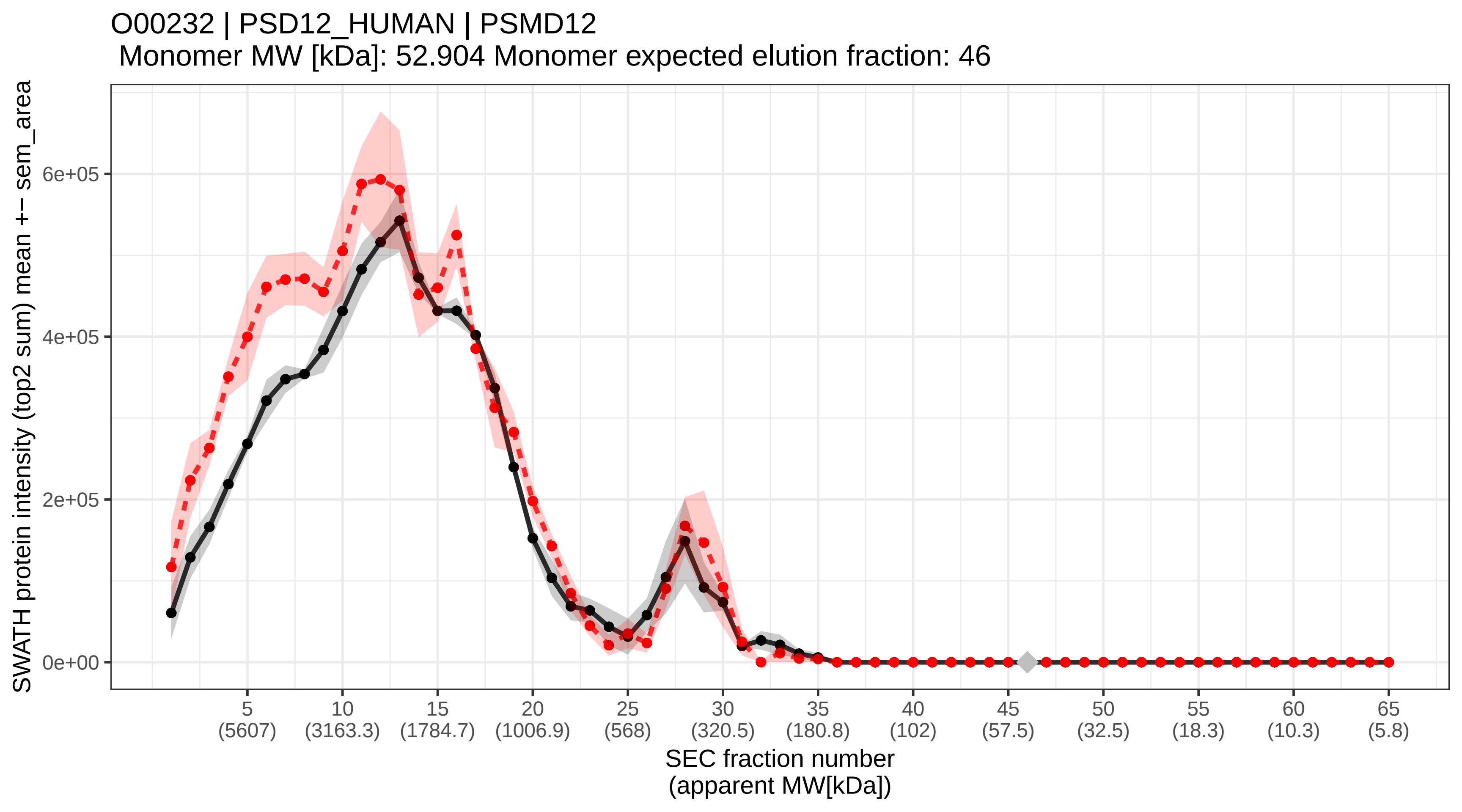

### SECchrom_O00233_PSMD9_HUMAN_PSMD9.pdf

O00233 | PSMD9_HUMAN | PSMD9
Monomer MW [kDa]: 24.682 Monomer expected elution fraction: 52

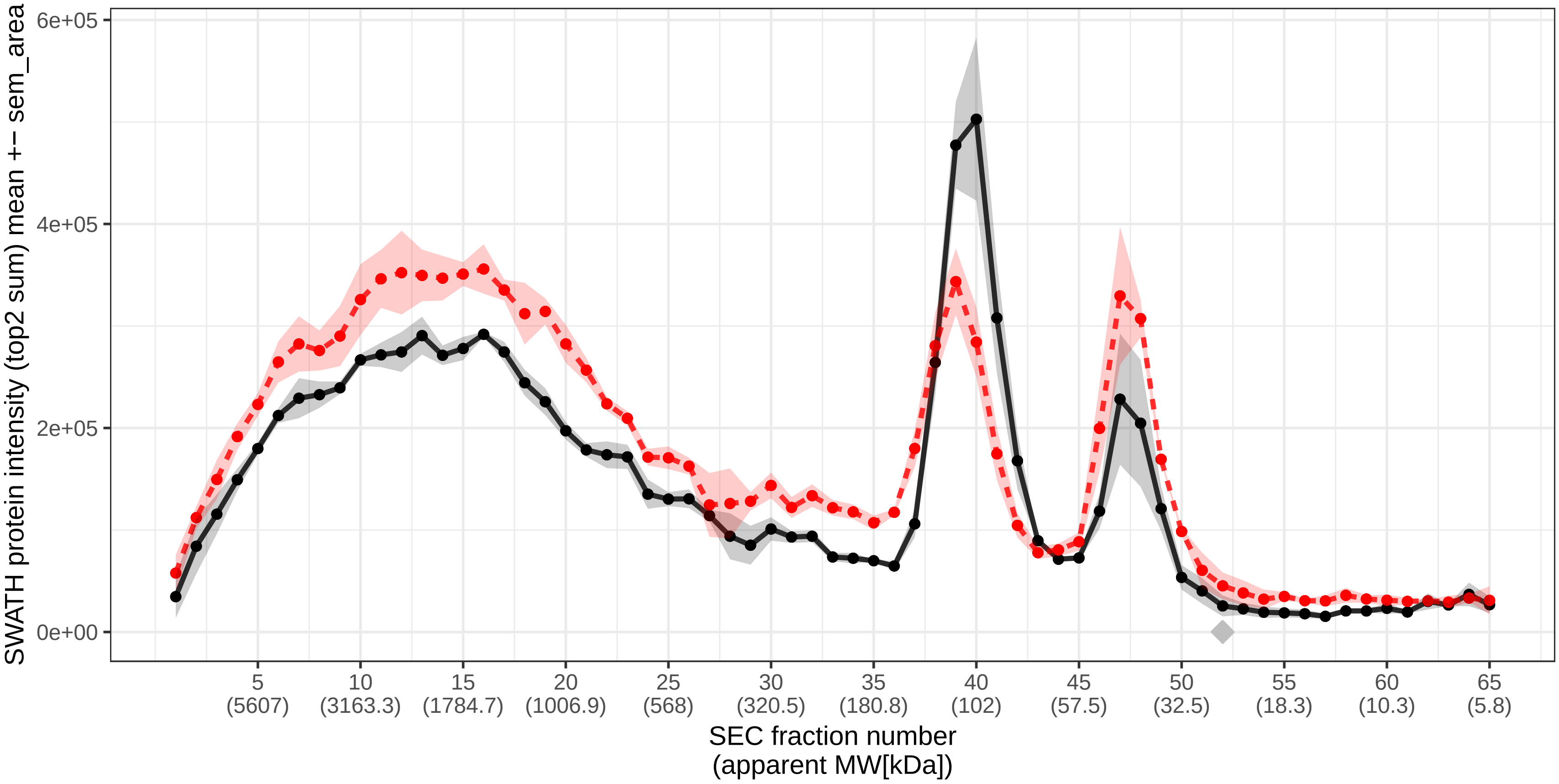

### SECchrom_O00244_ATOX1_HUMAN_ATOX1_HAH1.pdf

O00244 | ATOX1_HUMAN | ATOX1 HAH1
Monomer MW [kDa]: 7.402 Monomer expected elution fraction: 63

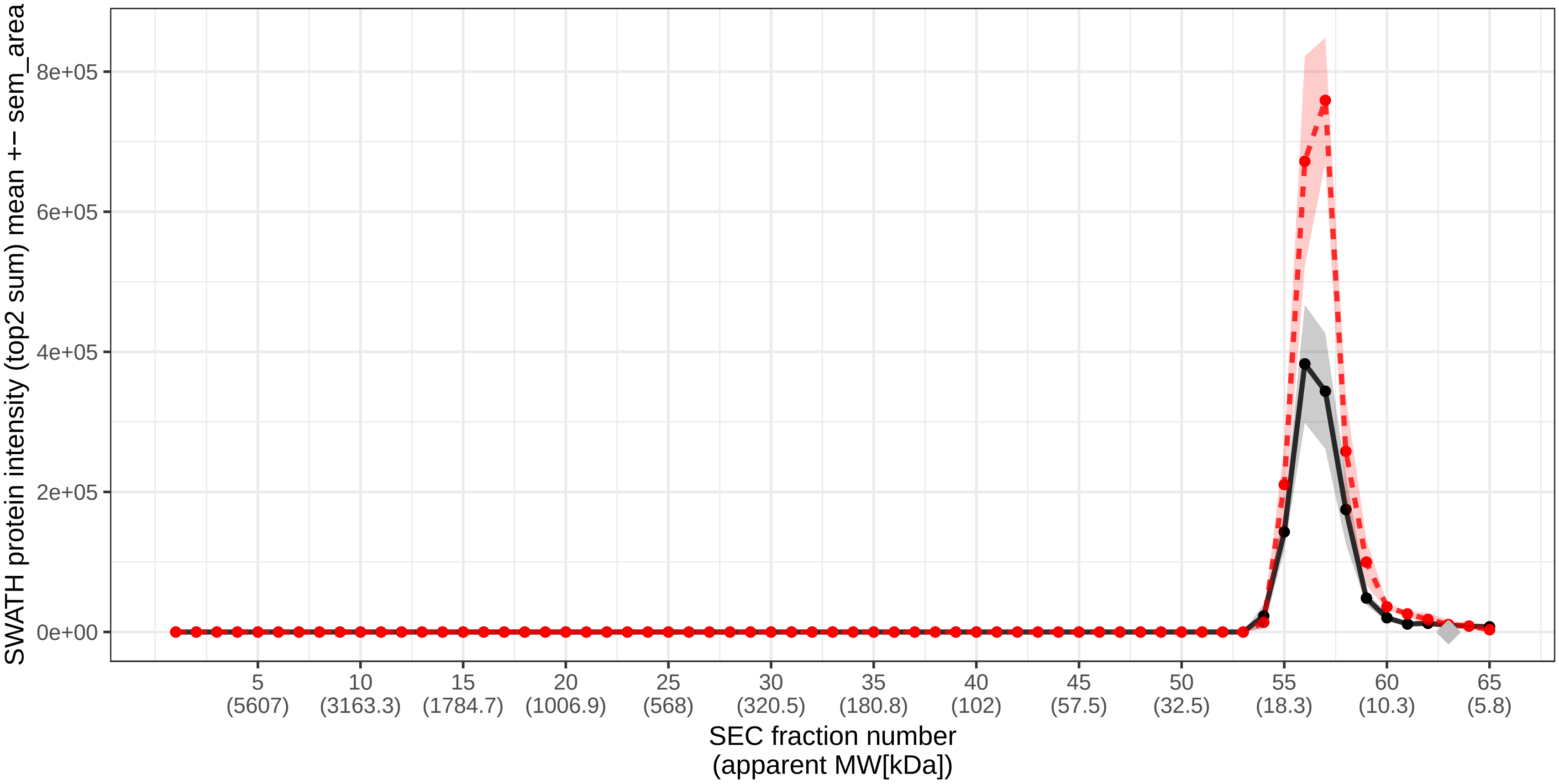

### SECchrom_O00264_PGRC1_HUMAN_PGRMC1_HPR6.6_PGRMC.pdf

O00264 | PGRC1_HUMAN | PGRMC1 HPR6.6 PGRMC
Monomer MW [kDa]: 21.671 Monomer expected elution fraction: 54

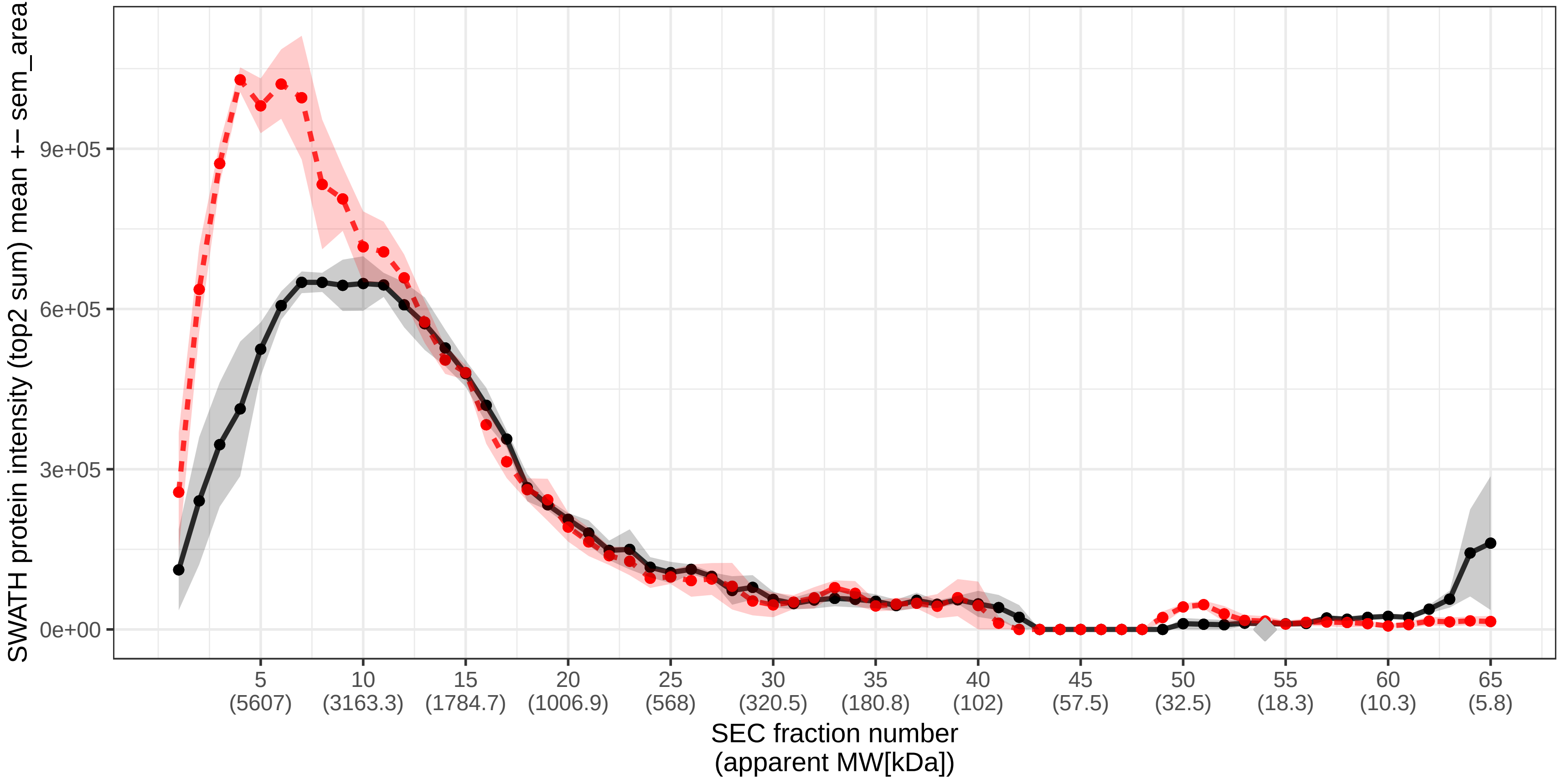

### SECchrom_O00267_SPT5H_HUMAN_SUPT5H_SPT5_SPT5H.pdf

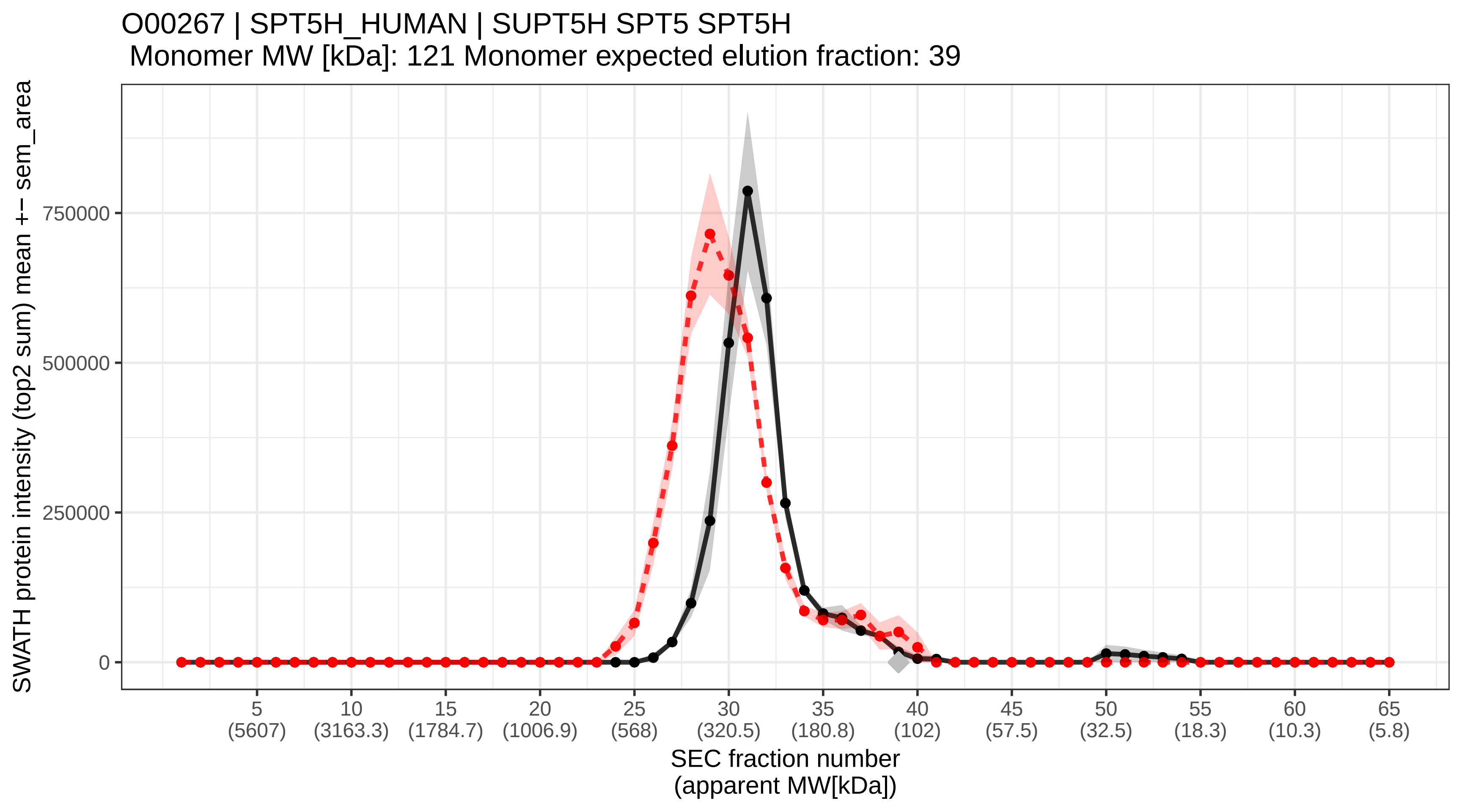

### SECchrom_O00268_TAF4_HUMAN_TAF4_TAF2C_TAF2C1_TAF4A_TAFII130.pdf

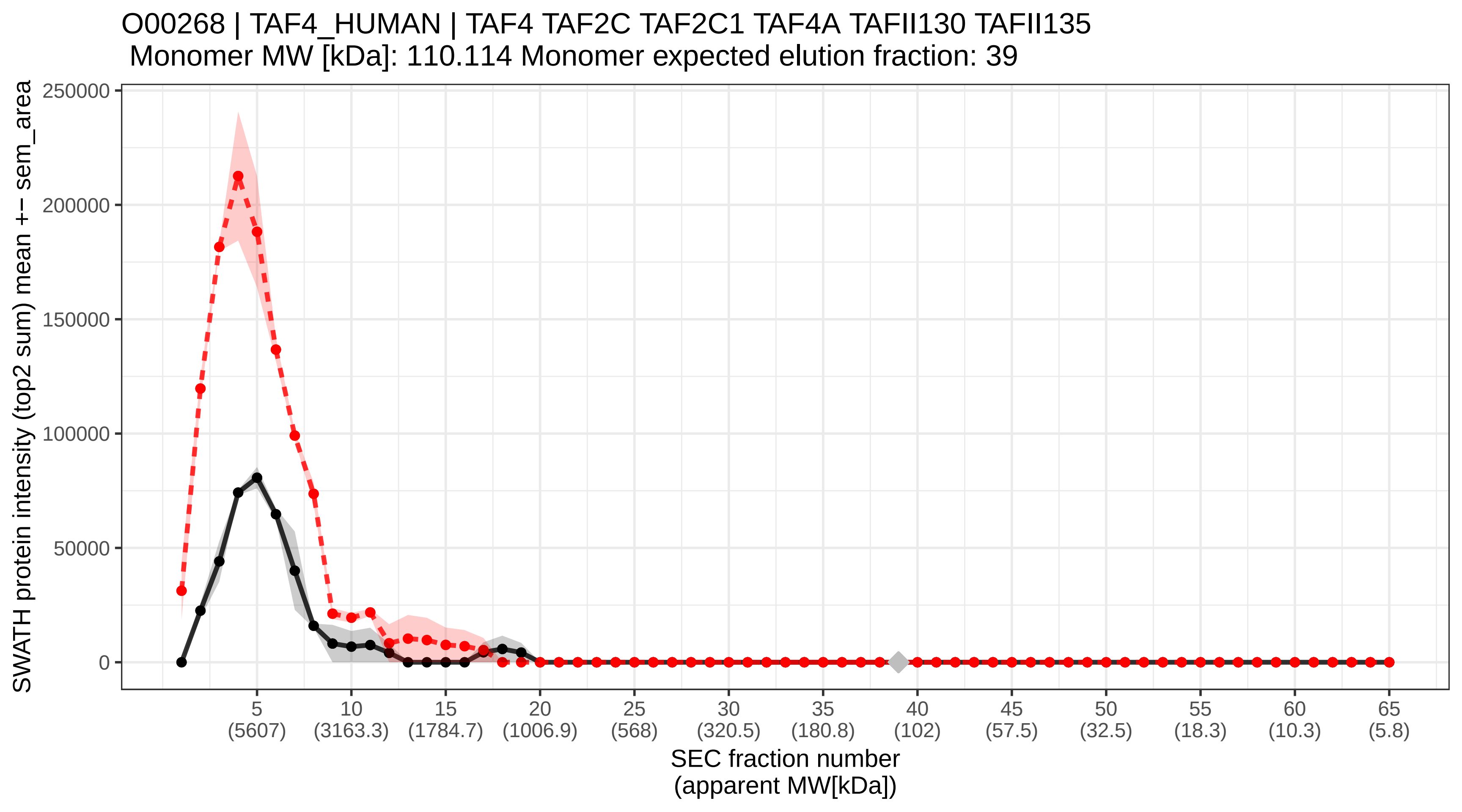

### SECchrom_O00273_DFFA_HUMAN_DFFA_DFF1_DFF45_H13.pdf

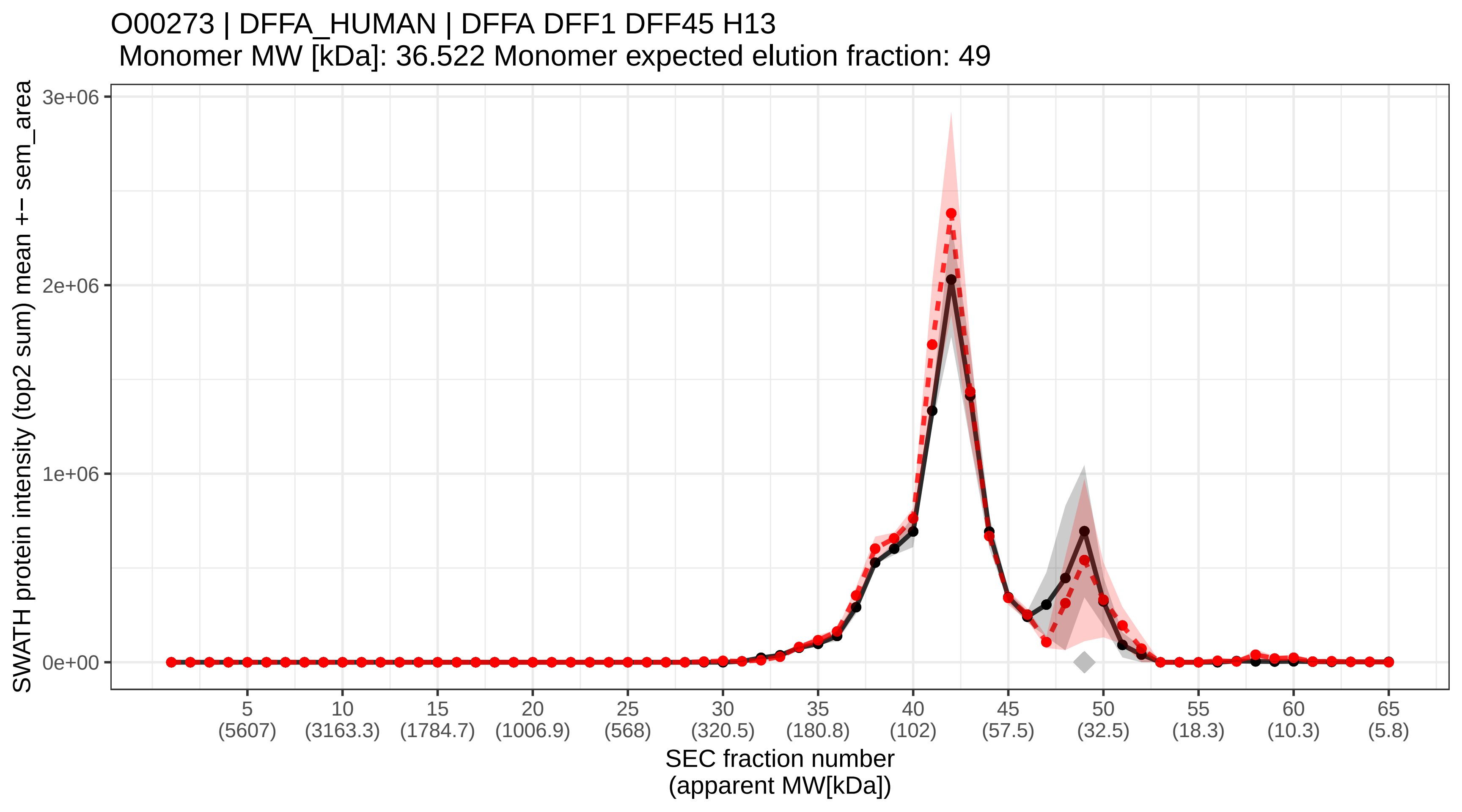

### SECchrom_O00287_RFXAP_HUMAN_RFXAP.pdf

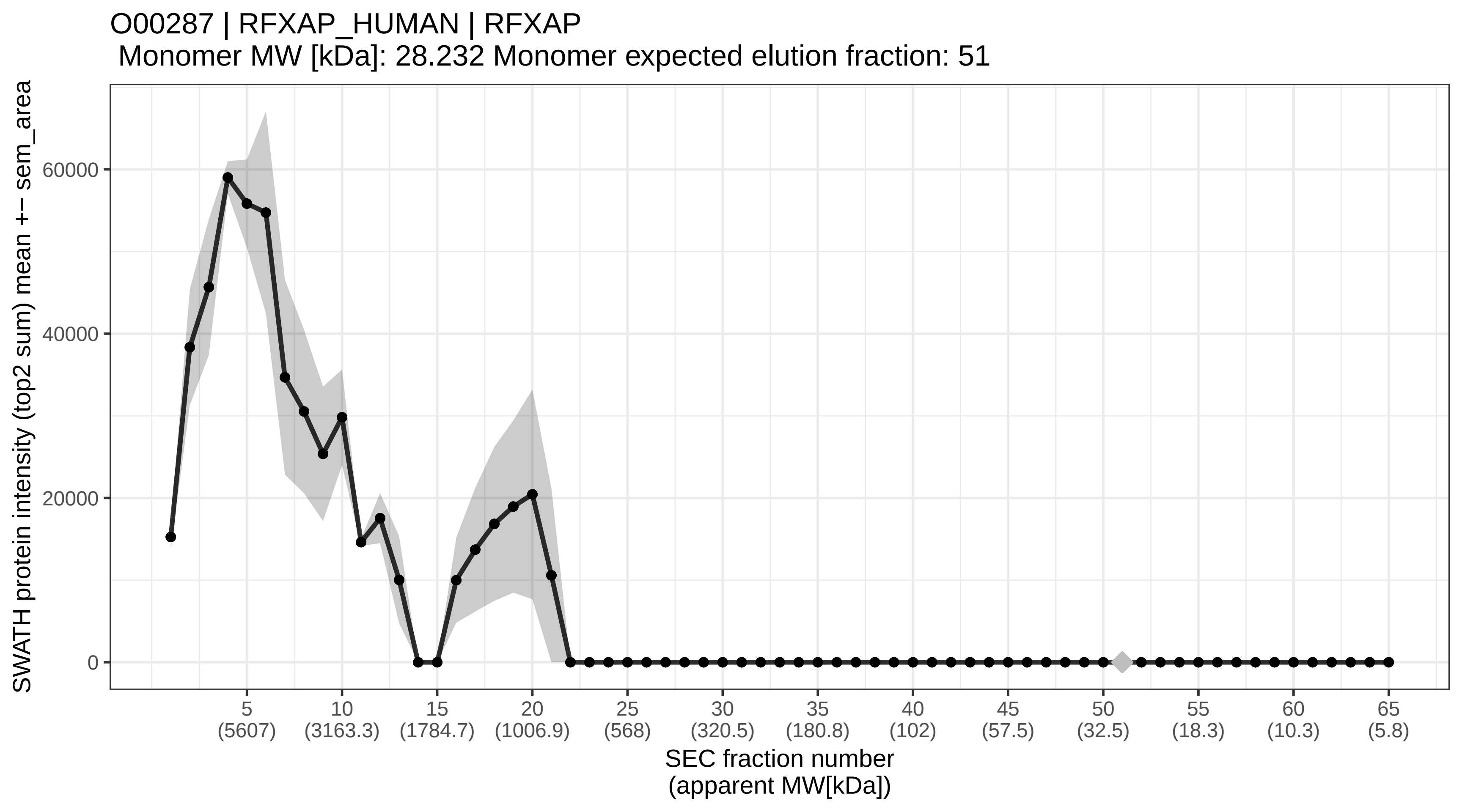

### SECchrom_O00291_HIP1_HUMAN_HIP1.pdf

O00291 | HIP1_HUMAN | HIP1
Monomer MW [kDa]: 116.221 Monomer expected elution fraction: 39

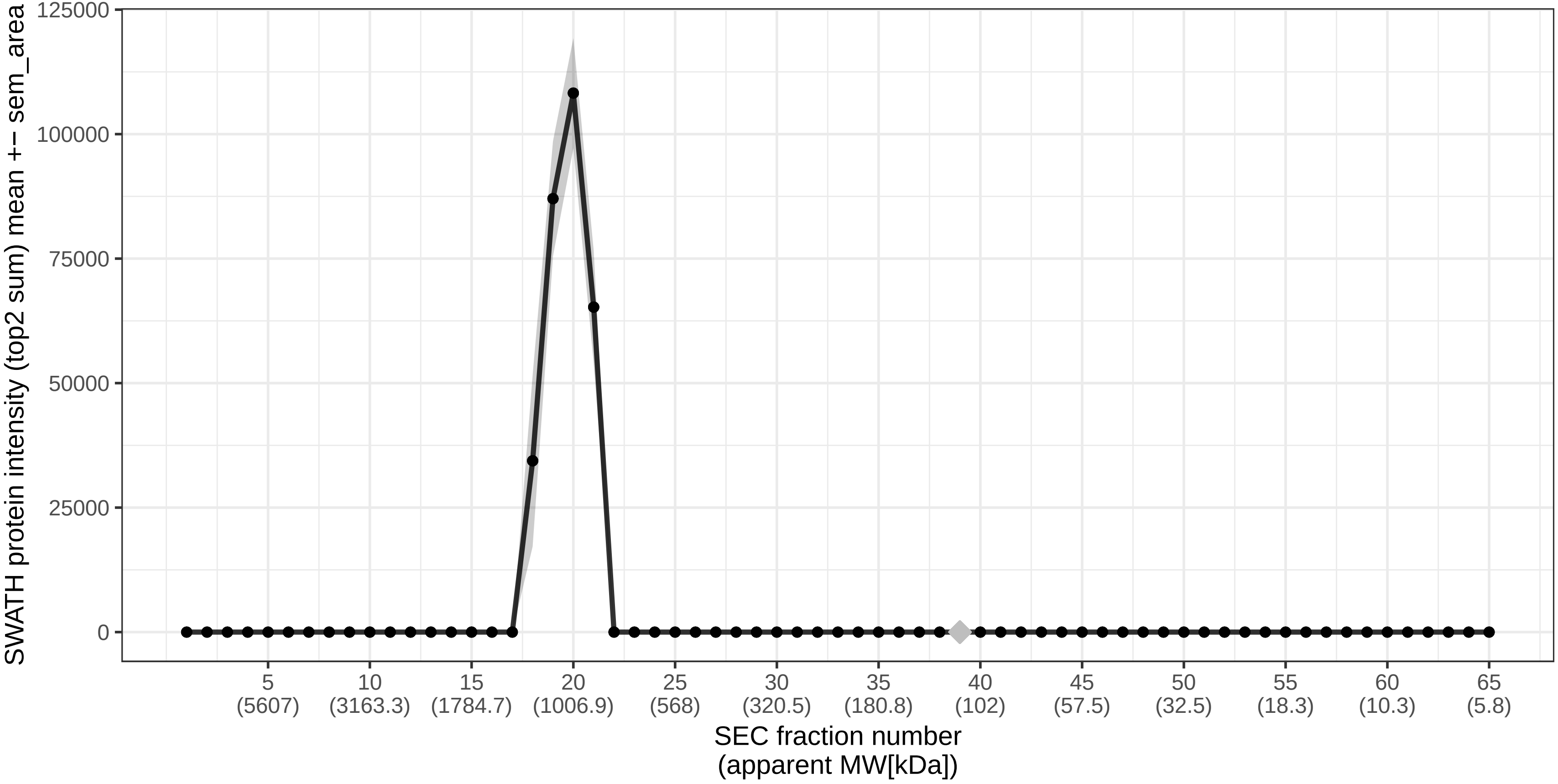

### SECchrom_O00299_CLIC1_HUMAN_CLIC1_G6_NCC27.pdf

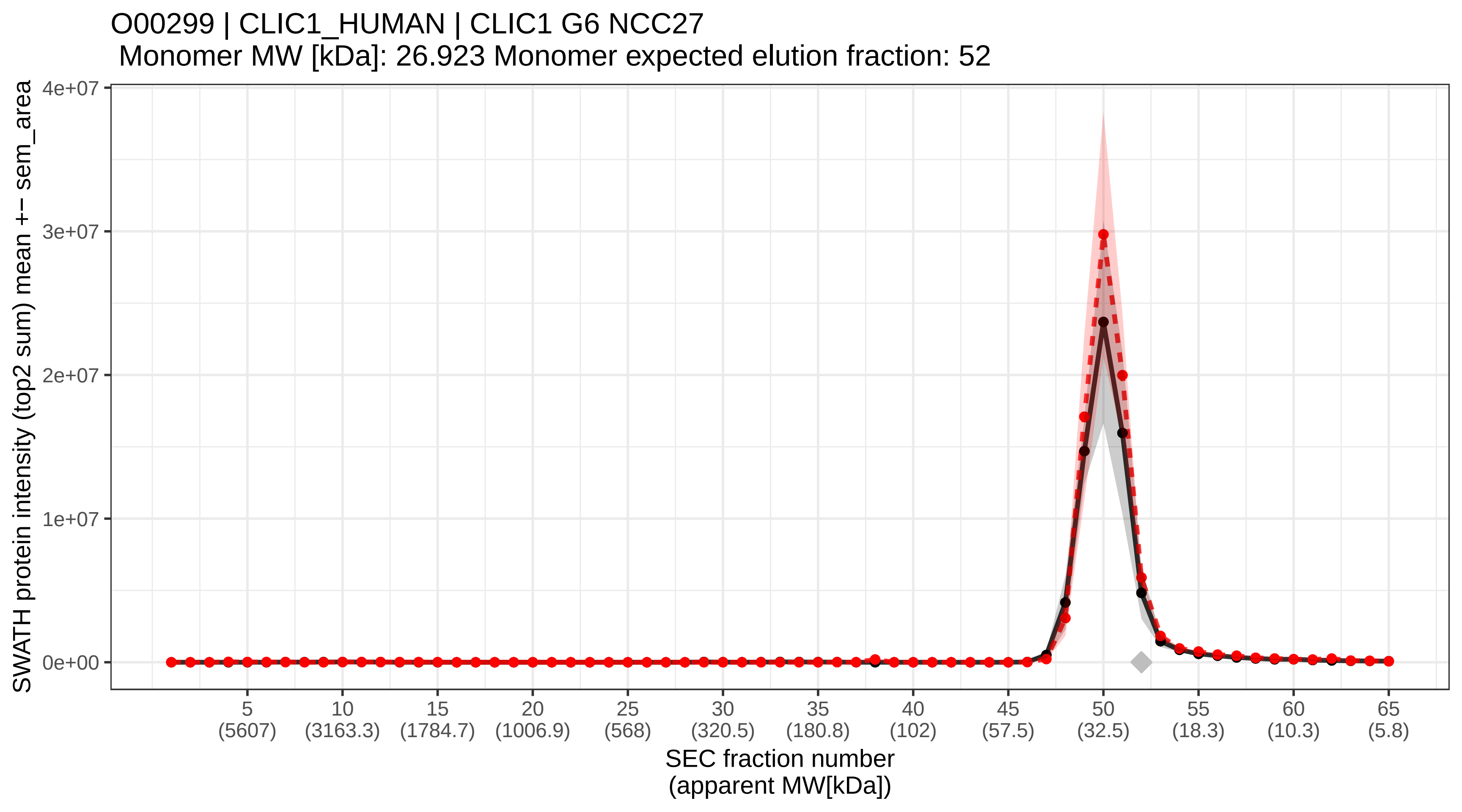

### SECchrom_O00303_EIF3F_HUMAN_EIF3F_EIF3S5.pdf

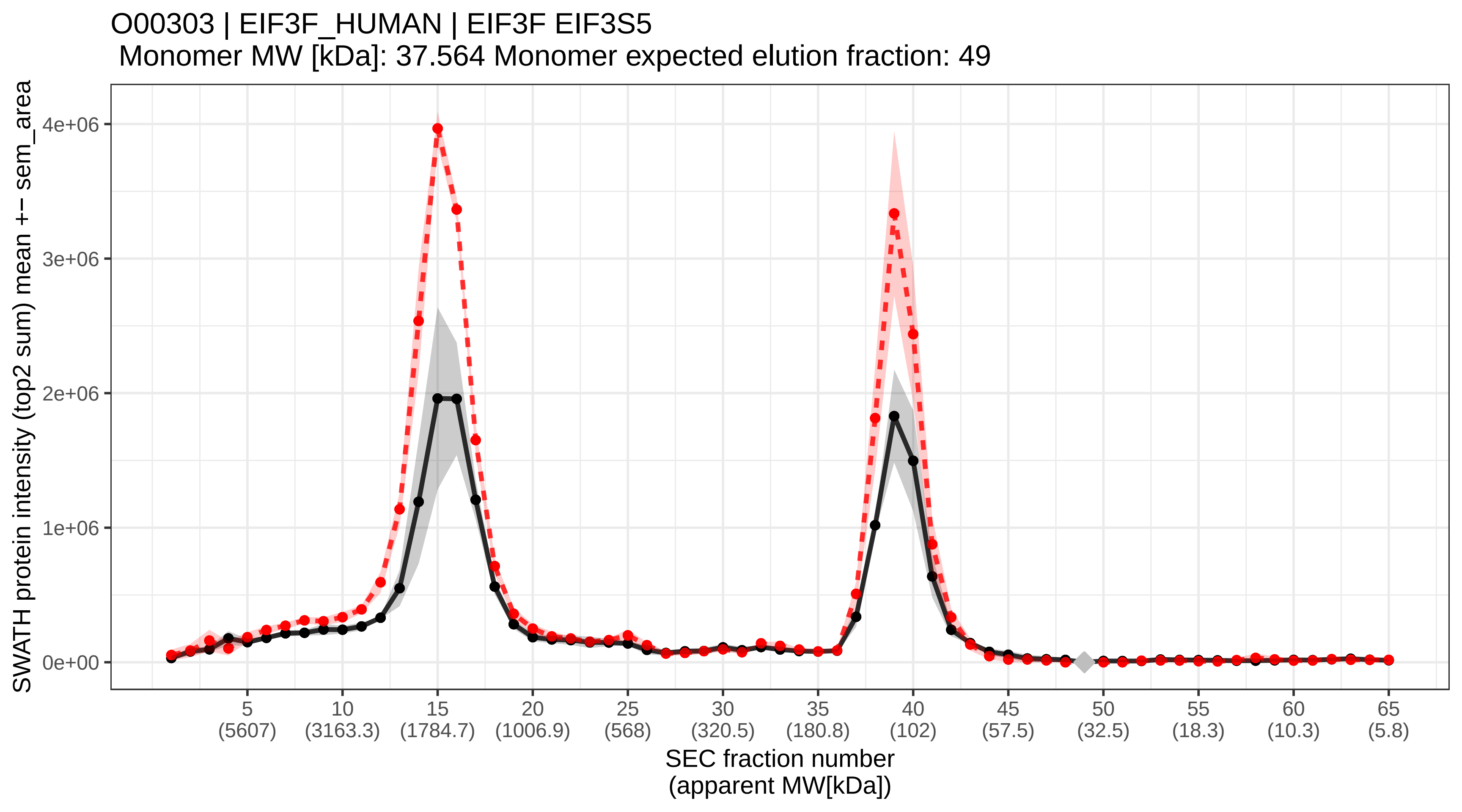

### SECchrom_O00330_ODPX_HUMAN_PDHX_PDX1.pdf

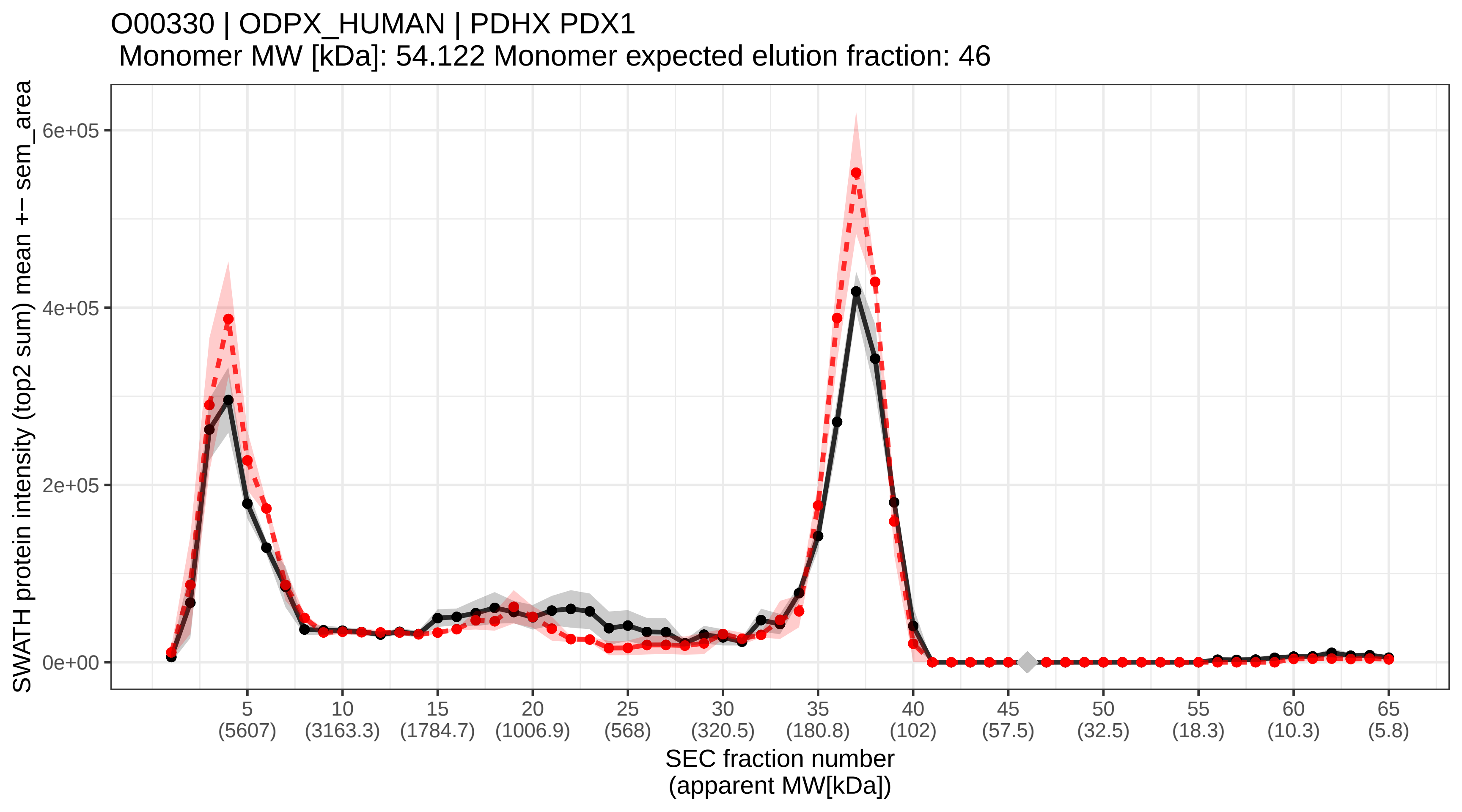

### SECchrom_O00399_DCTN6_HUMAN_DCTN6_WS3.pdf

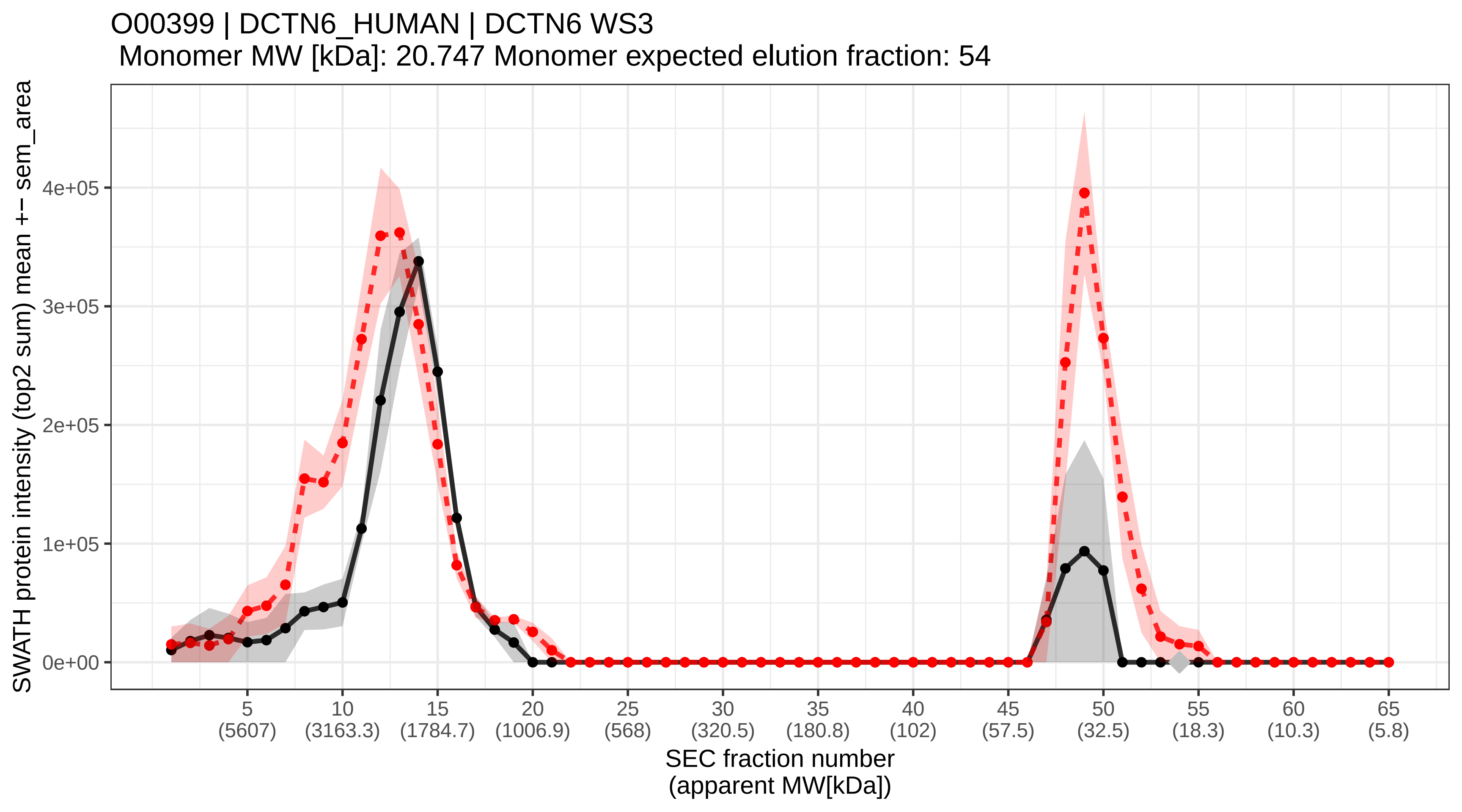

### SECchrom_O00400_ACATN_HUMAN_SLC33A1_ACATN_AT1.pdf

O00400 | ACATN_HUMAN | SLC33A1 ACATN AT1
Monomer MW [kDa]: 60.909 Monomer expected elution fraction: 45

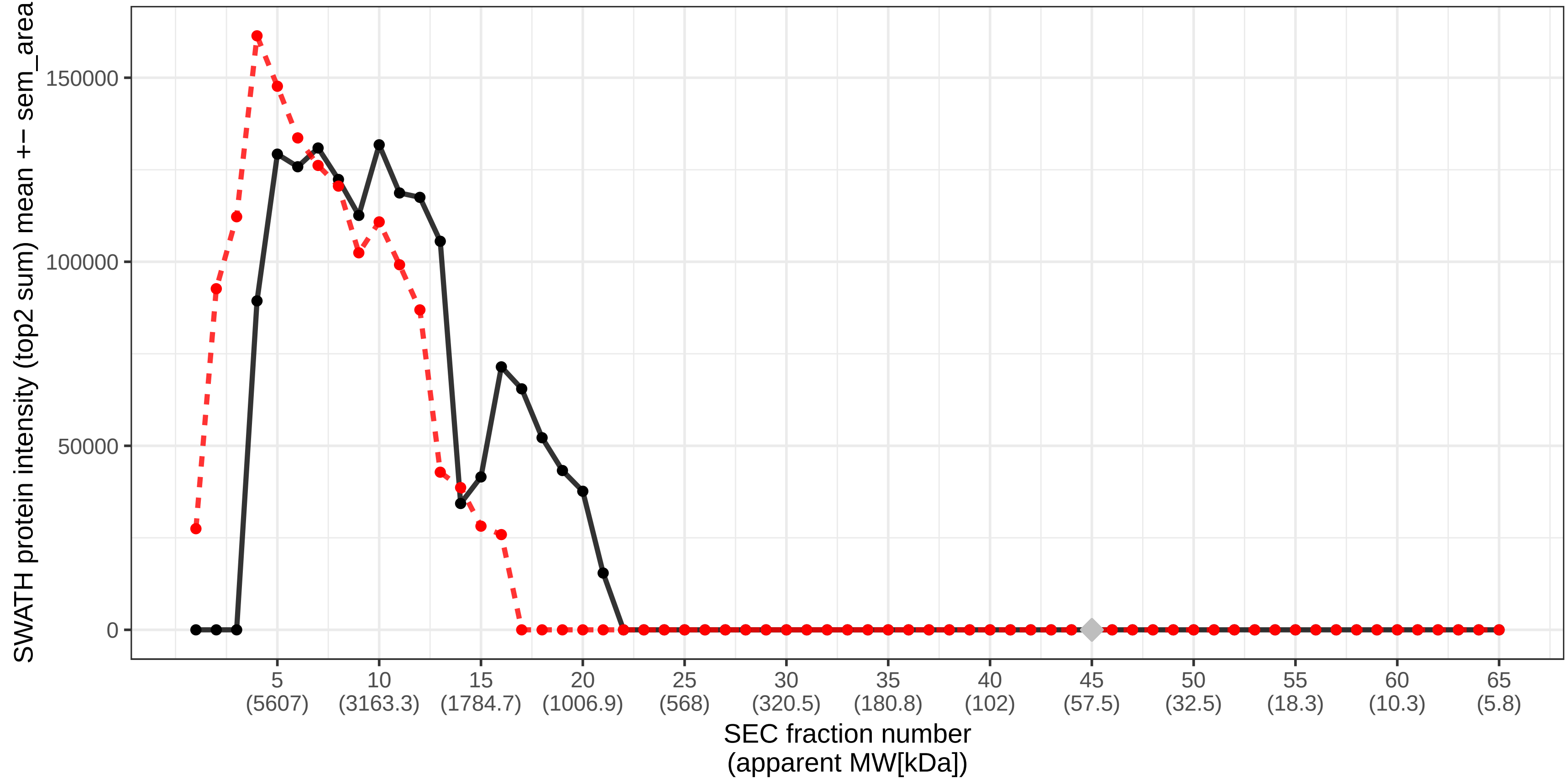

### SECchrom_O00401_WASL_HUMAN_WASL.pdf

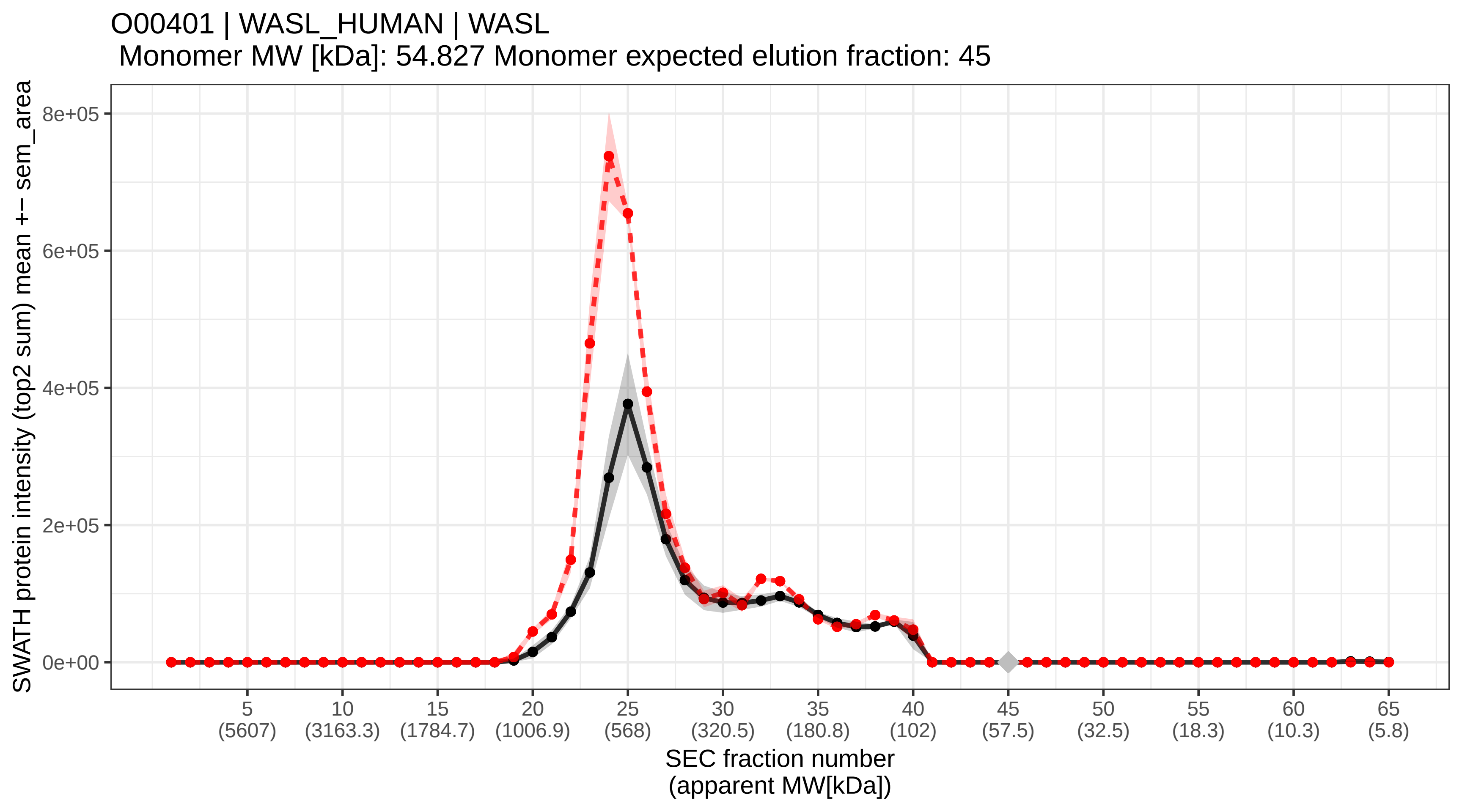

### SECchrom_O00410_IPO5_HUMAN_IPO5_KPNB3_RANBP5.pdf

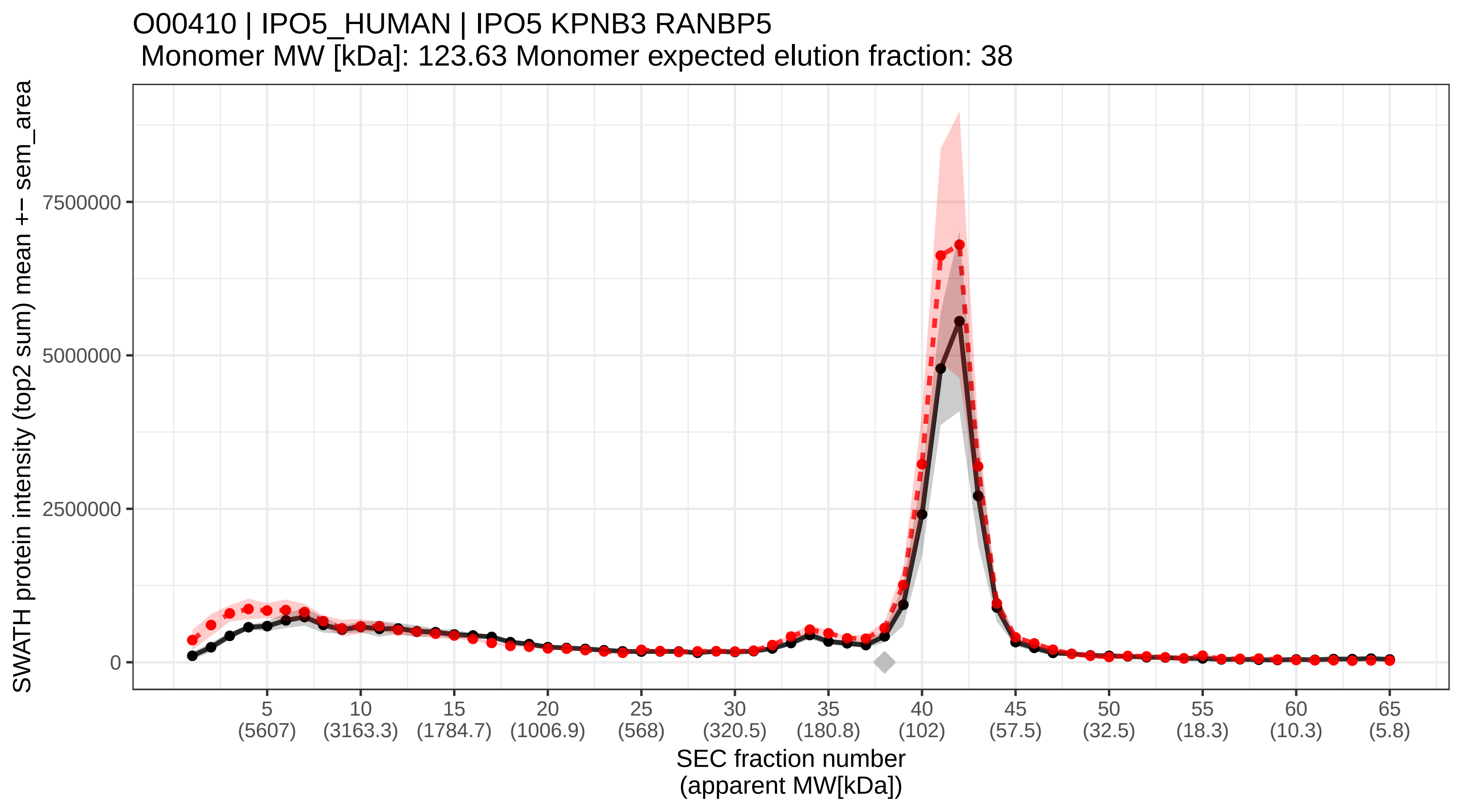

### SECchrom_O00411_RPOM_HUMAN_POLRMT.pdf

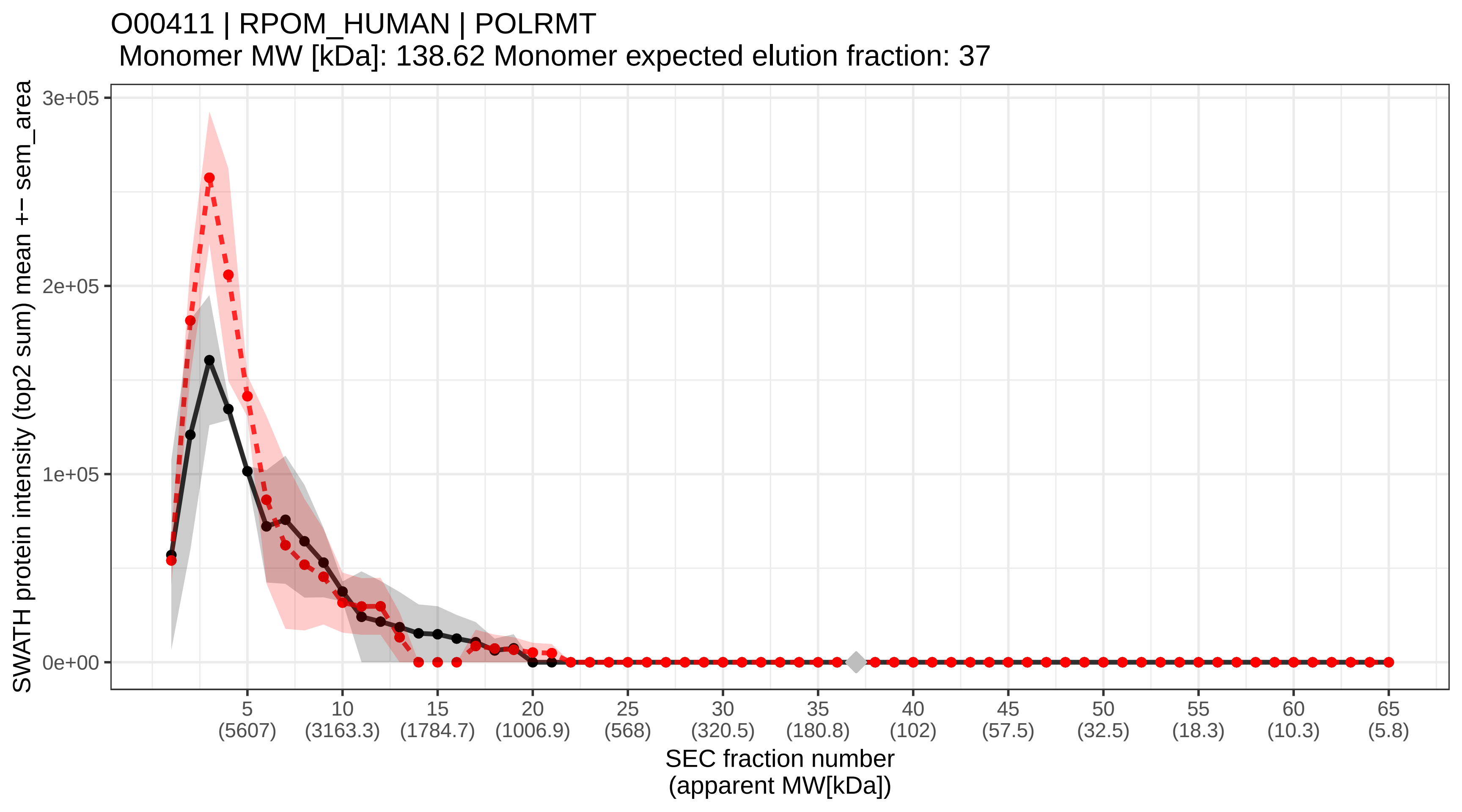

### SECchrom_O00418_EF2K_HUMAN_EEF2K.pdf

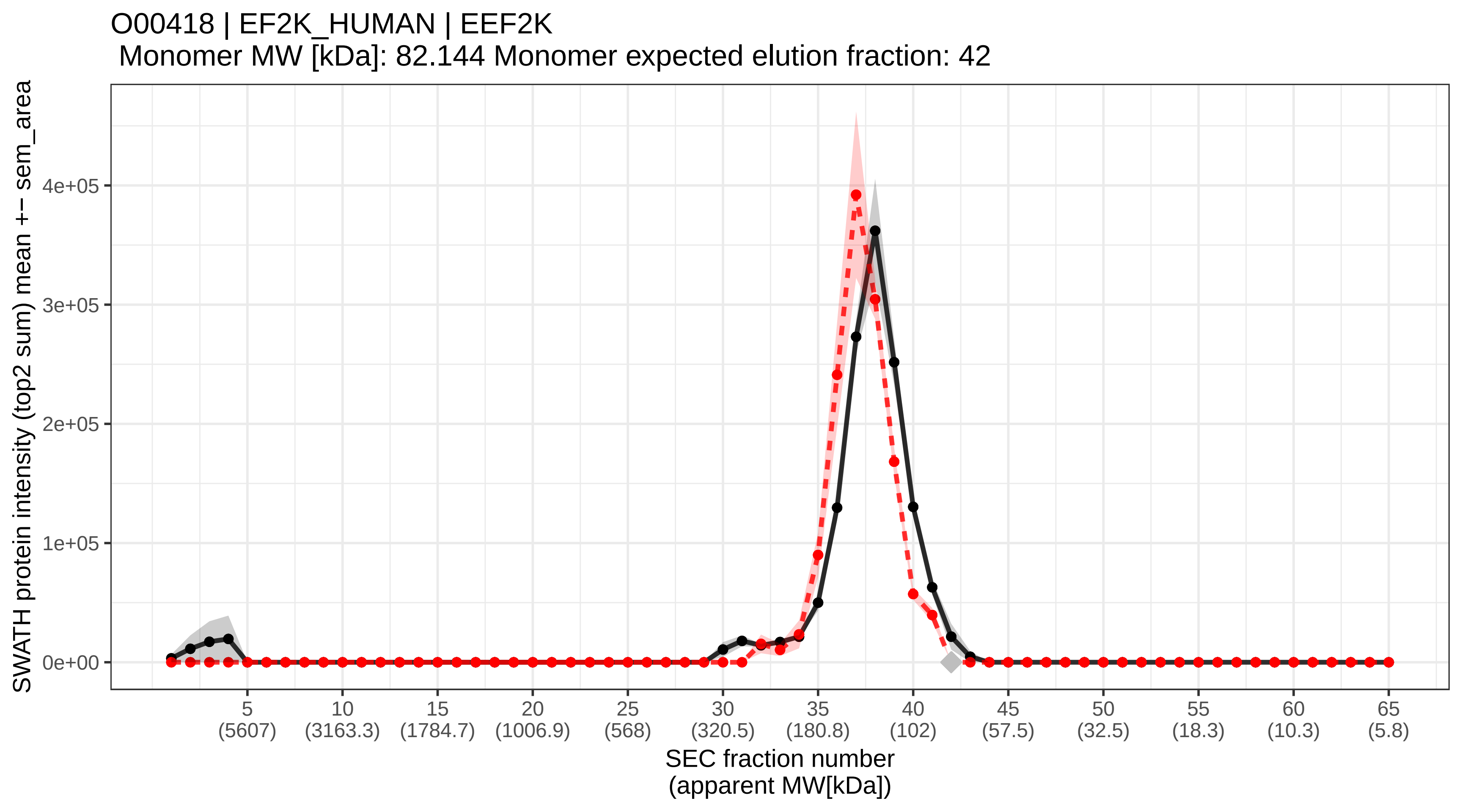

### SECchrom_O00425_IF2B3_HUMAN_IGF2BP3_IMP3_KOC1_VICKZ3.pdf

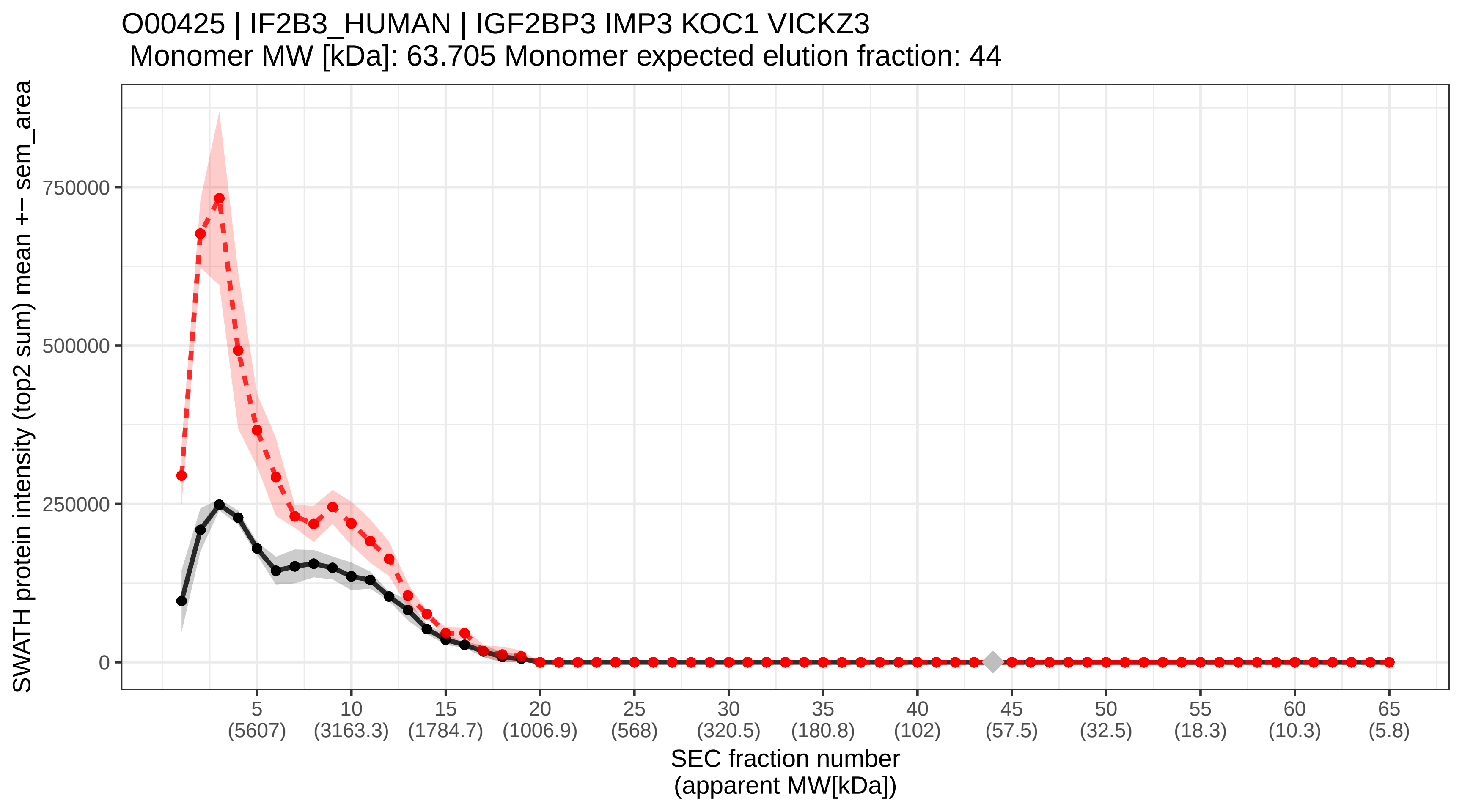

### SECchrom_O00429_DNM1L_HUMAN_DNM1L_DLP1_DRP1.pdf

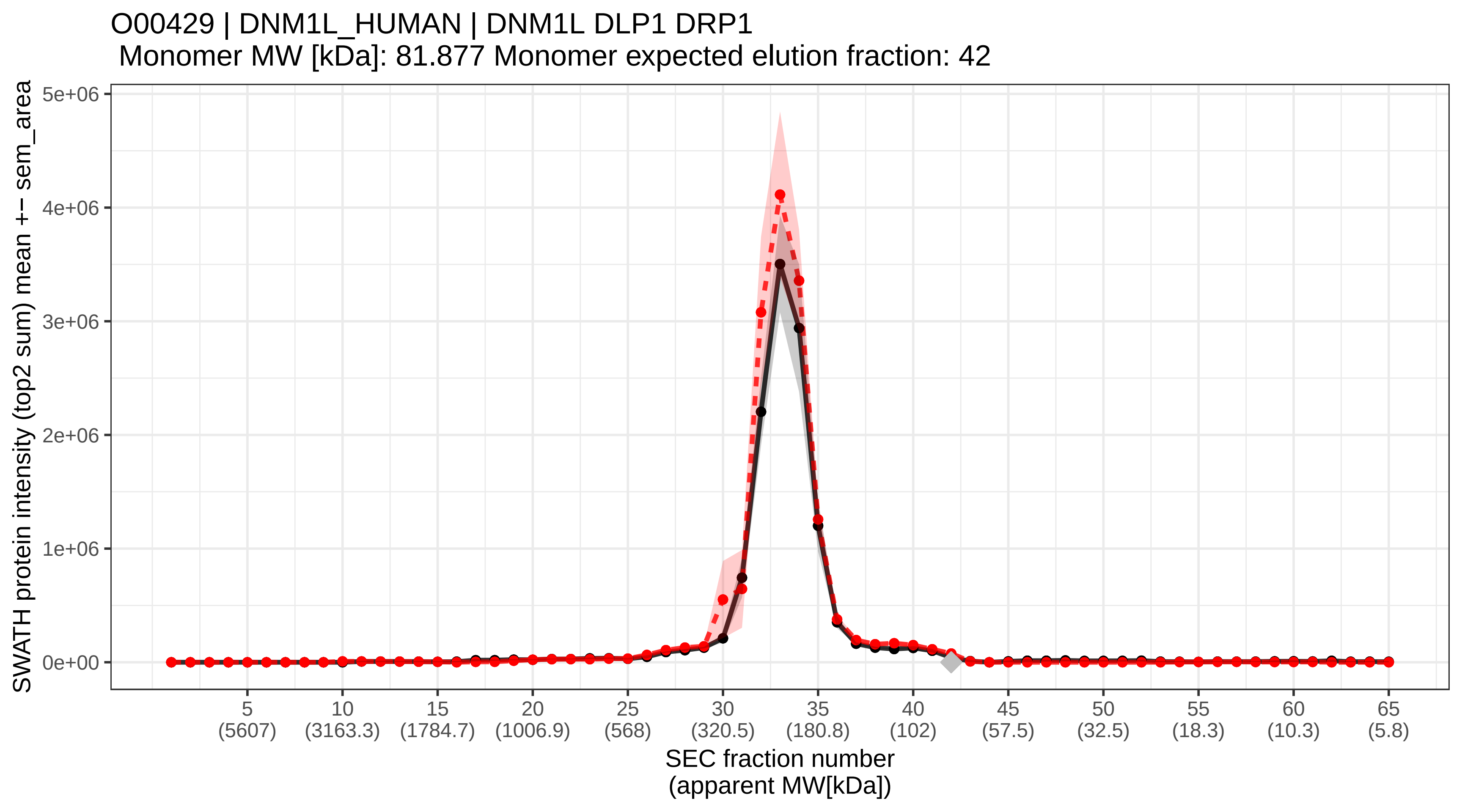

### SECchrom_O00442_RTCA_HUMAN_RTCA_RPC_RPC1_RTC1_RTCD1.pdf

O00442 | RTCA_HUMAN | RTCA RPC RPC1 RTC1 RTCD1
Monomer MW [kDa]: 39.337 Monomer expected elution fraction: 48

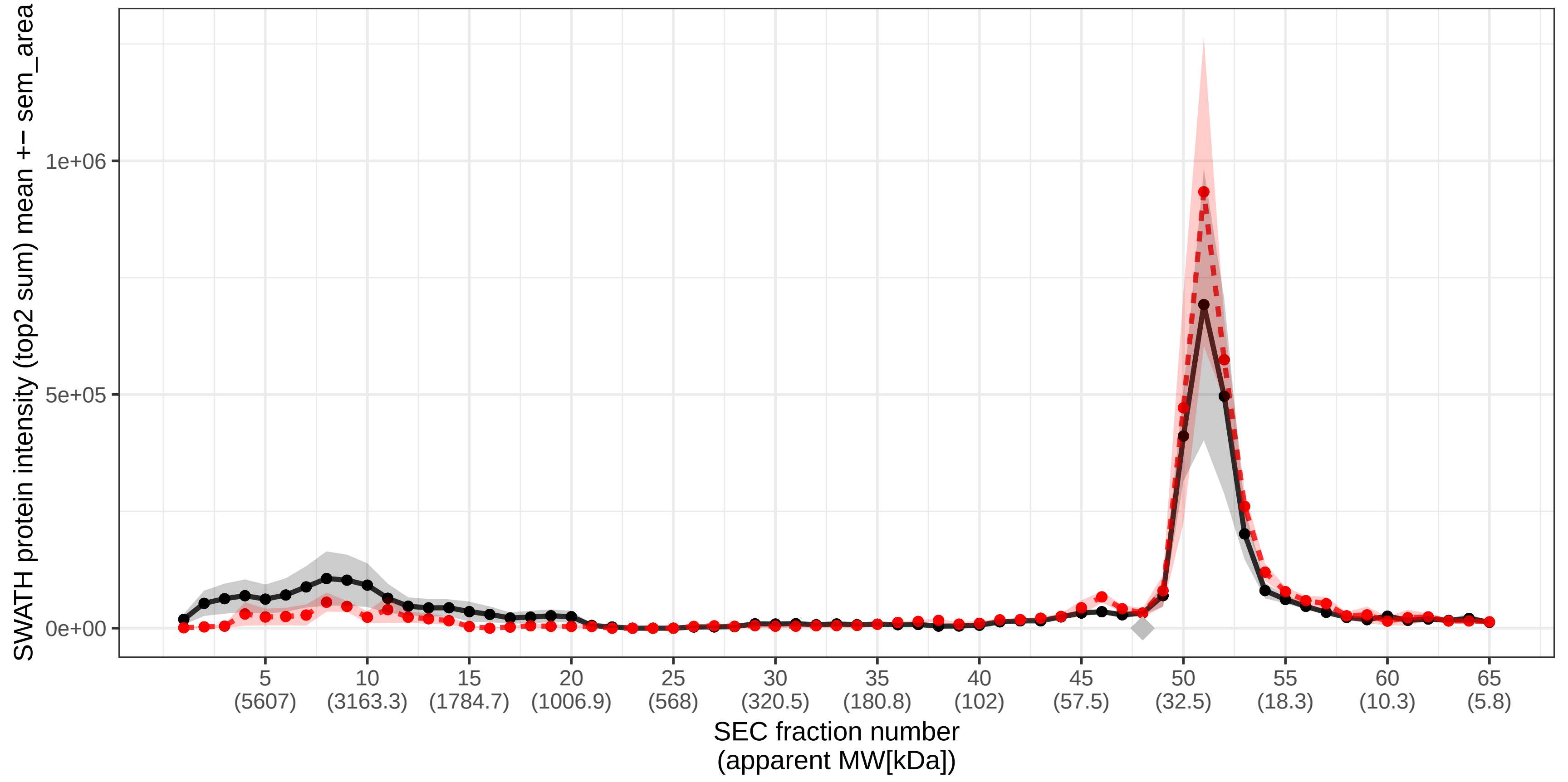

### SECchrom_O00459_P85B_HUMAN_PIK3R2.pdf

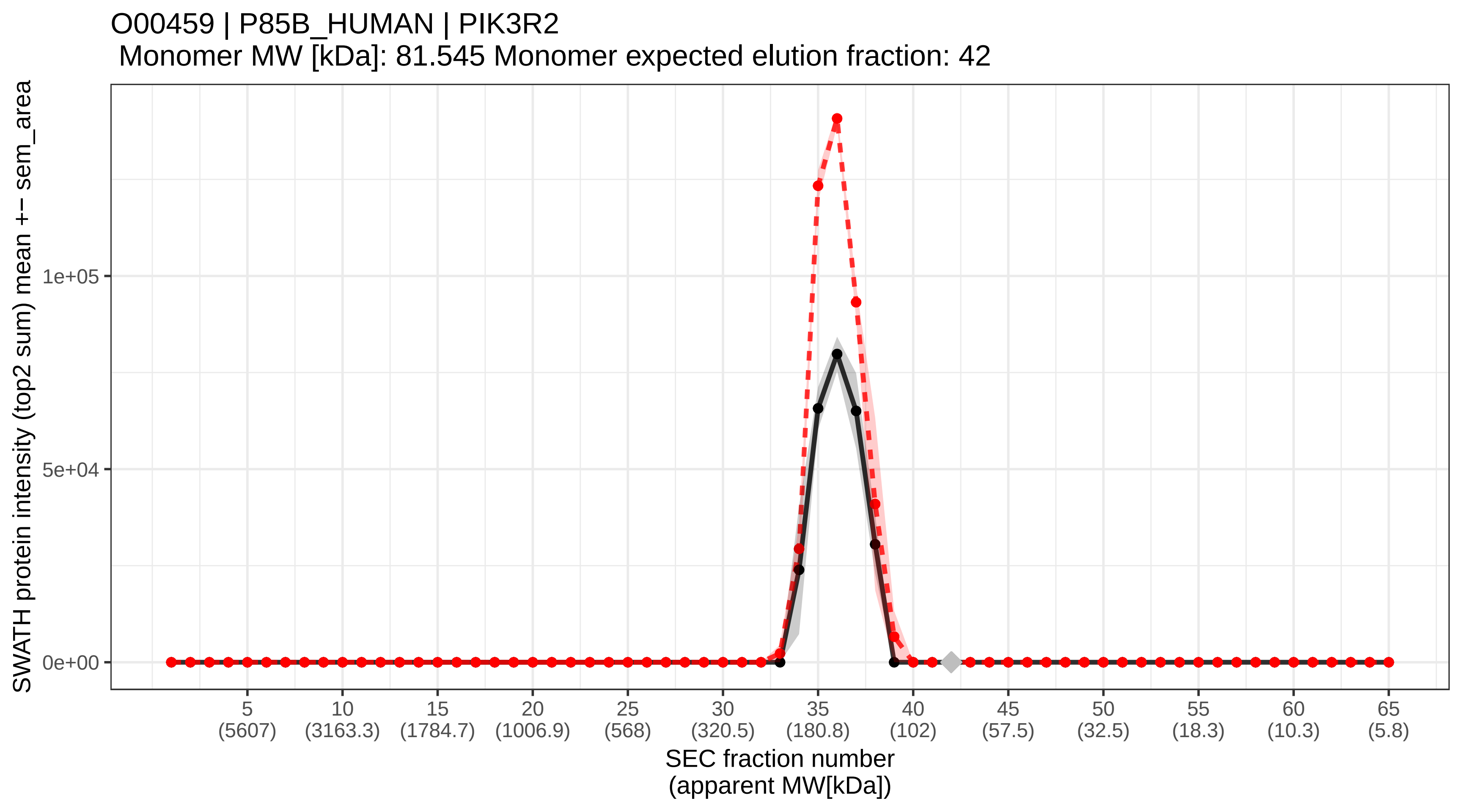

### SECchrom_O00461_GOLI4_HUMAN_GOLIM4_GIMPC_GOLPH4_GPP130.pdf

O00461 | GOLI4_HUMAN | GOLIM4 GIMPC GOLPH4 GPP130
Monomer MW [kDa]: 81.88 Monomer expected elution fraction: 42

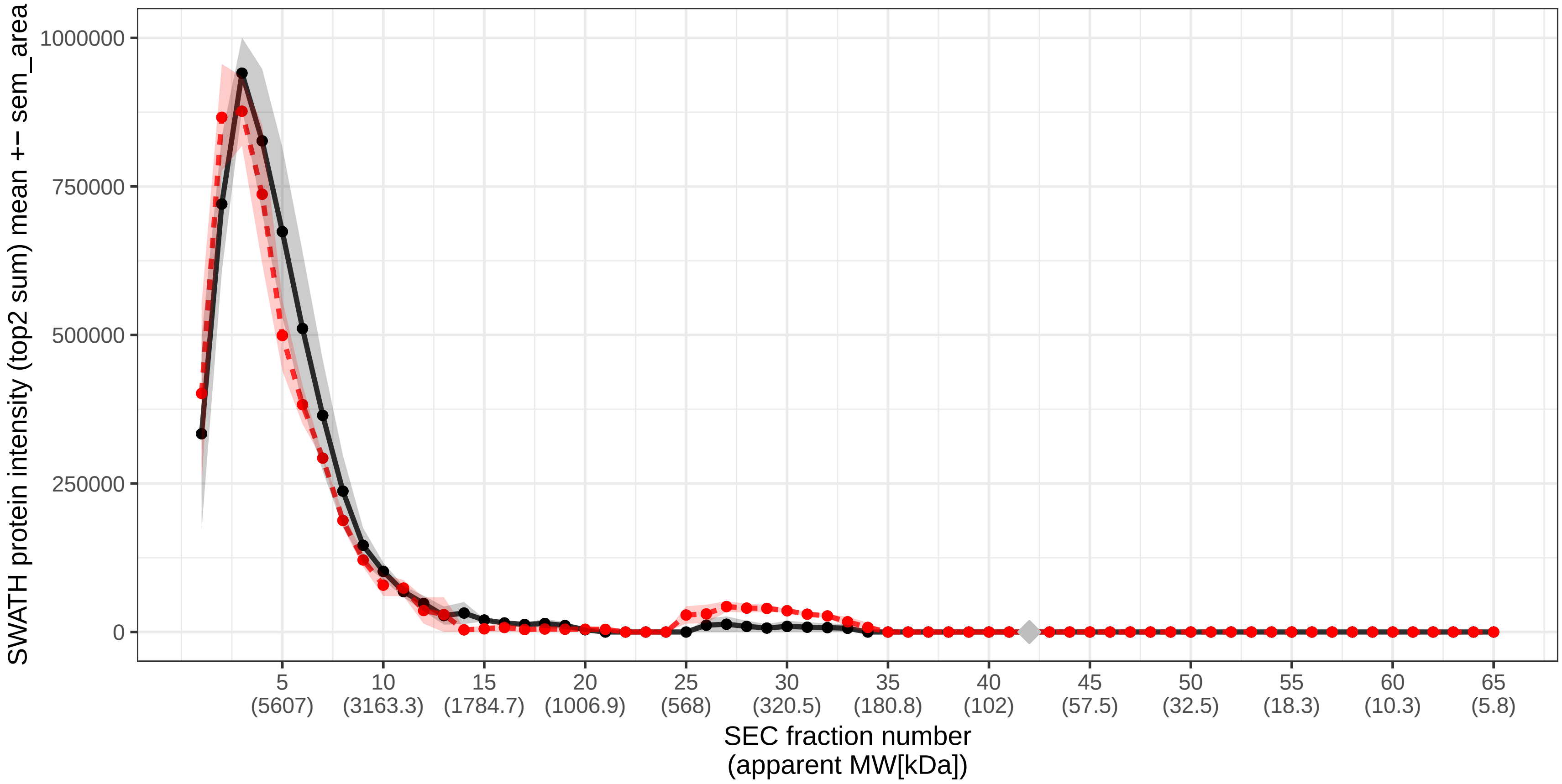

### SECchrom_O00462_MANBA_HUMAN_MANBA_MANB1.pdf

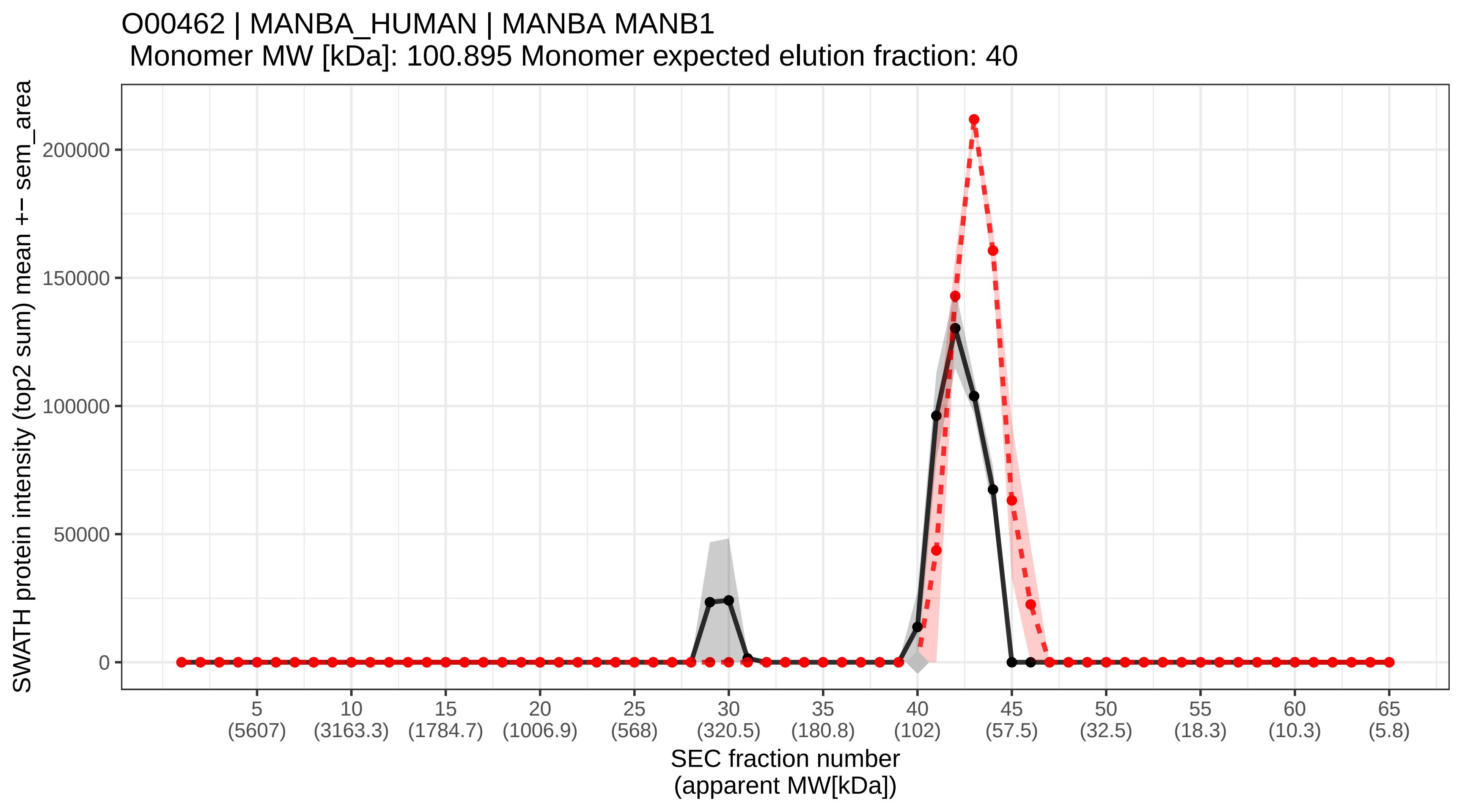

### SECchrom_O00468_AGRIN_HUMAN_AGRN_AGRIN.pdf

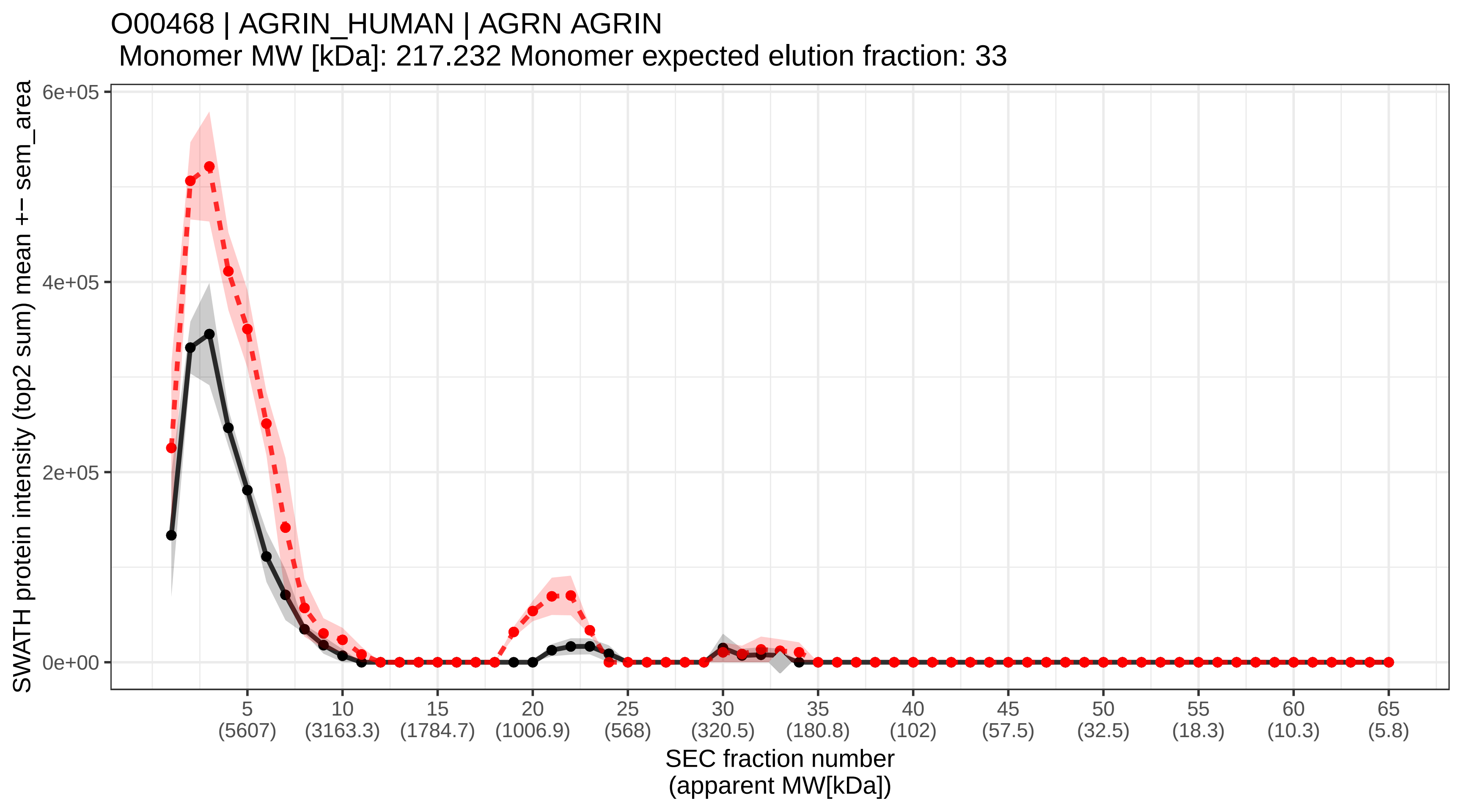

### SECchrom_O00469_PLOD2_HUMAN_PLOD2.pdf

O00469 | PLOD2_HUMAN | PLOD2
Monomer MW [kDa]: 84.686 Monomer expected elution fraction: 42

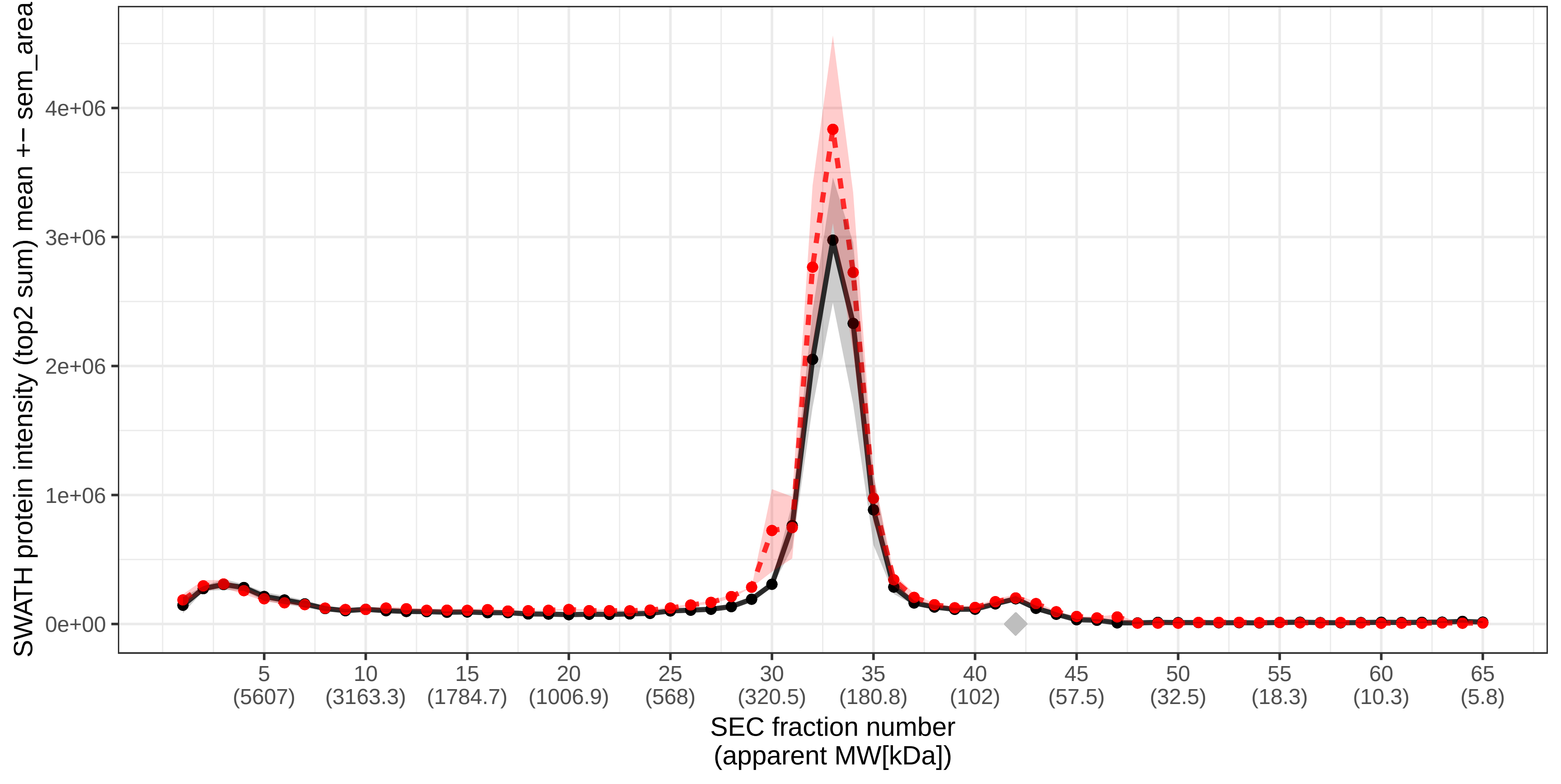

### SECchrom_O00470_MEIS1_HUMAN_MEIS1.pdf

#### O00470 | MEIS1_HUMAN | MEIS1
Monomer MW [kDa]: 43.016 Monomer expected elution fraction: 48

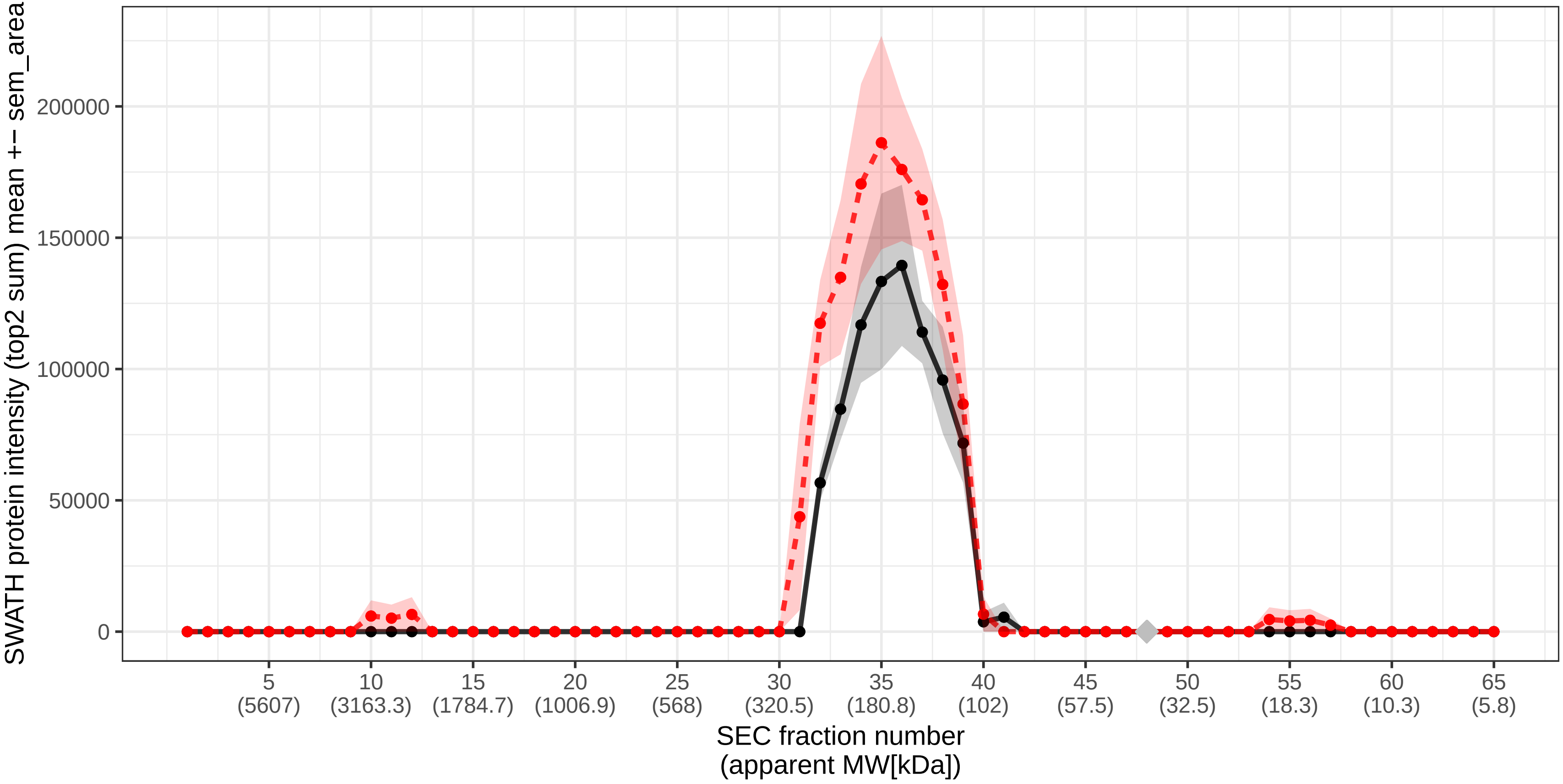

### SECchrom_O00471_EXOC5_HUMAN_EXOC5_SEC10_SEC10L1.pdf

O00471 | EXOC5_HUMAN | EXOC5 SEC10 SEC10L1
Monomer MW [kDa]: 81.853 Monomer expected elution fraction: 42

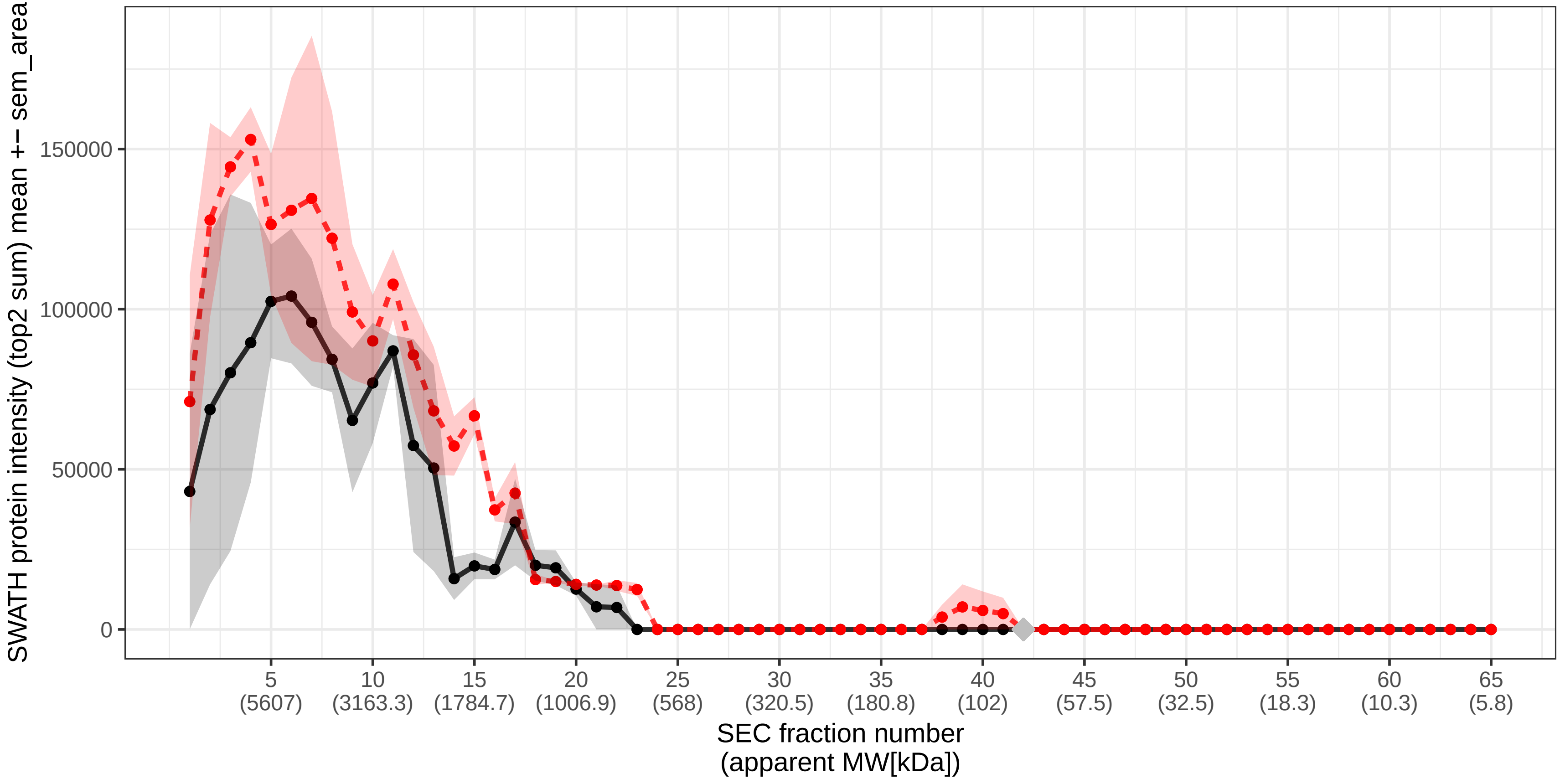

### SECchrom_O00483_NDUA4_HUMAN_NDUFA4.pdf

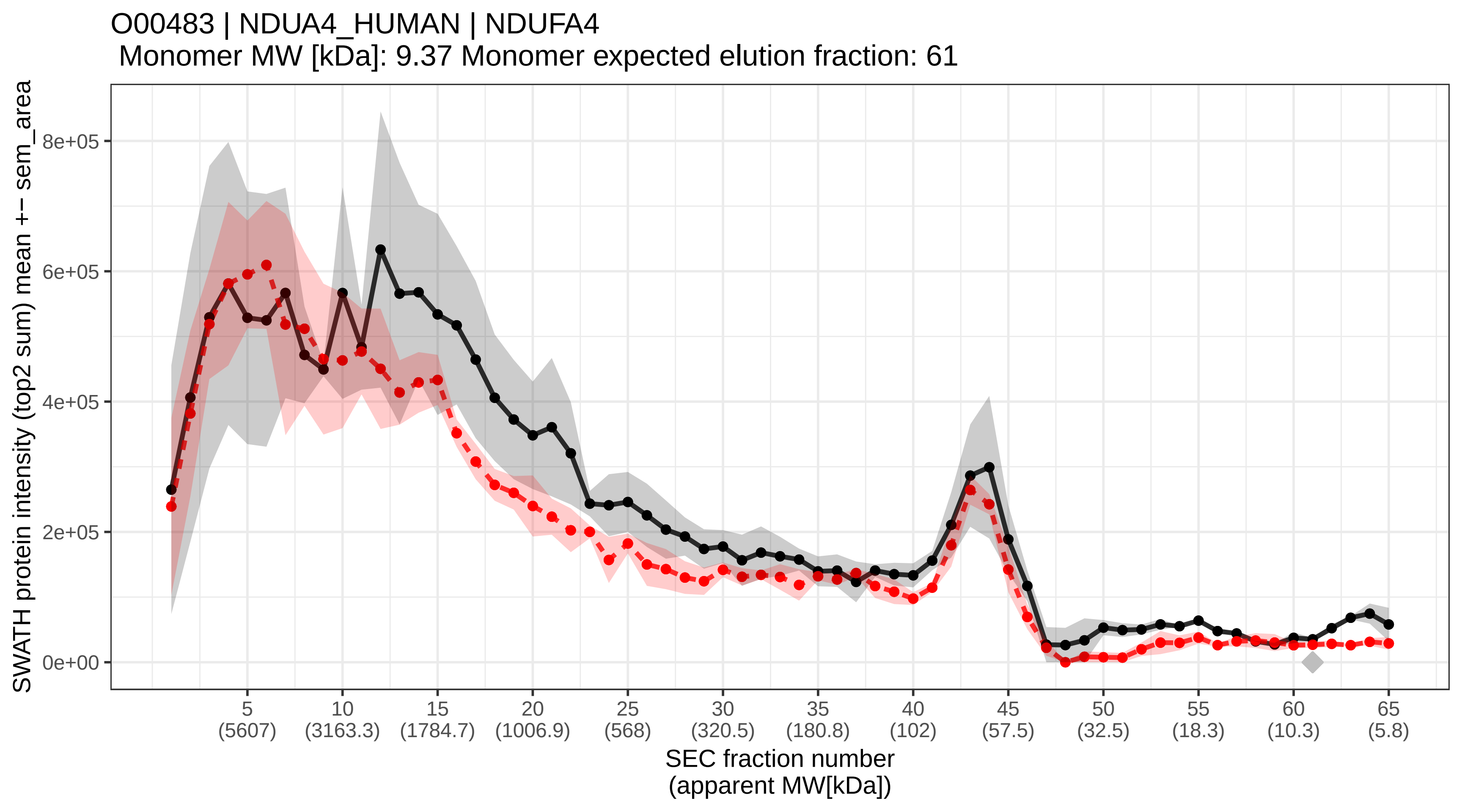

### SECchrom_O00487_PSDE_HUMAN_PSMD14_POH1.pdf

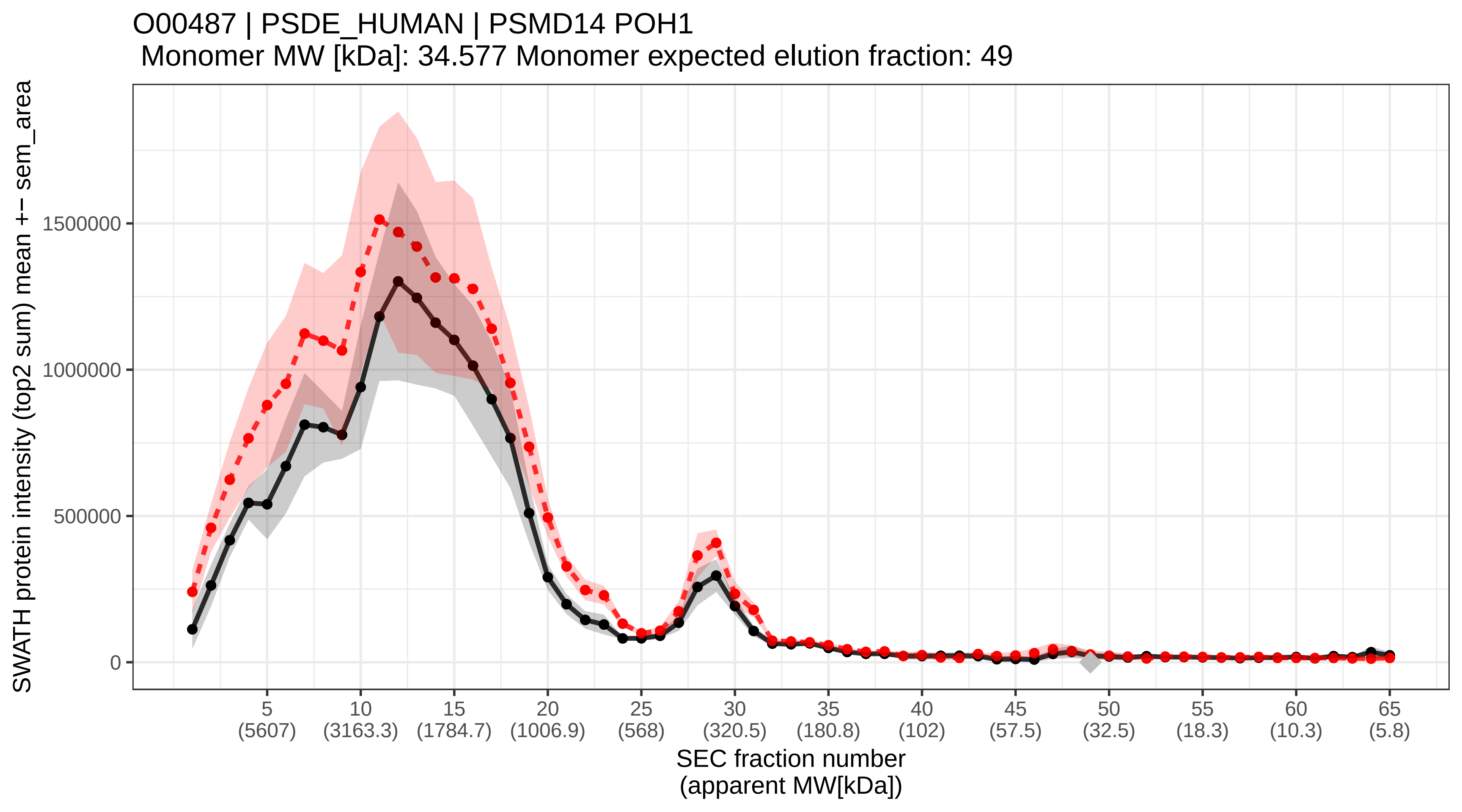

### SECchrom_O00499_BIN1_HUMAN_BIN1_AMPHL.pdf

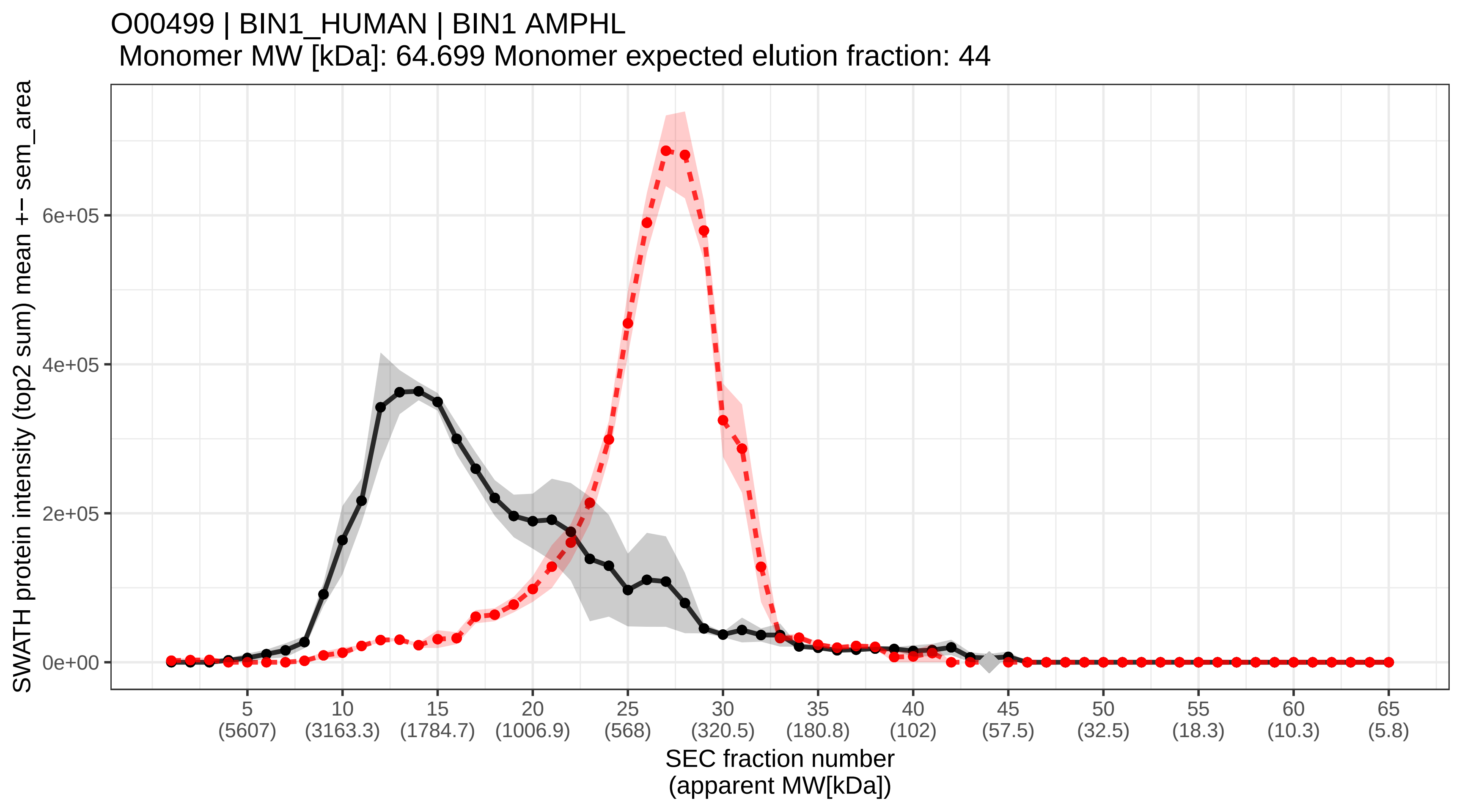

### SECchrom_O00505_IMA4_HUMAN_KPNA3_QIP2.pdf

#### O00505 | IMA4_HUMAN | KPNA3 QIP2
Monomer MW [kDa]: 57.811 Monomer expected elution fraction: 45

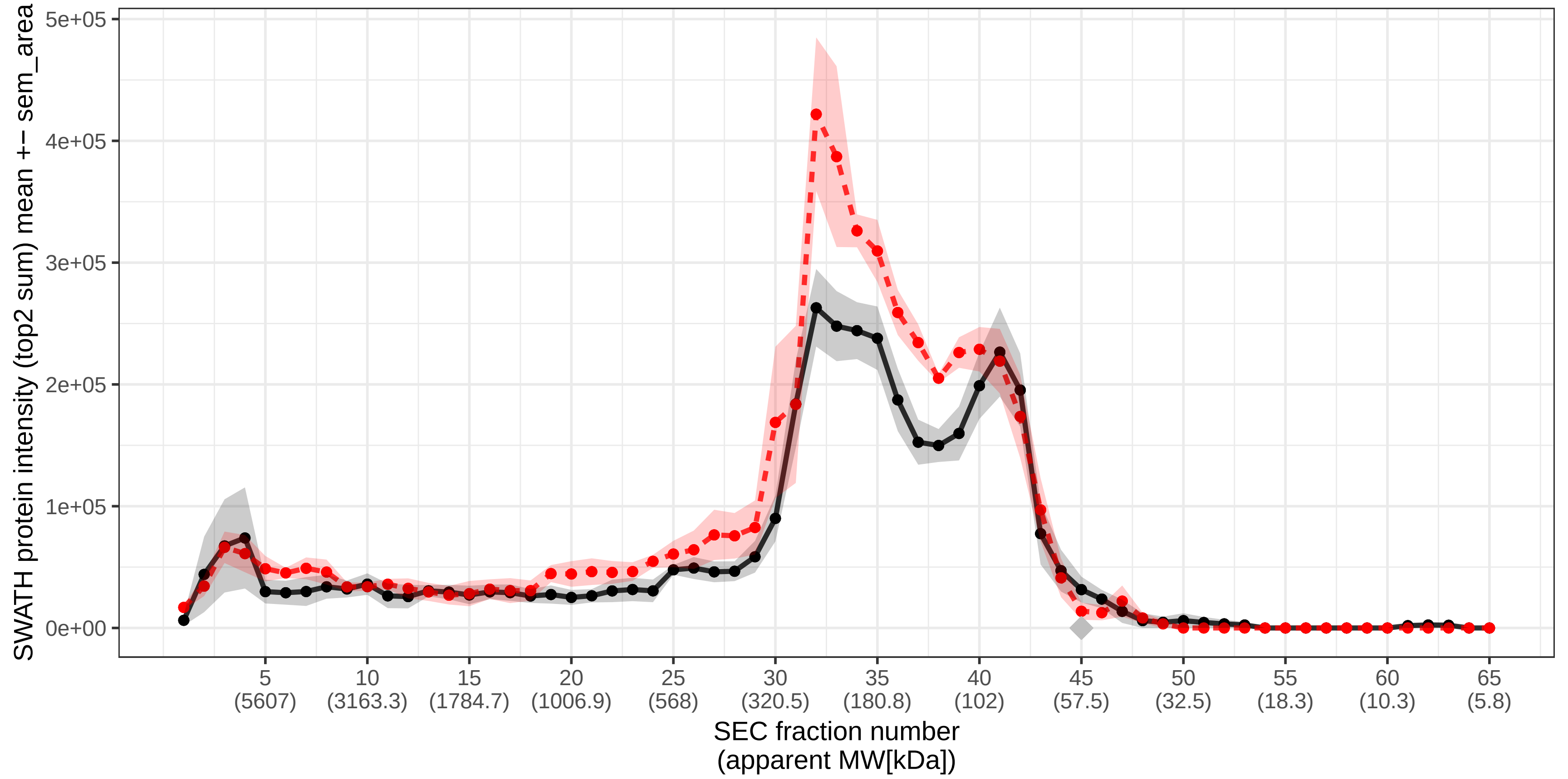

### SECchrom_O00506_STK25_HUMAN_STK25_SOK1_YSK1.pdf

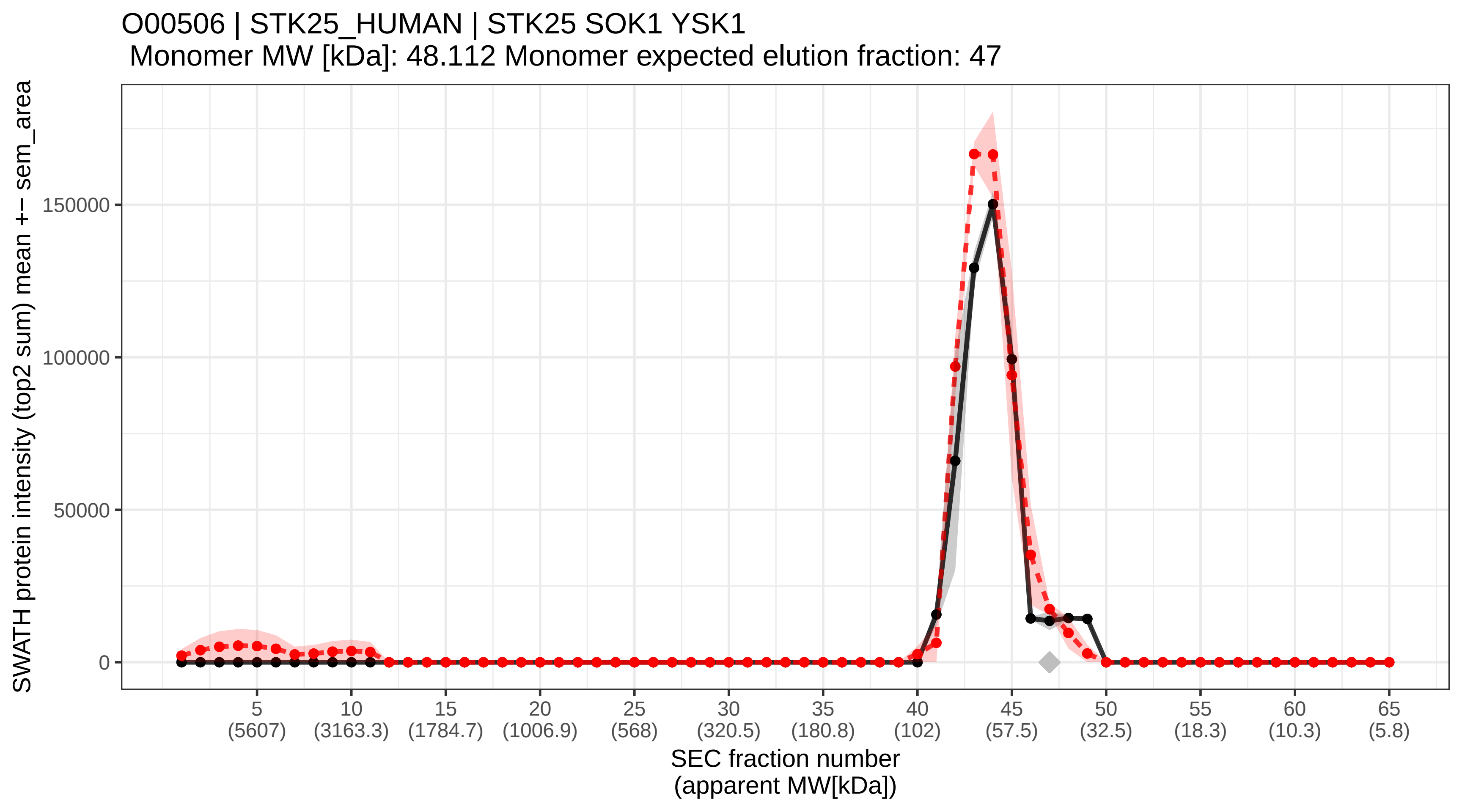

### SECchrom_O00541_PESC_HUMAN_PES1.pdf

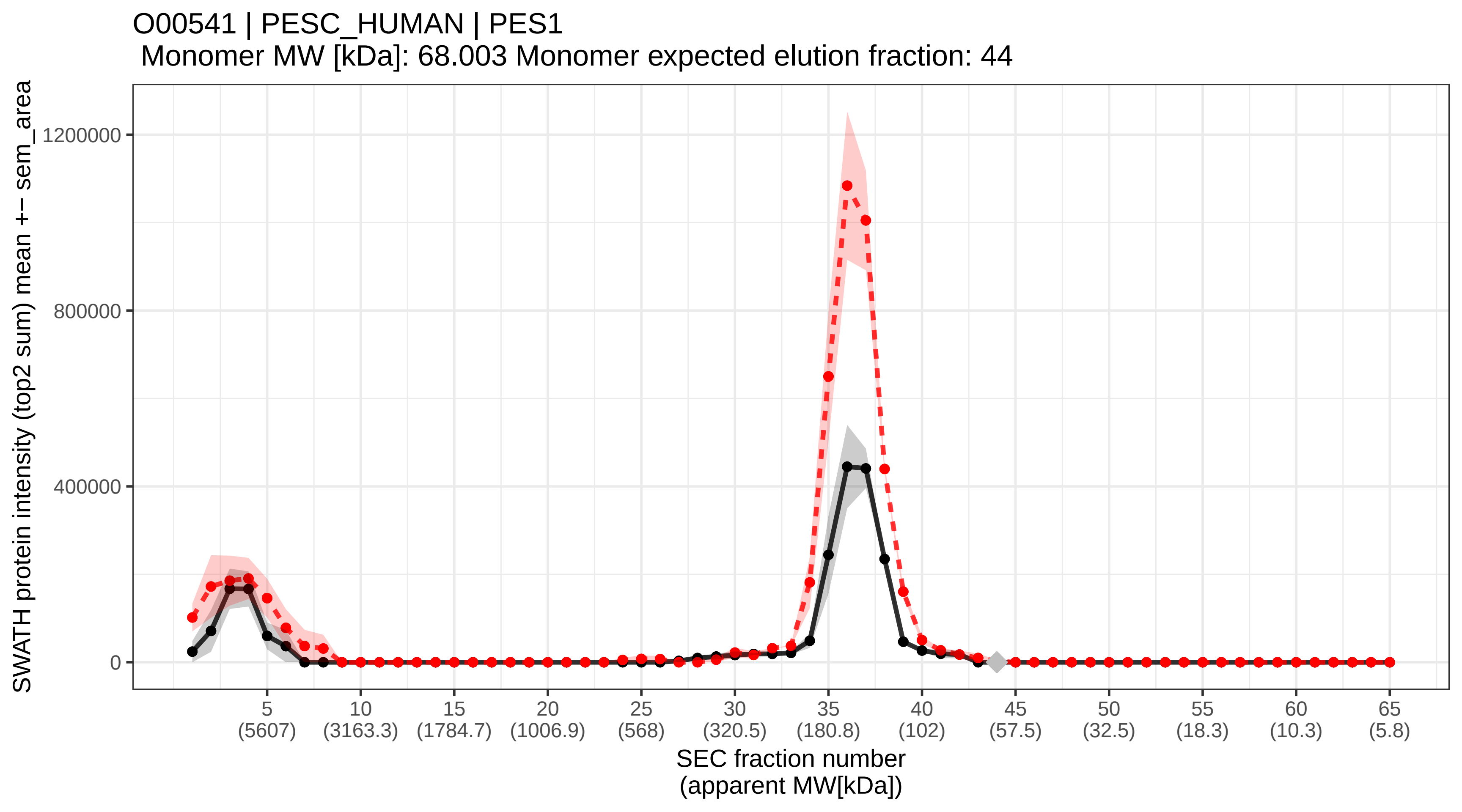

### SECchrom_O00560_SDCB1_HUMAN_SDCBP_MDA9_SYCL.pdf

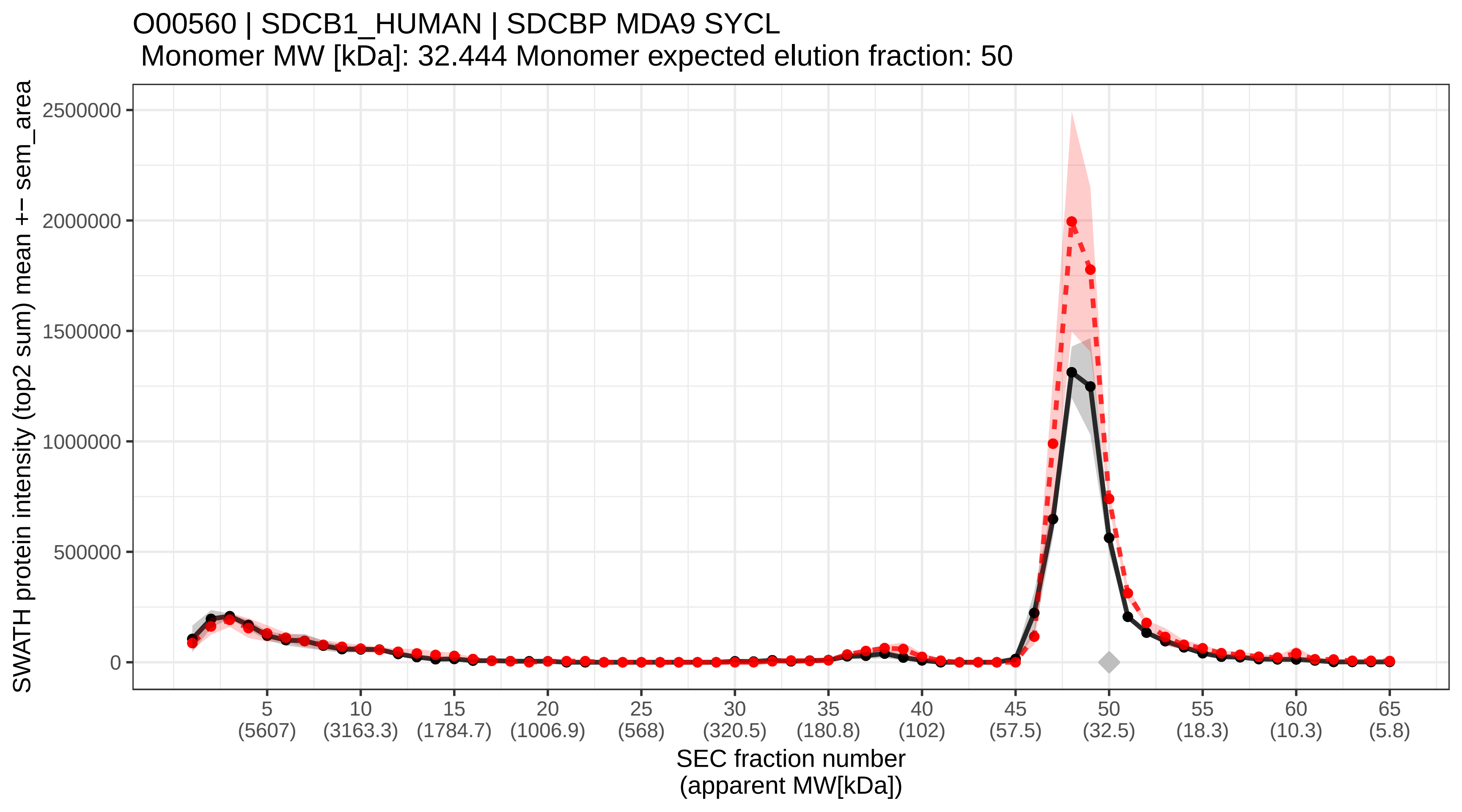

### SECchrom_O00567_NOP56_HUMAN_NOP56_NOL5A.pdf

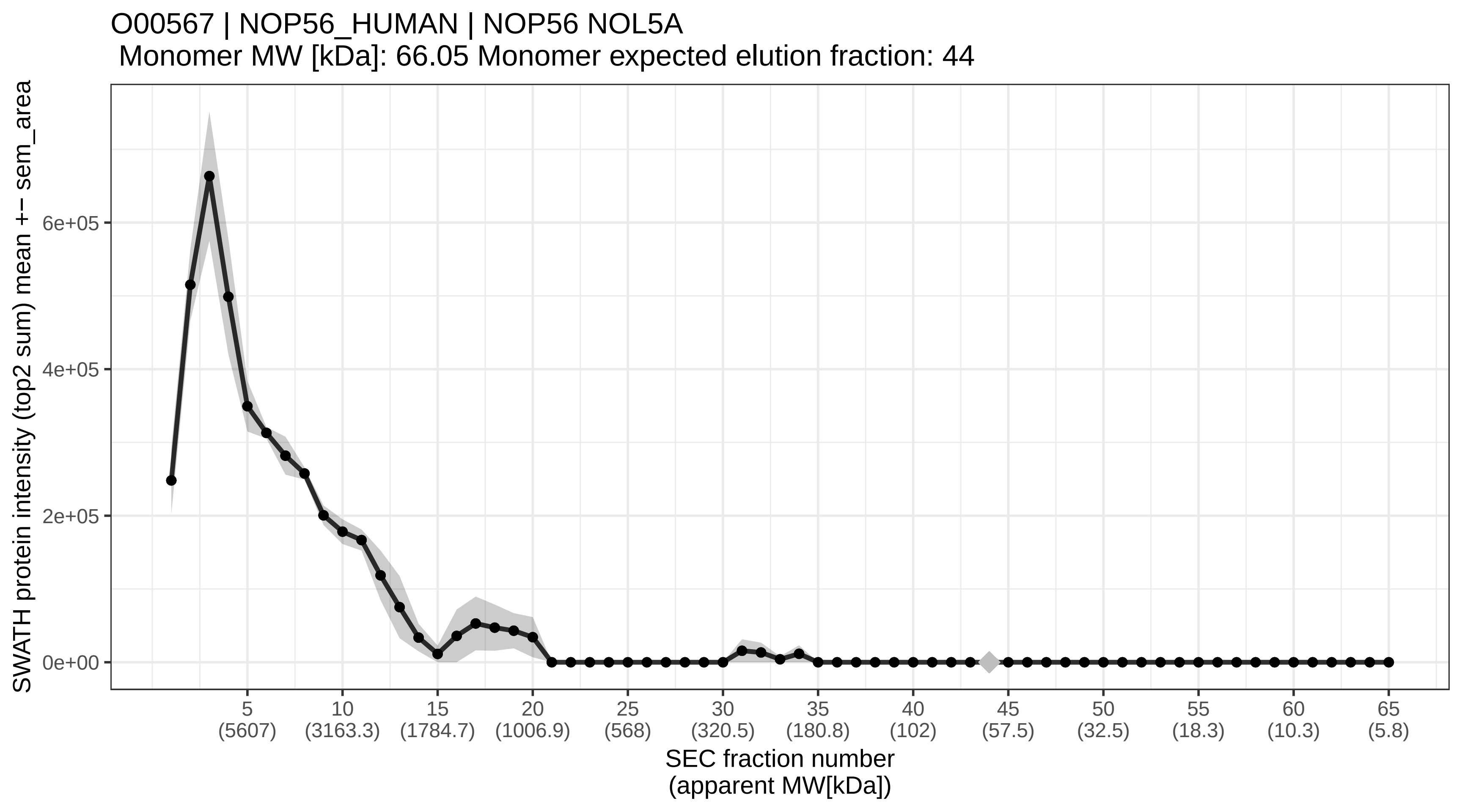

### SECchrom_O00584_RNT2_HUMAN_RNASET2_RNASE6PL.pdf

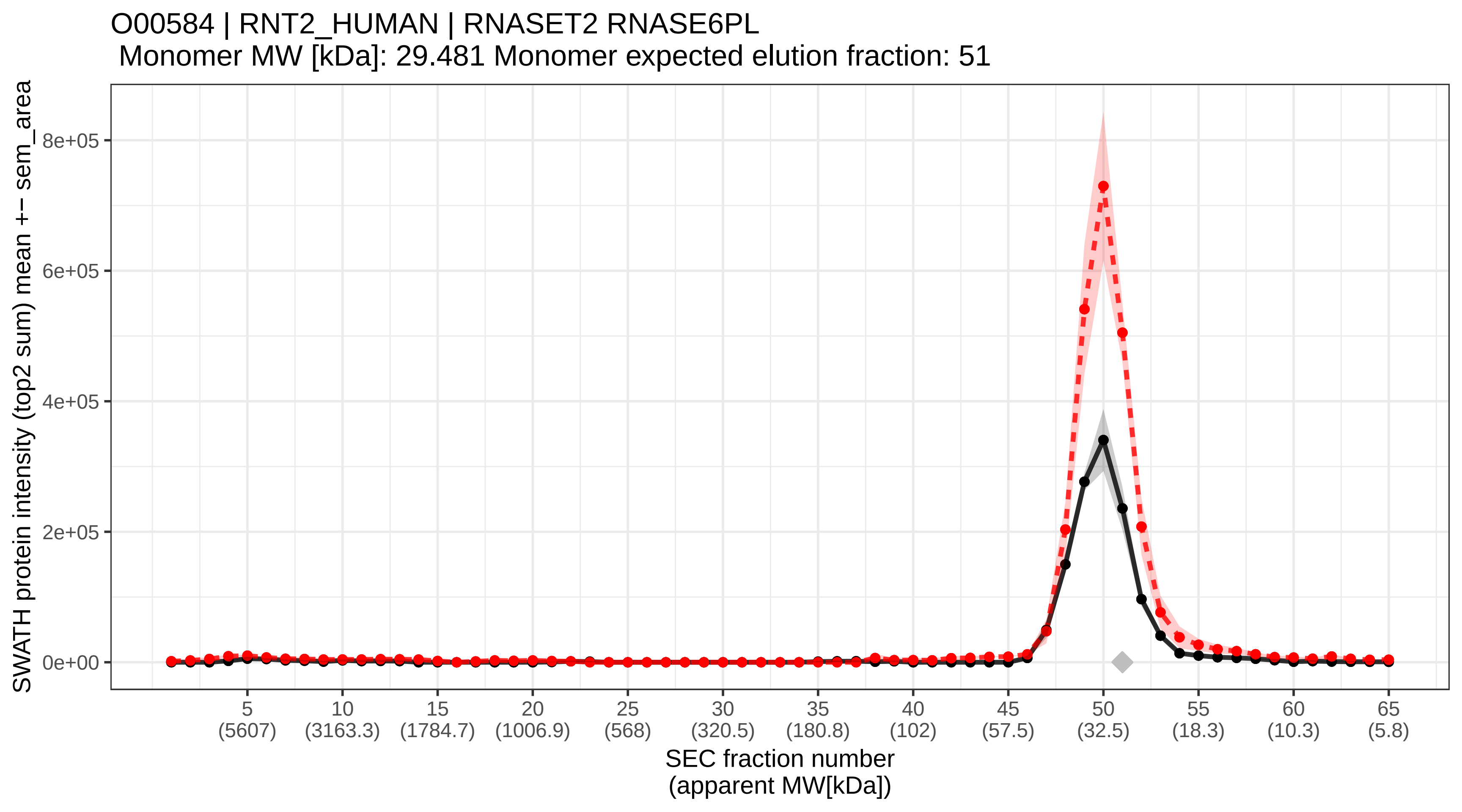

### SECchrom_O00592_PODXL_HUMAN_PODXL_PCLP_PCLP1.pdf

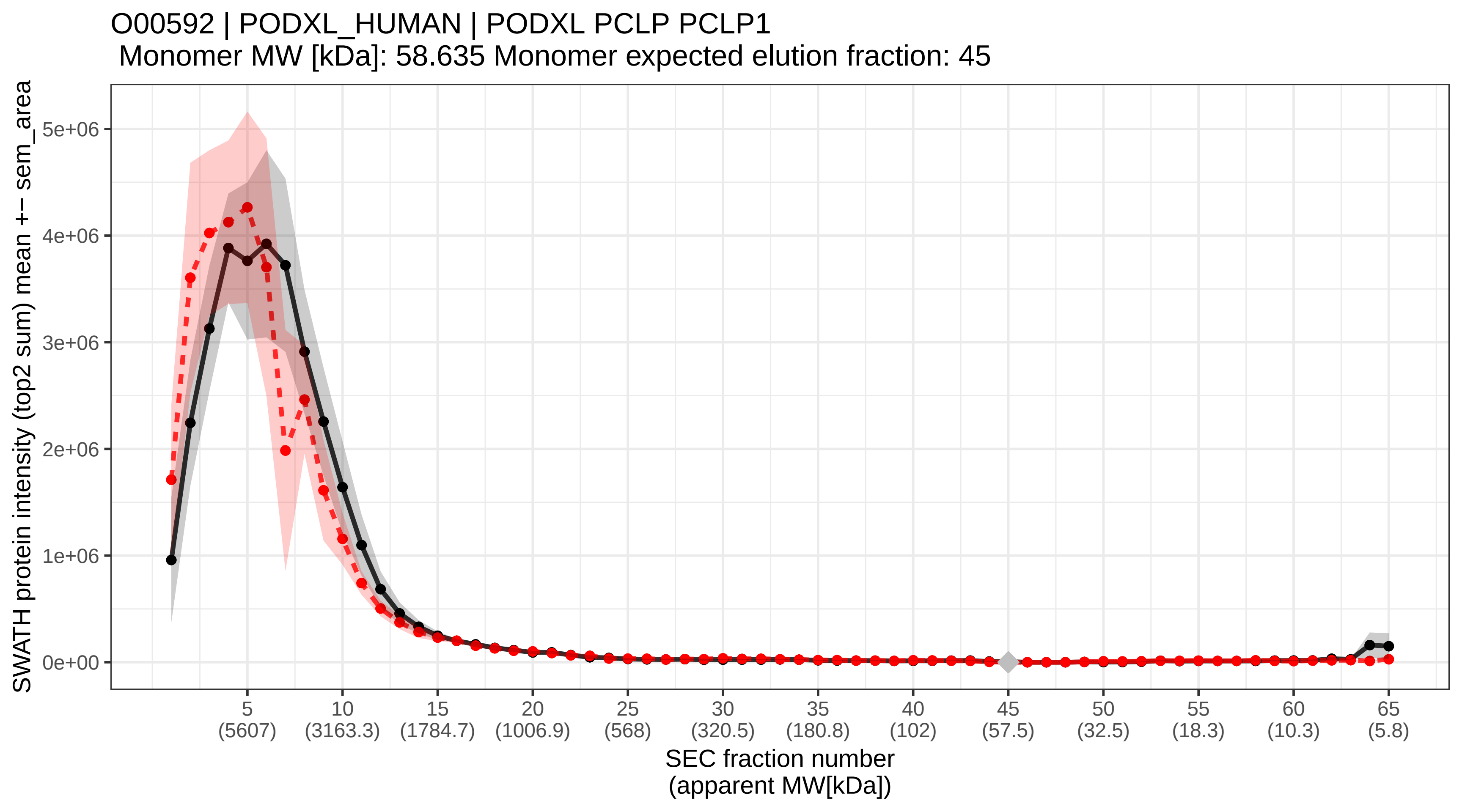

### SECchrom_O00622_CYR61_HUMAN_CYR61_CCN1_GIG1_IGFBP10.pdf

O00622 | CYR61_HUMAN | CYR61 CCN1 GIG1 IGFBP10
Monomer MW [kDa]: 42.027 Monomer expected elution fraction: 48

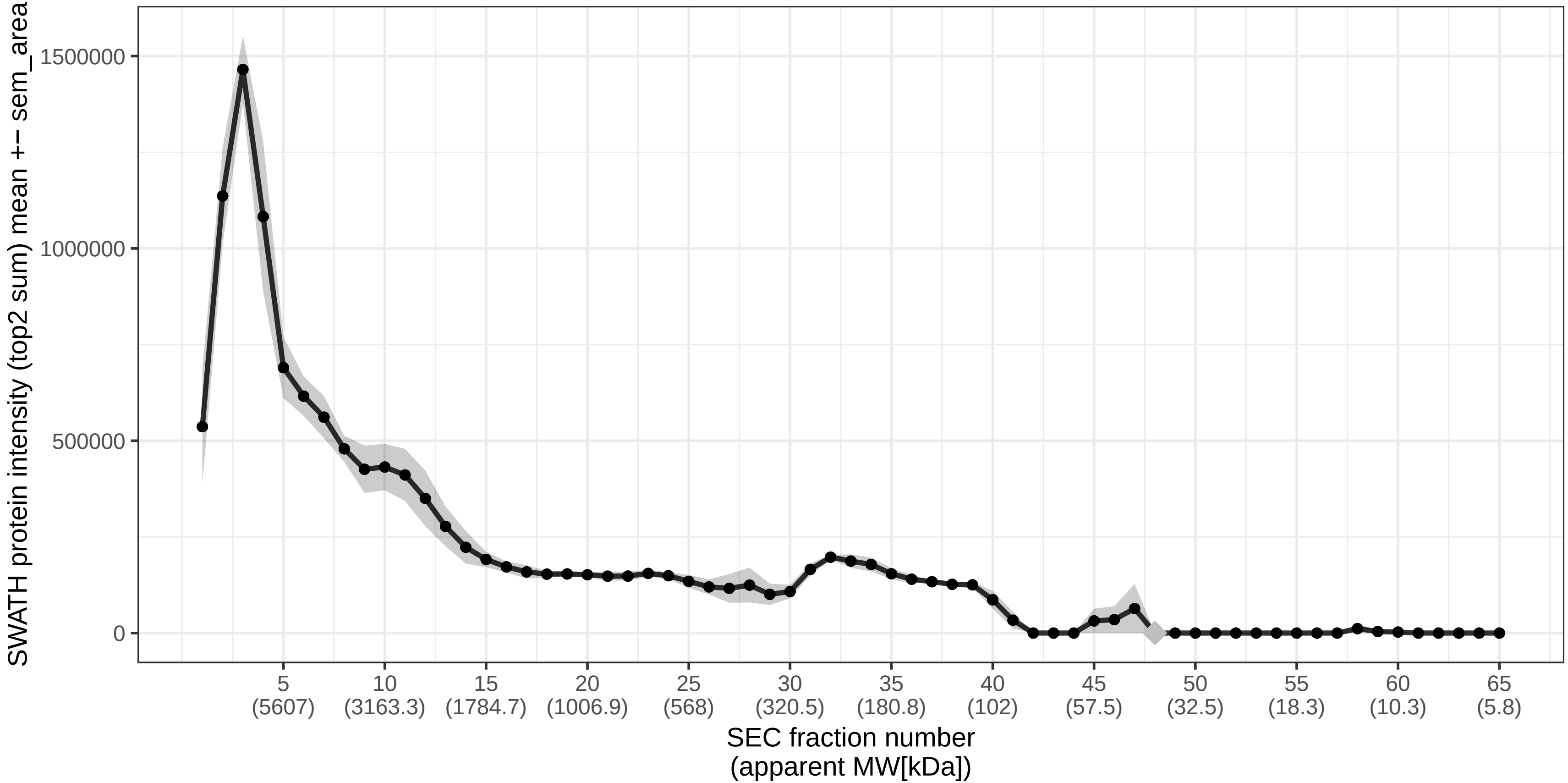

### SECchrom_O00625_PIR_HUMAN_PIR.pdf

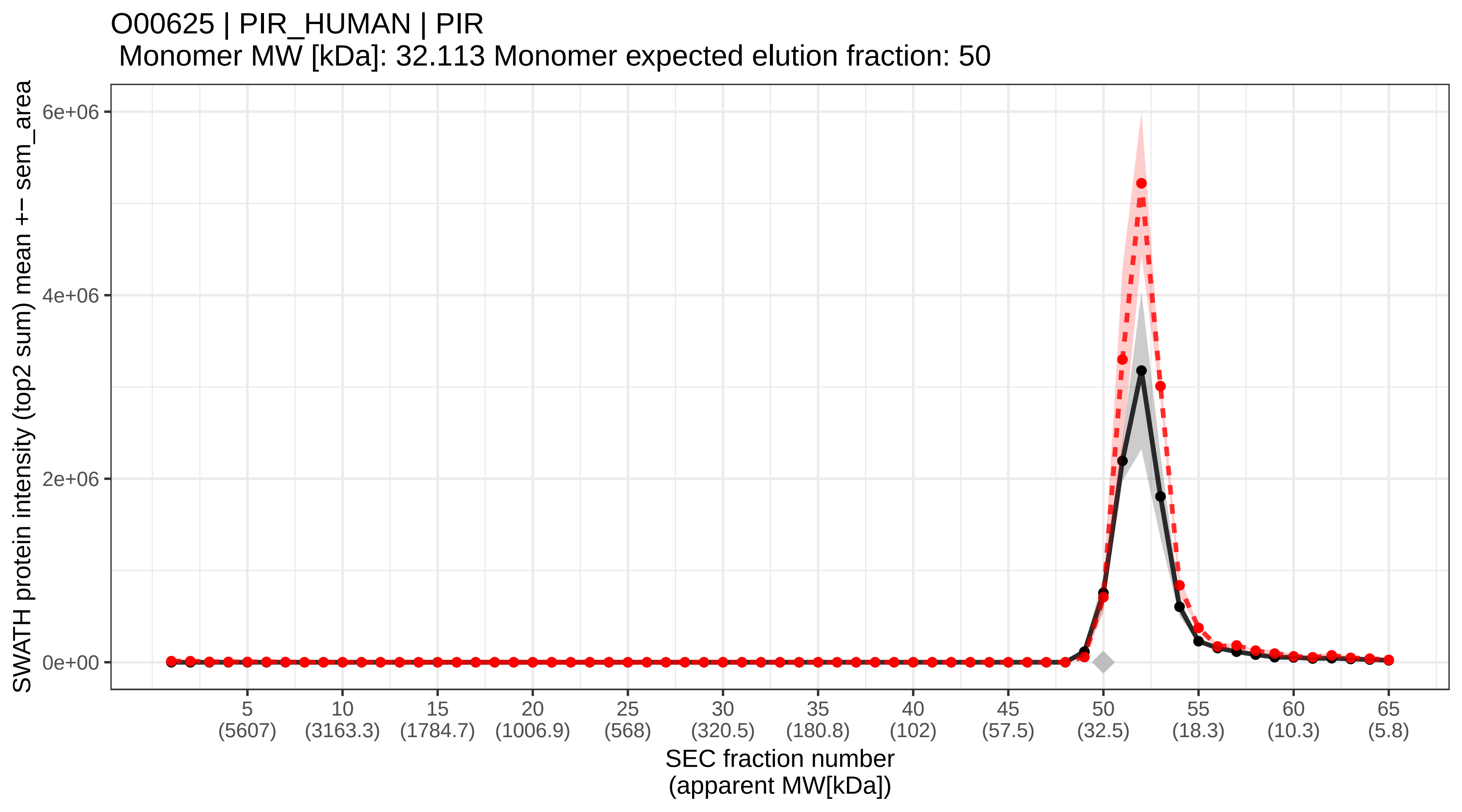

### SECchrom_O00629_IMA3_HUMAN_KPNA4_QIP1.pdf

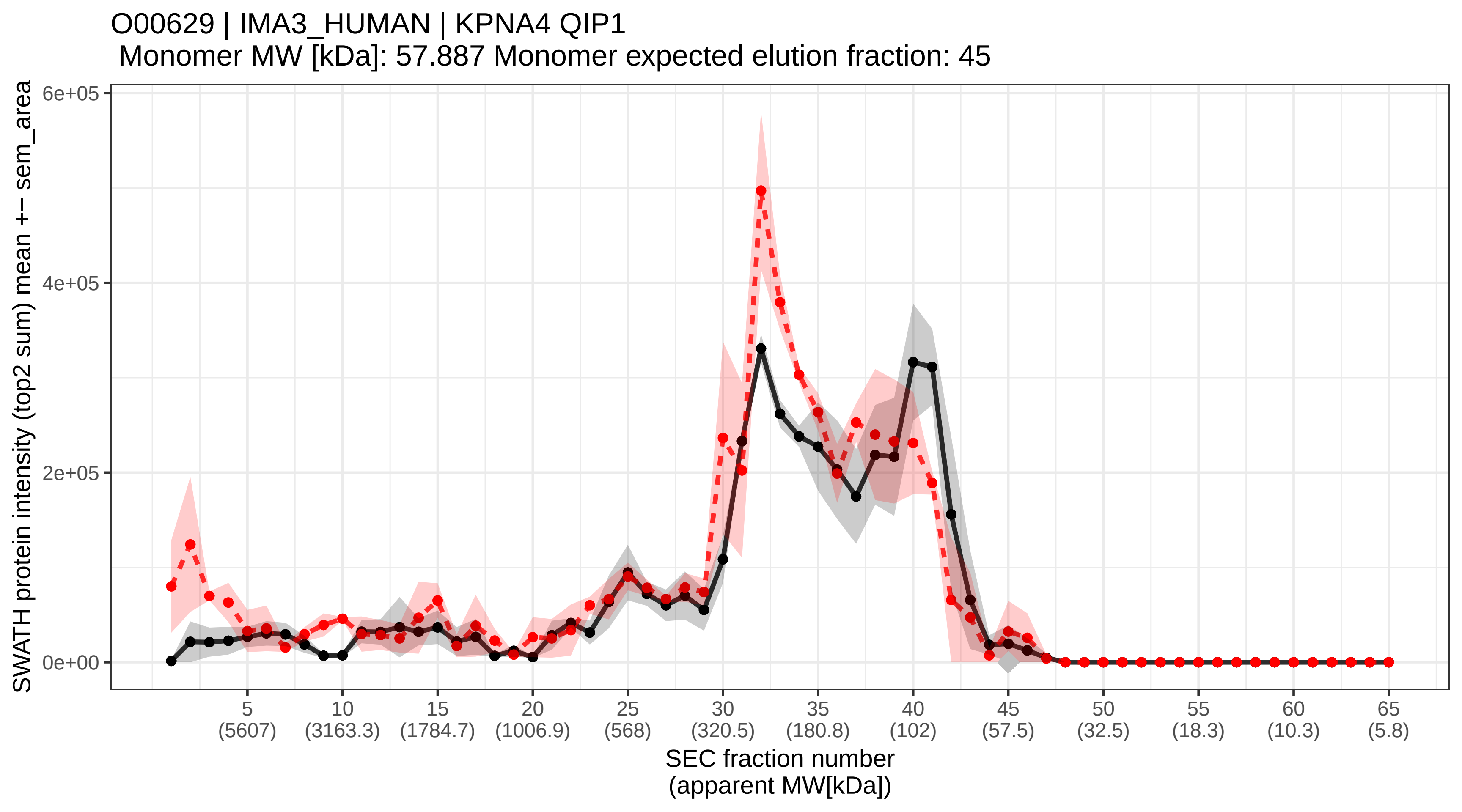

### SECchrom_O00635_TRI38_HUMAN_TRIM38_RNF15_RORET.pdf

O00635 | TRI38_HUMAN | TRIM38 RNF15 RORET
Monomer MW [kDa]: 53.416 Monomer expected elution fraction: 46

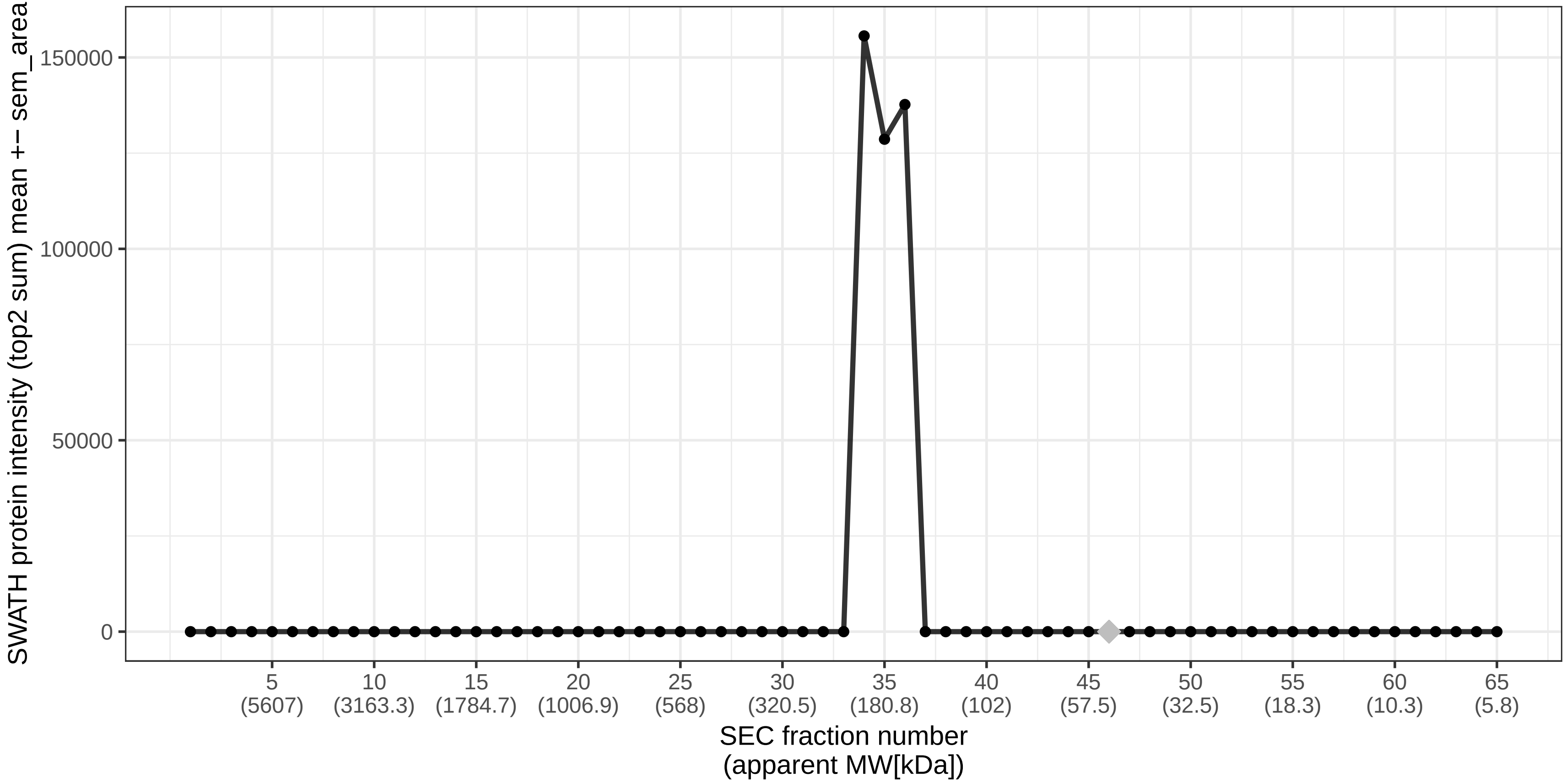

### SECchrom_O00712_NFIB_HUMAN_NFIB.pdf

O00712 | NFIB_HUMAN | NFIB
Monomer MW [kDa]: 47.442 Monomer expected elution fraction: 47

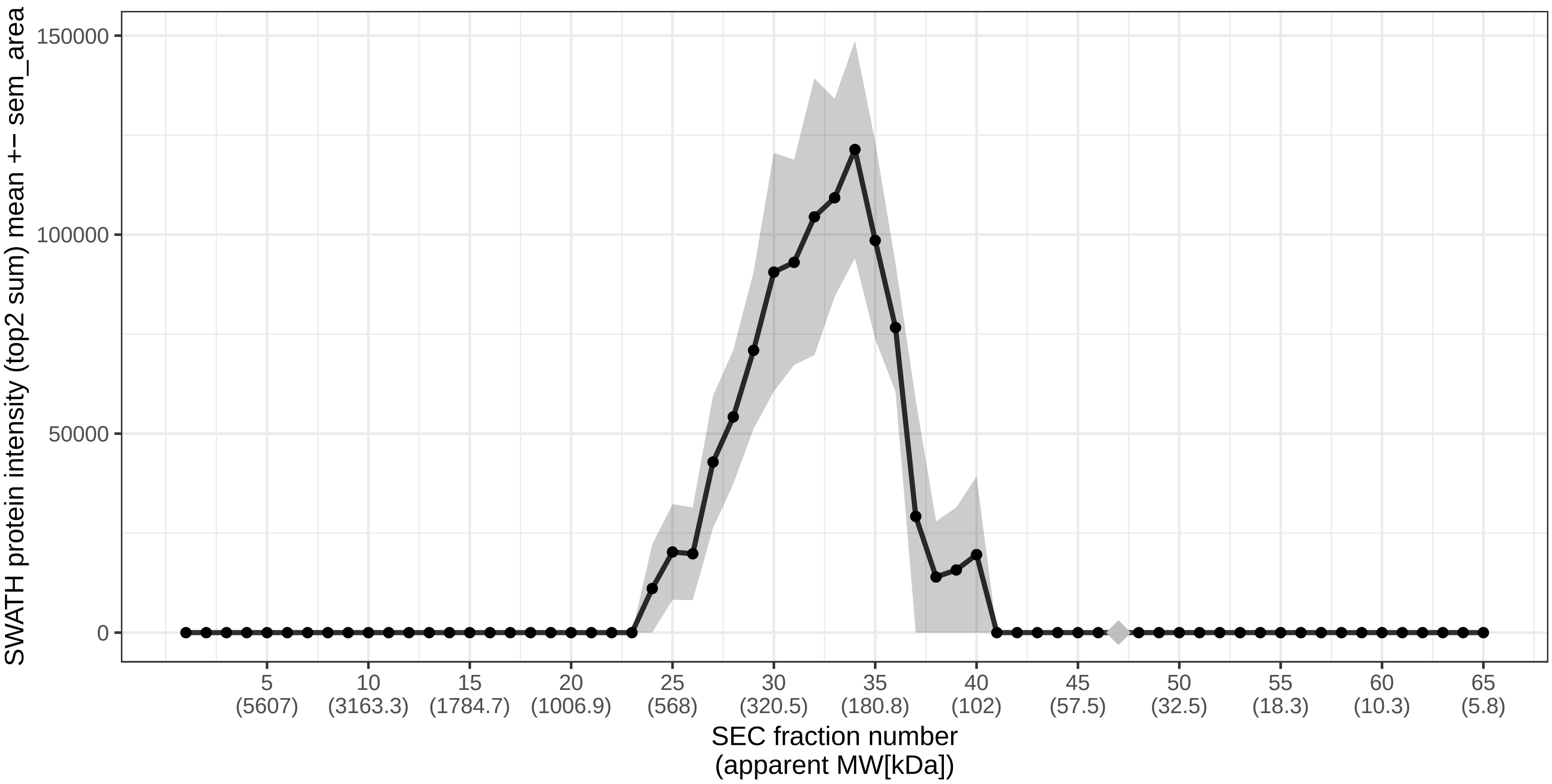

### SECchrom_O00743_PPP6_HUMAN_PPP6C_PPP6.pdf

#### O00743 | PPP6_HUMAN | PPP6C PPP6
Monomer MW [kDa]: 35.144 Monomer expected elution fraction: 49

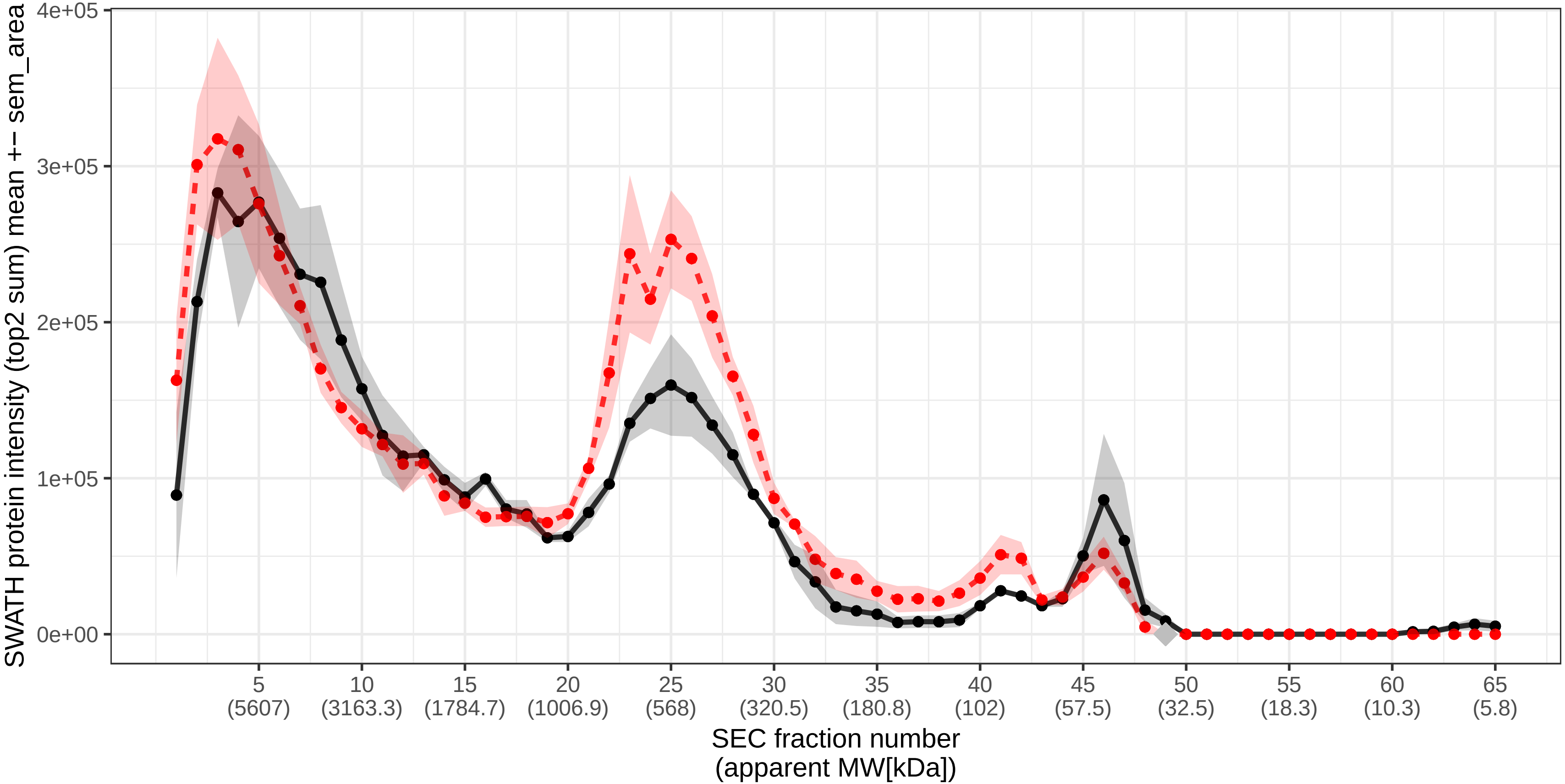

### SECchrom_O00750_P3C2B_HUMAN_PIK3C2B.pdf

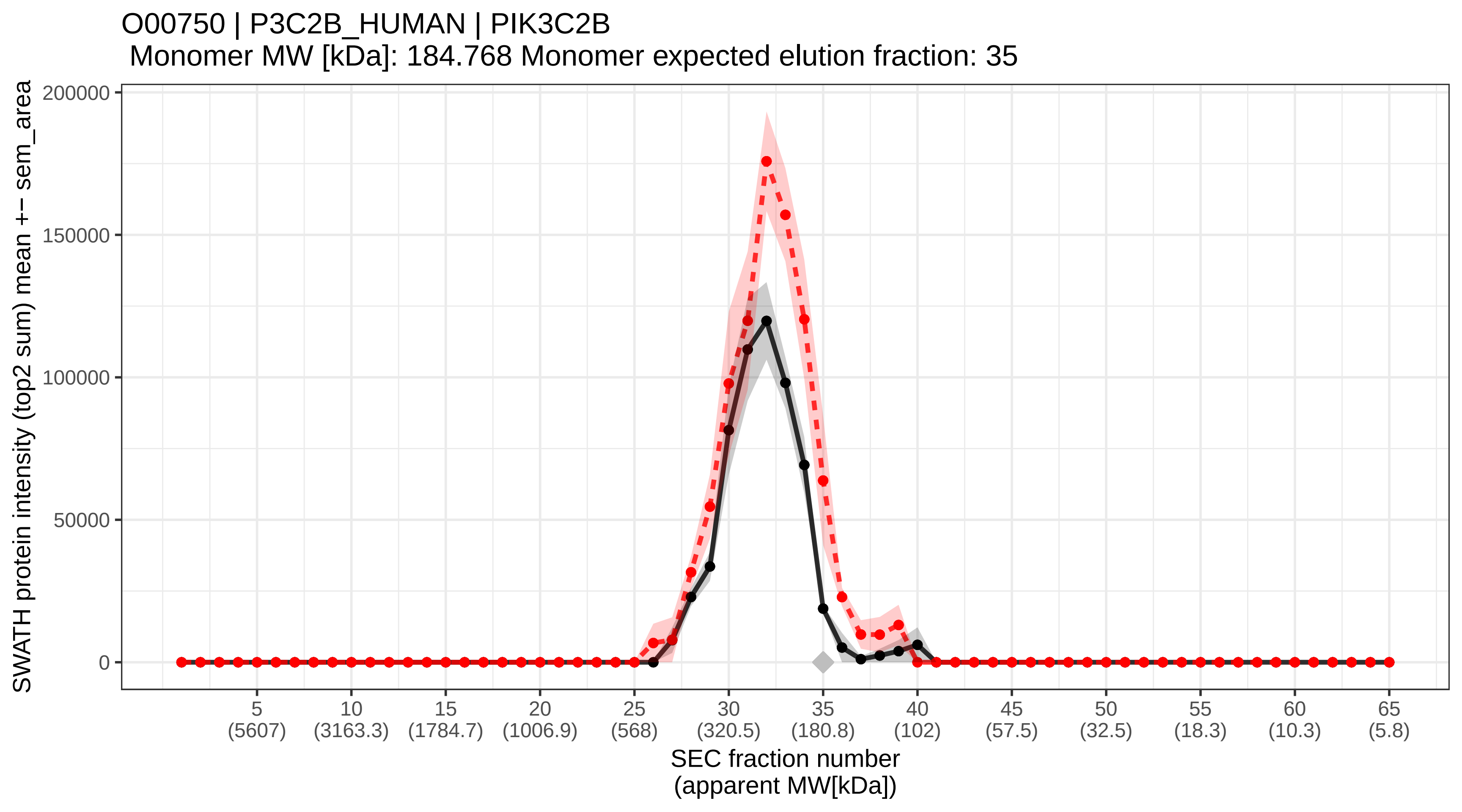

### SECchrom_O00754_MA2B1_HUMAN_MAN2B1_LAMAN_MANB.pdf

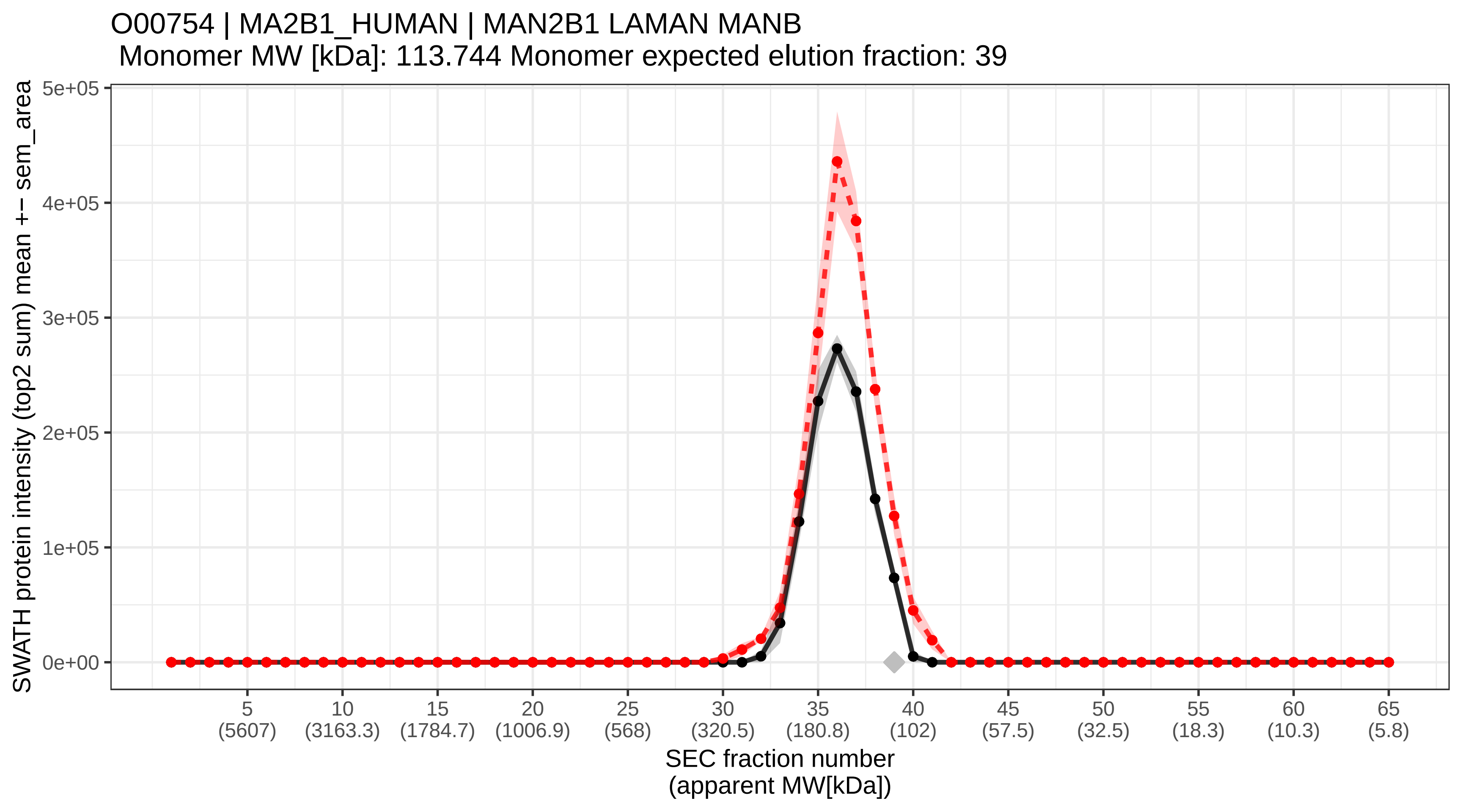
