## Supplemental Information and Figures S1-6 for "A global screen for assembly state changes of the mitotic proteome by SEC-SWATH-MS"

The supplemental information entails

- Supplemental Figures (S1-S6)

  Supplemental Figures are provided in this .pdf file,
  190506_AEB2019_supplemental_information.pdf

- one Supplemental Item (SI1):

  Supplemental Item SI1: SEC-SWATH-MS protein chromatogram plots

  enclosed as file "Supplemental Item SI1 Protein Chromatogram Plots.zip".

- three Supplemental Tables (ST1-3) with self-contained legends where applicable and the following titles.

  Supplemental Table ST1: Global protein quantification summary table

  Supplemental Table ST2: Global protein elution peak detection and statistical scoring summary table

  Supplemental Table ST3: Global complex remodeling summary table

  enclosed as file "Supplemental Tables ST1-3.zip".

### Supplemental Figures S1-6

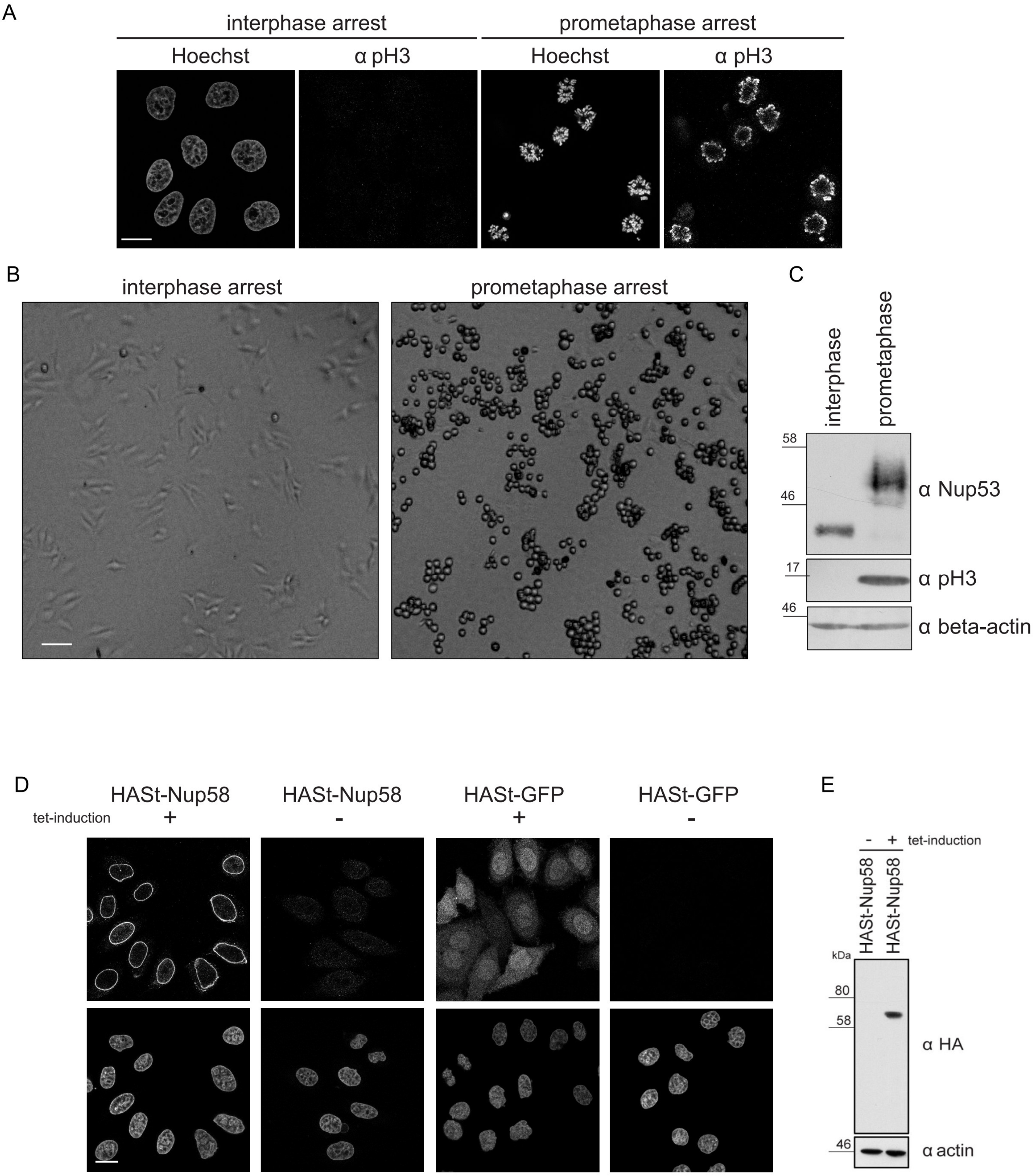


*Supplemental Figure S1: Validation of homogenous cell cycle arrest and of inducible Nup58-expressing HeLa cell lines.*

**A** Validation of homogenous interphase and prometaphase mitotic cell cycle arrest by visualizing chromatin structure (Hoechst staining) and detecting mitotic phosphorylation of histone H3 by immunofluorescence. **B** Phase contrast microscopy of HeLa CCL2 cells in interphase before harvest and in prometaphase before mitotic shake-off to retrieve only non-attached, rounded mitotic cells. **C** Homogeneity of collected cell populations assessed by immunoblotting, showing quantitative hyper-phosphorylation of Nup53 by the associated molecular weight gain. **D/E** Validation of inducible expression of HA-Strep-tagged NUP58 in HeLa cell lines using anti-HA immunofluorescence (compared to HA-Strep-GFP) and immunoblotting to confirm inducible expression of the HA-tagged bait protein.

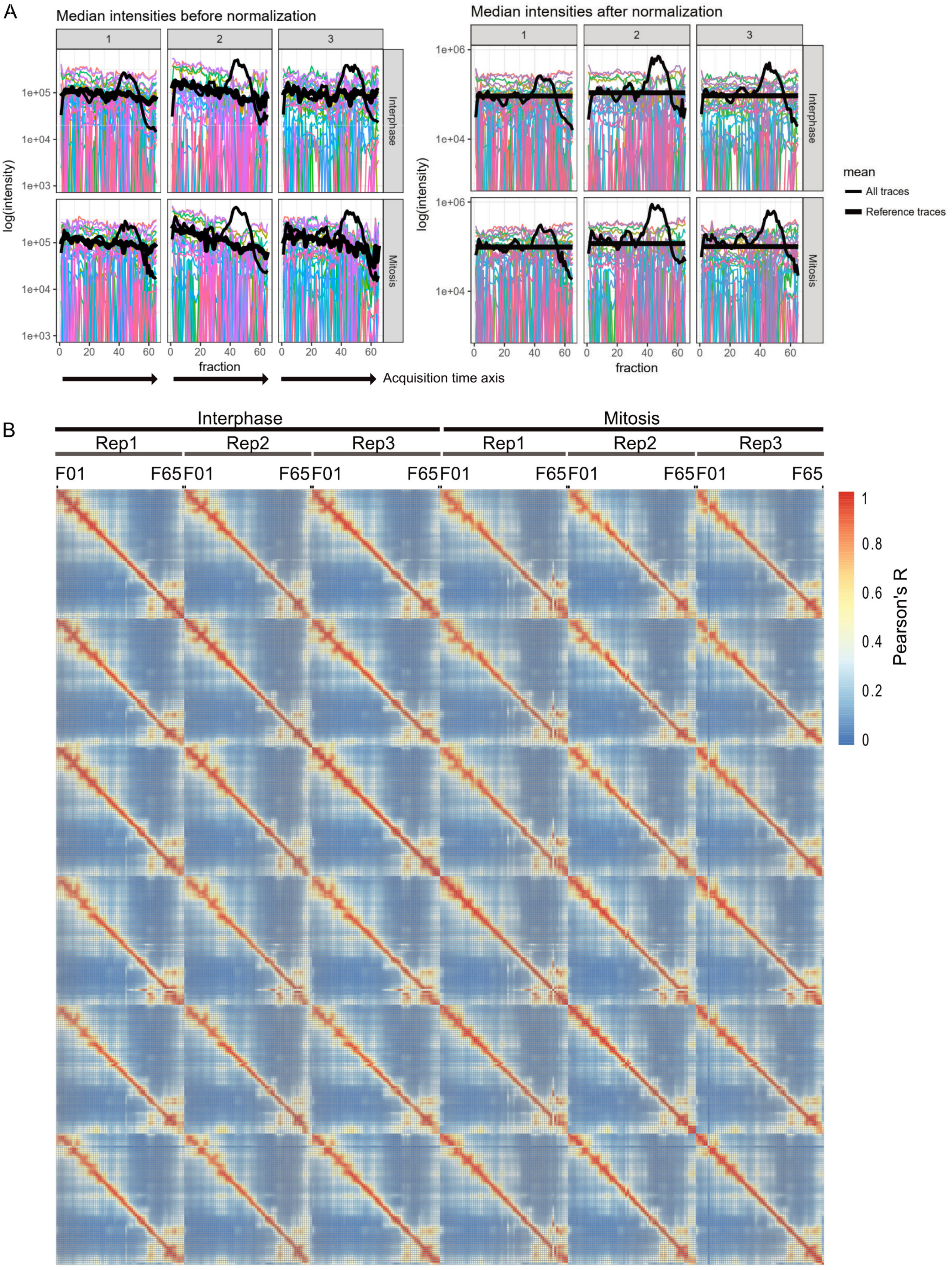


*Supplemental Figure S2: Benchmarking the SEC-SWATH-MS workflow - correction of longitudinal effects and replicate intensity correlation.*

**A** Normalization of SWATH-MS intensities based on reference peptides from a E. coli β-galactosidase tryptic digest spiked into all fractions at equal concentration. The thick black line represents the mean intensity obtained from these spike-in peptides and indicates longitudinal deterioration of mass spectrometer performance in all replicates (columns) and conditions (rows) (left panels). Peptide intensities over all 390 SWATH-MS measurements were corrected by scaling to the internal standard spike-in. Impact on summed global peptide traces is given in the thin black line. Furthermore, scaling and global intensity sum smoothing along fractions was applied. For details see experimental procedures. Arrows indicate the order of MS acquisition, alternating between different conditions in blocks of four consecutive fractions (i.e. measuring Interphase(Frac1,2,3,4)-Mitosis(Frac1,2,3,4)-Interphase(Frac5,6,7,8), etc.). The acquisition scheme can be obtained from the annotation tables available with the dataset made available via ProteomeXchange (PXD010288). **B** Protein intensity correlation analysis visualized in symmetric heatmap. Proteins were quantified based on the sum of the top2 most-intense peptides across the full dataset. Peptides were pre-filtered based on sibling peptide correlation along the SEC dimension (For details, see experimental procedures). Protein-level correlation analysis was performed on the global intersect set of proteins observed in all fractions. Resulting protein intensities are highly correlated across replicates and adjacent fractions with an average Pearson's $R > 0.98$ between replicate fractions of the same biological condition. Labels are given on the top, sample order is identical in x- and y- directions. Sample-sample correlation further shows highest resolution ("sharpness" of diagonal) in the SEC fraction range 40-50 and lower resolution in the early 'void volume' fractions 1-5 where many analytes greater than or equal to the exclusion limit of the 500 Å diameter pores of the Sepax SRT-C SEC 500 column elute.

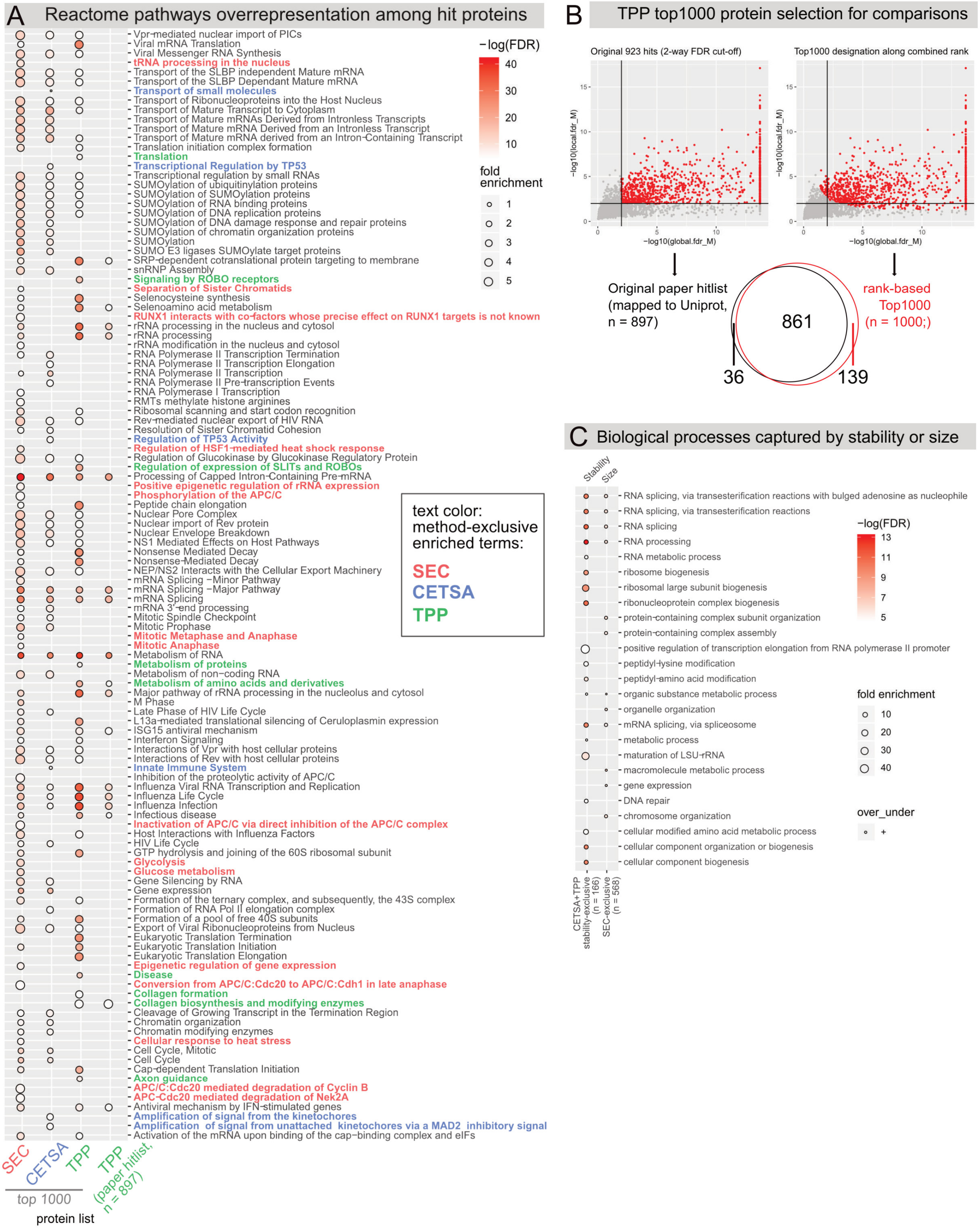


*Supplemental Figure S3: Pathway enrichment among top 1000 proteins recovered by size or thermostability profiling along the cell cycle.*

**A** Pathway statistical overrepresentation analysis among the top 1000 proteins (compare main **Figure 4**) against background of all 6,010 proteins characterized by at least one of the methods using the Panther gene ontology classification system (http://pantherdb.org/, Reactome pathways) and filtering results to 0.01 FDR. Labels are color-coded if recovered exclusively by one method. For comparison, the original list of hit proteins from Becher et al. 2018 (hit_M, mapped to Uniprot) is included. Results largely converge in terms of the enriched underlying pathways. Only SEC and CETSA result sets show significant overrepresentation of proteins associated with terms "Cell Cycle / Mitotic Cell Cycle". Multiple expectable pathways are exclusively recovered by SEC-SWATH-MS. **B** Ranking and selection of Top 1000 proteins from TPP results in comparison to hit protein designation in the original study. Rank-based selection f the top 1000 proteins covers most proteins designated as hits in the original study (96 %, 861 of 897 Uniprot identifiers). Protein ranking enables rank comparisons such as presented in **Figure 4F**. **C** Biological process statistical overrepresentation analysis among the proteins reported to change size (exclusively by SEC-SWATH-MS, n = 568) or proteins reported to change stability (with high confidence, reported by both CETSA and TPP, n = 166). Sets were assigned as in in **Figure 4C**. Overrepresentation was assessed against the full genome background using the Panther gene ontology classification system (http://pantherdb.org/) and filtering results to 0.01 FDR. Both RNA splicing and different instances of protein complex biogenesis are retrieved by either protein property. Also metabolic and enzymatic processes are detectable by either stability or size. Despite the difference in the protein sets the results suggest convergence on the functional level.

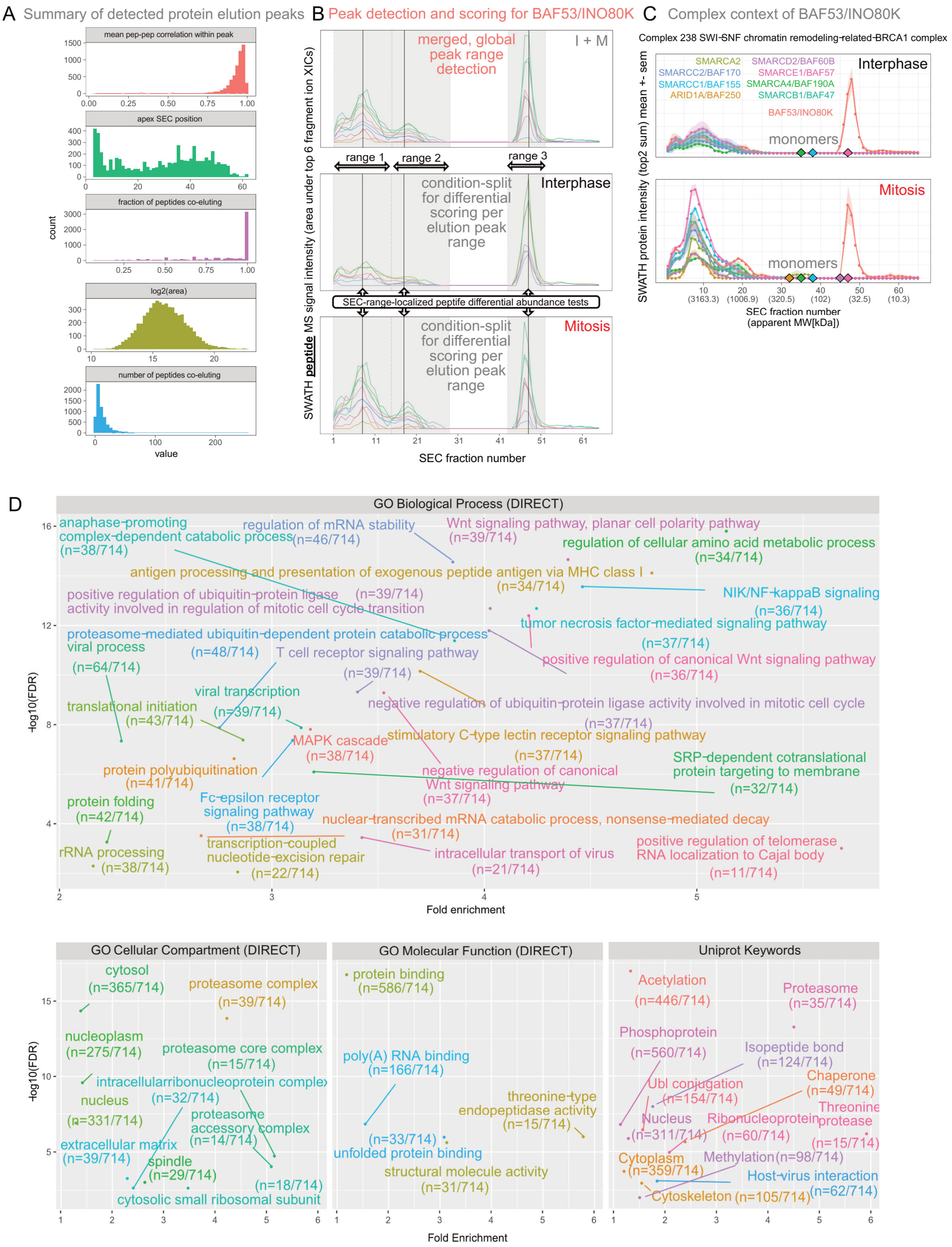


*Supplemental Figure S4: Protein elution peak detection and properties of multi-complex signal proteins.*

**A** Summary of the 6,040 distinct elution peaks detected from 4,515 proteins. The analysis was performed using the summed peptide SEC profiles as input to the *CCprofiler* protein-centric module and under strict error control against randomized peptide-to-protein associations (*CCprofiler* protein-centric q-value = 5%, compare **Figure 2B** and see experimental procedures for details). From top to bottom, detected protein elution peaks show i) high intra-peak peptide-peptide correlation. ii) Peak apex distribution across fractions with crowding in the void peak of proteins or -assemblies sized equal to or larger than the 500 Å pores (ca. 10 MDa). iii) High fraction of co-eluting peptides participating in peaks displays overall agreement but also delineates incomplete peak groups where some peptides show distinct SEC elution patterns. This could arise from either technical noise or biological signal such as alternative splicing or co- or post-translational processing or modification (Compare Uniprot annotation for Acetylation, Phosphoprotein and Ubl conjugation enriched among multi-complex proteins in panel C). iv) Large dynamic range and near-normal distribution of intra-peak-peptide MS intensity and v) Distribution of peptides co-eluting per peak detected. **B** Protein-centric peak detection and differential association testing (SEC-localized differential abundance testing) exemplified on the protein BAF53. Protein elution peaks are detected from peak groups of unique peptides of BAF53, defining the ranges for SEC-localized differential abundance testing. Differential abundance in the protein elution peaks indicated differential association of BAF53 to different protein modules, as apparent from a global shift of its parent complex and sibling subunits (panel C). Upper panel, one globally observable set of protein elution peaks is detected in a protein-centric fashion based on co-eluting protein-specific peptides from an artificially generated, merged 'master map' of summed peptide intensities across replicates and conditions. The detected peak ranges (3 distinct ranges 1-3 detected for BAF53) then serve as stencil of defined ranges for condition-dependent differential peptide abundance tests as proxy for differential parent protein association with the underlying entity of distinct molecular weight (lower panels). **C** BAF53 in the context of a parent complex shows joint re-arrangement and mitotic concentration of protein mass in a ca. ~ 3.5 MDa assembly. **D** DAVID functional annotation enrichment testing of proteins observed in two or more distinct complex-assembled states against the background of all 4515 proteins for which at least one elution peak was detected (performed at https://david.ncifcrf.gov/). Multi-complex proteins are enriched in signaling factors (despite global underrepresentation of signaling molecules, see **Supplemental Figure S3B**) and proteins known to have binding functions, as well as proteins known to function in relation to post-translational modifications such as acetylation and phosphorylation.

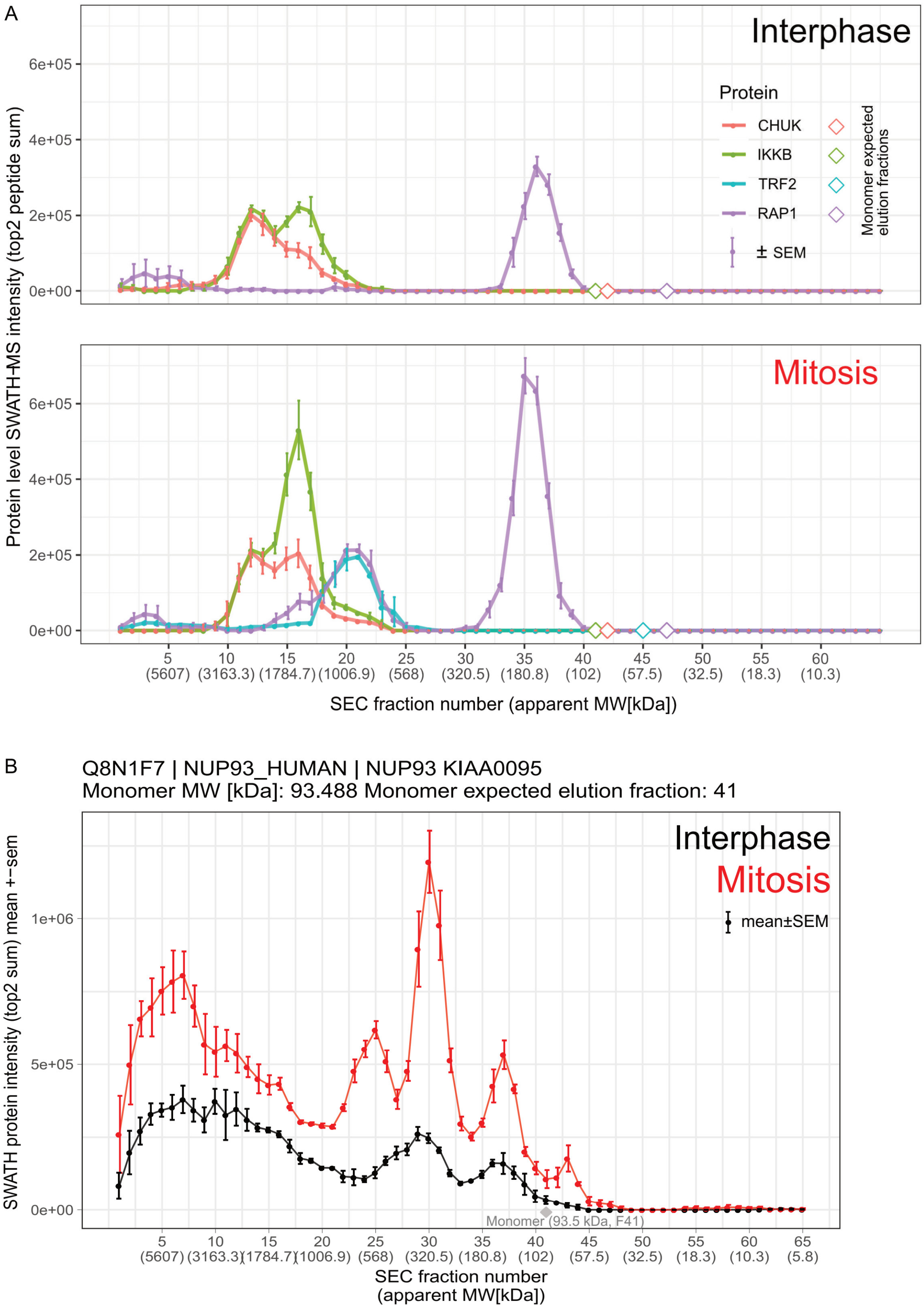


*Supplemental Figure S5: Investigation of RAP1-TRF2-IKK cross-complex interaction and zoom-in to NUP93 subpopulations resolved across mitotic states.*

**A** Related to Figure 5F. SEC chromatogram overlay to assess complex-complex interaction between RAP1-TRF2 complex and the IKK complex (represented by subunits CHUK and IKKB). IKK complex appears in two subpopulations, one of ca. 2.5 MDa (apex fraction 12) and one of ca. 1.7 MDa (apex fraction 16), with a peak shoulder in the mitotic 950 kDa-elution signal of RAP1 in principle consistent with recruitment of a fraction of RAP1 but not TRFII to the 1.7 MDa instance of the IKK complex. Equivalent custom analyses can rapidly be performed via *SECexplorer-cc*. **B** Related to Figure 6. Zoom-in to NUP93 elution profiles in interphase and mitosis (compare Figure 6) highlights distinct sub-complex signals arising in mitosis. This includes a new signal of ca. 568 kDa (elution range F21-F27) and two signals of increased intensity when compared to the interphase pattern at apparent MW of ca. 320 kDa (elution range F27-F33) and 145 kDa (elution range F35-F39).

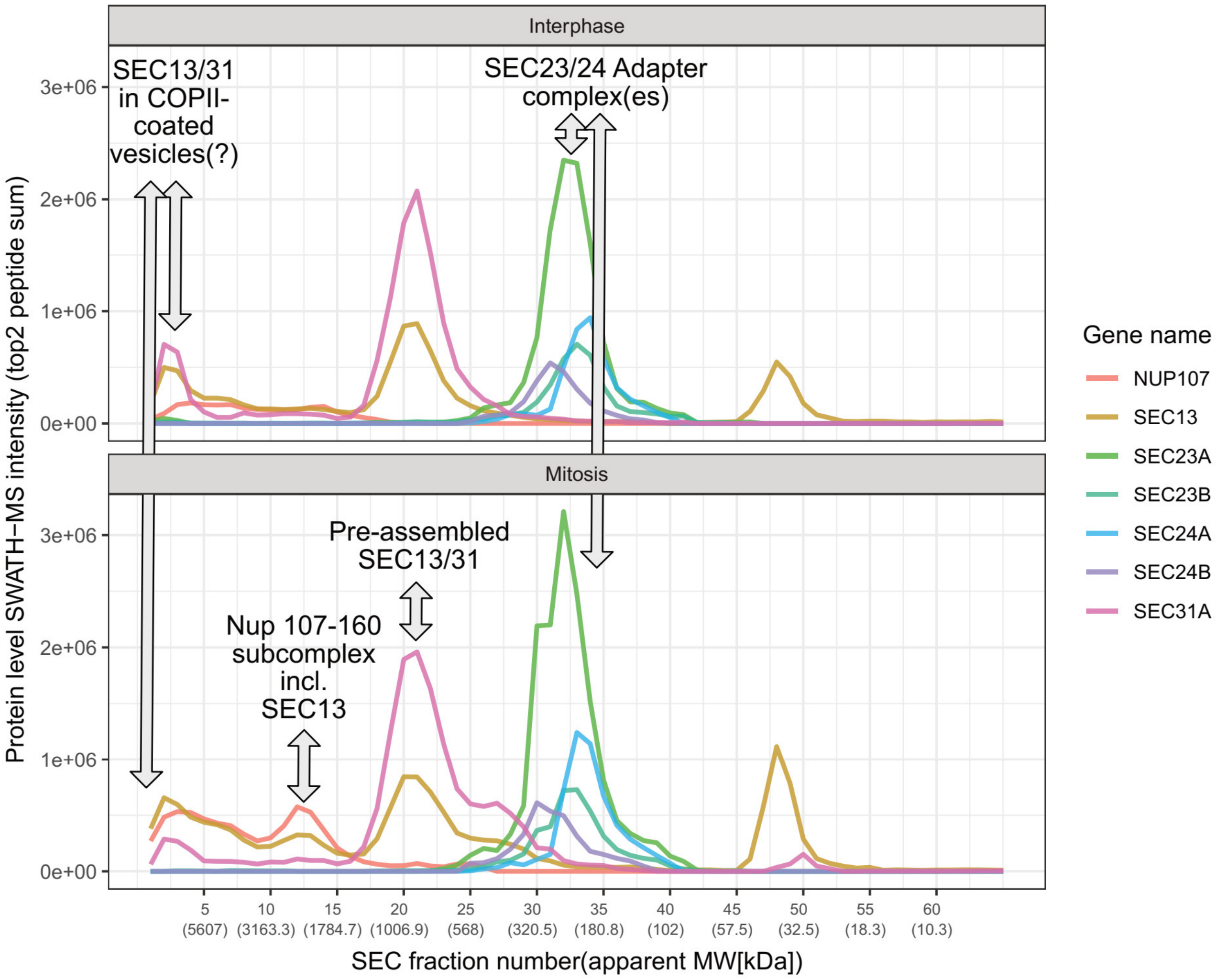


*Supplemental Figure S6: Investigation of SEC13 moonlighting across Nup107-160 subcomplex and COPII vesicle transport related complexes.*

Related to Figure 5H. Protein-level mean SEC-SWATH-MS chromatograms of Nup107-160 subcomplex subunits NUP107 and SEC13 and additional protein complex partners of SEC13 from its additional functional context as part of COPII coatomer complexes involved in anterograde vesicle-mediated transport, SEC23(A/B), SEC24(A/B) and SEC31(A). From left to right, large to small assemblies, the profiles suggest: (i) Potential SEC13/31 interaction in and elution as part of COPII-coated vesicles in the void volume; (ii) NUP107 and SEC13 appearance as part of the Nup107-160 subcomplex eluting with apex in fraction 12/13, ca. 2.8 MDa; (iii) SEC13-SEC31 co-elution with ca. 1 MDa, apex fraction 20-21, potentially pre-assembled when not participation in coatomers and (iv) SEC23A/B and SEC24A/B co-elution in fractions 30-35, 320 - 180 kDa, likely preassembled into a mixed population of variant adaptor complexes.
